# Joint single-cell capture of Cas9 edits and transcriptomes reveals on- and off-target effects on gene expression

**DOI:** 10.1101/2025.02.07.636966

**Authors:** Michael H. Lorenzini, Brad Balderson, Shanna N. Lavalle, Karthyayani Sajeev, Aaron J. Ho, Madelyn Light, Olalekan H. Usman, Gavin Kurgan, Graham McVicker

**Author notes:** These authors contributed equally (Michael H. Lorenzini, Brad Balderson).

## Abstract

A longstanding barrier in genome engineering with CRISPR-Cas9 has been the inability to measure editing outcomes and their functional effects at single-cell resolution. Here we present “Superb-seq”, which captures single-cell edits and measures their effects on the transcriptome by integrating Cas9 editing, T7 transcription, and single-cell RNA sequencing. We performed Superb-seq on over 34,000 cells from three cell lines. Using seven guide RNAs to target four chromatin remodeler genes in 10,000 K562 cells, Superb-seq detected 43 total edit sites, including 36 off-target sites, and 5761 edited cells (up to 5 edits per cell). Superb-seq improved estimation of gene perturbation effects compared to Perturb-seq and its edit detection sensitivity was comparable to bulk off-target detection methods. We identified 19 off-target edits associated with differential gene expression, nine of which were cell type specific. In summary, Superb-seq illuminates the consequences of Cas9 genome editing by connecting detected single-cell edits with changes in gene expression.

## Introduction

Cas9, the RNA-guided CRISPR nuclease, is a fundamental tool for programmable genome editing, with broad applications in functional genomics and therapeutics^1^. Assays that combine Cas9 editing with high-throughput single-cell RNA-seq, broadly referred to as Perturb-seq, have made it possible to measure the effects of gene perturbations on the transcriptome across millions of cells^2–7^. However, a key gap in current single-cell CRISPR methods is that they lack the ability to directly read genome editing outcomes. Perturb-seq reads the guide RNAs of each cell as a proxy for edit results^5,8^ (although it can occasionally record edit events in transcribed regions^9^) and downstream analysis methods associate detected guides with molecular effects^10^. However, this “guide capture” approach is confounded by the ambiguous association between detected guides and underlying edit events. Major sources of ambiguity include the unreliable detection of guides by some Perturb-seq methods^8^, and the inherent variability in on- and off-target activity of guides^11,12^. In a Perturb-seq experiment, up to 50% of guides can exhibit no effect on the intended target gene, despite a perfectly matching guide sequence^13^ (although this may vary by cell type and experimental conditions). This on-target inactivity produces unedited guide-positive cells and dampens sensitivity to detect the effects of functional sequences^14^, especially non-coding sequences that have modest effects on gene expression^15^. Furthermore, most guides exhibit off-target activity—the editing of unintended sequences with guide similarity^16^. Off-target edits that affect gene expression confound interpretation of on-target effects and biological mechanisms^6,12^.

Cas9 editing has also emerged as a powerful therapeutic tool in medicine^17^, with on-going clinical trials testing Cas9 editing to directly correct the genetic causes of diseases^18^ such as sickle-cell anemia^19^. However, a key concern is whether Cas9 off-targeting generates undesirable biological effects^20^. Currently, to manage off-targets, researchers computationally predict a large number of off-target genome sites^21^, refine computational predictions by bulk sequencing screens in cells^22,23^, and validate off-targets by bulk amplicon sequencing^24^. However, these methods lack the ability to assess how edits affect genome functions like gene expression. Bulk off-target sequencing does not read gene expression, and bulk functional assays (e.g. RNA-seq) aggregate effects across on- and off-target edits instead of isolating them. In addition to limited functional characterization, current bulk methods lack sufficient scalability to identify Cas9 off-targets in personal genomes^25^. This limitation could be addressed by single-cell edit capture, which would make it possible to identify patient-specific editing at scale by de-multiplexing cells by donor^26^.

Here we present **“Superb-seq”**, a scalable genome perturbation method for single-cell measurement of genome-wide Cas9 edits and associated transcriptome changes. In contrast to Perturb-seq’s single-cell guide capture, Superb-seq performs “single-cell edit capture” using a three-step protocol that requires only standard laboratory equipment and off-the-shelf reagents (**Fig. 1A**). First, live cells are edited with Cas9 ribonucleoprotein (RNP) and phage T7 promoter to generate promoter-labeled edits by homology-free insertion. Second, T7 in situ transcription^27^ (IST) is performed on fixed cells to generate edit-reporting T7 transcripts in each cell. Third, single-cell T7 transcripts and endogenous transcripts are jointly acquired by combinatorial scRNA-seq and analyzed. This work describes the development and validation of Superb-seq, two Superb-seq experiments in a total of 35,000 cells, and a companion software package for Superb-seq data processing named **“Sheriff”**.

**Figure 1.**
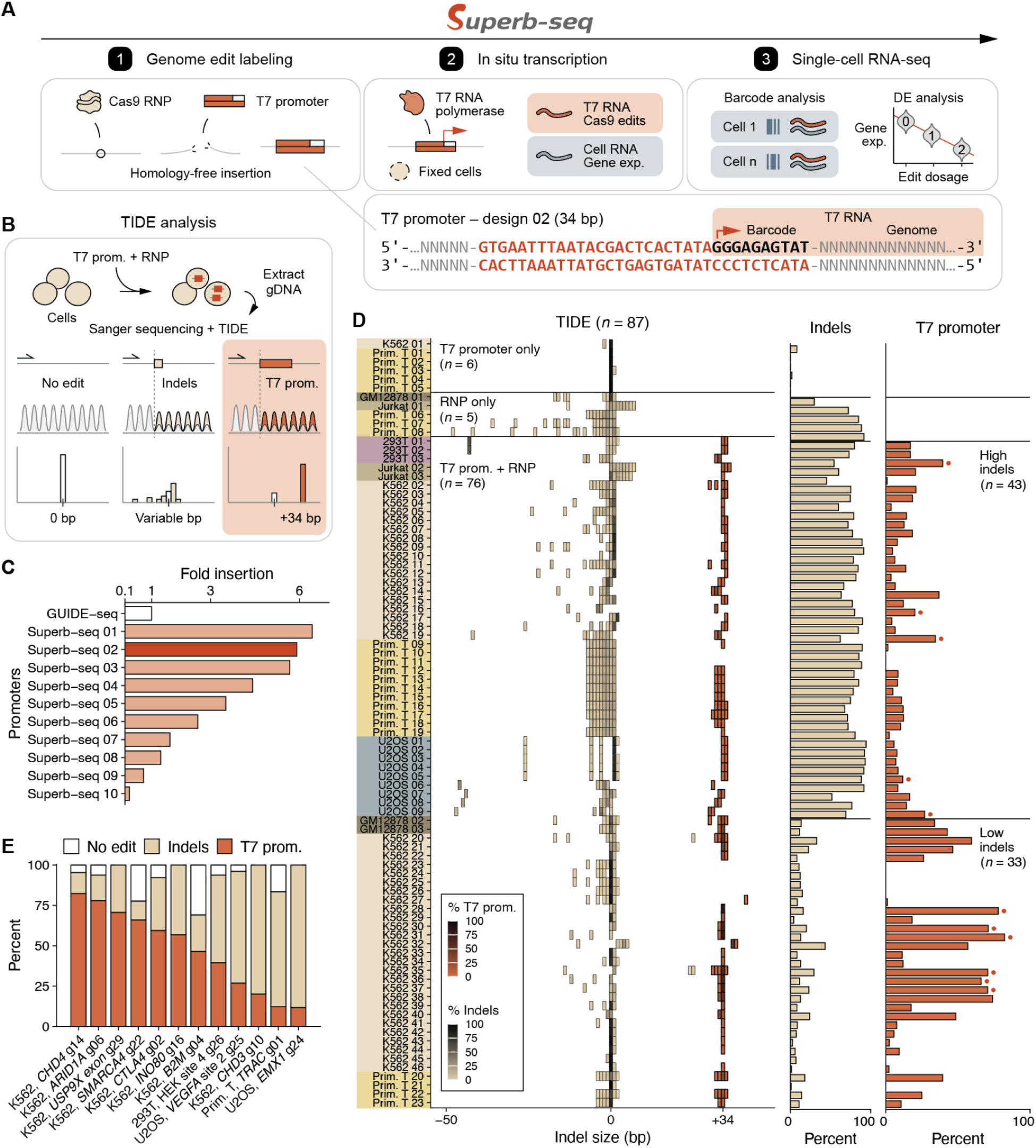
Homology-free labeling of genome edits with phage T7 promoter. (**A**) Superb-seq edit labeling with T7 promoter design 02, in situ transcription, and joint single-cell analysis of T7 and cellular transcripts. (**B**) TIDE analysis^37^ of T7 promoter labeling. (**C**) Relative insertion efficiency of ten T7 promoter designs compared to GUIDE-seq “dsODN”, by TIDE. Design 02 value is the mean of four samples. (**D**) TIDE analysis of 87 human cell samples. T7 promoter events are defined as > 30 bp insertions. All other events are defined as unlabeled (indels). Red dots indicate cell lines included in Superb-seq analyses. (**E**) T7 promoter labeling outcomes across multiple cell types and genome regions.

## Results

### Homology-free labeling of genome edits with T7 promoter sequences

To develop Superb-seq, we first established genome edit labeling with T7 promoter sequences. We chose a homology-free labeling strategy by cell electroporation with Cas9-guide RNA ribonucleoprotein (RNP) and double-stranded donor DNA encoding the T7 promoter. RNPs open double-strand breaks in the genome^28,29^, and the T7 promoter is inserted by end joining repair^30,31^ (**Fig. 1A**). Compared to homology-directed repair, two advantages of this approach are modular labeling for different guides and genome sites^22^, and broader compatibility with non-dividing cell types^32^.

To design the Superb-seq T7 promoter, we screened a panel of 27 constructs encoding T7 and SP6 phage promoters^33–36^ (**Supp. Fig. 1A, Supp. Table 1)**. We quantified homology-free insertion efficiency of each promoter by TIDE^37^ (tracking indels by decomposition, **Fig. 1B**) and compared efficiency to the published GUIDE-seq “dsODN”. Insertion efficiency varied widely across designs, and over 60% of designs (17 of 27) did not generate detectable labeling (**Supp. Fig. 1B**), suggesting that labeling performance is sensitive to construct length and composition. However, three T7 promoters (designs 01, 02, 03) achieved strong edit labeling, with 5.7–6.4 fold higher efficiency than the GUIDE-seq dsODN (**Fig. 1C**). These top-performing designs combined sequence elements of the GUIDE-seq dsODN and optimized T7 promoters from Cel-seq+ (**Supp. Fig. 1C**)^36^. Among these, the design 02 T7 promoter was previously shown to be amenable to cDNA synthesis for scRNA-seq libraries^36^. Therefore we selected design 02 as the Superb-seq T7 promoter (**Fig. 1A**). Insertion of this T7 promoter into coding sequences is expected to introduce frameshifting mutations and induce nonsense-mediated decay (NMD, **Ext. Data Fig. 1, Supp. Note 1**), so that it can be used for gene knockout experiments.

To establish that the T7 promoter effectively tags Cas9 edits in a variety of genome contexts, we generated > 130 edited cell samples and controls, using six human cell types (HEK293, GM12878, Jurkat, K562, U2OS, and primary T cells) and 30 guide RNAs targeting 11 genome loci (**Supp. Table 1)**. We then characterized the editing outcomes in 87 samples (11 controls, 76 T7 promoter edits mostly with design 02) that yielded confident TIDE results with *R*^2^ > 0.5 (**Fig. 1D, Supp. Fig. 1D, Supp. Table 2**). Treatment with both T7 promoter and RNP resulted in frequent +34 bp insertion events, indicating successful labeling of Cas9 edits with the Superb-seq T7 promoter in a variety of cell and genome contexts (**Fig. 1D**). High T7 promoter labeling of 40–80% was achieved at six genome regions (**Fig. 1E**) and in all cell types except Jurkat and U2OS cells, in which insert efficiency was lower (although we also tested fewer guides in these cell types, **Supp. Fig. 2C**). Across 30 guide RNAs tested, we observed that guides generated T7 promoter insertions accompanied by a bifurcated low or high rate of background indels (**Supp. Fig. 2, Supp. Note 2**). The size of T7 promoter insertion events was consistently within a few base pairs of +34 bp (**Ext. Data Fig. 1C**), demonstrating successful full-length T7 promoter labeling in a variety of genome editing contexts.

**Figure 2.**
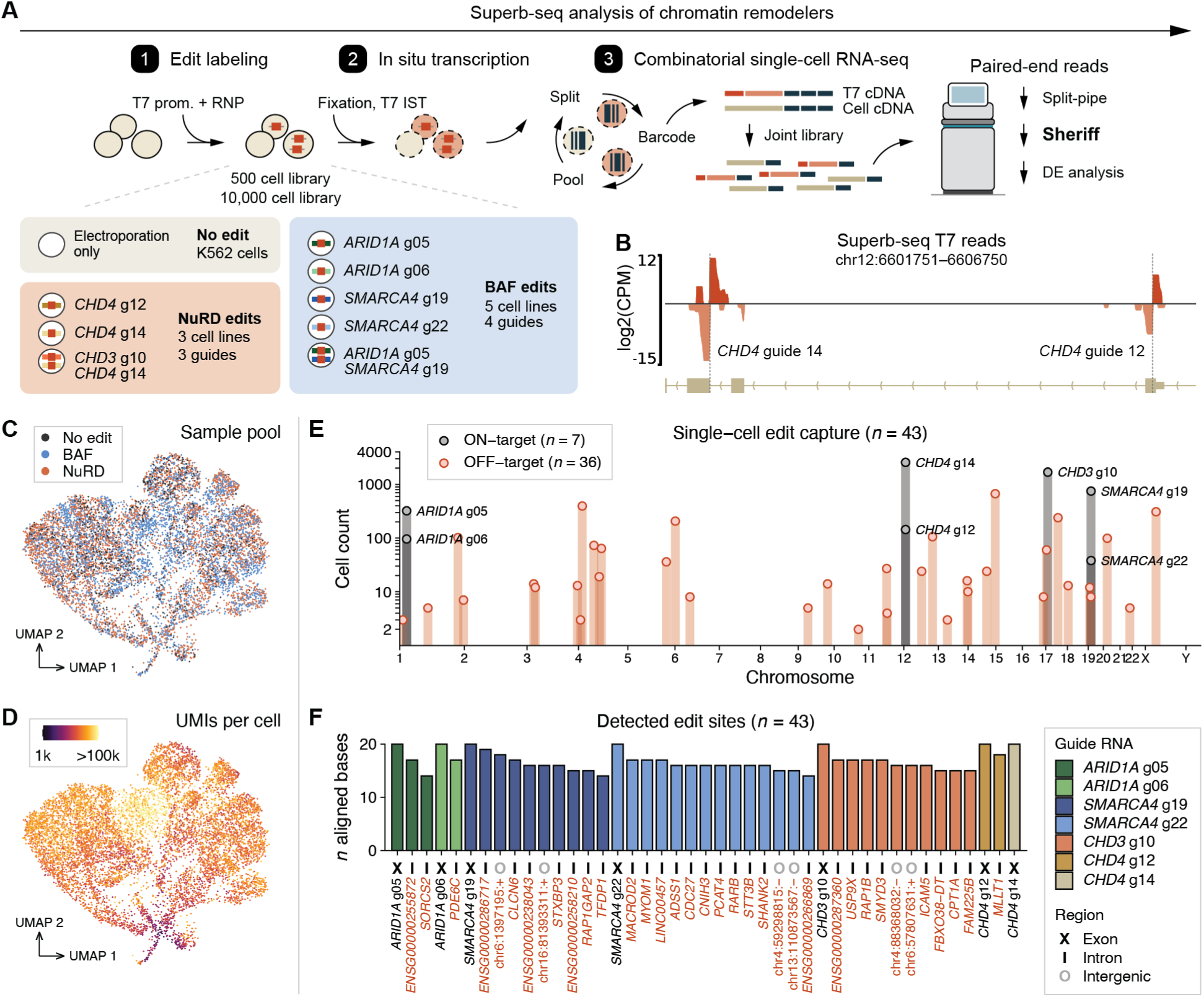
Joint single-cell capture of genome edits and transcriptomes by Superb-seq. (**A**) Superb-seq analysis of four chromatin remodeler genes with seven guide RNAs. In situ transcription and combinatorial indexing were performed on pools of unedited K562 cells, or K562 cells edited at *ARID1A*, *SMARCA4*, *CHD3*, or *CHD4*. Paired-end sequencing reads were analyzed by Split-pipe^40^ and custom Sheriff software. (**B**) Aligned T7 reads identified by Sheriff at the *CHD4* locus. Track shows mean count per million mapped reads (CPM) within 5 bp bins with positive and negative values corresponding to the read orientation. Dotted lines indicate the on-target cut site of each *CHD4* guide. (**C,D**) UMAP embedding of 9,500 K562 cells analyzed by Sheriff, colored by (**C**) sample pool or (**D**) number of unique molecular identifiers (UMIs) per cell. (**E**) Single-cell capture of on- and off-target Cas9 edits, determined by Sheriff analysis of Superb-seq reads. (**F**) Detected edit sites, colored by guide. Edit site names are colored by on-target (black) or off-target (red). Edit intersections with exons (“X”), introns (“I”), or intergenic regions (“O”) are indicated.

### In situ transcription enables direct edit detection by T7 transcript sequencing

For the next step of Superb-seq, we adapted T7 in situ transcription (IST) in fixed cells^27^, reasoning that IST would mark Cas9 edit sites with T7 transcripts, which could be captured alongside endogenous transcripts by fixed cell scRNA-seq. We performed a series of in vitro transcription (purified genomic DNA template) and IST experiments (nuclei or fixed cell template) that established that abundant edit-marking T7 transcripts are generated in SPLiT-seq fixed cells (**Ext. Data Fig. 2, Supp. Figs. 3–6, Supp. Note 3**). Additionally, we performed bulk RNA-seq on IST extracts and found that T7 transcript reads map precisely to expected Cas9 cut sites (**Ext. Data Fig. 3, Supp. Note 4**). Together, these results establish that Cas9 genome edits can be identified by IST and an RNA-seq readout.

**Figure 3.**
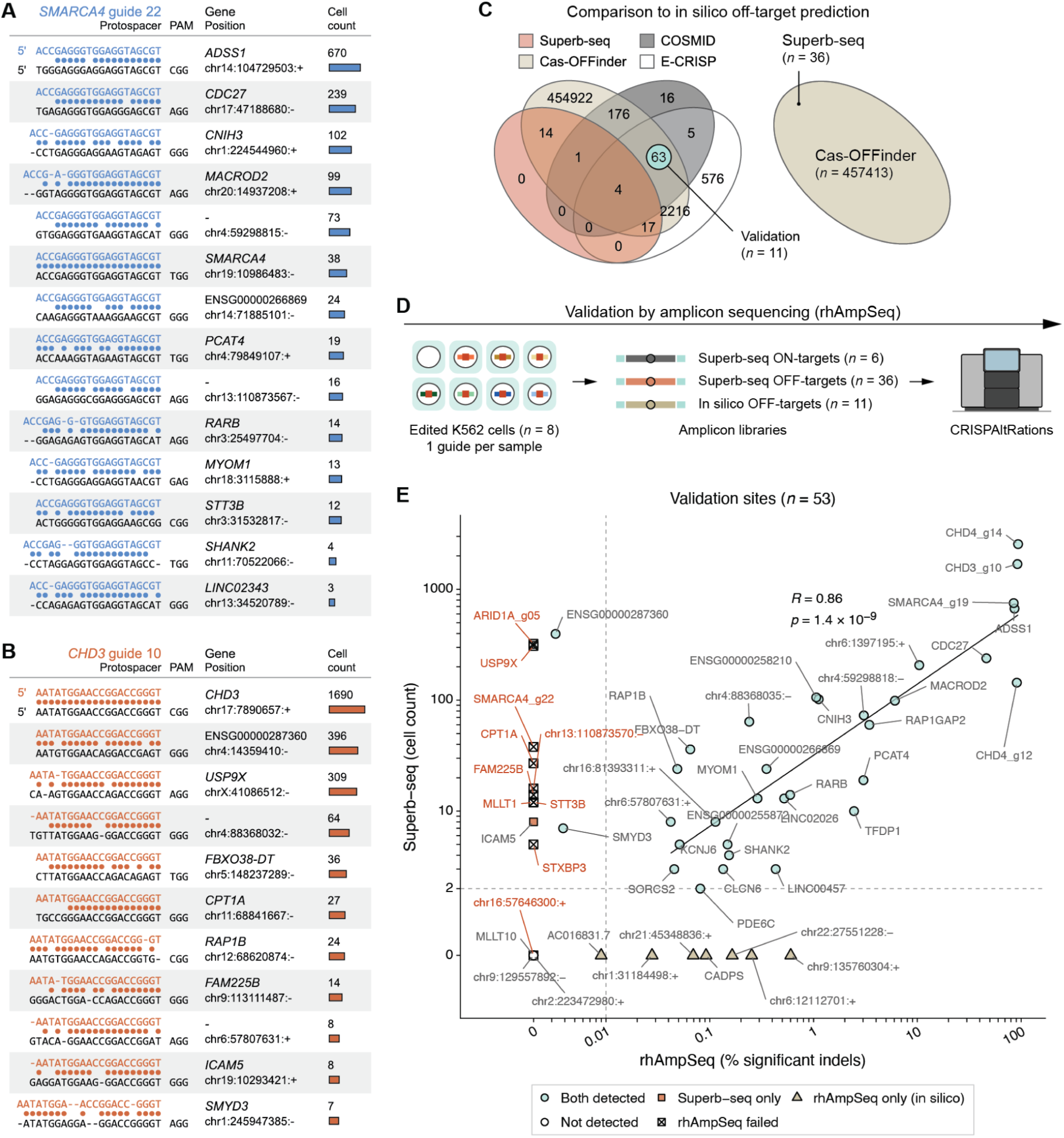
Superb-seq quantifies off-target editing in single cells. (**A,B**) Guide–reference genome sequence alignments and cell counts of Superb-seq detected edits by (**A**) *SMARCA4* guide 22 or (**B**) *CHD3* guide 10, determined by Sheriff. (**C**) Intersection of Superb-seq detected off-targets for the seven chromatin remodeler guides with in silico off-target predictions by Cas-OFFinder^42^, COSMID^43^, and E-CRISP^44^. Top predicted sites were selected for rhAmpSeq validation. (**D**) Validation of Superb-seq detected edit sites and 11 in silico off-targets by rhAmpSeq. (**E**) Association of Superb-seq edited cell counts and indel rates measured by rhAmpSeq. Log-log regression is shown for edit sites with > 2 cells and > 0.01% indels (*R* = 0.85, *p* = 2.9 × 10^−9^) . Red labels indicate Superb-seq edits that could not be confirmed due to sequencing failure (amplicons with < 50 reads in any sample).

**Figure 4.**
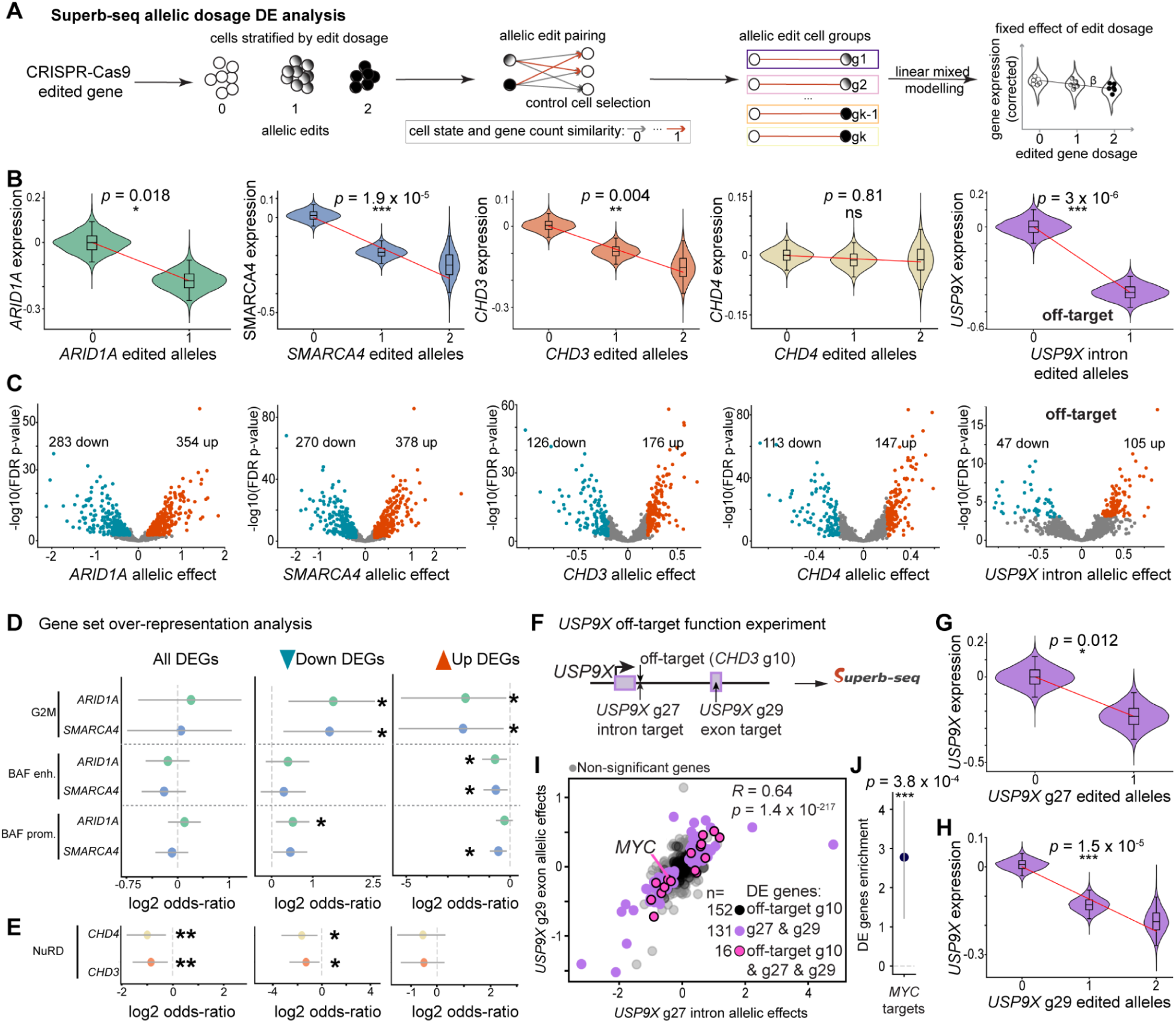
Superb-seq detects differential gene expression associated with on- and off-target edits. (**A**) Overview of the linear mixed model used to call differentially expressed genes (DEGs) associated with edit allelic dosage while accounting for cell state and gene detection rate per cell. (**B**) Violin plots of corrected gene expression from *n* = 10,000 bootstrap iterations stratified by edit allele dosage for each edit site. Lines indicate fitted effect of edit allele dosage on expression. Benjamini-Hochberg adjusted p-values are shown. (**C**) Volcano plots of DEGs associated with edits. (**D, E**) Functional enrichment analysis of BAF perturbations (**D**, *ARID1A* and *SMARCA4* edits) and NuRD perturbations (**E**, *CHD3* and *CHD4* edits), depicted as log2 odds ratios. Y-axis indicates the edit-associated DEG set and query gene set being compared. Query gene sets were hallmark G2M checkpoint genes (G2M), genes with BAF-engaged enhancers (BAF enh.), genes with BAF-engaged promoters (BAF prom.), and genes with NuRD engagement. (**F**) Overview of additional Superb-seq experiment used to confirm edit effect at *USP9X*. (**G,H**) Equivalent to B, for the *USP9X* intron targeting guide (**G**, g27) and the *USP9X* exon targeting guide (**H**, g29). (**I**) Scatterplot of *USP9X* gene expression effect sizes for the intron targeting guide (x-axis, g27) and the exon targeting guide (y-axis, g29). Colors indicate DEG sets including common DEGs between the *USP9X* intron and exon targeting guides (g27 & g29), and common DEGs for all three *USP9X* edit analyses (off-target g10 & g27 & g29). *MYC* is a DEG across all three analyses. (**J**) Enrichment (log2 odds-ratio) of *MYC* targets in the *USP9X* intron and exon DEGs. *p < 0.05, **p < 0.01,***p < 0.001.

**Figure 5.**
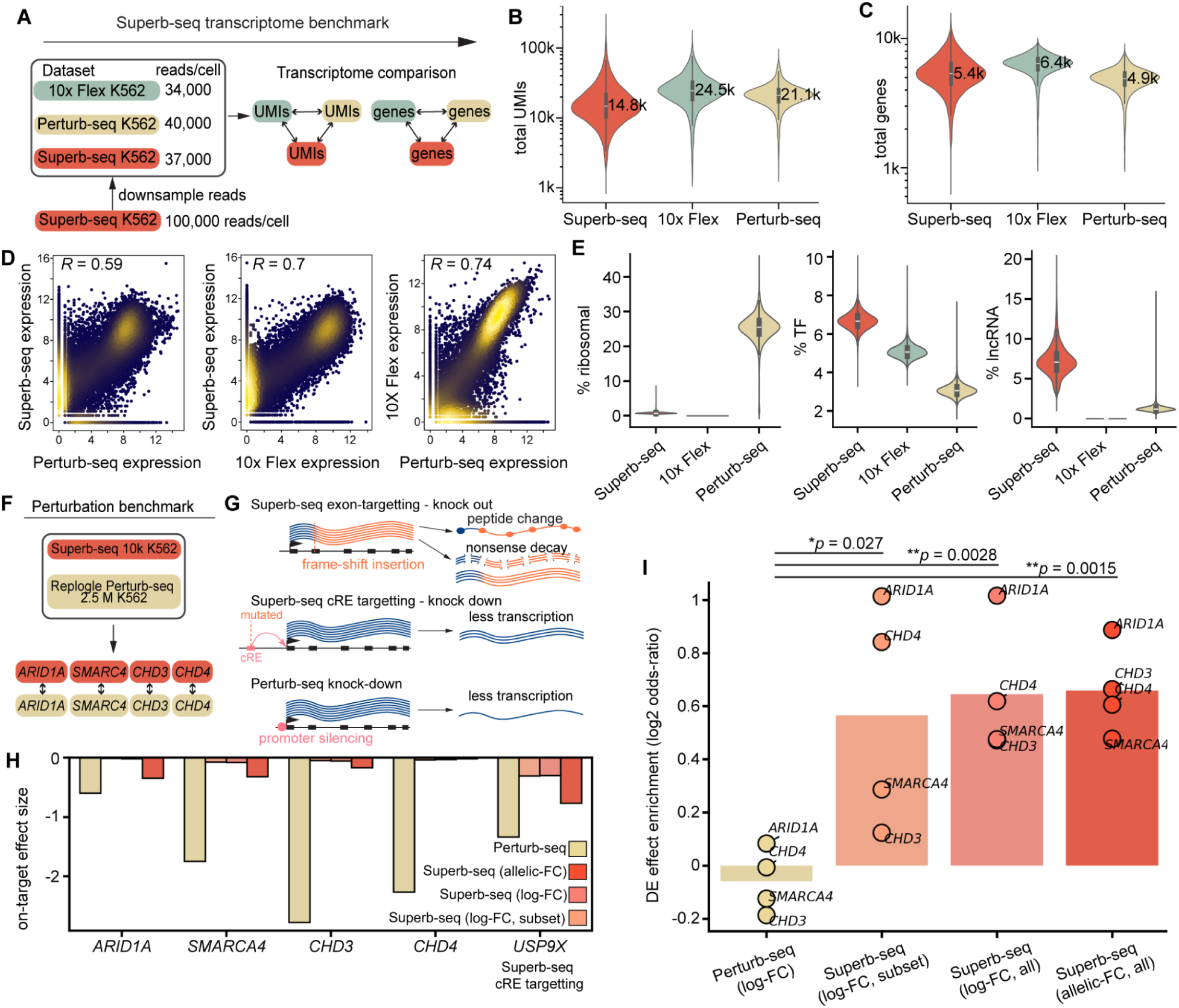
Superb-seq generates high quality transcriptomes and improves estimates of downstream expression effects. (**A**) Schematic of transcriptome comparison between Superb-seq, 10x scRNA-seq and 10x Perturb-seq. (**B**) Violin plots of total UMIs detected per cell in ∼10,000 K562 cells using Superb-seq, 10x Flex, and Perturb-seq. (**C**) Equivalent to B, except for the number of expressed genes detected per cell. (**D**) Pairwise comparisons of pseudobulked K562 gene expression between technologies. Points correspond to genes, colored by density, with Pearson’s *R* indicated. (**E**) Violin plots of the percentage of UMIs in each cell from ribosomal genes, transcription factors (TF) and long non-coding RNAs (lncRNA). (**F**) Schematic of the perturbation benchmark against published Perturb-seq data from 2.5 million K562 cells. (**G**) Schematic of the different mechanisms by which the CRISPR technologies perturb gene expression. (**H**) Gene expression effect sizes estimated from Perturb-seq or Superb-seq using 3 different methods: (log-FC, subset) log fold-change between edited and control cells computed after subsetting cells to the same number of guide-receiving cells detected in the Perturb-seq dataset; (log-FC, all) equivalent, but with all edited cells; (allelic-FC, all) effect sizes estimated from the linear mixed model (Fig. 4A). (**I**) Enrichment (log2 odds ratio) of downstream genes with gene expression perturbation effects in the expected direction.

**Figure 6.**
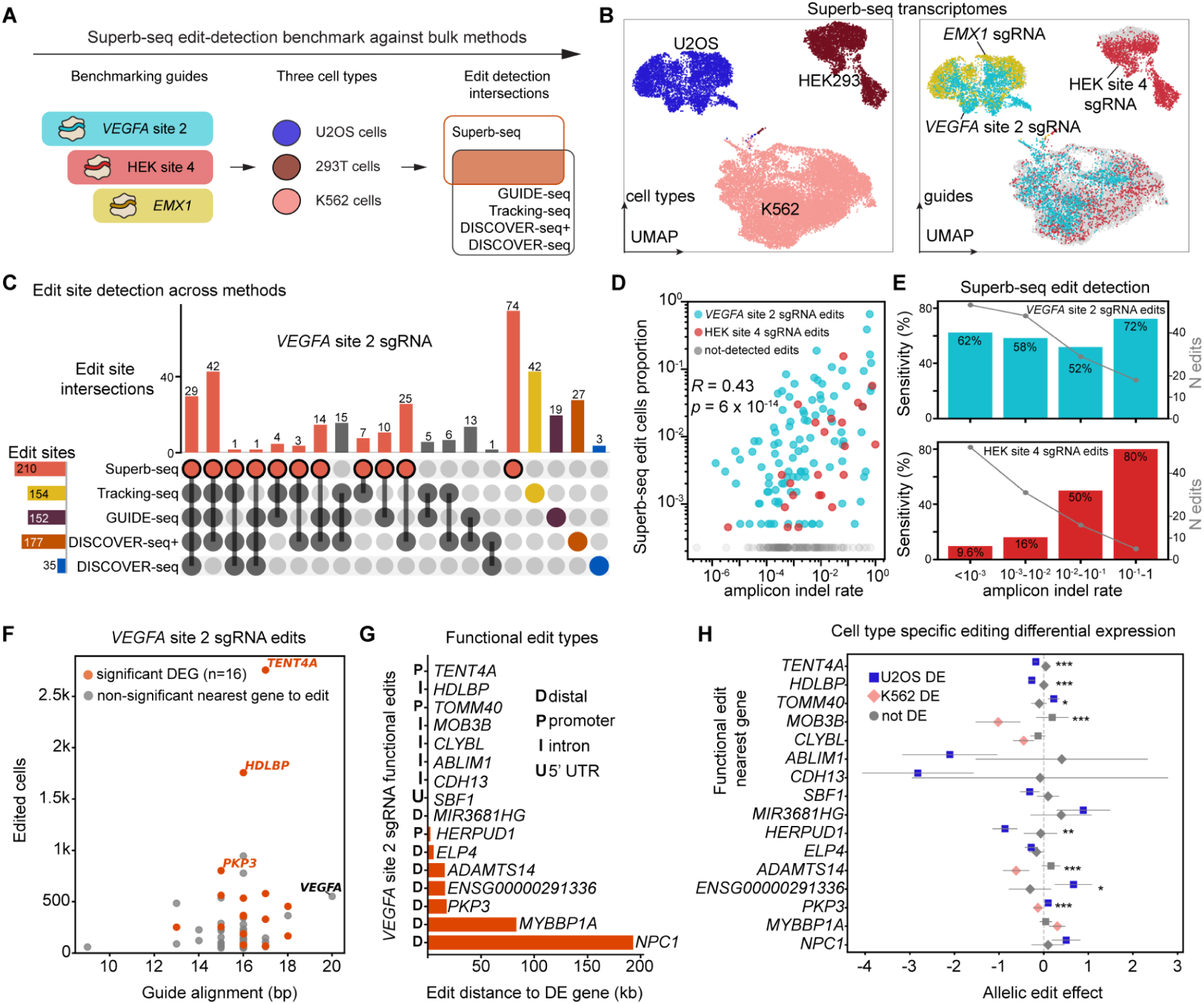
Cas9 edit-detection benchmark and cell type-specific edit effects with Superb-seq. (**A**) Schematic of Superb-seq experiment on three benchmark guides and three cell types. (**B**) UMAPs of Superb-seq single-cell transcriptomes, colored by cell type (left) or guide electroporation condition (right). (**C**) Upset plot of detected edit site intersections between Superb-seq and bulk-based methods for the benchmark guide “*VEGFA* site 2”. (**D**) Correlation between amplicon indel rate and the proportion of edited cells across edit sites for the VEGFA site 2 and HEK site 4 benchmark guides. (**E**) Edit detection sensitivity (bars) and total number of edits (lines) at different amplicon indel rates (x-axis) for the *VEGFA* site 2 (top) and HEK site 4 sgRNAs (bottom). (**F**) Scatterplot of number of edited cells (y-axis) vs. the number of matching basepairs between the edit site genome sequence and the *VEGFA* site 2 sgRNA spacer sequence. "Functional" edits with a significant association to the nearest expressed gene are highlighted. (**G**) Distance between functional edit sites and the associated gene, with labels indicating edit location: promoter (“P”), intron (“I”), 5’ UTR (“U”), or intergenic and distal to the gene body (“D”). (**H**) Cell type-specific effects of edits on the expression of the nearest gene (y-axis). Error bars are 95% confidence intervals. Colored points indicate significant differential expression in the respective cell type. Asterisks indicate significantly different editing effects between cell types (* = *p* < 0.05, ** = *p* < 0.01, *** = *p* <0.001).

### Superb-seq generates high-quality scRNA-seq reads

We next established the complete end-to-end protocol of T7 promoter labeling, IST, and scRNA-seq that we call **“Superb-seq”** (**Fig. 2A**). For scRNA-seq, we chose to use SPLiT-seq^38^, an off-the-shelf method that performs combinatorial indexing of RNA in fixed cells. Since SPLiT-seq uses a blend of oligo-dT and random hexamer primers, it can capture both polyadenylated endogenous transcripts and non-polyadenylated T7 transcripts. We generated K562 cells with T7 promoter-labeled edits using seven guides targeting components of chromatin remodeler complexes BAF (*ARID1A* and *SMARCA4*) and NuRD (*CHD3* and *CHD4*), since perturbing these complexes would result in a large number of transcriptomic changes that could be validated with existing ChIP-seq datasets^39^. We performed IST and SPLiT-seq library generation on unedited and edited K562 cells (**Supp. Figs. 7B,8, Supp. Note 5**), sequenced 500 cell and 10,000 cell libraries, and used Split-pipe^40^ for cell barcode processing and genome alignment. We observed high sequencing coverage and the expected number of cells for each library (**Supp. Fig. 7F**), indicating that the Superb-seq workflow generates high-quality scRNA-seq reads.

### Sheriff software quantifies transcriptomes and single-cell edit alleles at base pair resolution

To quantify single-cell Cas9 edits and transcriptomes from Superb-seq reads, we developed a custom software tool called “**Sheriff**”. Sheriff processes Split-pipe read alignments to output a cell-by-edit allele count matrix, and a standard cell-by-gene count matrix (**Fig. 2A, Ext. Data Fig. 4**). To do this, Sheriff first identifies high-quality T7 reads by the presence of a T7 barcode sequence in the 5’ soft-clipped portion of mapped reads, which is encoded in the inserted T7 promoter (**Fig. 1A, Ext. Data Fig. 4C**). This soft-clipped barcode feature distinguishes edit-marking T7 reads from background T7 reads and endogenous RNA reads. To reduce false-positive T7 read calls, Sheriff accounts for other sources of clipped reads and barcode-like sequences (e.g. sequencing errors). Next, Sheriff calls edit sites by aggregating T7 reads across cells and inspecting edit-specific features (e.g. divergent opposite-strand reads from bi-directional T7 promoter insertions, **Ext. Data Fig. 3B,C, Ext. Data Fig. 4B**). We observed variation in the position and sequence of T7 read alignments from the same cell, indicating multiple edit alleles (**Supp. Fig. 9**). Therefore, Sheriff analyzes this sequence variation to estimate the number of Cas9 edit copies per site per cell, which we term “edit allele dosage” (**Methods**, **Ext. Data Fig. 4F**). Finally, Sheriff corrects transcriptome counts by filtering out low-quality non-barcoded T7 reads from the vicinity of called edit sites (**Ext. Data Fig. 4E**), links off-target sites to their causal guide by sequence alignment (**Methods**), and outputs final count matrices and summary tables (**Ext. Data. Fig. 4G**). Since ambient T7 transcripts were observed in IST supernatants (**Ext. Data Fig. 2H, Supp. Fig. 5**), we additionally estimated and corrected for cross-contaminating ambient T7 reads (**Supp. Figs. 10,11, Supp. Note 6**).

We applied Sheriff to analyze the Superb-seq chromatin remodeler experiment. We observed deep pileups of T7 reads at the expected on-target edit sites of all seven guides (**Fig. 2B, Ext. Data Fig. 5A–D**), with high sensitivity and specificity of T7 read detection (**Ext. Data Fig. 5E**), and high-quality transcriptomes (**Fig. 2C,D, Supp. Fig. 12,13**). On-target edits were detected in 5,598 cells, including at *ARID1A*, where RT-qPCR was insensitive (**Supp. Fig. 6A**). Furthermore, Sheriff resolved Cas9 edit sites at base-pair resolution, positioning each on-target edit 0–3 bp from the expected cut position (**Ext. Data Fig. 5F**, **Supp. Table 1**). The edit allele counts were highly consistent between the higher-depth 500 cell library and the lower-depth 10,000 cell library (**Supp. Fig. 14, Supp. Note 7**).

### Superb-seq detects off-target events in single cells

In the 10,000 cell library, Sheriff not only identified all seven on-target edits, but also identified 36 off-target edit sites in 2,620 cells (**Fig. 2E, Supp. Fig. 15, Supp. Table 3**). These off-targets occurred across 18 chromosomes, were observed in a total of 2098 cells, and varied in frequency from 2 to 2,557 cells per site (0.05%–50% of the edited cell sample). All off-target sites were within non-coding sequences, and > 80% (30 of 36) were within introns (**Fig. 2F**). The high number of detected off-targets was unexpected, since each guide was selected for high predicted specificity and unique sequence match to the intended on-target site (**Supp. Table 1)**. Off-target protospacer sequences exhibited 14 to 19 base matches to the guide sequences (**Fig. 2F**) and a variety of mismatch types (non-canonical base pairing, DNA bulges, RNA bulges. Mismatches occurred across the protospacer, including the “seed” region that strongly influences Cas9 activity^41^ (**Fig. 3A,B, Ext. Data Fig. 6**). Most off-target edit sites were detected at a lower frequency than their corresponding on-target sites (**Supp. Fig. 15**). However, *SMARCA4* guide 22 generated five off-target edit sites that were more frequent than the on-target site (**Fig. 3A, Supp. Fig. 15D**). The most frequent off-target, in an intron of *ADSS1*, was detected in 18% (670 of 3730) of cells in the BAF sample, and the *ADSS1* guide alignment contained four mismatches between the guide and protospacer sequences (**Fig. 3A, Supp. Table 3**). This suggests that guide similarity to the protospacer does not perfectly predict Cas9 efficiency or specificity.

To compare off-targets detected by Superb-seq to those predicted by in silico tools, we applied Cas-OFFinder^42^, COSMID^43^, and E-CRISP^44^ to all seven guides, and relaxed search parameters until all Superb-seq off-targets were found. The in silico tools predicted 458,010 total sites, with the vast majority contributed by Cas-OFFinder (**Fig. 3C**). COSMID and E-CRISP predicted 14% (5 of 36) and 58% (21 of 36) of Superb-seq off-targets, respectively. Only Cas-OFFinder was able to predict all 36 Superb-seq off-targets, but at the cost of very poor specificity (**Fig. 3C**).

### Validation of Superb-seq off-target edit detection by target amplicon sequencing

To validate off-target events detected by Superb-seq, we performed rhAmpSeq, a method for targeted amplicon sequencing of Cas9 editing outcomes^24^. We designed rhAmpSeq primers to target all 36 off-target sites, 6 on-target sites, and 11 in silico off-target sites (not detected by Superb-seq), and performed rhAmpSeq on unedited and edited K562 cells (**Fig. 3D**). Abundant sequencing reads were generated for 81% (43 of 53) of amplicons (**Supp. Fig. 16**), including 27 Superb-seq off-target sites. Aligned rhAmpSeq reads confirmed T7 promoter insertions at off-target sites (**Supp. Fig. 17**) and Superb-seq edited cell counts were highly correlated with rhAmpSeq indel frequencies (*R* = 0.86, *p* = 1.4 × 10^−9^, **Fig. 3E**), indicating accurate edit quantification. One-third (12 of 36) of Superb-seq off-target sites were undetected or underestimated by rhAmpSeq due to sequencing failure or elevated background noise (e.g. mispriming and mismapped reads, **Supp. Fig. 18**), indicating that Superb-seq captures off-targets in difficult-to-amplify genome regions. Multiple in silico sites showed weak indel signals (0.009–0.6%, **Fig. 3E**), suggesting Superb-seq may miss rare off-target events, and that increasing Superb-seq library cell count could be beneficial. For 96% (26 of 27) of analyzed off-targets, rhAmpSeq indels were exclusively observed in cells treated with the inferred guide (**Ext. Data Fig. 7, Supp. Figs. 19,20**), validating the alignment-based identification of causal guides. Only *SORCS2* showed elevated indels from conflicting *SMARCA4* guide 22, suggesting unconfident guide assignment (**Supp. Fig. 20A**). In summary, these results validate that Superb-seq accurately identifies and quantifies Cas9 off-target events.

### Superb-seq associates on- and off-target genome edits with differential gene expression

Next we tested whether Superb-seq can associate genome edits with changes in the cell transcriptome. We developed a linear mixed model to test for associations between edit allele dosage and differential gene expression in Superb-seq data (**Methods, Fig. 4A**). We first applied this model to test whether on-target Cas9 edits affect the expression of the four targeted chromatin remodeler genes. On-target edits to *ARID1A*, *SMARCA4*, and *CHD3* significantly decreased gene expression as edit dosage increased (FDR *p* < 0.05, two-sided Wald test, **Fig. 4B, Supp. Table 4**), consistent with edit-induced NMD of these genes observed by RT-qPCR (**Ext. Data Fig. 1D)**. The change in expression of *CHD4* was not significant (FDR-adjusted *p* = 0.81, two-sided Wald test), consistent with modest *CHD4* decrease by RT-qPCR, possibly due to less efficient NMD (**Ext. Data Fig. 2D**). These results demonstrate that Superb-seq detects the expected effects of on-target Cas9 editing on target gene expression.

We next tested whether frequent intronic off-target edits were associated with differential expression of the genes they were contained within (requiring ≥ 30 edited cells and expression of the gene in K562 cells). Four edit sites met these criteria, within the genes *CDC27*, *CNIH3*, *FBXO38-DT*, and *USP9X*. Among these, off-target editing significantly decreased expression of *USP9X* (FDR-adjusted *p* = 3 × 10^−6^, two-sided Wald test), a deubiquitinase with diverse cellular roles^45^ (**Fig. 4B, Supp. Table 4**). The *USP9X* off-target edit was generated by *CHD3* guide 10 (**Fig. 3B**) and occurred within the first intron, 400 bp downstream of the transcription start site. This position is 2 bp from a candidate K562 enhancer, and intersects a motif and K562 ChIP-seq peak for the transcription factor SP1^39^ (**Supp. Fig. 21**). Since knockdown of SP1 reduces *USP9X* expression^46^, the decrease in *USP9X* expression is likely due to disruption of SP1 binding by the off-target edit. These results show that Superb-seq enables functional characterization of off-target effects, including non-coding edits.

We then tested for associations between Cas9 edits and the expression levels of other, downstream genes. We found that Cas9 editing at all four on-target genes was associated with hundreds of differentially expressed genes (DEGs, **Fig. 4C, Supp. Table 4**). DEGs associated with BAF editing (*ARID1A* or *SMARCA4*) or NuRD editing (*CHD3* or *CHD4*) were enriched in genes regulated by these complexes, determined by ChIP-seq (**Fig. 4D,E, Supp. Tables 5,6,7**). BAF regulates cell proliferation^47,48^ and G2/M cell cycle checkpoint genes, which were also enriched among BAF edit DEGs. These results demonstrate that Superb-seq captures the downstream transcriptomic effects of individual Cas9 edits.

In addition to associations between on-target edits and DEGs, off-target *USP9X* editing was associated with 152 DEGs (47 down-regulated, 105 upregulated), including the oncogene *MYC*^49^. However, since the *USP9X* edits frequently co-occurred with on-target *CHD3* edits (**Supp. Fig. 22E**), it was uncertain if the observed DEGs were caused by *USP9X* off-targeting. Therefore, we generated a second Superb-seq dataset on K562 cells that were edited with 1) a guide targeting the *USP9X* off-target site directly (i.e. perfect sequence match); 2) a guide targeting the first exon of *USP9X*; or 3) *CHD3* guide 10 (**Fig. 4F, Supp. Fig. 23**). All guides efficiently edited their intended target sites, with the *USP9X* intron guide editing 126 cells at the *USP9X* off-target site while reducing *CHD3* editing by 16 fold compared to *CHD3* guide 10 (**Supp. Table 8**). The *USP9X* exon guide edited 1,173 cells at the intended on-target site, with zero off-targets detected (**Supp. Table 8**). As expected, both the intron-targeting and exon-targeting *USP9X* guides decreased *USP9X* expression (**Fig. 4G,H**). The downstream effects on gene expression between the *USP9X* intron and exon edits were well correlated (*R* = 0.64, *p* = 1.4 × 10^-217^), with 131 DEGs associated with edits from both guides (**Fig. 4I, Supp. Table 9**). Of these 131 DEGs, 16 were also discovered in our initial analysis of the *USP9X* off-target edit site, including down-regulation of the oncogene *MYC*^49^ (**Fig. 4I, Supp. Table 9**). *MYC* targets were also significantly enriched in the 131 DEGs (*p* = 3.8 × 10^−4^, OR = 6.9, two-sided Fisher’s exact test, **Fig. 4J, Supp. Table 9**). These results show that Superb-seq captures the downstream gene expression effects of Cas9 on- and off-target edits.

### Superb-seq improves the detection of downstream gene perturbations

To assess Superb-seq transcriptome quality, we compared our 10,000 cell Superb-seq dataset to a 10x Genomics Perturb-seq dataset in K562 cells (5’ CRISPR chemistry), and a 10x Flex scRNA-seq dataset in fixed K562 cells (**Fig. 5A**). These datasets were similar in scale (7,632–11,645 cells), but had lower sequencing depth (34,000–40,000 reads per cell) than our Superb-seq dataset. We therefore downsampled our Superb-seq dataset to 37,000 reads per cell for comparison of transcriptomes (**Fig. 5A**). The downsampled Superb-seq dataset had comparable total UMIs and genes detected per cell (**Fig. 5B,C**). The expression per gene was well correlated across datasets (**Fig. 5D**). Superb-seq showed a stronger correlation to Flex (*R* = 0.70) than to Perturb-seq (*R* = 0.59), and the correlations with Superb-seq were comparable to the correlation between the two 10x Genomics chemistries (*R* = 0.74). We also examined the types of genes captured, such as ribosomal genes which are unfavourable due to extreme abundance and ubiquitous expression, and transcription factors (TFs) and long non-coding RNAs (lncRNAs) which are cell type specific but lowly expressed. Superb-seq had the most favourable capture of these genes, with a low percentage of total UMIs due to ribosomal genes and the highest percentage of TFs and lncRNAs (**Fig. 5E**). These results are consistent with benchmarks comparing commercial SPLiT-seq and 10x scRNA-seq^50^. In summary, the quality of the Superb-seq transcriptome is high and competitive with those from popular scRNA-seq methods.

We next compared perturbation effects detected by our 10,000 cell Superb-seq dataset to those from a large Perturb-seq dataset of 2.5 million K562 cells that included guides targeting the same BAF and NuRD chromatin remodeler genes^6^ (**Fig. 5F**). The Perturb-seq study used promoter-targeting CRISPR interference (CRISPRi, inactive Cas9 fused to KRAB) to repress transcription, while our Superb-seq experiment used frameshifting coding sequence insertions to induce NMD (**Fig. 5G**). Perturb-seq consistently detected larger effects to on-target genes than Superb-seq (**Fig. 5H**). We hypothesize this is because CRISPRi directly represses target gene transcription, whereas NMD permits transcription and instead represses through co-translational degradation (**Fig. 5G**). We next compared the detection of downstream effects on gene expression using target genes of the chromatin remodelers determined from ChIP-seq. Superb-seq showed greater enrichment of downstream genes with effects in the expected direction (i.e., BAF targets downregulated upon BAF perturbation, and NuRD targets upregulated upon NuRD perturbation, **Fig. 5I**). Two factors may contribute to this increased enrichment: differences in perturbation mechanism (transcriptional repression vs. frameshifting insertions), and direct detection of edited cells by Superb-seq, rather than guide detection by Perturb-seq^13^. In summary, despite smaller RNA-level effects on target genes, Superb-seq achieves more sensitive detection of downstream perturbation outcomes.

### Superb-seq edit detection sensitivity is comparable to bulk methods

To further evaluate off-target edit detection by Superb-seq, we performed a “benchmark” Superb-seq experiment in order to and compare results against four bulk off-target profiling methods: GUIDE-seq^22^, Tracking-seq^23^, DISCOVER-seq^51^, and DISCOVER-seq+^52^. For the benchmark experiment, we performed editing using three well-characterized benchmarking guides (*VEGFA* site 2, HEK site 4, and *EMX1*) in the same cell types as prior studies (HEK293 and U2OS), as well as in K562 cells (**Fig. 6A, Supp. Fig. 23**), and generated a 16,000 cell Superb-seq dataset at a sequencing depth of 72,000 reads per cell. Since Superb-seq measures single-cell transcriptomes, we could de-multiplex cell types based on transcriptome differences (**Fig. 6B, Supp. Table 10**).

For the *VEGFA* site 2 guide, Superb-seq detected the most edit sites (*n* = 210, **Supp. Figs. 24,25,26A**), followed by DISCOVER-seq+ (*n* = 177, **Fig. 6C, Supp. Table 11**). Superb-seq detected the highest number of unique edits (*n* = 74, **Fig. C**), but among “high-confidence” edits that were replicated by two or more methods, Superb-seq was the second most sensitive method (*n* = 136) behind DISCOVER-seq+ (*n* = 150 sites). For the HEK site 4 guide, Superb-seq detected fewer edits (*n* = 27, **Supp. Fig. 26B**) than Tracking-seq (*n* = 88) and GUIDE-seq (*n* = 134, **Ext. Data Fig. 8A, Supp. Table 11**). For the *EMX1* guide, only GUIDE-seq was available for comparison, and it detected more edits (*n* = 16) than Superb-seq (*n* = 6, **Ext. Data Fig. 8B, Supp. Fig. 26C, Supp. Table 11**). The proportion of edited cells detected by Superb-seq was well correlated with edit quantification metrics from each bulk method (**Supp. Fig. 27**), and well correlated to published amplicon indel rates from the Tracking-seq study^23^ (*R* = 0.43, *p* = 6 × 10^−14^**, Fig. 6D**). Among the off-targets with amplicon sequencing, Superb-seq achieved high sensitivity of 72–80% for frequent edit sites (>10% indel rate, **Fig. 6E**), but lower sensitivity for rare edits (**Fig. 6E**). However, this likely underestimates Superb-seq sensitivity because the Tracking-seq study did not include sites detected only by Superb-seq. Moreover, Superb-seq sensitivity can be increased by applying the scalable protocol to more cells. In this experiment, we sequenced 2,308–3,832 cells per guide, which limited the probability of sampling the low-frequency edits. Overall, Superb-seq detects Cas9 edits with a detection sensitivity and quantification that is comparable to bulk-based methods.

### Superb-seq identifies cell type-specific effects of on- and off-target editing

We next used Superb-seq to functionally characterize the on- and off-target edit events from the benchmark experiment. Across the three benchmark guides, we tested whether frequent edits (≥50 cells, *n* = 64), were associated with gene expression of the nearest expressed gene. To identify cell type-specific effects on gene expression, we added fixed effects for cell type and cell type × edit interactions to our linear mixed model (see **Methods**). We detected 19 on- and off-target edits associated with differential gene expression (**Ext. Data Fig. 8C, Supp. Fig. 28–30, Supp. Table 12**), of which 16 were generated by the *VEGFA* site 2 guide, and three were generated by the HEK site 4 guide (**Fig. 6F, Ext. Data Fig. 8D**). No differential expression was detected for edits generated by the *EMX1* guide (*n* = 3, **Ext. Data Fig. 8E**).

For the HEK site 4 guide, promoter edits to *DNMT3B* (on-target) and *CEP135* (off-target) were associated with lower expression of the corresponding gene in K562 cells, but not in HEK293 cells. A distal off-target edit (∼260 kb away) was associated with decreased expression of *SEC61G* in HEK293 cells but not K562 cells (**Ext. Data Fig. 8F, Supp. Fig. 28**). Of the 16 expression-associated edits generated by the *VEGFA* site 2 guide, three were substantially more frequent than the on-target edit (**Fig. 6F**). Three edits were in promoters, five were in introns, seven were intergenic and distal, and one was in a 5’ UTR (**Fig. 6G**). Eight of the 16 edits had gene expression effects that were specific to one cell type (**Fig. 6H**), which in many cases reflected cell type-specific expression of the gene (**Supp. Figs. 31,32**). For example, one edit was only associated with the expression of *MOB3B* in K562 cells, because the gene is not expressed in U2OS cells (**Fig. 6H, Supp. Fig. 31)**. One distal off-target edit had opposite effects across cell types, and was associated with increased expression of *PKP3* in U2OS cells and decreased expression of *PKP3* in K562 cells (**Fig. 6H, Supp. Fig. 30**). Overall, this demonstrates that Superb-seq can identify cell type-specific effects on gene expression.

## Discussion

Superb-seq represents several key advances to single-cell CRISPR screening and off-target edit detection. The first advance is that Superb-seq detects edits in individual cells, unlike Perturb-seq methods which detect guides but not edits^5,8^. This enables Superb-seq to remove unedited guide-positive cells that dampen Perturb-seq performance^14^ and improve estimation of gene-regulatory effects (**Fig. 5I**). In addition, Superb-seq can distinguish between the transcriptional effects of on- and off-target editing.

A second advance is that Superb-seq can determine the regulatory consequences of Cas9 off-target events. These off-target effects are an undetected confounding factor in Perturb-seq^6,12^, and a major concern for therapeutic genome editing^20^. In our chromatin remodeler experiment, Superb-seq detected frequent off-target editing of the *USP9X* promoter and downstream dysregulation of *MYC* (**Fig. 4I,J**). Oncogene activation is a major safety concern in gene therapy^53^, however, a downstream effect like this would likely be undetected by off-target analysis pipelines that rely on bulk cell assays. While off-target function can be assessed in single cells using the Tapestri platform^54^, this approach is limited to ∼40 surface proteins, and would not detect dysregulation of downstream oncogenes. Additionally, patient-specific off-target editing is a growing concern for CRISPR-therapies^25^. Multi-donor pooled Superb-seq experiments could help determine how genetic background affects the on- and off-target activity of Cas9 and other genome editors^25^.

A third advance is the optimization of IST for single-cell applications. Integration of T7 IST with scRNA-seq has been used for optical CRISPR screens^55–57^, for detecting epigenetic factors associated with prime editing^58^, and for determining the effects of structural variants^59^. However, these methods have limited cell and read throughput, and require a combination of bulk- and single-cell sequencing workflows. In contrast, Superb-seq achieves scalable joint sequencing of T7 and endogenous transcripts in a single experiment through the use of a strong T7 promoter variant, optimized IST conditions, and blended-primer combinatorial indexing.

A fourth advance is that Superb-seq does not require plasmid or viral vectors, which expands the ability to perform gene perturbation studies in diverse cell types. In addition, Superb-seq is performed with standard laboratory equipment and off-the-shelf supplies which will enable easy adoption of the technology.

A limitation of Superb-seq is that the detection of edits depends on the efficiency of T7 promoter labeling and IST. At some sites, Tracking-seq detected off-target events that were missed by Superb-seq, and vice versa. In our experiments, T7 promoter labeling was associated with the identity of the guide RNA, but not with cell type or genome region (**Supp. Fig. 2**), consistent with the observation that editing outcomes are more strongly associated with the protospacer sequence than cell type or flanking genome context^60^. Future Superb-seq datasets with more guides and adaptation of the Superb-seq workflow for bulk RNA-seq readout could further illuminate factors that affect T7 promoter labeling and improve Superb-seq edit detection.

A limitation of this study is that we did not perform a full Superb-seq experiment in primary cells, such as hematopoietic and liver cell types that are the targets of CRISPR therapies. However, Superb-seq is likely to be feasible in primary cells because the main components of Superb-seq have been applied to primary cells (homology-free labeling, T7 transcription, scRNA-seq), and we achieved efficient T7 promoter labeling in primary human T cells (**Fig. 1E**).

In summary, Superb-seq achieves scalable single-cell sequencing of genome editing outcomes, and can determine the regulatory consequences of on- and off-target edits. We envision that the Superb-seq protocol and companion “Sheriff” analysis software will be broadly useful for functional genomics research, and will enable the development of safer genome-edited therapeutics.

## Methods

### Superb-seq protocol

A detailed protocol for performing Superb-seq experiments and analysis is available on Protocols.io (https://doi.org/10.17504/protocols.io.81wgbwb31gpk/v1).

### Oligonucleotides

Sequences of all DNA and RNA oligonucleotides are listed in **Supp. Table 1**. All oligonucleotides were synthesized by Integrated DNA Technologies (IDT).

### Cell culture

All cell cultures were incubated at 37°C, 5% CO_2_. Human suspension cell lines K562, Jurkat (clone E6-1), and GM12878 were grown in RPMI-1640 (Gibco #11875-093) supplemented with 10% heat-inactivated fetal bovine serum (FBS, GeminiBio #100-500), 100 U/mL penicillin, and 100 µg/mL streptomycin (Gibco #15140-122). Suspension cells were maintained at cell densities of 0.2–1.5 million cells/mL, or 0.1–0.5 million cells/mL for K562 cells. Human adherent cell lines 293FT (Invitrogen #R70007) and U2OS were grown in DMEM high glucose GlutaMAX pyruvate (Gibco #10569-010) supplemented with 10% heat-inactivated fetal bovine serum, 100 U/mL penicillin, and 100 µg/mL streptomycin. Adherent cells were detached by aspirating cell medium, gently washing with 5–10 mL Dulbecco’s phosphate-buffered saline without calcium and magnesium (DPBS, Sigma #D8537-500ML), incubating in 1–3 mL TrypLE (Gibco #12604013) at 37°C until fully detached, then resuspending in the above complete DMEM medium containing 10% heat-inactivated FBS. Adherent cells were maintained below 80% confluency.

For primary human T-cell culture, peripheral blood mononuclear cells were isolated from healthy donor buffy coats by centrifugation on Ficoll-Paque Plus (Cytiva #17144002) or Lymphoprep (StemCell Technologies #07851) density gradient media, and mononuclear fractions were cryopreserved in heat-inactivated FBS with 10% DMSO (Sigma #D8418). Frozen mononuclear cells were thawed and cultured in IMDM with 25 mM HEPES (Gibco #12440-053), supplemented with 10% heat-inactivated FBS, 50 U/mL recombinant human IL-2 (Miltenyi Biotec #130-097-746), 100 U/mL penicillin, and 100 µg/mL streptomycin.

To activate T cells, 6-well plates were coated with anti-human CD3 antibody (clone OKT3, BioLegend # 317326) by adding 1 mL per well of 1 µg/mL anti-CD3 in DPBS, incubating at room temperature for 1 hour, and aspirating. Frozen mononuclear cells were thawed, rested for least 1 hour at 37°C, and seeded in coated plates at 1 million cells/mL in the above T-cell medium supplemented with 1 µg/mL anti-human CD28 antibody (clone CD28.2, BioLegend #302934), then expanded in T-cell medium without anti-CD28.

### T7 promoter design

The 34 base pair (bp) Superb-seq T7 promoter (design 02) was generated as follows. An optimized phage T7 promoter sequence was obtained from the Cel-seq+ method^36^. This sequence is composed of a 20 bp core T7 promoter sequence^61^ (5’-TAATACGACTCACTATAGGG-3’) and flanking 5’ and 3’ sequences that maximize promoter activity^36,62^. To this sequence, we added 2 bp terminal sequences, a 5’ phosphate and two 3’ phosphorothioate modified bases used in the GUIDE-seq “dsODN”^63^ (**Fig. 1A, Supp. Fig. 1C, Supp. Table 1)**.

We designed 26 additional phage promoter constructs by using alternative T7 and SP6 promoter variants and flanking sequences^33–36^, removing flanking sequence, or incorporating dual promoters and alternative polyA or tracrRNA capture sequences^8^ (**Supp. Table 1)**. To maximize gene-disrupting frameshifts upon construct insertion in coding sequence, all constructs were made to length *3n + 1* to yield +1 shifts upon precise insertion and +2 shifts upon common 1-bp templated insertion due to staggered Cas9 cleavage^64–69^. All construct sequences were ordered with the above GUIDE-seq base modifications as pre-duplexed lyophilized oligonucleotides from IDT, reconstituted to 100 µM in electroporation buffer (MaxCyte #EPB-1), and stored at −20°C.

### Labeling of Cas9 edits with T7 promoters

To label Cas9 edits with T7 promoter sequences, we performed homology-free knock-in. Live cells were treated with the Superb-seq T7 promoter and CRISPR-Cas9 ribonucleoprotein (RNP) by electroporation^28^. Single guide RNAs (sgRNAs) were assembled with 20 bp guide sequences, a modified “tracrV2” scaffold sequence^70^, and the common 2’ O-methyl 3’ phosphorothioate modifications on terminal bases^71^. Guides with specificity scores > 0.45 and efficiency scores > 0.3 were selected using GuideScan^72^ or GuideScan2^12^. Guide sequences targeting *TRAC* and *B2M* were previously reported^11,73^. All sgRNAs were synthesized by IDT, reconstituted to 125 µM (4 µg/µL) in electroporation buffer (MaxCyte #EPB-1), and stored at −20°C. All sgRNA sequences are listed in **Supp. Table 1**.

RNP was prepared by mixing 1 µL of 10 µg/µL purified *S. pyogenes* Cas9 protein (IDT #1081059) and 1 µL of 4 µg/µL of sgRNA (2 to 1 sgRNA to Cas9 molar ratio), incubating at room temperature for 15 minutes, and storing on ice until electroporation. Cells were harvested, washed once with 5–10 mL electroporation buffer, and resuspended in electroporation buffer to 100–125 million cells/mL. Per 25 µL electroporation, 5 µL of transfection mix was prepared with 2 µL RNP, 1 µL of 100 µM T7 promoter, 1 µL of 100 µM electroporation enhancer (IDT #1075916), and 1 µL electroporation buffer. Next, 20 µL of cells (2–2.5 M) was mixed with 5 µL of transfection mix, transferred to an OC-25x3 process assembly (MaxCyte #SOC-25x3), and immediately transfected with an ExPERT ATx electroporation system (MaxCyte) using cell type-specific instrument protocols provided by MaxCyte (**Supp. Table 1**). Final electroporation concentrations were 80–100 million cells/mL, 2.5 µM Cas9, 5 µM sgRNA, 4 µM T7 promoter, and 4 µM electroporation enhancer. After electroporation, cells were immediately seeded at 0.5 million cells/mL (K562) or 1 million cells/mL (all other cell types) in a 6-well plate with warm cell culture medium, expanded, and cryopreserved.

### TIDE quantification of T7 promoter-labeled Cas9 edits

On day 3 post-electroporation, genomic DNA (gDNA) was extracted from edited and control cells using a Quick-DNA Microprep kit (Zymo Research #D3020) and eluted in nuclease-free water. PCR amplicons were generated with primer pairs designed by Primer-BLAST^74^ with the following custom parameters: size 900–1200 bp, melting temperature 53–55–57°C (minimum–optimum–maximum), database genomes for selected eukaryotes, organism homo sapiens, max size 30, 40–60% GC, GC clamp of 1, max poly-X of 4, max GC in 3’ end of 4, human repeat filter, and low complexity filter on. Primer pairs were designed to amplify ∼1 kilobase genome regions from ∼150 bp upstream of edit sites to ∼850 bp downstream. Per 20 µL PCR reaction, ∼50 ng gDNA was combined with 0.5 µM of each primer, nuclease-free water, and either Phusion Plus (Thermo Scientific #F631S) or Phusion High-Fidelity (Thermo Scientific #F531S) master mix. PCR reactions were run on a SimpliAmp thermal cycler (Applied Biosystems) with the following protocol: 98°C for 2 minutes, 35 cycles of [98°C for 10 seconds, 60°C for 10 seconds, 72°C for 30 seconds], 72°C for 5 minutes, and then 10°C hold (ramp rate 2°C per second). PCR reactions were purified by ExoSAP-IT Express reagent (Applied Biosystems #75001.200.UL) or by NucleoSpin PCR cleanup kit (Takara #740609.25). Sanger sequencing of PCR amplicons was performed by Eton Bioscience (San Diego, CA) or Genewiz (San Diego, CA), and sequence traces (AB1 files) were analyzed by TIDE (tracking indels by decomposition)^37^ using the batch analysis web tool (https://apps.datacurators.nl/tide-batch/) with the following parameters: left boundary 50, indel size range 50 (−50 to +50), automatic decomposition window, and default p-value threshold (p < 0.001). All primer sequences and automatic web tool parameters are listed in the **Supp. Table 1**.

### Flow cytometry of target protein knockdown

On day 7 post-electroporation, unedited and T7 promoter edited GM12878 cells were harvested into a 96-well round-bottom plate, washed twice with 200 µL cell staining buffer (BioLegend #420201), spun down at 200 × *g* for 2 minutes, flick-decanted, gently vortexed, and stained with 50 µL of a 1 to 200 dilution of PE anti-human B2M antibody (clone 2M2, BioLegend #316305) in cell staining buffer. An isotype control was prepared by staining unedited cells with an equivalent concentration of diluted PE mouse IgG1 κ antibody (clone MOPC-21, BioLegend #400114). Cells were stained in the dark for 15 minutes at room temperature, washed twice with 150–200 µL cell stain buffer, and resuspended in 150 µL cell stain buffer. Forward and side scatter and PE fluorescence intensity were acquired using a BD FACSymphony A3 cell analyzer and HTS attachment. Live (FSC-A/SSC-A) and singlet (FSC-A/FSC-H) events were gated and plotted using R packages flowCore^75^ and ggcyto^76^ (**Supp. Fig. 33**).

### Nuclei isolation

Nuclei buffers were prepared in an RNase-free environment by the Omni-ATAC protocol^77^. A basal buffer of 10 mM Trizma HCl (Sigma #T2194-100ML), 10 mM NaCl (Sigma #S5150-1L), 3 mM MgCl_2_ (Sigma #M1028-100ML), 0.1% Tween-20 (Sigma #11332465001), and 48.75 mL nuclease-free water (Corning #46000CI) was prepared and stored at 4°C. For each nuclei isolation, fresh lysis and wash buffers were prepared. Lysis buffer was prepared with basal buffer supplemented with 0.1% IGEPAL CA-630 (Sigma #I8896-50ML), 0.01% digitonin (Invitrogen #BN2006, stock solution dissolved at 65°C), and 1 U/µL RiboLock RNase inhibitor (Thermo Scientific #EO0382). Wash buffer was prepared with basal buffer supplemented with 1 U/µL RiboLock. Fresh buffers were kept on ice until use.

To isolate human nuclei, cells were harvested, washed once with DPBS, filtered through 40 µm cell strainer (Fisherbrand #22363547), and counted. All steps and spins were performed on ice or 4°C with ice-cold buffers and low-bind tubes. To lyse cells, 2 million cells were lysed by gently resuspending in 45 µL lysis buffer (pipetting up-down exactly 3 times) and incubating on ice for 3 minutes. Nuclei were immediately washed twice by gently resuspending in 250 µL wash buffer, spinning down at 500 × *g* for 5 minutes, manually aspirating, resuspending in 100 µL wash buffer, filtering through a 40 µm Flowmi cell strainer (SP Bel-Art #H13680-0040), spinning down, aspirating, and resuspending in 35 µL wash buffer. Nuclei were counted and imaged by TC20 automated cell counter (Bio-Rad). To generate larger quantities of nuclei for formaldehyde fixation, cell and reagent amounts were scaled proportionally. Aliquots of 0.2–0.5 million nuclei were stored at −80°C.

### Cell and nuclei fixation

Fixation of cells or freshly isolated nuclei was performed in an RNase-free environment with Evercode kits (Parse Biosciences #ECF2001, #ECF2003, #ECF2101, #ECF2103, #ECFC3300). Per fixation, 1–3 million input cells or nuclei were used. Cells and nuclei were spun down at 200–300 × g for 5–10 minutes, according to kit protocol. Yields of fixed singlets was ∼50% of input. Aliquots of ∼0.5 million fixed cells or nuclei were stored at −80°C.

### T7 in vitro transcription

All T7 transcription was performed in an RNase-free environment using the HiScribe T7 Quick RNA synthesis kit (New England BioLabs #E2050S). For in vitro transcription (IVT) on purified gDNA, gDNA was extracted from T7 promoter edited and control cells using a Quick-DNA Microprep Plus kit (Zymo Research #D4074) and eluted in nuclease-free water. Each 20-µL IVT reaction was prepared with 0.5–1 µg gDNA, 10 µL of 2X HiScribe NTP and cofactor buffer mix (10 mM NTP final), 2 µL of 10X HiScribe T7 RNA polymerase (or water for “no polymerase” control), and nuclease-free water, and incubated at 37°C for 2 hour. For IST on unfixed nuclei, 40 µL reactions were prepared by gently mixing 16 µL of freshly isolated nuclei (∼100,000 nuclei) with a master mix of 20 µL of 2X HiScribe NTP buffer mix and 4 µL of 10X HiScribe T7 polymerase (or water), and incubated at 37°C for 2 hours.

### T7 in situ transcription

Prior in situ transcription (IST) methods have primarily used methanol-fixed cells^27,58^. In contrast, off-the-shelf SPLiT-seq uses another proprietary fixative (Parse Biosciences). Therefore, we evaluated compatibility between T7 IST and SPLiT-seq fixation by fixing T7 promoter-edited cells using commercial SPLiT-seq reagents, and performing immediate IST using the above IVT reaction conditions. We observed both T7 RNA at both edit sites and endogenous promoter-like sites (**Supp. Fig. 4E, Supp. Note 3**), indicating that the T7 RNA polymerase was active in these fixed cells. However, lower IST efficiency in fixed nuclei compared to unfixed nuclei from the same isolation suggested that these reaction conditions (37°C, 2 hours) were sub-optimal (**Supp. Fig. 4G,H**).

We systematically optimized IST conditions for commercial SPLiT-seq fixative and identified a specific temperature, time, and NTP concentration (40°C, 24 hours, 10 mM) that greatly improved reaction efficiency (**Supp. Fig. 5A–C**). Under these optimized conditions, levels of T7 RNA surpassed *RPL24* mRNA with minimal increase in background signal (**Ext. Data Fig. 2H, Supp. Fig. 5C**), with similar PCR efficiencies between assays (within 6.0%) estimated by LinRegPCR^78^ (**Supp. Table 2**), indicating surpassing T7 RNA abundance. Importantly, these levels of T7 RNA were observed in the cell pellet fractions of T7 IST reactions, suggesting that abundant T7 RNA is retained within fixed cells. To determine whether unfixed T7 RNA is resistant to washout, we performed IST on fixed cells and washed cell pellets, and then measured T7 RNA levels in post-wash supernatant and pellet fractions. T7 RNA was detected in the supernatant fraction, indicating some washout. However, the pellet fraction maintained a level of T7 RNA equivalent to *RPL24* (**Supp. Fig. 5D**). These results established that unfixed T7 transcripts are retained by fixed cells and suggested suitability for single-cell combinatorial indexing.

For IST on fixed cells (or nuclei), the optimal protocol was to gently add 15 µL of fixed cells (∼100,000 cells) in kit-provided storage buffer to a 25 µL master mix of 20 µL of 2X HiScribe NTP buffer mix (10 mM NTP final), 4 µL of 10X HiScribe T7 polymerase, and 1 µL of 40 U/µL RiboLock (1 U/µL final), and incubate at 40°C for up to 24 hours. This method was used for all Superb-seq samples. During IST optimization, we tested exchanging of fixed cell storage buffer with Omni-ATAC buffer (caused cell clumping), increasing NTP concentration to 20 mM (decreased IST efficiency), and incubating at 37°C or 42°C for 0.5–4 hours (less efficient and less specific, **Supp. Fig. 5**).

### Total RNA purification

Total RNA was extracted from T7 IVT and IST reactions by TRIzol and column purification. TRIzol LS reagent (Ambion #10296010) was added at a 3 to 1 ratio (120 µL TRIzol LS per 40 µL T7 reaction). To extract total RNA from IST pellet and supernatant fractions separately, IST reactions were spun down in a mini centrifuge (∼2,000 × *g* for 1 minute) and split into pellet and supernatant fractions. Pellet fractions were treated with 160 µL TRIzol reagent (Ambion #15596026), and 40 µL supernatant fractions were treated with 120 µL TRIzol LS. Total RNA was purified from TRIzol lysates using a Direct-zol RNA microprep kit (Zymo Research #R2060), with on-column DNase, and eluted in nuclease-free water. For sequencing, an additional 20 µL off-column DNase treatment was performed by mixing 10 µL eluted RNA, 1 µL of 2 U/µL Turbo DNase (Invitrogen #AM2238), 2 µL of 10X Turbo DNase buffer, 7 µL nuclease-free water and incubating at 37°C for 30 minutes before purifying by RNA Clean and Concentrator kit (Zymo Research #R1013) and eluting in nuclease-free water. Purified RNA was quantified by UV spectroscopy (A260/A280) using a DeNovix DS-11+ spectrophotometer, or by calibrated fluorometry using a Qubit 3 fluorometer (Invitrogen) and RNA HS kit (Invitrogen #Q32852) or RNA BR kit (Invitrogen # Q10210). For qPCR, purified total RNA was stored −20°C for up to one week. For sequencing, purified total RNA was stored at −80°C.

### Reverse transcription quantitative PCR (RT-qPCR)

For RT-qPCR of T7 RNA, custom primer pairs were designed using Primer-BLAST^74^ with the following custom parameters: size 80–120 bp, melting temperature 58–60–62°C (min-opt-max), max T_m_ difference of 2, database genomes for selected eukaryotes, organism homo sapiens, max size 30, 40–60% GC, GC clamp of 1, max poly-X of 4, max GC in 3’ end of 4, human repeat filter, and low complexity filter on. To measure on-target T7 RNA levels, primer pairs were designed to target flanking non-exon sequences of T7 promoter edited sites (to distinguish T7 signal from endogenous gene expression). Control primer pairs were designed for several intergenic sequences on different chromosomes, including two “genomic safe harbor” gene deserts (GSH1, GHS2)^79^. Additional primer pairs were designed to flank three sites in the human genome with high sequence similarity to the core 18 bp T7 promoter, discovered by BLAST^80^ (**Supp. Table 1)**. To measure gene expression (cellular mRNA) levels, intron-spanning primer pairs were selected from publications^81–85^ or pre-designed IDT PrimeTime assays.

Two-step RT-qPCR was performed as follows. First, 10 µL first-strand cDNA synthesis reactions were prepared with up to 250 ng total RNA, 2 µL of 5X Maxima H Minus master mix or no RT control mix (Thermo Scientific #M1661), and nuclease-free water, and then incubated on a SimpliAmp thermal cycler using the kit-recommended protocol: 25°C for 10 minutes, 50°C for 15 minutes, and 85°C for 5 minutes, and then 10°C hold. Notably, Maxima cDNA synthesis is primed by both random hexamer and oligo-dT, enabling reverse transcription of both non-polyadenylated T7 RNA and endogenous mRNA. For each 20 µL qPCR reaction, 2–25 ng of RNA-equivalent cDNA reaction was mixed with 0.5 µM of each primer and PowerUp SYBR Green master mix (Applied Biosystems #A25742) by combining 10 µL of diluted cDNA reaction with 10 µL of a 2X mix of 1 µM primers and 2X PowerUp mix. Quantitative PCR was run on a QuantStudio 6 Flex Real-Time PCR system (Applied Biosystems, software version 1.3) with the following protocol: 50°C for 2 minutes, 95°C for 2 minutes, 40 cycles of [95°C for 15 seconds, 60°C for 1 minute], 95°C for 15 seconds, 60°C for 1 minute, and then 0.1°C per second ramp to 95°C for 15 seconds.

Relative expression levels were determined from instrument-calculated cycle threshold (C_t_) values by the comparative C_t_ method^86,87^. Reference assays targeting GSH1 and GSH2 (background signal) were used for IVT and unfixed IST samples. For fixed IST samples, GSH1 and GSH2 signals were below the limit of detection, so assays targeting *RPL24* and *RPS10* mRNAwere used as reference. Results are represented as a difference of C_t_ (dC_t_, normalized to reference only) or a difference of differences (ddC_t_, normalized to reference and no knock-in control).

### Bulk RNA-seq analysis

Jurkat and K562 cells were edited at *CTLA4* or *B2M*, respectively, by treating with either RNP alone or with T7 promoter (**Supp. Table 1**). These four edited cell lines were subjected to nuclei isolation and immediate IST (37°C, 2 hours). An additional two “mock” IST samples without T7 RNA polymerase were generated, one per cell type , and the six total RNA samples were purified and quantified. A pooled, ribosomal RNA-depleted RNA-seq library was made using the QIAseq UPXome RNA library kit (QIAGEN #334782, #334782) and SimpliAmp thermal cyclers. The pooled library contained 12 barcoded samples, with two technical library replicates per total RNA sample. All library preparation steps were performed according to kit protocol, with three modifications. First, cDNA synthesis was performed with 50:50 random hexamer and oligo-dT primers. Second, all bead cleanups were performed with 0.8X QIAseq beads (QIAGEN #333923). Third, library amplification was performed with the following thermal cycler protocol: 98°C for 3 minutes, 18 cycles of [98°C for 30 seconds, 55°C for 10 seconds, 72°C for 40 seconds], 72°C for 5 minutes, and then 4°C hold (ramp rate 2°C per second). Library cDNA was analyzed on a TapeStation 4150 (Agilent) and High Sensitivity D1000 ScreenTape (Agilent #5067-5584) and quantified on a Qubit 3 fluorometer using a Qubit dsDNA HS kit (Invitrogen #Q32851).

All paired-end sequencing was performed at the Salk Institute NGS core facility. The RNA-seq library was run on an Illumina MiniSeq sequencer with a whole Mid Output flow cell, 150/10/10/150 cycles (Read 1, i7 index, i5 index, Read 2), without PhiX. Sample demultiplexing and adapter/poly sequence trimming was performed using Cutadapt^88^. Read quality was assessed by FastQC (https://github.com/s-andrews/FastQC) and MultiQC^89^. Reads were aligned to human reference genome hg38 using BWA-MEM^90^ and post-processed using Samtools^91^. Track visualizations were generated using IGV^92^. See provided code for command details (**Code availability**).

### Superb-seq on chromatin remodeler genes in human K562 cells

A detailed protocol for performing Superb-seq experiments and analysis is available on Protocols.io (https://doi.org/10.17504/protocols.io.81wgbwb31gpk/v1). Three K562 cell samples were prepared and cryopreserved: 1) unedited cells treated by electroporation only, 2) a “BAF” pool of five equally proportioned cell lines edited with guides targeting *ARID1A* and *SMARCA4* and Superb-seq edit labeling, and 3) a “NuRD” pool of three equally proportioned cell lines edit-labeled with guides targeting *CHD3* and *CHD4* (**Supp. Table 1**). Aliquots of each cell sample were thawed, fixed, and frozen. Frozen fixed samples were thawed and put into IST reactions as described above, using ∼100,000 fixed cells per 40 µL IST reaction. Duplicate reactions were performed per pool.

After IST incubation for 24 hours, duplicate reactions were pooled, spun down at 200 × *g* for 5 minutes at 4°C, gently aspirated, resuspended in 40 µL fixation kit storage buffer, and passed through a 40 µm Flowmi cell strainer (SP Bel-Art #H13680-0040) to ensure a single-cell suspension. Singlet cells were counted and immediately subjected to SPLiT-seq combinatorial indexing using an Evercode WT Mini v2 kit (Parse Biosciences #ECW02110, #UDI1001) and KAPA Pure beads (Roche #7983271001) according to kit protocol, targeting 20,000 barcoded cells. Sample proportions were 10% unedited cells, 45% *ARID1A- or SMARCA4*-edited cells, and 45% *CHD3*- or *CHD4*-edited cells. Barcoded cells were used to generate 500 cell and 10,000 (10k) cell sequencing libraries. Libraries were quantified by Qubit 3 fluorometer and dsDNA HS kit (Invitrogen #Q32851), and analyzed by TapeStation 4150 (Agilent) and D1000 ScreenTape (Agilent #5067-5582). Yields were ∼1 µg amplified cDNA library and ∼400 ng per sequencing library, with expected fragment size distributions (**Supp. Fig. 7C,D**).

The 500 cell library was run on an Illumina NextSeq 2000 sequencer with a whole P1 flow cell, 198/8/8/86 cycles, and 5% PhiX, targeting a read depth of 200,000 reads per cell. The 10k cell library was run on an Illumina NovaSeq 6000 with a whole SP flow cell, 98/8/8/124 cycles, and 5% PhiX, targeting 100,000 reads per cell.

### Sheriff processing of Superb-seq reads

A graphical overview of the Superb-seq read processing pipeline is detailed in **Ext. Data Fig. 4**. Briefly, the scRNA-seq FASTQ files were processed using Split-pipe^40^, and hg38 genome assembly and annotation files from Ensembl (https://ftp.ensembl.org/pub/) to generate barcode_headAligned_anno.bam, the Split-pipe BAM file of aligned and annotated reads (**Ext. Data Fig. 4A**). Split-pipe v1.1.1 with default settings and Ensembl release 110 were used for chromatin remodeler Superb-seq analysis. Split-pipe v1.6.0 with default settings and Ensembl release 114 were used for the benchmarking and validation Superb-seq analysis. T7 barcoded reads were identified by a *k*-mer match (*k* = 6) to the Superb-seq barcode in the 5’ soft-clipped sequence of mapped reads (**Ext. Data Fig. 4B,C**). These reads were considered “particular” edit events, due to occurrence in a particular cell (identified by read cell-barcode “CB” tag), the genomic mapping location of the base pair immediately upstream of the soft-clipped sequence (indicating the base-pair resolution repair site of the CRISPR-Cas9 edit event), and the soft-clip sequence (indicating part of inserted T7 promoter sequence transcribed by the T7 promoter). Due to a *k*-mer match between the Superb-seq barcode and the template switch oligo (TSO) sequence, we constructed a set of observed variations in the TSO sequence from the soft-clipped sequences of the reads, which were considered “blacklist” sequences (**Supp. Table 13**). We then matched the Superb-seq barcode to each of the blacklisted TSO sequences to generate a list of problematic Superb-seq barcode *k*-mers. If a read’s soft-clipped sequence contained a match to a blacklist *k*-mer, we then performed *k*-mer matching between the soft-clip sequence and each of the blacklisted sequences. If the read had at least one additional *k*-mer match to the blacklisted sequences, and no additional non-blacklist *k*-mer matches to the Superb-seq barcode, then it was not classified as a barcoded T7 read. After processing the mapped reads, we identified a list of particular edit events, from which we then called “canonical” edit sites.

### Sheriff identification of Cas9 edit sites

After calling T7 barcoded reads to identify particular edit events, we then collapsed particular edit events to canonical edit sites, each of which represents the most common version of an edit across cells. By associating particular edit sites with a canonical edit site, we were able to further filter candidate edit events for false-positives, based on metrics for each candidate edit site obtained from aggregating information across cells.

To call canonical edit events, we first constructed a graph per chromosome using Faiss (https://faiss.ai), where each node is a particular edit site, and edges between nodes represent the base-pair distance between the particular edit sites (**Ext. Data Fig. 4D**). Particular edit sites were then processed in order of the number of cells that have the particular edit, denoted as an ordered set *P* (**Ext. Data Fig. 4D**). The first particular edit in *P*, *P_0_*, is considered the first canonical edit site. All particular edits within a defined base-pair distance (140 bp) from the current canonical edit site (*P_0_* for the first iteration) are considered particular edits of the canonical edit site, denoted *P_c,0_*. The ordered set of particular edit sites is then updated, removing the current canonical edit site and the associated particular edit sites, such that *P = P - (P_c,0_* ⋃ *{ P_0_ })*. This process is repeated at each iteration, until all particular edit sites are associated with a canonical edit site, resulting in the stopping condition *P =* ∅ø (**Ext. Data Fig. 4D**).

Confident canonical edit sites were then determined by the following criteria; 1) the canonical edit site was supported by occurrence in at least three cells, 2) edit-marking T7-barcoded reads mapping in both the forward and reverse direction to the genome, indicating the Superb-seq T7 promoter inserted in either orientation, 3) there were no more than 15 bp between the two closest pairs of forward and reverse particular edit sites (indicating low variation of particular edits around the canonical edit site position), and 4) the canonical edit site did not overlap blacklisted genome regions (**Supp. Table 13**). For criteria 4, the blacklisted genome regions consisted of: 1) 17 genome regions in the unedited sample with excessive read mapping similar to ENCODE blacklist regions^93^, 2) genome locations with sequence similarity to the Superb-seq barcode in both the forward and reverse direction in close proximity identified from the alignment, which when combined with sequencing errors creates soft-clip sequences that resemble Superb-seq barcodes, and 3) simple repeat regions of poly-N sites of length greater than 50 bp, which accumulated reads with soft-clipped sequences resulting in false-positive canonical edit site calls. Excessive read mapping regions were identified by downsampling reads from the unedited sample to 10% of the total reads, and peak calling with Macs2^94^ using macs2 callpeak --broad --nomodel --extsize 98 --keep-dup all --broad-cutoff 0.0000000001 -g hs on the resulting BAM file. Simple repeat regions were downloaded (https://s3.amazonaws.com/igv.org.genomes/hg38/rmsk/hg38_rmsk_Simple_repeat.bed.gz) and subsetted to poly-N sites with length greater than 50bp.

Automated filtering of canonical edit sites as described above identified 63 candidate canonical edit sites from the chromatin remodeler-targeting guides in the 10k cell Superb-seq library. We manually examined the alignments of the called T7 barcoded reads at these edit sites in the Integrative Genome Viewer (IGV)^92^ (**Supp. Material 1**). Twenty of the 63 sites did not appear to be true edit sites, but rather were sites with similarity to the Superb-seq barcode where mismatches between the read and reference genome sequence (e.g., due to sequencing errors or mutations in the K562 genome) resulted in 5’ soft-clipping of the alignment and incorrect edit calls (**Supp. Material 1**). These false positives also differed by other metrics when compared to the final 43 confident edit sites, including specificity of the edit event for cells that were exposed to the same treatment (**Supp. Fig. 34A,B**), the frequency of the canonical edit across cells (**Supp. Fig. 34C**), the best-matching alignment rate to the set of guide sequences (determined below, **Supp. Fig. 34D**), the distance between the best-matching candidate Cas9 edit site based on similarity between guide and PAM sequences, and the called canonical edit site location (**Supp. Fig. 34E**), and the minimum distance between the forward and reverse T7 barcoded reads at the candidate edit site (**Supp. Fig. 34F**). Therefore only the 43 confident edit sites were kept for further downstream analyses.

### Performance assessment of T7 read processing

The sensitivity, specificity, and false discovery rate (FDR) of T7 read calling were estimated as follows. A positive set of T7 reads was approximated by taking all reads within a 200 bp window (± 100 bp) of the expected on-target Cas9 cut site of each guide RNA (**Supp. Table 1)**. A negative set of non-T7 reads was defined as non-edited sample reads, downsampled 100x. Sensitivity (true positive / positive set) was estimated from the proportion of positive set reads that were called T7 reads (true positives). Specificity, 1 – (false positives / negative set), was estimated from the proportion of negative set reads that were called T7 reads (false positives). The FDR (false positives / total positives) was estimated from the proportion of total positives (true positives + false positives) that were false positives.

### Sheriff quantification of single-cell edit allele dosage

At each called canonical edit site, we stratify the particular edits associated with the canonical edit by cell barcode (previously corrected for sequencing error by Split-pipe). To determine if particular edits were generated from different alleles, we performed pairwise comparison of all of the particular edits in each cell (**Ext. Data Fig. 4F**). Two particular edits were considered different if they 1) had a different mapping orientation, or 2) had a different reference genome mapping position. If two particular edits had the same orientation and the same reference genome position, we then examined the 5’ clipped region of their alignments to detect sequence variation within, or flanking, the inserted T7 promoter.

To compare the soft-clipped sequences of T7 barcoded reads, we accounted for several confounding factors that could introduce sequence variation, including read fragmentation during library preparation, the presence of the TSO within the 5’ end of the read and sequencing errors resulting in mis-called bases or incorrect homopolymer runs. We first performed local alignment between the sequences using pairwise2.align.localms with Biopython, with matching score +1, mismatch −1, −0.5 for gap open penalty, and −0.5 for gap extension penalty^95^. We then counted the number of mis-matched base pairs in the aligned sequences until all bases of the shorter soft-clip sequence were seen. This accounts for 5’ read fragmentation, since a mis-match count of 0 will occur if one soft-clip sequence is a subsequence of the other. If the mis-matches were 2 or more, we accounted for the possibility of homopolymer sequencing errors^96^ by re-calculating the mis-matches between the sequences after replacing homopolymer runs of length ≥ 3 with a single instance of the repeated base^96^. If the mis-matches between the two soft-clip sequences was still more than 2, we re-calculated by counting the number of mis-matches in the first 10 base pairs of the aligned sequences starting from the 3’ end of the alignment (to account for sequencing artifacts on the 5’ end of the read), and if this mis-match distance was 2 or less, we considered the particular soft-clip sequences from the T7 barcoded reads to be generated by the same edit events. Hence, the longer of the two soft-clip sequences was added to represent a candidate edit within the cell, and the shorter soft-clip sequence was removed from the list of candidate edits (if present). Alternatively, if the two soft-clip sequences had more than 3 mismatches at any point when comparing two sequences, or had different orientations or reference genome map positions, then both sequences were considered candidate edits. After completing all pairwise comparisons of soft-clipped sequences from T7-barcoded reads that occurred within a single cell, the cell edit allele count was determined from the number of significantly different candidate edits that had been called. This edit allelic dosage calling algorithm is implemented in Python within Sheriff (see **Code availability**).

To call edit allele dosage per gene, gene start and end positions were intersected with canonical edit sites (defined as canonical edit position ± 140 bp). Single-cell gene edit dosages were then determined as the sum of the edit alleles for each cell across all intersecting edit sites of a given gene. The number of called edit alleles was capped to the copy-number for the given gene in K562 cells, determined from deep whole genome sequencing of K562 cells^97^.

### Sheriff quantification of single-cell gene expression

Because the presence of T7 reads can bias estimates of gene expression, Sheriff corrects for T7 transcript read counts at Cas9 edit sites by removing all reads at these sites before performing UMI counting. Sheriff first determines sets of 8-base unique molecular identifiers (UMIs) per cell barcode per gene using “CB”, “GN”, and “GX” read tags from the aligned single-cell reads (annotated BAM file) generated by Split-pipe (**Ext. Data Fig. 4E**). Reads within ± 1000 bp of a canonical edit site across all cells, are then filtered to remove reads that could have been generated by T7 IST (**Ext. Data Fig. 4E**). The UMIs of remaining reads are then used to count the number of unique UMIs per gene per cell. Since sequencing errors can artificially inflate UMI diversity and final UMI counts, UMI sets were reduced to unique sequences based on a Hamming distance of 2 or more bases. To account for cases where one or more UMIs had a Hamming distance > 1 from one another but were within 1 base pair of a common UMI, we constructed UMI sets where within each set there existed a single edit distance path between the UMIs, similarly to previous methods^98^. The number of these sets was then counted as the number of unique molecules for a given gene and cell, effectively accounting for sequencing error and removing confounding non-barcoded reads generated from the T7 polymerase IST reaction (**Ext. Data Fig. 4E**). Normalization was then performed per cell by dividing the UMIs per gene by the total UMIs within the cell, multiplying by the median library size observed across cells, and log-scaling after adding a +1 pseudocount (as implemented in Scanpy)^99^. Scanpy with default parameters was also used to determine a UMAP of the cell gene expression using the cosine metric to construct the k-nearest-neighbour graph. The sc.tl.score_genes_cell_cycle function was used to determine the cell cycle annotations for each cell. Significantly different populations of cells were determined using Cytocipher^100^ with a p-value threshold of 0.01 and initial clusters determined using default Leiden clustering^99,101^.

### Edit to causal guide identification in the chromatin remodeler Superb-seq experiment

For the chromatin remodeler Superb-seq experiments, we determined the causal guide of each canonical edit site as the guide with the highest sequence similarity to the edit site across the 7 guides (*ARID1A* g05, *ARID1A* g06, *SMARCA4* g19, *SMARCA4* g22, *CHD3* g10, *CHD4* g12, and *CHD3* g14). To determine the sequence similarity, we first identified all PAM sites (with the canonical “NGG” or alternative “NAG” sequence^102^) on both the forward and reverse strand within ± 10 bp from each canonical edit site. We then extracted the 20 bp of reference genome sequence upstream of each PAM site as candidate target sequences. Each guide sequence was then aligned to each of the candidate target sequences using Biopython pairwise2.align.globalms, with scoring parameters: +1 for match, −1 for mismatch, −0.8 for gap, and −0.5 for gap extension^95^. The similarity between the guide sequence and candidate target sequences was determined as the number of base-pairs which were aligned. We also scored guides by taking the cosine similarity between the barcoded T7 UMI counts across cells for the guide on-target edit site and each canonical edit site.

### Correction of ambient T7 reads in the chromatin remodeler Superb-seq experiment

To correct for ambient T7 reads in the chromatin remodeler Superb-seq 500 cell and 10k cell libraries (**Supp. Figs. 10,11**), we used the set of edits associated with each guide to determine the guide(s) received by each cell. The sets of guides that were co-electroporated (**Fig. 2A, Supp. Table 1**) were used to determine possible combinations of guides that cells could have been received. The set of edits generated by each guide was used to score guides independently, before using the possible guide sets to determine guide set scores. Because edits had a wide range of detected T7 UMIs, and to account for the fact that more abundant RNA transcripts are likely to be the main sources of ambient RNA^103^, we weighted each edit by the inverse proportion of total T7 UMIs across cells that were found at each respective edit site. For a given guide (*g*) and cell (*c*), the guide score for the cell (*S_g,c_*) was determined from the T7 barcoded UMIs detected at the guide’s edit sites multiplied by the edit weights (*E_g,c_* , where 0 = edit missing, > 0 indicates the weighted T7 UMIs detected across edits in the cell), as follows:

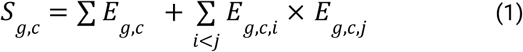

The first term of equation 1 is the weighted sum of the T7 UMIs within cell *c* for edits generated by guide *g*. The second term is the sum of the pairwise product of the edit scores. The second term upweights scores for cells that received a higher number of edits from a particular guide. Single guide scores were then used to calculate scores for each guide set (*G*) for each cell (*S_G,c_*), as follows:

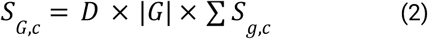

where *D* is an indicator that is set to 1 if all guides for the guide set had a non-zero guide score (*D = 0* if any guides in the set had a zero guide score, *S_g,c_*). The guide set with the highest score was then used to set a guide set label for each cell. Doublet cells were called if the highest guide set score for a given cell was less than 5% higher than the next best guide set score.

We used these initial guide set cell labels to determine foreground and background distributions of guide set scores, so that we could identify cells that did not receive any guides but nonetheless had T7 UMIs due to ambient reads. We fit two negative binomial models using statsmodels.discrete.discrete_model.NegativeBinomial to each guide set’s scores, one for cells initially called as receiving the guide set (foreground), and another for cells called as receiving a different guide set (background). We then calculated the probability of the guide set scores received by each cell using the negative binomial probability mass function, for both the foreground and background fitted distributions. New guide set scores for each cell were then calculated based on the log10 likelihood ratio of the foreground probability against the background probability for each guide set, and cells were allocated as receiving the guide set with the highest log10 likelihood ratio score. Cells where the maximum guide set log10 likelihood ratio score was smaller than −0.225 were considered as receiving no guides (**Supp. Figs. 10,11**).

After calling the received guides per cell, all other T7 barcoded UMIs that were not generated by the called guides were considered ambient reads. This was used to calculate ambient T7 UMIs per edit site, and all of these ambient T7 UMIs were removed from subsequent analyses.

### Comparison of Superb-seq chromatin remodeler libraries

The 500 cell and 10k cell Superb-seq libraries were processed with Split-pipe as described above. Sheriff was then used to process the 500 cell Split-pipe BAM file using the same parameters as the 10k cell library, except that a minimum of 1 cell (instead of 3 cells) was used for canonical edit site calling due to the smaller number of cells in the library. Since some of the Cas9 edit sites detected in the 10k cell library did not have T7 barcoded reads indicating T7 promoter insertion in both orientations in the 500 cell library, we generated a white list of confident Cas9 edit sites (*n* = 43) from the 10k cell library. We then called Cas9 edit sites at these locations in the 500 cell library if any T7 barcoded read was detected within 140 bp of each white listed edit site detected in the 10k cell library. This made it possible to determine the frequency of edited cells for an additional 18 edit sites in the 500 cell library (**Supp. Fig. 14C**). As with the 10k cell Superb-seq library, we observed some false positive edit site calls due to read sequencing errors that caused soft-clipped reads with similarity to the Superb-seq barcode. These were also determined by examining the alignment of barcoded T7 reads as for the 10k cell library. Confident edit sites constituted 30 of 40 called edit sites (**Supp. Table 3**). To compare whether the expected proportion of edited cells was maintained between the 500 cell and the 10k cell library, despite the higher sequencing depth of the former, we plotted the number of edited cells per edit site detected in the 10k cell library, filling in edit sites that were not detected in the 500 cell library with zeroes (**Supp. Fig. 14G**). We also plotted the expected number of edited cells in the 500 cell library against the observed number of edited cells. The expected number of edited cells (*E* [*editedcells*_500*cell*_ ]) was determined by equation 3,

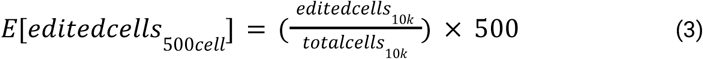

where *editedcells*_10*k*_ indicates the number of edited cells for the respective edit site in the 10k cell library, and *totalcells*_10*k*_ is the total number of cells in the 10k cell library. The same approach was also used to compare the number of edited alleles detected per edit site between Superb-seq libraries (**Supp. Fig. 14H**). After calculating these expected frequencies, we then computed Pearson’s correlation between the expected and observed measurements (for both the estimated number of edited cells and total edited alleles) and the corresponding two-sided p-value using scipy.stats.pearsonr.

### Comparison of Superb-seq and in silico predicted off-targets

The seven chromatin remodeler guides were analyzed with off-target prediction web tools COSMID^43^, E-CRISP^44^ , and command-line Cas-OFFinder^42^. COSMID (https://crispr.bme.gatech.edu/) was run on August 28, 2025 with target genome hg38, PAM “NRG”, and maximally relaxed no indel mismatch of 3, 1-base deletion mismatch of 2, and 1-base insertion mismatch of 3. E-CRISP Evaluation (http://www.e-crisp.org/reannotate_crispr.html) was run on July 31, 2024 with organism “Homo sapiens GRCh38”, 5’ mismatch of 0 to 7, and edit distance of 3. Cas-OFFinder 3.0.0b3 (https://github.com/snugel/cas-offinder/releases/tag/3.0.0b3) was run with the unmasked hg38 genome assembly (https://hgdownload.soe.ucsc.edu/goldenPath/hg38/bigZips/hg38.chromFa.tar.gz), PAM “NRG”, mispair 5, DNA bulge 1, RNA bulge 1. Intersects between Superb-seq and predicted off-target sites were defined as sites occurring within 5 bp between methods.

### Validation of Superb-seq off-target sites by amplicon sequencing

Amplicon sequencing was performed by the rhAmpSeq method^24^. A custom pool of 52 multiplexed rhAmpSeq primer pairs was designed to generate amplicons at: 6 on-target sites and 35 off-target sites (all except *ADSS1*) from the Superb-seq analysis of chromatin remodelers, and an additional 11 in silico predicted off-target sites that were nominated by all three off-target prediction tools and had at least 16 matching bases between the guide and protospacer, but were not detected by Superb-seq (**Supp. Table 2**). To avoid overlapping amplicons, one of the two *ARID1A* on-target sites was omitted. A separate singleplex rhAmpSeq primer pair was designed for the *ADSS1* off-target site, due to the presence of repetitive sequence and common variants that required additional primer optimization. Genomic DNA was isolated from seven K562 cell lines each edited with one chromatin remodeler guide (and T7 promoter, **Supp. Table 1**), and from one unedited (mock electroporation) K562 cell line using a DNeasy Blood and Tissue kit (QIAGEN #13323), according to kit protocol. Sixteen sequencing libraries were generated according to the rhAmpSeq CRISPR Library Preparation protocol (IDT), using 50 ng of gDNA from each cell line, either the multiplex 52-target primer pool or singleplex *ADSS1* primers, and unique rhAmpSeq indexing primers per sample. Raw amplified libraries were combined into one pool of multiplex samples and one pool of *ADSS1* samples, purified using AMPure XP beads (Beckman Coulter #A63880), and quantified by Qubit. After purification, the *ADSS1* pool was added to the multiplex pool at a ratio of 1 to 4 by mass, for a final library composition of 20% *ADSS1* library to avoid dropout. The final rhAmpSeq library was run on an Element AVITI sequencer using one lane of a High Output flow cell, 150/8/8/150 cycles, and 2% PhiX (in parallel with the 20,000 cell benchmark Superb-seq library).

Confirmation of off-target editing at nominated amplicon sites was performed using CRISPAltRations v1.2.1^104^. To identify indels, the window for event quantification was centered on the canonical cut site and events were quantified utilizing the default window size for Cas9 (8 bp). To determine whether indels found in the sequencing data could result from bona fide off-target cleavage, we first grouped indels by location relative to the cut site (prioritizing minimum distance to cut site). We then constructed contingency tables counting the indel events in the edited samples relative to the matched control sample, and performed a hypergeometric test for significance with Bonferroni correction on these tables for each location bin and edit site. For classification of indel off-target editing, the tool requires: 1) sufficient read coverage for the site (>1,000x) in all replicates, 2) significant edits to occur at or adjacent to the cut site after optimal alignment, 3) the classified cumulative significant edits to exceed 0.01%, 4) the comparison of treatment and control samples at the site to have a significant adjusted p-value (p < 0.05), and 5) an average coverage frequency of at least 5x the ascribed cumulative frequency observed (e.g., for 0.1% editing, at least 5,000x coverage). T7 promoter insertion was quantified by aligning > 10 bp insertions from the CRISPAltRations output to the expected T7 promoter sequence using the biopython implementation of the Needleman-Wunsch global alignment algorithm^95,105^. Alignments with scores > 50 were labeled as T7 promoter insertions.

### Differential gene expression analysis for chromatin remodeler edits

To identify associations between edit allele calls and gene expression, we developed an approach that was inspired by Perturb-seq linear models to call single-cell edit effects^3^, but that also allowed for non-linear effects of confounding variables similar to Mixscape^14^. For a given edited gene, we first determine a set of cells consisting of the unedited control cells and all cells with at least one edit allele call, *cells_edited_* , in the condition associated with the given edited gene (e.g. NuRD sample cells if *CHD4* is the edited gene). For each edited cell in *cells_edited_* , an example cell from the unedited control population is selected with the most similar cell state and number of genes detected. Only alleles with more than 25 example cells were considered. The most similar control cell was determined as the first cell with less than 1,000 detected genes different to the query edited cell within the top 10 nearest neighbor cells. The top 10 nearest neighbor cells were determined by Manhattan distance comparing the z-score standardized UMAP coordinates (cell-state) and z-score normalized number of genes detected within the cell. This effectively constructs “allelic pairings” of cell groups (*g*), whereby within each group cells are matched by cell state and genes detected, but differ by gene edit allele dosage.

To call differentially expressed genes with respect to a given edited gene, a linear mixed model was used to model the expression *y_ij_* of query gene *i* in cell *j* , as per equation 4,

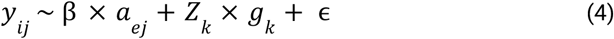

where β is the fixed effect of the number of edit alleles *ɑ_ej_* , of gene *e* in cell *j*, and *Z_k_* is the random effect of cell group *g_k_*. Gene expression was normalized using Sctransform Pearson residuals^106^ prior to the linear mixed modeling. This procedure effectively removed correlation between edit allele detection and the number of genes detected per cell (**Supp. Fig. 35A–C**). However, for some genes (*SMARCA4* in particular), we estimated higher gene expression in cells with 2 edit alleles compared with 1 (**Supp. Fig. 35D**), and in these cells the number of detected genes was substantially higher (**Supp. Fig. 35E**). To account for the systematic gene detection difference between cells with 1 or 2 edit alleles, for edits with > 1 edited allele detected, we used equation 5,

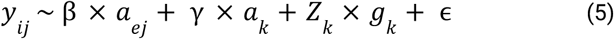

where *ɑ_k_* is the edit alleles of the grouping *k* (i.e., *k* is a shared allele group label applied to both the edit and control cells) and γ is the fixed effect of *ɑ_k_* on gene expression. In this way, control and edited cells are additionally adjusted for the increased gene detection rate that co-occurs with increased edit allele rate (**Fig. 5C, Supp. Fig. 35E**). We used a two-sided Wald test to calculate p-values, and adjusted p-values for multiple testing using the Benjamini-Hochberg method. Genes were called as differentially expressed if the adjusted *p*-value < 0.01 and β > 0.2. This approach enabled calling DEGs across the observed range of gene expression (**Supp. Fig. 22F–J**).

We tested for association between edit alleles and gene expression for edits that were present in ≥ 30 cells and where the gene’s mean expression was > 0.2. In our analysis of the chromatin remodeler Superb-seq experiment, we only tested for associations when the edit was in the body of the corresponding gene (i.e. within exon or intron). By these criteria, we tested for associations between edit allele count and the expression of the four on-target genes (*CHD3*, *CHD4*, *ARID1A*, and *SMARCA4*) and four off-target genes (*USP9X*, *CDC27*, *FBXO38-DT*, and *CNIH3*). For the subset of genes with significant associations (*CHD3*, *CHD4*, *ARID1A*, *SMARCA4*, and *USP9X*), we then tested whether the edit alleles were associated with differential expression of other genes, restricting the analysis to "highly variable" genes, determined using Scanpy with parameters min_mean=0.2 min_disp=0.6 max_mean=5, and NuRD- or BAF-regulated genes in K562 cells (details below), with min_mean=0.03 min_disp=0 for NuRD targets, and min_mean=0.2 min_disp=0.3 for BAF targets.

### Gene set enrichment analysis

We constructed gene sets of NuRD and BAF targets for gene set enrichment analysis (**Supp. Table 5**). NuRD targets were determined by downloading ENCODE hg38 IDR-thresholded *CHD4* ChIP-seq peaks from K562 cells (file accession <u>ENCFF985QBS</u>)^39^. These peaks were then intersected with transcription start sites (TSSs, ± 200 bp from the stranded gene start positions) from Ensembl release 110 gene annotations, to determine NuRD target genes (**Supp. Table 5**). For BAF target genes, we downloaded ENCODE hg38 pseudo-replicated ATAC-seq peaks (<u>ENCFF333TAT</u>) and IDR-thresholded *SMARCA4* ChIP-seq peaks from K562 cells (<u>ENCFF267OGF</u>)^39^. The K562 ATAC-seq peaks and the *SMARCA4* ChIP-seq peaks were then intersected to obtain K562 open *SMARCA4* binding sites. The open *SMARCA4* bindings sites were then intersected with gene TSSs to obtain BAF promoter target genes. Since BAF can also act at enhancers^107,108^, we additionally downloaded hg19 K562 enhancer-gene associations from EnhancerAtlas 2.0^109^, converted coordinates to hg38 using LiftOver^110^, and intersected with the open SMARCA4 peaks. We considered genes associated with the enhancer annotations that overlapped open SMARCA4 ChIP-seq peaks to be BAF enhancer target genes (**Supp. Table 5**).

We tested whether DEGs were enriched within each gene set using two-sided Fisher’s Exact Test and used a Benjamini-Hochberg FDR significance threshold of 5% to account for multiple testing (**Supp. Tables 6,7**). Odds ratios and associated two-sided 95% confidence intervals were calculated from the contingency tables of overlaps between DEGs and gene sets (**Supp. Table 6**) using the conditional maximum likelihood estimation method.

### Comparison of Superb-seq and Perturb-seq transcriptomes and perturbation effects

UMI count matrices for the 10x Perturb-seq and 10x Flex scRNA-seq from K562 cells were downloaded from: 10x Perturb-seq: https://www.10xgenomics.com/datasets/10k-k562-transduced-with-small-guide-library-5-ht-v2-0-chrom ium-x-2-standard 10x Flex: https://www.10xgenomics.com/datasets/10k-human-k562-r-cells-singleplex-sample-1-standard These count matrices were combined with the Superb-seq 10k cell K562 count matrix, and genes detected in less than 3 cells and cells with less than 1,000 total UMIs were excluded from further analysis. This resulted in a count matrix with 28,254 cells and 43,772 genes for comparison. This unified count matrix was used to quantify the total UMIs and genes detected per cell. Before comparison of transcriptome similarity, UMIs across all cells for each dataset were summed (pseudobulked) and normalized by sequencing depth. Sequencing depth normalization was performed by dividing the UMIs of each gene by the total UMIs of each dataset, and multiplying by the median total UMIs observed across the datasets. Library size normalized gene expression values were then log10 transformed with a +1 pseudocount. Pearson’s correlations for these normalized gene expression values between datasets were computed with scipy.stats.pearsonr.

The UMI count matrix and cell guide annotations for the Replogle Perturb-seq dataset^6^ were downloaded from Figshare (https://plus.figshare.com/ndownloader/files/35775507). The UMI counts were subsetted from the full 1,989,578 cells and 8,248 genes matrix to only cells that received guides targeting *SMARCA4*, *ARID1A*, *CHD3*, *CHD4*, *USP9X* or non-targeting control guides. This resulted in 76,225 cells for further analysis (132 *SMARCA4* cells, 271 *ARID1A* cells, 174 *CHD3* cells, 89 *CHD4* cells, 231 *USP9X* cells, 75,328 non-targeting control cells). Normalization was performed by dividing the UMIs detected per gene by the total UMIs detected per cell, multiplying by the median observed total UMIs across cells, followed by log10 transformation with a +1 pseudocount. The log2 fold change of the guide-receiving cells was calculated against the non-targeting control cells using scanpy.tl.rank_genes_groups. Genes tested in the Superb-seq chromatin remodeler differential expression analysis were intersected with expressed genes in the Replogle dataset and the intersecting gene set was used to compare the two datasets. Genes were classified as up- or down-regulated based on the log2 fold change against the non-targeting guide control cells. For the BAF gene perturbations (*SMARCA4*, *ARID1A*), the target-enrichment was quantified as the odds ratio for the intersection between the down-regulated genes and BAF target genes determined from *SMARCA4* ChIP-seq (detailed above). For the NuRD gene perturbations (*CHD3, CHD4*), the target-enrichment was quantified as the odds ratio between the up-regulated genes and the NuRD target genes determined from *CHD4* ChIP-seq (detailed above).

For Superb-seq, perturbation effects were quantified using three approaches, either to utilize the allelic edit information or control for different numbers of cells between the datasets. In the "Superb-seq logFC-all" approach, we computed log2 fold change between cells that had target gene editing and no-edit control cells. In the "Superb-seq logFC-subset" approach we also computed log2 fold change, but subset the edited Superb-seq cells to match the number of edited cells in the Perturb-seq dataset for each gene (there were more edited cells for each target gene in the Superb-seq dataset). To account for sampling variance, we repeated the downsampling 10 times, and recorded the median log2 fold change for each gene across re-samplings. In the "Superb-seq allelicFC-all" approach we estimated effects of allelic edit count on gene expression and called significant associations between edits and gene expression using our linear mixed model.

### Superb-seq on benchmarking guides and *USP9X* validation guides

We prepared and cryopreserved 12 cell lines (**Supp. Table 1**): unedited 293FT (HEK293) , U2OS, and K562 cells treated by electroporation only; cells with T7 promoter editing using benchmarking guides *EMX1*, HEK site 4, and *VEGFA* site 2^22^ (**Supp. Table 1)** and one or two cell types per guide; K562 cells treated with T7 promoter only; K562 cells with T7 promoter editing using *CHD3* guide 10, *USP9X* guide 27, or *USP9X* guide 29. Aliquots of each cell line were thawed, pooled by guide or cell type into 10 samples, fixed, and stored at −80°C. Fixed samples were thawed and put into duplicate IST reactions. After IST incubation for 24 hours, duplicate reactions were pooled, spun down at 300 × *g* for 5 minutes at 4°C, gently aspirated, resuspended in 40 µL fixation kit storage buffer, and passed through 40 µm cell strainers (Fisherbrand #22363547). Singlet cells were counted and immediately subjected to SPLiT-seq combinatorial indexing using an Evercode WT Mini v3 kit (Parse Biosciences #ECWT3100, #UDI1001) and SPRIselect beads (Beckman Counter #B23317) according to kit protocol, targeting 20,000 barcoded cells. Samples were proportioned evenly, 8.3% per condition per cell type. Barcoded cells were used to generate two 10,000 cell libraries. Libraries were quantified by Qubit and analyzed by TapeStation and D1000 high sensitivity ScreenTape (Agilent #5067-5584). Yields were ∼2 µg amplified cDNA library and ∼400 ng of sequencing library of the expected fragment size distribution for Split-seq libraries, similar to the chromatin remodeler libraries (**Supp. Fig. 7C,D**).

The two libraries were pooled and run on Illumina NovaSeq X Plus and Element AVITI sequencers, to a target read depth of 100,000 reads per cell. Two sequential NovaSeq runs were each performed with one lane of a 10B flow cell, 100/8/8/122 cycles, and 5% PhiX. The AVITI run was performed with one lane of a High Output flow cell, 150/8/8/150 cycles, and 5% PhiX.

### Superb-seq benchmark library preprocessing and edit site quality control

FASTQ files from the three sequencing runs were jointly processed using Split-pipe^40^ (**Supp. Fig. 7F**) to generate output BAM files. Cell barcodes were prefixed by the library number to ensure non-overlapping cell barcodes across libraries, prior to merging the BAM files with Samtools. Sheriff was used to process the merged annotated BAM file, using the same approach as for the 10k cell chromatin remodeler library described above, but with edit_site_min_cells set to 1 and Ensembl Release 115 hg38 used as the GTF input file of genome annotations.

Quality control of putative edit sites called by Sheriff was first performed by counting the total proportion of all edit alleles at each putative edit site that were from cells specific to each edit sample (*CHD3* g10, *USP9X* g27, *USP9X* g29, *VEGFA* site 2 guide, HEK site 4 guide, *EMX1* guide). The read alignment of called T7-barcoded reads at putative edit sites with a maximum edit*-*sample proportion greater than 50% were manually examined for sequencing errors and the characteristic Superb-seq signal (**Fig. 2B, Ext. Data Fig. 5A–D**). Putative edit sites that were false-positives, evident from sequencing errors in the 5’ soft-clip sequence and present across few cells, were removed from the curated edits (**Supp. Tables 8,10**). The curated edit sites were further validated by aligning the guide protospacer sequences for the corresponding sample of the edited cells to each 20 bp reference sequence adjacent to PAMs within 30 bp of the called canonical edit site. Alignment was performed using the biopython implementation of global alignment with +1 for matches, −1 for mis-matches, −3.05 for gap opening, and −0.5 for gap extension. For the *USP9X-*related guides, this revealed protospacer sequences with high sequence similarity (15–20 bp matches) with expected cut sites 0–1 bp from the called canonical edit sites (**Supp. Table 8**). For the benchmark guides, the median sequence similarity for protospacer sequences and guide spacer sequences across the > 200 sites was 16 bp, with expected cutsites ≤ 2 bp from the called canonical edit sites for all except 8 sites (**Supp. Table 10**).

### Validation analysis of *USP9X* off-target edit effects

*USP9X* differential expression was called with respect to on-target edits generated by the *USP9X* intron 1 (g27) and *USP9X* exon (g29) guides, using the same approach described previously for the chromatin remodeler guides. The K562 unedited sample was used as the control sample, and p-values were adjusted for multiple hypothesis testing using the Benjamini-Hochberg false discovery rate method. Downstream differentially expressed genes were called with respect to each *USP9X* edit across the same set of genes that were tested for the chromatin remodeler edits, with p-values adjusted using the Benjamini-Hochberg method. Three sets of differentially expressed genes were intersected: DEGs from the *USP9X* intron edit induced by *CHD3* g10, DEGs from the *USP9X* intron edit induced by *USP9X* g27, and DEGs from the *USP9X* exon edit induced by *USP9X* g29. This identified 16 DEGs in common between all three DEG sets, which included the oncogene *MYC*. *MYC* target gene perturbation enrichment was assessed using a Fisher’s exact test for the intersection between tested *MYC*-target genes and the 131 DEGs in common between the *USP9X* g27 intron editing and the *USP9X* g29 exon editing (**Supp. Table 9**).

### Benchmarking of Superb-seq edit detection

Benchmark on- and off-target Cas9 edit detection results were downloaded from supplemental tables of the respective publications for Tracking-seq^23^, GUIDE-seq^22^, DISCOVER-seq^51^, and DISCOVER-seq+^52^. Edit site coordinates for GUIDE-seq, DISCOVER-seq, and DISCOVER-seq+ were converted from hg19 to hg38, using the Liftover Python package (https://github.com/jeremymcrae/liftover). To account for differences in the conventions for reporting edit site coordinates (coordinates as either inferred cut site, PAM, or start of protospacer sequence), edit sites were harmonized by identifying sets of edits that were within 10 bp. This was achieved by first constructing an adjacency matrix of base-pair distance between all edit sites, and binarizing this matrix to 1 for edit sites within 10 bp, and 0 if more distant. A depth-first search on this binarized adjacency matrix was then used to find all sets of edits that were connected in this matrix. Between the sets of harmonized edits, the smallest observed distance was 199 bp, confirming accurate grouping of distinct edit sets. The “edit_id” in the provided edits across technologies identifies these edit sets in **Supp. Table 11**. These harmonized edits were used to construct all upset plots displaying intersections of edits detected across technologies per guide.

For edits quantified by amplicon sequencing in the Tracking-seq study, the mean amplicon indel rate across three replicates was used for additional benchmarking of Superb-seq edit site quantification and detection sensitivity. To compare the mean indel rates to Superb-seq, edits that were uniquely detected in Superb-seq were removed, and zero was used as the Superb-seq quantification for amplicon edits that were not detected. For detected edits, the proportion of edited cells for the corresponding cell type within the edit sample was used to quantify the Superb-seq editing rate. Across the amplicon-sequenced edit sites, the mean indel rate and these Superb-seq quantifications were correlated (**Fig. 6D**). To quantify edit detection performance, high confidence edits were defined as those with a mean indel rate greater than 0.01%, consistent with the Tracking-seq study^23^. Edits were then binned by indel rate, and the proportion of detected edits for the guides with amplicon-sequencing data detected by Superb-seq was used to quantify sensitivity at different editing frequencies (**Fig. 6E**).

### Cell type-specific differential expression analysis for benchmark edits

For each edit site detected by Superb-seq across the benchmark guides, the allelic edit dosage matrix was filtered for high-quality T7 reads by identifying cells with edits from multiple guides, and setting the T7 UMI count of all edits from untreated guides to 0. The maximum allelic edit dosages for each edit in each cell type was capped to the copy number of the corresponding genome segment for the cell type, determined from the Cancer Cell Line Encyclopedia (CCLE)^111^. Absolute copy number calls from CCLE were utilized for K562 and U2OS cells, while the relative segmented copy number calls from CCLE were used to estimate absolute copy numbers for each segment for HEK293 cells. Relative copy numbers were reported as the log2-transformed relative copy ratio for segments. Absolute copy numbers were estimated as 2 × 2^relative^ ^copy^ ^number^ rounded to the nearest whole number, capping the lower limit to 1 and upper limit to 6. Edits detected in ≥ 50 cells were kept for differential expression testing. The nearest expressed gene to each edit site was then determined. Expressed genes were defined as being detected in more than 30% of cells for a given cell type. For each edit, the nearest expressed gene was found using the PyRanges package^112^ nearest function on the Ensembl release 115 hg38 GTF file, subsetted to the genes expressed across any cell type that received the edit. Gene expression was normalized using Sctransform^106^ Pearson residuals prior to differential expression testing.

The cell type-specific differential expression analysis was performed utilizing a similar approach as for the chromatin remodeler edits, with adjustments to the allelic pairing strategy to account for cell type and with additional cell type terms added to the linear mixed model. The allelic pairing strategy was modified so that control cells were chosen from the corresponding cell type no-edit control samples. The additional terms in the linear mixed model included a term to account for differences in baseline gene expression between cell types (*c*) and an interaction term to account for cell type-specific editing (*c* : *ɑ_ej_*), as shown in equations 6 and 7,

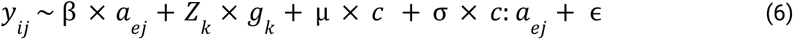

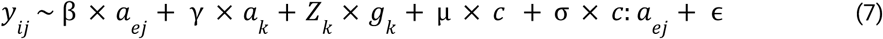

where equation 6 and equation 7 are equivalent to equations 4 and 5, respectively, with the additional cell type term (µ × *c*) and cell type/edit interaction term (σ × *c* : *ɑ_ej_*). The additional coefficient µ adjusts the baseline expression of the gene depending on the cell type, and σ corresponds to an additional change in gene expression if editing occurred in a different cell type from the baseline cell type. For edits from the *VEGFA* site 2 guide, the baseline cell type (*c* = 0) was encoded as K562 (*c* = 1 for U2OS). For edits from the HEK site 4 guide, the baseline cell type (*c* = 0) was encoded as HEK293 (*c* = 1 for K562). The *EMX1* guide was only electroporated into U2OS cells, so the additional cell type terms were not included for these edits. After fitting the linear mixed models described above, a Wald-test was used to determine p-values for the edit coefficient for the baseline cell type (β). For the non-baseline cell type, the editing effect size was determined as the sum of the baseline cell type edit effect size and the additional edit effect size from the interaction term (β + σ). A Wald-test applied to the sum of these coefficients was used to determine the p-value for the non-baseline cell type. To determine if the cell type had significantly different editing effects between cell types, a Wald test was also applied to the interaction effect size coefficient (σ). The estimated effect sizes per edit and cell type are provided in **Supp. Table 12**.

## Data availability

All sequencing datasets is deposited in the NCBI GEO repository^113^ as a SuperSeries (accession number <u>GSE314191</u>) containing SubSeries for Superb-seq (<u>GSE284207</u>), rhAmpSeq (<u>GSE313955</u>), and bulk RNA-seq experiments (<u>GSE313958</u>). The following K562 cell datasets were downloaded from the ENCODE data portal (https://www.encodeproject.org)^39^: SMARCA4 ChIP-seq (accession number <u>ENCFF267OGF</u>), CHD4 ChIP-seq (<u>ENCFF985QBS</u>), and ATAC-seq (<u>ENCFF333TAT</u>).

## Code availability

Sheriff is available on GitHub (https://github.com/BradBalderson/Sheriff). This software package quantifies single-cell CRISPR-Cas9 edit allele dosage and gene expression from Superb-seq datasets. Code for additional Superb-seq analyses, bulk RNA-seq, and data visualization in this work are also available on GitHub (https://github.com/BradBalderson/superb_analysis).

## Acknowledgements

This work was supported by NIH-NHGRI grant HG011315 to G.M. and the Frederick B. Rentschler Developmental Chair to G.M. B.B. was supported by a Hewitt Foundation Fellowship and an Ekhart Scholarship.

This work was also supported by the The Razavi Newman Integrative Genomics and Bioinformatics Core Facility of the Salk Institute (RRID:SCR_014842 and SCR_014846) with funding from NIH-NCI CCSG P30 CA014195, NIH-NIA San Diego Nathan Shock Center P30 AG068635, the NIH-NIA Liver Cancer P01 AG073084-04, the Howard and Maryam Newman Family Foundation and the Helmsley Trust. Raw sequencing data was prepared by Nina Tonnu and April Williams.

This work was also supported by the Single Cell & Spatial Omics Core Facility of the Salk Institute with funding from NIH-NCI CCSG P30 CA014195, NIH-NIA San Diego Nathan Shock Center P30 AG068635, the Chapman Foundation and the Helmsley Charitable Trust. Elsa Molina, Ling Ouyang, and Tzuwen Wang assisted in planning and coordinating sequencing runs.

This work was also supported by the Flow Cytometry Core Facility of the Salk Institute (RRID:SCR_014839) with funding from NIH-NCI CCSG: P30 CA01495, and Shared Instrumentation Grants S10-OD023689 (Aria Fusion cell sorter), and S10 OD034268 (Thermo Fisher Bigfoot).

We thank Diana Hargreaves, Alex Jones, and Helen McRae for assistance with selection of BAF and NuRD complex genes, and for advice on determining BAF-regulated genes.

## Author contributions

M.H.L. and B.B. contributed equally. M.H.L. and G.M. conceived of the project. M.H.L. designed, performed, and analyzed experiments, developed software, made figures, and wrote the manuscript. B.B. analyzed experiments, designed algorithms, developed software, made figures, and wrote the manuscript. Under direction of M.H.L., K.S. performed and analyzed experiments. Under direction of B.B., A.J.H. developed software. M.L. (Madelyn Light) designed the rhAmpSeq primers. S.N.L performed the rhAmpSeq experiment. Under direction of G.K, O.H.U. analyzed rhAmpSeq data. G.M. supervised the project, designed and analyzed experiments, and revised the manuscript.

---

**Extended Data Figure 1.**
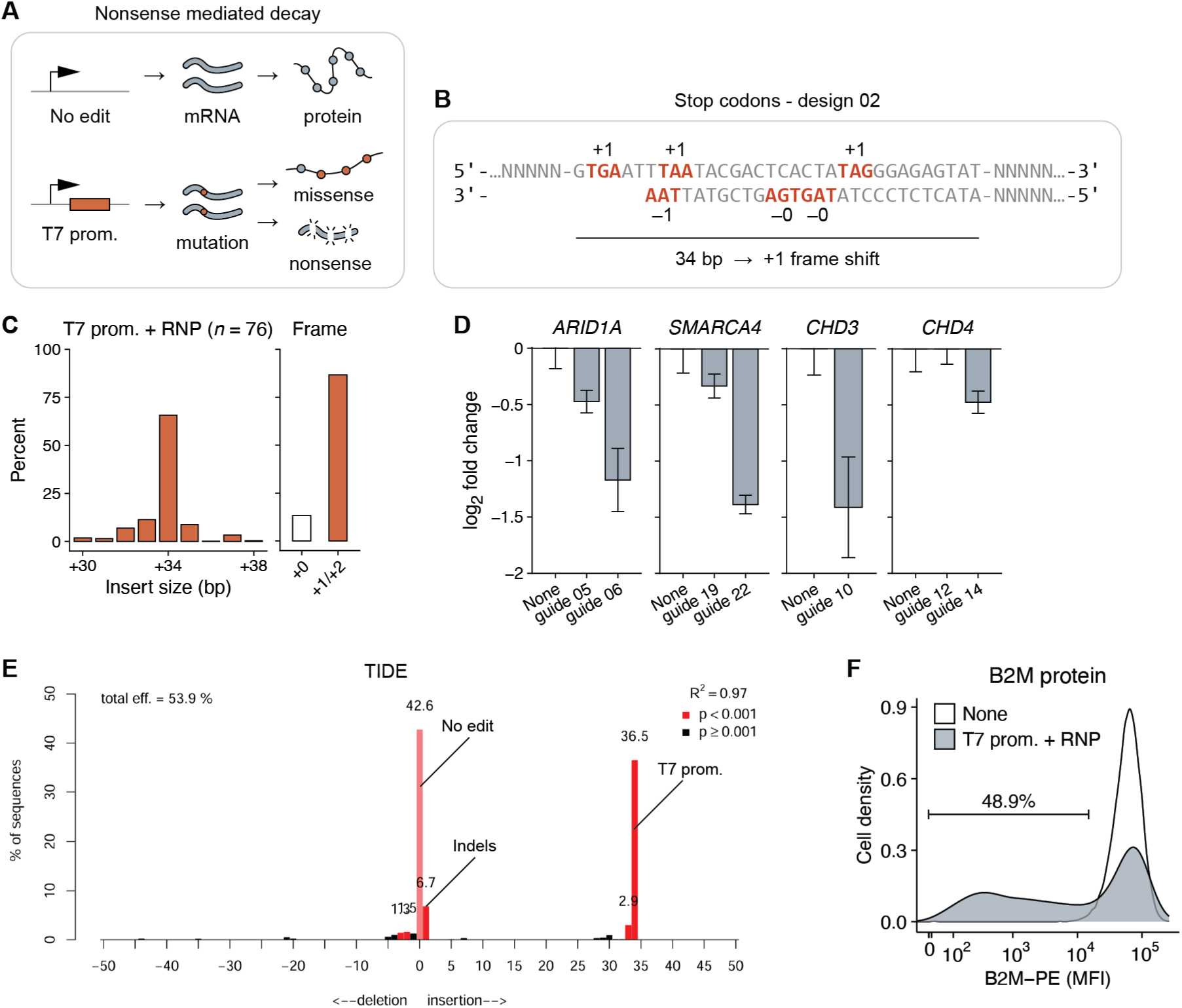
Superb-seq T7 promoter insertion disrupts target gene expression. (**A**) Mechanisms of gene disruption by T7 promoter insertion. (**B**) Stop codon and frameshift mutations inserted by the T7 promoter. Strand and frame of each stop codon is indicated. (**C**) Distribution of T7 promoter insertion size in 76 samples with detectable insertion events, and cumulative frequency of in-frame (+0) or frameshift (+1 and +2) events. (**D**) Relative levels of chromatin remodeler gene expression in K562 cells edited with the indicated guide (and T7 promoter) compared to unedited cells (None), measured by RT-qPCR and the comparative C_t_ method^86^, and represented as log_2_ fold change (log_2_FC) normalized to reference genes *RPL24* and *RPS10*, and the control condition. Bars represent the mean of technical replicates (*n* = 3), and whiskers represent 2 × standard error in the mean (SEM). (**E**) Indel distribution in GM12878 cells with T7 promoter insertion at *B2M*, measured by TIDE. (**F**) Median fluorescence intensity (MFI) of unedited (None) and edited (T7 prom. + RNP) GM12878 cells immunostained with anti-human B2M–PE antibody, measured by flow cytometry. Events are pre-gated on live singlets (**Supp.** Fig. 33).

**Extended Data Figure 2.**
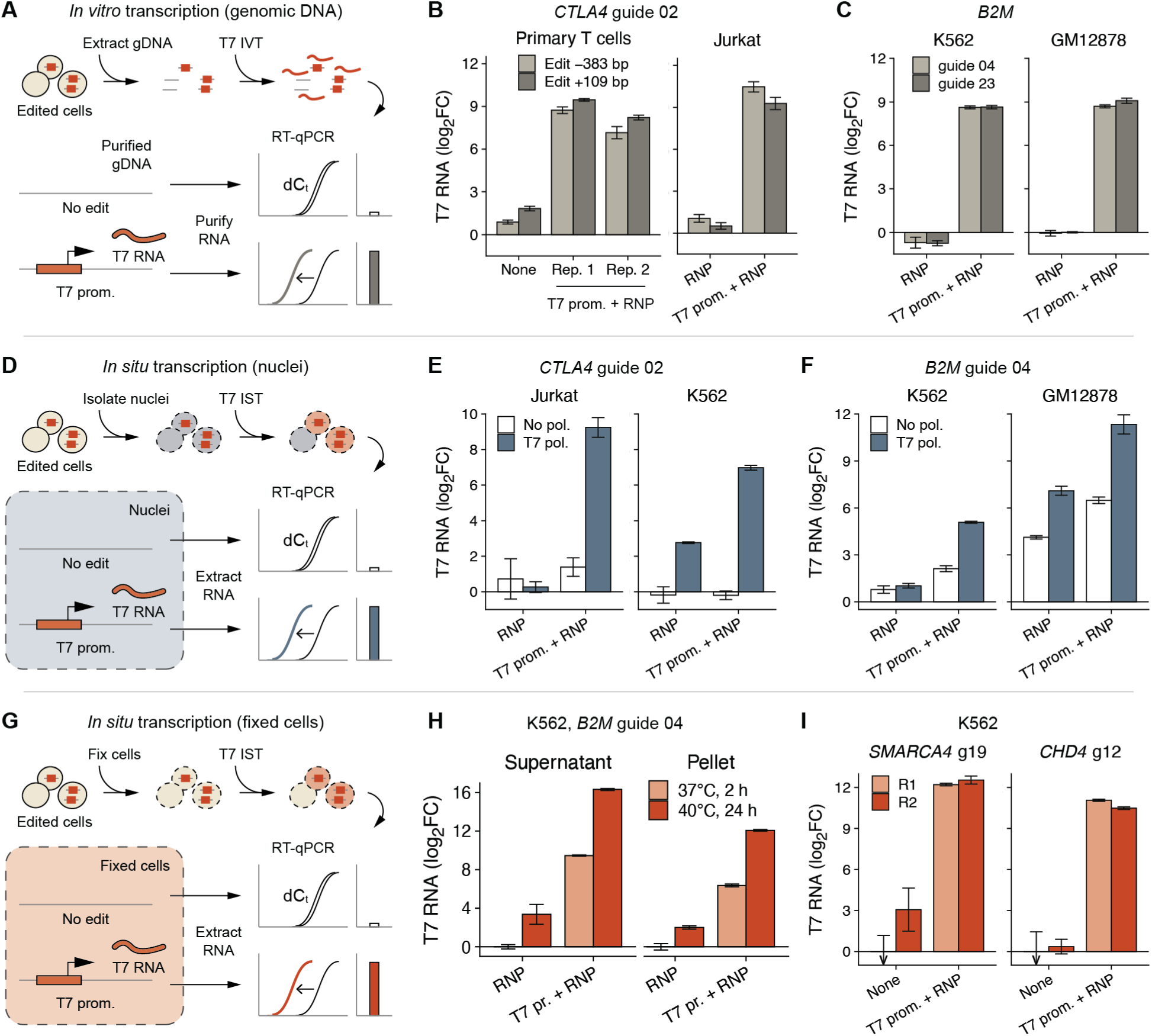
Development of in situ transcription of genome edits. (**A**) IVT of genomic DNA, and RT-qPCR. (**B,C**) Relative levels of IVT transcripts from genomic DNA treated with Cas9 ribonucleoprotein only (RNP), RNP and T7 promoter (T7 prom. + RNP), or none, determined by RT-qPCR. Intron-targeted Cas9 edits were generated at the (**B**) *CTLA4* or (**C**) *B2M* locus in the indicated cell types. Additional labels indicate IVT replicates (R1, R2), PCR assay position relative to the edit site (e.g. Edit +109 bp), and guide RNA (e.g. guide 04). (**D**) IST of unfixed nuclei. (**E,F**) Levels of T7 transcripts from nuclei IST with T7 RNA polymerase (T7 pol) or mock IST without polymerase (No pol). (**G**) IST on fixed cells. (**H**) Levels of T7 transcripts in IST supernatant or cell pellet fractions, before and after optimizing IST incubation. (**I**) Levels of T7 transcripts at *SMARCA4* and *CHD4* edit sites. Transcript levels are normalized to background signal at two non-transcribed genomic safe harbor (GSH) loci^79^ (**B,C,E,F**), or normalized to reference genes *RPL24* and *RPS10*, and the control (no T7 promoter) condition (**H,I**). Bars represent the mean of technical replicates (*n* = 3), and whiskers represent 2 × SEM.

**Extended Data Figure 3.**
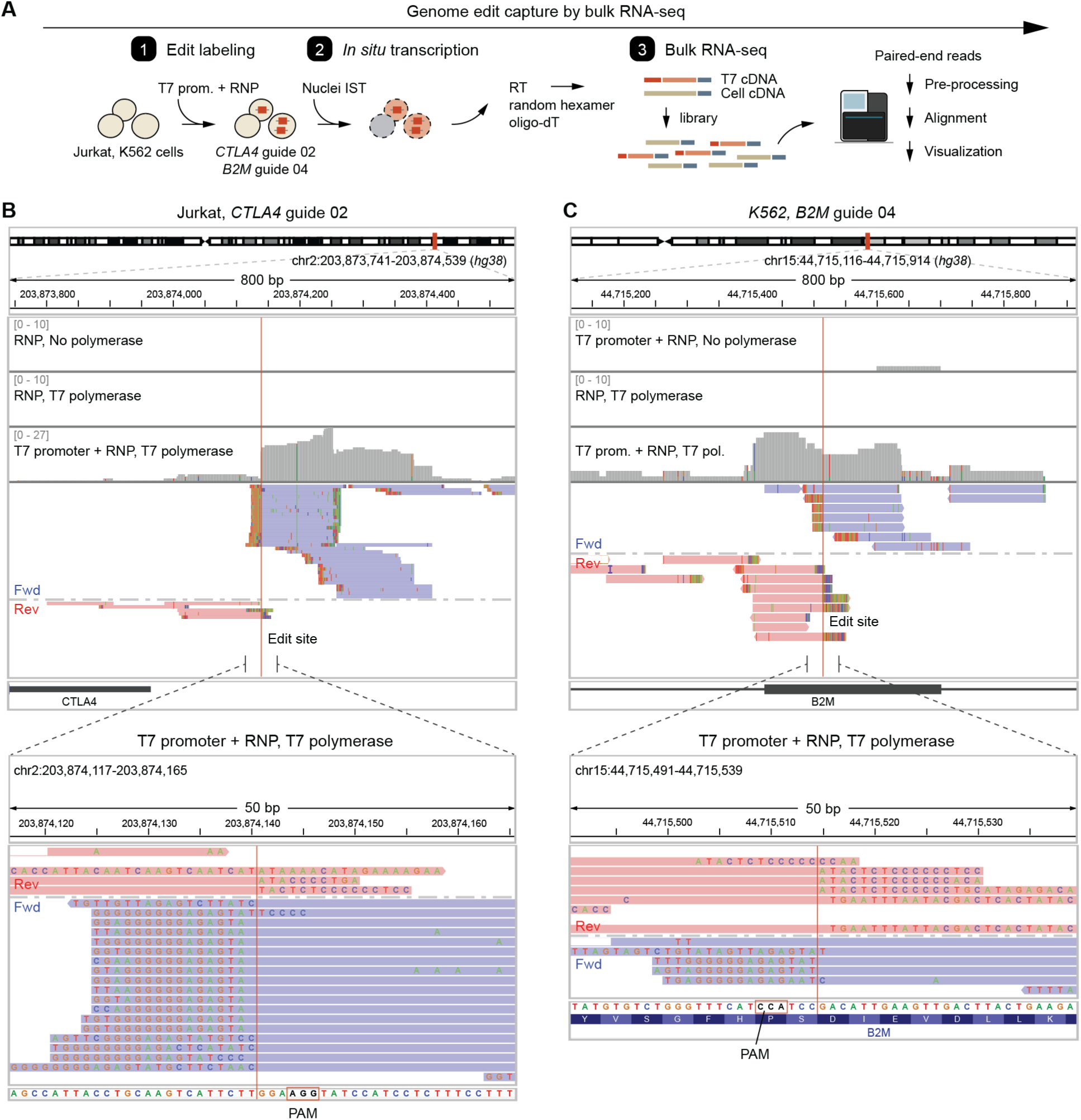
Sequencing of in situ T7 transcripts identifies genome edits. (**A**) Diagram of the bulk RNA-seq experiment. Jurkat and K562 cells were treated with T7 promoter and Cas9 RNP targeting *CTLA4* or *B2M.* Nuclei isolation and T7 IST were then performed, followed by total RNA extraction, RNA-seq library preparation, and paired-end sequencing. (**B,C**) Sequence alignments at *CTLA4* (**B**) and *B2M* (**C**) from samples treated with RNP alone or with T7 promoter, and IST reactions with or without T7 RNA polymerase. Read pileups (gray), individual forward strand (blue) and reverse strand (red) reads, the expected Cas9 cut position (red line), unaligned positions (visible bases), and PAM are shown.

**Extended Data Figure 4.**
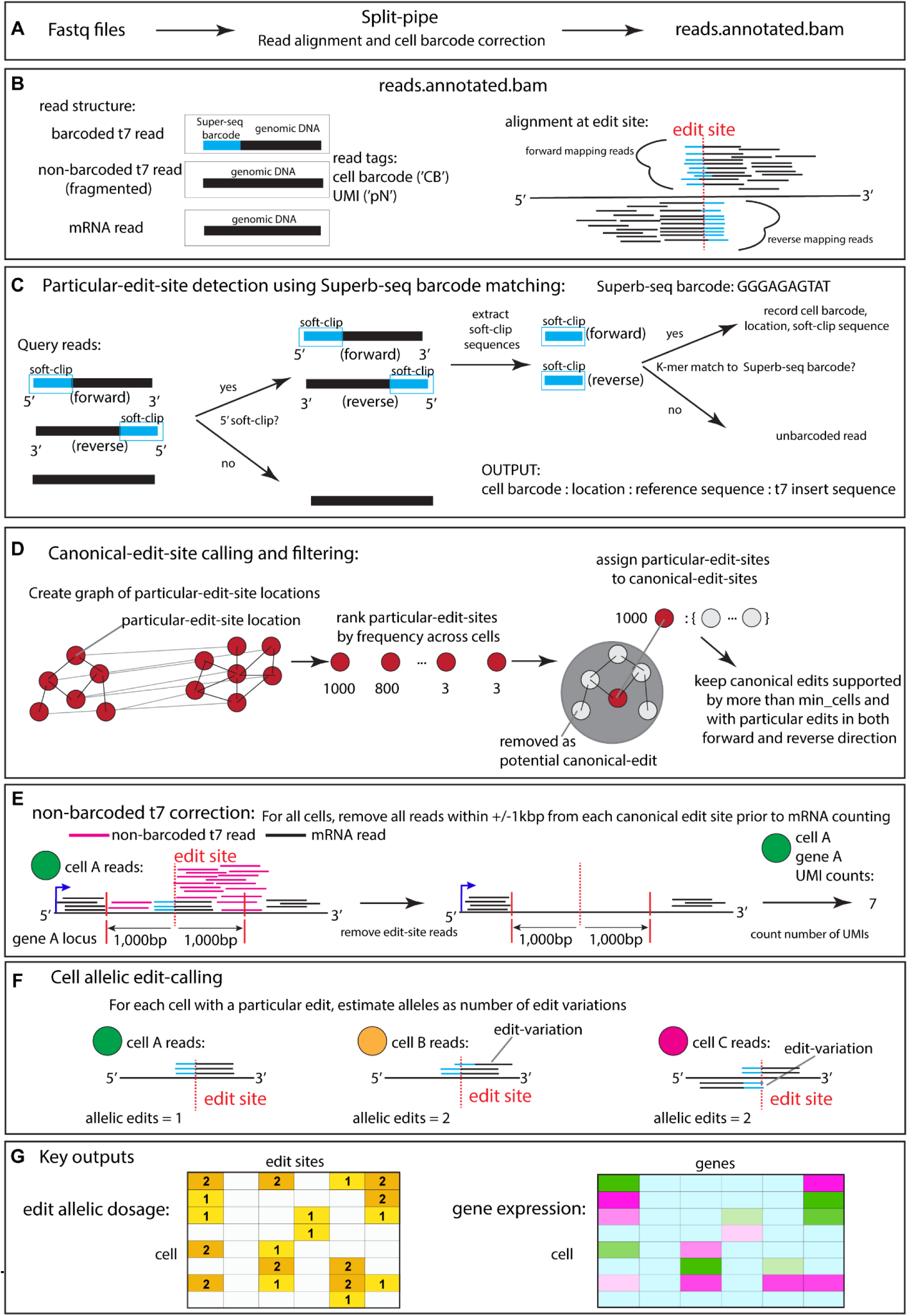
Sheriff, a command line tool to quantify Superb-seq edits and transcriptomes. (**A**) FASTQ files generated from Superb-seq are processed using Split-pipe^40^ to generate an annotated BAM file. (**B**) Three kinds of reads are present within the annotated BAM file; 1) T7 polymerase generated reads with a Superb-seq barcode, evident in the 5’ soft-clipped region of the mapped read, 2) reads generated by T7 polymerase that have been fragmented, and hence without Superb-seq barcode, and 3) reads generated from normal RNA pol II transcription within expressed genic regions. Edit sites have a distinct T7 barcoded read pattern. (**C**) Particular-edit-sites are determined by using cell barcodes on classified T7 barcoded reads, identified by a *k*-mer match on the 5’ soft-clip sequence of reads. (**D**) Particular-edits represent slight edit variation around an edit site. Canonical-edit-sites are determined by assigning particular-edits to more commonly occurring versions of the edit, by first constructing a graph of particular-edit locations for each chromosome, and then rank-ordering particular-edits by the frequency that they occur across cells, then progressively assigning particular-edits to canonical-edit-sites. (**E**) All remaining reads within ± 1000 bp from canonical-edit-sites are removed prior to cell and gene UMI counting, since otherwise non-barcoded T7 reads will confound gene expression estimation. (**F**) For each canonical-edit-site, the number of edited alleles for that site is called per cell. (**G**) The key outputs from the Superb-seq read processing. See **Methods** for details.

**Extended Data Figure 5.**
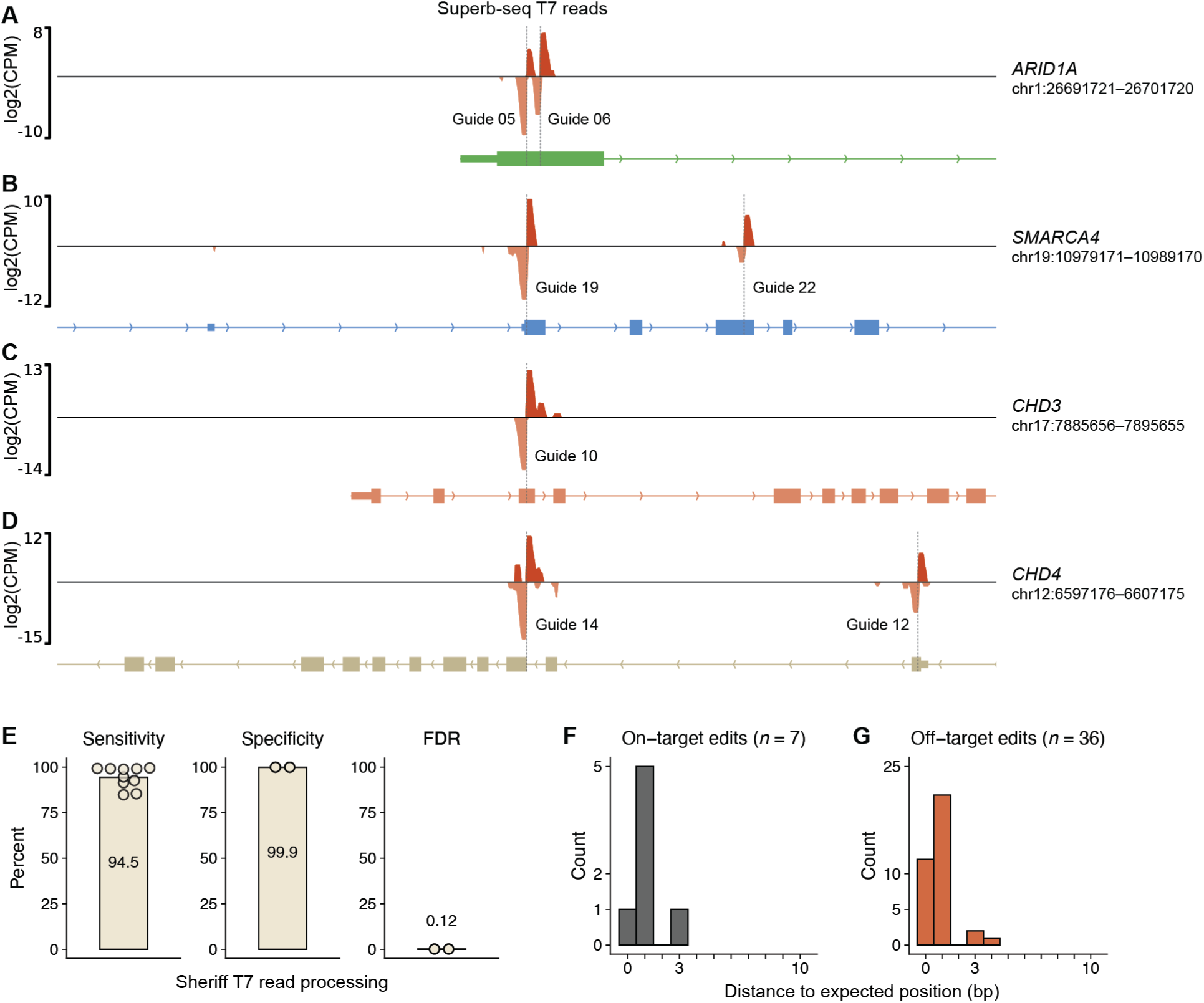
Performance of T7 read processing by Sheriff. (**A–D**) Coverage of T7 reads called by Sheriff at the seven on-target edit sites (count per million mapped reads, CPM). Expected on-target sites are indicated for each guide RNA (dotted lines). (**E**) Sensitivity, specificity, and false discovery rate (FDR) of T7 read calling in 10 technical cell barcoding replicates of BAF- or NuRD-edited sample pools. Sensitivity was defined as the rate of T7 reads detected within ± 100 bp of the seven expected on-target edit sites. Specificity was defined as the rate of non-T7 reads classified correctly in the unedited sample. FDR was defined as the rate of non-T7 reads that were classified as T7 reads. (**F**) Absolute distance between expected on-target edit sites and called edit sites by Sheriff. (**G**) Absolute distance between called edit sites and guide-targeted PAM sequences for the 36 off-target sites, determined by Sheriff.

**Extended Data Figure 6.**
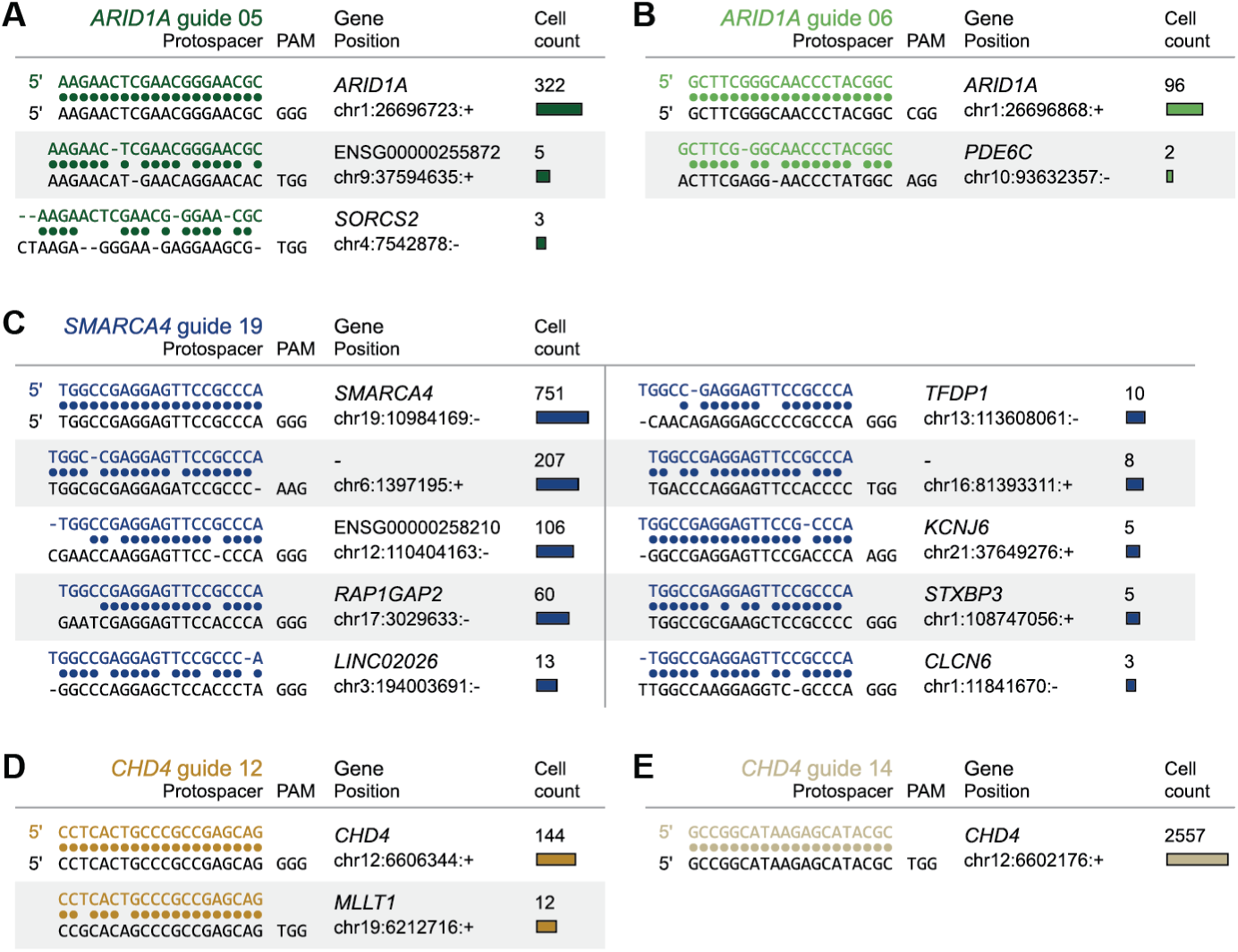
Guide alignments at Superb-seq edit sites. (**A–E**) Guide–reference genome sequence alignments and cell counts of Superb-seq detected edits by (**A**) *ARID1A* guide 05, (**B**) *ARID1A* guide 06, (**C**) *SMARCA4* guide 19, (**D**) *CHD4* guide 12, and (**E**) *CHD4* guide 14. Colored dots represent matched bases. The protospacer adjacent motif (PAM) and the number of edited cells is shown for each edit site.

**Extended Data Figure 7.**
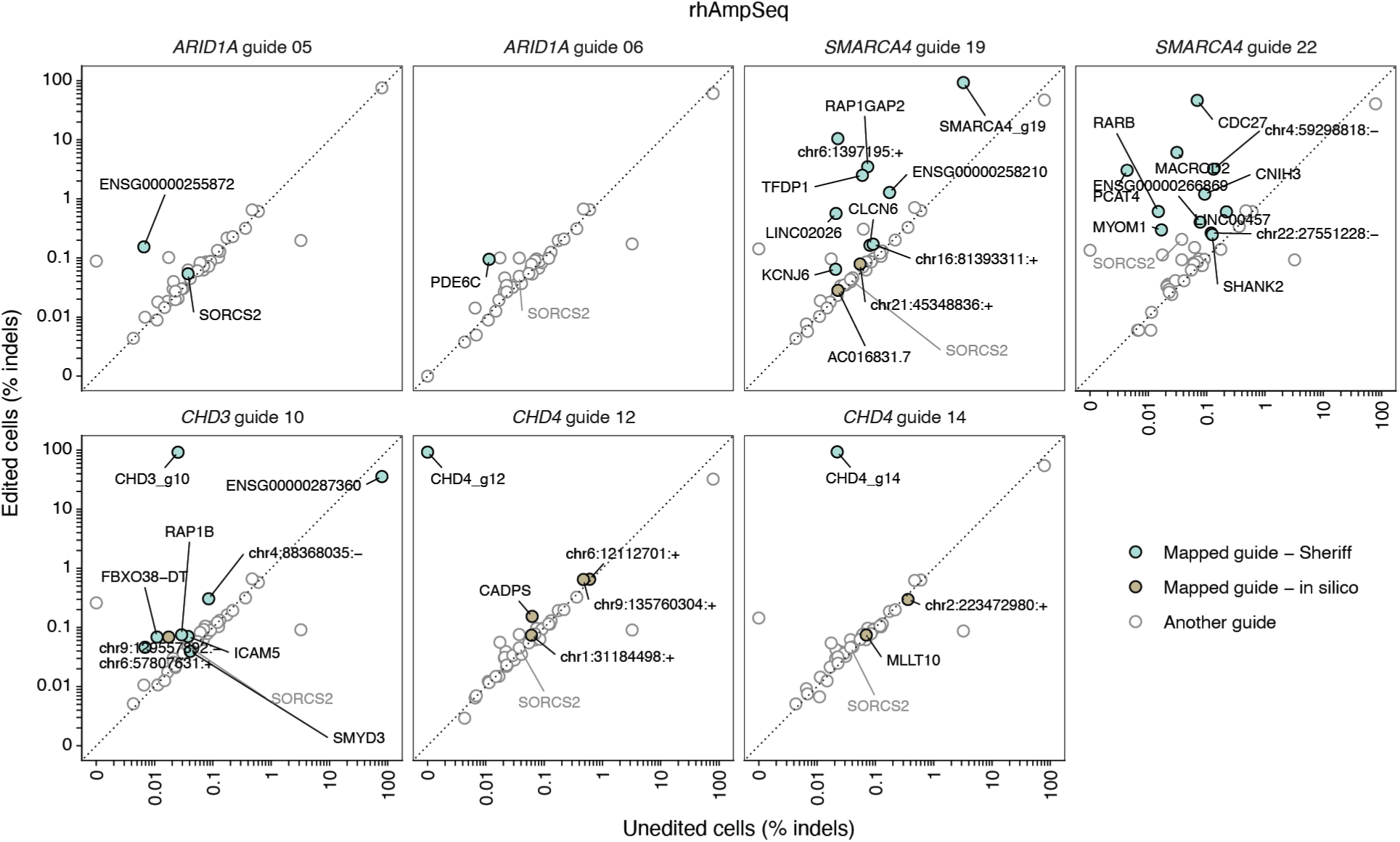
Validation of Sheriff assignment of causal guides by rhAmpSeq. Comparison of indel frequencies between edited and unedited K562 cell samples, determined by CRISPAltRations^104^. Each panel represents a different sample treated with the indicated guide and the panel of 52 multiplexed rhAmpSeq primers. Each point represents a different amplicon with sufficient sequencing coverage (*n* = 43, **Supp.** Fig. 16). Color indicates Sheriff guide association (green), in silico guide association (khaki), or association to another guide (white).

**Extended Data Figure 8.**
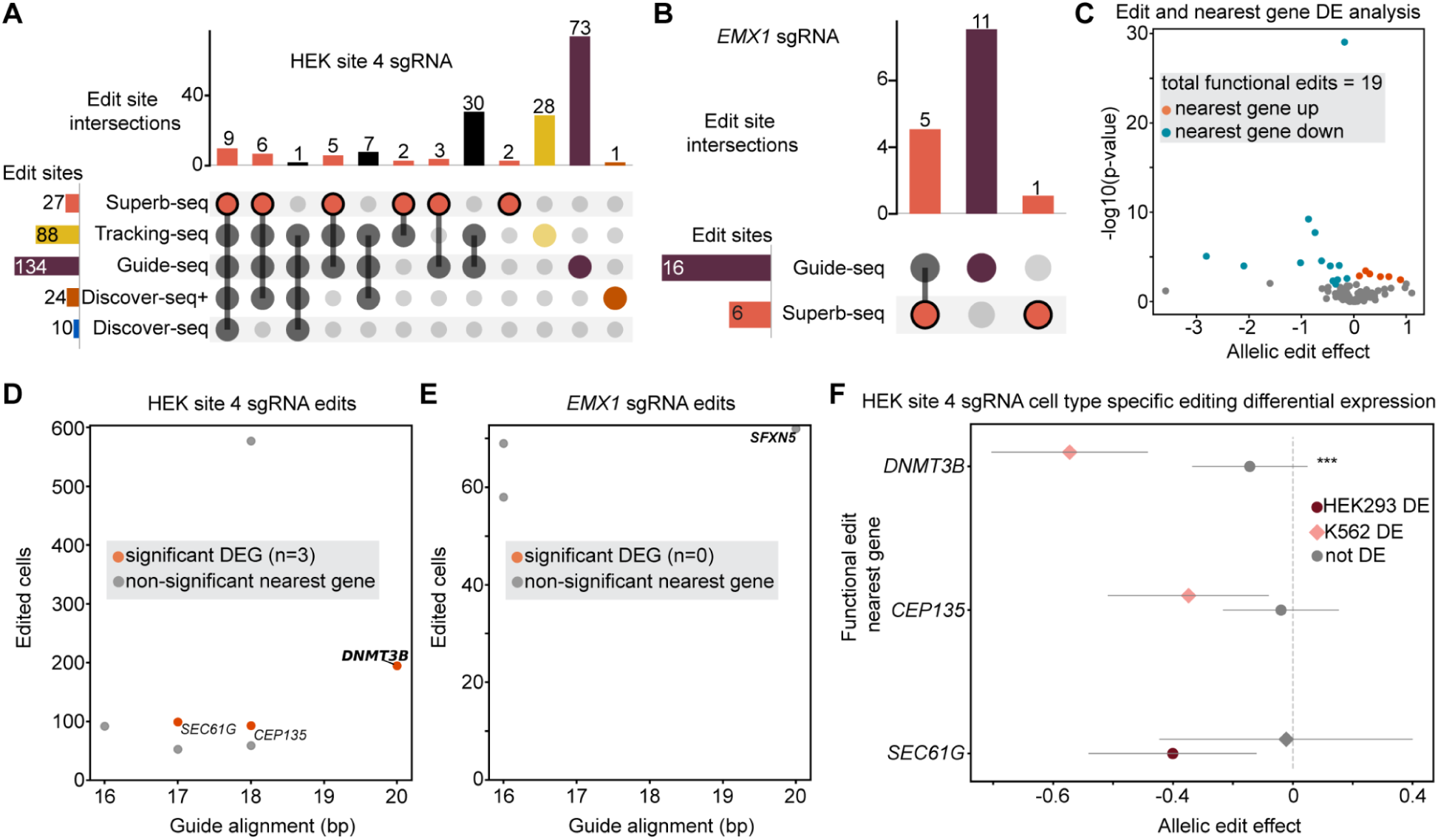
Superb-seq on benchmark guides enables comparison with bulk edit detection methods. (**A**) Upset plot of edits detected across technologies for the HEK site 4 guide. (**B**) Equivalent to A for the *EMX1* guide. (**C**) Volcano plot of frequent edits (*n* ≥ 50 cells) and nearest expressed gene differential expression analysis, with each point representing a test of an edit effecting expression of the nearest gene in a particular cell type. The x-axis is the effect size of one edit allele, and the y-axis is the -log10 transformed p-values of significance. Edits that significantly affect the expression of the nearest gene are highlighted. (**D**) Scatter plot of frequent HEK site 4 guide edits, with guide sequence similarity to the edit site on the x-axis and the number of edited cells on the y-axis. Edits with significant detected effects on the nearest gene expression are highlighted. (**E**) Equivalent to D, except for the *EMX1* guide. (**F**) Forest plot of cell type-specific editing effect sizes (x-axis) across multiple edits (y-axis). Error bars are the 95% confidence intervals of the effect size estimates. Colored points indicate significant differential expression in the respective cell type. Asterisks indicate significantly different editing effects between cell types (*** = *p* < 0.001).

## References

1. Pacesa, M., Pelea, O. & Jinek, M. Past, present, and future of CRISPR genome editing technologies. Cell 187, 1076–1100 (2024).

2. Adamson, B. et al. A Multiplexed Single-Cell CRISPR Screening Platform Enables Systematic Dissection of the Unfolded Protein Response. Cell 167, 1867–1882.e21 (2016).

3. Dixit, A. et al. Perturb-Seq: Dissecting Molecular Circuits with Scalable Single-Cell RNA Profiling of Pooled Genetic Screens. Cell 167, 1853–1866.e17 (2016).

4. Jaitin, D. A. et al. Dissecting Immune Circuits by Linking CRISPR-Pooled Screens with Single-Cell RNA-Seq. Cell 167, 1883–1896.e15 (2016).

5. Datlinger, P. et al. Pooled CRISPR screening with single-cell transcriptome readout. Nat. Methods 14, 297–301 (2017).

6. Replogle, J. M. et al. Mapping information-rich genotype-phenotype landscapes with genome-scale Perturb-seq. Cell 185, 2559–2575.e28 (2022).

7. Peidli, S. et al. scPerturb: harmonized single-cell perturbation data. Nat. Methods 21, 531–540 (2024).

8. Replogle, J. M. et al. Combinatorial single-cell CRISPR screens by direct guide RNA capture and targeted sequencing. Nat. Biotechnol. 38, 954–961 (2020).

9. Zheng, X. et al. Massively parallel in vivo Perturb-seq reveals cell-type-specific transcriptional networks in cortical development. Cell 187, 3236–3248.e21 (2024).

10. Barry, T., Mason, K., Roeder, K. & Katsevich, E. Robust differential expression testing for single-cell CRISPR screens at low multiplicity of infection. Genome Biol. 25, 124 (2024).

11. Doench, J. G. et al. Optimized sgRNA design to maximize activity and minimize off-target effects of CRISPR-Cas9. Nat. Biotechnol. 34, 184–191 (2016).

12. Schmidt, H. et al. Genome-wide CRISPR guide RNA design and specificity analysis with GuideScan2. Genome Biol. 26, 41 (2025).

13. Southard, K. M. et al. Comprehensive transcription factor perturbations recapitulate fibroblast transcriptional states. Nat. Genet. 57, 2323–2334 (2025).

14. Papalexi, E. et al. Characterizing the molecular regulation of inhibitory immune checkpoints with multimodal single-cell screens. Nat. Genet. 53, 322–331 (2021).

15. Ray, J. et al. An unbiased survey of distal element-gene regulatory interactions with direct-capture targeted Perturb-seq. bioRxivorg 2025.09.16.676677 (2025) doi:10.1101/2025.09.16.676677.

16. Morgens, D. W. et al. Genome-scale measurement of off-target activity using Cas9 toxicity in high-throughput screens. Nat. Commun. 8, 15178 (2017).

17. Musunuru, K. et al. Patient-specific in vivo gene editing to treat a rare genetic disease. N. Engl. J. Med. 392, 2235–2243 (2025).

18. Bhokisham, N. et al. CRISPR-Cas system: The current and emerging translational landscape. Cells 12, (2023).

19. Frangoul, H. et al. Exagamglogene autotemcel for severe sickle cell disease. N. Engl. J. Med. 390, 1649–1662 (2024).

20. Center for Biologics Evaluation & Research. Human gene therapy products incorporating human genome editing. *U.S. Food and Drug Administration* https://www.fda.gov/regulatory-information/search-fda-guidance-documents/human-gene-therapy-products-inc orporating-human-genome-editing (2024).

21. Yan, J. et al. Benchmarking and integrating genome-wide CRISPR off-target detection and prediction. Nucleic Acids Res. 48, 11370–11379 (2020).

22. Tsai, S. Q. et al. GUIDE-seq enables genome-wide profiling of off-target cleavage by CRISPR-Cas nucleases. Nat. Biotechnol. 33, 187–197 (2015).

23. Zhu, M. et al. Tracking-seq reveals the heterogeneity of off-target effects in CRISPR-Cas9-mediated genome editing. Nat. Biotechnol. 1–12 (2024).

24. Jacobi, A. M. et al. Simplified CRISPR tools for efficient genome editing and streamlined protocols for their delivery into mammalian cells and mouse zygotes. Methods 121-122, 16–28 (2017).

25. Cancellieri, S. et al. Human genetic diversity alters off-target outcomes of therapeutic gene editing. Nat. Genet. 55, 34–43 (2023).

26. Neavin, D. et al. Demuxafy: improvement in droplet assignment by integrating multiple single-cell demultiplexing and doublet detection methods. Genome Biol. 25, 94 (2024).

27. Askary, A. et al. In situ readout of DNA barcodes and single base edits facilitated by in vitro transcription. Nat. Biotechnol. 38, 66–75 (2020).

28. Kim, S., Kim, D., Cho, S. W., Kim, J. & Kim, J.-S. Highly efficient RNA-guided genome editing in human cells via delivery of purified Cas9 ribonucleoproteins. Genome Res. 24, 1012–1019 (2014).

29. Zuris, J. A. et al. Cationic lipid-mediated delivery of proteins enables efficient protein-based genome editing in vitro and in vivo. Nat. Biotechnol. 33, 73–80 (2015).

30. Suzuki, K. et al. In vivo genome editing via CRISPR/Cas9 mediated homology-independent targeted integration. Nature 540, 144–149 (2016).

31. Geisinger, J. M., Turan, S., Hernandez, S., Spector, L. P. & Calos, M. P. In vivo blunt-end cloning through CRISPR/Cas9-facilitated non-homologous end-joining. Nucleic Acids Res. 44, e76 (2016).

32. Chen, X. et al. Recent advances in CRISPR-Cas9-based genome insertion technologies. Mol. Ther. Nucleic Acids 35, 102138 (2024).

33. Lieber, A., Sandig, V. & Strauss, M. A mutant T7 phage promoter is specifically transcribed by T7-RNA polymerase in mammalian cells. Eur. J. Biochem. 217, 387–394 (1993).

34. Paul, S., Stang, A., Lennartz, K., Tenbusch, M. & Überla, K. Selection of a T7 promoter mutant with enhanced in vitro activity by a novel multi-copy bead display approach for in vitro evolution. Nucleic Acids Res. 41, e29 (2013).

35. Komura, R., Aoki, W., Motone, K., Satomura, A. & Ueda, M. High-throughput evaluation of T7 promoter variants using biased randomization and DNA barcoding. PLoS One 13, e0196905 (2018).

36. Conrad, T., Plumbom, I., Alcobendas, M., Vidal, R. & Sauer, S. Maximizing transcription of nucleic acids with efficient T7 promoters. Commun Biol 3, 439 (2020).

37. Brinkman, E. K., Chen, T., Amendola, M. & van Steensel, B. Easy quantitative assessment of genome editing by sequence trace decomposition. Nucleic Acids Res. 42, e168 (2014).

38. Rosenberg, A. B. et al. Single-cell profiling of the developing mouse brain and spinal cord with split-pool barcoding. Science 360, 176–182 (2018).

39. Luo, Y. et al. New developments on the Encyclopedia of DNA Elements (ENCODE) data portal. Nucleic Acids Res. 48, D882–D889 (2020).

40. Tran, V. et al. High sensitivity single cell RNA sequencing with split pool barcoding. bioRxiv 2022.08.27.505512 (2022) doi:10.1101/2022.08.27.505512.

41. Jiang, F., Zhou, K., Ma, L., Gressel, S. & Doudna, J. A. A Cas9-guide RNA complex preorganized for target DNA recognition. Science 348, 1477–1481 (2015).

42. Bae, S., Park, J. & Kim, J.-S. Cas-OFFinder: a fast and versatile algorithm that searches for potential off-target sites of Cas9 RNA-guided endonucleases. Bioinformatics 30, 1473–1475 (2014).

43. Cradick, T. J., Qiu, P., Lee, C. M., Fine, E. J. & Bao, G. COSMID: A web-based tool for identifying and validating CRISPR/Cas off-target sites. Mol. Ther. Nucleic Acids 3, e214 (2014).

44. Heigwer, F., Kerr, G. & Boutros, M. E-CRISP: fast CRISPR target site identification. Nat. Methods 11, 122–123 (2014).

45. Macrae, T. A. & Ramalho-Santos, M. The deubiquitinase Usp9x regulates PRC2-mediated chromatin reprogramming during mouse development. Nat. Commun. 12, 1865 (2021).

46. Song, J. et al. Regulation of alternative polyadenylation by the C2H2-zinc-finger protein Sp1. Mol. Cell 82, 3135–3150.e9 (2022).

47. van der Vaart, A., Godfrey, M., Portegijs, V. & van den Heuvel, S. Dose-dependent functions of SWI/SNF BAF in permitting and inhibiting cell proliferation in vivo. Sci. Adv. 6, eaay3823 (2020).

48. Braun, S. M. G. et al. BAF subunit switching regulates chromatin accessibility to control cell cycle exit in the developing mammalian cortex. Genes Dev. 35, 335–353 (2021).

49. Dhanasekaran, R. et al. The MYC oncogene - the grand orchestrator of cancer growth and immune evasion. Nat. Rev. Clin. Oncol. 19, 23–36 (2022).

50. Xie, Y. et al. Comparative analysis of single-cell RNA sequencing methods with and without sample multiplexing. Int. J. Mol. Sci. 25, 3828 (2024).

51. Wienert, B. et al. Unbiased detection of CRISPR off-targets in vivo using DISCOVER-Seq. Science 364, 286–289 (2019).

52. Zou, R. S. et al. Improving the sensitivity of in vivo CRISPR off-target detection with DISCOVER-Seq. Nat. Methods 20, 706–713 (2023).

53. Duncan, C. N. et al. Hematologic cancer after gene therapy for cerebral adrenoleukodystrophy. N. Engl. J. Med. 391, 1287–1301 (2024).

54. Kalter, N. et al. Precise measurement of CRISPR genome editing outcomes through single-cell DNA sequencing. Mol. Ther. Methods Clin. Dev. 33, 101449 (2025).

55. Kudo, T. et al. Multiplexed, image-based pooled screens in primary cells and tissues with PerturbView. Nat. Biotechnol. (2024) doi:10.1038/s41587-024-02391-0.

56. Fandrey, C. I. et al. NIS-Seq enables cell-type-agnostic optical perturbation screening. Nat. Biotechnol. (2024) doi:10.1038/s41587-024-02516-5.

57. Binan, L. et al. Simultaneous CRISPR screening and spatial transcriptomics reveal intracellular, intercellular, and functional transcriptional circuits. Cell 188, 2141–2158.e18 (2025).

58. Li, X. et al. Chromatin context-dependent regulation and epigenetic manipulation of prime editing. Cell 0, (2024).

59. Pinglay, S. et al. Multiplex generation and single cell analysis of structural variants in a mammalian genome. bioRxiv 2024.01.22.576756 (2024) doi:10.1101/2024.01.22.576756.

60. van Overbeek, M. et al. DNA repair profiling reveals nonrandom outcomes at Cas9-mediated breaks. Mol. Cell 63, 633–646 (2016).

61. Dunn, J. J. & Studier, F. W. Complete nucleotide sequence of bacteriophage T7 DNA and the locations of T7 genetic elements. J. Mol. Biol. 166, 477–535 (1983).

62. Tang, G.-Q., Bandwar, R. P. & Patel, S. S. Extended upstream A-T sequence increases T7 promoter strength. J. Biol. Chem. 280, 40707–40713 (2005).

63. Malinin, N. L. et al. Defining genome-wide CRISPR–Cas genome-editing nuclease activity with GUIDE-seq. Nat. Protoc. 16, 5592–5615 (2021).

64. Allen, F. et al. Predicting the mutations generated by repair of Cas9-induced double-strand breaks. Nat. Biotechnol. 37, 64–72 (2018).

65. Lemos, B. R. et al. CRISPR/Cas9 cleavages in budding yeast reveal templated insertions and strand-specific insertion/deletion profiles. Proc. Natl. Acad. Sci. U. S. A. 115, E2040–E2047 (2018).

66. Shou, J., Li, J., Liu, Y. & Wu, Q. Precise and predictable CRISPR chromosomal rearrangements reveal principles of Cas9-mediated nucleotide insertion. Mol. Cell 71, 498–509.e4 (2018).

67. Chakrabarti, A. M. et al. Target-specific precision of CRISPR-mediated genome editing. Mol. Cell 73, 699–713.e6 (2019).

68. Chen, W. et al. Massively parallel profiling and predictive modeling of the outcomes of CRISPR/Cas9-mediated double-strand break repair. Nucleic Acids Res. 47, 7989–8003 (2019).

69. Gisler, S. et al. Multiplexed Cas9 targeting reveals genomic location effects and gRNA-based staggered breaks influencing mutation efficiency. Nat. Commun. 10, 1598 (2019).

70. Sanson, K. R. et al. Optimized libraries for CRISPR-Cas9 genetic screens with multiple modalities. Nat. Commun. 9, 5416 (2018).

71. Hendel, A. et al. Chemically modified guide RNAs enhance CRISPR-Cas genome editing in human primary cells. Nat. Biotechnol. 33, 985–989 (2015).

72. Perez, A. R. et al. GuideScan software for improved single and paired CRISPR guide RNA design. Nat. Biotechnol. 35, 347–349 (2017).

73. Stadtmauer, E. A. et al. CRISPR-engineered T cells in patients with refractory cancer. Science 367, eaba7365 (2020).

74. Ye, J. et al. Primer-BLAST: a tool to design target-specific primers for polymerase chain reaction. BMC Bioinformatics 13, 134 (2012).

75. Hahne, F. et al. flowCore: a Bioconductor package for high throughput flow cytometry. BMC Bioinformatics 10, 106 (2009).

76. Van, P., Jiang, W., Gottardo, R. & Finak, G. ggCyto: next generation open-source visualization software for cytometry. Bioinformatics 34, 3951–3953 (2018).

77. Corces, M. R. et al. An improved ATAC-seq protocol reduces background and enables interrogation of frozen tissues. Nat. Methods 14, 959–962 (2017).

78. Untergasser, A., Ruijter, J. M., Benes, V. & van den Hoff, M. J. B. Web-based LinRegPCR: application for the visualization and analysis of (RT)-qPCR amplification and melting data. BMC Bioinformatics 22, 398 (2021).

79. Aznauryan, E. et al. Discovery and validation of human genomic safe harbor sites for gene and cell therapies. Cell Rep Methods 2, 100154 (2022).

80. Ye, J., McGinnis, S. & Madden, T. L. BLAST: improvements for better sequence analysis. Nucleic Acids Res. 34, W6–9 (2006).

81. Laurent, S. et al. The engagement of CTLA-4 on primary melanoma cell lines induces antibody-dependent cellular cytotoxicity and TNF-α production. J. Transl. Med. 11, 108 (2013).

82. Wilson, M. R. et al. ARID1A and PI3-kinase pathway mutations in the endometrium drive epithelial transdifferentiation and collective invasion. Nat. Commun. 10, 3554 (2019).

83. Geigges, M. et al. Reference genes for expression studies in human CD8+ naïve and effector memory T cells under resting and activating conditions. Sci. Rep. 10, 9411 (2020).

84. Xu, N. et al. CHD4 mediates proliferation and migration of non-small cell lung cancer via the RhoA/ROCK pathway by regulating PHF5A. BMC Cancer 20, 262 (2020).

85. Yao, B. et al. PRMT1-mediated H4R3me2a recruits SMARCA4 to promote colorectal cancer progression by enhancing EGFR signaling. Genome Med. 13, 58 (2021).

86. Schmittgen, T. D. & Livak, K. J. Analyzing real-time PCR data by the comparative C(T) method. Nat. Protoc. 3, 1101–1108 (2008).

87. Taylor, S. C. et al. The Ultimate qPCR Experiment: Producing Publication Quality, Reproducible Data the First Time. Trends Biotechnol. 37, 761–774 (2019).

88. Martin, M. Cutadapt removes adapter sequences from high-throughput sequencing reads. EMBnet.journal 17, 10–12 (2011).

89. Ewels, P., Magnusson, M., Lundin, S. & Käller, M. MultiQC: summarize analysis results for multiple tools and samples in a single report. Bioinformatics 32, 3047–3048 (2016).

90. Li, H. Aligning sequence reads, clone sequences and assembly contigs with BWA-MEM. arXiv [q-bio.GN] (2013) doi:10.48550/arXiv.1303.3997.

91. Li, H. et al. The Sequence Alignment/Map format and SAMtools. Bioinformatics 25, 2078–2079 (2009).

92. Robinson, J. T. et al. Integrative genomics viewer. Nat. Biotechnol. 29, 24–26 (2011).

93. Amemiya, H. M., Kundaje, A. & Boyle, A. P. The ENCODE blacklist: Identification of problematic regions of the genome. Sci. Rep. 9, 9354 (2019).

94. Zhang, Y. et al. Model-based analysis of ChIP-Seq (MACS). Genome Biol. 9, R137 (2008).

95. Cock, P. J. A. et al. Biopython: freely available Python tools for computational molecular biology and bioinformatics. Bioinformatics 25, 1422–1423 (2009).

96. Stoler, N. & Nekrutenko, A. Sequencing error profiles of Illumina sequencing instruments. NAR Genom. Bioinform. 3, lqab019 (2021).

97. Zhou, B. et al. Comprehensive, integrated, and phased whole-genome analysis of the primary ENCODE cell line K562. Genome Res. 29, 472–484 (2019).

98. Smith, T., Heger, A. & Sudbery, I. UMI-tools: modeling sequencing errors in Unique Molecular Identifiers to improve quantification accuracy. Genome Res. 27, 491–499 (2017).

99. Wolf, F. A., Angerer, P. & Theis, F. J. SCANPY: large-scale single-cell gene expression data analysis. Genome Biol. 19, 15 (2018).

100. Balderson, B., Piper, M., Thor, S. & Bodén, M. Cytocipher determines significantly different populations of cells in single-cell RNA-seq data. Bioinformatics 39, (2023).

101. Traag, V. A., Waltman, L. & van Eck, N. J. From Louvain to Leiden: guaranteeing well-connected communities. Sci. Rep. 9, 5233 (2019).

102. Hsu, P. D. et al. DNA targeting specificity of RNA-guided Cas9 nucleases. Nat. Biotechnol. 31, 827–832 (2013).

103. Fleming, S. J. et al. Unsupervised removal of systematic background noise from droplet-based single-cell experiments using CellBender. Nat. Methods 20, 1323–1335 (2023).

104. Kurgan, G. et al. CRISPAltRations: a validated cloud-based approach for interrogation of double-strand break repair mediated by CRISPR genome editing. Mol. Ther. Methods Clin. Dev. 21, 478–491 (2021).

105. Needleman, S. B. & Wunsch, C. D. A general method applicable to the search for similarities in the amino acid sequence of two proteins. J. Mol. Biol. 48, 443–453 (1970).

106. Hafemeister, C. & Satija, R. Normalization and variance stabilization of single-cell RNA-seq data using regularized negative binomial regression. Genome Biol. 20, 296 (2019).

107. Hodges, C., Kirkland, J. G. & Crabtree, G. R. The many roles of BAF (mSWI/SNF) and PBAF complexes in cancer. Cold Spring Harb. Perspect. Med. 6, a026930 (2016).

108. McDonald, B. et al. Canonical BAF complex activity shapes the enhancer landscape that licenses CD8+ T cell effector and memory fates. Immunity 56, 1303–1319.e5 (2023).

109. Gao, T. & Qian, J. EnhancerAtlas 2.0: an updated resource with enhancer annotation in 586 tissue/cell types across nine species. Nucleic Acids Res. 48, D58–D64 (2020).

110. Hinrichs, A. S. et al. The UCSC Genome Browser Database: update 2006. Nucleic Acids Res. 34, D590–8 (2006).

111. Barretina, J. et al. The Cancer Cell Line Encyclopedia enables predictive modelling of anticancer drug sensitivity. Nature 483, 603–607 (2012).

112. Stovner, E. B. & Sætrom, P. PyRanges: efficient comparison of genomic intervals in Python. Bioinformatics 36, 918–919 (2020).

113. Barrett, T. et al. NCBI GEO: archive for functional genomics data sets--update. Nucleic Acids Res. 41, D991–5 (2013).

