## Supplementary Information for "Joint single-cell capture of Cas9 edits and transcriptomes reveals on- and off-target effects on gene expression"

Supplementary Note 1. Superb-seq T7 promoter insertion disrupts target gene expression.
Supplementary Note 2. T7 promoter labeling results in distinct rates of background indels.
Supplementary Note 3. Development of in vitro and in situ transcription of genome edit sites.
Supplementary Note 4. Bulk RNA-seq of in situ T7 transcripts.
Supplementary Note 5. Superb-seq on chromatin remodeler complexes.
Supplementary Note 6. Correction of ambient T7 reads.
Supplementary Note 7. Assessment of edit allele quantification.

Supplementary Figure 1. Development of phage T7 promoter labeling of genome edits.
Supplementary Figure 2. Bimodal background indel rate associates with guide RNA.
Supplementary Figure 3. Generation of in vitro T7 transcripts at genome edit sites.
Supplementary Figure 4. Generation of in situ T7 transcripts in nuclei and fixed cells.
Supplementary Figure 5. Optimization of in situ transcription for SPLiT-seq fixed cells.
Supplementary Figure 6. In situ transcription at edit sites in chromatin remodeler genes.
Supplementary Figure 7. Generation of Superb-seq libraries.
Supplementary Figure 8. Structure of Superb-seq libraries.
Supplementary Figure 9. Correction of ambient T7 reads in the 10,000 cell Superb-seq library.
Supplementary Figure 10. Correction of ambient T7 reads in the 500 cell Superb-seq library.
Supplementary Figure 11. UMI and gene count distributions across Superb-seq sample pools.
Supplementary Figure 12. Transcriptional heterogeneity of Superb-seq transcriptomes.
Supplementary Figure 13. T7 read features indicate multiple edit alleles within individual cells.
Supplementary Figure 14. Reproducible edit capture between 500 cell and 10,000 Superb-seq libraries.
Supplementary Figure 15. Superb-seq captures genome-wide off-target events.
Supplementary Figure 16. Sequencing coverage of rhAmpSeq amplicons.
Supplementary Figure 17. Validation of off-target T7 promoter insertion by rhAmpSeq.
Supplementary Figure 18. Read mismapping dampens rhAmpSeq detection of T7 promoter insertion.
Supplementary Figure 19. Validation of Sheriff guide assignments by rhAmpSeq.
Supplementary Figure 20. Validation of Sheriff guide assignments by rhAmpSeq indel ratios.
Supplementary Figure 21. Differential gene expression associates with edit allele dosage.
Supplementary Figure 22. Off-target *USP9X* edit site intersects an SP1 binding motif.
Supplementary Figure 23. Quality control metrics in the Superb-seq benchmarking and validation library.
Supplementary Figure 24. *VEGFA* site 2 guide alignments at Superb-seq edit sites.
Supplementary Figure 25. *VEGFA* site 2 guide alignments at Superb-seq edit sites, continued.
Supplementary Figure 26. Benchmark guide alignments at Superb-seq edit sites.
Supplementary Figure 27. Superb-seq edited cell counts are correlated with bulk methods.
Supplementary Figure 28. Nearest gene differential expression results by cell type.
Supplementary Figure 29. Nearest gene differential expression results by cell type, continued 1.
Supplementary Figure 30. Nearest gene differential expression results by cell type, continued 2.
Supplementary Figure 31. Cell type-specific expression of nearest genes.
Supplementary Figure 32. Cell type-specific expression of nearest genes, continued.
Supplementary Figure 33. Flow cytometry gating.
Supplementary Figure 34. Multiple metrics confirm confident Superb-seq edit detection.
Supplementary Figure 35. Correction of gene expression level estimation by allelic pairing.

### Additional files

Supplementary Table 1 (.xlsx)

| | |
|---|---|
| guide_rnas | Guide RNA sequences. |
| t7_promoters | T7 promoter and additional donor DNA sequences. |
| tide_pcr_primers | PCR primers for Tracking indels by decomposition (TIDE). |
| t7_pcr_primers | PCR primers for RT-qPCR of T7 transcripts. |
| gene_pcr_primers | PCR primers for RT-qPCR of gene expression. |
| cell_samples | Cell sample information and editing treatment conditions. |

Supplementary Table 2 (.xlsx)

| | |
|---|---|
| tide_settings | TIDE analysis parameters. |
| tide_summary | TIDE sample information and summary statistics. |
| tide_long | TIDE raw data and calculations. |
| tide_wide | TIDE summary data. |
| qubit_rnahs | Qubit RNA HS fluorometry on T7 in vitro transcription. |
| blast_hit_table | BLAST hits of T7 promoter sequence alignment to hg38 reference genome. |
| blast_alignment | BLAST sequence alignments. |
| rtqpcr_t7 | RT-qPCR data on T7 transcription. |
| rtqpcr_genes | RT-qPCR data on gene expression. |
| linregpcr | LinRegPCR data on PCR efficiencies of RT-qPCR assays. |
| flow_cytometry | Flow cytometry data on B2M protein expression. |
| splitpipe_remod500 | Split-pipe quality check metrics, 500 cell chromatin remodeler library. |
| splitpipe_remod10k | Split-pipe quality check metrics, 10,000 cell chromatin remodeler library. |
| splitpipe_bench10krep1 | Split-pipe quality check metrics, first 12,000 cell benchmarking library. |
| splitpipe_bench10krep2 | Split-pipe quality check metrics, second 12,000 cell benchmarking library. |
| insilico_offtargets | Off-target sites predicted by Cas-OFFinder, COSMID, and E-CRISP ($n$ = 63). |
| rhampseq | RhAmpSeq results from CRISPAltRations analysis. |

Supplementary Table 3. (.xlsx) Sheriff called edit sites for the chromatin remodeler Superb-seq libraries.
Supplementary Table 4. (.xlsx) DE statistics for chromatin remodeler Superb-seq experiment.
Supplementary Table 5. (.xlsx) NuRD, BAF, and G2/M functional gene sets.
Supplementary Table 6. (.xlsx) Contingency tables intersecting DEGs with functional gene sets.
Supplementary Table 7. (.xlsx) Functional gene set and DEG enrichment analysis summary statistics.
Supplementary Table 8. (.xlsx) Edit sites for *USP9X* guides
Supplementary Table 9. (.xlsx) Summary statistics for *USP9X* guide DE and *MYC* enrichment analyses.
Supplementary Table 10. (.xlsx) Edit sites for guides in benchmark Superb-seq experiment
Supplementary Table 11. (.xlsx) Summary statistics of benchmark results between sequencing methods.
Supplementary Table 12. (.xlsx) DE statistics for the benchmarking Superb-seq experiment.
Supplementary Table 13. (.xlsx) Blacklisted sequences and genome regions for edit site calling.

Supplementary Material 1. Barcoded T7 read alignments, 10,000 cell chromatin remodeler library.

**Supplementary Note 1. Superb-seq T7 promoter insertion disrupts target gene expression.**

To maximize gene disruption upon insertion of T7 promoter into coding sequences (**Ext. Data. 1A**), we set the length of all donor DNA designs to *3n+1* to yield +1 shifts upon precise insertion and +2 shifts upon the common one base pair (bp) templated insertion by Cas9[1] (**Supp. Table 1**). Additionally, the Superb-seq T7 promoter (design 02) encodes stop codons in one of three forward reading frames and two of three reverse reading frames (**Ext. Data Fig. 1B**). Consistent with its design, > 90% of observed T7 promoter insertions generated +1 or +2 frame-shifts (**Ext. Data Fig. 1C**). To confirm that insertion events triggered nonsense-mediated decay (NMD) of target genes, we measured gene and protein expression by reverse-transcription qPCR (RT-qPCR) and flow cytometry. Promoter-edited cells exhibited target transcript knockdown consistent with NMD (**Ext. Data Fig. 1D**). Furthermore, in a sample of cells with 40% T7 promoter alleles and < 10% background indel alleles (**Ext. Data Fig. 1E**), we observed knockdown of B2M target protein in 50% of cells (**Ext. Data Fig. 1F**). In summary, homology-free insertion of the Superb-seq T7 promoter achieves efficient labeling of Cas9 edits and knockdown of targeted genes.

**Supplementary Note 2. T7 promoter labeling results in distinct rates of background indels.**

We observed that delivery of RNP and T7 promoter resulted in two discrete outcomes: a "high indel" outcome with a high rate of small indels but a low rate of T7 promoter insertions, and a "low indel" outcome with a low rate of small indels and the potential for a very high rate of T7 promoter insertions (**Fig. 1D, Supp. Fig. 1E, Supp. Fig. 2A**). On average, the low indel group had a 4 fold lower rate of background indels (18.0% vs. 72.8%) and a 2.5 fold higher rate of T7 promoter insertions (34.9% vs. 6.8%) (**Supp. Fig. 1E**). To explain this bifurcated editing fate, we looked for associations of edit outcome with guide RNA, cell type, and genome site. In the subset of 63 samples treated with guide RNAs that were used in two or more samples (excluding Jurkat samples), we observed that a given guide always generated the same type of outcome (**Supp. Fig. 2B,E**). Editing outcome was not associated with cell type or genome site (**Supp. Fig. 2C,D,F,G**). We compared guide sequences within each outcome group and observed that most low-indel guides contained a guanine at protospacer position -2 (**Supp. Fig. 2H**), a feature previously associated with increased Cas9 cleavage efficiency[2,3]. High-indel guides exhibited limited sequence similarity (**Supp. Fig. 2I**). These results suggest that certain guides may minimize background indel formation and maximize homology-free T7 promoter insertion.

**Supplementary Note 3. Development of in vitro and in situ transcription of genome edit sites.**

As a first step toward IST of inserted phage promoters, we started with in vitro transcription (IVT) on purified genomic DNA (**Ext. Data Fig. 2A**). RNA was only generated in the presence of T7 polymerase and not in its absence, indicating functional T7 transcription (**Supp. Fig. 3A**). However, RNA was generated in the absence of T7 promoter insertion (**Supp. Fig. 3A**). Since T7 polymerase activity is promoter-specific[4], we suspected that RNA was generated from endogenous human sequences that are similar to the T7 promoter. To examine this, we aligned the canonical 18 bp T7 promoter sequence (5'-TAATACGACTCACTATAG-3')[4] to the human reference genome (hg38) using BLAST[5] and identified 14 promoter-like sequences with at least 90% sequence similarity (16–18 bp), and five sequences containing an terminal guanine that initiates T7 transcription (**Supp. Table 2**)[6]. To determine whether these endogenous sites engage the T7 RNA polymerase, we selected the three most similar sequences (17–18 bp similarity and terminal guanine) and quantified T7 transcription at these sites in three cell types by reverse-transcription quantitative PCR (RT-qPCR). Two of the three sequences gave 50 fold induction of T7 RNA compared to five control sites without sequence similarity (**Supp. Fig. 3B,C**). This indicates that endogenous T7 promoters are a potential source of background T7 transcription.

Next, we performed T7 IVT on genomic DNA from four cell types (including primary T cells) with promoter-labeled edits at two genome sites (*B2M* and *CTLA4*). We quantified T7 RNA by RT-qPCR and observed consistent 500 fold induction of T7 RNA at promoter-labeled Cas9 edit sites (**Ext. Data Fig. 2B,C**), which was 10 fold higher than the strongest promoter-like site on chromosome 6 identified by BLAST (**Supp. Fig. 3C,D**). Next we characterized the length and direction of T7 RNA extension. Most transcripts extended < 1 kilobase (kb) from the edit site, with a small fraction of transcripts extending > 10 kb away (**Supp. Fig. 3C,D**), indicating a concentration of short T7 transcripts near to Cas9 cleavage sites. At background sites, we observed T7 RNA only downstream (3') of the promoter sequence. At Cas9 edits however, T7 RNA extended in both directions with similar abundance, indicating homology-free insertion of the T7 promoter sequence in both orientations (**Supp. Fig. 3B–D**). This bi-directional transcriptional signature distinguished genuine Cas9 edit sites from background T7 transcription occurring at endogenous promoter-like sequences that resembled the T7 promoter sequence. Together, these IVT results demonstrate that promoter-labeled Cas9 edits encode a functional T7 promoter that can be used to report Cas9 edit events with distinct RNA features.

Next, to determine the performance of T7 RNA generation on intact chromatin within nuclei, we performed T7 IST on nuclei isolated in ATAC-seq buffer[7] (**Ext. Data Fig. 2D**). We performed nuclei isolation on three cell lines with promoter-labeled Cas9 edits at *B2M* or *CTLA4*, followed by immediate IST, total RNA extraction, and RT-qPCR. Our results were comparable to those from IVT on genomic DNA, with up to 500 fold induction of T7 RNA at T7 promoter insertions (**Ext. Data Fig. 2E,F, Supp. Fig. 4A–D**), indicating effective IST on chromosomal DNA. IST transcripts were not detectable at the 10 kb range at either loci (**Supp. Fig. 4A–C**), suggesting that nuclei IST generates shorter, more localized T7 RNA compared to IVT (**Supp. Fig. 3C,D**). Next we compared T7 RNA abundance to that of endogenous mRNA from *RPL24*, an extremely expressed ribosome subunit. We observed that levels of T7 RNA at *CTLA4* approached those of *RPL24* mRNA (**Supp. Fig. 4A,B**). LinRegPCR[8] determined similar PCR efficiencies between T7 RNA and *RPL24* assays (within 7.4%, **Supp. Table 2**), indicating similar RNA abundance. Levels of T7 RNA at the *B2M* locus were more difficult to compare due to elevated *B2M* mRNA background (**Supp. Fig. 4C,D**), indicating that downstream sequencing analysis would require distinguishing reads from T7 and endogenous transcripts. These results establish that IST can generate high levels of edit-marking T7 RNAs.

Next we developed T7 IST in fixed cells (**Ext. Data Fig. 2G**) and established IST compatibility for SPLiT-seq[9]. To adapt IST reaction conditions for SPLiT-seq, we performed cell fixation using an off-the-shelf SPLiT-seq kit, and optimized IST incubation temperature, time, and NTP concentration. Under optimal IST incubation conditions (40°C, 24 hours, 10 mM NTPs), we achieved levels of T7 RNA up to 16 fold higher than *RPL24* mRNA, an extremely expressed ribosome subunit, with minimal increase in background signal (**Supp. Fig. 5**). High levels of T7 RNA were retained in the cell pellet fractions of T7 IST reactions and were resistant to washout (**Ext. Data Fig. 2H**, **Supp. Fig. 5D**), suggesting that T7 transcripts would be retained through downstream single-cell barcoding. To determine whether IST is efficient at different genome sites, we performed T7 IST with seven guide RNAs targeting the coding exons of four chromatin remodeler genes (*ARID1A*, *SMARCA4*, *CHD3*, *CHD4*). We observed high T7 transcription at all loci except *ARID1A* (**Ext. Data Fig. 2I, Supp. Fig. 6**), where both edit sites exhibited low T7 signal, likely because our RT-qPCR primers were designed to amplify sequences ~500 bp further from the edit sites at this locus (due to GC-rich sequences).

**Supplementary Note 4. Bulk RNA-seq of in situ T7 transcripts.**

Next we established that Cas9 edit sites can be identified by bulk RNA sequencing of T7 transcripts (**Ext. Data Fig. 3A**). We took two cell types (Jurkat and K562) edited with T7 promoter labeling at two genome sites (*CTLA4* and *B2M*) and performed nuclei isolation and IST. We then generated a 3' random hexamer-primed RNA-seq library to match the capture chemistry of SPLiT-seq libraries, and performed low-depth, paired-end sequencing. In the absence of T7 promoter or T7 RNA polymerase, no reads were observed at either Cas9 edit site (**Ext. Data Fig. 3B,C**). However, cells that received both T7 promoter and T7 polymerase exhibited read pileups centered at expected Cas9 cut sites. Reads began with T7 barcode sequences encoded by inserted T7 promoters (`5'-GGGAGAGTAT-3'`), followed by genome-aligned sequence that began at the exact position of expected Cas9 cleavage, 3 bp upstream of the protospacer activation motif (PAM)[10] (**Ext. Data Fig. 3B,C**). Each edit site showed divergent, opposite-stranded reads that mapped within 200 bp of expected edit sites, consistent with RT-qPCR observations of short, bi-directional T7 RNAs (**Supp. Fig. 3B–D**). Some reads began less than 100 bp from the edit site but lacked T7 barcodes, likely from truncated T7 cDNA caused by library fragmentation. Together, these results establish that Cas9 genome edits can be identified by IST and T7 transcript sequencing.

**Supplementary Note 5. Superb-seq on chromatin remodeler complexes.**

We designed guide RNAs to target the chromatin remodeler complexes BAF and NuRD, since perturbing these complexes would result in a large number of transcriptomic changes that could be validated with existing ChIP-seq datasets[11]. We prepared K562 cell lines with T7 promoter-labeled Cas9 edits by RNP electroporation, using seven guide RNAs targeting early coding sequences of four genes (**Fig. 2A, Supp. Table 1**): BAF complex members *ARID1A* and *SMARCA4*, and NuRD complex members *CHD3* and *CHD4*[12]. We generated three sample pools consisting of: 1) unedited electroporated K562 cells, 2) five cell lines that were edited with either at *ARID1A*, *SMARCA4*, or both genes (BAF pool), 3) three cell lines that were edited at either *CHD4* or both *CHD3* and *CHD4* (NuRD pool, **Fig. 2A**). We verified that these chromatin remodeler edit sites generate T7 transcripts (**Supp. Fig. 6**).

To perform Superb-seq, we performed IST reactions on thawed aliquots of fixed sample pools, followed by combinatorial indexing of single-cell RNA with an off-the-shelf SPLiT-seq kit (**Fig. 2A**, **Supp. Figs. 7B,8**). We applied the same IST treatment to both unedited and edited sample pools so that we could control for background transcription by T7 RNA polymerase (**Supp. Fig. 4A–D**) when quantifying gene expression. We generated two Superb-seq libraries, a 500 cell library and a 10,000 cell library (**Supp. Fig. 7C,D**). T7 cDNA was detected in both cDNA libraries (**Supp. Fig. 7E**), indicating successful capture of T7 fragments. To assess library quality, we performed deep paired-end sequencing (200,000 reads per cell) of the 500 cell library, and used Split-pipe[13] to align reads to the human reference genome, correct cell barcodes for sequencing errors, call high-quality cells, and perform initial read-gene assignments. In total, there were 126 million high-quality reads with a mapping rate of 66.7%, and 583 high-quality cells with 217,000 reads per cell (**Supp. Fig. 7F**). We then performed paired-end sequencing and Split-pipe analysis of the 10,000 cell library at a depth of ~100,000 reads per cell. We observed 937 million high-quality reads with a mapping rate of 78.8%, and 9,500 cells with 98,600 mean reads per cell (**Supp. Fig. 7F**). These results demonstrate that the Superb-seq workflow generates high-quality combinatorial scRNA-seq reads.

**Supplementary Note 6. Correction of ambient T7 reads.**

Since ambient T7 transcripts were observed in IST supernatants (**Ext. Data Fig. 2H, Supp. Fig. 5**), we estimated and corrected for cross-contaminating ambient T7 reads. In short, we determined the guide associated with each edited cell, and defined ambient T7 reads as those at edit sites associated with other guides (**Methods**, **Supp. Fig. 10A,B, 11A,B, Supp. Table 3**). We observed an elevated rate of ambient RNA in the BAF sample pool in both the 500 cell and 10,000 cell libraries (29–38% of T7 UMIs) compared to the NuRD sample pool (0.6–0.7% of T7 UMIs, **Supp. Fig. 10A,B, 11A,B**). This elevated ambient rate was due to both an elevated editing rate (29 BAF edit sites versus 14 NuRD edits) and an abundant off-target edit event in the intron of *ADSS1* which accounted for 73–85% of ambient T7 UMIs in the BAF sample (**Supp. Fig. 10A,B, 11A,B, Supp. Table 3**). Because a small number of edit sites were responsible for most ambient T7 UMIs, removing these reads had only a modest effect on the edit events called across cells (**Supp. Fig. 10C,D, 11C,D, Supp. Table 3**). Thus T7 ambient reads can be abundant at high efficiency sites, similar to ambient RNA from highly expressed genes in standard scRNA-seq[14], and ambient guide RNA in Perturb-seq[15].

**Supplementary Note 7. Assessment of edit allele quantification.**

To assess edit allele quantification, we used edit allele counts from the lower-depth 10,000 cell library to estimate expected edited cell counts and edit allele counts in the higher-depth 500 cell library. The expected and observed counts in the 500 cell library were highly correlated (edited cell Pearson's $R$ = 0.985, two-sided $p = 6.9 \times 10^{-33}$; edit allele Pearson's $R$ = 0.976, two-sided $p = 6.8 \times 10^{-29}$; **Supp. Fig. 14G,H**). At the *CHD4* on-target edit site, however, edited cell and allele counts were higher than expected. This suggests that *CHD4* edit alleles are undercounted in the 10,000 cell library, likely due to the high rate of editing at this site (**Fig. 1E**). These results establish that Sheriff effectively quantifies the number of edited cells and single-cell edit allele dosage, although edit allele dosage was underestimated at one of the 43 edit sites.

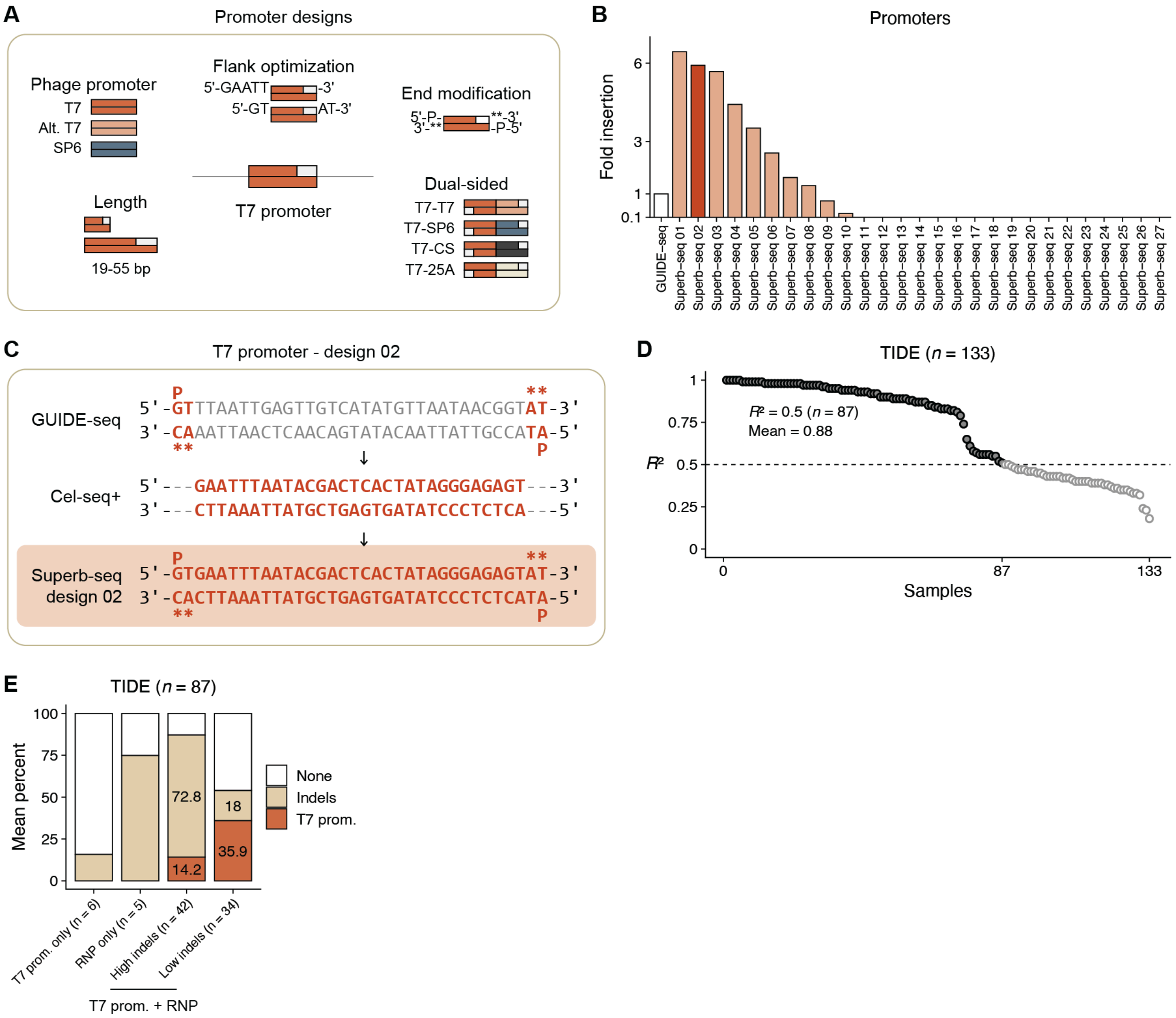


**Supplementary Figure 1. Development of phage T7 promoter labeling of genome edits.** (**A**) Design categories of 27 candidate Superb-seq phage promoter constructs. (**B**) Insertion rate of the 27 candidate Superb-seq T7 promoter designs relative to the GUIDE-seq "dsODN", measured by TIDE. Value for design 2 is the mean of four samples. (**C**) Sequences of the GUIDE-seq dsODN[16], Cel-seq+ T7 promoter[17], and the derived Superb-seq T7 promoter. Terminal 5' phosphate (P) and 3' phosphorothioate (askerisk) modifications are indicated. (**D**) TIDE R-squared ($R^2$) values from 112 Sanger sequencing samples. Samples with $R^2$ > 0.5 ($n$ = 70) were used for downstream analyses, and are indicated in red. (**E**) Frequency of editing outcomes (mean percent) for indicated sample groups. Possible editing outcomes are unedited (none), background edits (Indels), and T7 promoter insertion (+30 bp insertion or greater).

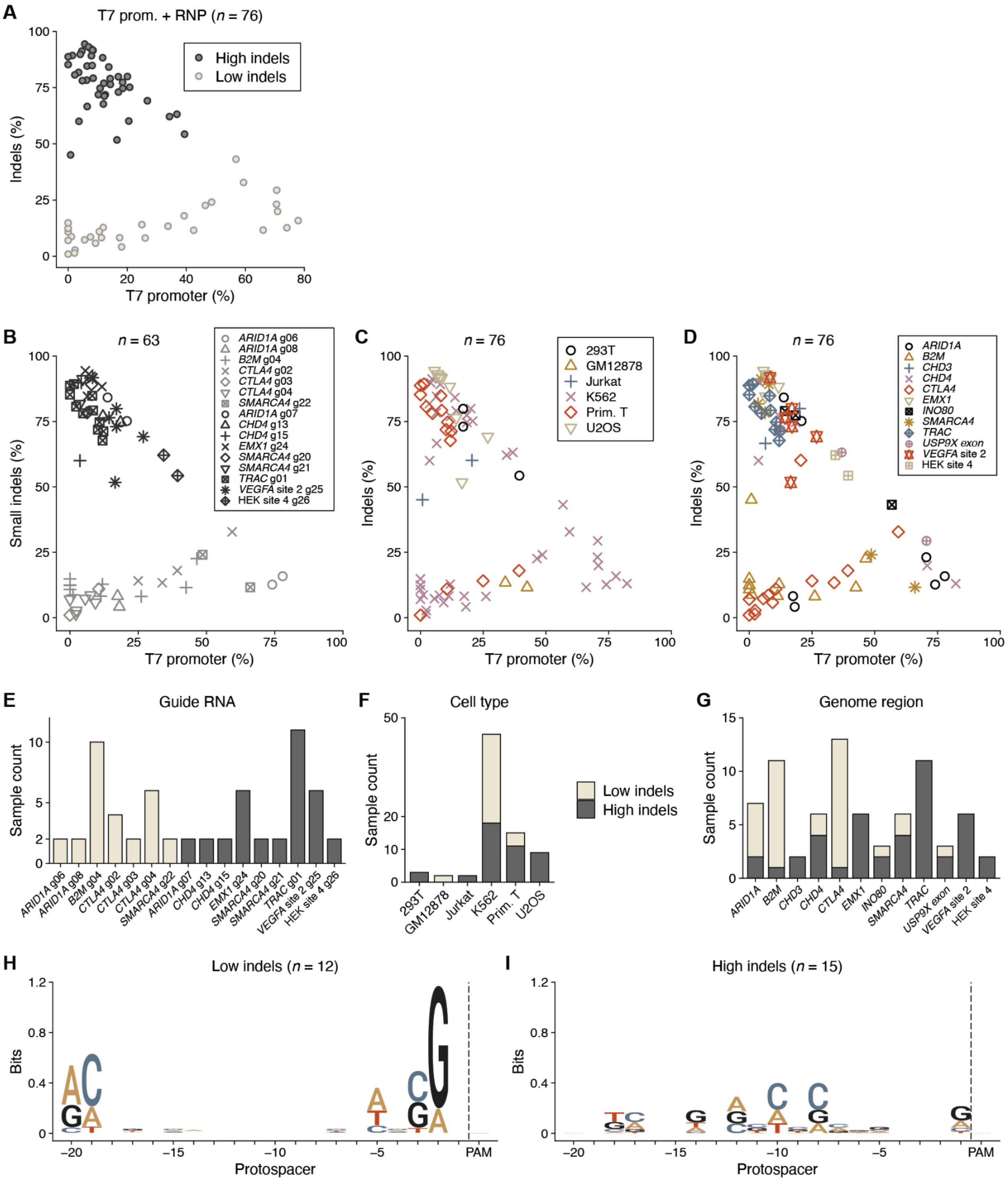


**Supplementary Figure 2. Bimodal background indel rate associates with guide RNA.** (**A–D**) Frequency of T7 promoter insertion and variable small indel events in 76 T7 promoter edited samples, labeled by (**A**) outcome group (High or Low indels), (**B**) guide RNA, (**C**) cell type, or (**D**) target genome region. (**E–G**) Frequency of high and low indel samples, grouped as in **B–D**. (**H,I**) Positional Shannon entropy (bits) of guide sequences for (**H**) the 10 guide RNAs used in the 32 "low indel" samples, or (**I**) the 10 guides used in the 27 "high indel" samples, calculated using ggseqlogo[18].

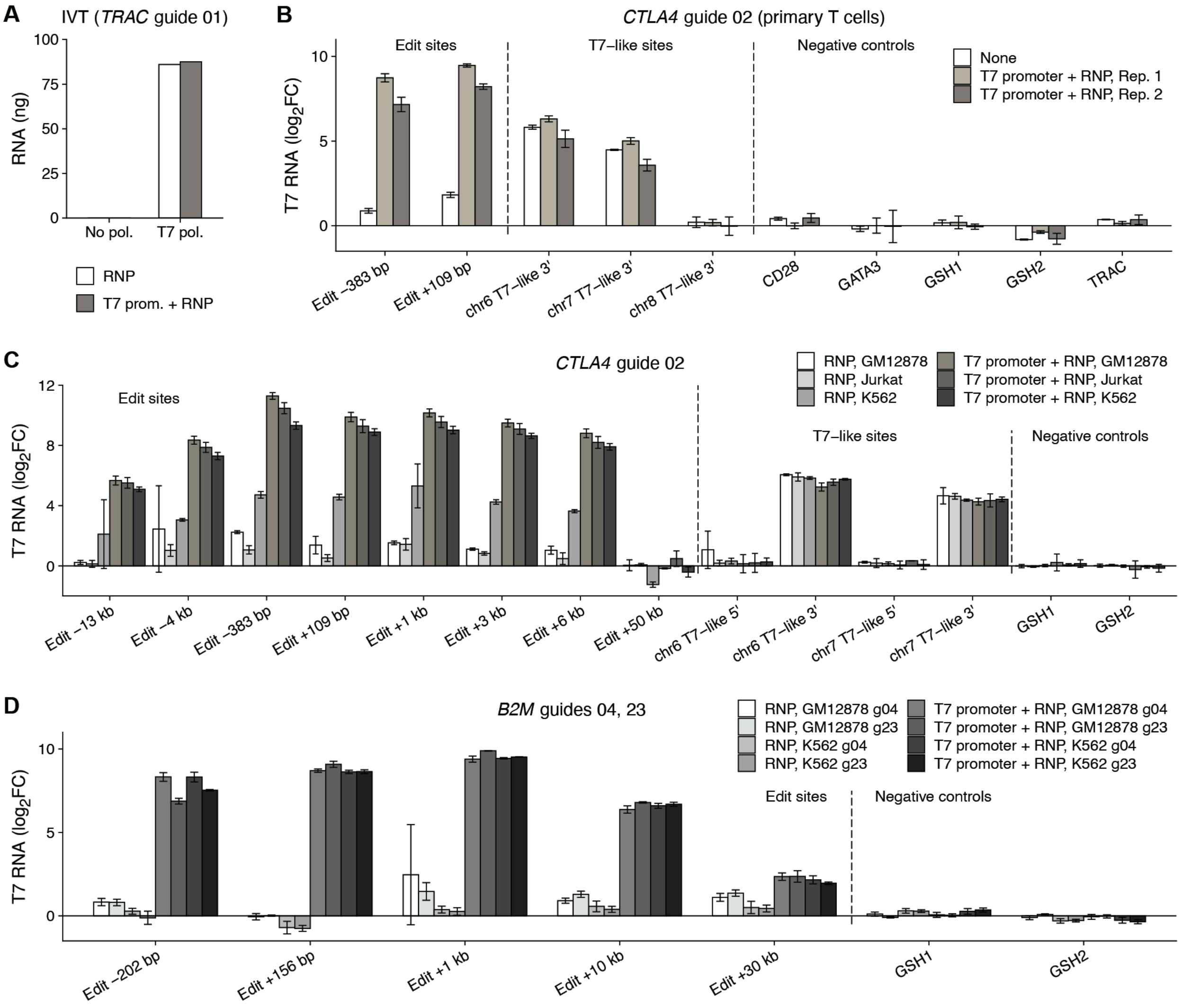


**Supplementary Figure 3. Generation of in vitro T7 transcripts at genome edit sites.** (**A**) RNA yield from IVT on genomic DNA from primary human T cells, measured by fluorometry. Cells were treated with RNP alone (RNP) or with T7 promoter and RNP (T7 promoter + RNP). IVT was performed with T7 RNA polymerase (T7 pol) or without polymerase (No pol). (**B–D**) Relative levels of IVT transcripts at T7 promoter-labeled genome edits, measured by RT-qPCR and the comparative $C_t$ method[19]. Target genome locus, guide RNA, and cell type are indicated. Edit-targeted PCR assays are named by their position relative to the edit site (e.g. Edit +109 bp). Endogenous promoter-targeted assays are named by their position relative to promoter orientation (e.g. chr6 T7-like 3'). Reference assays targeted non-coding sequences at three genome loci (*CD28*, *GATA3*, *TRAC*) and two intergenic "genomic safe harbor" sites (GSH1, GHS2)[20]. Results are represented as $\log_2$ fold change ($\log_2$FC), normalized to GSH assays. Bars represent the mean of technical replicates (*n* = 3), and whiskers represent 2 × standard error in the mean (SEM).

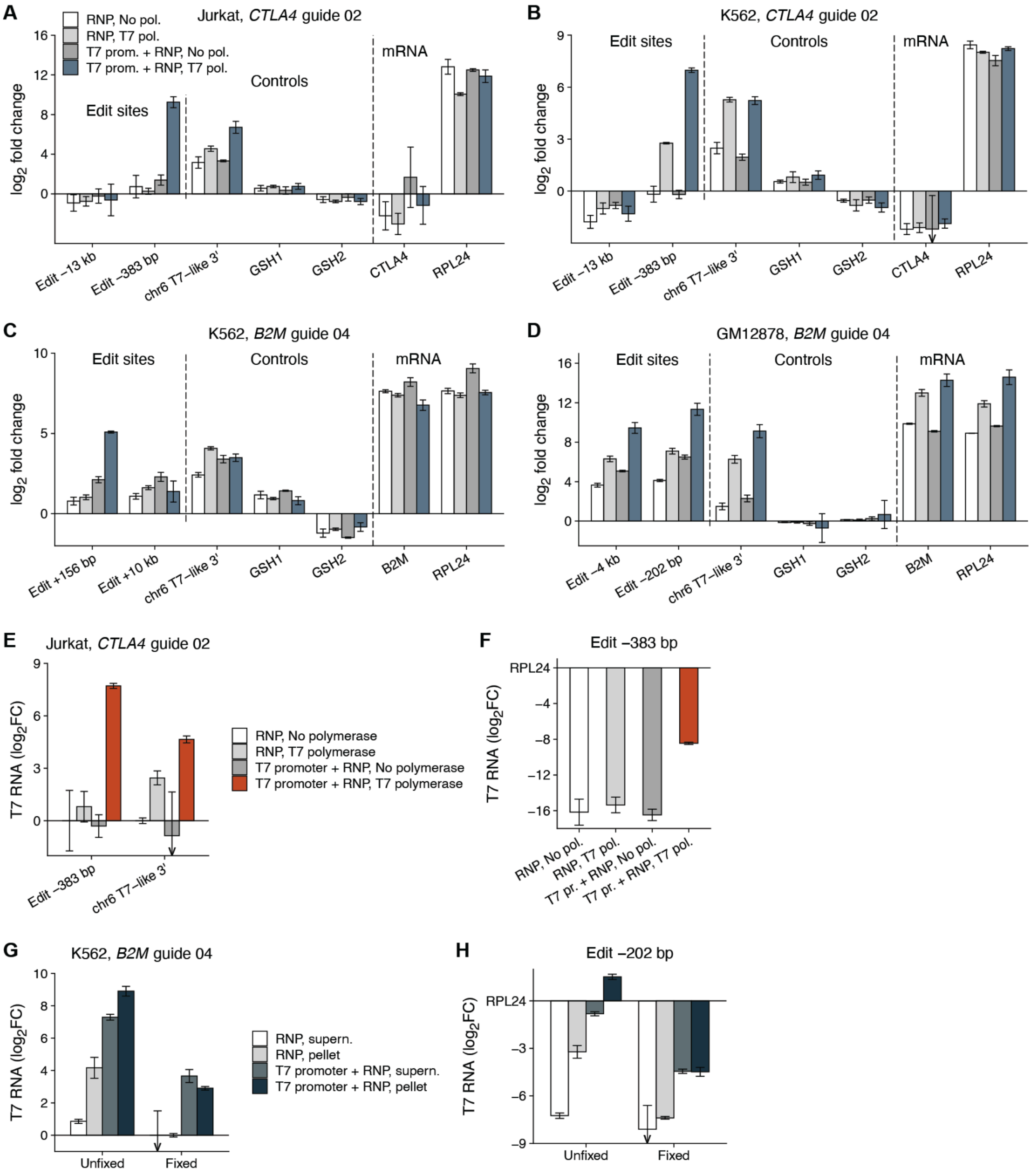


**Supplementary Figure 4. Generation of in situ T7 transcripts in nuclei and fixed cells.** (**A–D**) Relative RNA levels from IST on unfixed nuclei, measured by RT-qPCR and comparative $C_t$. Results are represented as $\log_2$ fold change ($\log_2$FC) normalized to GSH1 and GHS2 assays. (**E,F**) Levels of T7 RNA from IST on fixed cells. (**G,H**) Level of T7 RNA from IST on unfixed or fixed nuclei from the same nuclei sample. Results are normalized to (**E,G**) reference genes *RPL24* and *RPS10*, and the control cell treatment, or (**F,H**) *RPL24* gene expression. Bars represent the mean of technical replicates ($n$ = 3), and whiskers represent 2 × SEM.

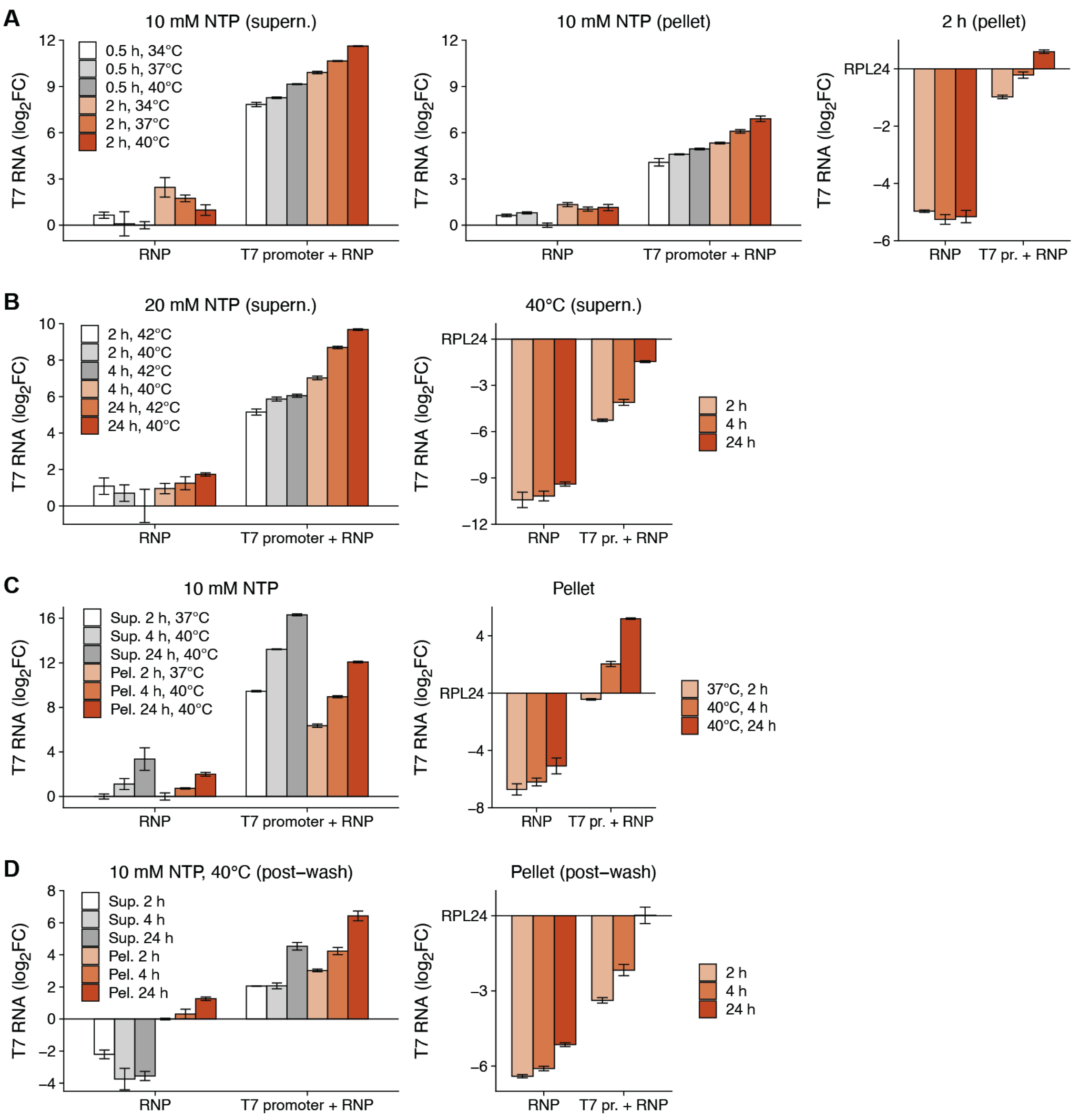


**Supplementary Figure 5. Optimization of in situ transcription for SPLiT-seq fixed cells.** (**A–D**) Relative level of IST transcripts generated in fixed K562 cells under variable IST reaction conditions (NTP concentration, incubation time, incubation temperature), measured by RT-qPCR and comparative $C_t$. K562 cells were treated with Cas9 RNP alone (RNP) or with T7 promoter (T7 promoter + RNP) using *B2M* guide 4. RT-qPCR was performed on total RNA extracted from the supernatant (Sup.) or cell pellet (pellet) fraction of IST reactions. For **D**, IST cell pellets were washed once by resuspending in SPLiT-seq cell buffer, and total RNA fractions were isolated from this resuspension. Results are represented in two ways. First, as $\log_2$ fold change ($\log_2$FC) normalized to reference genes *RPL24* and *RPS10*, and sample treated with RNP only (no T7 promoter). Second, as $\log_2$FC normalized to *RPL24* gene expression. All bars represent the mean of technical replicates ($n$ = 3), and whiskers represent 2 × SEM.

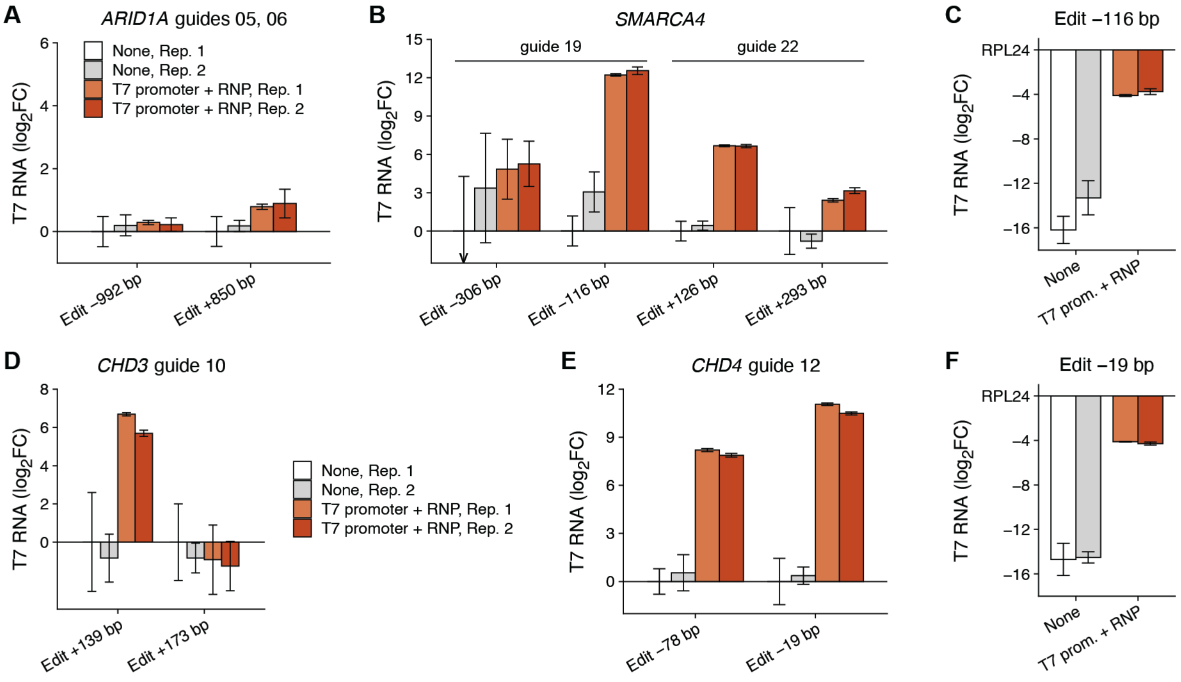


**Supplementary Figure 6. In situ transcription at edit sites in chromatin remodeler genes.** (**A,B**) Relative level of IST transcripts generated in pools of fixed K562 cell lines containing T7 promoter-labeled Cas9 edits in chromatin remodeler genes, measured by RT-qPCR. PCR assays targeted edit sites at (**A**) *ARID1A* or (**B,C**) *SMARCA4* of the BAF complex, and (**D**) *CHD3* or (**E,F**) *CHD4* of the NuRD complex. Three cell pools were used: Mock-electroporated cells (None), *ARID1A*- and *SMARCA4*-edited cells (**A–C** T7 promoter + RNP), and CHD3- and CHD4-edited cells (**D–F** T7 promoter + RNP). IST replicates (R1, R2) are indicated. Results are represented as (**A,B,D,E**) $\log_2$ fold change ($\log_2$FC) normalized to reference genes *RPL24* and *RPS10* and a mock-treated sample, or (**C,F**), as $\log_2$FC normalized to *RPL24* gene expression. Bars represent the mean of technical replicates ($n$ = 3), and whiskers represent 2 × SEM.

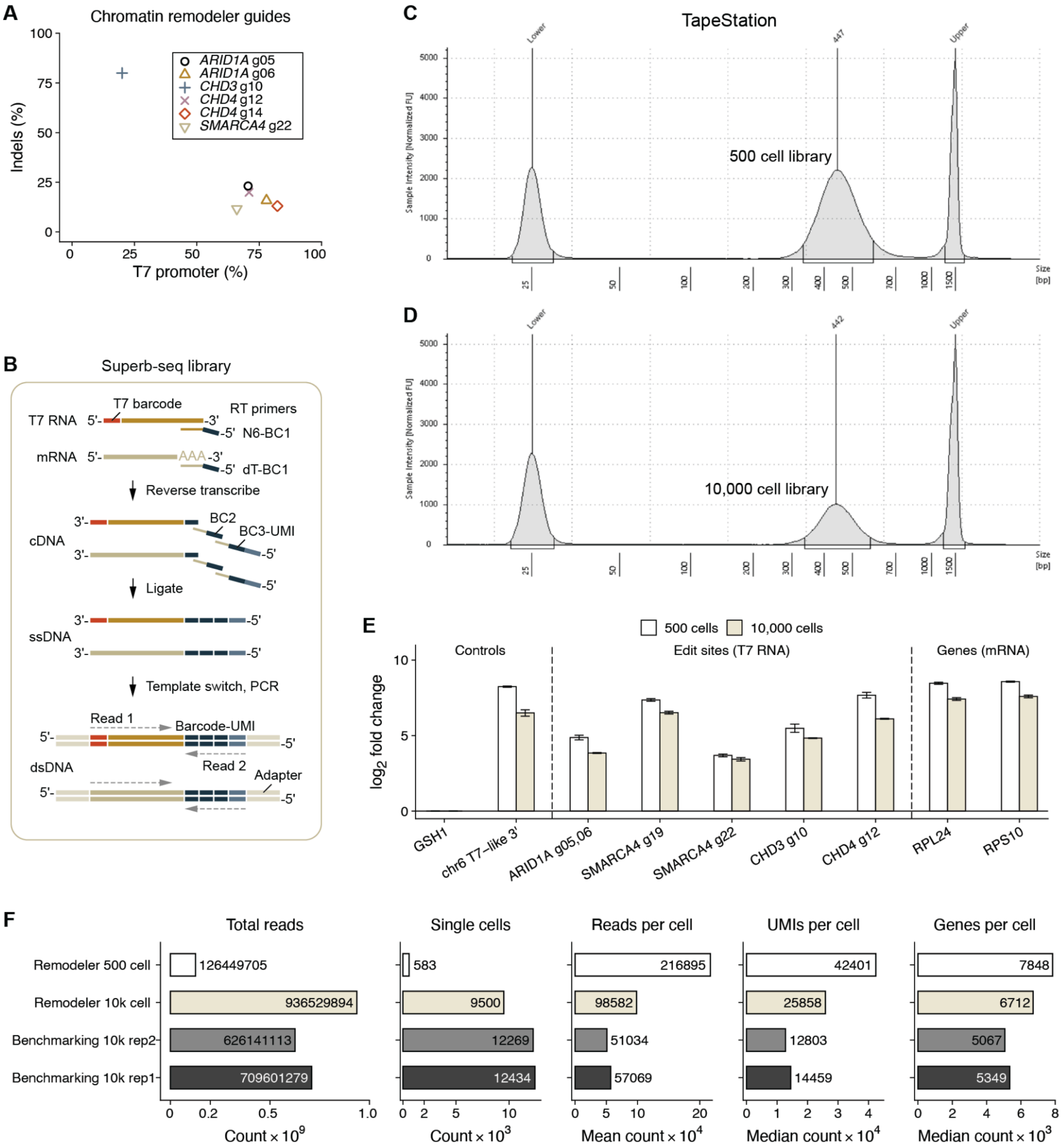


**Supplementary Figure 7. Generation of Superb-seq libraries.** (**A**) Frequency of T7 promoter insertion and background indels in K562 from six guides used in the Superb-seq chromatin remodeler experiment. (**B**) Joint combinatorial indexing of T7 and endogenous cDNA fragments with cell barcodes (BC) and unique molecular identifiers (UMI). Detailed library sequence structure is shown in **Supp. Fig. 8**. (**C,D**) Distribution of library fragment size in the (**C**) 500 cell and (**D**) 10,000 cell libraries, measured by TapeStation. (**E**) Relative levels of T7 RNA and mRNA in the Superb-seq libraries, measured by RT-qPCR and comparative $C_t$, and represented as $\log_2$ fold change ($\log_2$FC). Control sites are GSH1 (negative) and a T7 promoter-like sequence on chromosome 6 (positive). Bars represent the mean of technical replicates ($n$ = 3), and whiskers represent 2 × SEM. (**F**) Split-pipe summary statistics from chromatin remodeler and benchmarking Superb-seq libraries. Additional Split-pipe quality metrics are listed in **Supp. Table 2**.

**A** **T7 cDNA**

```
     READ 1|-------------------->
            T7 barcode  genome
5'-[TruSeq]-GGGAGAGTAT-XXXX…XXXX-NNNNNN-XXXXXXXX-[L1]-XXXXXXXX-[L2]-XXXXXXXX-NNNNNNNNNN-[TruSeq]-3'
3'-[TruSeq]-CCCTCTCATA-XXXX…XXXX-NNNNNN-XXXXXXXX-[L1]-XXXXXXXX-[L2]-XXXXXXXX-NNNNNNNNNN-[TruSeq]-5'
                                 N6-RT  Cell BC1      Cell BC2      Cell BC3    UMI
                                        <-----------------------------------------------|READ 2
  | 70 bp  | 10 bp    | ~200 bp | 6 bp | 8 bp    | 22 | 8 bp    | 30 | 8 bp    | 10 bp    | 66 bp  |  (v2)
                                                 | 12 |         | 12 |                               (v3)
```

**B** **Cell cDNA**

```
     READ 1|--------------------------->
              mRNA          polyA
5'-[TruSeq]-XXXX…XXXX-NVAAAAAAAAAAAAAAA-XXXXXXXX-[L1]-XXXXXXXX-[L2]-XXXXXXXX-NNNNNNNNNN-[TruSeq]-3'
3'-[TruSeq]-XXXX…XXXX-NVTTTTTTTTTTTTTTT-XXXXXXXX-[L1]-XXXXXXXX-[L2]-XXXXXXXX-NNNNNNNNNN-[TruSeq]-5'
                            dT-RT       Cell BC1      Cell BC2      Cell BC3    UMI
                                        <-----------------------------------------------|READ 2
  | 70 bp  | ~200 bp | 17 bp           | 8 bp    | 22 | 8 bp    | 30 | 8 bp    | 10 bp    | 66 bp  |  (v2)
                                                 | 12 | 8 bp    | 12 |                               (v3)
```

```
        Key                                           Linker sequence (Read 2)
        BC1-3  = Cell + sample barcodes               L1  5'-CCACAGTCTCAAGCACGTGGAT-3'   (v2)
        UMI    = Unique molecular identifier              5'-TCCAACCACCTC-3'             (v3)
        N6-RT  = Random hexamer RT primer
        dT-RT  = Oligo-dT RT primer                   L2  5'-AGTCGTACGCCGATGCGAAACATCGGCCAC-3'  (v2)
        TruSeq = Dual index adapters                      5'-ATGAGNGGTCAG-3'                (v3)
```

**C** **Adapters**

```
                                                                   READ 1|----->
  P5 adapter                                TruSeq R1 adapter
5'-AATGATACGGCGACCACCGAGATCTACAC-NNNNNNNN-ACACTCTTTCCCTACACGACGCTCTTCCGATCT-[cDNA]…
3'-TTACTATGCCGCTGGTGGCTCTAGATGTG-NNNNNNNN-TGTGAGAAAGGGATGTGCTGCGAGAAGGCTAGA-[cDNA]…
                                     i5
                                  <-------|INDEX 2
  | 29 bp                         | 8 bp   | 33 bp                            |~200 bp|


                                                             INDEX 1|------->
                                   TruSeq R2 adapter                   i7
                         …[BC-UMI]-AGATCGGAAGAGCACACGTCTGAACTCCAGTCAC-NNNNNNNN-ATCTCGTATGCCGTCTTCTGCTTG-3'
                         …[BC-UMI]-TCTAGCCTTCTCGTGTGCAGACTTGAGGTCAGTG-NNNNNNNN-TAGAGCATACGGCAGAAGACGAAC-5'
                                                                                           P7 adapter
                           <-------|READ 2
                    (v2)  | 86 bp  | 34 bp                         | 8 bp   | 24 bp                 |
                    (v3)  | 58 bp  |
```

**Supplementary Figure 8. Structure of Superb-seq libraries.** (**A**) T7 RNA-derived cDNA fragments. Read 1 contains the 5' T7 barcode and genome sequence at the Cas9 edit site. Read 2 contains the UMI and SPLiT-seq three-part combinatorial cell barcode ( v2, Parse Biosciences). Cell barcode 1 (BC1) is the sample barcode. (**B**) Cell mRNA-derived cDNA fragments. Read 1 contains mRNA sequence. These are representative fragments, as both T7 RNA and mRNA can be captured by both types of RT primers. (**C**) Structure of the TruSeq dual-index sequencing adapters (Illumina). Sequences were inferred from library documentation and the Single Cell Genomics Library Structure resource (https://teichlab.github.io/scg_lib_structs/methods_html/SPLiT-seq.html)[21].

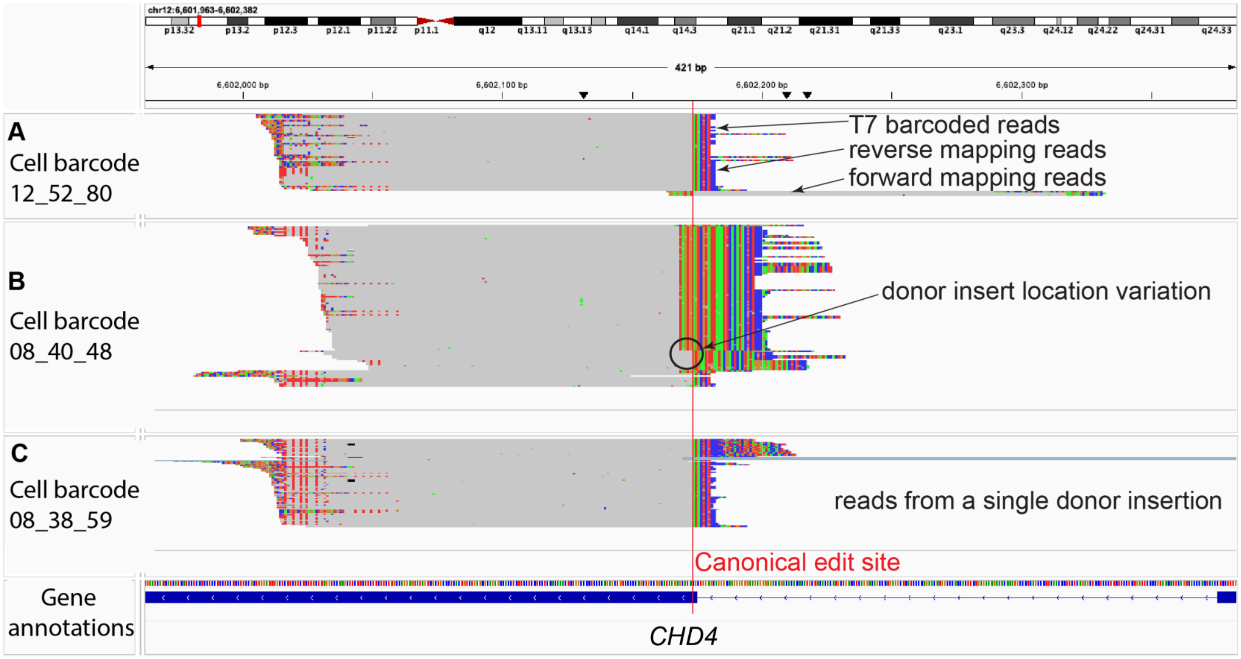


**Supplementary Figure 9. T7 read features indicate multiple edit alleles within individual cells.** Integrative Genomics Viewer (IGV) snapshot of T7 barcoded reads from the Superb-seq 500 cell library, with soft-clipped sequences and base mismatches to the reference genome visualised. The *CHD4* on-target edit site is at the locus centre, with the called canonical edit site indicated with a vertical line. (**A**) T7 barcoded reads called in the cell 12_52_80, which had both forward and reverse mapping T7 reads, mutually exclusive events that indicate more than one allele has the Superb-seq T7 promoter insertion. (**B**) T7 barcoded reads for cell 08_40_48. Differences in barcoded T7 read mapping position indicating variation in the sequence of T7 promoter insertion site, and indicating that more than one allele is edited in this cell. (**C**) T7 barcoded reads in cell 08_38_59, with no clear variation in T7 promoter insertion sequence, providing evidence indicating only one edited allele.

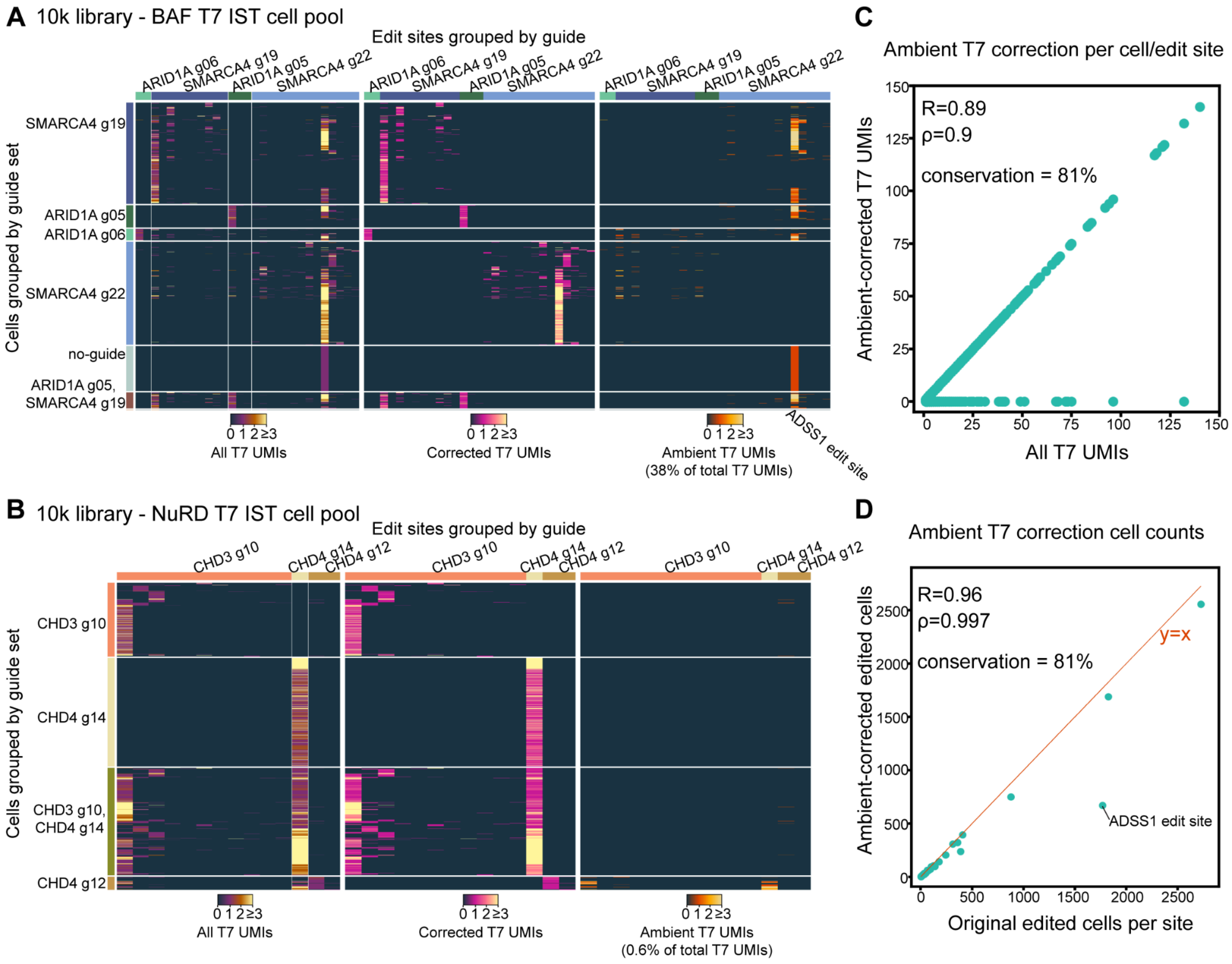


**Supplementary Figure 10. Correction of ambient T7 reads in the 10,000 cell Superb-seq library.** (**A**) Heatmaps showing the BAF in situ transcription (IST) edit pool T7 UMI counts across cells (rows) and detected edit sites (columns). Edits are colored by the causal guide, and cells are colored by the inferred guide the cell received. The first heatmap indicates all T7 UMIs, the second indicates the expected T7 UMIs based on guide combinations that were co-electroporated, and the third indicates ambient T7 UMIs where cells are detected with T7 UMIs that mark edits from guides which were not co-electroporated. (**B**) Equivalent to A, except for the NuRD T7 IST cells. (**C**) Scatter plot indicating the original T7 UMIs detected across cells and edit sites, and the y-axis indicates the ambient-corrected T7 UMIs. Pearson's $R$ and Spearman's $\rho$ are indicated. "Conservation" indicates the percentage of sites which did not change. (**D**) Equivalent to C, except showing the number of edited cells per edit site, before and after removing the ambient T7 UMI counts.

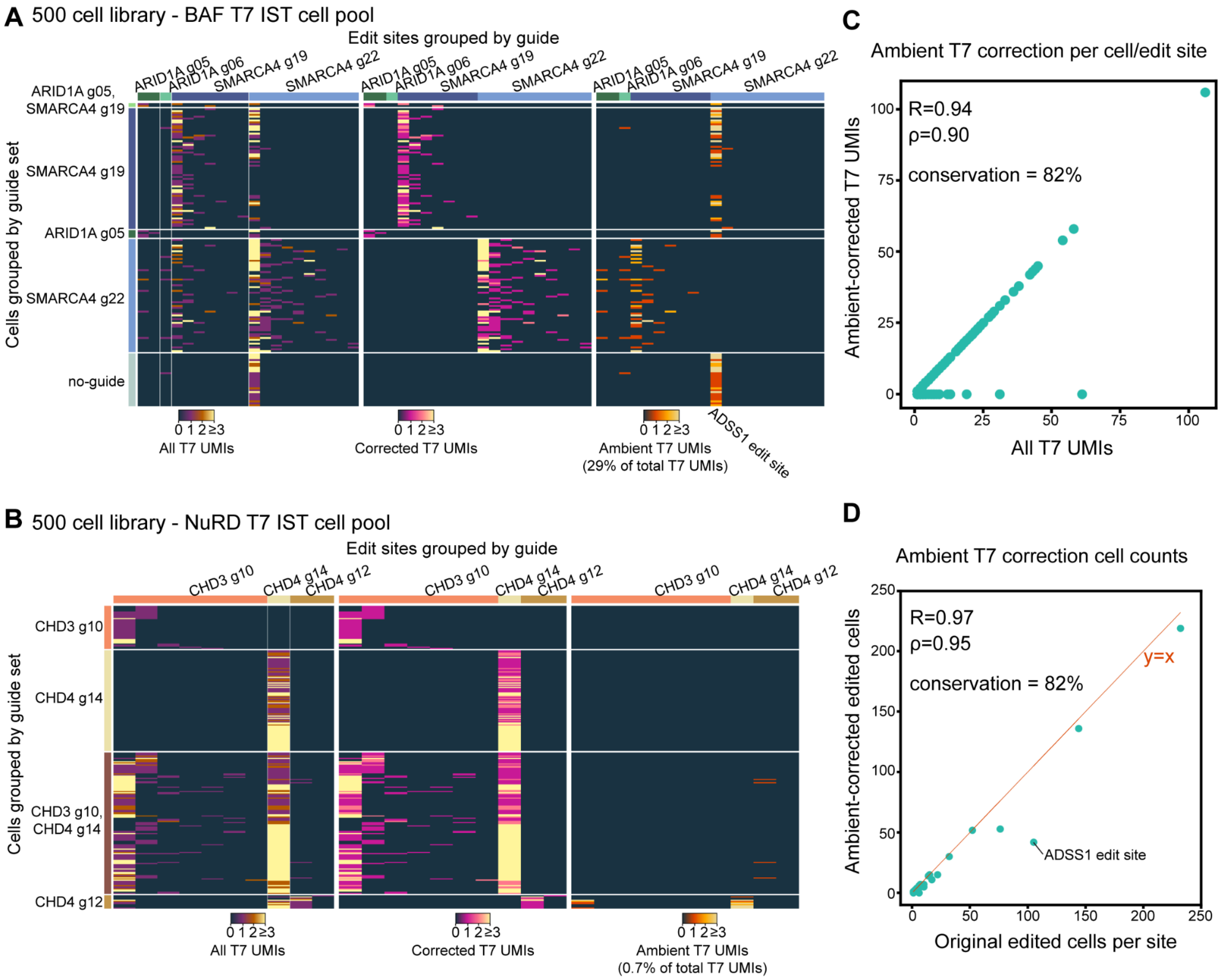


**Supplementary Figure 11. Correction of ambient T7 reads in the 500 cell Superb-seq library.** (**A**) Heatmaps showing the BAF in situ transcription (IST) edit pool T7 UMI counts across cells (rows) and detected edit sites (columns). Edits are colored by the causal guide, and cells are colored by the inferred guide the cell received. The first heatmap indicates all T7 UMIs, the second indicates the expected T7 UMIs based on guide combinations that were co-electroporated, and the third indicates ambient T7 UMIs where cells are detected with T7 UMIs that mark edits from guides which were not co-electroporated. (**B**) Equivalent to A, except for the NuRD T7 IST cells. (**C**) Scatter plot indicating the original T7 UMIs detected across cells and edit sites, and the y-axis indicates the ambient-corrected T7 UMIs. Pearson's R and Spearman's ρ are indicated. "Conservation" indicates the percentage of sites which did not change. (**D**) Equivalent to C, except showing the number of edited cells per edit site, before and after removing the ambient T7 UMI counts.

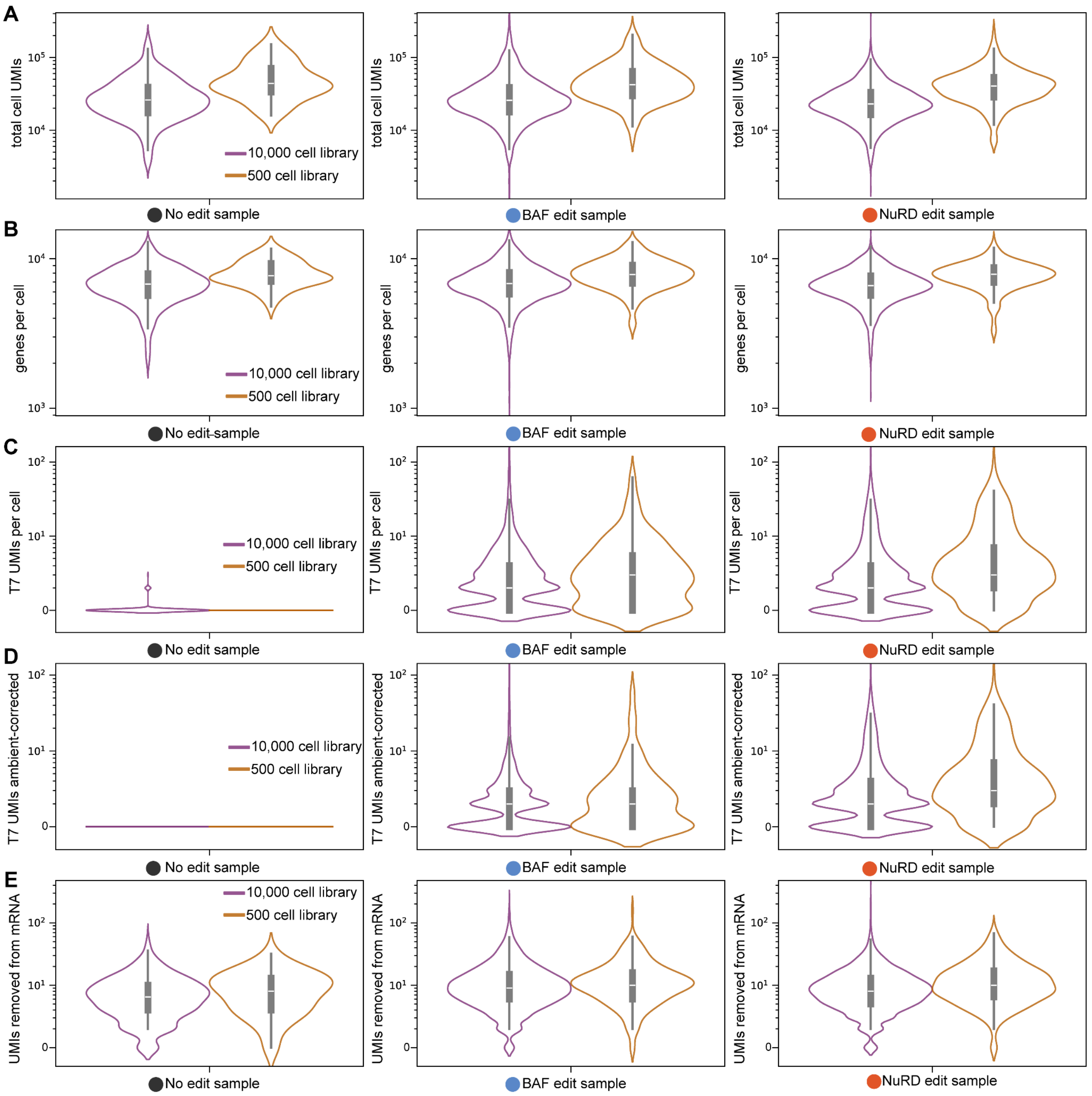


**Supplementary Figure 12. UMI and gene count distributions across Superb-seq sample pools.** (**A**) Violin plots of single-cell total UMIs captured, stratified by the chromatin remodeler library (10,000 cells or 500 cell library), with panels from left to right indicating cells from the no-edit cell pool, the BAF edited pool, and the NuRD edited pool. (**B**) Equivalent to A, except for the total genes detected. (**C**) Equivalent to A except for the T7 barcoded UMIs detected across all edit sites per cell. (**D**) Equivalent to C, except after T7 ambient UMI removal. (**E**) Equivalent to A except for the total removed UMIs occurring ± 1000 bp from detected edit sites.

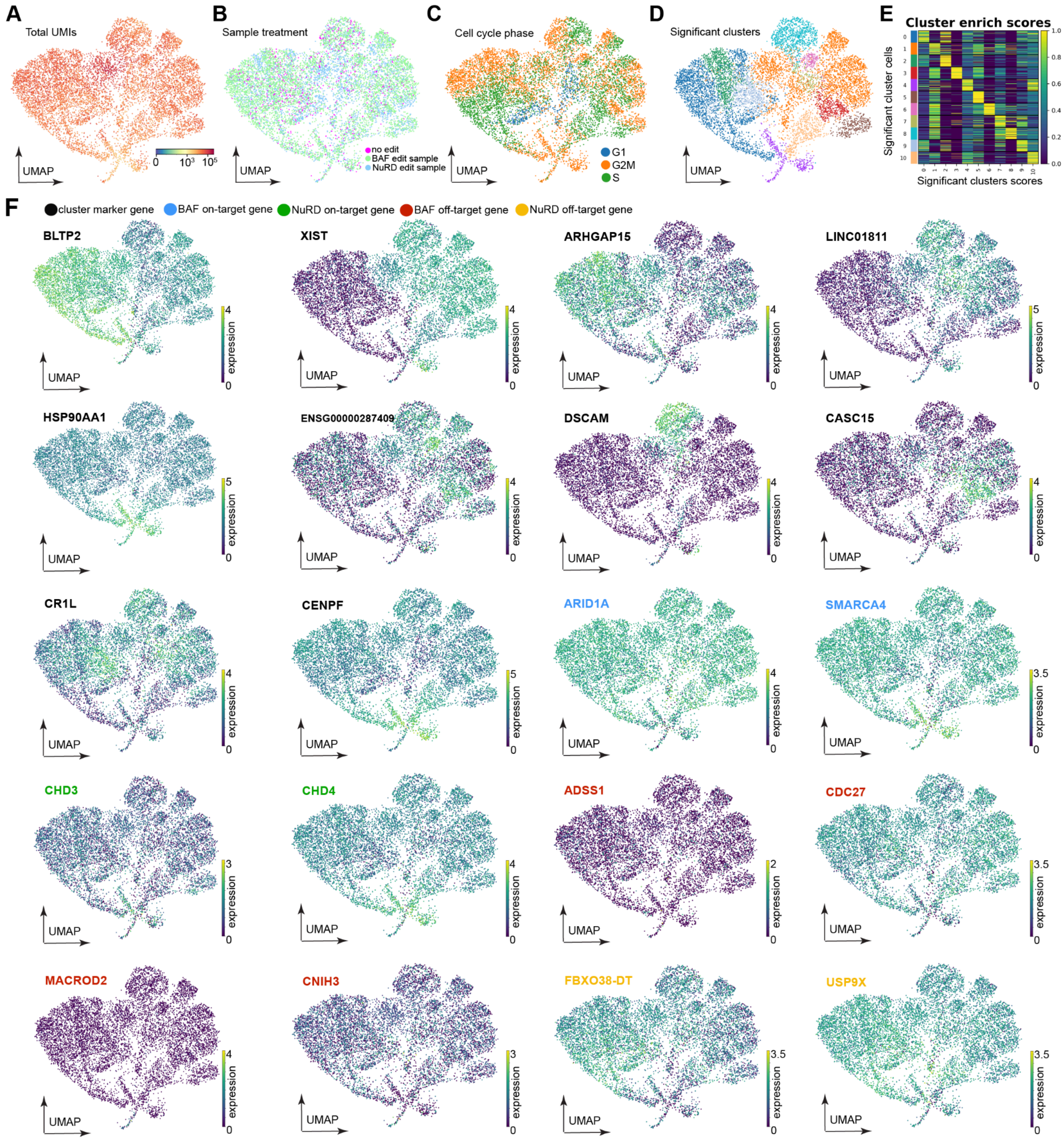


**Supplementary Figure 13. Transcriptional heterogeneity of Superb-seq transcriptomes.** (**A**) Uniform manifold approximation and projection (UMAP) of Superb-seq single-cell gene expression with total cell UMIs annotated. (**B**) UMAP with cell edit sample treatments annotated (no edit sample, BAF edit sample, NuRD edit sample). (**C**) UMAP with predicted cell cycle phase annotations. (**D**) UMAP with significantly different single-cell clusters annotated. (**E**) Heatmap with cells on the rows grouped by significant clusters, and significant clusters on the columns. Higher values indicate higher expression of the significant clusters marker genes in the single cells, showing distinct gene expression between the single-cell clusters. (**F**) UMAPs with gene expression annotated. Genes displayed are either marker genes of distinctly different clusters of cells, BAF editing on-target genes (*ARID1A*, *SMARCA4*), NuRD editing sample on-target genes (*CHD3*, *CHD4*), or off-target edited genes (*ADSS1*, *CDC27*, *MACROD2*, *CNIH3*, *FBXO38-DT*, *USP9X*).

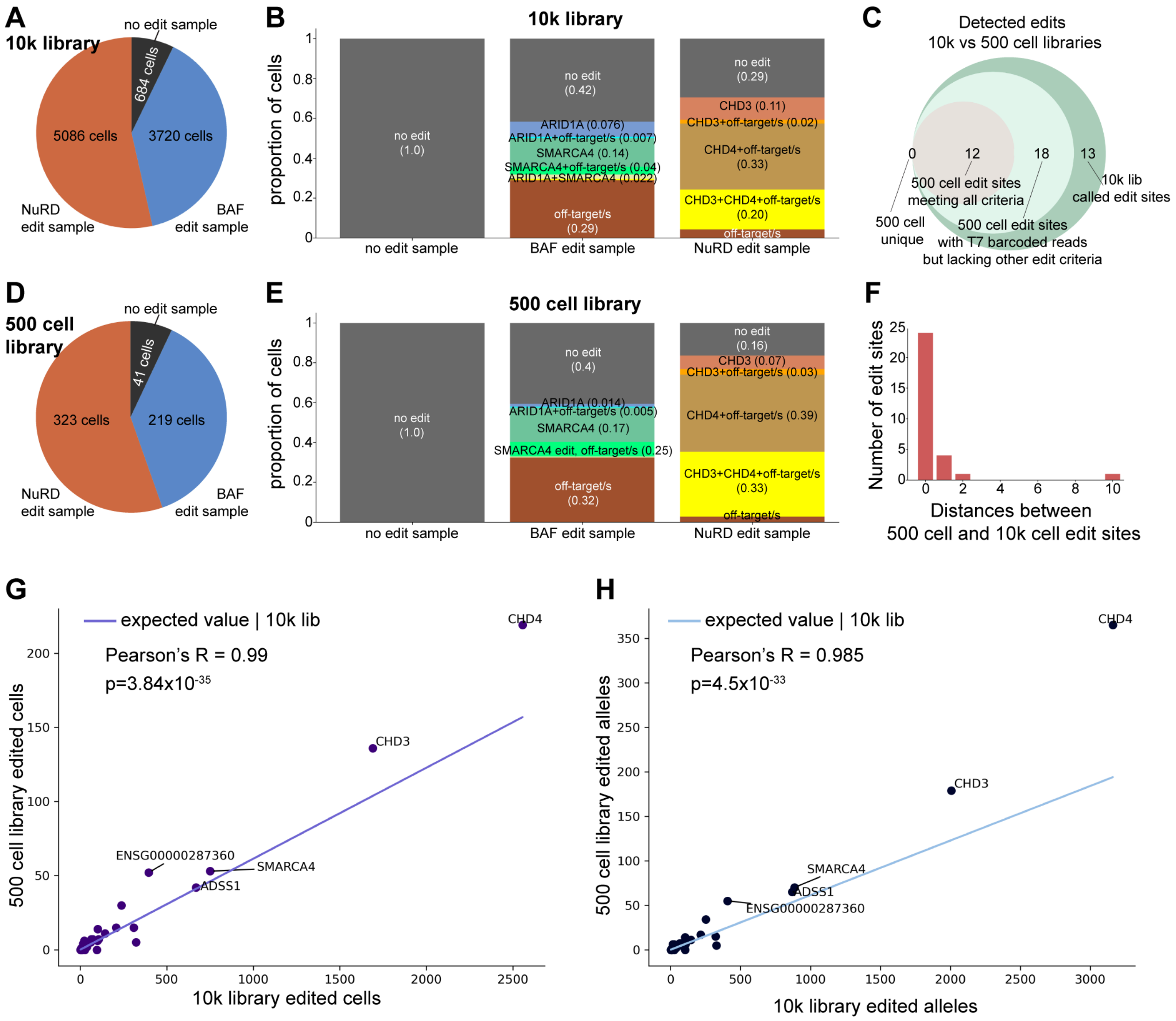


**Supplementary Figure 14. Reproducible edit capture between 500 cell and 10,000 cell Superb-seq libraries.** (**A**) Proportion of cells per sample for the 10,000 cell library. (**B**) Proportion of cells stratified by no detected cas9 editing (no edit), cells with one or more off-target edits but no on-target edits (off-targets), cells with one or more on-target gene edited (*SMARCA4* or *ARID1A* for BAF sample, *CHD3* or *CHD4* for NuRD sample), and cells with both on-target and off-target edits. (**C**) Intersection of Cas9 edits detected in the 500 cell and 10,000 cell libraries. Eighteen edits in the 500 cell library had T7 barcoded reads, but lacked additional features that Sheriff uses to call edit sites (e.g. both forward and reverse mapping T7 barcoded reads). Twelve edit sites met all criteria for *de novo* edit calling in the 500 cell library. Thirteen edit sites called in the 10,000 cell library were not observed in the 500 cell library. However the frequency of these edits was below the sampling power of 500 cells (mean expectation of 0.5 edited cells, range 0.2–1.4). (**D,E**) Equivalent to **A,B** in the 500 cell library. (**F**) Distance between equivalent edit sites detected in both Superb-seq libraries. (**G**) The number of cells edited for each edit site in the 10,000 cell library on the x-axis and the 500 cell library on the y-axis. The plotted line indicates the expected number of cells edited in the 500 cell library given the observed proportion of cells edited in the 10,000 cell library. (**H**) The equivalent to G, except with respect to the total number of edited alleles at each edit site within each respective library.

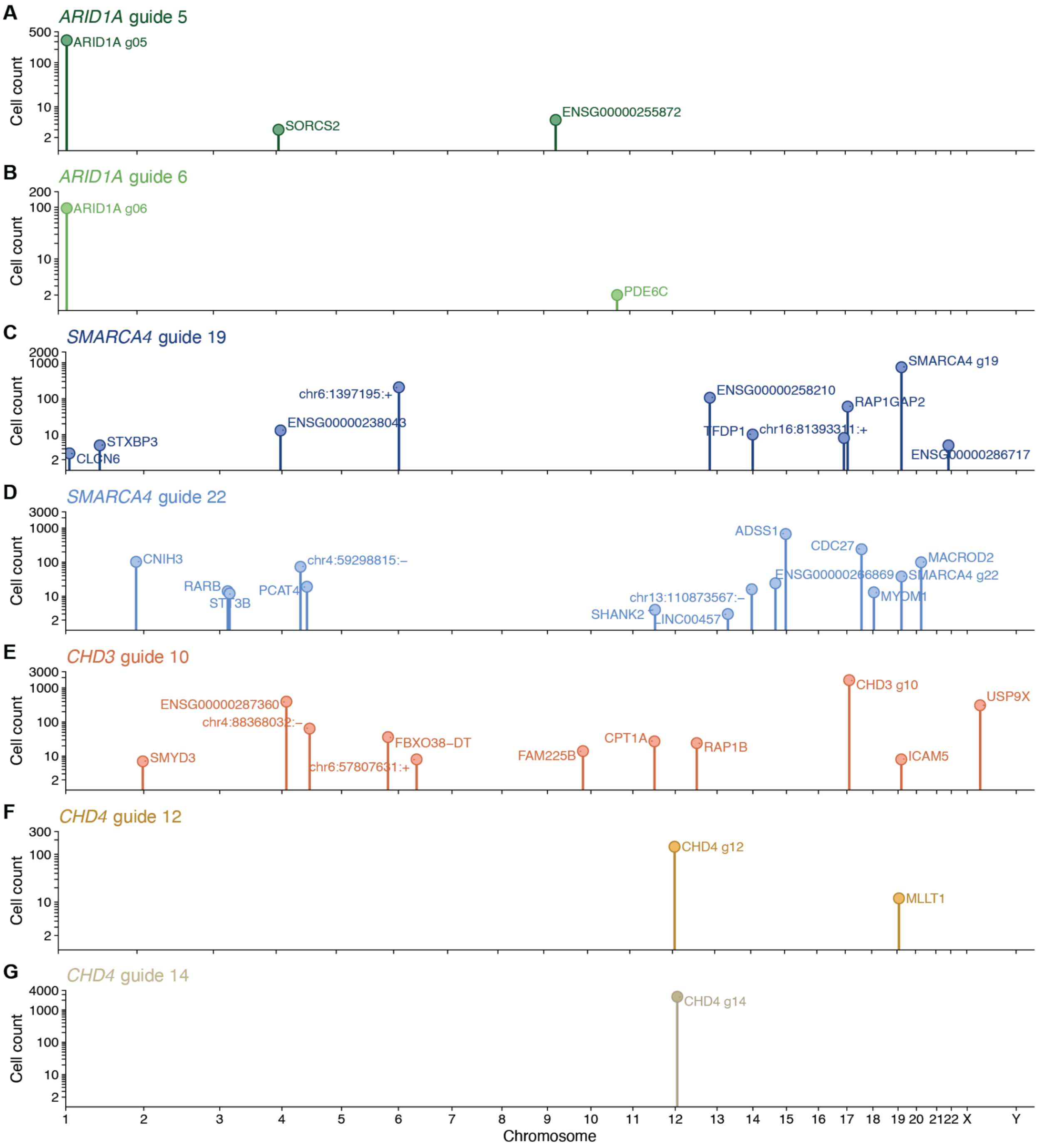


**Supplementary Figure 15. Superb-seq captures genome-wide off-target events.** (**A–G**) On-target and off-target Cas9 edit sites identified in the 10,000 cell library, separated by causal guide RNA. Genome positions (hg38) and cell counts are indicated.

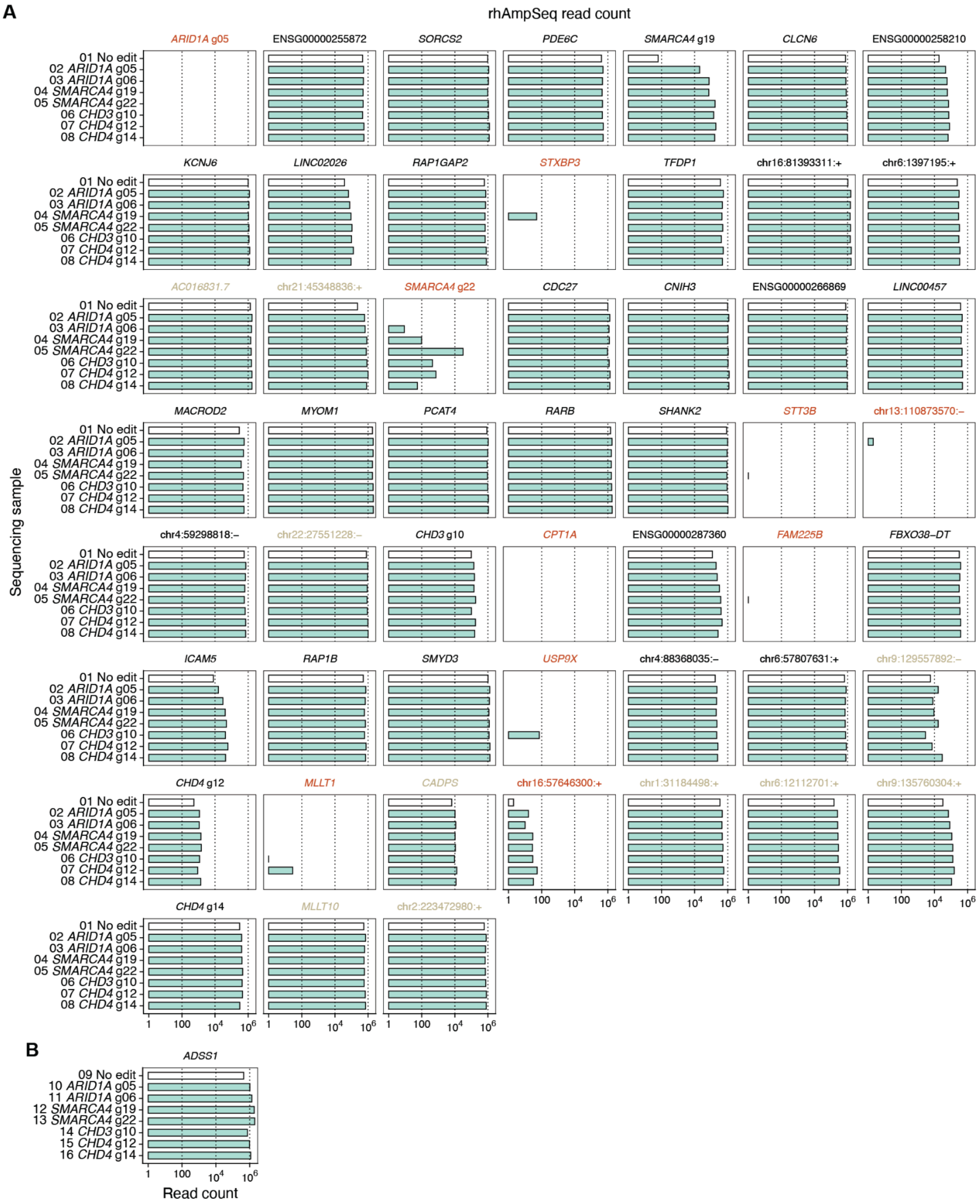


**Supplementary Figure 16. Sequencing coverage of rhAmpSeq amplicons.** Read count per amplicon and per sample (y axis). Red labels indicate sequencing failure (amplicons with < 50 reads in any sample). Gray labels indicate in silico predicted off-target sites.

**A**

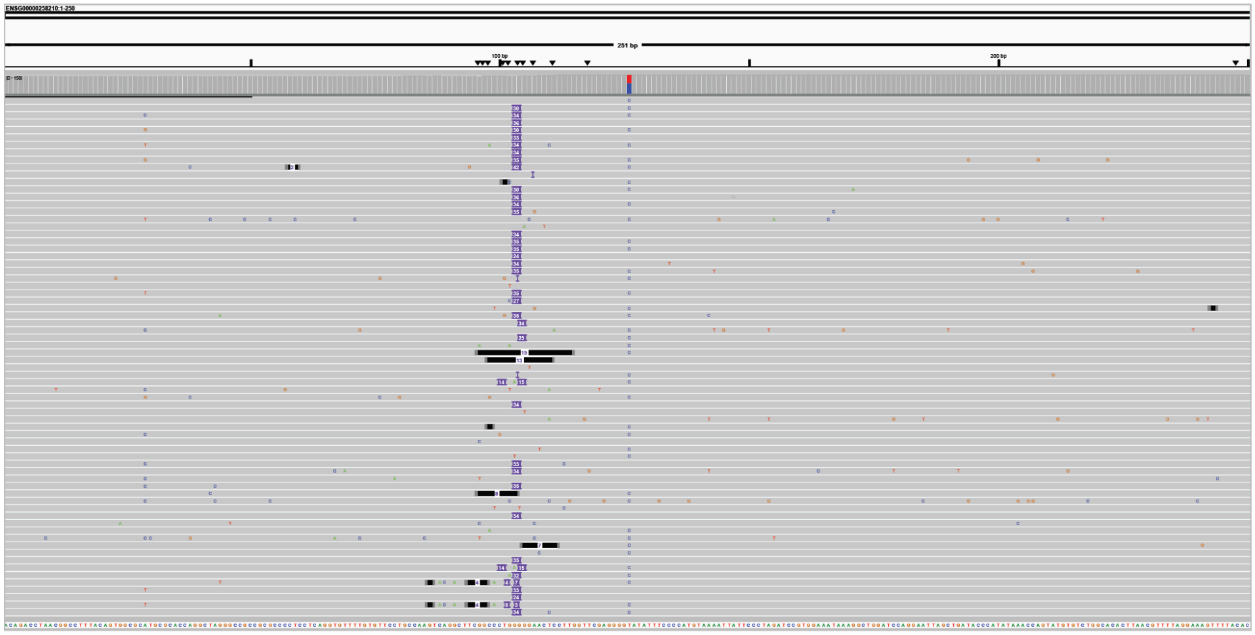


**B**

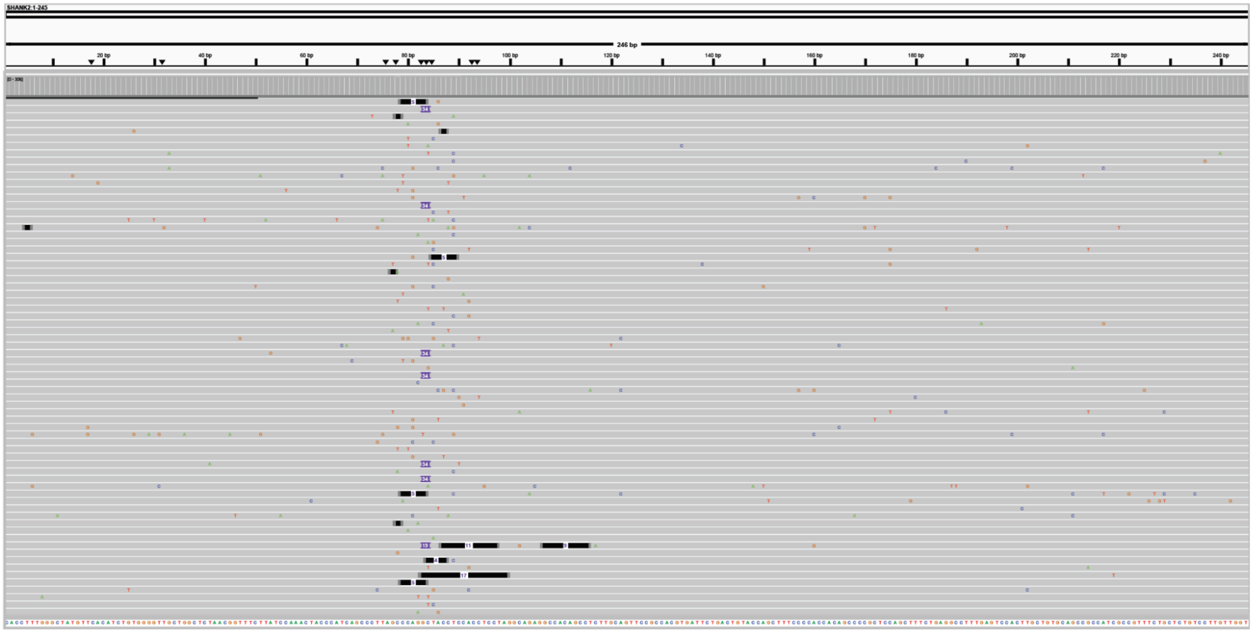


**Supplementary Figure 17. Validation of off-target T7 promoter insertion by rhAmpSeq. (A,B)** RhAmpSeq read alignments visualized by IGV at Superb-seq off-target sites (**A**) ENSG00000258210 and (**B**) *SHANK2*. Purple segments with white numbers indicate position and size of insertion events. The expected size of T7 promoter insertions is 34 bp.

ENSG00000287360

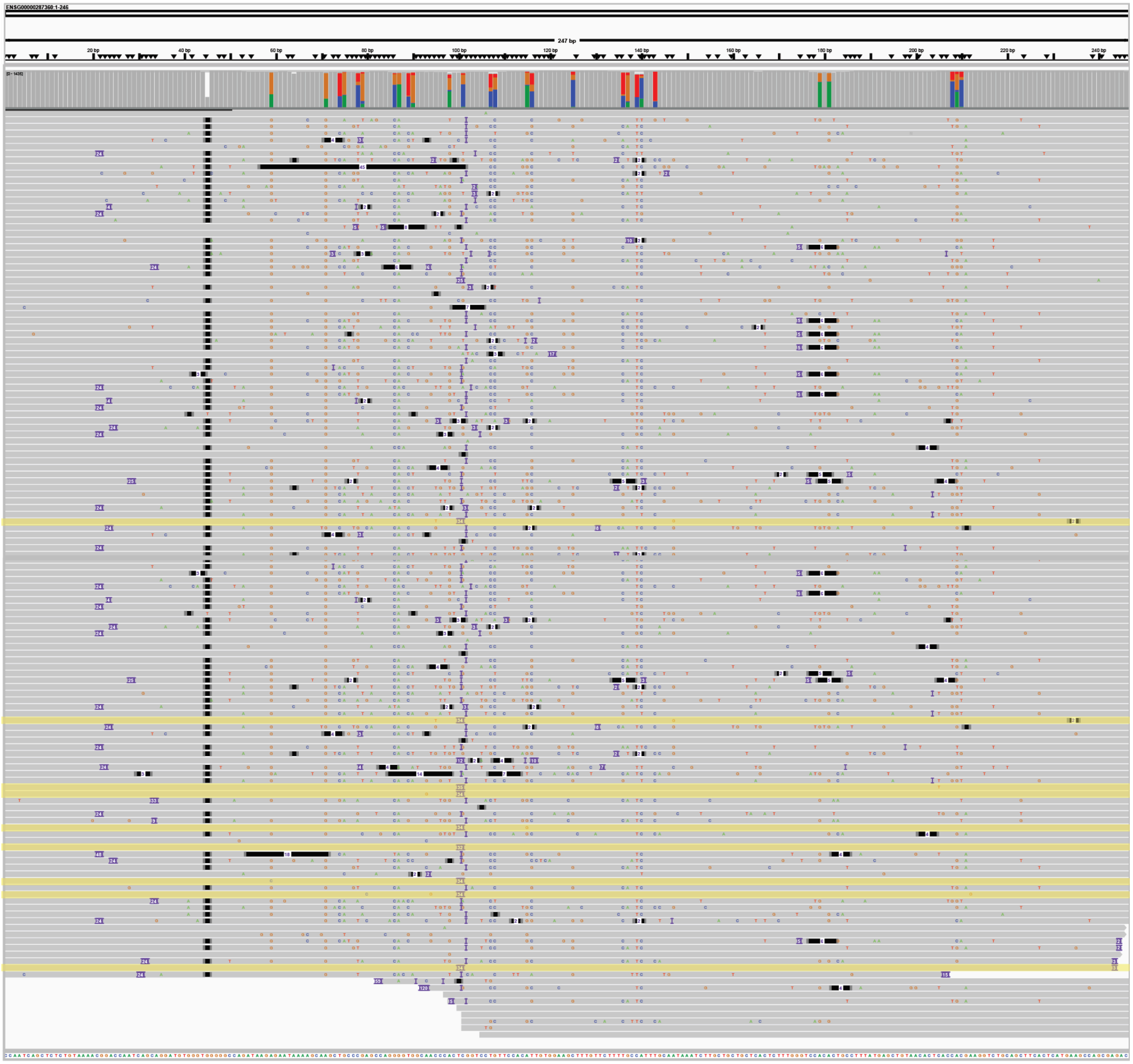

**Supplementary Figure 18. Read mismapping dampens rhAmpSeq detection of T7 promoter insertion.** RhAmpSeq read alignments visualized by IGV at Superb-seq off-target site ENSG0000287360. Highlights indicate reads containing T7 promoter (34 bp insertion) and strong reference genome alignment. Other sequences exhibit conserved mismatches to the expected genome sequence, indicating misprimed and mismapped background reads.

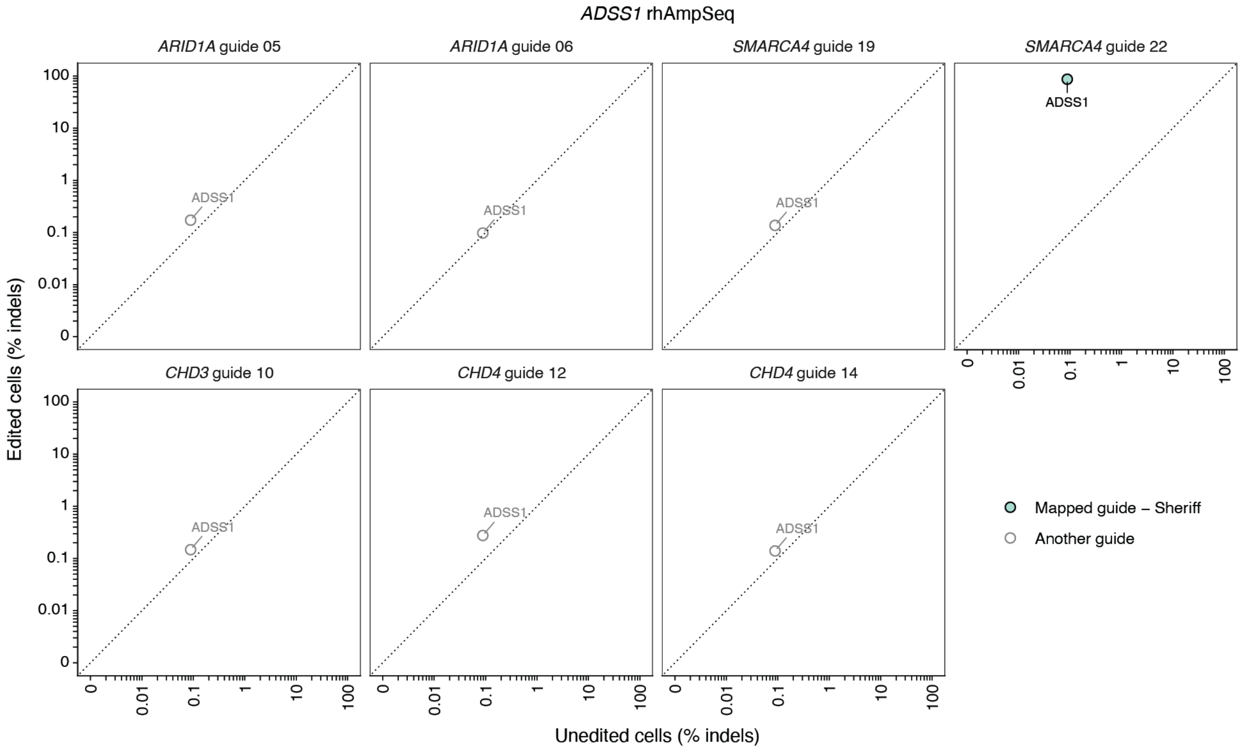


**Supplementary Figure 19. Validation of Sheriff guide assignments by rhAmpSeq.** Comparison of indel frequencies between edited and unedited K562 cell samples, determined by CRISPAltRations[22]. Each panel represents a different sample treated with the indicated guide and the singleplexed rhAmpSeq primers targeting *ADSS1*. Each point represents a different amplicon with sufficient sequencing coverage (*n* = 43, **Supp. Fig. 16**). Color indicates Sheriff guide association (green) or association to another guide (white).

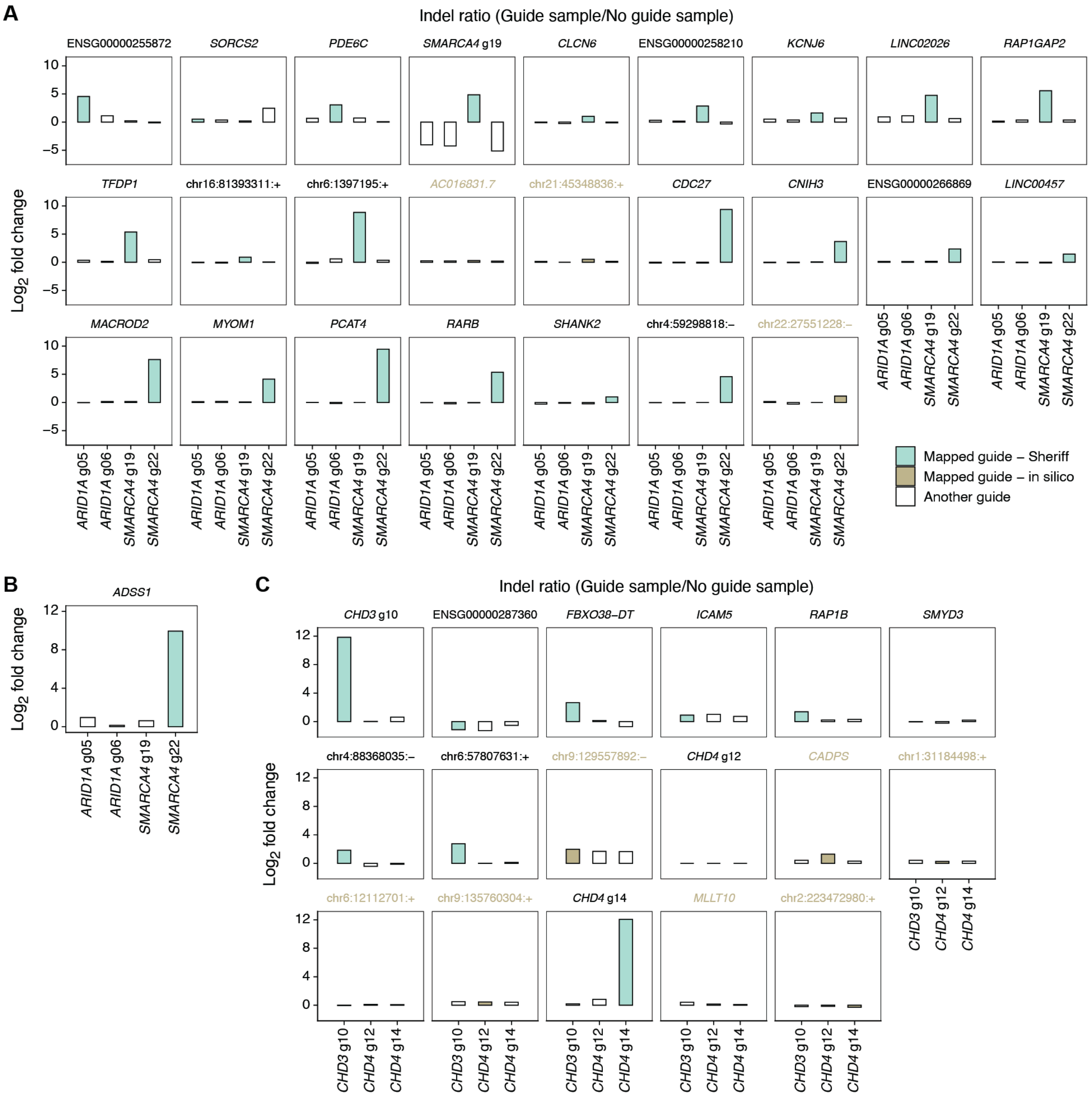


**Supplementary Figure 20. Validation of Sheriff guide assignments by rhAmpSeq indel ratios.** (**A–C**) Ratio of indel frequencies ($\log_2$ fold change) between edited and unedited K562 cell samples. (**A,B**) Samples treated with *ARID1A* or *SMARCA4* guides and (**A**) the panel of 52 multiplexed rhAmpSeq primers or (**B**) the singleplexed *ADSS1* primers. (**C**) Samples treated with *CHD3* or *CHD4* guides and multiplexed primers. Each panel represents an amplicon with sufficient sequencing coverage (**Supp. Fig. 16**). Color indicates Sheriff guide association (green), in silico guide association (khaki), or association to another guide (white).

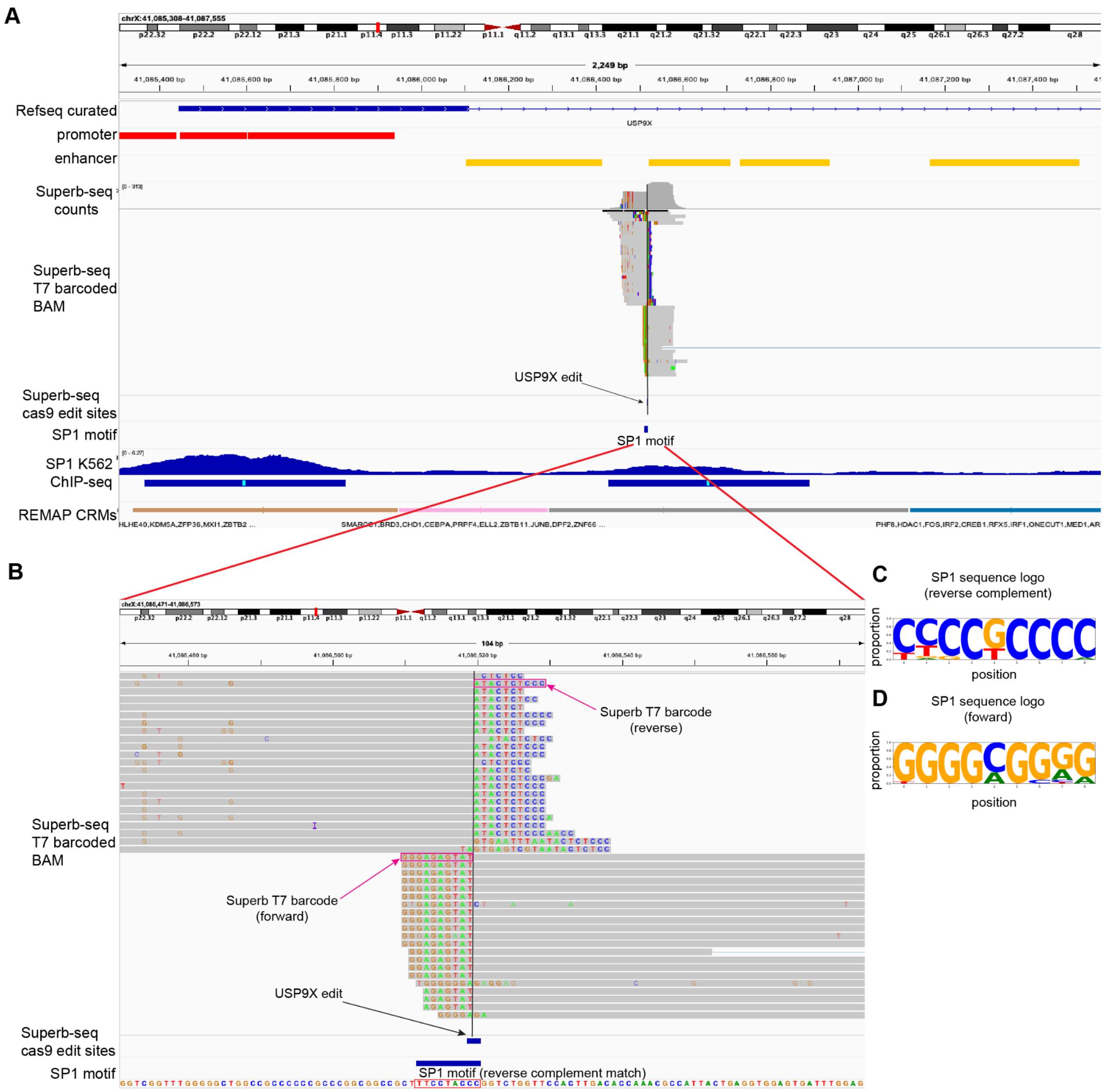


**Supplementary Figure 21. Off-target *USP9X* edit site intersects an SP1 binding motif.** (**A**) Integrative Genome Viewer (IGV) track view showing (from top to bottom) Refseq gene annotations, ENCODE promoter annotations, ENCODE candidate enhancer annotations, Superb-seq T7 barcoded read count pile-ups, Superb-seq T7 reads BAM (soft-clipped mismatched base pairs due to Superb barcode are highlighted either side of the *USP9X* Cas9 edit site), Superb-seq canonical edit site locations, annotation of SP1 motif that intersects the *USP9X* edit site, SP1 K562 ChIP-seq with fold-change over control and IDR conserved peaks annotated, and REMAP[23] cis regulatory module (CRM) annotations. (**B**) Zoomed in view of A, except only showing the Superb-seq T7 barcoded read BAM with the Superb barcoded highlighted, the annotated canonical edit site for *USP9X*, and the identified SP1 motif with the reference sequence highlighted (corresponding to the reverse complement of the actual SP1 motif). (**C**) Sequence logo of reverse complement of SP1 motif. (**D**) Sequence logo for the forward strand version of the SP1 motif.

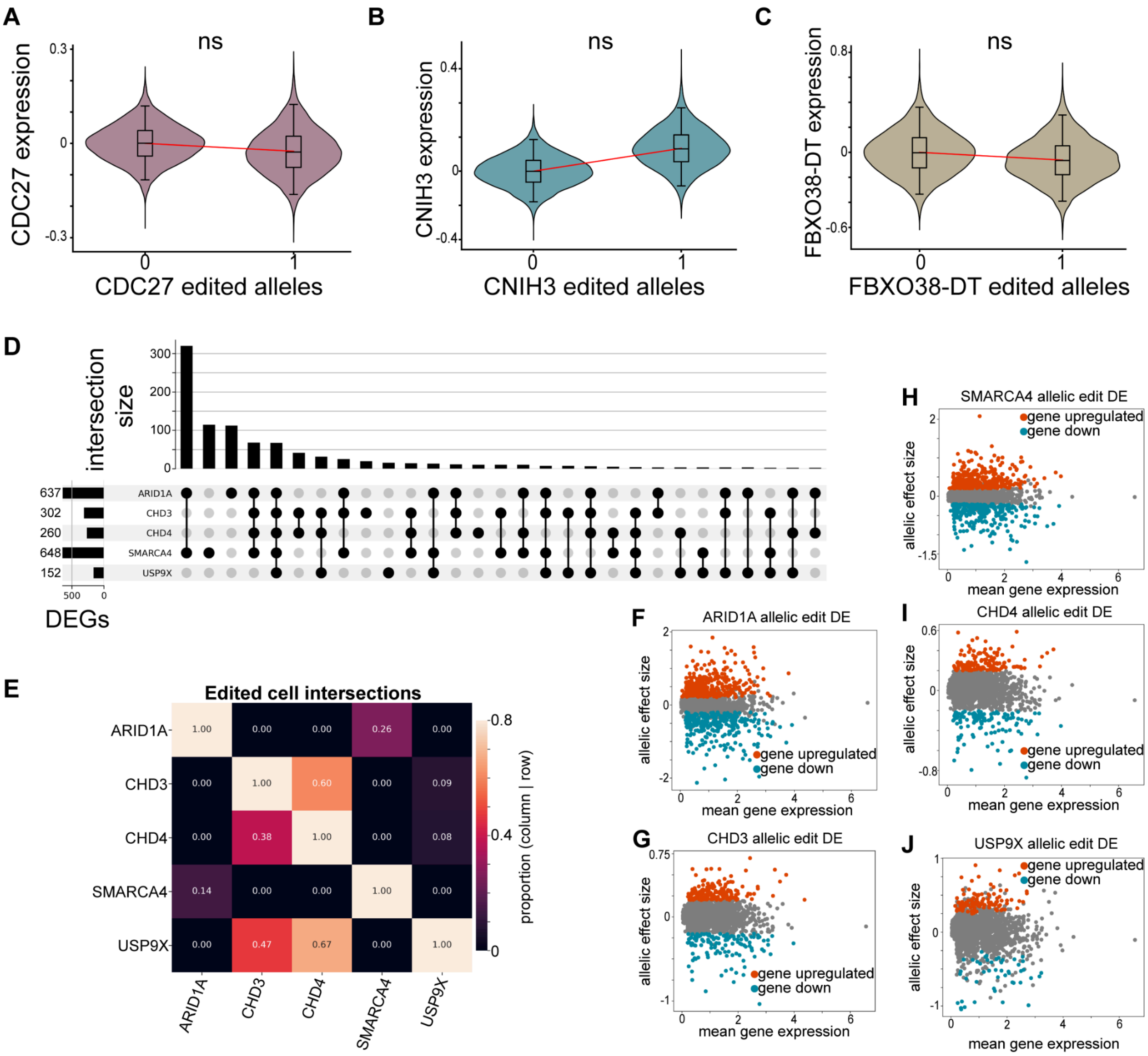


**Supplementary Figure 22. Differential gene expression associates with edit allele dosage.** (**A–C**) Violin plots of corrected gene expression estimates from *n* = 10,000 bootstrap iterations, grouped by edit allele dosage for off-target edits within the gene body of the tested gene. Overlaid box-plots show the interquartile range with the whiskers indicating the full data range (minimum and maximum). Lines indicate the effect on expression of edit allele dosage estimated by the linear mixed model. P-values are from performing a two-sided Wald test using the linear mixed model. (**D**) Upset plot indicating the intersection sizes of DEGs associated with the on- and off-target edit. (**E**) Heatmap indicating the proportion of cells within combinations of selected Cas9 edits. Proportions are conditioned on the edited gene listed on each row. For example, row one of the heatmap indicates 36% of cells with *ARID1A* edits also have *SMARCA4* edits. (**F–J**) Scatter plots with allelic effect size on the y-axis and mean gene expression on the x-axis, indicating differentially expressed genes are detected across a range of gene expression levels without bias based on mean expression level.

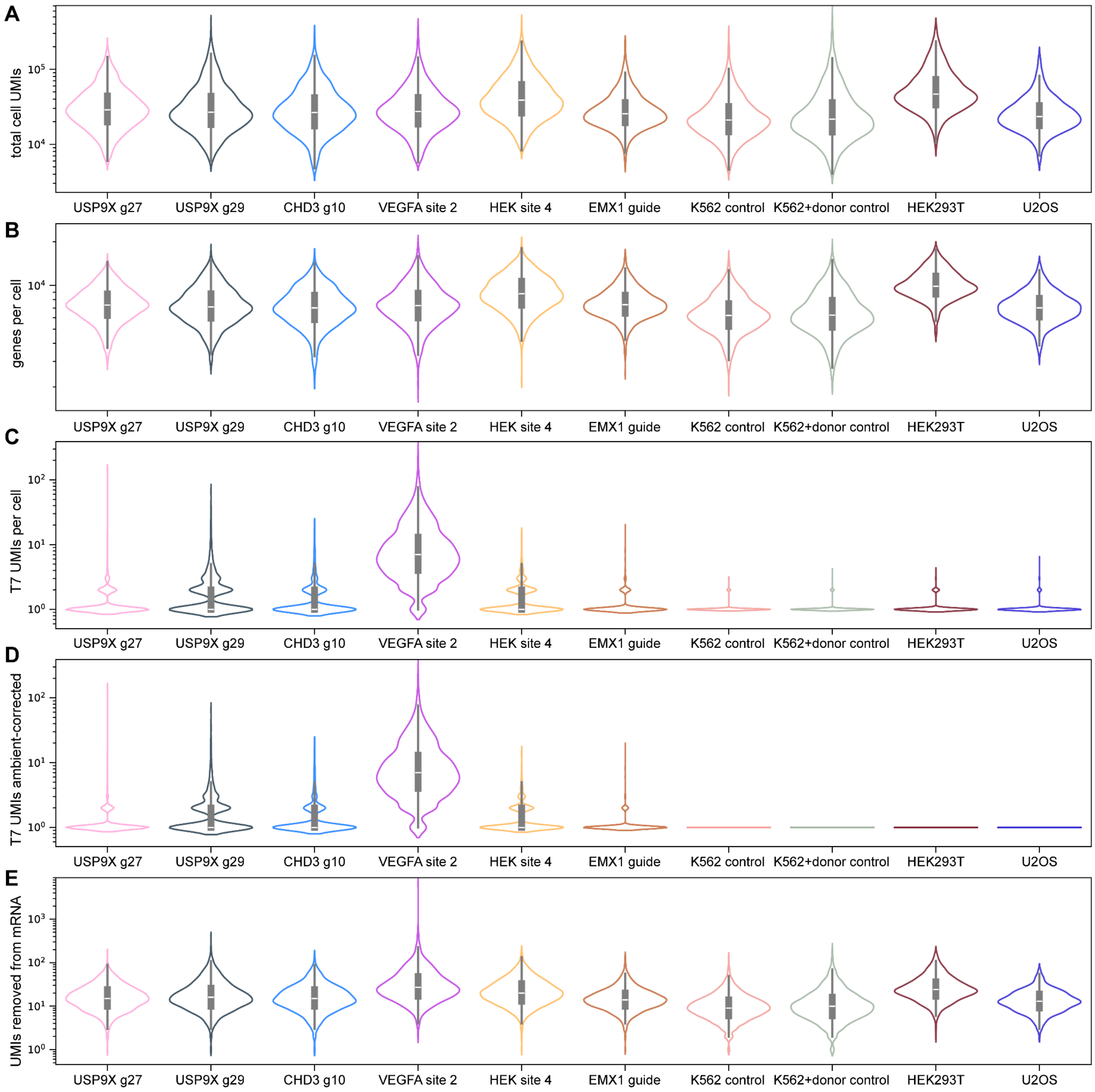


**Supplementary Figure 23. Quality control metrics in the Superb-seq benchmarking and validation library.** (**A**) Violin plots of single-cell total UMIs captured, with cells stratified by sample: *USP9X* g27 edited condition, *USP9X* g29 edited condition, *CHD3* g10 condition, *VEGFA* site 2 guide condition, HEK site 4 guide condition, *EMX1* guide condition, K562 no edit control condition, K562 no edit+T7 promoter control condition, HEK293T control condition, and the U2OS control condition. Because the y-axis is plotted on the log10 scale, a +1 pseudocount was added to the metrics to prevent infinity values. (**B**) Equivalent to A, except for the total genes detected. (**C**) Equivalent to A except for the T7 barcoded UMIs detected across all edit sites per cell. (**D**) Equivalent to C, except after T7 ambient UMI removal. (**E**) Equivalent to A except for the total removed UMIs occurring ± 1000 bp from detected edit sites.

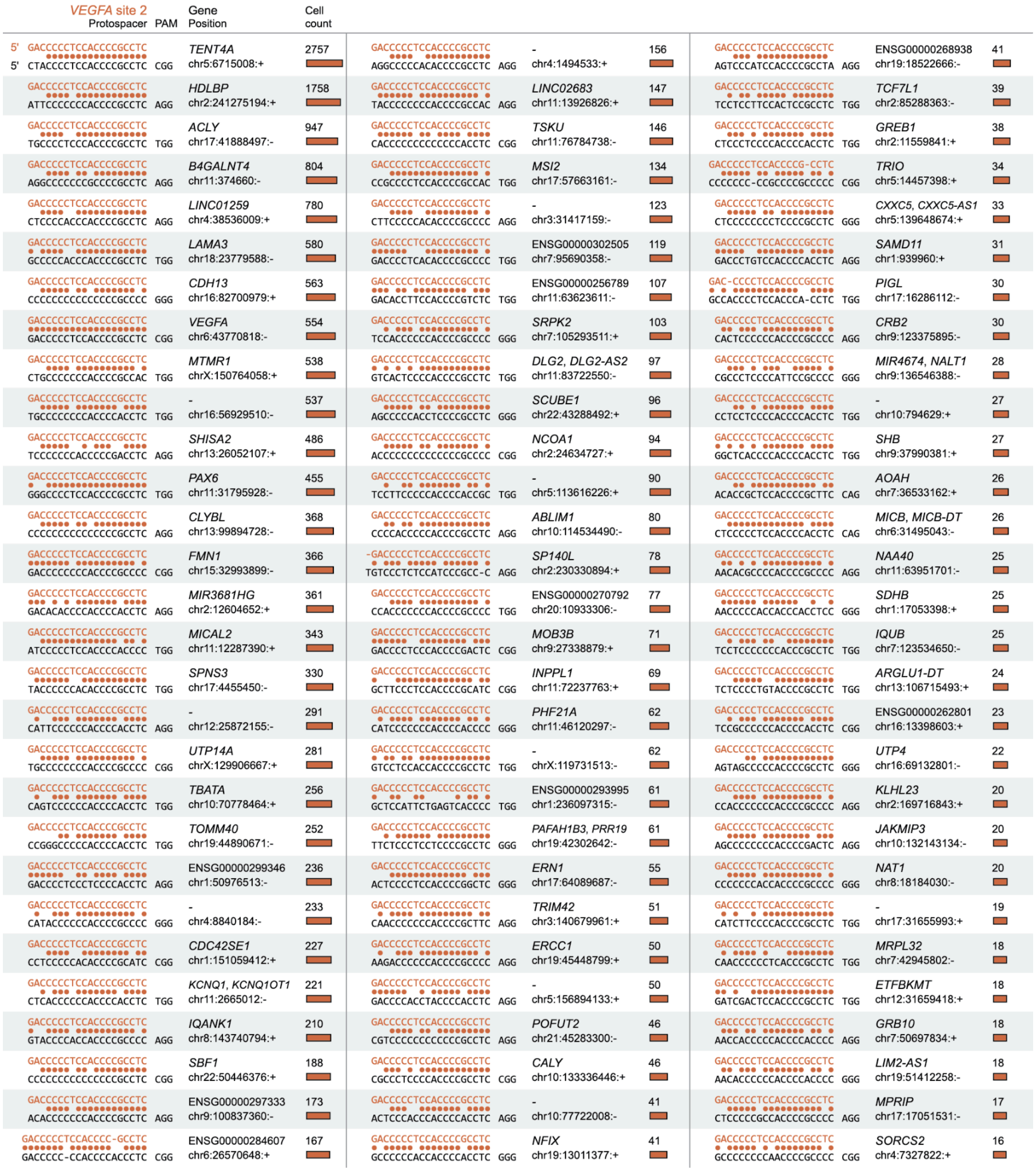


**Supplementary Figure 24. *VEGFA* site 2 guide alignments at Superb-seq edit sites.** Guide–reference genome alignments of the *VEGFA* site 2 guide at Superb-seq detected edit sites. Colored dots represent matched bases. The protospacer adjacent motif (PAM) and the number of edited cells is shown for each edit site. The most frequent edit sites out of a total of 210 sites are shown.

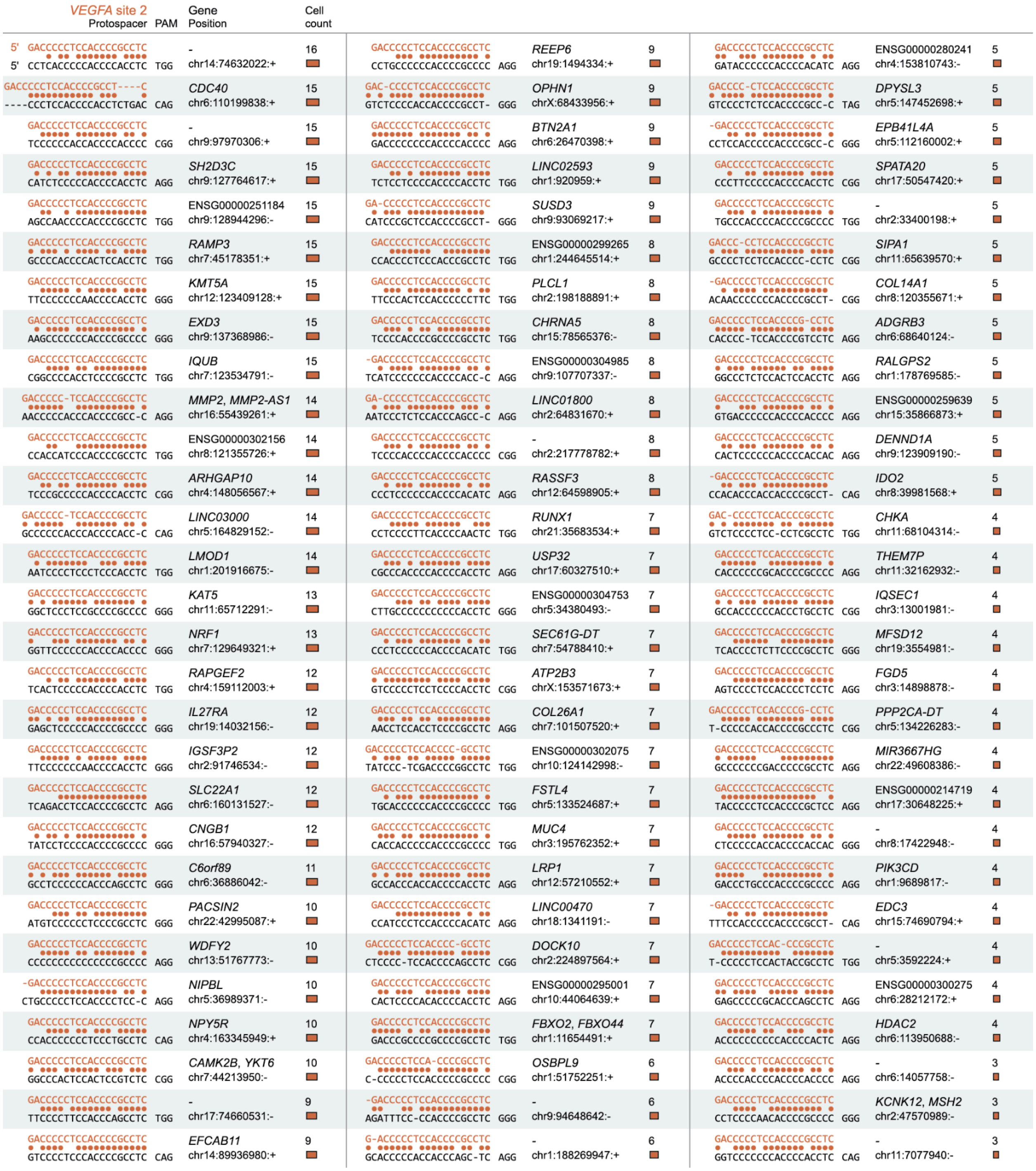


**Supplementary Figure 25. *VEGFA* site 2 guide alignments at Superb-seq edit sites, continued.** The next most frequent sequence alignments of the *VEGFA* site 2 guide at 210 Superb-seq detected edit sites.

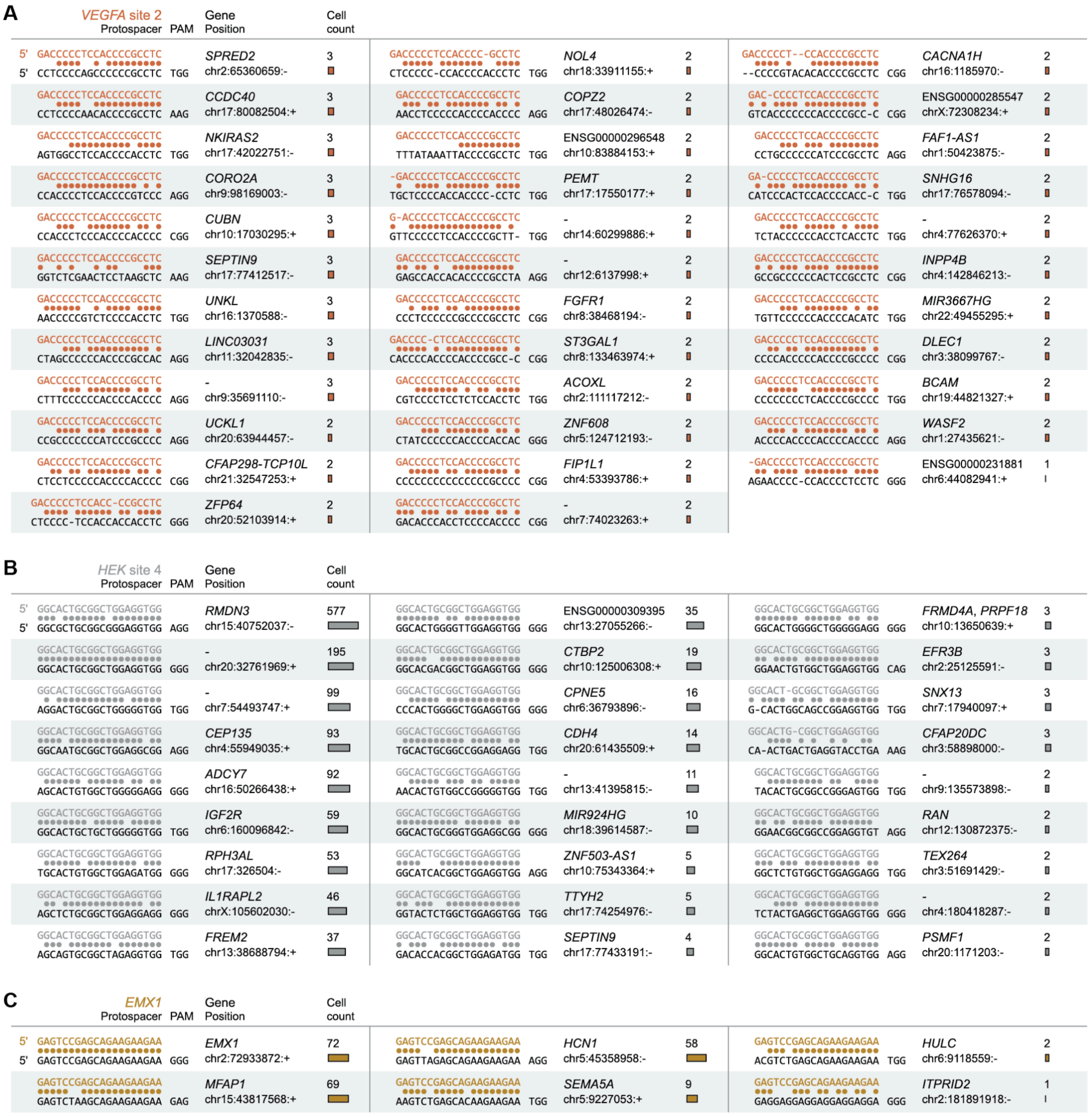


**Supplementary Figure 26. Benchmark guide alignments at Superb-seq edit sites.** (**A-C**) Guide–reference genome sequence alignments of (**A**) remainder of 210 edit sites of the *VEGFA* site 2 guide, (**B**) 27 edit sites of the HEK site 4 guide, and (**C**) six edit sites of the *EMX1* guide. The protospacer adjacent motif (PAM) and the number of edited cells is shown for each edit site.

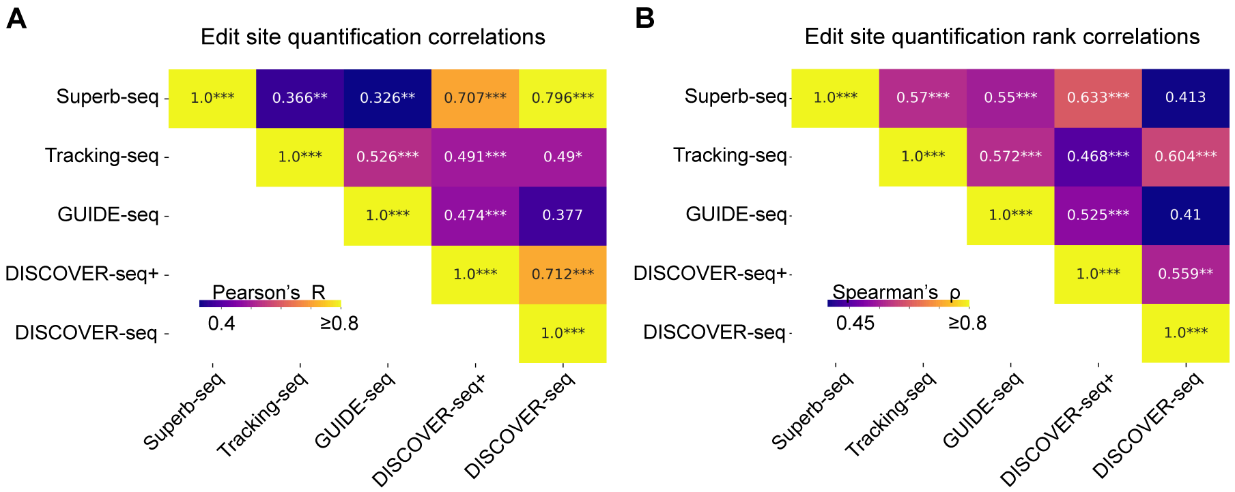


**Supplementary Figure 27. Superb-seq edited cell counts are correlated with bulk methods.** (**A**) Heatmap showing Pearson correlations for edit site quantification across various Cas9 editing detection methods. For each comparison, edit sites detected by both methods are utilized. (**B**) Equivalent to A, except using Spearman's correlation. ***Bonferroni p-value<0.001, **Bonferroni p-value<0.01, *Bonferroni p-value<0.05.

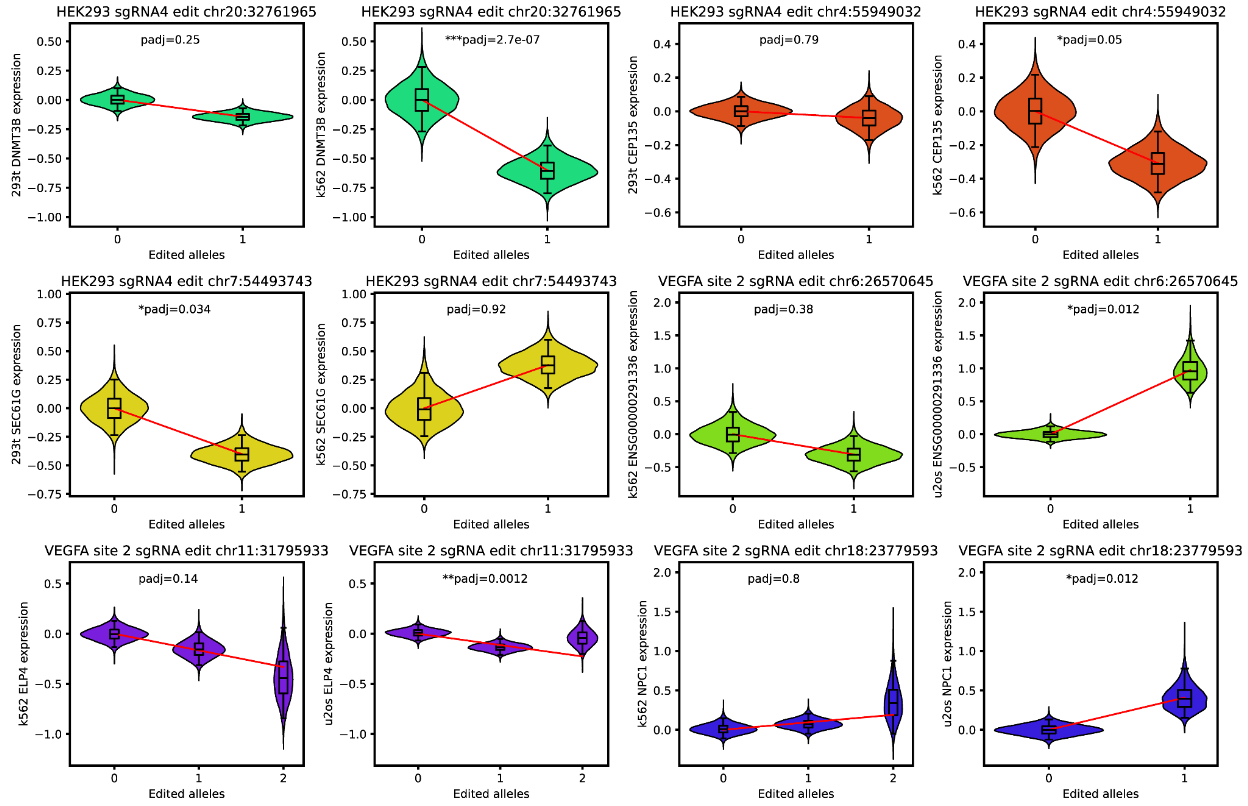


**Supplementary Figure 28. Nearest gene differential expression results by cell type.** Each violin plot shows the co-variate corrected expression of the tested gene in the particular cell type on the y-axis, and the number of edited alleles for the tested edit site on the x-axis. The violin-plot title indicates the causal guide of the edit and the edit location. Benjamini-Hochberg adjusted p-values indicate the significance of the association from our linear mixed model. Edits that were tested across cell types are shown adjacent to one another with the cell type indicated on the y-axis. The nearest expressed gene across any cell types electroporated with the guide was tested for differential expression with respect to each edit site.

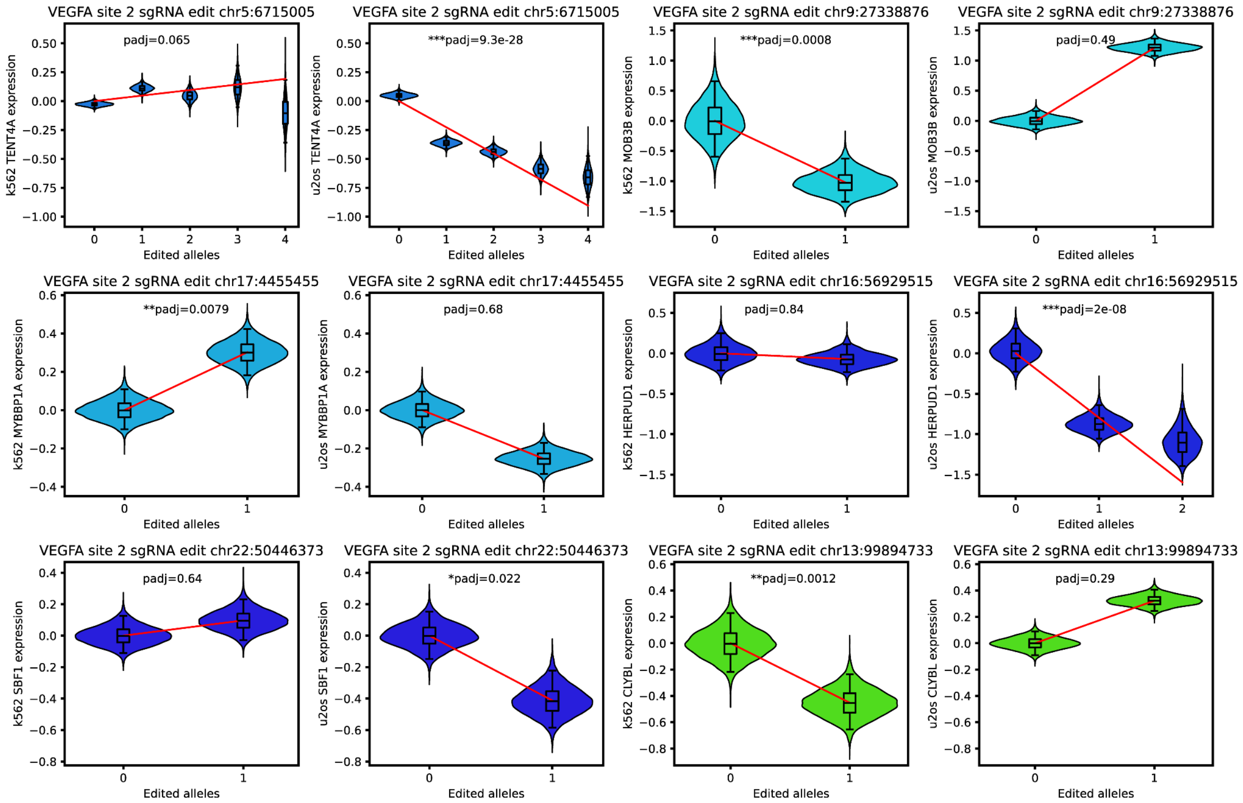


**Supplementary Figure 29. Nearest gene differential expression results by cell type, continued 1.** Each violin plot shows the co-variate corrected expression of the tested gene in the particular cell type on the y-axis, and the number of edited alleles for the tested edit site on the x-axis. The violin-plot title indicates the causal guide of the edit and the edit location. Benjamini-Hochberg adjusted p-values indicate the significance of the association from our linear mixed model. Edits that were tested across cell types are shown adjacent to one another with the cell type indicated on the y-axis. The nearest expressed gene across any cell types electroporated with the guide was tested for differential expression with respect to each edit site.

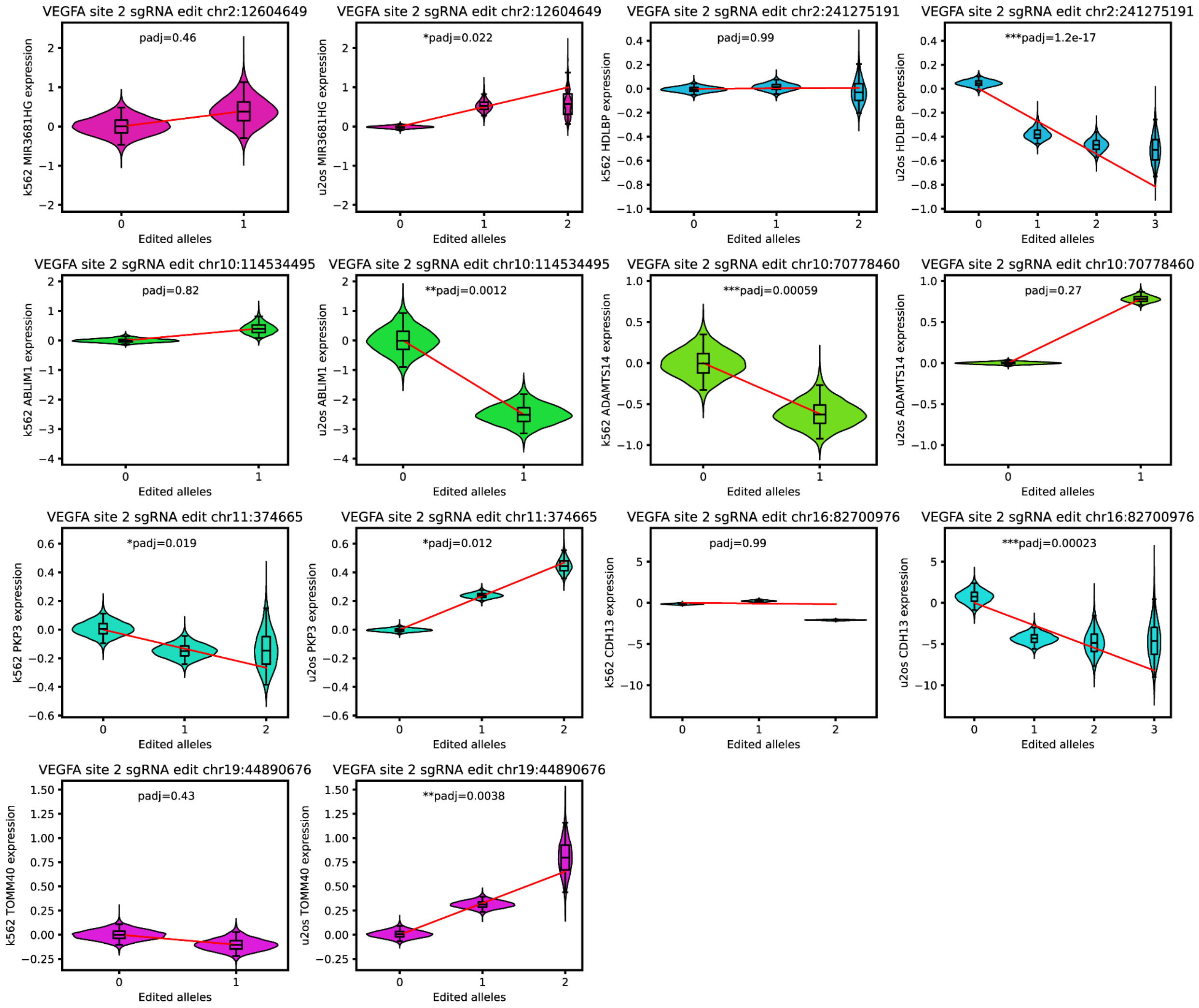


**Supplementary Figure 30. Nearest gene differential expression results by cell type, continued 2.** Each violin plot shows the co-variate corrected expression of the tested gene in the particular cell type on the y-axis, and the number of edited alleles for the tested edit site on the x-axis. The violin-plot title indicates the causal guide of the edit and the edit location. Benjamini-Hochberg adjusted p-values indicate the significance of the association from our linear mixed model. Edits that were tested across cell types are shown adjacent to one another with the cell type indicated on the y-axis. The nearest expressed gene across any cell types electroporated with the guide was tested for differential expression with respect to each edit site.

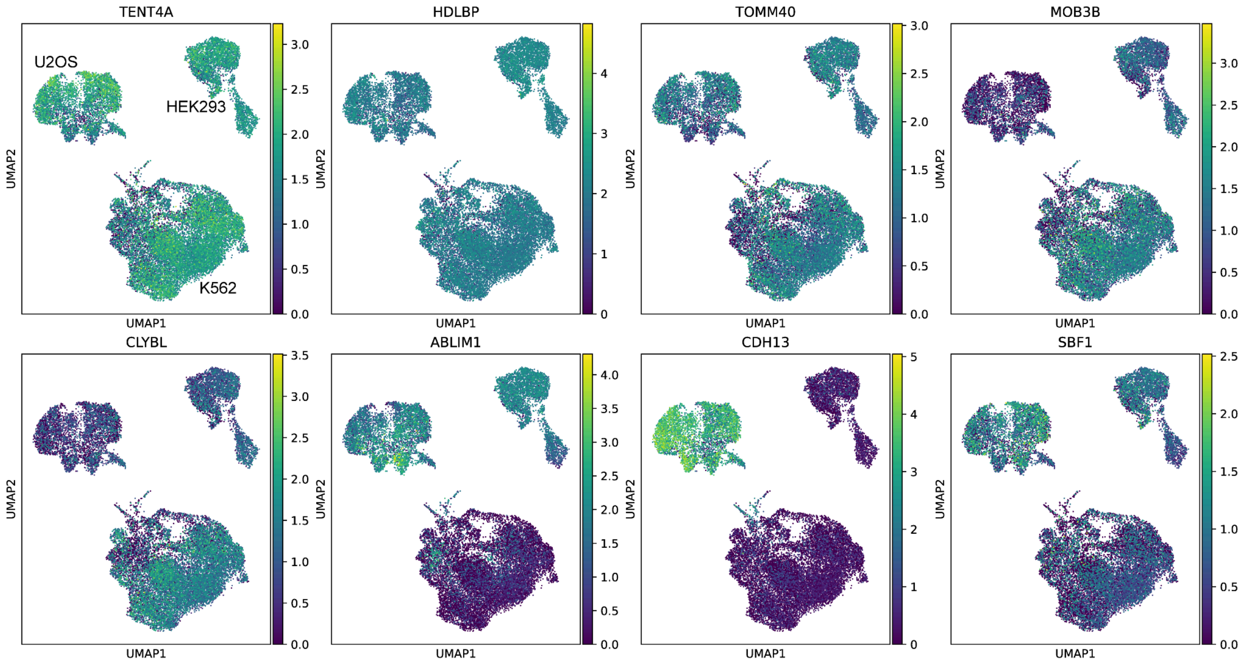


**Supplementary Figure 31. Cell type-specific expression of nearest genes.** Uniform manifold approximation (UMAP) plots of Superb-seq single cell transcriptomes for U2OS, HEK293, and K562 cells. Cells are colored by the log-transformed library-size normalized expression values of different genes. The genes shown are those that had differential expression in response to Cas9 editing at particular sites.

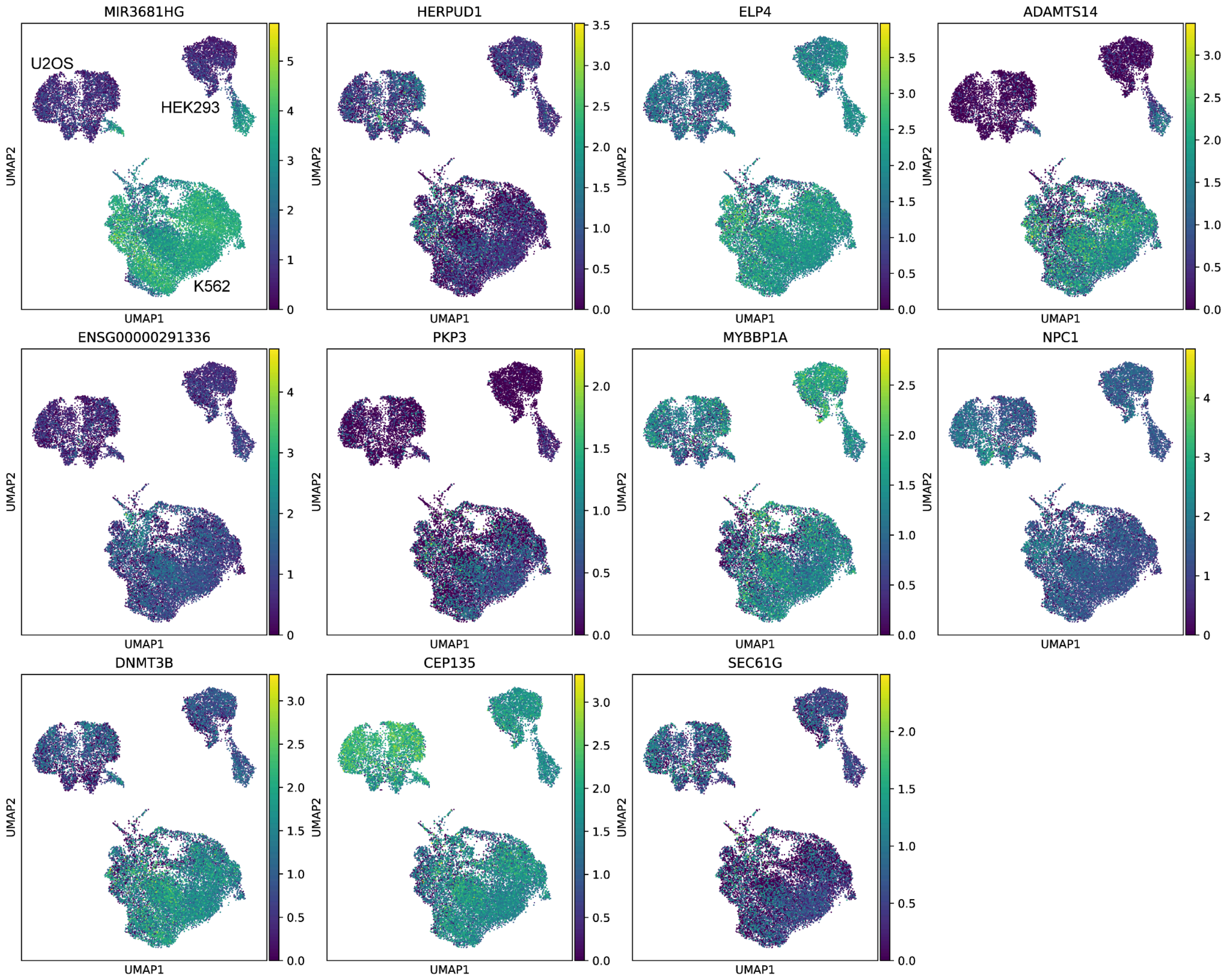


**Supplementary Figure 32. Cell type-specific expression of nearest genes, continued.** Uniform manifold approximation (UMAP) plots of Superb-seq single cell transcriptomes for U2OS, HEK293, and K562 cells. Cells are colored by the log-transformed library-size normalized expression values of different genes. The genes shown are those that were tested for differential expression in response to Cas9 editing at particular sites.

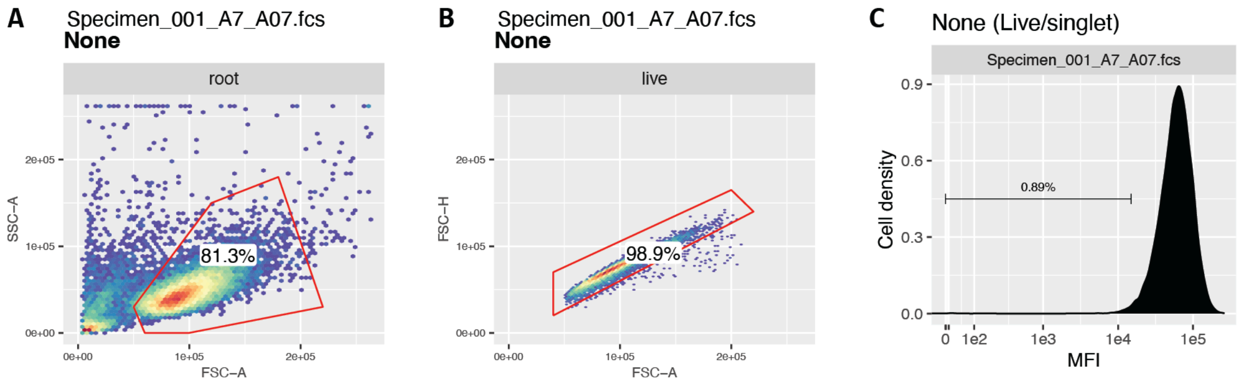


**Supplementary Figure 33. Flow cytometry gating.** (**A–C**) Flow cytometry gating of unedited GM12878 cells. Example gating of (**A**) live, (**B**) singlet, and (**C**) B2M-knockdown gating, used in **Ext. Data Fig. 1F**.

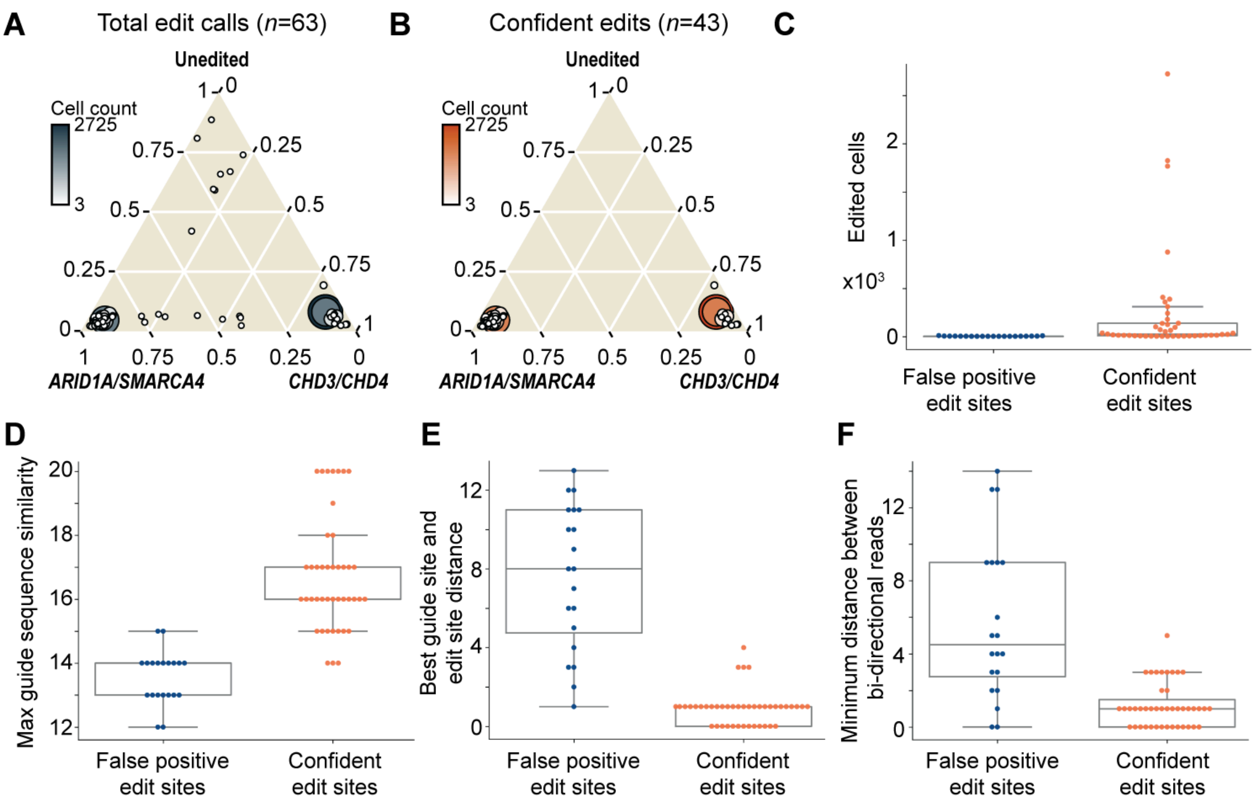


**Supplementary Figure 34. Multiple metrics confirm confident Superb-seq edit detection.** (**A**) Ternary plot, with each dot indicating a called canonical edit site (total *n* = 63). Dots are colored and scaled by the number of cells called with T7 barcoded reads for the edit site. Edit sites are plotted according to the proportion of cells called with the edit in the different treatment samples (unedited, *ARID1A/SMARCA4* guide treated cells, *CHD3* and *CHD4* guide treated cells). True edit sites are expected to be specific to a particular treatment sample. (**B**) Equivalent to A, except subsetted to confident edit sites, called based on manual examination of the T7 barcoded read alignment at each edit site. (**C**) Swarm- and box- plots overlaid, with each point indicating a called edit site, with edit sites stratified into confident and false positive edit sites on the x-axis. Overlaid box-plots show the interquartile range with the whiskers indicating the full data range (minimum and maximum). The y-axis indicates the number of cells within which the respective edit was detected prior to T7 ambient read correction. (**D**) Equivalent to C, except with the y-axis indicating the maximum sequence similarity across the 7 guide sequences and the reference genome at the called edit sites. (**E**) Equivalent to C, except the y-axis indicates the distance between the best candidate Cas9 cleavage site based on sequence similarity to one of the 7 guide sequences, and the called canonical edit site position. (**F**) Equivalent to C, except the y-axis indicates the minimum base pair distance between the closest pair of forward and reverse mapping reads, with true edit sites expected to have bi-directional reads immediately either side of the Cas9 cleavage site for true edit events.

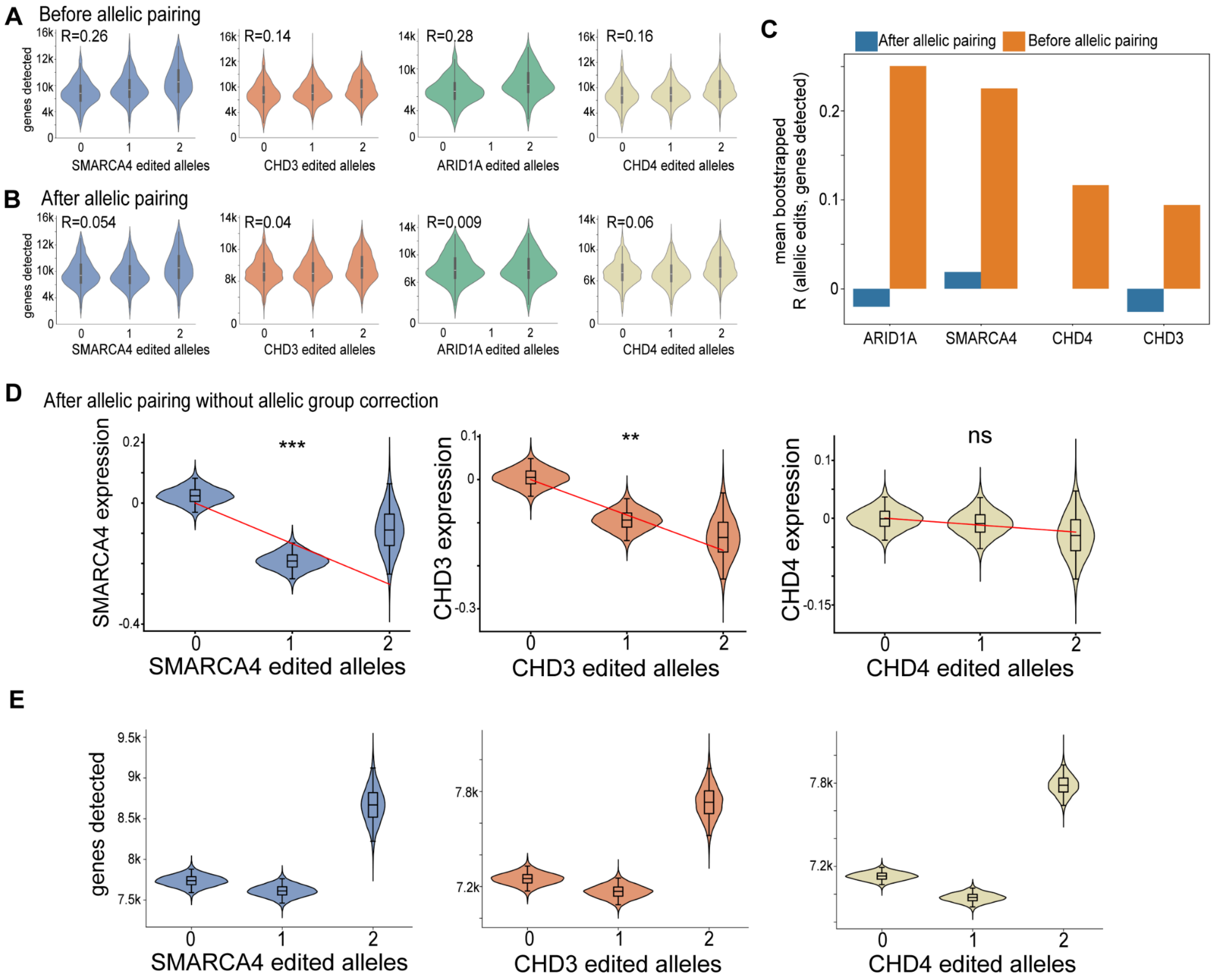


**Supplementary Figure 35. Correction of gene expression level estimation by allelic pairing.** (**A**) Correlation of the number of detected genes per cell with the number of detected edit alleles per cell at the indicated on-target gene (*SMARCA4*, *CHD3*, *ARID1A*, *CHD4*). 'R' indicates the Pearson correlation. Overlaid box-plots show the interquartile range (IQR) with the whiskers indicating the full data range (minimum and maximum), excluding outliers that fall outside 1.5x the IQR above or below the 25th and 75th percentiles. (**B**) Equivalent to A after performing allelic pairing. (**C**) Pearson correlations between the gene detection rate and the edit alleles detected. Correlations are stratified by the edited gene and before or after performing the allelic pairing. (**D**) Violin plots with boot-strapped mean estimates (10,000 boot-straps) of expression for genes fitted with a linear mixed model that does not consider allelic groups (groups of unedited and 1 allele cells, unedited and 2 allele cells) when performing the linear mixed modeling. **Fig. 4B** has the version of this panel with the allelic group correction. Overlaid box-plots show the interquartile range with the whiskers indicating the full data range (minimum and maximum). (**E**) Equivalent to D, except showing the genes detected in each group of cells.

### Supplementary references


1. Shou, J., Li, J., Liu, Y. & Wu, Q. Precise and predictable CRISPR chromosomal rearrangements reveal principles of Cas9-mediated nucleotide insertion. *Mol. Cell* **71**, 498–509.e4 (2018).
2. Wang, T., Wei, J. J., Sabatini, D. M. & Lander, E. S. Genetic screens in human cells using the CRISPR-Cas9 system. *Science* **343**, 80–84 (2014).
3. Xu, H. *et al.* Sequence determinants of improved CRISPR sgRNA design. *Genome Res.* **25**, 1147–1157 (2015).
4. Dunn, J. J. & Studier, F. W. Complete nucleotide sequence of bacteriophage T7 DNA and the locations of T7 genetic elements. *J. Mol. Biol.* **166**, 477–535 (1983).
5. Ye, J., McGinnis, S. & Madden, T. L. BLAST: improvements for better sequence analysis. *Nucleic Acids Res.* **34**, W6–9 (2006).
6. Rosa, M. D. Four T7 RNA polymerase promoters contain an identical 23 bp sequence. *Cell* **16**, 815–825 (1979).
7. Corces, M. R. *et al.* An improved ATAC-seq protocol reduces background and enables interrogation of frozen tissues. *Nat. Methods* **14**, 959–962 (2017).
8. Untergasser, A., Ruijter, J. M., Benes, V. & van den Hoff, M. J. B. Web-based LinRegPCR: application for the visualization and analysis of (RT)-qPCR amplification and melting data. *BMC Bioinformatics* **22**, 398 (2021).
9. Rosenberg, A. B. *et al.* Single-cell profiling of the developing mouse brain and spinal cord with split-pool barcoding. *Science* **360**, 176–182 (2018).
10. Mali, P. *et al.* RNA-guided human genome engineering via Cas9. *Science* **339**, 823–826 (2013).
11. Luo, Y. *et al.* New developments on the Encyclopedia of DNA Elements (ENCODE) data portal. *Nucleic Acids Res.* **48**, D882–D889 (2020).
12. Hargreaves, D. C. & Crabtree, G. R. ATP-dependent chromatin remodeling: genetics, genomics and mechanisms. *Cell Res.* **21**, 396–420 (2011).
13. Tran, V. *et al.* High sensitivity single cell RNA sequencing with split pool barcoding. *bioRxiv* 2022.08.27.505512 (2022) doi:10.1101/2022.08.27.505512.
14. Fleming, S. J. *et al.* Unsupervised removal of systematic background noise from droplet-based single-cell experiments using CellBender. *Nat. Methods* **20**, 1323–1335 (2023).
15. Replogle, J. M. *et al.* Mapping information-rich genotype-phenotype landscapes with genome-scale Perturb-seq. *Cell* **185**, 2559–2575.e28 (2022).
16. Malinin, N. L. *et al.* Defining genome-wide CRISPR–Cas genome-editing nuclease activity with GUIDE-seq. *Nat. Protoc.* **16**, 5592–5615 (2021).
17. Conrad, T., Plumbom, I., Alcobendas, M., Vidal, R. & Sauer, S. Maximizing transcription of nucleic acids with efficient T7 promoters. *Commun Biol* **3**, 439 (2020).
18. Wagih, O. ggseqlogo: a versatile R package for drawing sequence logos. *Bioinformatics* **33**, 3645–3647 (2017).
19. Schmittgen, T. D. & Livak, K. J. Analyzing real-time PCR data by the comparative C(T) method. *Nat. Protoc.* **3**, 1101–1108 (2008).
20. Aznauryan, E. *et al.* Discovery and validation of human genomic safe harbor sites for gene and cell therapies. *Cell Rep Methods* **2**, 100154 (2022).
21. Booeshaghi, A. S., Chen, X. & Pachter, L. A machine-readable specification for genomics assays. *Bioinformatics* **40**, (2024).
22. Kurgan, G. *et al.* CRISPAltRations: a validated cloud-based approach for interrogation of double-strand break repair mediated by CRISPR genome editing. *Mol. Ther. Methods Clin. Dev.* **21**, 478–491 (2021).
23. Hammal, F., de Langen, P., Bergon, A., Lopez, F. & Ballester, B. ReMap 2022: a database of Human, Mouse, Drosophila and Arabidopsis regulatory regions from an integrative analysis of DNA-binding sequencing experiments. *Nucleic Acids Res.* **50**, D316–D325 (2022).
