## Additional Files for "Joint single-cell capture of Cas9 edits and transcriptomes reveals on- and off-target effects on gene expression": README.md

Sheriff 
==================
## Identification of CRISPR/cas9 edit sites in single cells
<img src="https://github.com/BradBalderson/Sheriff/blob/main/img/sheriff2.png" alt="Sheriff Badge" width="600">

**Sheriff processes aligned Superb-seq data to call edit sites and quantify gene expression in single cells** 

The inputted bam must be the annotated bam file outputted from split-pipe, which is available after creating an account at [Parse Biosciences](https://support.parsebiosciences.com/hc/en-us/articles/17200056667924-Pipeline-Download-Current-Version).

For split-pipe version and commands used to process the fastq files to the annotated bam file, please see the [superb_analysis repository](https://github.com/BradBalderson/superb_analysis/tree/main).

Install
-------
Please replace 'mamba' with 'conda' if not installed, mamba is much faster however (we recommend installing mamba).

Expected install time is approximately 1-minute. 

The current version has been tested with python 3.10 using the conda environment setup specified below, 
on both linux (Rocky 9 distro), macOS Sonoma 14.1, & macOS Ventura.

To install from source:

    mamba create -n sheriff_env python=3.10
    mamba activate sheriff_env
    mamba install conda-forge::scipy conda-forge::numpy==1.26.4 bioconda::gtfparse conda-forge::faiss-cpu conda-forge::numba conda-forge::biopython=1.81 typing_extensions typer bioconda::pyranges bioconda::pysam 

    git clone https://github.com/BradBalderson/Sheriff.git
    cd Sheriff
    pip install .

    # ensure zlib is found, can be an issue on some machines.
    export LDFLAGS="-L$CONDA_PREFIX/lib"
    export CPPFLAGS="-I$CONDA_PREFIX/include"
    export LD_LIBRARY_PATH="$CONDA_PREFIX/lib:$LD_LIBRARY_PATH"

Usage
-----
NOTE running ***sheriff --help*** the first time is slow. 

    sheriff --help

     Usage: sheriff [OPTIONS] BAM_FILE REF_FILE BARCODE_FILE GTF_FILE
    
    ╭─ Arguments ────────────────────────────────────────────────────────────────────────────────────────────────────────────────────────────────────────────────────────────────────────────────────────────────────────╮
    │ *    bam_file          TEXT  BAM file [default: None] [required]                                                                                                                                                   │
    │ *    ref_file          TEXT  Fasta containing ref genome [default: None] [required]                                                                                                                                │
    │ *    barcode_file      TEXT  Text file containing whitelisted barcode per line [default: None] [required]                                                                                                          │
    │ *    gtf_file          TEXT  GTF file containing relevant gene data [default: None] [required]                                                                                                                     │
    ╰────────────────────────────────────────────────────────────────────────────────────────────────────────────────────────────────────────────────────────────────────────────────────────────────────────────────────╯
    ╭─ Options ──────────────────────────────────────────────────────────────────────────────────────────────────────────────────────────────────────────────────────────────────────────────────────────────────────────╮
    │ --t7,--t7_barcode                                                                                  TEXT     Target/query barcode sequence to denote t7 reads. [default: GGGAGAGTAT]                                │
    │ --blacklist,--blacklist_file                                                                       TEXT     Bed file that species the location of blacklist regions, these generate alot of endogenuous t7 reads   │
    │                                                                                                             that can lead to slow processing time and false-positive edit-site calling.                            │
    │                                                                                                             [default: None]                                                                                        │
    │ --whitelist,--whitelist_file                                                                       TEXT     Bed file that species the location of whitelist regions, which are known edit sites and so will call   │
    │                                                                                                             any barcoded reads implying an edit site intersecting these regions as canonical edit sites.           │
    │                                                                                                             [default: None]                                                                                        │
    │ --kmer,--kmer_size                                    -k                                           INTEGER  Size of kmers used to pattern match read barcodes to the t7 barcode. [default: 6]                      │
    │ --edit_dist,--edist,--dist                                                                         INTEGER  +/- distance from edit site to be grouped as same edit. [default: 140]                                 │
    │ --bidirectional_inserts                                              --no-bidirectional_inserts             Candidate edit site must have evidence of bi-directional donor insertion to be called as a canonical   │
    │                                                                                                             edit site.Highly recommended criteria. If turned off, it also turns off --stranded_edit_dist criteria, │
    │                                                                                                             but this information about the edit sites are still recorded in the output via the                     │
    │                                                                                                             'stranded_edit_dist' columnin edit_site_info.txt                                                       │
    │                                                                                                             [default: bidirectional_inserts]                                                                       │
    │ --stranded_edit_dist                                                                               INTEGER  Maximum allowed distance between the nearest forward and reverse edit sites at a given canonical edit  │
    │                                                                                                             site to qualify as real edit.                                                                          │
    │                                                                                                             [default: 15]                                                                                          │
    │ --edit_site_min_cells                                                                              INTEGER  Minimum cells in edit site to be considered true edit. [default: 3]                                    │
    │ --nonbc_edit_dist,--nonbc_edist,--nonbc_dist,--nonbc                                               INTEGER  +/- distance from edit to mop up the non-barcoded reads. [default: 1000]                               │
    │ --ploidy                                                                                           INTEGER  Ploidy/Number of chromosomes in the genome. [default: 2]                                               │
    │ --cnv,--cnv_file,--copy_number_variant_file                                                        TEXT     A bedGraph file that specifies copy-number-variation sites, that deviate from the ploidy number.       │
    │                                                                                                             [default: None]                                                                                        │
    │ --blacklist_seqs                                                                                   TEXT     Text file of sequences, with a new sequence on each line, that may be present in read soft-clip        │
    │                                                                                                             sequencescan confound t7 barcoded read calls. Currently only the TSO, which is a common left-over      │
    │                                                                                                             artifact.                                                                                              │
    │                                                                                                             [default: None]                                                                                        │
    │ --mrna_count_mode                                                                                  TEXT     Mode for quantifying gene expression,'all' is to count all reads associated with a gene, 'polyT' is to │
    │                                                                                                             only count polyT reads, indicating mature mRNA transcripts.                                            │
    │                                                                                                             [default: all]                                                                                         │
    │ --uncorrected_count                                                  --no-uncorrected_count                 Whether to also output the gene counts WITHOUT removing reads around inferred edit sites.              │
    │                                                                                                             [default: no-uncorrected_count]                                                                        │
    │ --out,--outdir,--out_dir                              -o                                           TEXT     Write output files to this location. Defaults to Current Working Directory [default: None]             │
    │ --v,--verbosity                                       -v,-verbosity                                INTEGER  Verbosity levels. 0 errors only, 1 prints processing progress, 2 prints debugging information.         │
    │                                                                                                             [default: 1]                                                                                           │
    │ --cpu                                                 -cpu                                         INTEGER  Number of CPUs to use for processing, necessary to increase this for fast UMI counting. [default: 1]   │
    │ --chunk                                               -chunk                                       INTEGER  Number of mega-bases to process at a time for gene UMI counting. Set this lower if get memoryissues,   │
    │                                                                                                             currently set to > hg38 chr1 size (249 Mb), so is parallelized by chromosome.Trade-off is can cause    │
    │                                                                                                             minor double-counting of UMIs if a genome chunk intersects a gene.Very small / negligible difference   │
    │                                                                                                             in counts.                                                                                             │
    │                                                                                                             [default: 250]                                                                                         │
    │ --install-completion                                                                                        Install completion for the current shell.                                                              │
    │ --show-completion                                                                                           Show completion for the current shell, to copy it or customize the installation.                       │
    │ --help                                                                                                      Show this message and exit.                                                                            │
    ╰────────────────────────────────────────────────────────────────────────────────────────────────────────────────────────────────────────────────────────────────────────────────────────────────────────────────────╯

Example
------

Expected run-time for data preparation below is ~2 minutes.

Assumes are in the Sheriff directory.

#### Preparing reference genome
    wget http://ftp.ensembl.org/pub/release-110/fasta/homo_sapiens/dna/Homo_sapiens.GRCh38.dna.primary_assembly.fa.gz -O example_data/Homo_sapiens.GRCh38.dna.primary_assembly.fa.gz
    gzip -d example_data/Homo_sapiens.GRCh38.dna.primary_assembly.fa.gz

    mamba install bioconda::samtools
    samtools faidx example_data/Homo_sapiens.GRCh38.dna.primary_assembly.fa

#### Preparing reference annotations
    wget http://ftp.ensembl.org/pub/release-110/gtf/homo_sapiens/Homo_sapiens.GRCh38.110.gtf.gz -O example_data/Homo_sapiens.GRCh38.110.gtf.gz
    gzip -d example_data/Homo_sapiens.GRCh38.110.gtf.gz

#### Running Sheriff
***NOTE*** The Superb-seq paper included three sequencing libraries. Two of these libraries were on the same 
edited K562 cell lines, with edits generated by 7 guides targeting the exons of chromatin remodeler genes CHD3, CHD4, SMARCA4, and ARID1A. 
One library sequenced 10,000 cells at 100,000 reads/cell, and the other library sequenced 500 cells at 200,000 reads/cell.

The bam file used here is for the more deeply sequenced 500 cell library, with reads subsetted to those occurring
with 200kb of true edit sites that were identified in the 10,000 cell library. 

Therefore the edit site calling will be accurate, ***BUT*** the gene counts will NOT be
accurate, because most reads have been filtered from the example bam file. 
This filtering was performed so the smaller bam could be easily made available via github.

For more details on the Superb-seq libraries (including Sheriff outputs on GEO) and the inner processes of Sheriff, 
please refer to the manuscript listed at the end of this document.

#### Setting parameters
***NOTE*** 
For minimum cells to call canonical edit site, a minimum of 3 cells was used for the 10,000 library.
The CNV file input shown is only necessary if copy number variantion is present in the genome of the edited cells. 
Exclusion will result in 2 gene copies assumed throughout the genome. See the 'ploidy' parameter.
If pooling multiple cell lines with different copy number variation, will need to account for this in down-stream analysis. 

Expected run time for this demo is <30 seconds.

    dir_="example_data/"
    bam_="${dir_}barcode_headAligned_anno.sorted.edit_regions_200kb.bam"
    ref_="${dir_}Homo_sapiens.GRCh38.dna.primary_assembly.fa"
    cells_="${dir_}barcode_whitelist.500-cell.txt"
    gtf_="${dir_}Homo_sapiens.GRCh38.110.gtf"
    cnv_="${dir_}K562_CNV_hg19.tsv"
    blacklist_="${dir_}black_100x_peaks_by_qval.simple_repeats_50N.EXTRA.bed"
    blacklist_seqs="${dir_}blacklist_seqs.txt"
    min_="1" 
    cpu="1"
    out_dir="./subset_500_cell_sheriff_output/"

    sheriff ${bam_} ${ref_} ${cells_} ${gtf_} -cpu ${cpu} --cnv_file ${cnv_} --blacklist_file ${blacklist_} --blacklist_seqs ${blacklist_seqs} --edit_site_min_cells ${min_} -o ${out_dir} 

Output
------

***NOTE*** the parquet.gz files listed below are a binary format (even when unzipped). 
Example code to read these files in Python and R is provided below.

The output directory for this example is: 

    ./subset_500_cell_sheriff_output/

Output files include:

- **edit_site_info.txt**: Has the edit site information, name, location, window size, and overlapping genes.
- **edit_sites.bed**: Bed file of the edit site locations.
- **t7_barcode_edits.tsv**: Particular edit events, with detail on which canonical-edit-site the edit belongs to,
                            and the ref and alt sequence at the particular edit site.
- **cell_allelic_dosage.canonical-edit-sites.parquet.gz**: (cell X edit-site) matrix of counts, for called allelic edits.
- **cell_allelic_dosage.canonical-edit-sites.gene-collapsed.parquet.gz**: (cell X edit-site OR edit-gene) matrix of allelic edits, but collapse to the gene level for edit-sites that intersect genes.
- **cell_gene_mrna_counts.parquet.gz**: (cell X gene) The UMI counts per gene, excluding confounding T7 reads.
- **t7_barcoded_counts.parquet.gz**: (cell X edit-site) matrix with number of t7 barcoded UMIs per cell.
- **t7_nonbarcoded_counts.parquet.gz**: (cell X edit-site) matrix with number of UMIs per cell for reads that were +/- some distance from canonical edit sites, and were subsequently filtered from gene counting.
- **t7_all_counts.parquet.gz**: (cell X edit-site) matrix of all T7 related UMIs, addition of both t7_barcoded_counts.parquet.gz AND t7_nonbarcoded_counts.parquet.gz.

#### Below are just for sanity checking / visualisation, but less important for downstream analysis
- **t7_reads.txt**: List of call t7 read names in bam.
- **t7_barcoded_reads.txt**: List of t7 barcoded reads in bam.
- **t7_non-barcoded_reads.txt**: List of t7 non-barcoded reads in bam.
- **t7_filt.bam**: Bam file containing only the mRNA reads, i.e. removes reads +/- a certain bp range from the canonical edit sites.
- **t7_only.bam**: Bam file containing only the t7 reads, which includes barcoded reads as well as some reads that could potentially be t7 reads.
- **t7_non-barcoded_only.bam**: Bam containing all reads that were within some +/- bp distance from canonical edit sites, potential non-barcoded T7 reads.
- **t7_barcoded_only.bam**: Bam file containing only the BARCODED t7 reads, called be the kmer match in the 5' soft-clip.

Reading output
------
***NOTE*** for the parquet.gz files, these can be rapidly read with pandas in Python:

    python

    import pandas as pd

    cell_by_gene_edit_dosage = pd.read_parquet("subset_500_cell_sheriff_output/cell_allelic_dosage.canonical-edit-sites.gene-collapsed.parquet.gz")

Or in R:

    mamba install r-base

    R

    install.packages("arrow")
    library(arrow)

    cell_by_gene_edit_dosage <- as.data.frame( read_parquet("subset_500_cell_sheriff_output/cell_allelic_dosage.canonical-edit-sites.gene-collapsed.parquet.gz") )
    rownames(cell_by_gene_edit_dosage) <- cell_by_gene_edit_dosage[,'__index_level_0__']
    cell_by_gene_edit_dosage <- cell_by_gene_edit_dosage[, 1:dim(cell_by_gene_edit_dosage)[2]-1]

Citation
--------

Joint single-cell profiling of CRISPR-Cas9 edits and transcriptomes reveals widespread off-target events and their effects on gene expression
Michael H. Lorenzini, Brad Balderson, Karthyayani Sajeev, Aaron J. Ho, Graham McVicker
bioRxiv 2025.02.07.636966; doi: https://doi.org/10.1101/2025.02.07.636966

Contact
-------

Authors: Brad Balderson, Mickey Lorenzini, Aaron Ho, Graham McVicker

Contact: 

Change log
-------

    * v1.1.4 Fixed slight non-determinism in the outputted dosages, due to set usage causing random ordering in get_longest_edits function.
    * v1.1.3 Fixed UMI counting bugs, that were introduced during the run-speed optimization of v1.1.0.
    * v1.1.1 Minor-change in the gene-counting logic, to account for different split-pipe version outputs.
    * v1.1.0 Run-speed optimized and parallelisation for gene counting.
    * v1.0.0 Initial version used for manuscript that was uploaded to bioarchive.
    
    
