## Additional Files for "Joint single-cell capture of Cas9 edits and transcriptomes reveals on- and off-target effects on gene expression": setup.py

```
from setuptools import setup, find_packages

with open("README.md", "r") as fh:
    long_description = fh.read()

setup(
    name='Sheriff',
    version='1.1.4',
    author='Brad Balderson, Michael Lorenzini, Aaron Ho',
    author_email='',
    packages=find_packages(),
    #license=TBD #'GPL-3.0',
    long_description_content_type="text/markdown",
    long_description=long_description,
    scripts=['bin/sheriff'],
    # install_requires = [#'pandas',
    #                     'gtfparse==2.5.0', 'faiss-cpu==1.10.0',
    #                     #"numpy<2"
    #                     ],
    #install_requires = ['numpy','pandas'],
    #install_requires = ['pandas', 'gtfparse==2.5.0', 'faiss-cpu', "numpy<2"],
    entry_points={
        'console_scripts': [
        'sheriff=sheriff.__main__:main',
        'Sheriff=sheriff.__main__:main'
        ]
    },
    python_requires='>=3',
    description=("Sheriff calls CRISPR/cas9 edit sites, and quantifies the number of edited alleles at each site along "
                 "with gene expression in single cells using Superb-seq data."),
    keywords=['Sheriff', 'sheriff', 'Superb-seq', 'superb-seq'],
    url='https://github.com/BradBalderson/Sheriff',
    classifiers=[
        "Programming Language :: Python :: 3",
        "Development Status :: 5 - Production/Stable",
        "Topic :: Scientific/Engineering :: Bio-Informatics :: CRISPR",
        # license TBD "License :: OSI Approved :: GNU General Public License v3 or later (GPLv3+)",
    ],
)
```
