## Additional Files for "Joint single-cell capture of Cas9 edits and transcriptomes reveals on- and off-target effects on gene expression": __main__.py

```
from pathlib import Path
from typing import List, Optional
from typing_extensions import Annotated

import typer
import sys

# Local Imports
from .count_t7 import run_count_t7

# app = typer.Typer()
# app = typer.Typer(pretty_exceptions_show_locals=False)
app = typer.Typer(pretty_exceptions_short=False)

@app.command()
def get_t7_edits(
    bam_file: Annotated[str, typer.Argument(help="BAM file")],
    ref_file: Annotated[str, typer.Argument(help="Fasta containing ref genome")],
    barcode_file: Annotated[str, typer.Argument(help="Text file containing whitelisted barcode per line")],
    gtf_file: Annotated[str, typer.Argument(help="GTF file containing relevant gene data")],
    t7_barcode: Annotated[
        Optional[str],
        typer.Option(
            "--t7",
            "--t7_barcode",
            help=(
                "Target/query barcode sequence to denote t7 reads."
                )
            )
        ] = "GGGAGAGTAT",
blacklist_file: Annotated[
        Optional[str],
        typer.Option(
            "--blacklist",
            "--blacklist_file",
            help=(
                "Bed file that species the location of blacklist regions, these generate alot of endogenuous t7 reads "
                "that can lead to slow processing time and false-positive edit-site calling."
                )
            )
        ] = None,
whitelist_file: Annotated[
        Optional[str],
        typer.Option(
            "--whitelist",
            "--whitelist_file",
            help=(
                "Bed file that species the location of whitelist regions, which are known edit sites and so will call "
                "any barcoded reads implying an edit site intersecting these regions as canonical edit sites."
                )
            )
        ] = None,
    k: Annotated[
        Optional[int],
        typer.Option(
            "-k",
            "--kmer",
            "--kmer_size",
            help=("Size of kmers used to pattern match read barcodes to the t7 barcode."
                  )
            )
        ] = 6,
    edit_dist: Annotated[
        Optional[int],
        typer.Option(
            "--edit_dist",
            "--edist",
            "--dist",
            help=("+/- distance from edit site to be grouped as same edit."
                  )
            )
        ] = 140,
    edit_site_rev_comp_filt: Annotated[
        Optional[bool],
        typer.Option(
            "--bidirectional_inserts/--no-bidirectional_inserts",
            help=(
                "Candidate edit site must have evidence of bi-directional donor insertion to be called as a canonical edit site."
                "Highly recommended criteria. If turned off, it also turns off --stranded_edit_dist criteria, but this "
                "information about the edit sites are still recorded in the output via the 'stranded_edit_dist' column"
                "in edit_site_info.txt"
                )
            )
        ] = True,
    stranded_edit_dist: Annotated[
        Optional[int],
        typer.Option(
            "--stranded_edit_dist",
            help=("Maximum allowed distance between the nearest forward and reverse edit sites at a given canonical edit site to qualify as real edit."
                  )
            )
        ] = 15,
    edit_site_min_cells: Annotated[
        Optional[int],
        typer.Option(
            "--edit_site_min_cells",
            help=("Minimum cells in edit site to be considered true edit."
                  )
            )
        ] = 3,
    nonbc_edit_dist: Annotated[
        Optional[int],
        typer.Option(
            "--nonbc_edit_dist",
            "--nonbc_edist",
            "--nonbc_dist",
            "--nonbc",
            help=("+/- distance from edit to mop up the non-barcoded reads."
                  )
            )
        ] = 1000,
    ploidy: Annotated[
        Optional[int],
        typer.Option(
            "--ploidy",
            help=("Ploidy/Number of chromosomes in the genome."
                  )
            )
        ] = 2,
    cnv_file: Annotated[
        Optional[str],
        typer.Option(
            "--cnv",
            "--cnv_file",
            "--copy_number_variant_file",
            help=(
                "A bedGraph file that specifies copy-number-variation sites, "
                "that deviate from the ploidy number."
                )
            )
        ] = None,
    blacklist_seqs: Annotated[
        Optional[str],
        typer.Option(
            "--blacklist_seqs",
            help=(
                "Text file of sequences, with a new sequence on each line, that may be present in read soft-clip sequences"
                "can confound t7 barcoded read calls. Currently only the TSO, which is a common left-over artifact."
                )
            )
        ] = None,
    mrna_count_mode: Annotated[
        Optional[str],
        typer.Option(
            "--mrna_count_mode",
            help=(
                "Mode for quantifying gene expression,"
                "'all' is to count all reads associated with a gene, 'polyT' is to only count polyT reads, indicating mature mRNA transcripts."
                )
            )
        ] = "all",
    uncorrected_gene_count: Annotated[
        Optional[bool],
        typer.Option(
            "--uncorrected_count/--no-uncorrected_count",
            help=(
                "Whether to also output the gene counts WITHOUT removing reads around inferred edit sites."
                )
            )
        ] = False,
    outdir: Annotated[
        Optional[str],
        typer.Option(
            "-o",
            "--out",
            "--outdir",
            "--out_dir",
            help=(
                "Write output files to this location. "
                "Defaults to Current Working Directory"
                ),
            )] = None,
    verbosity: Annotated[
            Optional[int],
            typer.Option(
                "--v",
                "-v",
                "--verbosity",
                "-verbosity",
                help=("Verbosity levels. 0 errors only, 1 prints processing progress, 2 prints debugging information."
                      )
                )
            ] = 1,
    n_cpus: Annotated[
            Optional[int],
            typer.Option(
                "--cpu",
                "-cpu",
                help=("Number of CPUs to use for processing, necessary to increase this for fast UMI counting."
                      )
                )
            ] = 1,
chunk_size_mb: Annotated[
            Optional[int],
            typer.Option(
                "--chunk",
                "-chunk",
                help=("Number of mega-bases to process at a time for gene UMI counting. Set this lower if get memory"
                      "issues, currently set to > hg38 chr1 size (249 Mb), so is parallelized by chromosome."
                      "Trade-off is can cause minor double-counting of UMIs if a genome chunk intersects a gene."
                      "Very small / negligible difference in counts."
                      )
                )
            ] = 250,
):
    
    # Run
    
    # Test outputs
    run_count_t7(bam_file=bam_file,
                 ref_file=ref_file,
                 barcode_file=barcode_file,
                 gtf_file=gtf_file,
                 t7_barcode=t7_barcode,
                 blacklist_file=blacklist_file,
                 whitelist_file=whitelist_file,
                 k=k,
                 edit_dist=edit_dist,
                 stranded_edit_dist=stranded_edit_dist,
                 edit_site_min_cells=edit_site_min_cells,
                 nonbc_edit_dist=nonbc_edit_dist,
                 ploidy=ploidy,
                 copy_number_variant_file=cnv_file,
                 blacklist_seqs=blacklist_seqs,
                 mrna_count_mode=mrna_count_mode,
                 outdir=outdir,
                 edit_site_rev_comp_filt=edit_site_rev_comp_filt,
                 max_gene_count_reads=None,
                 uncorrected_gene_count=uncorrected_gene_count, # For testing
                 constrain_allele_calls=False,
                 verbosity=verbosity,
                 n_cpus=n_cpus,
                 chunk_size_mb=chunk_size_mb,
                 )

def main():
    root_dir = Path(__file__).parent
    sys.path.append(str(root_dir))
    app()

if __name__ == "__main__":
    main()
```
