## Additional Files for "Joint single-cell capture of Cas9 edits and transcriptomes reveals on- and off-target effects on gene expression": count_t7.py

```
import sys
import timeit 
from pathlib import Path
from collections import defaultdict, namedtuple

import os
import numpy as np
import pandas as pd
import polars as pl

import pyranges as pr

import faiss
import gtfparse # Maybe put on top for now until optional working

import contextlib

import pysam
from pysam.libcalignmentfile import AlignmentFile

# Same directory, shouldn't need relative import. Throws error otherwise
from .helpers import (get_t7_count_matrix, get_cell_counts_from_umi_dict, bam_count_gene_umis,
                      get_longest_edits, bed_file_flag_edits)

# Import of helper functions !
# from .helpers import get_t7_count_matrix, get_cell_counts_from_umi_dict, bam_count_gene_umis, bio_edit_distance, \
#                     get_edit_sets, get_longest_edits

@contextlib.contextmanager
def stdoutsuppress():
    """Suppress stdout and stderr globally, including C-level output."""
    with open(os.devnull, 'w') as fnull:
        original_stdout_fd = os.dup(sys.stdout.fileno())
        original_stderr_fd = os.dup(sys.stderr.fileno())

        os.dup2(fnull.fileno(), sys.stdout.fileno())
        os.dup2(fnull.fileno(), sys.stderr.fileno())

        try:
            yield
        finally:
            os.dup2(original_stdout_fd, sys.stdout.fileno())
            os.dup2(original_stderr_fd, sys.stderr.fileno())

# class that handles kmer matching
class KmerMatcher:

    def __init__(self, k, sequences=None):
        
        self.k = k
        self.hash_symbol = {"A":0, "C":1, "G":2, "T":3} # symbol to num
        self.hash_num = ["A", "C", "G", "T"] # Num to symbol
        self.match_set = set() # not properly matching as a set, need a hash
        self.match_hash = []
        
        if sequences is not None:
            # Load in kmer initial input
            self.update_matches(sequences)
    
    def update_matches(self, sequences):
        if isinstance(sequences, str):
            self.add_kmers(sequences)
        else:
            for seq in sequences:
                self.add_kmers(seq)

    def add_kmers(self, seq):
        # NOTE that this was updated NOT to include the reverse compliment, since need to separate this, only do rev-comp
        # for reverse reads!
        #rev_seq = self.revcomp(seq) # get reverse complement
        
        # Get forward and reverse kmers
        kmers = [seq[i: i + self.k] for i in range(len(seq) - self.k + 1)]
        #rev_kmers = [rev_seq[i: i + self.k] for i in range(len(rev_seq) - self.k + 1)]

        # Update the match set
        # self.match_set.update(kmers + rev_kmers)
        # self.match_hash.extend([self.kmer_to_num(i) for i in kmers + rev_kmers])
        self.match_set.update( kmers )
        self.match_hash.extend([self.kmer_to_num(i) for i in kmers])
        self.match_hash.sort() # Keep the hash list sorted

    
    def kmer_to_num(self, kmer):

        if len(kmer) < 1:
            return 0

        return (4*self.kmer_to_num(kmer[:-1:])) + self.hash_symbol[kmer[-1]]

    def num_to_kmer(self, num, k):

        if k == 1:
            return ["A", "C", "G", "T"][num]

        prefix_num, remainder = divmod(num, 4)

        return self.num_to_kmer(prefix_num, k-1) + self.hash_num[remainder]

    @staticmethod
    def revcomp( seq ):
        """ Helper method if also want to make a kmer-matcher for the reverse complement sequence.
        """
        comp_table = str.maketrans('ATCG', 'TAGC')  # used for revcomp
        return seq[::-1].translate( comp_table )

# GOTTA BE A BETTER WAY
# TRY DOING A DECORATOR, MIGHT MAKE MOST SENSE
def reformat_chr_name(read):
    """ Reformats the chromosome name attached to the read to be conistent with the reference genome naming.
    In this case, the bam stores the chromosome names as hg38_1 (for chr1). But the fasta stores as 1. So just 
    reformats like that.
    """
    return read.reference_name.replace('hg38_', '')

def match_kmer(bc_kmer_matcher, indel_seq, output_kmer_hash):
    """Gets kmer matches
    """
    k = bc_kmer_matcher.k
    match_kmers = bc_kmer_matcher.match_hash

    # Will count every barcode occurance in N time, uses numerical hashes
    # Scalable and allows for mismatches amongst other things
    freq_array = np.zeros((4 ** k), dtype=np.uint8)

    try:
        freq_array[[bc_kmer_matcher.kmer_to_num(indel_seq[i: i + k])
                    for i in range(len(indel_seq) - k + 1)]] += 1
    except KeyError:
        # We ran into an N, which can't be hashed
        # Not ideal way to handle, but seems rare enough
        freq_array[
            [
                bc_kmer_matcher.kmer_to_num(kmer) for kmer in
                [
                    indel_seq[i: i + k] for i in range(len(indel_seq) - k + 1)
                ]
                if 'N' not in kmer
            ]
        ] += 1

    kmer_matches = freq_array.nonzero()[0]

    # For our use match_kmers won't be None
    if match_kmers is not None:
        kmer_matches = kmer_matches[np.isin(kmer_matches, match_kmers)]

        if kmer_matches.size == 0:
            kmer_matches = None
        elif output_kmer_hash:
            # Show matches as hash index instead of kmers
            # Not sure if this is best idea, but needed to hash into set
            kmer_matches = tuple(kmer_matches)
        else:
            # Show kmer matches instead of hash
            kmer_matches = tuple(bc_kmer_matcher.num_to_kmer(i, k) for i in kmer_matches)

    return kmer_matches

# Updated process forward and reverse
def match_barcode_forward(read, fasta, bc_kmer_matcher, output_kmer_hash=False):

    # Need enough space for insert
    if read.query_alignment_start < bc_kmer_matcher.k:
        return None
    
    indel_seq = read.query_sequence[:read.query_alignment_start]
    
    kmer_matches = match_kmer(bc_kmer_matcher, indel_seq, output_kmer_hash)

    # n_matches = freq_array[match_kmers].nonzero()[0].shape[0] # if match kmers not zero

    # t7_barcode = 'GGGAGAGTAT'  # barcode
    # rev_comp = 'ATACTCTCCC'  # rev cop, just for reference

    # Get ref seq from fasta
    chr_name = reformat_chr_name(read)

    ref_pos = read.reference_start

    try:
        ref_seq = fasta.fetch(chr_name, ref_pos, ref_pos+1)#ref_pos+3)
    except:
        print(f"Warning could not load from reference sequence to construct REF: {chr_name}: {ref_pos}-{ref_pos+1}",
              file=sys.stdout, flush=True)
        return None
    
    alt_seq = indel_seq + ref_seq

    # If defined as ReadEdit(chrom, ref_pos, ref_seq, alt_seq, forward, kmer_matches)
    
    # Stick this here if it's being wonky
    ReadEdit = namedtuple("ReadEdit", ["chrom", "ref_pos", "ref_seq", "alt_seq", "forward", "kmer_matches"])

    return ReadEdit(read.reference_name, ref_pos, ref_seq, alt_seq, read.is_forward, kmer_matches)

    
def match_barcode_reverse(read, fasta, bc_kmer_matcher, output_kmer_hash=False):

    if read.query_length - read.query_alignment_end < bc_kmer_matcher.k:
        return None
    
    indel_seq = read.query_sequence[read.query_alignment_end:]

    kmer_matches = match_kmer(bc_kmer_matcher, indel_seq, output_kmer_hash)

    # n_matches = freq_array[match_kmers].nonzero()[0].shape[0] # if match kmers not zero

    # t7_barcode = 'GGGAGAGTAT'  # barcode
    # rev_comp = 'ATACTCTCCC'  # rev cop, just for reference

    # Get ref seq from fasta
    chr_name = reformat_chr_name(read) # TEMPORARY SOLUTION UNTIL CHECK
    
    ref_pos = read.reference_end - 1

    # Added this after I got an error during processing, not sure of problem YET
    try:
        ref_seq = fasta.fetch(chr_name, ref_pos, read.reference_end)
    except:
        print(f"Warning could not load from reference sequence to construct REF: {chr_name}: {ref_pos}-{read.reference_end}",
              file=sys.stdout, flush=True)
        return None
    
    alt_seq =  ref_seq + indel_seq # Reference sequence occurs to the right for reverse-reads!

    # If defined as ReadEdit(chrom, ref_pos, ref_seq, alt_seq, forward, kmer_matches)
    ReadEdit = namedtuple("ReadEdit", ["chrom", "ref_pos", "ref_seq", "alt_seq", "forward", "kmer_matches"])

    return ReadEdit(read.reference_name, ref_pos, ref_seq, alt_seq, read.is_forward, kmer_matches)

def get_blacklist_kmer_matches(edit_data, blacklist_seq_kmers_rev, blacklist_matcher_rev):
    """ Gets blacklist kmers that match the soft-clip seq of a particular read!
    """
    #### Will score against reverse of matched blacklist seqs
    if edit_data.forward:
        indel_seq = edit_data.alt_seq[:-len(edit_data.ref_seq)]
    else:
        indel_seq = edit_data.alt_seq[len(edit_data.ref_seq):]

    blacklist_kmer_matches = set()
    for kmer_ in edit_data.kmer_matches:
        if kmer_ in blacklist_seq_kmers_rev:  # Has the barcode kmer match to the blacklist seq!
            blacklist_match = blacklist_seq_kmers_rev[kmer_]

            blacklist_matcher_ = blacklist_matcher_rev[blacklist_match]
            kmer_matches = match_kmer(blacklist_matcher_, indel_seq, False)
            blacklist_kmer_matches = blacklist_kmer_matches.union(set(kmer_matches))

    return blacklist_kmer_matches

# UPDATED FOR EFFICIENCY
def get_barcoded_edits(bam_file, cell_barcodes,
                       ref_fasta_file,
                       k, t7_barcode,
                       output_kmer_hash=False, blacklist_seqs=None,
                       print_freq=1000000, verbosity=1):

    # This dictionary keeps track of faiss indices PER chromosome, for fast neighbourhood lookup.
    # This is used downstream to group edits by 'canonical' edit!
    chr_edit_loc_indices = {}
    # For each chromosome, this stores a list of the edit_data's as they were added to the chromosome indices
    # for fast nearest-neighbour lookup. This is neccesary so that when we lookup the nearest-neighbours, will
    # only get the indices of these neighbours in the chromosome index. Therefore, need to map these back
    # to the global set of edit_datas !
    chr_edits_order_added = defaultdict( list )

    ### Old version, which was more 'cell-centric' metadata. Is tricky to do posthoc filtering with this form
    ### of storing the data, so will make it all 'edit-centric', and reverse it to be 'cell-centric' metadata
    ### after filtering out initially called edits which are not well supported.
    # t7_barcoded_reads = []
    # read_edits = []
    # bc_edits = defaultdict(list)  # list iteration faster than set for part 2
    # bc_umi = defaultdict(set)

    # Edit centric data structures.
    edit_counts = {} # ReadEdit to number of unique cell barcodes, updated throughout
    edit_bc_cell_umis = defaultdict(dict) # edit to barcode to umis
    edit_reads = defaultdict(list) # Edit to list of supporting barcoded t7 reads

    # Holds edit data if read is t7...doesn't seem to work if defined here and called in match func
    # ReadEdit = namedtuple("ReadEdit", ["chrom", "ref_pos", "ref_seq", "alt_seq", "forward", "kmer_matches"])

    bc_kmer_matcher_forward = KmerMatcher(k, t7_barcode)
    bc_kmer_matcher_reverse = KmerMatcher(k, KmerMatcher.revcomp(t7_barcode))

    if type(blacklist_seqs)!=type(None): # Need to handle blacklist sequences
        print("Handling blacklist seqs: ", blacklist_seqs,
              file=sys.stdout, flush=True) if verbosity >= 1 else None
        blacklist_seqs = [line.strip('\n') for line in open(blacklist_seqs, 'r')]
        # Get if and what kmers in the barcode match those in the blacklist sequences,
        #  any reads that have these kmers below will need to be scored for the blacklist seqs as well,
        #  and the set difference taken to see if there are still additional kmers as evidence for the barcode being
        #  still being present in the read soft-clip.
        # The blacklist seqs will only be considered if have kmers that match the actual barcode!
        blacklist_seq_kmers_forward = {} # Kmer objects for inputted blacklist sequences that have a match for the barcode
        blacklist_seq_kmers_rev = {}
        blacklist_matcher_forward = {}
        blacklist_matcher_rev = {}
        for seq_ in blacklist_seqs:
            blacklist_matcher_forward[ seq_ ] = KmerMatcher(k, seq_)
            blacklist_matcher_rev[ seq_ ] = KmerMatcher(k, KmerMatcher.revcomp(seq_))

            kmer_matches_forward = match_kmer(bc_kmer_matcher_forward, seq_, False)
            kmer_matches_rev = match_kmer(bc_kmer_matcher_reverse, KmerMatcher.revcomp(seq_), False)
            for kmer in kmer_matches_forward:
                blacklist_seq_kmers_forward[kmer] = seq_
            for kmer in kmer_matches_rev:
                blacklist_seq_kmers_rev[kmer] = seq_

    with AlignmentFile(bam_file, "rb") as bam, pysam.FastaFile(ref_fasta_file) as fasta:

        start_time = timeit.default_timer()
        
        idx_stats = bam.get_index_statistics()
        total_reads = np.sum([stat.total for stat in idx_stats])
        
        for i, read in enumerate(bam):

            cell_barcode = read.get_tag('CB')
            # If the cell barcode is NOT in the cell barcode white list, do not neeed to process.
            if cell_barcode not in cell_barcodes:
                continue

            if i % print_freq == 0 and verbosity==1:
                print(f"PROCESSED {i} / {total_reads} reads in {(timeit.default_timer()-start_time)/60:.3f} minutes",
              file=sys.stdout, flush=True)

            if read.is_forward:
                edit_data = match_barcode_forward(read, fasta, bc_kmer_matcher_forward, output_kmer_hash)
            else:
                edit_data = match_barcode_reverse(read, fasta, bc_kmer_matcher_reverse, output_kmer_hash)

            # If edit data contained barcode kmer match, then it's a t7 read
            if (edit_data is not None) and (edit_data.kmer_matches is not None):

                # HERE is where I additionally check for TSO match...
                if type(blacklist_seqs) != type(None):  # Need to handle blacklist sequences

                    ### Checking for match with blacklist kmers:
                    potential_blacklist = False # Need to keep track if went into below if-statements.
                    if read.is_forward and np.any([kmer_ in blacklist_seq_kmers_forward
                                                   for kmer_ in edit_data.kmer_matches]):
                        blacklist_kmer_matches = get_blacklist_kmer_matches(edit_data, blacklist_seq_kmers_forward,
                                                                                                  blacklist_matcher_forward)
                        potential_blacklist = True

                    elif read.is_forward==False and np.any([kmer_ in blacklist_seq_kmers_rev
                                                            for kmer_ in edit_data.kmer_matches]):
                        blacklist_kmer_matches = get_blacklist_kmer_matches(edit_data, blacklist_seq_kmers_rev,
                                                                                                      blacklist_matcher_rev)
                        potential_blacklist = True

                    ###### Now will get the set difference between the blacklist seq matching kmers and the barcode
                    ###### kmers, if there are no barcode kmers left as evidence for a match, AND there are still a ton
                    ##### of blacklist kmer matches, then this indicates is actually a match to the TSO. BUT if has no
                    ##### other kmer matches to the TSO, then will still consider it a barcode match.
                    if potential_blacklist:
                        non_blacklist_kmers = set(edit_data.kmer_matches).difference( blacklist_kmer_matches )
                        blacklist_kmers = blacklist_kmer_matches.difference( set(edit_data.kmer_matches) )
                        if len(non_blacklist_kmers)==0 and len(blacklist_kmers) > 0: # Meets criteria for TSO ONLY match
                            continue # Continue without recording as barcoded read, since actually ONLY a TSO artifact

                # Add unique edits to dict
                # if edit_data not in bc_edits[cell_barcode]:
                #     bc_edits[cell_barcode].append(edit_data)
                if cell_barcode not in edit_bc_cell_umis[edit_data]:
                    edit_bc_cell_umis[edit_data][cell_barcode] = set() # umi as set
                    # edit_bc_cell_umis[edit_data][cell_barcode] = []

                    if edit_data in edit_counts: # Logged this before
                        edit_counts[edit_data] += 1

                    else: # Have not logged, will need to add it to the index as well!
                        edit_counts[edit_data] = 1

                        if edit_data.chrom not in chr_edit_loc_indices: # Need to start a new index!
                            chr_index = faiss.IndexFlat1D()
                            chr_edit_loc_indices[edit_data.chrom] = chr_index
                        else:
                            chr_index = chr_edit_loc_indices[edit_data.chrom]

                        # Not implemented for relevant index
                        #edit_id = list( edit_counts.keys() ).index( edit_data ) # edit ID will be the location of the edit in edit_count
                        # chr_index.add_with_ids(np.array(edit_data.ref_pos).reshape(-1, 1),
                        #                        np.array(edit_id).reshape(-1) )

                        #### Adding to index, can keep track of edit_data to this index since will be the
                        #### same order as the keys in edit_counts.
                        chr_index.add( np.array(edit_data.ref_pos).reshape(-1, 1) )
                        # Log what edit was added to this chromosome, so can look it up from the nearest-neighbour
                        # indices later.
                        chr_edits_order_added[ edit_data.chrom ].append( edit_data )
                        # Example of getting all of the other edits within a distance of 1 position from itself!
                        # chr_index.range_search(np.array(edit_data.ref_pos).reshape(-1,1), 1)

                # Old cell-centric version
                # t7_barcoded_reads.append(read.query_name)
                # read_edits.append( edit_data )
                # bc_umi[cell_barcode].add(read.get_tag('pN'))

                # Edit-centric version, for ease of post-hoc filtering once collapsed to canonical edit sites.
                edit_bc_cell_umis[edit_data][cell_barcode].add(read.get_tag('pN'))
                # edit_bc_cell_umis[edit_data][cell_barcode].append( read.get_tag('pN') )
                edit_reads[edit_data].append( read.query_name )

    # Edit-centric version, for ease of post-hoc filtering once collapsed to canonical edit sites.
    return edit_counts, edit_bc_cell_umis, edit_reads, chr_edit_loc_indices, chr_edits_order_added


def get_nonbarcoded_edits(bam_file, canonical_to_edits, canonical_to_edited_cells,
                          cells_to_canonical_and_edits, edit_reads_filtered,
                          dist=1000, verbosity=1):
    """ MODIFIED from the original above to be more edit-centric, i.e. INSTEAD of iterating through every read in the
    bam file and seeing if it matches the criteria for an edited bam (extremely slow), will iterate through every
    EDIT SITE, pulling out reads within a certain distance from that edit site using pysam fetch, and then checking
    each of these reads against each of the particular single cell edits that have been called to see if it matches a
    non-t7 edited read criteria!

    :param bam_file:
    :param t7_barcoded_reads:
    :param bc_edits:
    :param dist:
    :return:
    """
    # Iterating through each canonical edit site, pulling out all reads in the bam the overlap this site to call as
    # non-barcoded t7 edit or not!
    bam_file_handle = pysam.AlignmentFile(bam_file, 'rb')
    start_time = timeit.default_timer()

    # Storing the non-barcoded read information !
    t7_nonbarcoded_reads = []
    nonbarcoded_umi = defaultdict(set)

    # edit_no_bc_cell_umis = defaultdict(dict)
    canonical_edit_no_bc_cell_umis = defaultdict(lambda: defaultdict(set))

    for i, (edit_site, edits) in enumerate( canonical_to_edits.items() ):

        edit_chr = edit_site.chrom
        edit_pos = edit_site.ref_pos
        edit_window_start = max([edit_pos-dist, 0]) # Accounting for edge of chromosome.
        edit_window_end = edit_pos+dist # pysam is OK with trying to fetch greater than chromosome length.

        # Getting set of cells that are called as edited for this edit site
        edit_cells = canonical_to_edited_cells[edit_site]

        # Getting set of barcoded reads that overlap this edit site
        t7_barcoded_read_set = []
        [t7_barcoded_read_set.extend(edit_reads_filtered[edit_data]) for edit_data in edits]
        t7_barcoded_read_set = set( t7_barcoded_read_set )

        read_positions = []
        for read in bam_file_handle.fetch(edit_chr, edit_window_start, edit_window_end):
            ### Only need to filter if is a edited cell.
            cell_barcode = read.get_tag('CB')

            ### Already counted as barcoded, so don't need to filter.
            if read.query_name in t7_barcoded_read_set:
                continue

            read_positions.append( read.pos )

            # NOTE DECIDED to completely relax this criteria, so that we essentially black-list around all
            # canonical edit sites so they do not contribute to the mRNA counting downstream.
            edit_pos = edit_site.ref_pos  # This is actually where the insert starts
            read_edit_dist = read.pos - edit_pos

            if (abs(read_edit_dist) <= dist):
                #### Meets criteria for non-barcoded t7 edit!!!
                t7_nonbarcoded_reads.append(read.query_name)
                nonbarcoded_umi[cell_barcode].add( read.get_tag('pN') )

                # Keeping track of all non-bc t7s, associating with each canonical edit-site
                canonical_edit_no_bc_cell_umis[edit_site][cell_barcode].add(  read.get_tag('pN') )

        ### For debugging to make sure the windowing strategy is correct !!
        ### NOTE observed SOME reads that are slightly outside the query region, reading here it appears that this occurs
        ### if there is a partial alignment that DOES overlap with the position. This is handled by still applying the
        ### distance criteria above
        # import matplotlib.pyplot as plt
        # fig, ax = plt.subplots()
        # ax.hist(read_positions, bins=50)
        # ax.vlines(edit_window_start, 0, 100, color='red')
        # ax.vlines(edit_window_end, 0, 100, color='red')
        # plt.show()
        if i % 50 == 0 and verbosity==1:
            print(f"PROCESSED {i} canonical-edit-sites in {(timeit.default_timer() - start_time) / 60:.3f} minutes",
                                                                                            file=sys.stdout, flush=True)

    return t7_nonbarcoded_reads, nonbarcoded_umi, canonical_edit_no_bc_cell_umis


def run_count_t7(bam_file,
                 ref_file,
                 barcode_file,
                 gtf_file=None, # GTF file to call allelic edits per gene
                 t7_barcode="GGGAGAGTAT",
                 blacklist_file=None, # bed file specifying regions to exclude as potential edit sites, generally sites of high endogenuous t7 expression.
                 whitelist_file=None,
                 k=6,
                 edit_dist=140, # Distance from canonical edit site, for edit to be grouped with that edit!
                 stranded_edit_dist=15, # distance between the closest forward and reverse reads from the canonical edit site to classify as a true edit event (real edits tend to have a higly localised cut sequence!
                 edit_site_rev_comp_filt=True, # For a given edit site, IF there does NOT exist a reverse-complement version of the edit then exclude.
                 edit_site_min_cells=3, # Minimum number of cells the edit site must replicate in, to be considered a True edit.
                 nonbc_edit_dist=500, # Distance from edit to mop up the non-barcoded reads...
                 max_gene_count_reads=None, # Just for testing purposes, so does not become a bottle-neck.
                 uncorrected_gene_count=False, # Whether to also do the UMI counting WITHOUT correcting for t7...
                 ploidy=2, # Specifies overall ploidy number genome wide
                 copy_number_variant_file = None, # a bedGraph file that specifies copy-number-variation sites, that deviate from the ploidy number
                 constrain_allele_calls=True, # Whether or not to constrain the number alleles called to the inputted max alleles
                 blacklist_seqs = None, # Seqs to check for in the soft-clip sequences, that may collide with barcode.
                 mrna_count_mode = 'all', # Mode for quantifying gene expression,
                                            # 'all' is to count all reads associated with a gene,
                                            # 'polyT' is to only count polyT reads, indicating mature mRNA transcripts.
                 outdir=None, verbosity=1, n_cpus=1,
                 chunk_size_mb=15, #Mb to process at a time, in parallel.
                 ):
    
    # Process output data stuff
    if outdir is None:
        outdir = Path.cwd()
    
    out_path = Path(outdir)
    out_path.mkdir(parents=True, exist_ok=True) # Create dir before making output files

    # Load whitelisted barcodes
    with open(barcode_file) as file:
        cell_barcodes_list = [line.rstrip() for line in file]
        cell_barcodes_dict = {key: value for value, key in enumerate(cell_barcodes_list)}
        cell_barcodes = set(cell_barcodes_list)
    
    
    # Step 1: Get barcoded t7 edits
    print_freq = 1000000 # for testing
    
    start_bc = timeit.default_timer()
    print("Counting barcoded edits...", file=sys.stdout, flush=True) if verbosity >= 1 else None

    ## Edit-centric version, much easier to filter out edits that are not well-supported!
    edit_counts, edit_bc_cell_umis, edit_reads, chr_edit_loc_indices, chr_edits_order_added = get_barcoded_edits(
        bam_file,
        cell_barcodes,
        ref_file,
        k, t7_barcode,
        output_kmer_hash=False,
        blacklist_seqs=blacklist_seqs,
        print_freq=print_freq, verbosity=verbosity,
        )
    
    print(f"Processed barcoded edits in {(timeit.default_timer()-start_bc)/60:.3f} minutes\n",
          file=sys.stdout, flush=True) if verbosity>= 1 else None

    ####################################################################################################################
    # Step 2: Adding canonical 'edit site' labels to each of the identified edits, so can easily define a window around
    #       which to do additional secondary filtering for non-barcoded reads below!
    ####################################################################################################################
    edit_datas = list(edit_counts.keys()) # This defines the order of the edit_datas...
    edit_count_values = np.array(list(edit_counts.values()))

    EditSite = namedtuple("EditSite", ["chrom", "ref_pos"])

    edit_order = np.argsort(-edit_count_values)
    edit_indices_processed = [] # Keeping track of the indices of the edits already processed, so don't assign multiple labels!
    canonical_edit_sites = [] # Final set of canonical edit sites, with small variations collapsed
    canonical_edit_reversed = [] # For each edit site, is there the reverse compliment for that edit site.
    canonical_edit_cell_counts = [] # Counting number of cells with an edit at this canonical site.
    #edit_labels = np.full((len(edit_datas)), fill_value=np.nan) # Labels for all edits, indicating index of the canonical edit site for the edit.
    edits_to_canonical = {}
    canonical_to_edits = {}
    for orderi, editi in enumerate(edit_order):
        edit_data = edit_datas[editi]
        if editi not in edit_indices_processed:

            # Retrieving all edits within a pre-defined distance from this edit site!
            chr_edit_datas = chr_edits_order_added[ edit_data.chrom ]

            chr_index = chr_edit_loc_indices[ edit_data.chrom ]
            _, dists, close_neighbour_indices = chr_index.range_search(np.array(edit_data.ref_pos).reshape(-1, 1),
                                                                       edit_dist**2) # ^2 since faiss will use L2 norm.
            # Filtering to the indices that have not already been assigned!
            close_neighbour_indices_GLOBAL = np.array([edit_datas.index( chr_edit_datas[index] ) for index in close_neighbour_indices])
            keep_bool = [index not in edit_indices_processed for index in close_neighbour_indices_GLOBAL]
            dists, close_neighbour_indices = dists[keep_bool], close_neighbour_indices[keep_bool]
            close_neighbour_indices_GLOBAL = close_neighbour_indices_GLOBAL[keep_bool]

            # Logging the processed sites, so that we don't re-label by a less abundant nearby site!
            # OLD version, does not work if running on multiple chromosomes at same time!
            #edit_indices_processed.extend( close_neighbour_indices ) # NOTE editi is within these indices.
            # NEW version that considers the GLOBAL index of the edit_datas.
            edit_indices_processed.extend( close_neighbour_indices_GLOBAL )

            #### Sanity check; are the edit_data corresponding to each other?
            edits_match = [edit_datas[global_index] == chr_edit_datas[chr_index] for global_index, chr_index in
                           zip(close_neighbour_indices_GLOBAL, close_neighbour_indices)]
            edits_match_overall = np.all(edits_match)
            if not edits_match_overall:
                print("WARNING: Edits dont match between global and chromosome-specific indices!!!", file=sys.stdout,
                                                                                                             flush=True)

            #### Sanity check; do the edits at these indices have distances that are consistent with the dists listed?
            #manual_dists = np.array([abs(edit_data.ref_pos-edit_datas[neighi].ref_pos) for neighi in close_neighbour_indices_GLOBAL])
            manual_dists = np.array([abs(edit_data.ref_pos - chr_edit_datas[neighi].ref_pos) for neighi in close_neighbour_indices])
            #### This does not seem to be matching, I think Faiss is using some approximation for a
            #### larger index! let's just
            expected_range = np.all(manual_dists <= edit_dist)
            # dists_match = np.all(manual_dists == np.sqrt(dists))
            # if not dists_match:
            #     print("Warning: Distances don't match!",
            #   file=sys.stdout, flush=True)
                # I checked the few case where this is happening, and the distances are APPROXIMATELY correct;
                # it seems that the faiss index is doing some kind of approximate distance lookup which results in
                # some heuristic distance. They round in factors of 2 to the correct distance!
            if not expected_range:
                print("Warning: Distances NOT within expected range!", file=sys.stdout, flush=True)

            # Saving the canonical edit site, will just reference the position!
            edit_site = EditSite(edit_data.chrom, edit_data.ref_pos)
            canonical_edit_sites.append( edit_site )
            # Adding in the label for these nearby edits as this most common edit site!
            #edit_labels[close_neighbour_indices] = len(canonical_edit_sites)
            edits_to_this_canonical = {chr_edit_datas[neighi]: edit_site for neighi in close_neighbour_indices}
            edits_to_canonical.update( edits_to_this_canonical )
            canonical_to_edits[edit_site] = list( edits_to_this_canonical.keys() )

            # Determining if the edits associated with the canonical site has both forward and reverse direction,
            # which is good evidence of t7 insert site since can insert in either direction!
            canonical_site_directions = []
            canonical_edit_site_cell_set = set()
            #canonical_edit_cell_counts.append( 0 )
            for neighi in close_neighbour_indices:
                # OLD version, this works fine when parallelised by chromosome, BUT if running on a full bam across
                # chromosomes it introduces a bug!
                #neigh_edit = edit_datas[neighi]
                neigh_edit = chr_edit_datas[neighi]

                # Keeping track if there is evidence of an edit variation in both directions.
                canonical_site_directions.append( neigh_edit.forward )

                # Counting number of cells UNIQUE cells
                canonical_edit_site_cell_set = canonical_edit_site_cell_set.union( set(
                                                                          list(edit_bc_cell_umis[neigh_edit].keys()) ) )

            canonical_edit_cell_counts.append( len( canonical_edit_site_cell_set ) )

            # Has both forward and reverse direction!
            canonical_edit_reversed.append( len(set(canonical_site_directions)) == 2 )

        if orderi % 5000 == 0:
            print(f"Processed edit {orderi} / {len(edit_datas)}", file=sys.stdout, flush=True) if verbosity>= 1 else None

    print("Done calling canonical edits", file=sys.stdout, flush=True) if verbosity>= 1 else None

    ####################################################################################################################
    print(f"Filtering canonical edits to those with criteria: min_cells: {edit_site_min_cells}, "
          f"reverse_reads: {edit_site_rev_comp_filt}.", file=sys.stdout, flush=True) if verbosity>= 1 else None
    ####################################################################################################################
    keep_sites = np.array(canonical_edit_cell_counts)>=edit_site_min_cells
    print(f"{sum(keep_sites)} / {len(canonical_edit_sites)} kept after min cells criteria",
          file=sys.stdout, flush=True) if verbosity >= 1 else None
    if edit_site_rev_comp_filt:
        keep_sites = np.logical_and(keep_sites, canonical_edit_reversed)
        print(f"{sum(keep_sites)} / {len(canonical_edit_sites)} kept after reversed reads criteria",
              file=sys.stdout, flush=True) if verbosity >= 1 else None

    # Now will filter sites that intersect black list regions, if present.
    if type(blacklist_file)!=type(None):
        print("\nFiltering identified edit sites that intersect inputted blacklist regions.",
              file=sys.stdout, flush=True) if verbosity >= 1 else None
        bed_file_flag_edits(blacklist_file, canonical_edit_sites, keep_sites, False, # False indicate to blacklist
                            edit_dist, verbosity=verbosity)
        print(f"{sum(keep_sites)} / {len(canonical_edit_sites)} kept after remove edit sites overlapping blacklist regions\n",
              file=sys.stdout, flush=True) if verbosity >= 1 else None

    # WHITELIST adding back in of edit sites, i.e. we know these are edit sites and want to mark them as such, regardless
    # of the calling criteria applied.
    whitelist_canonical_edit_indices = []
    if type(whitelist_file)!=type(None):
        print("\nAdding edit sites that intersect inputted whitelist regions.",
              file=sys.stdout, flush=True) if verbosity>= 1 else None
        whitelist_canonical_edit_indices = bed_file_flag_edits(whitelist_file, canonical_edit_sites, keep_sites, True, # False indicate to blacklist
                            0, # Set this to 0 since this should be precise.
                                    verbosity=verbosity,
                            )
        print(f"{sum(keep_sites)} / {len(canonical_edit_sites)} kept after ADD edit sites overlapping whitelist regions\n",
              file=sys.stdout, flush=True) if verbosity>= 1 else None

    # Now also filtering based on stranded edit distance!!!
    #if edit_site_rev_comp_filt and type(stranded_edit_dist)!=type(None):
    keep_indices = np.where(keep_sites)[0]
    canonical_to_stranded_edit_dist = {}
    for index in keep_indices:
        edit_site = canonical_edit_sites[index]

        # With this logic, will still record information about the stranded edit site distance IF we are not using
        # the bi-directional insert criteria OR the edit site has been white-listed.
        if not canonical_edit_reversed[index] and \
            (index in whitelist_canonical_edit_indices or edit_site_rev_comp_filt==False): # If is a
            canonical_to_stranded_edit_dist[edit_site] = np.nan
            continue

        particular_edits = canonical_to_edits[edit_site]
        forward_edit_locs = np.array([edit_.ref_pos for edit_ in particular_edits if edit_.forward])
        reverse_edit_locs = np.array([edit_.ref_pos for edit_ in particular_edits if not edit_.forward])
        closest_forward = forward_edit_locs[ np.argmin( np.abs( forward_edit_locs-edit_site.ref_pos ) ) ]
        closest_reverse = reverse_edit_locs[ np.argmin( np.abs( reverse_edit_locs-edit_site.ref_pos ) ) ]

        # Will keep the whitelisted regions regardless of the stranded edit distance, OR keep all the called edit sites
        # if we are not considering the bi-directional inserts criteria.
        edit_stranded_dist = np.abs( closest_forward - closest_reverse )
        if type(stranded_edit_dist)!=type(None) and edit_stranded_dist >= stranded_edit_dist and \
                (index not in whitelist_canonical_edit_indices or edit_site_rev_comp_filt==False):
            keep_sites[index] = False # The distance between the forward and reverse edits is far, so likely not true edit.
        else:
            canonical_to_stranded_edit_dist[edit_site] = edit_stranded_dist

    if edit_site_rev_comp_filt and type(stranded_edit_dist)!=type(None):
        print(f"{sum(keep_sites)} / {len(canonical_edit_sites)} kept after remove edit sites with >= {stranded_edit_dist} stranded edit distance.\n",
              file=sys.stdout, flush=True) if verbosity >= 1 else None

    # Now need to filter all information for association with unsupported canonical edits...
    called_edit_sites = [edit for i, edit in enumerate(canonical_edit_sites) if keep_sites[i]]
    
    # # for debugging with CHD4 edit site as example..
    # chd4_edits = [edit for edit in called_edit_sites if str(edit.ref_pos).startswith('6602')]
    print(f"Finished edit site calling.", file=sys.stdout, flush=True) if verbosity>= 1 else None

    print("Filtering called t7 reads that do not fit a canonical edit site with minimum criteria cutoffs.",
              file=sys.stdout, flush=True) if verbosity >= 1 else None
    called_edit_sites_counts = np.array(canonical_edit_cell_counts)[keep_sites]
    edits_to_canonical_filtered = {edit_data: edit_site for edit_data, edit_site in edits_to_canonical.items()
                                    if edit_site in called_edit_sites}
    edit_counts_filtered = {}
    edit_bc_cell_umis_filtered = {}
    edit_reads_filtered = {}
    for edit_data in edits_to_canonical_filtered:
        edit_counts_filtered[ edit_data ] = edit_counts[edit_data]
        edit_bc_cell_umis_filtered[ edit_data ] = edit_bc_cell_umis[ edit_data ]
        edit_reads_filtered[ edit_data ] = edit_reads[ edit_data ]

    # Becomes important downstream for doing overlaps with other features, such as determining copy number of edit site
    # and also assigning these edit sites to genes via the GTF file.
    print(f"Writing edit sites to bed...", file=sys.stdout, flush=True) if verbosity >= 1 else None

    edit_sites_bed_file = open(out_path/"edit_sites.bed", 'w')
    
    for edit_site in called_edit_sites:
        edit_sites_bed_file.write(
            f"{edit_site.chrom}\t{edit_site.ref_pos - edit_dist}\t{edit_site.ref_pos + edit_dist}\t{edit_site.chrom}:{edit_site.ref_pos}\n")
    edit_sites_bed_file.close()

    ####################################################################################################################
    # Step 3: Get NON-Barcoded t7 edits;
    #   3.1: IF cell has known edit, mop up non-BC reads as those within X bp that are in-line with edit.
    ####################################################################################################################
    print("Counting Non-barcoded edits...", file=sys.stdout, flush=True) if verbosity >= 1 else None

    # Let's construct this so we put the specific edits at a particular site as a group!
    canonical_to_edits = defaultdict(list)
    [canonical_to_edits[edit_site].append( edit_data ) for edit_data, edit_site in edits_to_canonical_filtered.items()]

    # Previously using a list to lookup was making nonbarcoded super slow...
    canonical_to_edited_cells = defaultdict(set)
    for edit_data, edit_site in edits_to_canonical_filtered.items():
        canonical_to_edited_cells[edit_site].update(edit_bc_cell_umis_filtered[edit_data].keys())

    # Nested defaultdict
    cells_to_canonical_and_edits = defaultdict(lambda: defaultdict(list)) # Would set be better?
    # cells_to_canonical_and_edits = defaultdict(dict) # Nested dictionary; cell_barcodes -> edit_site -> particular edit
    
    for edit_site, edits in canonical_to_edits.items():
        for edit in edits:
            cell_barcodes_to_umis = edit_bc_cell_umis_filtered[edit]

            for cell_barcode in cell_barcodes_to_umis:
                
                cells_to_canonical_and_edits[cell_barcode][edit_site].append(edit)

    # Edit-site-centric method; subset reads from bam to those that are within Xbp of a canonical edit site, then proces
    t7_nonbarcoded_reads, nonbarcoded_umi, canonical_edit_no_bc_cell_umis = get_nonbarcoded_edits(
            bam_file,
            canonical_to_edits,
            canonical_to_edited_cells,
            cells_to_canonical_and_edits,
            edit_reads_filtered,
            verbosity=verbosity,
            dist=nonbc_edit_dist,
            )
    
    print(f"Processed Non-barcoded edits in {(timeit.default_timer()-start_bc)/60:.3f} minutes\n",
                                                                file=sys.stdout, flush=True) if verbosity >= 1 else None
    
    
    # CREATE T7 FILT BAMS EARLIER TO ASSIST WITH DOWNSTREAM PROCESSING STEPS
    ################################# Writing out the custom bam files #################################################
    
    # Create list of t7 barcoded reads
    t7_barcoded_reads_list = []
    for t7_reads in edit_reads_filtered.values():
        t7_barcoded_reads_list.extend(t7_reads)
    
    # Create Sets for t7 read categories
    t7_barcoded_reads_set = set(t7_barcoded_reads_list)
    t7_nonbarcoded_reads_set = set(t7_nonbarcoded_reads)
    
    # Combined t7 read list and set
    allt7_reads = t7_barcoded_reads_set.union(t7_nonbarcoded_reads_set)

    # Write ALL t7 reads to file
    t7_read_file = out_path / "t7_reads.txt"
    with open(t7_read_file, "w") as file:
        # Not sure if using the set and removing the dups would be better...
        file.write("\n".join([*t7_barcoded_reads_list, *t7_nonbarcoded_reads]))
    
    # Write barcoded t7 reads to file
    t7_barcoded_read_file = out_path / "t7_barcoded_reads.txt"
    with open(t7_barcoded_read_file, "w") as file:
        file.write("\n".join(t7_barcoded_reads_list))
    
    # Write non-barcoded t7 reads to file
    t7_non_barcoded_read_file = out_path / "t7_non-barcoded_reads.txt"
    with open(t7_non_barcoded_read_file, "w") as file:
        file.write("\n".join(t7_nonbarcoded_reads))
    
    
    # Create filtered and split BAM outputs!
    t7_bam = out_path / "t7_only.bam" # Both barcoded and non-barcoded called reads
    filt_bam = out_path / "t7_filt.bam" # All non-t7 reads, ideally just mRNA
    t7_barcode_bam = out_path / "t7_barcoded_only.bam" # All called t7 reads based on the barcode
    t7_non_barcode_bam = out_path / "t7_non-barcoded_only.bam" # All called t7 reads WITHOUT barcode

    # Split into t7 and non-t7 bam
    if mrna_count_mode == 'all': # filt_bam contains ALL reads for quantification of gene expression
        pysam.view("-N", str(t7_read_file), "-o", str(t7_bam), "-U", str(filt_bam), str(bam_file), catch_stdout=False)

    elif mrna_count_mode == 'polyT': # filt_bam contains only polyT reads for quantification of gene expression
        # Extracting all of the non-t7 reads
        all_filt_bam = out_path / "t7_filt.all-reads.bam" # All non-t7 reads, ideally just mRNA
        pysam.view("-N", str(t7_read_file), "-o", str(t7_bam), "-U", str(all_filt_bam), str(bam_file), catch_stdout=False)
        pysam.index(str(all_filt_bam), catch_stdout=False)

        # Isolating the polyT mRNA reads
        os.system(f"samtools view {all_filt_bam} | cut -f1 > {out_path}/all_mRNA_reads.txt")
        os.system(f"cat {out_path}/all_mRNA_reads.txt | grep __T__ > {out_path}/all_mRNA_polyT_reads.txt")

        # Creating the filt bam with only the polyT mRNA reads, downstream will result in only using these to
        # construct the count matrix.
        pysam.view("-N", f"{out_path}/all_mRNA_polyT_reads.txt", "-o", str(filt_bam), str(all_filt_bam),
                   catch_stdout=False)

    pysam.index(str(t7_bam), catch_stdout=False) # I think this should already be sorted
    pysam.index(str(filt_bam), catch_stdout=False)
    
    # Split t7 bam into barcoded and non barcoded
    pysam.view("-N", str(t7_barcoded_read_file), "-o", str(t7_barcode_bam), "-U", str(t7_non_barcode_bam), str(t7_bam), catch_stdout=False)
    pysam.index(str(t7_barcode_bam), catch_stdout=False)
    pysam.index(str(t7_non_barcode_bam), catch_stdout=False)
    
    # TODO should also write-out the state of the input arguments, just for record-keeping purposes (i.e. log inputs that generated the runs outputs)

    ####################################################################################################################
    # Step 4: T7 (barcode and/or non-barcoded) UMI counting per canonical edit-site
    ####################################################################################################################
    #### t7 barcoded umi counting
    # Need to collapse the particular-edit UMIs to the canonical-edit.
    canonical_edit_bc_cell_umis_filtered = defaultdict(lambda: defaultdict(set))
    for edit in edit_bc_cell_umis_filtered:
        edit_site = edits_to_canonical[edit]
        for cell_barcode in edit_bc_cell_umis_filtered[edit]:
            union_ = canonical_edit_bc_cell_umis_filtered[edit_site][cell_barcode].union( edit_bc_cell_umis_filtered[edit][cell_barcode] )
            canonical_edit_bc_cell_umis_filtered[edit_site][cell_barcode] = union_

    t7_barcoded_counts = get_t7_count_matrix(cell_barcodes_dict, canonical_to_edits, canonical_edit_bc_cell_umis_filtered)

    #### t7 non-barcoded umi counting
    t7_non_barcoded_counts = get_t7_count_matrix(cell_barcodes_dict, canonical_to_edits, canonical_edit_no_bc_cell_umis)

    ### all t7 counts.
    t7_all_counts = t7_barcoded_counts + t7_non_barcoded_counts

    ####################################################################################################################
    # Step 5: Call cell allelic edits for each canonical-edit-site.
    ####################################################################################################################
    # First create a mapping from edit_site to ploidy/copy number. If is a single number, is easy! But if is a copy
    # number bedGraph file, then need to figure out which location the edit-site overlaps with, to indicate the copy
    # number fo that particular genome location.
    # Just specifying single ploidy across genome, easy to handle
    print("Determining expected number of alleles for each edit site before counting allelic edits...",
              file=sys.stdout, flush=True) if verbosity >= 1 else None
    print(f"Inputted ploidy: {ploidy}", file=sys.stdout, flush=True) if verbosity >= 1 else None
    edit_sites_to_copy_number = {edit_site: ploidy for edit_site in called_edit_sites}

    if type(copy_number_variant_file)!=type(None) and len(called_edit_sites) > 0: # Trickier case, have specified a ploid file, so need to overlap with edit_site locations...
        print(f"Using inputted copy-number-variant file to update expected allele number at edit sites: {copy_number_variant_file}",
              file=sys.stdout, flush=True) if verbosity >= 1 else None
        copy_number_variants = pd.read_csv(copy_number_variant_file, sep='\t', header=None)
        copy_number_variants.columns = ['Chromosome', 'Start', 'End', 'Copy_number']
        copy_number_variants.index = copy_number_variants.apply(lambda x: ':'.join([str(x_) for x_ in x[0:3]]), axis=1)

        copy_number_locs = copy_number_variants.copy().iloc[:,0:3]
        copy_number_locs['Name'] = copy_number_locs.index
        copy_number_locs.index = list(range(copy_number_locs.shape[0]))
        copy_number_locs.columns = ['Chromosome', 'Start', 'End', 'Name']

        copy_number_ranges = pr.PyRanges( copy_number_locs )

        # Determining what edit_sites overlap the copy number variants, to update the ploidy number!
        edit_sites = pd.read_csv(out_path/"edit_sites.bed", sep='\t', header=None)
        edit_sites.iloc[:, 0] = [chrom.replace('hg38_', 'chr') for chrom in edit_sites.iloc[:, 0]]
        edit_sites.columns = ['Chromosome', 'Start', 'End', 'Name']

        # Extending out in either direction by the distance edits were collapsed to the edit site!
        edit_site_ranges = pr.PyRanges(edit_sites)  # .extend( edit_dist ) # ended up extending up there ^

        # Need to first get the edits that intersect genes
        closest_copy_variant_regions = edit_site_ranges.nearest(copy_number_ranges).df
        # Handling case where there are no copy number variants if we are processing split by chromosome.
        if closest_copy_variant_regions.shape[0] != 0:
            intersecting_copy_variant_regions = closest_copy_variant_regions.loc[closest_copy_variant_regions['Distance'].values==0,:]

            # Updating the ploidy number for each edit intersecting a copy-number-variant!
            for edit_site_name, copy_variant_loc in zip(intersecting_copy_variant_regions['Name'],
                                                        intersecting_copy_variant_regions['Name_b']):
                edit_site_ = EditSite(chrom=edit_site_name.split(':')[0], ref_pos=int(edit_site_name.split(':')[1]))
                edit_sites_to_copy_number[edit_site_] = int( copy_number_variants.loc[copy_variant_loc, 'Copy_number'] )
        else:
            print("No copy number variants found that match edit sites...",
              file=sys.stdout, flush=True) if verbosity >= 1 else None

    print("\nCalling allelic edits per cell per canonical-edit-site...",
              file=sys.stdout, flush=True) if verbosity >= 1 else None
    # Below considers the known copy-number variation across the genome if inputted, but otherwise assumes the
    # ploidy number of copy-number information is missing at the edit_site.
    cell_allelic_edits = np.zeros((len(cell_barcodes_list), len(called_edit_sites)), dtype=np.uint16)
    cells_edited = np.zeros((len(cell_barcodes_list), len(called_edit_sites)), dtype=np.uint16)

    for cell_barcode, edit_sites_to_edits in cells_to_canonical_and_edits.items():

        celli = cell_barcodes_dict[cell_barcode]

        for edit_site, edits in edit_sites_to_edits.items():

            edit_site_copy_number = edit_sites_to_copy_number[edit_site]
            
            # .index might be kinda slow, but seems correct
            edit_sitei = called_edit_sites.index( edit_site )

            cells_edited[celli, edit_sitei] += 1

            # Making sure is a unique set
            edit_set = list( set(edits) )

            if len(edit_set) == 1:
                # Only 1 edit here so only 1 allelic edit detected.
                cell_allelic_edits[celli, edit_sitei] = 1 # min( [cell_allelic_edits[celli, edit_sitei]+1, 2])
                continue

            # V3 of method, tries to get the longest t7 insertion sequences that are distinct from each other,
            # to avoid subsequences of a longer edit inflating allelic edit estimation.
            unique_edits = get_longest_edits(edit_set)

            n_alleles_called = len(unique_edits)

            # for debugging check...
            if n_alleles_called > edit_site_copy_number:
                print(f"WARNING: allelic edit number called at edit site: {edit_site} is {n_alleles_called} "
                      f"but should be maximum {edit_site_copy_number}",
                      file=sys.stdout, flush=True) if verbosity >= 2 else None
                print("Could be real, or could be due to cases not currently handled, "
                      "e.g. unaccounted seq-error, adapter barcodes on soft-clip, ...",
                      file=sys.stdout, flush=True) if verbosity >= 2 else None
                print("If below does not look to match these cases, may have non-diploid or CNV at edit site.",
                                              file=sys.stdout, flush=True) if verbosity >= 2 else None
                print(f"Cell barcode: {cell_barcode}", file=sys.stdout, flush=True) if verbosity >= 2 else None
                print(f"Unique allelic edits called: ", file=sys.stdout, flush=True) if verbosity >= 2 else None
                for edit_set_ in  unique_edits:
                    print(f"\t {edit_set_}", file=sys.stdout, flush=True) if verbosity >= 2 else None
                if constrain_allele_calls:
                    print(f"Will call as maximum {edit_site_copy_number} allelic edits anyhow, due to inputted constrain_allele_calls=True\n",
                    file=sys.stdout, flush=True) if verbosity >= 2 else None

            if constrain_allele_calls:
                # Here I make sure that our edit site allelic edit estimates does not exceed the inputted copy-number-variation
                # for the given genome location.
                cell_allelic_edits[celli, edit_sitei] = min([n_alleles_called, edit_site_copy_number])
            else: # If we don't constrain it, can let it over-estimate. Useful for downstream evaluation of how often this occurs.
                cell_allelic_edits[celli, edit_sitei] = n_alleles_called

    print("\nFinished allelic calling.\n", file=sys.stdout, flush=True) if verbosity >= 1 else None

    ####################################################################################################################
    # Step 6 **Optional**: If GTF provided on input; call allelic edits PER GENE
    ####################################################################################################################
    print(f"Calling allelic edits per gene, loading gtf..", file=sys.stdout, flush=True) if verbosity >= 1 else None

    gtf_cols = ["seqname", "start", "end", "gene_id", "gene_name"]
    with stdoutsuppress(): # To suppress the output of gtfparse...
        gtf = gtfparse.read_gtf(gtf_file,
                                features={"gene"},
                                usecols=gtf_cols,
                                result_type="pandas"
                                )

    gtf["gene_dup"] = gtf.duplicated(subset=['gene_name'], keep=False)
    
    gene_names = np.where(((gtf["gene_dup"] == True) | (gtf["gene_name"] == "")),
                          gtf["gene_id"], gtf["gene_name"]
                          )
    
    gtf["name"] = gene_names

    id_to_gene = dict(zip(gtf["gene_id"], gtf["name"]))

    genes_bed = gtf[['seqname', 'start', 'end', 'name']]
    genes_bed.columns = ['Chromosome', 'Start', 'End', 'Name']

    gene_ranges = pr.PyRanges( genes_bed )

    if len(called_edit_sites) == 0: # make blank versions of the variables so still runs through.
        genes_to_edits = defaultdict(list)
        genes_to_copy_number = defaultdict(int)
        edit_names = []
        genic_edits = []

        called_edit_site_names = [f'{edit_.chrom}:{edit_.ref_pos}' for edit_ in called_edit_sites]
        genes_and_edits = []
        cell_allelic_gene_edits = np.zeros((len(cell_barcodes_list), len(genes_and_edits)), dtype=np.uint16)

    else:
        edit_sites = pd.read_csv(out_path/"edit_sites.bed", sep='\t', header=None)
        edit_sites.iloc[:,0] = [chrom.replace('hg38_', '') for chrom in edit_sites.iloc[:,0]]
        edit_sites.columns = ['Chromosome', 'Start', 'End', 'Name']

        # Extending out in either direction by the distance edits were collapsed to the edit site!
        edit_site_ranges = pr.PyRanges( edit_sites ) #.extend( edit_dist ) # ended up extending up there ^

        # Need to first get the edits that intersect genes
        intersection = edit_site_ranges.intersect( gene_ranges )
        if len(intersection) > 0:
            genic_edits = np.unique( intersection.df['Name'] )

            # Getting the edits that intersect with the genes
            genes_to_edits = defaultdict(list)
            genes_to_copy_number = defaultdict(int)
            ### Changed this from the orignial edit_sites, since within the edit_site_ranges it re-arranged the order to
            ### reflect the order within the chromosomes, so if processing a bam with reads ACROSS chromosomes this will
            ### result in a different order!!
            edit_names = list( edit_site_ranges.df['Name'] )
            for i, edit_name in enumerate(genic_edits):

                n_intersects = sum( edit_site_ranges.intersect( gene_ranges ).df['Name'].values==edit_name )
                edit_entry = edit_names.index( edit_name )

                edit_site = called_edit_sites[ edit_entry ]
                edit_site_copy_number = edit_sites_to_copy_number[edit_site]

                edit_loc = pr.PyRanges( edit_site_ranges.df.iloc[edit_entry:edit_entry+1,:] )
                edit_genes = edit_loc.k_nearest(gene_ranges, k=n_intersects)
                if edit_name != edit_site_ranges.df.iloc[edit_entry,:]['Name']:
                    print("ERROR indexing problem, pulled edit entry does not match edit entry: ")
                    print(edit_name)
                    print(edit_site_ranges.df.iloc[edit_entry,:]['Name'])

                if np.any(edit_genes.df['Distance'].values != 0):
                    err_bool = edit_genes.df['Distance'].values != 0
                    err_dists = list(edit_genes.df['Distance'].values[err_bool])
                    err_genes = list(edit_genes.df['Name_b'].values[err_bool])

                    print("ERROR have somehow selected genes that are non-intersecting...", file=sys.stdout, flush=True)
                    print(f"This called edit: {edit_site}")
                    print(f"These genes: {err_genes}")
                    print(f"At these distances: {err_dists}")

                for gene in edit_genes.df['Name_b']: # Determining gene-copy-number as the max of the edit-site copy number..
                    genes_to_edits[gene].append(edit_name)
                    genes_to_copy_number[gene] = max([genes_to_copy_number[gene], edit_site_copy_number]) # Saves re-overlaps

        else: # No genic overlaps, so just need to initialise with blanks.
            genes_to_edits = defaultdict(list)
            genes_to_copy_number = defaultdict(int)
            edit_names = []
            genic_edits = []

            called_edit_site_names = [f'{edit_.chrom}:{edit_.ref_pos}' for edit_ in called_edit_sites]
            genes_and_edits = []
            cell_allelic_gene_edits = np.zeros((len(cell_barcodes_list), len(genes_and_edits)), dtype=np.uint16)

        # Now need to go through, and construct new allelic edit matrix that adds the allelic edicts
        #  called at the per canonical-edit-site level, and put collapse these to the gene level !!!!
        # For every edit that overlaps a gene, we will exclude these from our final list, and replace them with the
        # gene level allelic edit calls!
        called_edit_site_names = [f'{edit_.chrom}:{edit_.ref_pos}' for edit_ in called_edit_sites]
        genes_and_edits = list(genes_to_edits.keys()) + [edit_name for edit_name in called_edit_site_names
                                                                                            if edit_name not in genic_edits]
        cell_allelic_gene_edits = np.zeros((len(cell_barcodes_list), len(genes_and_edits)), dtype=np.uint16)
        for loci, gene_or_edit in enumerate(genes_and_edits):
            if gene_or_edit in genes_to_edits: # Is a gene, need to collapse it's edits to make a gene-level allelic call.
                gene_edit_indices = [called_edit_site_names.index(genic_edit)
                                     for genic_edit in genes_to_edits[ gene_or_edit ]]
                # Debug check
                #print( np.all(np.array(called_edit_site_names)[gene_edit_indices] == genes_to_edits[ gene_or_edit ]) )
                # returned True for all cases during testing!

                gene_copy_number = genes_to_copy_number[ gene_or_edit ]

                # Accounting for inputted ploidy number OR CNV file to cap the number of called alleles.
                gene_allelic_edits = cell_allelic_edits[:, gene_edit_indices].sum(axis=1)
                gene_allelic_edits[gene_allelic_edits > gene_copy_number] = gene_copy_number

                cell_allelic_gene_edits[:, loci] = gene_allelic_edits

            else: # Is an edit site that does not overlap the gene, so no need to collapse !
                cell_allelic_gene_edits[:, loci] = cell_allelic_edits[:, called_edit_site_names.index(gene_or_edit)]

    # Diagnostic check, still calling same number of cells edited for each edit site?
    if verbosity >= 2:
        nongenic_edit_indices_gene_alleles = [loci for loci, gene_or_edit in enumerate(genes_and_edits)
                                              if gene_or_edit in called_edit_site_names]
        nongenic_edit_indices_edit_alleles = [loci for loci, edit_ in enumerate(called_edit_site_names)
                                              if edit_ in genes_and_edits]
        correct_cell_edit_number = np.all(cell_allelic_gene_edits[:,nongenic_edit_indices_gene_alleles] ==
                                            cell_allelic_edits[:,nongenic_edit_indices_edit_alleles] )
        print(f"MAINTINED CORRECT EDITED CELL CALLS (full check)? {correct_cell_edit_number}", file=sys.stdout, flush=True)
        # Returned True during check

        # Total number of cells called as edited retained?
        correct_cell_edit_number = (cell_allelic_gene_edits.sum(axis=1)>0).sum() == (cell_allelic_edits.sum(axis=1)>0).sum()
        print(f"MAINTINED CORRECT EDITED CELL CALLS (SUM method)? {correct_cell_edit_number}", file=sys.stdout, flush=True)
        # Returned True during check!!!

    cell_allelic_edits = pd.DataFrame(cell_allelic_edits, index=cell_barcodes_list, columns=called_edit_site_names)
    cell_allelic_gene_edits = pd.DataFrame(cell_allelic_gene_edits, index=cell_barcodes_list, columns=genes_and_edits)

    print("Finished gene level allelic edit-calling.", file=sys.stdout, flush=True) if verbosity >= 1 else None

    ####################################################################################################################
    # Step 7 **Optional**: If GTF provided on input; implement UMI counting per gene
    # (mRNA-counting)
    ####################################################################################################################
    print("Performing mRNA counting, removing the confounding t7 reads.. Most time consuming step.",
              file=sys.stdout, flush=True) if verbosity >= 1 else None

    # Run using prefiltered non-t7 bam
    cell_by_gene_umi_counts = bam_count_gene_umis(filt_bam, cell_barcodes_dict, gene_names, id_to_gene,
                                                    n_cpus=n_cpus, verbose=(verbosity >= 1), chunk_size_mb=chunk_size_mb,
                                 )

    if uncorrected_gene_count and len(allt7_reads)>0: # only makes sense if there was actual edit sites.
        print("Performing mRNA counting, INCLUDING the confounding t7 roudns on user request "
              "(via 'uncorrect_gene_count' input).",
              file=sys.stdout, flush=True) if verbosity>= 1 else None
        cell_by_gene_umi_counts_t7_confounded = bam_count_gene_umis(bam_file, cell_barcodes_dict, gene_names, id_to_gene,
                                                                    n_cpus=n_cpus, verbose=(verbosity >= 1),
                                                                    chunk_size_mb=chunk_size_mb
                                                                    )

    elif uncorrected_gene_count:
        cell_by_gene_umi_counts_t7_confounded = cell_by_gene_umi_counts

    if uncorrected_gene_count: # Will still write two outputs, just for consistency (even if the same thing!)
        # gene counts (mRNA) counts, confounded by t7 reads.
        cell_by_gene_umi_counts_t7_confounded.to_parquet(out_path/'cell_gene_mrna_counts.t7-uncorrected.parquet.gz',
                                                         compression="gzip")

    # Will also do UMI counting on our edit-sites, since hypothetically we have corrected for the t7 reads so
    # any remaining t7 reads indicating false-negative calls. Useful diagnostic to evaluate sensitivity downstream.
    # edit_site_ranges # Will use these ranges to pull out reads overlapping edit sites.
    # TODO NOTE if an edit site overlaps a gene, the UMIs for the edit site will be double-counted with the UMIs counted above!
    #  Am not worried about this, is just mean to be a diagnostic, where will calculate an enrichment score to see
    #  if the edit site appears to be have more reads than the rest of the gene body (indicating false-negatives),
    #  but also for non-genic edit sites any left-over will indicate definite false-negatives!
    print("Performing UMI counting at edit sites, after t7 read correction. Used for downstream diagnostic checks.",
              file=sys.stdout, flush=True) if verbosity >= 1 else None
    
    # Run using prefiltered non-t7 bam
    # after_t7_edits_to_bc_to_umis = defaultdict(dict) # Will also count the reads remaining per edit site !
    after_t7_edits_to_bc_to_umis = defaultdict(lambda: defaultdict(set))
    with AlignmentFile(filt_bam, "rb") as bam:
        for edit_site in called_edit_sites:

            edit_chr = edit_site.chrom
            edit_pos = edit_site.ref_pos
            edit_window_start = max([edit_pos - edit_dist, 0])  # Accounting for edge of chromosome.
            edit_window_end = edit_pos + edit_dist  # pysam is OK with trying to fetch greater than chromosome length.

            # This is most likely the culprit of the really long runtime
            for read in bam.fetch(edit_chr, edit_window_start, edit_window_end):

                ### Only need to filter if is a edited cell.
                cell_barcode = read.get_tag('CB')
                
                # Using filt bam means I shouldnt have to check against allt7
                # I should also try prefilt using barcode whitelist
                if (cell_barcode not in cell_barcodes):
                    continue

                read_edit_dist = read.pos - edit_site.ref_pos
                
                # I SHOULD FIX THIS TO NOT BE NESTED AND TO ALSO USE DEFAULT DICT TO AVOID INIT CHECK
                if abs(read_edit_dist) <= edit_dist:
                    after_t7_edits_to_bc_to_umis[edit_site][cell_barcode].add(read.get_tag('pN'))

    # Now doing the read counting for this.
    after_t7_edits_umi_counts = np.zeros((len(cell_barcodes_list), len(called_edit_sites)), dtype=np.uint16)
    for edit_index, edit_site in enumerate(called_edit_sites):
        after_t7_edits_umi_counts[:, edit_index] = get_cell_counts_from_umi_dict(
                                                            after_t7_edits_to_bc_to_umis[edit_site], cell_barcodes_dict)

    after_t7_edits_umi_counts = pd.DataFrame(after_t7_edits_umi_counts, index=cell_barcodes_list,
                                             columns=called_edit_site_names)


    print("Finished all UMI counting.", file=sys.stdout, flush=True) if verbosity >= 1 else None

    ####################################################################################################################
    # Step 8: Writing outputs.
    ####################################################################################################################
    print(f"Final step of writing all outputs to {outdir}.", file=sys.stdout, flush=True) if verbosity >= 1 else None
    # Going to Write out two main outputs! T7 Edit Info and Filtered Bams
    # out_path.mkdir(parents=True, exist_ok=True)

    ################################# Writing out the UMI counts at each edit site #################################
    # DATAFRAMES SEEM LIKE A BAD IDEA, SHOULD PROBS CHANGE TO SPARSE MATRICES + OUTPUT INTS NOT FLOATS
    # Anndata / H5 format may be a good contender for I/O speed and memory, but might add complexity
    
    # Cell X canonical-edit-site allelic dosage count matrices
    cell_allelic_edits.to_parquet(out_path/'cell_allelic_dosage.canonical-edit-sites.parquet.gz', compression="gzip")
    cell_allelic_gene_edits.to_parquet(out_path/'cell_allelic_dosage.canonical-edit-sites.gene-collapsed.parquet.gz',
                                       compression="gzip")

    # T7 barcoded counts
    t7_barcoded_counts.to_parquet(out_path/'t7_barcoded_counts.parquet.gz', compression="gzip")

    # T7 non-barcoded cell counts
    t7_non_barcoded_counts.to_parquet(out_path/'t7_nonbarcoded_counts.parquet.gz', compression="gzip")

    # T7 barcoded and non-barcoded counts
    t7_all_counts.to_parquet(out_path/'t7_all_counts.parquet.gz', compression="gzip")

    # gene counts (mRNA) counts
    cell_by_gene_umi_counts.to_parquet(out_path/'cell_gene_mrna_counts.parquet.gz', compression="gzip")

    # count of reads that intersect the edit site counts, AFTER removing all of the potential T7 reads.
    # Not really relevant for new version.
    after_t7_edits_umi_counts.to_parquet(out_path/'cell_canonical-edit-site_counts.t7-removed.parquet.gz',
                                         compression="gzip")

    ################################# Writing out bed file of the canonical edit sites #################################
    # Already wrote out the bed file above, but will include metadata about the edit site, particularly if overlaps a gene...

    # Should probably use pandas/polars/csvwriter to avoid this manual writing situation
    edit_site_info = open(out_path/'edit_site_info.txt', 'w')
    edit_site_info.write(f'name\tchr\tpos\tstart_window\tend_window\tintersecting_genes\tcopy-number\tn_cells_edited\tstranded_edit_dist\n')
    for i in range(len(called_edit_sites)):

        edit_bed = edit_sites.values[i, :]
        edit_info = [edit_bed[-1], edit_bed[0], edit_bed[-1].split(':')[-1], str(edit_bed[1]), str(edit_bed[2])]
        edit_info.append( [] )

        for gene, edit_names in genes_to_edits.items():
            if edit_info[0] in edit_names:
                edit_info[-1].append( gene )

        edit_info[-1] = ','.join( edit_info[-1] )

        # Now also adding the copy number for the edit, in-case overlaps a copy number variant.
        edit_info.append( str(edit_sites_to_copy_number[ EditSite(chrom=edit_info[0].split(':')[0],
                                                              ref_pos=int(edit_info[0].split(':')[1])) ]) )

        edit_info.append( str(len(canonical_to_edited_cells[ called_edit_sites[i] ])) ) # Adding number of edited cells

        edit_info.append( str(canonical_to_stranded_edit_dist[ called_edit_sites[i] ]) ) # Adding the stranded edit distance

        edit_info_str = "\t".join(edit_info)
        edit_site_info.write(f'{edit_info_str }\n')
    edit_site_info.close()

    ################################# Writing out custom TSV file with per cell edits ##################################
    
    # First Create a TSV file for the data
    edit_tsv = out_path / "t7_barcode_edits.tsv"

    # TSV column schema
    schema = {"barcode": pl.Categorical, "edit_name": pl.Categorical, "chrom": pl.Categorical,
              "ref_pos": pl.UInt32, "ref_seq": pl.Categorical,
              "alt_seq": pl.Utf8, "forward": pl.Boolean,
              "canonical_edit_site_name": pl.Categorical,
              "kmer_matches": pl.Utf8,
              "t7_barcoded_umis": pl.Utf8,
              't7_nonbarcoded_umis': pl.Utf8}

    data_list = []
    
    # Fix to not be triple nested loop if this gets too slow
    for barcode, edit_site_to_edits in cells_to_canonical_and_edits.items():

        for edit_site, edits in edit_site_to_edits.items():

            for edit in edits:

                bc_umis = edit_bc_cell_umis_filtered[edit][barcode]

                data_list.append(
                    [
                        barcode, ':'.join(np.array(edit[:-1]).astype(str)),
                        *edit[:-1],
                        f'{edit_site.chrom}:{edit_site.ref_pos}',
                        '-'.join(edit[-1]),
                        '-'.join(bc_umis),
                        '-'.join(nonbarcoded_umi[barcode])
                    ]
                )

    # Convert into dataframe - Polars is faster than pandas when performance matters...
    df = pl.DataFrame(data_list, schema=schema)
    df.write_csv(edit_tsv, separator="\t") # Maybe including reads could also be useful

    print(f"Wrote T7 Edit data and filtered alignments to {outdir}!", file=sys.stdout,
                                                                                 flush=True) if verbosity >= 1 else None
```
