## Additional Files for "Joint single-cell capture of Cas9 edits and transcriptomes reveals on- and off-target effects on gene expression": helpers.py

```
""" Helper functions for processing the t7 sequencing
"""

import math
import sys
import re
import itertools # Used to do pairwise comparisons of umis for similarity

from collections import defaultdict

import numpy as np
import pandas as pd

from pysam.libcalignmentfile import AlignmentFile

import warnings
from Bio import BiopythonDeprecationWarning

# numba-0.61.2 when developing this on mac.
from numba import jit
from numba import prange
from numba.typed import List

# Important for maintaining good memory performance
#from scipy.sparse import dok_array as sparse_array # Generalise so can test different implementations easily.
from scipy.sparse import csr_array

# Suppress only BiopythonDeprecationWarning
warnings.simplefilter("ignore", BiopythonDeprecationWarning)
from Bio import pairwise2  # pip install biopython

def get_t7_count_matrix(cell_barcodes_dict, canonical_to_edits, canonical_edit_bc_cell_umis_filtered):
    """ Performs the UMI counting per edit site, either for the barcoded or non-barcoded reads!
    """

    t7_barcoded_counts = np.zeros((len(cell_barcodes_dict), len(canonical_to_edits)),
                                  dtype=np.uint32)

    #edit_bc_cell_umis_filtered # edit to cell barcodes to umis
    #canonical_to_edits # canonical-edits to edits
    
    # REQUIRE DICT SINCE IT MAINTAINS ORDER
    # cell_barcodes_dict = {key: value for value, key in enumerate(cell_barcodes_list)}
    
    edit_site_names = []
    for edit_site_index, (edit_site, edits) in enumerate(canonical_to_edits.items()):

        edit_site_names.append( f'{edit_site.chrom}:{edit_site.ref_pos}' )

        # for edit in edits:
        #
        # NOTE edit_bc_cell_umis_filtered[edit] MUST NOW BE A DICTIONARY THAT STORES SETS AS VALUES
        t7_barcoded_counts[:, edit_site_index] += get_cell_counts_from_umi_dict(
                                                    canonical_edit_bc_cell_umis_filtered[edit_site], cell_barcodes_dict)

    t7_barcoded_counts = pd.DataFrame(t7_barcoded_counts, index=list(cell_barcodes_dict.keys()), columns=edit_site_names)

    return t7_barcoded_counts

def get_cell_counts_from_umi_dict(cell_bc_to_umis, cell_barcodes_dict):
    """Gets the counts across cells for a given barcode to umi dict.

    VERY slow. Needs to be optimized for run speed because right now this eats most of the processing time.
    """
    if False:
        return cell_umi_counts_original(cell_bc_to_umis, cell_barcodes_dict)
    else: # Trying a speed optimized version.

        # First need to filter the list of cell barcodes to those in the white list...
        cell_bc_to_umis = {cell_bc: umis for cell_bc, umis in cell_bc_to_umis.items() if cell_bc in cell_barcodes_dict}

        if len(cell_bc_to_umis) == 0:
            return np.zeros(( len(cell_barcodes_dict) ), dtype=np.uint32)

        # Now making more Numba friendly data structures.
        # Re-processing these to make it simpler and also compilable with numba - namely will represent the UMI data as
        # 2 numpy arrays and one list:
        cell_bc_indexes = np.array([cell_barcodes_dict[cell_barcode] for cell_barcode in cell_bc_to_umis], dtype=np.uint32)
        umi_len = len( list( list(cell_bc_to_umis.values())[0] )[0] )
        cell_umis = [np.array(list(umis), dtype=f'<U{umi_len}') for umis in cell_bc_to_umis.values()]

        return cell_umi_counts_FAST(len(cell_barcodes_dict), cell_bc_indexes, cell_umis)

@jit(nopython=True)
def cell_umi_counts_FAST(total_cells, cell_bc_indexes, cell_umis):
    """ Counts the cell UMIs FAST
    """

    cell_umi_counts = np.zeros((total_cells), dtype=np.uint32)

    for i in range(len(cell_bc_indexes)):

        cell_index = cell_bc_indexes[i]
        umi_array = cell_umis[i]

        num_unique_umi = len(umi_array)

        if num_unique_umi == 1:
            cell_umi_counts[cell_index] += 1
        elif num_unique_umi == 2:
            umi_1, umi_2 = umi_array

            if within_single_mismatch_int(umi_1, umi_2):
                cell_umi_counts[cell_index] += 1
            else:
                cell_umi_counts[cell_index] += 2
        else:

            cell_umi_counts[cell_index] = deduplicated_umi_count_FAST( umi_array )

    return cell_umi_counts

@jit(nopython=True)
def deduplicated_umi_count_FAST( umi_array ):
    """ Gets the deduplicated UMI count FAST!
    """

    # Now let's do this, get adjacency matrix between the umis first, and then a quick algorithm to get the disconnected
    # sets.
    adjacency_matrix = get_adjacency_matrix(umi_array)

    # If none are neighbours, just return the length!
    umi_neighbors = adjacency_matrix.sum(axis=1)
    if np.all(umi_neighbors==0): # They are all different!
        return len(umi_array)

    elif sum(umi_neighbors > 0)==2: #Only 2 connect UMIs, so only solution would be to collapse these.
        return len(umi_array) - 1

    else: # More complicated, many alternatives for the remaining UMIs.
        umi_count = sum(umi_neighbors==0) # These are our base disconnected set.

        # Now we can use a depth first search to get the dis-connected components.
        remaining_umi_indices = np.where( umi_neighbors>0 )[0]
        n_connected_sets = 0 # Number of iterations indicates the number of unique sets of connected UMIs.
        while len(remaining_umi_indices) > 0: # While there are remaining indices

            connected_umi_indices = [remaining_umi_indices[0]]
            depth_first_search(remaining_umi_indices[0], connected_umi_indices, adjacency_matrix)

            n_connected_sets += 1

            # Implemented this way to prevent typechange errors with numba.
            keep_indices = np.array([index_ for index_ in range(len(remaining_umi_indices))
                                     if remaining_umi_indices[index_] not in connected_umi_indices], dtype=np.int64)
            remaining_umi_indices = remaining_umi_indices[keep_indices]

        # umi_count is all of the indices with no neighbours, while n_connected_sets is the set of connected umis
        # that are NOT connected to each other, so they are called as PCR duplicates.
        return umi_count + n_connected_sets

@jit(nopython=True)
def get_adjacency_matrix(umi_array):
    """Gets adjacency matrix between the UMIs, filled with 1 if the UMIs are within 1 edit distance from each other.
    """
    adjacency_matrix = np.zeros((len(umi_array), len(umi_array)), dtype=np.int64)

    for umi_i in range(len(umi_array)):

        umi_1 = umi_array[umi_i]

        for umi_j in range(umi_i + 1, len(umi_array)):
            umi_2 = umi_array[umi_j]

            adjacency_matrix[umi_i, umi_j] = within_single_mismatch_int(umi_1, umi_2)
            adjacency_matrix[umi_j, umi_i] = adjacency_matrix[umi_i, umi_j]  # Make it square matrix.

    return adjacency_matrix

@jit(nopython=True)
def within_single_mismatch_int(seq1, seq2):
    diff_score = 0
    #for i1, i2 in zip(seq1, seq2):
    for i_ in range(len(seq1)):
        i1, i2 = seq1[i_], seq2[i_]

        if i1 != i2:
            if diff_score > 0:
                return 0
            diff_score += 1
    return 1

@jit(nopython=True)
def depth_first_search(current_index, current_connected_indices, adjacency_matrix):
    """Adds all indexes to current_connected_indices from the inputted current_index"""

    neighbour_indices = np.where(adjacency_matrix[current_index, :] > 0)[0]
    for neighbour_index in neighbour_indices:
        if neighbour_index not in current_connected_indices:
            # Add it to the set of connections, then perform depth-search on this neighbour.
            current_connected_indices.append( neighbour_index )
            depth_first_search(neighbour_index, current_connected_indices, adjacency_matrix)
        else:
            continue # Continue to other neighbours, this one already accounted for.

def cell_umi_counts_original(cell_bc_to_umis, cell_barcodes_dict):
    """Counts the cell UMIs using the original and SLOW method"""
    cell_umi_counts = np.zeros((len(cell_barcodes_dict)), dtype=np.uint32)
    for cell_barcode in cell_bc_to_umis:

        cell_index = cell_barcodes_dict[cell_barcode]

        umi_set = cell_bc_to_umis[cell_barcode]

        # umi_set = set(umis) # stored as set automatically now
        num_unique_umi = len(umi_set)

        if num_unique_umi == 1:
            cell_umi_counts[cell_index] += 1
        elif num_unique_umi == 2:
            umi_1, umi_2 = umi_set

            if within_single_mismatch(umi_1, umi_2):
                cell_umi_counts[cell_index] += 1
            else:
                cell_umi_counts[cell_index] += 2
        else:

            unique_umi_sets = deduplicate_umis(umi_set)

            cell_umi_counts[cell_index] += len(unique_umi_sets)

    return cell_umi_counts

def deduplicate_umis(umi_set):
    """ De-duplicates the UMIs for a gene in a single cell
    """
    # Kinda complex, fix if it becomes an issue later
    umis_to_match_umis = {umi: {umi} for umi in umi_set}

    for umi_1, umi_2 in itertools.combinations(umi_set, 2):

        if within_single_mismatch(umi_1, umi_2):
            umis_to_match_umis[umi_1] = umis_to_match_umis[umi_1].union(umis_to_match_umis[umi_2])
            umis_to_match_umis[umi_2] = umis_to_match_umis[umi_2].union(umis_to_match_umis[umi_1])

            # syncing!
            # Need to also pull in the umis with edit distance 1 that are also edit distance 1 from
            # this umi!
            for umi in umis_to_match_umis[umi_1]:
                umis_to_match_umis[umi_1] = umis_to_match_umis[umi_1].union(umis_to_match_umis[umi])
                umis_to_match_umis[umi_2] = umis_to_match_umis[umi_2].union(umis_to_match_umis[umi])

            # Now need to update all the other umis with these umis edit dist 1
            for umi in umis_to_match_umis[umi_1]:
                umis_to_match_umis[umi] = umis_to_match_umis[umi].union(umis_to_match_umis[umi_1])

    unique_umi_sets = []
    for umi_set_ in umis_to_match_umis.values():
        if umi_set_ not in unique_umi_sets:
            unique_umi_sets.append(umi_set_)

    return unique_umi_sets

def within_single_mismatch(seq1, seq2):
    diff_score = 0
    for i1, i2 in zip(seq1, seq2):
        if i1 != i2:
            if diff_score > 0:
                return False
            diff_score+=1
    return True

def bam_count_gene_umis(bam_file, cell_barcodes_dict, gene_names, id_to_genes, n_cpus=1, verbose=True,
                        chunk_size_mb=15, # Measured in mb
                        ):
    """ Count all the gene UMIs across the bam file!
    """
    if n_cpus == 1:
        cell_by_gene_umi_counts_SPARSE = bam_count_gene_umis_contig(bam_file, cell_barcodes_dict,
                                                                    gene_names, id_to_genes, verbose,None)
        # Convert from sparse format to dense for output.
        cell_by_gene_umi_counts = pd.DataFrame(cell_by_gene_umi_counts_SPARSE.toarray(),
                                               index=list(cell_barcodes_dict.keys()),
                                               columns=list(gene_names))

        return cell_by_gene_umi_counts

    else: #### Parallel processing
        # Determining the contig names so can parallelize across contigs.
        bam_ = AlignmentFile(bam_file, "rb")
        chrom_names = [contig['SN'] for contig in bam_.header['SQ']]
        chrom_sizes = np.array([contig['LN'] for contig in bam_.header['SQ']])
        bam_.close()

        # Let's set the chunk size to be 15Mbp, which is about 1/16th of chromosome 1.
        chunk_size = chunk_size_mb * (10**6)

        # Using this to determine a set of loci to be counted independently in parallel:
        genome_chunks = []
        for chr_, size_ in zip(chrom_names, chrom_sizes):
            if size_ < chunk_size:
                genome_chunks.append( [chr_] )
            else:
                start_ = 0
                for end_ in range(chunk_size, size_, chunk_size):
                    genome_chunks.append( (chr_, start_, end_) )
                    start_ = end_ # re-set the start point!

                if end_ < size_: # Truncated end of the contig
                    genome_chunks.append((chr_, end_, size_))

        #### Processing in parallel
        from concurrent.futures import ProcessPoolExecutor, as_completed

        from functools import partial
        partial_func = partial(bam_count_gene_umis_contig, bam_file, cell_barcodes_dict,
                               gene_names, id_to_genes, verbose)

        with ProcessPoolExecutor(max_workers=n_cpus) as executor:
            futures = {executor.submit(partial_func, chunk): chunk for chunk in genome_chunks}
            contig_counts = []
            for future in as_completed(futures):
                try:
                    result = future.result()  # will re-raise exceptions from worker
                    contig_counts.append( result )
                except Exception as e:
                    chunk = futures[future]
                    raise Exception(f"Error in genome chunk {chunk}: {e}")

        # OLD version, had problem of SILENTLY erroring out, leaving certain chromosomes without counts!
        # with ProcessPoolExecutor(max_workers=n_cpus) as executor:
        #     contig_counts = list(executor.map(partial_func, genome_chunks))

        # DENSE version
        # cell_by_gene_umi_counts = contig_counts[0].values
        # for counts_ in contig_counts[1:]:
        #     cell_by_gene_umi_counts += counts_.values

        # SPARSE version, add these together FIRST, more memory efficient.
        cell_by_gene_umi_counts = contig_counts[0] # sparse csr array.
        for counts_ in contig_counts[1:]:
            cell_by_gene_umi_counts += counts_

        # Now we have just one array, make it dense, and return as a dataframe.
        cell_by_gene_umi_counts = pd.DataFrame(cell_by_gene_umi_counts.toarray(),
                                               index=list(cell_barcodes_dict.keys()),
                                               columns=list(gene_names))

        return cell_by_gene_umi_counts

def bam_count_gene_umis_contig(bam_file, cell_barcodes_dict, gene_names, id_to_genes, verbose, contig,
                        ):
    """Get's gene UMI counts per cell, returning as sparse array of counts! Sparse is important to prevent memory issues
    from returning a ton of cells X genes sized arrays from each thread.
    """

    #### Testing raise error!!! Before it would silently error.
    # if 'hg38_1' in contig:
    #     raise Exception("Successfully raise exeption from a thread!!")

    gene_set = set(gene_names)

    if verbose:
        print(f"Counting UMIs for contig: {contig}", file=sys.stdout, flush=True)

    # NOT numba friendly
    genes_to_bcs_to_umis = defaultdict(lambda: defaultdict(set))

    # If the bam has not reads for a given region, just doesn't iterate
    with AlignmentFile(bam_file, "rb") as bam:
        nreads = 0
        if type(contig)!=type(None):
            iterator_ = bam.fetch(*contig)
        else:
            iterator_ = bam

        for i, read in enumerate( iterator_ ):
            nreads += 1

            cell_barcode = read.get_tag('CB')
            
            # Using filt bam means I shouldnt have to check against allt7
            if (cell_barcode not in cell_barcodes_dict):
                continue

            # NEW logic, more robust to how the reads are tagged.
            gene_id_tag = read.get_tag('GX')

            if (gene_id_tag != '') and (gene_id_tag in id_to_genes):
                gene_or_id = id_to_genes[ gene_id_tag ]
            elif gene_id_tag != '': # Is tagged with something unexpected here....
                for file_ in [sys.stdout, sys.stderr]:
                    print(f"WARNING: found read with unexpected GX tag given INPUT GTF file: {gene_id_tag}",
                                                                                                 file=file_, flush=True)
                    print(f"\tCHECK used the same GTF file as the split-pipe processing", file=file_, flush=True)
            else:
                continue

            # OLD logic, this was sensitive to different versions of split-pipe.
            # gene_name_tag = read.get_tag('GN')
            # gene_id_tag = read.get_tag('GX')
            #
            # if (gene_name_tag != '') and (gene_name_tag in gene_set):
            #     gene_or_id = gene_name_tag
            # elif (gene_id_tag != '') and (gene_id_tag in gene_set):
            #     gene_or_id = gene_id_tag
            # else:
            #     continue

            # Should be cleaner and faster
            genes_to_bcs_to_umis[gene_or_id][cell_barcode].add( read.get_tag('pN') )

    if len(genes_to_bcs_to_umis) == 0: # No genic reads found on this contig, so just return an empty count matrix.

        # Sparse version
        return csr_array((len(cell_barcodes_dict), len(gene_names)), dtype=np.uint32)

        # OLD version
        # return pd.DataFrame(np.zeros((len(cell_barcodes_dict), len(gene_names)), dtype=np.uint16),
        #              index=list(cell_barcodes_dict.keys()),
        #              columns=list(gene_names))

    # Now converting this to a more Numba friendly format
    gene_names_list = list(gene_names)
    gene_indices = []
    gene_cell_indices = []
    gene_cell_umis = []
    gene_names_extracted = []
    for genei in range(len(gene_names)):

        gene_name = gene_names_list[genei]

        if gene_name in genes_to_bcs_to_umis:
            gene_names_extracted.append( gene_name )
            gene_indices.append(genei)

            cell_barcodes = list( genes_to_bcs_to_umis[gene_name].keys() )

            gene_cell_indices.append( np.array([cell_barcodes_dict[cell_barcode] for cell_barcode in cell_barcodes],
                                                dtype=np.int64)
                                       )

            umi_len = len( list( genes_to_bcs_to_umis[gene_name][cell_barcodes[0]])[0] )

            gene_cell_umis.append( List([np.array(list(genes_to_bcs_to_umis[gene_name][cell_barcode]), dtype=f"<U{umi_len}")
                                        for cell_barcode in cell_barcodes])
                                 )

    gene_indices = np.array(gene_indices, dtype=np.int64)

    # Performing the counting with Numba speed-up
    cell_by_gene_umi_counts_SPARSE_indices = get_cell_by_gene_umi_counts(len(cell_barcodes_dict), #len(gene_names),
                                                                            gene_indices, gene_cell_indices, gene_cell_umis)

    # Constructing the sparse array from the collated sparse indices, need to do here because sparse array unsupported in numba.
    cell_by_gene_umi_counts_SPARSE = csr_array((cell_by_gene_umi_counts_SPARSE_indices[:, 0], # The counts
                                                    (cell_by_gene_umi_counts_SPARSE_indices[:, 1],
                                                      cell_by_gene_umi_counts_SPARSE_indices[:, 2])), # row, col indices
                                                    shape=(len(cell_barcodes_dict),
                                                           len(gene_names)), # Total shape of sparse array.
                                                    dtype = np.uint32
                                                )

    # cell_by_gene_umi_counts = pd.DataFrame(cell_by_gene_umi_counts,
    #                                        index=list(cell_barcodes_dict.keys()),
    #                                        columns=gene_names_list)

    return cell_by_gene_umi_counts_SPARSE

@jit(nopython=True)
def get_cell_by_gene_umi_counts(total_cells,
                                gene_indices, gene_cell_indices, gene_cell_umis):

    # OLD dense version
    #cell_by_gene_umi_counts = np.zeros((total_cells, total_genes), dtype=np.uint16)

    # Will instead store the required Sparse format to create a sparse array:
    row_cell_indices = []
    col_gene_indices = []
    counts = []
    for genei in range(len(gene_indices)): # SHOULD NOT BE DONE IN PARALLEL COULD CREATE RACE CONDITION...

        gene_index = gene_indices[ genei ]

        cell_bc_indexes = gene_cell_indices[ genei ]
        cell_umis = gene_cell_umis[ genei ]

        # OLD dense version
        #cell_by_gene_umi_counts[:, gene_index] = cell_umi_counts_FAST(total_cells, cell_bc_indexes, cell_umis)
        cell_counts = cell_umi_counts_FAST(total_cells, cell_bc_indexes, cell_umis)
        cell_indices = np.where(cell_counts > 0)[0]
        cell_counts = cell_counts[ cell_indices ]

        row_cell_indices.extend( list(cell_indices) )
        col_gene_indices.extend( [gene_index]*len(cell_indices) )
        counts.extend( list(cell_counts) )

    # Outputting as a single array
    cell_by_gene_umi_SPARSE_indices = np.zeros((len(counts), 3), dtype=np.uint32)
    cell_by_gene_umi_SPARSE_indices[:, 0] = np.array(counts, dtype=np.uint32)
    cell_by_gene_umi_SPARSE_indices[:, 1] = np.array(row_cell_indices, dtype=np.uint32)
    cell_by_gene_umi_SPARSE_indices[:, 2] = np.array(col_gene_indices, dtype=np.uint32)

    return cell_by_gene_umi_SPARSE_indices

def bio_edit_distance(seqA, seqB, start_from_first_smallest_seq_aln=True, alns_to_compare=None):
    """More biologically relevant edit distance, which does a global alignment between two strings, then takes the
        edit distance as the number of gaps introduced minus the difference in sequence length. Will result
        in a 0 score if one sequence is a subset of the other.
    """
    ### Puts the sequences together better, at the 5' end
    aln = pairwise2.align.localms(seqA, seqB,
                                  1, # score for match
                                  -1, # mismatch penalty
                                  -.5, # gap-open penalty
                                  -.5, # gap-extension penalty
                                  one_alignment_only=True)[0]

    seq_lens = [len(seqA), len(seqB)]
    shorter_seq_index = np.argmin(seq_lens)
    #### Counting mismatches till all of shorter sequence accounted for
    aln_seqs = [aln.seqA, aln.seqB]
    n_seen = 0
    edit_dist = 0
    start_count = start_from_first_smallest_seq_aln == False # Only start counting once the alignment of the shorter sequence starts...
    if type(alns_to_compare)==type(None):
        alns_to_compare = len(aln.seqA) # Compare all of the global alignment, not just from one-side
    else: # Min between the max alignment length and the requested number to compare.
        alns_to_compare = min([len(aln.seqA), alns_to_compare])

    for i in range( alns_to_compare ):

        if start_count and aln.seqA[i] != aln.seqB[i]:
            edit_dist += 1

        if aln_seqs[shorter_seq_index][i] != '-': # Aligned character
            n_seen += 1
            start_count = True

            if n_seen == seq_lens[shorter_seq_index]: # seen all characters
                break

    return edit_dist

def get_edit_sets(edit_set):
    """Gets set of edits, version which did not considered subsequence problem, where partial overlap of subsequences
        between larger t7 edits results in very bad over-estimation of allelic number.
    """
    # Keep track of which edits look like each other, so only have a count of 1 for these edits.
    # This de-duplicates for edits that might represent slightly offset t7 priming, which can be quite common.
    # Basically the criteria is that the end of the read (forward read) or start of the read (reverse read)
    # is different between the two edit sites, indicating extra sequence content not present in the other.
    # Otherwise, could indicate different priming / read truncation due to the library fragmentation.

    edits_to_edit_matches = {edit: {edit} for edit in edit_set}  # initialise with just a set of itself

    for edit_1, edit_2 in itertools.combinations(edit_set, 2):

        # If they map in opposite directions, definitely different alleles edited
        # (donor-insert went in different direction at both alleles)
        # Also different alleles if the genomic proportion of the read is different, indicates different insert site
        reflen = len(edit_1.ref_seq)
        if (edit_1.forward != edit_2.forward) or (edit_1.ref_pos != edit_2.ref_pos):
            continue

        # Both forwards, so could be from the same allele with different variation in start site
        elif edit_1.forward and edit_2.forward:  # Reverse the direction to get the end of transcription
            edit_1_seq = edit_1.alt_seq[::-1][0:-reflen]  # Cutting off the added reference sequence
            edit_2_seq = edit_2.alt_seq[::-1][0:-reflen]

        elif edit_2.forward == False and edit_2.forward == False:
            edit_1_seq = edit_1.alt_seq[0:-reflen]  # Already in opposite direction of transcription
            edit_2_seq = edit_2.alt_seq[0:-reflen]

        #### The same insert at a different position is clear case of different insert
        if edit_1.alt_seq == edit_2.alt_seq and edit_1.ref_pos != edit_2.ref_pos:
            continue

        # Getting the common sequence at the end of the read in the direction of transcription
        # edit_seqs = [edit_1_seq, edit_2_seq]

        ##### Original method, does not consider sequencing error in either edit, below version does, even
        ####    for cases where the sequences are of different length!
        # longest_common_seq = os.path.commonprefix( edit_seqs )
        # # if the longest common sequence between the two is equal to one of the sequences, and ref
        # # position is the same, indicating the genomic part of the read is the same, the one sequence
        # # is likely a truncated read generated of the other, but generated from the same allele
        # if longest_common_seq in edit_seqs:
        #     edits_to_edit_matches[edit_1] = edits_to_edit_matches[edit_1].union({edit_2})
        #     edits_to_edit_matches[edit_2] = edits_to_edit_matches[edit_2].union({edit_1})
        #     continue

        # dist_between_seqs = nltk.edit_distance(edit_1_seq, edit_2_seq) - abs(len(edit_1_seq) - len(edit_2_seq))

        # The levenshtein distance above can under-estimate the difference between biological sequences, because
        # mutates them in ways that don't quite make biological sense which resulted in edit distances that
        # didn't make sense in some cases.
        # Below does a global alignment and gets the gap count in both aligned sequences as the edit distance,
        # subtracting the expected number of gaps due to differences in sequence length.
        dist_between_seqs = bio_edit_distance(edit_1_seq, edit_2_seq)

        if dist_between_seqs > 1:
            # HOMO-OLIGO sequencing error correction.
            # Need to correct for homo-oligo sequencing error. Can see cases where it OVERESTIMATES the number
            # of alleles, where the only different BETWEEN the edits is the NUMBER of a particular base IF there are
            # consecutive versions of that base. Illumina sequences is prone to this error. SO if that is the source
            # of difference, will remove this consecuative run of bases and re-calculate edit distance.
            # Example seen during debugging:
            # ATAATACTCTCCCTATTCACTCTGCGT
            # ATAATACTCTCCCCCCATTCACTC
            # -> Aside from the consecutive runs of 'CC', there is only an edit distance of 1 between these two sequences.
            #  and the 'CC' is not accurate, so would remove these consecutive runs and re-calculate the edit distance.
            # Find subsequences with more than 2 identical characters in a row, and replace them with one-copy of the
            # character BEFORE the edit-distance calculation between the sequences, to handle the homo-polymer
            # illumina sequencing error
            edit_1_consecs = re.findall(r"((.)\2{2,})", edit_1_seq)
            edit_2_consecs = re.findall(r"((.)\2{2,})", edit_2_seq)
            # Only a problem if one of them has a consecuative run of a particular base, the 'GGG' or 'CCC' of the
            # superb-seq barcode in particular seems to sometimes cause this issue, if is a non-canonical donor
            # sequence.
            if len(edit_1_consecs) > 0 or len(edit_2_consecs) > 0:
                # Example case am trying to catch here:
                # edit_1_seq: ATAATACTCTCCCCTTCACTCTGCGTTGATAC
                # edit_2_seq: ATAATACTCTCCCCCCATTCACT
                # edit_1_consecs: [('CCCC', 'C')]
                # edit_2_consecs: [('CCCCCC', 'C')]
                edit_1_seq_homo_fix = edit_1_seq
                for consec, base in edit_1_consecs:
                    #### I noticed that where there is a homopolymer, the next base
                    edit_1_seq_homo_fix = edit_1_seq_homo_fix.replace(consec, base)  # make it not a polymer

                edit_2_seq_homo_fix = edit_2_seq
                for consec, base in edit_2_consecs:
                    edit_2_seq_homo_fix = edit_2_seq_homo_fix.replace(consec, base)  # make it not a polymer
                # edit_1_seq_homo_fix: ATAATACTCTCTTCACTCTGCGTTGATAC
                # edit_2_seq_homo_fix: ATAATACTCTCATTCACT
                # the edit dist here is now:
                # nltk.edit_distance(edit_1_seq_homo_fix, edit_2_seq_homo_fix) - abs( len(edit_1_seq_homo_fix)-len(edit_2_seq_homo_fix) )
                # 2
                # due to the substitution cost, it prefers to flip two characters rather than create a gap,
                # which would yield a 1 edit distance. To enable this, will lower the substitution cost
                # dist_between_seqs = (nltk.edit_distance(edit_1_seq_homo_fix, edit_2_seq_homo_fix, substitution_cost=0.5) -
                #                      abs(len(edit_1_seq_homo_fix) - len(edit_2_seq_homo_fix)))
                dist_between_seqs = bio_edit_distance(edit_1_seq_homo_fix,
                                                      edit_2_seq_homo_fix) - 1  # -1 because the homopolymer extra error

        # Base-pair sequencing error correction
        # Some cases where the edits are VERY similar, off by 1bp, likely sequencing error!
        # Below is using the levenstein distance, so number of edits used to make one sequence into the other.
        # Taking the difference with the sequence lengths, to account for extra bp which would be added
        # to make the sequences the same if they are of different lengths. So would give 0 if 1 seq is a subset
        # of the other !
        if dist_between_seqs <= 1:  # Only one edit distance from each other, accounts for short indel at start also
            edits_to_edit_matches[edit_1] = edits_to_edit_matches[edit_1].union({edit_2})
            edits_to_edit_matches[edit_2] = edits_to_edit_matches[edit_2].union({edit_1})

    # Unique edits will be the number of unique sets of edits generated from the above dictionary!
    edit_sets = [set(edits_) for edits_ in edits_to_edit_matches.values()]
    # These unique edit sets will be inflated in terms of the unique edits, IF there are two edits that have
    # shared subsequence, with sub-edits shared between them. It causes them to be added to shared sets,
    # but they will be non-overlapping.
    unique_edit_sets = []
    [unique_edit_sets.append(edit_set_) for edit_set_ in edit_sets if edit_set_ not in unique_edit_sets]

    if len(unique_edit_sets) == 1:
        return unique_edit_sets
        # cell_allelic_edits[celli, edit_sitei] = len(unique_edit_sets)
        # continue

    # Need to de-duplicate these edit sets, since shared subsequences between longer edits can cause them to
    # create unique sets. So to de-duplicate, need to go through and REMOVE edit sets which represent
    # combinations of other edit sets!
    unique_edit_sets_deoverlapped = unique_edit_sets.copy()
    for edit_set_1, edit_set_2 in itertools.combinations(unique_edit_sets, 2):

        union_set = edit_set_1.union(edit_set_2)
        if union_set in unique_edit_sets_deoverlapped:
            unique_edit_sets_deoverlapped.pop(unique_edit_sets_deoverlapped.index(union_set))

    return unique_edit_sets_deoverlapped

def get_longest_edits(edit_set):
    """This method tries to get the longest t7 edits that are distinct from one another, overcomes subsequence problem
        of previous method.
    """
    # This ensures the inputted edit_set has a deterministic order, so between runs there is not a difference in the
    # outputs.
    edit_set = sorted(edit_set, key=lambda e: (len(e.alt_seq), len(e.kmer_matches), e.ref_pos, id(e)))

    # In this version, just for looking
    edits_to_subedits = {edit: set() for edit in edit_set}  # initialise with list

    ##### Will keep track of the longer edits in a set, where if two edits are matched, only store the longer
    ##### edit, and if the shorter edit is currently called as a longer edit from previous comparisons, pop
    ##### it from the long-edit list.
    edit_set = list(edit_set)
    edit_lens = np.array([len(edit.alt_seq) for edit in edit_set])
    edit_order = np.argsort(edit_lens)  # Will go from smallest to largest

    # By starting with the shortest edits, comparing to everything larger than it, the shorter sequences will initially
    # be called the longer edits, but if a longer match is found will downgrade to a sub-edit, with the final
    # set of long edits being the edits that could not be matched to a longer edit.

    longest_edits = []  # These will be our final edit set!
    sub_edits = [] # These are the edits that have already been called subedits, so don't add them as longer edit!
    for i, edit_1_i in enumerate(edit_order):
        edit_1 = edit_set[edit_1_i]

        for edit_2_i in edit_order[i+1:]: # Compare against all longer edits not yet seen
            edit_2 = edit_set[edit_2_i]

            # If they map in opposite directions, definitely different alleles edited
            # (donor-insert went in different direction at both alleles)
            # Also different alleles if the genomic proportion of the read is different, indicates different insert site
            reflen = len(edit_1.ref_seq)
            if (edit_1.forward != edit_2.forward) or (edit_1.ref_pos != edit_2.ref_pos):
                if edit_1 not in sub_edits: # Hasn't been logged as a subedit.
                    longest_edits.append( edit_1 )
                if edit_2 not in sub_edits:
                    longest_edits.append( edit_2 )

                continue

            # Both forwards, so could be from the same allele with different variation in start site
            elif edit_1.forward == False and edit_2.forward == False:  # Reverse the direction to get the end of transcription
                edit_1_seq = edit_1.alt_seq[reflen:]  # Cutting off the added reference sequence
                edit_2_seq = edit_2.alt_seq[reflen:]
                # Need to reverse the direciton, since the 'bio_edit_distance' function will ONLY count the edit distance
                # alignement correctly if the shorter subsequence maps to the FRONT of the longer subsequence.
                edit_1_seq = edit_1_seq[::-1]
                edit_2_seq = edit_2_seq[::-1]

            elif edit_2.forward and edit_2.forward:
                edit_1_seq = edit_1.alt_seq[0:-reflen]  # Already in opposite direction of transcription
                edit_2_seq = edit_2.alt_seq[0:-reflen]

            # Below does a global alignment and gets the gap count in both aligned sequences as the edit distance,
            # subtracting the expected number of gaps due to differences in sequence length.
            dist_between_seqs = bio_edit_distance(edit_1_seq, edit_2_seq)

            if dist_between_seqs > 1:
                # HOMO-OLIGO sequencing error correction.
                # Need to correct for homo-oligo sequencing error. Can see cases where it OVERESTIMATES the number
                # of alleles, where the only different BETWEEN the edits is the NUMBER of a particular base IF there are
                # consecutive versions of that base. Illumina sequences is prone to this error. SO if that is the source
                # of difference, will remove this consecuative run of bases and re-calculate edit distance.
                # Example seen during debugging:
                # ATAATACTCTCCCTATTCACTCTGCGT
                # ATAATACTCTCCCCCCATTCACTC
                # -> Aside from the consecutive runs of 'CC', there is only an edit distance of 1 between these two sequences.
                #  and the 'CC' is not accurate, so would remove these consecutive runs and re-calculate the edit distance.
                # Find subsequences with more than 2 identical characters in a row, and replace them with one-copy of the
                # character BEFORE the edit-distance calculation between the sequences, to handle the homo-polymer
                # illumina sequencing error
                edit_1_consecs = re.findall(r"((.)\2{2,})", edit_1_seq)
                edit_2_consecs = re.findall(r"((.)\2{2,})", edit_2_seq)
                # Only a problem if one of them has a consecuative run of a particular base, the 'GGG' or 'CCC' of the
                # superb-seq barcode in particular seems to sometimes cause this issue, if is a non-canonical donor
                # sequence.
                if len(edit_1_consecs) > 0 or len(edit_2_consecs) > 0:
                    # Example case am trying to catch here:
                    # edit_1_seq: ATAATACTCTCCCCTTCACTCTGCGTTGATAC
                    # edit_2_seq: ATAATACTCTCCCCCCATTCACT
                    # edit_1_consecs: [('CCCC', 'C')]
                    # edit_2_consecs: [('CCCCCC', 'C')]
                    edit_1_seq_homo_fix = edit_1_seq
                    for consec, base in edit_1_consecs:
                        #### I noticed that where there is a homopolymer, the next base
                        edit_1_seq_homo_fix = edit_1_seq_homo_fix.replace(consec, base)  # make it not a polymer

                    edit_2_seq_homo_fix = edit_2_seq
                    for consec, base in edit_2_consecs:
                        edit_2_seq_homo_fix = edit_2_seq_homo_fix.replace(consec, base)  # make it not a polymer
                    # edit_1_seq_homo_fix: ATAATACTCTCTTCACTCTGCGTTGATAC
                    # edit_2_seq_homo_fix: ATAATACTCTCATTCACT
                    # the edit dist here is now:
                    # nltk.edit_distance(edit_1_seq_homo_fix, edit_2_seq_homo_fix) - abs( len(edit_1_seq_homo_fix)-len(edit_2_seq_homo_fix) )
                    # 2
                    # due to the substitution cost, it prefers to flip two characters rather than create a gap,
                    # which would yield a 1 edit distance. To enable this, will lower the substitution cost
                    dist_between_seqs = bio_edit_distance(edit_1_seq_homo_fix, edit_2_seq_homo_fix) - 1  # -1 because the homopolymer adds extra error

            # They are different from one another in the sense that one is not a subset of the other (i.e. not
            # different due to different t7 primering).
            # Need to now consider if they only differ at the 5' end of the read, if so likely only different
            # due to sequencing adapters / TSO stil present at the end of the read!
            # Getting the edit distance from the first 5 base-pairs of the 3' end of the read
            if dist_between_seqs > 2:
                # They are different from one another in the sense that one is not a subset of the other (i.e. not
                # different due to different t7 primering).
                # Need to now consider if they only differ at the 5' end of the read, if so likely only different
                # due to sequencing adapters / TSO stil present at the end of the read!
                # Getting the edit distance from the first 5 base-pairs of the 3' end of the read
                three_prime_edit_dist = bio_edit_distance(edit_1_seq[::-1], edit_2_seq[::-1],
                                                          start_from_first_smallest_seq_aln=False,
                                                          alns_to_compare=10)

                # Only 1 mismatch at the 3' end! so let's consider them the same sequence, allocating shorter as sub...
                if three_prime_edit_dist <= 1:
                    dist_between_seqs = three_prime_edit_dist

            # Base-pair sequencing error correction
            # Some cases where the edits are VERY similar, off by 1bp, likely sequencing error!
            # Below is using the levenstein distance, so number of edits used to make one sequence into the other.
            # Taking the difference with the sequence lengths, to account for extra bp which would be added
            # to make the sequences the same if they are of different lengths. So would give 0 if 1 seq is a subset
            # of the other !
            if dist_between_seqs <= 2:  # edit distance cutoff from each other, accounts for short indel at start also

                # Recording the longer edit, assigning the shorter edit to the sub-edit list
                edit_pair = [edit_1, edit_2]
                edit_pair_lens = np.array([len(edit_1.alt_seq), len(edit_2.alt_seq)])
                if edit_pair_lens[0] != edit_pair_lens[1]: # different lengths, so clear which is the subedit
                    long_index = np.argmax([len(edit_1.alt_seq), len(edit_2.alt_seq)])
                else: # same length, choose one with more kmer matches to barcode
                    n_kmers = [len(edit_1.kmer_matches), len(edit_2.kmer_matches)]
                    if n_kmers[0] != n_kmers[1]: # Different kmer matches
                        long_index = np.argmax(n_kmers)
                    else: # Match from this criteria to, just choose the first then
                        long_index = 0

                long_edit = edit_pair.pop(long_index)
                long_edits = [long_edit] # So can process as a list below!

                subedit = edit_pair[0] # remaining edit is the subedit

                # Add the subedit!
                edits_to_subedits[long_edit] = edits_to_subedits[long_edit].union({subedit})

                while subedit in longest_edits: # Need to remove the subedit from list of long edits, found a longer match
                    longest_edits.pop( longest_edits.index(subedit) )

                if subedit not in sub_edits: # Need to add to subedit list
                    sub_edits.append( subedit )

            else: # They are different by all our criteria! including at 3' end. So let's consider them both long edits
                long_edits = [edit_1, edit_2]

            # Now adding to the longest_edits list if any long_edits found, if they were not already categorised as
            # subedits!!!!
            for long_edit in long_edits:
                # not previously found as a subedit, and found here is a longer edit, so add it to longest edits list!
                if long_edit not in longest_edits and long_edit not in sub_edits:
                    longest_edits.append( long_edit )

    # The surviving longest edits did not have a match with a longer t7 insert, so they are the set of unique edit sites
    seen = set()
    deduped_longest_edits = []
    for e in longest_edits:
        if e not in seen:
            seen.add(e)
            deduped_longest_edits.append(e)

    return deduped_longest_edits

def bed_file_flag_edits(bed_file, canonical_edit_sites, keep_sites, whitelist,
                          edit_dist, # Extra error around the edit site specification to allow for overlap with the bed regions
                          verbosity=1,
                        ):
    """ whitelist = True flag to flag any of the canonical edit sites for REMOVAL if intersect with the bed regions.
        whitelist = False to flag any canonical edit sites that intersect with the bed regions to KEEP.
    """
    if whitelist:
        filter_type = "whitelist"
        keep_indices = np.array( list(range(len(canonical_edit_sites))) ) # Since need to iterate through every site.
    else:
        filter_type = "blacklist"
        # Updating the keep_sites to remove those that intersect blacklisted regions...
        keep_indices = np.where(keep_sites)[0]

    blacklist_regions = pd.read_csv(bed_file, sep='\t', header=None)
    blacklist_chroms = blacklist_regions.values[:, 0].astype(str)
    blacklist_starts = blacklist_regions.values[:, 1].astype(int)
    blacklist_ends = blacklist_regions.values[:, 2].astype(int)

    black_chrom_set = set(list(blacklist_chroms))
    blacklist_chrom_to_indices = {chrom: np.where(blacklist_chroms == chrom)[0]
                                  for chrom in black_chrom_set}

    flagged_indices = []
    for index in keep_indices:
        edit_site = canonical_edit_sites[index]

        # Checking if intersects blacklist region.
        edit_chrom, edit_pos = edit_site.chrom, edit_site.ref_pos
        if edit_chrom in black_chrom_set:
            black_chrom_indices = blacklist_chrom_to_indices[edit_chrom]
            starts_, ends_ = blacklist_starts[black_chrom_indices], blacklist_ends[black_chrom_indices]
            # +/- edit_dist to the edit site, so can account for a level of error in edit site determination.
            overlap_bool = np.logical_and(starts_ <= (edit_pos + edit_dist), ends_ >= (edit_pos - edit_dist))

            if np.any(overlap_bool):
                keep_sites[index] = whitelist # Setting the keep_site to false, so now we filter this edit site !!!
                flagged_indices.append( index )
                print(f"Overlap found: {edit_site}: ", file=sys.stdout, flush=True) if verbosity >= 2 else None
                for overlap_index in np.where(overlap_bool)[0]:
                    print(f"{filter_type} region {edit_chrom}: {starts_[overlap_index]}-{ends_[overlap_index]}",
                                               file=sys.stdout, flush=True) if verbosity >= 2 else None

    return flagged_indices
```
