## Additional Files for "Joint single-cell capture of Cas9 edits and transcriptomes reveals on- and off-target effects on gene expression": README.md

# Superb-seq Manuscript code

Analysis code for the Superb-seq manuscript after processing the data with [Sheriff](https://github.com/BradBalderson/Sheriff)

Mostly intended as a detailed record of methods utilised.

Note that all directory paths have been removed, simply listing the name of the input/output file, in order to not
leak sensitive information about file structures in remote servers.
    
## Quick index
Quick explanation of respository structure.

scripts/

    split_pipe/ -> Code for running ParseBiosciences split-pipe to generated annotated bam files.

        split_pipe_10k_cell.sh -> Runs split-pipe with Sun-gride-engine (SGE) on the 10k cell library paired-end fastq 
                                    files.

        split_pipe_500_cell.sh -> Runs split-pipe with Sun-gride-engine (SGE) on the 500 cell library paired-end fastq 
                                    files.

    run_sheriff/ -> Code which ran Sheriff, not that used a slightly different API and was named superb_count.

        blacklist.ipynb -> Code for generating blacklist regions for calling canonical edit sites with Sheriff.

        # 10k cell library processing
        superb_count_multi.sh -> SGE script that processed the split-pipe annotated bam for 10k library, 
                                which had been split by chromosome, and also wrote out the output by chromosome. 

        combine_results.py -> Used to combine all of the count matrices and bam files for the processing across 
                                chromosomes, also compiles all of this into an Anndata object for downstream analysis.

        # 500 cell library processing
        superb_count_multi_500-cells.sh -> Bash script for the read processing for the 500 cell library, not done in 
                                            parallel.

        create_anndata.py -> Used to compile the 500 cell count matrices into an AnnData for downstream analysis.

    downstream_analysis/ -> Code related to downstream analysis after the Superb-seq annotated bam processing.

        superb_analysis_guide_inference.py -> Adds the manual edit calls to the edit site information in the 
                                        AnnData, and also perform the edit-to-guide inference and adds this information
                                        to the edit site info also.

            superb_guide_inference.py -> Supporting helper functions for superb_analysis_barcode-correction_v2.py with
                                          regards to the edit-to-guide inference analysis.

        cell_qc_10k-lib.ipynb -> Creates the cell QC plots for the 10k library. Also includes the motif analysis around
                                    the USP9X edit site.

        cell_qc_5k-lib.ipynb -> Creates the cell QC plots for the 500 cell library, but also compares the 500 cell library
                                to the 10k library in terms of the detected edits cells and alleles.

        compile_baf-nurd-targets.ipynb -> Performs the ENCODE analysis to determine genes regulated by BAF, NuRD, and 
                                            NRF1.

        superb_differential_expression_analysis.py -> Performs the differential expression analysis and over-representation analysis with 
                                        respect to allelic edit dosage. Comments within report the R code used for the 
                                        Pearson's residuals normalisation with Sctransform.

            superb_helpers.py -> Supporting helper functions for performing the differential expression analysis.

            utils.py -> Some extra auxillary functions to help with saving plots and other miscallaneous repeated task.
        
