## Additional Files for "Joint single-cell capture of Cas9 edits and transcriptomes reveals on- and off-target effects on gene expression": cell_qc_5k-lib.ipynb

```
# %% [markdown]
# Looking at the cell QC metrics, 500 cell library.

# INPUT:

    * superb_count_500_cells1/superb_data.h5ad

# OUTPUT:

    * cell_qc_tables/superb_500cell_v1/*

# Environment Setup

# %% [code]
work_dir = './'
import os
os.chdir( work_dir )

import numpy as np
import pandas as pd

import scanpy as sc

import seaborn as sb
import matplotlib.pyplot as plt

data_dir = 'superb_count_500_cells1/'
data_dir2 = 'superb_count_all/combined/'
out_dir = data_dir+'cell_qc_tables/'
out_plots = 'superb_500cell_v1/'

# %% [markdown]
# Loading the data

# %% [code]
data = sc.read_h5ad(f'{data_dir}superb_data.h5ad')
data

# %% [code]
data.shape

# %% [code]
data.var.keys()

# %% [code]
##### Counting the mitochondrial reads per cell.
mito_genes = [gene for gene in data.var_names if gene.startswith('MT-')]
mito_counts = data[:,mito_genes].layers['gene_counts'].toarray().sum(axis=1)
mito_perc = (mito_counts.astype(float) / data.obs['total_counts'].values)*100

data.obs['mito_perc'] = mito_perc

# %% [code]
sum(mito_counts)

# %% [markdown]
***Need to label which are the real edits or not by looking at the t7 barcoded read bam file***

# %% [code]
real_edits = ['hg38_12:6602172', 'hg38_14:104729499', 'hg38_17:7890653', 'hg38_4:14359416', 'hg38_19:10984175',
             'hg38_17:47188686', 'hg38_6:1397192', 'hg38_1:224544957', 'hg38_X:41086519', 'hg38_12:110404169',
             'hg38_12:6606341', 'hg38_5:148237295', 'hg38_4:59298822', 'hg38_20:14937204', 'hg38_17:3029640',
             'hg38_1:26696719', 'hg38_13:110873574', 'hg38_14:71885108', 'hg38_12:68620881',
             'hg38_4:88368039', 'hg38_4:79849103', 'hg38_3:194003699', 'hg38_9:37594621', 'hg38_1:26696864',
             'hg38_1:245947392', 'hg38_18:3115881', 'hg38_19:6212712', 'hg38_19:10986490',
             'hg38_13:113608068', 'hg38_16:81393308']

# %% [code]
edit_site_info = data.uns['edit_site_info']

edit_site_info['manual_edit_call'] = [edit_site in real_edits for edit_site in edit_site_info['name']]

edit_site_info

# %% [markdown]
***Think in the blacklist regions, should basically call all of the MT as blacklisted, since has a high number of FP calls.***

# %% [code]
##### Making a manual edit call version!
data.obsm.keys()

# %% [code]
t7_keys = ['dosages', 't7_all', 't7_no_bc_umis', 't7_umis']
for key in t7_keys:

    data.obsm[f"{key}_manual-edits-only"] = data.obsm[key].loc[:,real_edits]

# %% [code]
data.obsm[f"{key}__manual-edits-only"]

# %% [code]
manual_gene_edits_ = edit_site_info['intersecting_genes'].loc[edit_site_info['manual_edit_call']]
manual_edit_names = edit_site_info['name'].loc[edit_site_info['manual_edit_call']]

manual_gene_edits = []
for entry_name, entry in zip(manual_edit_names, manual_gene_edits_):
    if entry!='nan':
        manual_gene_edits.extend( entry.split(',') )
    else:
        manual_gene_edits.append( entry_name )

manual_gene_edits = np.unique( manual_gene_edits )
len(manual_gene_edits), manual_gene_edits[0:3]

# %% [code]
data.obsm['dosages_genes_manual-edits-only'] = data.obsm['dosages_genes'].loc[:,manual_gene_edits]
data.obsm['dosages_genes_manual-edits-only']

# %% [markdown]
***Let's count the t7 barcoded reads!***

# %% [code]
data.obsm.keys()

# %% [code]
t7_umis = data.obsm['t7_umis_manual-edits-only'].sum(axis=1).values

# %% [code]
t7_umis

# %% [code]
data.obs['t7_umis'] = t7_umis

# %% [code]
data.obs['t7_all'] = data.obsm['t7_all_manual-edits-only'].sum(axis=1).values

# %% [code]
data.obs['t7_no_bc_umis'] = data.obsm['t7_no_bc_umis_manual-edits-only'].sum(axis=1).values

# %% [markdown]
# Per cell metrics

# %% [code]
out_plots

# %% [code]
data.obs

# %% [code]
#### Need to label by sample first:
data_dir_orig = 'DGE_filtered/'
cell_meta = pd.read_csv(f"{data_dir_orig}cell_metadata.csv")
cell_meta.index = cell_meta['bc_wells'].values
cell_meta = cell_meta.loc[data.obs_names.values, :]

cell_meta.shape, cell_meta.head(3)

# %% [code]
data.obs['sample'] = cell_meta['sample'].values
data.obs['sample'] = data.obs['sample'].astype('category')

# %% [code]
def pie(labels: np.array, label_set: np.array=None,
        colors: dict=None, min_perc: float=3, autopct: str='',
        labeldistance=1.05, explode_f=0.8,
        filter_perc=0, show_perc=False, show_size=False, print_filt=False, figsize=(5,5),
        show=True):
    """Piechart, but formatted nicely, e.g. will order pies & explode out
        low frequency labels.

    Args:

        min_perc (float): Percentage below which labels will 'explode'.
        filter_perc (float): Percentage below which labels will not be shown.

    """
    if type(label_set)==type(None):
        label_set = np.unique(labels)

    sizes = np.array([len(np.where(labels == dataset)[0])
                      for dataset in label_set])
    percs = np.array([(size / sum(sizes)) * 100 for size in sizes])

    if filter_perc > 0:
        perc_filt = percs > filter_perc
        if print_filt:
            print( label_set[perc_filt==False] )

        remain_size = sum(sizes[perc_filt == False])
        sizes = np.array(list(sizes[perc_filt]) + [remain_size])
        remain_perc = sum(percs[perc_filt == False])
        percs = np.array(list(percs[perc_filt]) + [remain_perc])

        label_set = np.array(list(label_set[perc_filt]) + [f'other'])

    if show_perc:
        label_set = np.array([f'{label} ({round(perc, 2)}%)'
                              for label, perc in zip(label_set, percs)])
    elif show_size:
        label_set = np.array([f'{label} (n={round(size_, 2)})'
                              for label, size_ in zip(label_set, sizes)])

    order = np.argsort(-sizes)
    sizes = sizes[order]
    label_set = label_set[order]
    percs = percs[order]

    ### Getting the colors, will add this to scripts.helpers so consistent ###
    if type(colors)==type(None):
        colors = vhs.getColors(label_set)

    colors_ = [colors[dataset] for dataset in label_set]
    end = sum(percs<min_perc)
    explode = [0] * (len(label_set) - end) + [explode_f * (i + 1) for i in list(
        range(end))]  # Expand out less numerous data...

    fig1, ax1 = plt.subplots(figsize=figsize)
    ax1.pie(sizes, labels=label_set, autopct=autopct,  # '%0.1f%%',
            shadow=False, startangle=90, colors=colors_, explode=explode,
            labeldistance=labeldistance)
    ax1.axis('equal')  # Equal aspect ratio ensures that pie is drawn as a circle.

    if show:
        plt.show()
    else:
        return fig1, ax1


# %% [code]
#### Pie chart of the cells captured in the different samples
sample_colors = {'nurd_ki': 'palegreen', 'baf_ki': 'lightskyblue', 'no_ki': 'magenta'}
sample_colors_wSizes = {'nurd_ki (n=323)': sample_colors['nurd_ki'], 
                        'baf_ki (n=219)': sample_colors['baf_ki'], 
                        'no_ki (n=41)': sample_colors['no_ki']}

pie(data.obs['sample'].values, colors=sample_colors_wSizes, 
        show_size=True, show=False)
plt.show()

# %% [markdown]
# Let's also get the proportion of these that have the on-target edit in each cell, and evidence of off-target in different combinations..

# %% [code]
sample_labels = data.obs['sample'].values

dosages_genes = data.obsm['dosages_genes_manual-edits-only']
arid1a_edited = dosages_genes['ARID1A'].values > 0
smarca4_edited = dosages_genes['SMARCA4'].values > 0
chd3_edited = dosages_genes['CHD3'].values > 0
chd4_edited = dosages_genes['CHD4'].values > 0

any_edited = dosages_genes.sum(axis=1) > 0

# %% [code]
np.unique( sample_labels )

# %% [code]
len(any_edited)

# %% [code]
def double_label(edit_bool_1, edit_bool_2, gene_1, gene_2, any_edited_bool):
    
    cell_label = 'no-edit'
    if edit_bool_1 and not edit_bool_2:
        cell_label = f'{gene_1}-edit'
    elif not edit_bool_1 and edit_bool_2:
        cell_label = f'{gene_2}-edit'
    elif edit_bool_1 and edit_bool_2:
        cell_label = f'double-edit'

    if any_edited_bool and cell_label!='no-edit':
        cell_label = cell_label + ', off-target/s'
    elif any_edited_bool:
        cell_label = 'off-target/s'
        
    return cell_label

# %% [code]
cell_labels = []
for i in range(len(sample_labels)):

    if sample_labels[i] == 'no_ki' and any_edited[i]:
        cell_labels.append( 'false-edit-call' )

    elif sample_labels[i] == 'no_ki':
        cell_labels.append( 'no-edit' )

    elif sample_labels[i] == 'baf_ki':
        cell_labels.append( double_label(arid1a_edited[i], smarca4_edited[i], 'ARID1A', 'SMARCA4', any_edited[i]) )

    elif sample_labels[i] == 'nurd_ki':
        cell_labels.append( double_label(chd3_edited[i], chd4_edited[i], 'CHD3', 'CHD4', any_edited[i]) )

# %% [code]
np.unique(cell_labels)

# %% [code]
data.obs['edit_cell_labels'] = cell_labels
data.obs['edit_cell_labels'] = data.obs['edit_cell_labels'].astype('category')

# %% [code]
def get_overlap_counts(data, groupby1: str, groupby2: str,
                       uns_key: str = 'annot_overlaps',
                       verbose: bool = True):
    """ Calculates no. of cells belonging to the different categories of groupby1 and groupby2.
    """

    labels = data.obs[groupby1].values.astype(str)
    label_set = np.array(list(data.obs[groupby1].cat.categories))

    labels2 = data.obs[groupby2].values.astype(str)
    label_set2 = np.array(list(data.obs[groupby2].cat.categories))

    overlap_counts = np.zeros((len(label_set), len(label_set2)))
    for i, label in enumerate(label_set):
        label_bool = labels == label
        for j, label2 in enumerate(label_set2):
            label_bool2 = labels2 == label2

            overlap_counts[i, j] = len(
                np.where(np.logical_and(label_bool, label_bool2))[0])

    overlap_counts_df = pd.DataFrame(overlap_counts, index=label_set,
                                     columns=label_set2)
    data.uns[uns_key] = overlap_counts_df
    if verbose:
        print(f"Added data.uns['{uns_key}']")

# %% [code]
get_overlap_counts(data, 'sample', 'edit_cell_labels')

# %% [code]
data.uns['annot_overlaps']

# %% [code]
#### Want to see the actual proportions:
totals = data.uns['annot_overlaps'].values.sum(axis=1)
proportions = np.apply_along_axis(np.divide, 0, data.uns['annot_overlaps'].values, totals)

pd.DataFrame(proportions, index=data.uns['annot_overlaps'].index, columns=data.uns['annot_overlaps'].columns)

# %% [code]
proportions.sum(axis=1)

# %% [code]
sc.pl.umap(data, color='edit_cell_labels')

# %% [code]
data.obs['edit_cell_labels'].cat.categories

# %% [code]
def getColors(labels, labelSet=None, colorMap='tab20', rgb=False):
	""" Gets an OrderedDict of colors; the order indicates the frequency of \
	labels from largest to smallest.

	Args:
		labels (numpy.array<str>): Indicates a set of labels for observations.

		labelSet (list-like<str>): Indicates the set of labels in labels. \
									If None, calculated based on labels.

		colorMap (str): A matplotlib colormap.

		rgb (bool): If True, colors indicated by rgb value, if false hexcode.

	Returns:
		dict<str, tuple or str>: An ordered dict indicating the labels which \
					occur most to least frequently and the associated colors.
	"""
	# Determining the set of labels #
	labelSet = input.returnDefaultIfNone(labelSet,
										 calcs.getOrderedLabelSet(labels))

	# Initialising the ordered dict #
	cellTypeColors = {}

	# Ordering the cells according to their frequency and obtaining colors #
	nLabels = len(labelSet)
	cmap = plt.cm.get_cmap(colorMap, nLabels)
	rgbs = [cmap(i)[:3] for i in range(nLabels)]
	#rgbs = list(numpy.array(rgbs)[order]) # Make sure color order is the same.

	# Populating the color dictionary with rgb values or hexcodes #
	for i in range(len(labelSet)):
		cellType = labelSet[i]
		rgbi = rgbs[i]
		if not rgb:
			cellTypeColors[cellType] = matplotlib.colors.rgb2hex(rgbi)
		else:
			cellTypeColors[cellType] = rgbi

	return cellTypeColors

# %% [code]
color_dict = getColors(cell_labels)

color_dict['no-edit'] = 'silver'
color_dict['false-edit-call'] = 'dimgrey'
color_dict['off-target/s'] = 'sienna'#'crimson'
color_dict['CHD3-edit, off-target/s'] = 'yellow'
color_dict['CHD4-edit, off-target/s'] = 'goldenrod'
color_dict['ARID1A-edit, off-target/s'] = 'lime' #'#20DF7F'
color_dict['SMARCA4-edit, off-target/s'] = 'green' #'#45BA45'
color_dict['double-edit, off-target/s'] = 'palegoldenrod'
data.uns['edit_cell_labels_colors'] = [color_dict[label_] for label_ in data.obs['edit_cell_labels'].cat.categories]

# %% [code]
color_dict

# %% [code]
len(color_dict)

# %% [code]
data.uns['annot_overlaps']

# %% [code]
def dataframe_annot_overlap_barplot(overlap_counts_df, groupby1, groupby2, colors=None,
                          bar_width: float = .85,
                          labelset1 = None, labelset2 = None,
                          figsize=(10, 5), scale: bool = True,
                          transpose: bool = False, edge_color=None,
                          remove_labelling: bool = False):
    """Plots relationship between annotations by showing proportions of each type found within
        each of the other categories.
    """
    if transpose:  ##### Looking at relationship in opposite direction.
        overlap_counts_df = overlap_counts_df.transpose()
        tmp = groupby1
        groupby1 = groupby2
        groupby2 = tmp

    if type(labelset1)==type(None):
        labelset1 = overlap_counts_df.index.values
    else:
        overlap_counts_df = overlap_counts_df.loc[labelset1, :]

    if type(labelset2) == type(None):
        labelset2 = overlap_counts_df.columns.values
    else:
        overlap_counts_df = overlap_counts_df.loc[:, labelset2]

    ##### Getting colors
    if type(colors)==type(None):
        colors = {val: 'grey' for i, val in enumerate(labelset2)}

    ##### Scaling the data to proportions.
    if scale:
        total_counts = overlap_counts_df.sum(axis=1)
        overlap_counts = np.apply_along_axis(np.divide, 0,
                                             overlap_counts_df.values,
                                             total_counts)
    else:
        overlap_counts = overlap_counts_df.values

    ##### Creating the barplots...
    fig, axs = plt.subplots(figsize=figsize)
    col_locs = list(range(len(labelset1)))
    for labeli, label in enumerate(labelset2):
        props = overlap_counts[:, labeli]

        if labeli == 0:
            bottom = [0] * len(props)

        axs.bar(col_locs, props, bottom=bottom,
                color=colors[label], edgecolor=edge_color,
                width=bar_width, )
        bottom = list(np.array(bottom) + props)

        # Custom x axis
        axs.set_xticks(list(range(len(labelset1))), list(labelset1),
                       rotation=90)
        axs.spines['right'].set_visible(False)
        axs.spines['top'].set_visible(False)
        if remove_labelling:
            axs.set_xticks([])
            axs.set_yticks([])
            axs.spines['left'].set_visible(False)
            axs.spines['bottom'].set_visible(False)
        else:
            axs.set_xlabel(f"{groupby1} Clusters")

def annot_overlap_barplot(data, groupby1: str, groupby2: str,
                          uns_key: str = 'annot_overlaps',
                          labelset1 = None, labelset2 = None,
                          bar_width: float = .85,
                          figsize=(10, 5), scale: bool = True,
                          transpose: bool = False, edge_color=None,
                          remove_labelling: bool = False):
    """Plots relationship between annotations by showing proportions of each type found within
        each of the other categories. Written to work with AnnData input.
    """
    if uns_key not in data.uns:
        print("Need to run get_overlap_counts() first.")

    overlap_counts_df = data.uns[uns_key].copy()

    # ##### Getting colors
    if f'{groupby2}_colors' in data.uns:
        colors = {val: data.uns[f'{groupby2}_colors'][i]
                  for i, val in enumerate(data.obs[groupby2].cat.categories)}
    else:
        colors = {val: 'grey' for
                  i, val in enumerate(data.obs[groupby2].cat.categories)}


    dataframe_annot_overlap_barplot(overlap_counts_df, groupby1, groupby2, colors=colors,
                          bar_width = bar_width,
                          labelset1 = labelset1, labelset2 = labelset2,
                          figsize=figsize, scale=scale,
                          transpose=transpose, edge_color=edge_color,
                          remove_labelling=remove_labelling)


# %% [code]
annot_overlap_barplot(data, 'sample', 'edit_cell_labels',
                         labelset1=np.array(['no_ki', 'baf_ki', 'nurd_ki']),
                         labelset2=np.array(['no-edit', #'false-edit-call',
                                            'ARID1A-edit, off-target/s', 'SMARCA4-edit, off-target/s',
                                            'CHD3-edit, off-target/s', 'CHD4-edit, off-target/s',
                                             'double-edit, off-target/s',
                                             'off-target/s'
                                            ][::-1])
                         )
plt.show()

# %% [markdown]
# Looking at dosages of allele calls

# %% [code]
on_targets = ['CHD3', 'CHD4', 'ARID1A', 'SMARCA4']
samples_ = ['nurd_ki', 'nurd_ki', 'baf_ki', 'baf_ki']
colors_  = ['yellow', 'goldenrod', '#20DF7F', '#45BA45']

# Constructing own overlaps...
allelic_counts = np.zeros((len(on_targets), 3))

for i, (sample_, gene_) in enumerate( zip(samples_, on_targets) ):

    color_ = colors_[i]
    data.obs[f"{gene_}_edit_dosages"] = dosages_genes[gene_].values.astype(str)
    data.obs[f"{gene_}_edit_dosages"] = data.obs[f"{gene_}_edit_dosages"].astype('category')

    get_overlap_counts(data, 'edit_cell_labels', f"{gene_}_edit_dosages")

    # This part only works cause no-cases where have a pure edit with no-off-targets detected.
    allele_counts = data.uns['annot_overlaps'].loc[f'{gene_}-edit, off-target/s'].values[1:]
    alleles = data.uns['annot_overlaps'].columns.values[1:]

    allelic_counts[i,0:len(alleles)] = allele_counts
    
    plt.bar(alleles, allele_counts,
       color=color_)
    plt.xlabel('allelic edits')
    plt.ylabel('n cells')

    plt.show()

# %% [code]
allelic_counts = pd.DataFrame(allelic_counts, index=on_targets, columns=list(range(1,allelic_counts.shape[1]+1)))
allelic_counts

# %% [code]
data.uns['annot_overlaps'] = allelic_counts

# %% [code]
totals = data.uns['annot_overlaps'].values.sum(axis=1)
proportions = np.apply_along_axis(np.divide, 0, data.uns['annot_overlaps'].values, totals)

pd.DataFrame(proportions, index=data.uns['annot_overlaps'].index, columns=data.uns['annot_overlaps'].columns)

# %% [code]
data.uns.keys()

# %% [code]
np.unique([-1]+list(allelic_counts.columns.values.astype(int)))

# %% [code]
allele_colors = getColors(np.unique([0]+list(allelic_counts.columns.values.astype(int))), colorMap='Blues')
allele_colors

# %% [code]
dataframe_annot_overlap_barplot(allelic_counts, 'on-targets', 'alleles-edited', colors=allele_colors)
plt.show()

# %% [code]
#### Some stats on the total counts per cell!
total_counts = data.obs['total_counts'].values

min_, median, max_ = np.quantile(total_counts, [0, 0.5, 1])

print("Min, median, max:", min_, median, max_)

# %% [code]
sample_labels = data.obs['sample'].values
sample_set = np.unique(sample_labels)

# %% [code]
def distrib(x, bins=100, x_label='', fig_title='', log=False, density=False,
            figsize=(6.4,4.8), file_name=None, add_mean=False, logbase=np.e,
            color='blue', alpha=1, ax=None, fig=None, show=True, cutoff=None,
            cutoff_color='r',
            label='', total=None, return_total=False, ylims=None, xlims=None,
            mean_fontsize='small', mean_fontweight='bold'):
    """Plots a histogram of values."""
    if type(ax)==type(None) or type(fig)==type(None):
        fig, ax = plt.subplots(figsize=figsize)

    # Getting the counts in desired format #
    counts, bins = np.histogram(x, bins=bins)
    logcounts = np.log(counts+1)/np.log(logbase) if log else counts
    if density and type(total)==type(None):
        total = sum(logcounts)
        logcounts = logcounts/total
    elif density:
        logcounts = logcounts/total

    ax.hist(bins[:-1], bins, weights=logcounts, color=color, alpha=alpha,
            label=label)
    ax.set_xlabel(x_label, fp)
    if not density:
        ax.set_ylabel(f'log{round(logbase, 2)}-counts' if log else 'counts', fp)
    else:
        ax.set_ylabel('density-'+f'log{round(logbase, 2)}-counts'
                                                      if log else 'density', fp)
    fig.suptitle(fig_title)

    if add_mean:
        mean = np.mean(x)
        y = ax.get_ylim()[1]*.5
        ax.vlines(mean, 0, y, colors=cutoff_color)
        ax.text(mean, y, f'mean:{round(mean, 4)}', c=cutoff_color,
                fontsize=mean_fontsize, fontweight=mean_fontweight)
    if cutoff:
        y = ax.get_ylim()[1] * .5
        ax.vlines(cutoff, 0, y, colors=cutoff_color)
        ax.text(cutoff, y, f'cutoff: {round(cutoff, 4)}', c=cutoff_color,
                fontsize=mean_fontsize, fontweight=mean_fontweight)

    # Add axes these limits #
    if type(xlims)!=type(None):
        ax.set_xlim(*xlims)
    if type(ylims)!=type(None):
        ax.set_ylim(*ylims)

    # Removing boxes outside #
    ax.spines['top'].set_visible(False)
    ax.spines['right'].set_visible(False)

    if show:
        #vhs.dealWithPlot(type(file_name)!=type(None), True, True, '',
        #                                                        file_name, 300)
        plt.show()
    elif not return_total:
        return fig, ax
    else:
        return fig, ax, total

# %% [code]
fig, ax = distrib(total_counts, color='c', log=True, logbase=10, add_mean=True, show=False)

cutoff = min_
cutoff_color = 'red'
y = ax.get_ylim()[1] * .2
mean_fontsize='small'
mean_fontweight='bold'
ax.vlines(cutoff, 0, y, colors=cutoff_color)
ax.text(cutoff, y, f'minimum: {round(cutoff, 4)}', c=cutoff_color,
        fontsize=mean_fontsize, fontweight=mean_fontweight)

plt.show()

# %% [code]
tenk_ranges = {'t7_umis': [0, 160],
 't7_no_bc_umis': [0, 477],
 't7_all': [0, 494],
 'total_counts': [1137.0, 409262.0],
 'pct_counts_in_top_100_genes': [8.095781071835804, 33.15808340303501],
 'n_genes_by_counts': [890, 15605]}

fiveh_ranges = {'t7_umis': [0, 107],
 't7_no_bc_umis': [0, 157],
 't7_all': [0, 215],
 'total_counts': [6906.0, 241211.0],
 'pct_counts_in_top_100_genes': [11.711418938132185, 45.63162184189079],
 'n_genes_by_counts': [3151, 13221]}

# %% [code]
### Collecting the min and maxes so can set the ranges.
stratified_ranges = {}

for metric in ['t7_umis', 't7_no_bc_umis', 't7_all', 'total_counts', 'pct_counts_in_top_100_genes', 'n_genes_by_counts']:

    measures = data.obs[metric].values

    stratified_ranges[metric] = [999999999, 0]

    tenk_xlims = tenk_ranges[metric]
    fiveh_xlims = fiveh_ranges[metric]
    xlims = (np.min([tenk_xlims[0], fiveh_xlims[0], -1]),
             np.max([tenk_xlims[1], fiveh_xlims[1]]))
    
    for sample in sample_set:
        sample_meaures = measures[sample_labels==sample]

        stratified_ranges[metric] = [np.min([np.min(sample_meaures), stratified_ranges[metric][0]]), 
                                     np.max([np.max(sample_meaures), stratified_ranges[metric][1]])]
        
        fig, ax = distrib(sample_meaures, color=sample_colors[sample], log=True, logbase=10, add_mean=False, 
                              cutoff_color='k', alpha=0.7, show=False, bins=50, xlims=xlims)
        fig.suptitle(f"{sample} {metric}")
        
        cutoff = np.round(min( sample_meaures ), 1)
        cutoff_color = 'black'
        y = ax.get_ylim()[1] * .2
        mean_fontsize='large'
        mean_fontweight='normal'
        ax.vlines(cutoff, 0, y, colors=cutoff_color)
        ax.text(cutoff, y, f'minimum: {round(cutoff, 4)}', c=cutoff_color,
                fontsize=mean_fontsize, fontweight=mean_fontweight)

        cutoff = np.round(np.mean( sample_meaures ), 3)
        y = ax.get_ylim()[1] * .8
        ax.vlines(cutoff, 0, y, colors=cutoff_color)
        ax.text(cutoff, y, f'mean: {round(cutoff, 4)}', c=cutoff_color,
                fontsize=mean_fontsize, fontweight=mean_fontweight)
        
        plt.show()

# %% [code]
stratified_ranges

# %% [code]
out_plots

# %% [markdown]
# Some UMAPs of this stuff

# %% [code]
sc.pl.umap(data, color='sample', show=False)
plt.show()

# %% [code]
!wget https://raw.githubusercontent.com/scverse/scanpy_usage/refs/heads/master/180209_cell_cycle/data/regev_lab_cell_cycle_genes.txt -O {data_dir}regev_lab_cell_cycle_genes.txt

# %% [code]
cycle_genes = [line.strip('\n') for line in open(f"{data_dir}regev_lab_cell_cycle_genes.txt", "r")]
s_genes = cycle_genes[:43]
g2m_genes = cycle_genes[43:]
s_genes, g2m_genes

# %% [code]
sc.tl.score_genes_cell_cycle(data, s_genes, g2m_genes)

# %% [code]
data.obs.keys()

# %% [code]
sc.pl.umap(data, color='S_score')

# %% [code]
sc.pl.umap(data, color='G2M_score')

# %% [code]
sc.pl.umap(data, color='phase')

# %% [code]
sc.pl.umap(data, color='XIST')

# %% [code]
sc.tl.leiden(data, resolution=1.5)

# %% [code]
sc.pl.umap(data, color='leiden')

# %% [code]
import cytocipher as cc

# %% [code]
cc.tl.get_markers(data, 'leiden', var_groups='highly_variable')

# %% [code]
cc.tl.code_enrich(data, 'leiden', n_cpus=10)

# %% [code]
cc.pl.enrich_heatmap(data, 'leiden', figsize=(4,4))

# %% [code]
cc.tl.merge_clusters(data, 'leiden', n_cpus=10, p_cut=.005)

# %% [code]
cc.pl.enrich_heatmap(data, 'leiden_merged', figsize=(4,4), scale_cols=True, show=False)
vhs.dealWithPlot(True, True, True, out_plots, f"leiden-merged_scores_heatmap.png", 300)

# %% [code]
cc.pl.volcano(data, 'leiden', p_cut=.005, p_adjust=True, figsize=(4,4))

# %% [code]
#cc.tl.merge(data, 'leiden', 0.01)

# %% [code]
sc.pl.umap(data, color='leiden_merged', show=False)
vhs.dealWithPlot(True, True, True, out_plots, f"leiden-merged_umap.png", 300)

# %% [code]
cc.pl.sig_cluster_diagnostics(data, 'leiden')

# %% [code]
fig, ax = plt.subplots(ncols=2, figsize=(10,3))
cc.pl.check_abundance_bias(data, 'leiden', p_cut=.045, p_adjust=True, show_legend=False, ax=ax[0], show=False)
cc.pl.check_total_abundance_bias(data, 'leiden', p_cut=.045, p_adjust=True, show_legend=False, ax=ax[1], show=False)
plt.show()

# %% [code]
markers = data.uns['leiden_merged_markers']
markers

# %% [code]
marker_genes = {key: values[0:2] for key, values in markers.items()}

# %% [code]
for clust, genes in marker_genes.items():

    sc.pl.umap(data, color=['leiden_merged']+list(genes), groups=[clust])

# %% [code]
data.layers["scaled"] = sc.pp.scale(data, copy=True).X

# %% [markdown]
# Some UMAPs of this stuff

# %% [code]
data.obsm.keys()

# %% [code]
data.obs['sample'].cat.categories

# %% [code]
data.uns['sample_colors']

# %% [code]
data.uns['sample_colors'] = [sample_colors[sample_] for sample_ in data.obs['sample'].cat.categories]

# %% [code]
data.obs['log10_total_counts'] = np.log10(data.obs['total_counts'].values)

# %% [code]
sc.pl.umap(data, color='log10_total_counts', show=False, size=150, cmap='Spectral_r', vmin=0)
plt.show()

# %% [code]
sc.pl.umap(data, color='phase', show=False)
plt.show()

# %% [code]
#### Selecting some example genes to save and show in the plot...
select_genes = ['BLTP2', 'XIST', 'ARHGAP15', 'CASC15', 'HSP90AA1', 'ENSG00000287409', 'LINC01811', 'DSCAM',
               'CR1L', 'CENPF']
for gene in select_genes:
    sc.pl.umap(data, color=gene, show=False)
    plt.show()

# %% [markdown]
# Now need to compare against the 10k lib, particularly in terms of the off-target edits detected.

# %% [code]
tenk_data = sc.read_h5ad(f'{data_dir2}superb_data.with-guide-imputation.h5ad')
tenk_data

# %% [code]
tenk_edit_info = tenk_data.uns['edit_site_info_guide_homology_and_coinc_final']
tenk_edit_info.head(2)

# %% [code]
tenk_edit_info = tenk_edit_info.loc[tenk_edit_info['manual_edit_call'].values,:]

# %% [code]
fiveh_edit_info = edit_site_info.loc[edit_site_info['manual_edit_call'].values,:]
fiveh_edit_info

# %% [code]
#### Determining the edits that are in-common between the 500 cell lib and the 10k lib.
chrs_ = np.unique( list(tenk_edit_info['chr'].values)+list(fiveh_edit_info['chr']) )
chrs_

# %% [code]
fiveh_edits_paired = [] # fiveh edits that were paired to a 10k
tenk_edits_paired = [] # tenk edit that were paired to a 5h cell
paired_edit_dists = []

dist = 20
for chr_ in chrs_:

    tenk_chr_edits = tenk_edit_info.loc[tenk_edit_info['chr'].values==chr_,:]
    fiveh_chr_edits = fiveh_edit_info.loc[fiveh_edit_info['chr'].values==chr_,:]

    for _, tenk_edit in tenk_chr_edits.iterrows():
        for _, fiveh_edit in fiveh_chr_edits.iterrows():

            dist_ = abs( tenk_edit['pos'] - fiveh_edit['pos'] )

            if dist_ < dist:
                fiveh_edits_paired.append( fiveh_edit['name'] )
                tenk_edits_paired.append( tenk_edit['name'] )
                paired_edit_dists.append( dist_ )

# %% [code]
fiveh_edit_info.shape

# %% [code]
len(fiveh_edits_paired), len(np.unique(fiveh_edits_paired))

# %% [code]
len(tenk_edits_paired), len(np.unique(tenk_edits_paired))

# %% [code]
for i in range(len(fiveh_edits_paired)):
    print(fiveh_edits_paired[i], tenk_edits_paired[i])

# %% [markdown]
***Wow they are essentially at the same position!***

# %% [code]
dist_set = np.unique(paired_edit_dists)
dist_counts = [paired_edit_dists.count(dist_) for dist_ in dist_set]

# %% [code]
dist_counts

# %% [code]
plt.bar(dist_set, dist_counts, color='indianred')
plt.xlabel('Distance between 10k lib edit site and 500 cell lib edit site')
plt.show()

# %% [markdown]
***Looking at the unique edits called***

# %% [code]
fiveh_edits_unique = set(list(fiveh_edit_info['name'].values)).difference( set(fiveh_edits_paired) )
tenk_edits_unique = set(list(tenk_edit_info['name'].values)).difference( set(tenk_edits_paired) )

fiveh_edits_unique, tenk_edits_unique

# %% [code]
from matplotlib_venn import venn2, venn3
?venn2

# %% [code]
venn2(subsets = {'10': len(fiveh_edits_unique), '01': len(tenk_edits_unique), 
                 '11': len(fiveh_edits_paired)}, set_labels = ('500 cell edit sites', '10k cell edit sites'),
     set_colors=['thistle', 'seagreen'])
plt.show()

# %% [markdown]
***Can also break down which of the edit sites were due to the white listing***

# %% [code]
no_reverse_read_edits = fiveh_edit_info['name'].values[ fiveh_edit_info['stranded_edit_dist'].values.astype(str)=='nan' ]
len(no_reverse_read_edits), no_reverse_read_edits

# %% [code]
?venn3

# %% [code]
len(fiveh_edits_paired)-len(no_reverse_read_edits)

# %% [code]
out_plots

# %% [code]
#100, 010, 110, 001, 101, 011, 111
venn3(subsets = {'010': len(fiveh_edits_unique), '001': len(tenk_edits_unique), 
                 '011': len(no_reverse_read_edits),
                '111': len(fiveh_edits_paired)-len(no_reverse_read_edits)}, 
      #set_labels = ('500 cell edit sites - no reverse reads', '500 cell edit sites', '10k cell edit sites'),
      set_labels=('', '', ''), # Will relabel in figure
     set_colors=['lightcoral', 'thistle', 'seagreen'])
plt.show()

# %% [markdown]
***Now seeing what the number of edited cells is like between the two libraries!***

# %% [code]
fiveh_edit_info.index = fiveh_edit_info['name']

# %% [code]
tenk_edits_unique_list = list(tenk_edits_unique)

# %% [code]
(np.array(tenk_edited_cells)/tenk_data.shape[0])*data.shape[0]

# %% [code]
fiveh_edit_info.loc[fiveh_edits_paired,:]

# %% [code]
import scipy.stats

# %% [code]
fiveh_edited_cells = list(fiveh_edit_info.loc[fiveh_edits_paired, 'n_cells_edited'])+[0]*len(tenk_edits_unique)
tenk_edited_cells = list(tenk_edit_info.loc[tenk_edits_paired, 'n_cells_edited'])+\
                    list(tenk_edit_info.loc[tenk_edits_unique_list, 'n_cells_edited'])

print(scipy.stats.pearsonr(fiveh_edited_cells, tenk_edited_cells))

fig, ax = plt.subplots()
ax.scatter(tenk_edited_cells, fiveh_edited_cells, color='seagreen')
plt.plot(tenk_edited_cells, (np.array(tenk_edited_cells)/tenk_data.shape[0])*data.shape[0],
        color='lime')

order = np.argsort(-np.array(fiveh_edited_cells[0:len(fiveh_edits_paired)]))
top_genes = fiveh_edit_info.loc[fiveh_edits_paired, 'intersecting_genes'].values

for i in order[0:5]:
    plt.text(tenk_edited_cells[i], fiveh_edited_cells[i], top_genes[i].split(',')[-1])

ax.spines['top'].set_visible(False)
ax.spines['right'].set_visible(False)

plt.xlabel('10k lib cells edited')
plt.ylabel('500 cell lib cells edited')
vhs.dealWithPlot(True, True, True, out_plots, f"lib_edit-site_edited-cells_scatter.png", 300)

# %% [code]
tenk_edit_info.loc[tenk_edits_unique_list,:]

# %% [code]
##### Every site we DON'T detect is actually at a frequency we DON'T expect to detect, given the total number of cells sequenced.
nondetected_edit_expected_counts = (tenk_edit_info.loc[tenk_edits_unique_list,'n_cells_edited'] / tenk_data.shape[0])*500
nondetected_edit_expected_counts

# %% [code]
detected_edit_expected_counts = (tenk_edit_info.loc[tenk_edits_paired,'n_cells_edited'] / tenk_data.shape[0])*500
detected_edit_expected_counts

# %% [code]
### Could also plot this distribution

#### Non-detected edits
fig, ax = plt.subplots()
ax.hist(nondetected_edit_expected_counts.values, color='red', alpha=0.5, bins=8)
plt.title("Undetected edits in 500 cell lib")
plt.xlabel("Expected edited cells in 500 cell lib | 10k cell lib proportions")

cutoff = np.round(min( nondetected_edit_expected_counts.values ), 1)
cutoff_color = 'black'
y = ax.get_ylim()[1] * .2
mean_fontsize='large'
mean_fontweight='normal'
ax.vlines(cutoff, 0, y, colors=cutoff_color)
ax.text(cutoff, y, f'minimum: {round(cutoff, 4)}', c=cutoff_color,
        fontsize=mean_fontsize, fontweight=mean_fontweight)

cutoff = np.round(np.mean( nondetected_edit_expected_counts.values ), 1)
y = ax.get_ylim()[1] * .8
ax.vlines(cutoff, 0, y, colors=cutoff_color)
ax.text(cutoff, y, f'mean: {round(cutoff, 4)}', c=cutoff_color,
        fontsize=mean_fontsize, fontweight=mean_fontweight)
plt.show()

#### Detected edits
fig, ax = plt.subplots()
ax.hist(detected_edit_expected_counts.values, color='seagreen', alpha=0.5, bins=30)
plt.title("Detected edits in 500 cell lib")
plt.xlabel("Expected edited cells in 500 cell lib | 10k cell lib proportions")

cutoff = np.round(min( detected_edit_expected_counts.values ), 1)
cutoff_color = 'black'
y = ax.get_ylim()[1] * .2
mean_fontsize='large'
mean_fontweight='normal'
ax.vlines(cutoff, 0, y, colors=cutoff_color)
ax.text(cutoff, y, f'minimum: {round(cutoff, 4)}', c=cutoff_color,
        fontsize=mean_fontsize, fontweight=mean_fontweight)

cutoff = np.round(np.mean( detected_edit_expected_counts.values ), 1)
y = ax.get_ylim()[1] * .8
ax.vlines(cutoff, 0, y, colors=cutoff_color)
ax.text(cutoff, y, f'mean: {round(cutoff, 4)}', c=cutoff_color,
        fontsize=mean_fontsize, fontweight=mean_fontweight)
plt.show()

# %% [markdown]
***Now seeing the relationship between the number of edited alleles across cells between the two libraries***

# %% [code]
fiveh_edit_info

# %% [code]
fiveh_edit_info['total_alleles'] = fiveh_edit_info['copy-number'].values * data.shape[0]
fiveh_edit_info

# %% [code]
fiveh_edit_info['total_edited_alleles'] = data.obsm['dosages_manual-edits-only'].loc[:,fiveh_edit_info.index.values].sum(axis=0)
fiveh_edit_info

# %% [code]
tenk_edit_info['total_alleles'] = tenk_edit_info['copy-number'].values * tenk_data.shape[0]
tenk_edit_info['total_edited_alleles'] = tenk_data.obsm['dosages'].loc[:,tenk_edit_info.index.values].sum(axis=0)

# %% [code]
tenk_edit_info

# %% [code]
fiveh_edited_alleles = list(fiveh_edit_info.loc[fiveh_edits_paired, 'total_edited_alleles'])+[0]*len(tenk_edits_unique)
tenk_edited_alleles = list(tenk_edit_info.loc[tenk_edits_paired, 'total_edited_alleles'])+\
                    list(tenk_edit_info.loc[tenk_edits_unique_list, 'total_edited_alleles'])

print(scipy.stats.pearsonr(fiveh_edited_alleles, tenk_edited_alleles))

order = np.argsort(-np.array(fiveh_edited_alleles[0:len(fiveh_edits_paired)]))
top_genes = fiveh_edit_info.loc[fiveh_edits_paired, 'intersecting_genes'].values

fig, ax = plt.subplots()
ax.scatter(tenk_edited_alleles, fiveh_edited_alleles, color='dodgerblue')
ax.plot(tenk_edited_alleles, (np.array(tenk_edited_alleles)/tenk_data.shape[0])*data.shape[0],
        color='cyan')

for i in order[0:5]:
    plt.text(tenk_edited_alleles[i], fiveh_edited_alleles[i], top_genes[i].split(',')[-1])

ax.spines['top'].set_visible(False)
ax.spines['right'].set_visible(False)

plt.xlabel('10k lib alleles edited')
plt.ylabel('500 cell lib alleles edited')
plt.show()

# %% [markdown]
# Creating and comparing and edit-site co-occurence matrix...

# %% [code]
full_edits = tenk_data.uns['edit_site_info_guide_homology_and_coinc_final']

final_edits = full_edits.index.values[ full_edits['manual_edit_call'].values ]
len(final_edits), final_edits

# %% [code]
final_edits_wGenes = [f"{final_edit} ({genes})" for final_edit, genes in zip(final_edits, full_edits.loc[final_edits,'intersecting_genes'])]

# %% [code]
final_edits_wGenes

# %% [code]
dosages = tenk_data.obsm['dosages'].loc[:, final_edits]
dosages

# %% [code]
import scipy.stats

# %% [code]
def pearsons(x,y):
    return scipy.stats.pearsonr(x,y)[0]

# %% [code]
edit_pearsons = np.zeros((len(final_edits), len(final_edits)))
for i in range(len(final_edits)):
    for j in range(len(final_edits)):

        edit_pearsons[i,j] = scipy.stats.pearsonr(dosages.values[:,i],dosages.values[:,j])[0]

edit_pearsons = pd.DataFrame(edit_pearsons, index=final_edits_wGenes, columns=final_edits_wGenes)

# %% [code]
g = sb.clustermap(edit_pearsons, vmax=.2, vmin=-.2, cmap='Spectral_r')
g.ax_heatmap.set_xticks(np.array(list(range(edit_pearsons.shape[0])))+.5, g.data2d.index.values)
g.ax_heatmap.set_yticks(np.array(list(range(edit_pearsons.shape[0])))+.5, g.data2d.columns.values)

vhs.dealWithPlot(True, True, True, out_plots, "10k-lib_dosage-correlation_heatmap.png", 300)

# %% [code]
data.obsm.keys()

# %% [code]
len(real_edits)

# %% [code]
dosages = data.obsm['dosages'].loc[:,real_edits]
dosages

# %% [code]
data.uns.keys()

# %% [code]
fiveh_edit_site_info = data.uns['edit_site_info']
fiveh_edit_site_info.index = fiveh_edit_site_info['name']

final_edits_wGenes = [f"{final_edit} ({genes})" for final_edit, genes in zip(real_edits, fiveh_edit_site_info.loc[real_edits,'intersecting_genes'])]

# %% [code]
edit_pearsons = np.zeros((len(real_edits), len(real_edits)))
for i in range(len(real_edits)):
    for j in range(len(real_edits)):

        edit_pearsons[i,j] = scipy.stats.pearsonr(dosages.values[:,i],dosages.values[:,j])[0]

edit_pearsons = pd.DataFrame(edit_pearsons, index=final_edits_wGenes, columns=final_edits_wGenes)

# %% [code]
g = sb.clustermap(edit_pearsons, vmax=.2, vmin=-.2, cmap='Spectral_r')
g.ax_heatmap.set_xticks(np.array(list(range(edit_pearsons.shape[0])))+.5, g.data2d.index.values)
g.ax_heatmap.set_yticks(np.array(list(range(edit_pearsons.shape[0])))+.5, g.data2d.columns.values)

vhs.dealWithPlot(True, True, True, out_plots, "500-cell-lib_dosage-correlation_heatmap.png", 300)

# %% [code]
out_plots

# %% [markdown]
# Saving the cell metadata to include with the GEO submission.

# %% [code]
data.obs.columns

# %% [code]
keep_cols = ['sample', 'n_genes_by_counts', 'log1p_n_genes_by_counts',
       'total_counts', 'log1p_total_counts', 'pct_counts_in_top_50_genes',
       'pct_counts_in_top_100_genes', 'pct_counts_in_top_200_genes',
       'pct_counts_in_top_500_genes', 'mito_perc',
       't7_all', 't7_no_bc_umis', 't7_umis', 'S_score', 'G2M_score', 'phase',
       'leiden', 'leiden_merged', 'log10_total_counts', 'edit_cell_labels']

cell_meta = data.obs.loc[:,keep_cols]
cell_meta

# %% [code]
data.obsm.keys()

# %% [code]
cell_meta['UMAP_1'] = data.obsm['X_umap'][:,0]
cell_meta['UMAP_2'] = data.obsm['X_umap'][:,1]

cell_meta

# %% [code]
!mkdir {out_dir}

# %% [code]
cell_meta.to_csv(f'{out_dir}cell_metadata_500_cell.tsv.gz', sep='\t')

# %% [code]
!zcat {out_dir}cell_metadata_500_cell.tsv.gz | head

# %% [code]
print(f"{out_dir}cell_metadata_500_cell.tsv.gz")

# %% [code]
data.uns['edit_site_info']

# %% [code]
site_specific_umi_count = data.obsm['t7_umis'].sum(axis=0).reset_index()
site_specific_umi_count.index = site_specific_umi_count['index']
site_specific_umi_count.columns = ['index', 't7_barcoded_umis']
site_specific_umi_count

# %% [code]
t7_umis = data.obsm['t7_umis']

site_specific_mean_umis = []
for i, edit_site in enumerate(t7_umis.columns):

    edit_site_counts = t7_umis.values[:,i]
    nonzero_bool = edit_site_counts > 0

    mean_ = np.mean( edit_site_counts[nonzero_bool] )
    site_specific_mean_umis.append( mean_ )

# %% [code]
site_specific_umi_count.loc[t7_umis.columns.values, 't7_barcoded_umis_per_edited_cell'] = site_specific_mean_umis

# %% [code]
site_specific_umi_count

# %% [code]
edit_site_info = data.uns['edit_site_info']
edit_site_info

# %% [code]
site_specific_umi_count = site_specific_umi_count.loc[edit_site_info.index.values]

# %% [code]
site_specific_umi_count.index.values == edit_site_info.index.values

# %% [code]
edit_site_info['t7_barcoded_umis'] = site_specific_umi_count['t7_barcoded_umis'].values
edit_site_info['t7_barcoded_umis_per_edited_cell'] = site_specific_umi_count['t7_barcoded_umis_per_edited_cell'].values

# %% [code]
edit_site_info.to_csv(f'{out_dir}edit_site_metadata_500_cell.tsv', sep='\t', index=False)

# %% [code]
!cat {out_dir}edit_site_metadata_500_cell.tsv

# %% [code]
```
