## Additional Files for "Joint single-cell capture of Cas9 edits and transcriptomes reveals on- and off-target effects on gene expression": cell_qc_10k-lib.ipynb

```
# %% [markdown]
# Looking at the cell QC metrics, may also compare to the 500 cell library from this notebook, but not yet.
                                          

# INPUT:

    * superb_data.with-guide-imputation.h5ad

# Environment Setup

# %% [code]
work_dir = './'
import os
os.chdir( work_dir )

import numpy as np
import pandas as pd

import scanpy as sc

import matplotlib.pyplot as plt

data_dir = 'superb_count_all/combined/'
out_dir = data_dir+'cell_qc_tables/'

# %% [markdown]
# Loading the data

# %% [code]
data = sc.read_h5ad(f'{data_dir}superb_data.with-guide-imputation.h5ad')
data

# %% [code]
data.var.keys()

# %% [code]
##### Counting the mitochondrial reads per cell.
mito_genes = [gene for gene in data.var_names if gene.startswith('MT-')]
mito_counts = data[:,mito_genes].layers['gene_counts'].toarray().sum(axis=1)
mito_perc = (mito_counts.astype(float) / data.obs['total_counts'].values)*100

data.obs['mito_perc'] = mito_perc

# %% [code]
sum(mito_counts)

# %% [markdown]
***No mitochondrial read counts..***

# %% [markdown]
***Let's count the t7 barcoded reads!***

# %% [code]
data.obsm.keys()

# %% [code]
data.uns.keys()

# %% [code]
#### Subsetting the t7 reads to just the confident edit calls!
edit_info = data.uns['edit_site_info_guide_homology_and_coinc_final']
edit_info

# %% [code]
conf_bool = edit_info[ 'manual_edit_call' ].values

conf_edits = edit_info.index.values[ conf_bool ]
len( conf_edits )

# %% [code]
data.obsm['t7_umis'].iloc[0:3,0:3]

# %% [code]
t7_umis = data.obsm['t7_umis'].loc[:,conf_edits].sum(axis=1).values

# %% [code]
t7_umis

# %% [code]
data.obs['t7_umis'] = t7_umis

# %% [code]
data.obs['t7_all'] = data.obsm['t7_all'].loc[:,conf_edits].sum(axis=1).values

# %% [code]
data.obs['t7_no_bc_umis'] = data.obsm['t7_no_bc_umis'].loc[:,conf_edits].sum(axis=1).values

# %% [markdown]
# Per cell metrics

# %% [code]
def pie(labels: np.array, label_set: np.array=None,
        colors: dict=None, min_perc: float=3, autopct: str='',
        labeldistance=1.05, explode_f=0.8,
        filter_perc=0, show_perc=False, show_size=False, print_filt=False, figsize=(5,5),
        show=True):
    """Piechart, but formatted nicely, e.g. will order pies & explode out
        low frequency labels.

    Args:

        min_perc (float): Percentage below which labels will 'explode'.
        filter_perc (float): Percentage below which labels will not be shown.

    """
    if type(label_set)==type(None):
        label_set = np.unique(labels)

    sizes = np.array([len(np.where(labels == dataset)[0])
                      for dataset in label_set])
    percs = np.array([(size / sum(sizes)) * 100 for size in sizes])

    if filter_perc > 0:
        perc_filt = percs > filter_perc
        if print_filt:
            print( label_set[perc_filt==False] )

        remain_size = sum(sizes[perc_filt == False])
        sizes = np.array(list(sizes[perc_filt]) + [remain_size])
        remain_perc = sum(percs[perc_filt == False])
        percs = np.array(list(percs[perc_filt]) + [remain_perc])

        label_set = np.array(list(label_set[perc_filt]) + [f'other'])

    if show_perc:
        label_set = np.array([f'{label} ({round(perc, 2)}%)'
                              for label, perc in zip(label_set, percs)])
    elif show_size:
        label_set = np.array([f'{label} (n={round(size_, 2)})'
                              for label, size_ in zip(label_set, sizes)])

    order = np.argsort(-sizes)
    sizes = sizes[order]
    label_set = label_set[order]
    percs = percs[order]

    ### Getting the colors, will add this to scripts.helpers so consistent ###
    if type(colors)==type(None):
        colors = vhs.getColors(label_set)

    colors_ = [colors[dataset] for dataset in label_set]
    end = sum(percs<min_perc)
    explode = [0] * (len(label_set) - end) + [explode_f * (i + 1) for i in list(
        range(end))]  # Expand out less numerous data...

    fig1, ax1 = plt.subplots(figsize=figsize)
    ax1.pie(sizes, labels=label_set, autopct=autopct,  # '%0.1f%%',
            shadow=False, startangle=90, colors=colors_, explode=explode,
            labeldistance=labeldistance)
    ax1.axis('equal')  # Equal aspect ratio ensures that pie is drawn as a circle.

    if show:
        plt.show()
    else:
        return fig1, ax1

# %% [code]
#### Pie chart of the cells captured in the different samples
sample_colors = {'nurd_ki': 'palegreen', 'baf_ki': 'lightskyblue', 'no_ki': 'magenta'}
sample_colors_wSizes = {'nurd_ki (n=5086)': sample_colors['nurd_ki'], 
                        'baf_ki (n=3730)': sample_colors['baf_ki'], 
                        'no_ki (n=684)': sample_colors['no_ki']}

pie(data.obs['sample'].values, colors=sample_colors_wSizes, show_size=True, show=False)
plt.show()

# %% [markdown]
# Let's also get the proportion of these that have the on-target edit in each cell, and evidence of off-target in different combinations..

# %% [code]
sample_labels = data.obs['sample'].values

dosages_genes = data.obsm['dosages_corrected_genes_manual-edits-only']
arid1a_edited = dosages_genes['ARID1A'].values > 0
smarca4_edited = dosages_genes['SMARCA4'].values > 0
chd3_edited = dosages_genes['CHD3'].values > 0
chd4_edited = dosages_genes['CHD4'].values > 0

any_edited = dosages_genes.sum(axis=1) > 0

# %% [code]
np.unique( sample_labels )

# %% [code]
len(any_edited)

# %% [code]
def double_label(edit_bool_1, edit_bool_2, gene_1, gene_2, any_edited_bool):
    
    cell_label = 'no-edit'
    if edit_bool_1 and not edit_bool_2:
        cell_label = f'{gene_1}-edit'
    elif not edit_bool_1 and edit_bool_2:
        cell_label = f'{gene_2}-edit'
    elif edit_bool_1 and edit_bool_2:
        cell_label = f'double-edit'

    if any_edited_bool and cell_label!='no-edit':
        cell_label = cell_label + ', off-target/s'
    elif any_edited_bool:
        cell_label = 'off-target/s'
        
    return cell_label

# %% [code]
cell_labels = []
for i in range(len(sample_labels)):

    if sample_labels[i] == 'no_ki' and any_edited[i]:
        cell_labels.append( 'false-edit-call' )

    elif sample_labels[i] == 'no_ki':
        cell_labels.append( 'no-edit' )

    elif sample_labels[i] == 'baf_ki':
        cell_labels.append( double_label(arid1a_edited[i], smarca4_edited[i], 'ARID1A', 'SMARCA4', any_edited[i]) )

    elif sample_labels[i] == 'nurd_ki':
        cell_labels.append( double_label(chd3_edited[i], chd4_edited[i], 'CHD3', 'CHD4', any_edited[i]) )

# %% [code]
np.unique(cell_labels)

# %% [code]
data.obs['edit_cell_labels'] = cell_labels
data.obs['edit_cell_labels'] = data.obs['edit_cell_labels'].astype('category')

# %% [code]
def get_overlap_counts(data, groupby1: str, groupby2: str,
                       uns_key: str = 'annot_overlaps',
                       verbose: bool = True):
    """ Calculates no. of cells belonging to the different categories of groupby1 and groupby2.
    """

    labels = data.obs[groupby1].values.astype(str)
    label_set = np.array(list(data.obs[groupby1].cat.categories))

    labels2 = data.obs[groupby2].values.astype(str)
    label_set2 = np.array(list(data.obs[groupby2].cat.categories))

    overlap_counts = np.zeros((len(label_set), len(label_set2)))
    for i, label in enumerate(label_set):
        label_bool = labels == label
        for j, label2 in enumerate(label_set2):
            label_bool2 = labels2 == label2

            overlap_counts[i, j] = len(
                np.where(np.logical_and(label_bool, label_bool2))[0])

    overlap_counts_df = pd.DataFrame(overlap_counts, index=label_set,
                                     columns=label_set2)
    data.uns[uns_key] = overlap_counts_df
    if verbose:
        print(f"Added data.uns['{uns_key}']")

# %% [code]
get_overlap_counts(data, 'sample', 'edit_cell_labels')

# %% [code]
data.uns['annot_overlaps']

# %% [code]
#### Want to see the actual proportions:
totals = data.uns['annot_overlaps'].values.sum(axis=1)
proportions = np.apply_along_axis(np.divide, 0, data.uns['annot_overlaps'].values, totals)

pd.DataFrame(proportions, index=data.uns['annot_overlaps'].index, columns=data.uns['annot_overlaps'].columns)

# %% [code]
proportions.sum(axis=1)

# %% [code]
sc.pl.umap(data, color='edit_cell_labels')

# %% [code]
import numpy
import matplotlib

def getOrderedLabelSet(labels):
	""" Gets the set of labels associated with labels ordered from most to \
		least frequent.
	"""
	labelSet = list(set(labels))
	freqs = numpy.array([len(numpy.where(labels == label)[0])
						 for label in labelSet])
	order = numpy.array(list((-freqs).argsort()))
	labelSetOrdered = list(numpy.array(labelSet)[order])

	return labelSetOrdered

def getColors(labels, labelSet=None, colorMap='tab20', rgb=False):
	""" Gets an OrderedDict of colors; the order indicates the frequency of \
	labels from largest to smallest.

	Args:
		labels (numpy.array<str>): Indicates a set of labels for observations.

		labelSet (list-like<str>): Indicates the set of labels in labels. \
									If None, calculated based on labels.

		colorMap (str): A matplotlib colormap.

		rgb (bool): If True, colors indicated by rgb value, if false hexcode.

	Returns:
		dict<str, tuple or str>: An ordered dict indicating the labels which \
					occur most to least frequently and the associated colors.
	"""
	# Determining the set of labels #
	labelSet = input.returnDefaultIfNone(labelSet,
										 getOrderedLabelSet(labels))

	# Initialising the ordered dict #
	cellTypeColors = {}

	# Ordering the cells according to their frequency and obtaining colors #
	nLabels = len(labelSet)
	cmap = plt.cm.get_cmap(colorMap, nLabels)
	rgbs = [cmap(i)[:3] for i in range(nLabels)]
	#rgbs = list(numpy.array(rgbs)[order]) # Make sure color order is the same.

	# Populating the color dictionary with rgb values or hexcodes #
	for i in range(len(labelSet)):
		cellType = labelSet[i]
		rgbi = rgbs[i]
		if not rgb:
			cellTypeColors[cellType] = matplotlib.colors.rgb2hex(rgbi)
		else:
			cellTypeColors[cellType] = rgbi

	return cellTypeColors

# %% [code]
color_dict = getColors(cell_labels)
data.uns['edit_cell_labels_colors'] = [color_dict[label_] for label_ in data.obs['edit_cell_labels'].cat.categories]

# %% [code]
data.obs['edit_cell_labels'].cat.categories

# %% [code]
color_dict = getColors(cell_labels)

color_dict['no-edit'] = 'silver'
color_dict['false-edit-call'] = 'dimgrey'
color_dict['off-target/s'] = 'sienna'#'crimson'
color_dict['CHD3-edit, off-target/s'] = 'yellow'
color_dict['CHD4-edit, off-target/s'] = 'goldenrod'
color_dict['ARID1A-edit, off-target/s'] = 'lime' #'#20DF7F'
color_dict['SMARCA4-edit, off-target/s'] = 'green' #'#45BA45'
color_dict['double-edit, off-target/s'] = 'palegoldenrod'
data.uns['edit_cell_labels_colors'] = [color_dict[label_] for label_ in data.obs['edit_cell_labels'].cat.categories]

# %% [code]
color_dict

# %% [code]
def dataframe_annot_overlap_barplot(overlap_counts_df, groupby1, groupby2, colors=None,
                          bar_width: float = .85,
                          labelset1 = None, labelset2 = None,
                          figsize=(10, 5), scale: bool = True,
                          transpose: bool = False, edge_color=None,
                          remove_labelling: bool = False):
    """Plots relationship between annotations by showing proportions of each type found within
        each of the other categories.
    """
    if transpose:  ##### Looking at relationship in opposite direction.
        overlap_counts_df = overlap_counts_df.transpose()
        tmp = groupby1
        groupby1 = groupby2
        groupby2 = tmp

    if type(labelset1)==type(None):
        labelset1 = overlap_counts_df.index.values
    else:
        overlap_counts_df = overlap_counts_df.loc[labelset1, :]

    if type(labelset2) == type(None):
        labelset2 = overlap_counts_df.columns.values
    else:
        overlap_counts_df = overlap_counts_df.loc[:, labelset2]

    ##### Getting colors
    if type(colors)==type(None):
        colors = {val: 'grey' for i, val in enumerate(labelset2)}

    ##### Scaling the data to proportions.
    if scale:
        total_counts = overlap_counts_df.sum(axis=1)
        overlap_counts = np.apply_along_axis(np.divide, 0,
                                             overlap_counts_df.values,
                                             total_counts)
    else:
        overlap_counts = overlap_counts_df.values

    ##### Creating the barplots...
    fig, axs = plt.subplots(figsize=figsize)
    col_locs = list(range(len(labelset1)))
    for labeli, label in enumerate(labelset2):
        props = overlap_counts[:, labeli]

        if labeli == 0:
            bottom = [0] * len(props)

        axs.bar(col_locs, props, bottom=bottom,
                color=colors[label], edgecolor=edge_color,
                width=bar_width, )
        bottom = list(np.array(bottom) + props)

        # Custom x axis
        axs.set_xticks(list(range(len(labelset1))), list(labelset1),
                       rotation=90)
        axs.spines['right'].set_visible(False)
        axs.spines['top'].set_visible(False)
        if remove_labelling:
            axs.set_xticks([])
            axs.set_yticks([])
            axs.spines['left'].set_visible(False)
            axs.spines['bottom'].set_visible(False)
        else:
            axs.set_xlabel(f"{groupby1} Clusters")

def annot_overlap_barplot(data, groupby1: str, groupby2: str,
                          uns_key: str = 'annot_overlaps',
                          labelset1 = None, labelset2 = None,
                          bar_width: float = .85,
                          figsize=(10, 5), scale: bool = True,
                          transpose: bool = False, edge_color=None,
                          remove_labelling: bool = False):
    """Plots relationship between annotations by showing proportions of each type found within
        each of the other categories. Written to work with AnnData input.
    """
    if uns_key not in data.uns:
        print("Need to run get_overlap_counts() first.")

    overlap_counts_df = data.uns[uns_key].copy()

    # ##### Getting colors
    if f'{groupby2}_colors' in data.uns:
        colors = {val: data.uns[f'{groupby2}_colors'][i]
                  for i, val in enumerate(data.obs[groupby2].cat.categories)}
    else:
        colors = {val: 'grey' for
                  i, val in enumerate(data.obs[groupby2].cat.categories)}


    dataframe_annot_overlap_barplot(overlap_counts_df, groupby1, groupby2, colors=colors,
                          bar_width = bar_width,
                          labelset1 = labelset1, labelset2 = labelset2,
                          figsize=figsize, scale=scale,
                          transpose=transpose, edge_color=edge_color,
                          remove_labelling=remove_labelling)

# %% [code]
annot_overlap_barplot(data, 'sample', 'edit_cell_labels',
                         labelset1=np.array(['no_ki', 'baf_ki', 'nurd_ki']),
                         labelset2=np.array(['no-edit', 'false-edit-call',
                                            'ARID1A-edit, off-target/s', 'SMARCA4-edit, off-target/s',
                                            'CHD3-edit, off-target/s', 'CHD4-edit, off-target/s',
                                             'double-edit, off-target/s',
                                             'off-target/s'
                                            ][::-1])
                         )
plt.show()

# %% [markdown]
# Now for the proportion of allele calls of each on-target, to understand allelic dosages.

# %% [code]
on_targets = ['CHD3', 'CHD4', 'ARID1A', 'SMARCA4']
samples_ = ['nurd_ki', 'nurd_ki', 'baf_ki', 'baf_ki']
colors_  = ['yellow', 'goldenrod', '#20DF7F', '#45BA45']

# Constructing own overlaps...
allelic_counts = np.zeros((len(on_targets), 3))

for i, (sample_, gene_) in enumerate( zip(samples_, on_targets) ):

    color_ = colors_[i]
    data.obs[f"{gene_}_edit_dosages"] = dosages_genes[gene_].values.astype(str)
    data.obs[f"{gene_}_edit_dosages"] = data.obs[f"{gene_}_edit_dosages"].astype('category')

    get_overlap_counts(data, 'edit_cell_labels', f"{gene_}_edit_dosages")

    # This part only works cause no-cases where have a pure edit with no-off-targets detected.
    allele_counts = data.uns['annot_overlaps'].loc[f'{gene_}-edit, off-target/s'].values[1:]
    alleles = data.uns['annot_overlaps'].columns.values[1:]

    allelic_counts[i,0:len(alleles)] = allele_counts
    
    plt.bar(alleles, allele_counts,
       color=color_)
    plt.xlabel('allelic edits')
    plt.ylabel('n cells')

    plt.show()

# %% [code]
allelic_counts = pd.DataFrame(allelic_counts, index=on_targets, columns=list(range(1,allelic_counts.shape[1]+1)))
allelic_counts

# %% [code]
data.uns['annot_overlaps'] = allelic_counts

# %% [code]
totals = data.uns['annot_overlaps'].values.sum(axis=1)
proportions = np.apply_along_axis(np.divide, 0, data.uns['annot_overlaps'].values, totals)

pd.DataFrame(proportions, index=data.uns['annot_overlaps'].index, columns=data.uns['annot_overlaps'].columns)

# %% [code]
data.uns.keys()

# %% [code]
np.unique([-1]+list(allelic_counts.columns.values.astype(int)))

# %% [code]
allele_colors = getColors(np.unique([0]+list(allelic_counts.columns.values.astype(int))), colorMap='Blues')
allele_colors

# %% [code]
dataframe_annot_overlap_barplot(allelic_counts, 'on-targets', 'alleles-edited', colors=allele_colors)
plt.show()

# %% [code]
#### Some stats on the total counts per cell!
total_counts = data.obs['total_counts'].values

min_, median, max_ = np.quantile(total_counts, [0, 0.5, 1])

print("Min, median, max:", min_, median, max_)

# %% [code]
sample_labels = data.obs['sample'].values
sample_set = np.unique(sample_labels)

# %% [code]
def distrib(x, bins=100, x_label='', fig_title='', log=False, density=False,
            figsize=(6.4,4.8), file_name=None, add_mean=False, logbase=np.e,
            color='blue', alpha=1, ax=None, fig=None, show=True, cutoff=None,
            cutoff_color='r',
            label='', total=None, return_total=False, ylims=None, xlims=None,
            mean_fontsize='small', mean_fontweight='bold'):
    """Plots a histogram of values."""
    if type(ax)==type(None) or type(fig)==type(None):
        fig, ax = plt.subplots(figsize=figsize)

    # Getting the counts in desired format #
    counts, bins = np.histogram(x, bins=bins)
    logcounts = np.log(counts+1)/np.log(logbase) if log else counts
    if density and type(total)==type(None):
        total = sum(logcounts)
        logcounts = logcounts/total
    elif density:
        logcounts = logcounts/total

    ax.hist(bins[:-1], bins, weights=logcounts, color=color, alpha=alpha,
            label=label)
    ax.set_xlabel(x_label, fp)
    if not density:
        ax.set_ylabel(f'log{round(logbase, 2)}-counts' if log else 'counts', fp)
    else:
        ax.set_ylabel('density-'+f'log{round(logbase, 2)}-counts'
                                                      if log else 'density', fp)
    fig.suptitle(fig_title)

    if add_mean:
        mean = np.mean(x)
        y = ax.get_ylim()[1]*.5
        ax.vlines(mean, 0, y, colors=cutoff_color)
        ax.text(mean, y, f'mean:{round(mean, 4)}', c=cutoff_color,
                fontsize=mean_fontsize, fontweight=mean_fontweight)
    if cutoff:
        y = ax.get_ylim()[1] * .5
        ax.vlines(cutoff, 0, y, colors=cutoff_color)
        ax.text(cutoff, y, f'cutoff: {round(cutoff, 4)}', c=cutoff_color,
                fontsize=mean_fontsize, fontweight=mean_fontweight)

    # Add axes these limits #
    if type(xlims)!=type(None):
        ax.set_xlim(*xlims)
    if type(ylims)!=type(None):
        ax.set_ylim(*ylims)

    # Removing boxes outside #
    ax.spines['top'].set_visible(False)
    ax.spines['right'].set_visible(False)

    if show:
        #vhs.dealWithPlot(type(file_name)!=type(None), True, True, '',
        #                                                        file_name, 300)
        plt.show()
    elif not return_total:
        return fig, ax
    else:
        return fig, ax, total

# %% [code]
fig, ax = distrib(total_counts, color='c', log=True, logbase=10, add_mean=True, show=False)

cutoff = min_
cutoff_color = 'red'
y = ax.get_ylim()[1] * .2
mean_fontsize='large'
mean_fontweight='bold'
ax.vlines(cutoff, 0, y, colors=cutoff_color)
ax.text(cutoff, y, f'minimum: {round(cutoff, 4)}', c=cutoff_color,
        fontsize=mean_fontsize, fontweight=mean_fontweight)

plt.show()

# %% [code]
edit_info = data.uns['edit_site_info_guide_homology_and_coinc_final']
edit_info.columns

# %% [code]
on_targets = ['CHD3', 'CHD4', 'ARID1A', 'SMARCA4']

on_target_edits = [edit_row['name'] for i, edit_row in edit_info.iterrows() if np.any([on_target in edit_row['intersecting_genes'] for on_target in on_targets])]
on_target_edits

# %% [code]
edit_info.loc[on_target_edits,['intersecting_genes', 'offset', 'abs_offset', 'homol_guide_name']]

# %% [code]
tenk_ranges = {'t7_umis': [0, 160],
 't7_no_bc_umis': [0, 477],
 't7_all': [0, 494],
 'total_counts': [1137.0, 409262.0],
 'pct_counts_in_top_100_genes': [8.095781071835804, 33.15808340303501],
 'n_genes_by_counts': [890, 15605]}

fiveh_ranges = {'t7_umis': [0, 107],
 't7_no_bc_umis': [0, 157],
 't7_all': [0, 215],
 'total_counts': [6906.0, 241211.0],
 'pct_counts_in_top_100_genes': [11.711418938132185, 45.63162184189079],
 'n_genes_by_counts': [3151, 13221]}

# %% [code]
### Collecting the min and maxes so can set the ranges.
stratified_ranges = {}

for metric in ['t7_umis', 't7_no_bc_umis', 't7_all', 'total_counts', 'pct_counts_in_top_100_genes', 'n_genes_by_counts']:

    measures = data.obs[metric].values

    stratified_ranges[metric] = [999999999, 0]

    tenk_xlims = tenk_ranges[metric]
    fiveh_xlims = fiveh_ranges[metric]
    xlims = (np.min([tenk_xlims[0], fiveh_xlims[0], -1]),
             np.max([tenk_xlims[1], fiveh_xlims[1]]))
    
    for sample in sample_set:
        sample_meaures = measures[sample_labels==sample]

        stratified_ranges[metric] = [np.min([np.min(sample_meaures), stratified_ranges[metric][0]]), 
                                     np.max([np.max(sample_meaures), stratified_ranges[metric][1]])]
        
        fig, ax = distrib(sample_meaures, color=sample_colors[sample], log=True, logbase=10, add_mean=False, 
                              cutoff_color='k', alpha=0.7, show=False, bins=50, xlims=xlims)
        fig.suptitle(f"{sample} {metric}")
        
        cutoff = np.round(min( sample_meaures ), 1)
        cutoff_color = 'black'
        y = ax.get_ylim()[1] * .2
        mean_fontsize='large'
        mean_fontweight='normal'
        ax.vlines(cutoff, 0, y, colors=cutoff_color)
        ax.text(cutoff, y, f'minimum: {round(cutoff, 4)}', c=cutoff_color,
                fontsize=mean_fontsize, fontweight=mean_fontweight)

        cutoff = np.round(np.mean( sample_meaures ), 3)
        y = ax.get_ylim()[1] * .8
        ax.vlines(cutoff, 0, y, colors=cutoff_color)
        ax.text(cutoff, y, f'mean: {round(cutoff, 4)}', c=cutoff_color,
                fontsize=mean_fontsize, fontweight=mean_fontweight)
        
        plt.show()

# %% [code]
stratified_ranges

# %% [markdown]
# Some UMAPs of this stuff

# %% [code]
data.obsm.keys()

# %% [code]
data.obs['sample'].cat.categories

# %% [code]
data.uns['sample_colors']

# %% [code]
data.uns['sample_colors'] = [sample_colors[sample_] for sample_ in data.obs['sample'].cat.categories]

# %% [code]
?sc.pl.umap

# %% [code]
sc.pl.umap(data, color='sample', show=False, size=13)
plt.show()

# %% [code]
data.obs.keys()

# %% [code]
data.obs['log10_total_counts'] = np.log10(data.obs['total_counts'].values)

# %% [code]
sc.pl.umap(data, color='log10_total_counts', show=False, size=13, cmap='Spectral_r', vmin=0)
plt.show()

# %% [code]
!wget https://raw.githubusercontent.com/scverse/scanpy_usage/refs/heads/master/180209_cell_cycle/data/regev_lab_cell_cycle_genes.txt -O {data_dir}regev_lab_cell_cycle_genes.txt

# %% [code]
cycle_genes = [line.strip('\n') for line in open(f"{data_dir}regev_lab_cell_cycle_genes.txt", "r")]
s_genes = cycle_genes[:43]
g2m_genes = cycle_genes[43:]
s_genes, g2m_genes

# %% [code]
sc.tl.score_genes_cell_cycle(data, s_genes, g2m_genes)

# %% [code]
data.obs.keys()

# %% [code]
sc.pl.umap(data, color='S_score', show=False)
plt.show()

# %% [code]
sc.pl.umap(data, color='G2M_score', show=False)
plt.show()

# %% [code]
sc.pl.umap(data, color='phase', show=False)
plt.show()

# %% [code]
sc.pl.umap(data, color='XIST')
plt.show()

# %% [code]
sc.tl.leiden(data, resolution=1)

# %% [code]
sc.pl.umap(data, color='leiden')

# %% [code]
!pip install cytocipher

# %% [code]
import cytocipher as cc

# %% [code]
cc.tl.get_markers(data, 'leiden', var_groups='highly_variable')

# %% [code]
cc.tl.code_enrich(data, 'leiden', n_cpus=10)

# %% [code]
cc.pl.enrich_heatmap(data, 'leiden', figsize=(4,4))

# %% [code]
cc.tl.merge_clusters(data, 'leiden', n_cpus=10, p_cut=.01)

# %% [code]
cc.pl.enrich_heatmap(data, 'leiden_merged', figsize=(4,4), scale_cols=True, show=False)
plt.show()

# %% [code]
cc.pl.volcano(data, 'leiden', p_cut=.01, p_adjust=True, figsize=(4,4))

# %% [code]
#cc.tl.merge(data, 'leiden', 0.01)

# %% [code]
sc.pl.umap(data, color='leiden_merged', show=False)
plt.show()

# %% [code]
cc.pl.sig_cluster_diagnostics(data, 'leiden')

# %% [code]
fig, ax = plt.subplots(ncols=2, figsize=(10,3))
cc.pl.check_abundance_bias(data, 'leiden', p_cut=.045, p_adjust=True, show_legend=False, ax=ax[0], show=False)
cc.pl.check_total_abundance_bias(data, 'leiden', p_cut=.045, p_adjust=True, show_legend=False, ax=ax[1], show=False)
plt.show()

# %% [code]
markers = data.uns['leiden_merged_markers']
markers

# %% [code]
marker_genes = {key: values[0:2] for key, values in markers.items()}

# %% [code]
for clust, genes in marker_genes.items():

    sc.pl.umap(data, color=['leiden_merged']+list(genes), groups=[clust])

# %% [code]
#### Selecting some example genes to save and show in the plot...
select_genes = ['BLTP2', 'XIST', 'ARHGAP15', 'CASC15', 'HSP90AA1', 'ENSG00000287409', 'LINC01811', 'DSCAM',
               'CR1L', 'CENPF']
for gene in select_genes:
    sc.pl.umap(data, color=gene, show=False)
    plt.show()

# %% [code]
len(select_genes)

# %% [code]
#### Also for the on-target and off-target genes of interest.
perturbed_genes = ['ARID1A', 'SMARCA4', 'CHD3', 'CHD4', 'CDC27', 'USP9X', 'ADSS1', 'MACROD2', 'CNIH3', 'FBXO38-DT']
for gene in perturbed_genes:
    sc.pl.umap(data, color=gene, show=False)
    vhs.dealWithPlot(True, True, True, out_plots, f"{gene}_umap.png", 300)

# %% [code]
len(perturbed_genes)

# %% [code]
data.layers["scaled"] = sc.pp.scale(data, copy=True).X

# %% [code]
sc.pl.matrixplot(
    data,
    markers,
    "leiden_merged",
    dendrogram=True,
    colorbar_title="mean z-score",
    layer="scaled",
    vmin=-2.5,
    vmax=2.5,
    cmap="bwr", show=False
)
plt.show()

# %% [code]
tcfs = [gene for gene in data.var_names if gene.startswith('TCF')]

# %% [code]
# Looking at WNT co-TFs
sc.pl.umap(data, color='LEF1')

# %% [code]
sc.pl.umap(data, color=tcfs)

# %% [code]
sc.pl.umap(data, color='EZH2')

# %% [code]
sc.pl.umap(data, color='USP9X')

# %% [code]
sc.pl.umap(data, color='MCL1')

# %% [code]
sc.pl.umap(data, color='NFYA')

# %% [code]
sc.pl.umap(data, color='PRICKLE2')

# %% [code]
sc.pl.umap(data, color='CDC27')

# %% [code]
genes_ = ['ADSS1', 'MACROD2', 'CNIH3', 'FBXO38-DT', 'USP9X', 'CDC27', 'ARID1A', 'SMARCA4', 'CHD3', 'CHD4']
for gene_ in genes_:
    fig, ax = plt.subplots(figsize=(6,5))
    sc.pl.umap(data, color=gene_, ax=ax)
    plt.show()

# %% [code]
data.shape

# %% [markdown]
# Saving the cell metadata to add to the GEO entry.

# %% [code]
data.obs.columns

# %% [code]
keep_cols = ['sample', 'n_genes_by_counts', 'log1p_n_genes_by_counts',
       'total_counts', 'log1p_total_counts', 'pct_counts_in_top_50_genes',
       'pct_counts_in_top_100_genes', 'pct_counts_in_top_200_genes',
       'pct_counts_in_top_500_genes', 'mito_perc',
       't7_all', 't7_no_bc_umis', 't7_umis', 'S_score', 'G2M_score', 'phase',
       'leiden', 'leiden_merged', 'log10_total_counts', 'edit_cell_labels']

cell_meta = data.obs.loc[:,keep_cols]
cell_meta

# %% [code]
data.obsm.keys()

# %% [code]
cell_meta['UMAP_1'] = data.obsm['X_umap'][:,0]
cell_meta['UMAP_2'] = data.obsm['X_umap'][:,1]

cell_meta

# %% [code]
out_dir

# %% [code]
?cell_meta.to_csv

# %% [code]
cell_meta.to_csv(f'{out_dir}cell_metadata_10k.tsv.gz', sep='\t')

# %% [code]
print( f'{out_dir}cell_metadata_10k.tsv.gz' )

# %% [code]
!zcat {out_dir}cell_metadata_10k.tsv.gz | head

# %% [markdown]
***Now the edit_site_info metadata!!!***

# %% [code]
data.uns.keys()

# %% [code]
data.obsm.keys()

# %% [code]
site_specific_umi_count = data.obsm['t7_umis'].sum(axis=0).reset_index()
site_specific_umi_count.index = site_specific_umi_count['index']
site_specific_umi_count.columns = ['index', 't7_barcoded_umis']
site_specific_umi_count

# %% [code]
site_specific_umi_count

# %% [code]
t7_umis = data.obsm['t7_umis']

site_specific_mean_umis = []
for i, edit_site in enumerate(t7_umis.columns):

    edit_site_counts = t7_umis.values[:,i]
    nonzero_bool = edit_site_counts > 0

    mean_ = np.mean( edit_site_counts[nonzero_bool] )
    site_specific_mean_umis.append( mean_ )

# %% [code]
site_specific_umi_count.loc[t7_umis.columns.values, 't7_barcoded_umis_per_edited_cell'] = site_specific_mean_umis

# %% [code]
site_specific_umi_count

# %% [code]
edit_site_info = data.uns['edit_site_info_guide_homology_and_coinc_final']
edit_site_info.columns

# %% [code]
site_specific_umi_count = site_specific_umi_count.loc[edit_site_info.index.values]

# %% [code]
edit_site_info.index

# %% [code]
site_specific_umi_count.index.values == edit_site_info.index.values

# %% [code]
keep_cols = ['name', 'chr', 'pos', 'start_window', 'end_window',
       'intersecting_genes', 'copy-number', 'n_cells_edited',
       'stranded_edit_dist', 'target_name', 'target_start', 'target_end',
       'target_seq', 'target_pam', 'strand', 'offset', 'abs_offset',
       'homol_guide_name', 'homol_guide_seq', 'homol_guide_pam', 'target_aln',
       'guide_aln', 'hamming_aln_distance', 'hamming_aln_indel_corrected',
       'match_aln_count', 'coinc_guide_name', 'coinc_guide_seq', 'coinc_count',
       'coinc_fraction', 'guide_score', 'confident_guide_call',
       'manual_edit_call']

edit_site_info_sub = edit_site_info.loc[:, keep_cols]
edit_site_info_sub['t7_barcoded_umis'] = site_specific_umi_count['t7_barcoded_umis'].values
edit_site_info_sub['t7_barcoded_umis_per_edited_cell'] = site_specific_umi_count['t7_barcoded_umis_per_edited_cell'].values
edit_site_info_sub

# %% [code]
out_dir

# %% [code]
edit_site_info_sub.to_csv(f'{out_dir}edit_site_metadata_10k_cell.tsv', sep='\t', index=False)

# %% [code]
print( f'{out_dir}edit_site_metadata_10k_cell.tsv' )

# %% [code]
!head {out_dir}edit_site_metadata_10k_cell.tsv

# %% [markdown]
# Doing some motif searching around the USP9X edit site

Since is an intronic edit but appears to have an effect, but also appears to occur in a CRE. From REMAP, can see a ton of TFs which are annotated as binding in this window in about a 1kb space. 

So will search to see which of these TFs specifically have a binding site at this edit position, and if these actually are expressed in K562 cells. 

First downloaded REMAP cis regulatory module (CRM) annotation from REMAP here: 

    https://remap2022.univ-amu.fr/download_page

Then loaded this into IGV with the T7 barcoded bam, and went to the USP9X edit site.

Then looked at the bed annotation, and copy-pasted the TFs listed at the CRM intersecting the USP9X edit site below:

# %% [code]
crm_loc = ('chrX', 41086293, 41087118) # Containg the USP9X binding site.

# %% [code]
tfs_around_edit = 'MYBL2,ZBTB26,RELA,KDM5B,ESR2,MXI1,EP300,OSR2,TBP,E2F1,IRF2,PRDM10,TBX2,NR3C1,TFAP2C,AGO1,KDM3A,FOXA2,YY2,STAG1,ARNTL,KDM4B,CREM,YY1,ZHX2,FOXA1,MYCN,ARID1B,HCFC1,STAT1,HOXA3,NFRKB,SIN3A,ARNT,MAX,TEAD4,CREB1,ZNF281,ZNF652,SFMBT1,JUND,ZFP42,PAX5,CXXC4,ZNF561,KMT2B,GATAD2B,ZNF467,ATF1,MCRS1,IFNA1,SP2,MED,SIRT6,ZNF711,RUVBL2,SMAD2-3,STAT3,SP1,KMT2D,KLF15,PGR,TAF1,ZNF148,AR,CEBPD,EHMT2,CBX4,INTS13,TP53,CEBPB,VEZF1,RBPJ,ZNF141,BCL6,SRSF1,ELF1,IKZF3,KMT2C,ASH2L,SP4,HEXIM1,SKI,FOXM1,SMC1,CDK9,SMARCC1,TARDBP,POU5F1,ARID1A,KLF12,ZSCAN30,ERG,ZNF695,GLIS2,BRD2,TCF4,PAXIP1,JUN,PATZ1,MLLT1,ARID2,MTA3,GLIS1,ETS1,PCGF1,RORC,HNF4A,DPF2,CCNT2,CTBP1,ZNF202,INTS11,SS18,KLF1,DRAP1,REST,ZBTB7A,CDKN1B,BACH1,BAF155,RUNX1,SMARCA5,GATA3,NONO,SMAD1,ESR2,TFE3,HMGN3,FLI1,SPIN1,KLF5,ZNF143,MORC2,MRTFB,MAZ,INO80,ZNF596,NEUROD1,BCL11A,KLF14,TWIST1,RB1,SIX5,BRD9,TBL1X,NIPBL,MTA1,GTF2F1,SMC1A,CREB3L1,EZH2,AFF4,NELFCD,MNX1,BCLAF1,GABPA,SMC3,ZNF423,ZNF660,PHF8,SKIL,KDM5B,STAG2,NUTM1,ATRX,RAD21,ZNF740,JUND,ZNF184,APC,ZBTB48,GRHL2,CTCF,ZNF501,ZNF189,BRD3,ZNF770,NELFA,CDK8,MYC-DAXX,SMARCB1,NFE2,MNT,HMGXB4,KDM1A,TBP,ZFX,ZNF276,ZNF765,BCOR,GATAD1,SMAD3,GATA4,E2F1,FOXP1,ZNF341,RCOR1,CREB1,ZFY,FOXK1,AR,MYNN,YY1,HNRNPLL,E2F7,EP300,TFDP2,HBP1,SMAD5,RNF2,ZNF175,KLF6,ZKSCAN1,ZBED1,GATA6,IRF1,TRIM24,EGR1,PAX5,TAF7,TBL1XR1,ZNF142,KLF1,ZFP69B,CDX2,ELL2,ZBED2,JMJD1C,E2F6,HCFC1,TRPS1,ZNF524,BRCA1,HDAC2,CBFA2T3,ZNF697,GATA2,TCF3,TCF12,MAFF,CHD2,ZNF24,ILF3,CEBPD,FOXP2,RBBP5,E2F4,ZBTB33,UBTF,IRF2,SMAD1,U2AF1,MSX2,NCOR1,GATA1,NEUROD1,HDAC3,CREM,NELFE,ZNF18,NR2F6,CREBBP,ELF1,ATF3,KLF4,NR2F2,XRCC5,TAL1,ATF7,CBFA2T2,HDAC1,TEAD4,BRD1,ZKSCAN2,ARID1B,TRIM28,RBBP4,GFI1B,EZH2,IRF4,ZNF282,MEF2B,ARID3A,PML,ZHX1,CHD1,BCLAF1,LDB1,KLF6,ZSCAN29,ZNF639,MED1,DEK,CTCFL,FOXA1,ZGPAT,JUN,ZNF324,ZFP37,SMARCC2,TBX21,PHIP,ZNF652,NBN,MLLT1,TAF1,CEBPB,RB1,ARNT,TRIM22,MEN1,MIER1,HIF1A,HDAC2,TBP,CHD1,SAP30,ZNF664,NFATC3'
tfs_around_edit = tfs_around_edit.split(',')
len(tfs_around_edit)

# %% [code]
data.shape

# %% [code]
data.var.keys()

# %% [code]
##### Checking which of the TFs are expressed.
tfs_present = [tf for tf in tfs_around_edit if tf in data.var_names]
len(tfs_present)

# %% [code]
tfs_means = data.var.loc[tfs_present, 'means']

tfs_means

# %% [code]
plt.hist(tfs_means.values, bins=100)
plt.vlines(0.2, 0, 35, color='red')
plt.show()

# %% [code]
tfs_expressed = tfs_means.index.values[ tfs_means.values >= 0.2 ]
len(tfs_expressed)

# %% [code]
##### Downloading the non-redundant vertebrate TFs from JASPAR
!wget https://jaspar.elixir.no/download/data/2024/CORE/JASPAR2024_CORE_vertebrates_non-redundant_pfms_jaspar.txt -O {out_dir}JASPAR2024_CORE_vertebrates_non-redundant_pfms_jaspar.txt

# %% [code]
from Bio import motifs

# Path to your JASPAR format file
file_path = f"{out_dir}JASPAR2024_CORE_vertebrates_non-redundant_pfms_jaspar.txt"

# Open the file and parse the motifs
with open(file_path) as f:
    jaspar_motifs = motifs.parse(f, "jaspar")

# %% [code]
jaspar_motifs[0].name

# %% [code]
tfs_motifs = [motif for motif in jaspar_motifs if np.any([motif.name==tf for tf in tfs_expressed])]
tfs_names = [motif.name for motif in tfs_motifs]
len(tfs_motifs), tfs_names

# %% [code]
crm_sequence = 'GTTGTGTGTGGAGCCGGTGTCCCGAGAGCGTGTCTGTGTGTTTTGGACTGTAAGGGAAAATGGCTGACGGCTCCGTCCCTCCCCCTCCTCCCCCTTCCCGTCCTGTCTGGGGCCGAGTAGGAGGATGGAGTCTCCGGGCGTGGCGGGCAGCCCCCCCGCCGTCCGCACCCGGGGGCCGGGTCGGTTTGGGGGCTGGCCGCCCCCCGCCCGGCGGCCGCTTTCCTACCCGGTCTGGTTCCACTTGACACCAAACGCCATTACTGAGGTGGAGTGATTTGGAGCTGGGGAGCTAGGCCTCGGCTACGCCTGGCTATCAGGCCTGACGACTTGTCCGCCTCCCCCCACCCTCCCGCGATAACCCGCAGTTTCTCCCCGGCCCTGGGCTCCTGGAGTGGGAGCTGGGTTATCGGCCTCCTACACCTCCTTCGCCCCGCTTTATTTGCAGAGGGCCAAAGCCGTGAAAGGGGTTTTTGAATGCAGAAATCGCTTCCAAGCTAGCTTGGACCTATTACCTTTCTATTTTTAACTGTTTCTGCAAATATTAGCATGTGCGAGTTTGGTAGTGATCAGGTCTTCAGATTCTTGTGACGTGAGGCTTAGGGTTGGATAAATGGCTGATTTTTCATTCATTGTGAAATAAGTTAAAGAGTTATACAGCAGGTGTGTAGATGGCTTCTTAAATTATTTGTGTGGATTTCTGTAGCGACTAGAACTTACACGAGAATTTGCAAAATTTCTAGTAGAACCTGCTTAAATTAAGCATGTTATTTTTTGGAAGTTCCATAGTTACAACGTTAATTGAATTTACACGGTGCAGTTTACAGGT'

# %% [code]
len(crm_sequence)

# %% [code]
from Bio.Seq import Seq

crm_seq = Seq(crm_sequence)

# %% [code]
crm_seq

# %% [code]
#### Let's reduce the CRM seq to just a few bp around the USP9X cut site!
edit_site = ('chrX', 41086519)

# %% [code]
crm_loc

# %% [code]
edit_site_in_crm = edit_site[1] - crm_loc[1]
edit_site_in_crm

# %% [code]
crm_seq[(edit_site_in_crm-2):(edit_site_in_crm+3)] # This makese sense!!!

# %% [code]
??pssm.search

# %% [code]
hits = []
hit_tfs = []
search_lens = []
motif_lens = []
motif_seqs = []
hit_seq = []
motifs_ = []
for tf_name, tf_motif in zip(tfs_names, tfs_motifs):
    pssm = tf_motif.pssm

    motif_len = tf_motif.length

    # Only search in the sequence that would intersect the cut site, i.e.
    # the donor insertion would have to BE immediately at the TF binding site. 
    search_seq = crm_seq[(edit_site_in_crm-motif_len-1):edit_site_in_crm+motif_len]

    for hit in pssm.search( search_seq ):

        hits.append( hit )
        search_lens.append( len(search_seq) )
        motif_lens.append( motif_len )
        hit_tfs.append( tf_name )
        motif_seqs.append( tf_motif.consensus )
        motifs_.append( tf_motif )
        
        if hit[0] < 0:
            seq_ = search_seq.reverse_complement()[abs(hit[0])-motif_len:abs(hit[0])]
            hit_seq.append( (seq_.reverse_complement(), seq_) )
        else:
            seq_ = search_seq[abs(hit[0]):abs(hit[0])+motif_len]
            hit_seq.append( (seq_, seq_.reverse_complement()) )

# %% [code]
search_seq.reverse_complement()

# %% [code]
hits, hit_tfs, search_lens, motif_lens, motif_seqs, hit_seq

# %% [code]
sp1_start = (edit_site[1]+(motif_lens[0]-1))-(abs(hits[0][0]))
sp1_loc = (sp1_start, sp1_start+motif_lens[0])
sp1_loc

# %% [markdown]
***Verifying if any of these have a peak in K562 cells in ENCODE***

SP1: https://www.encodeproject.org/experiments/ENCSR991ELG/
YES, directly at the USP9X edit site. AND the height of the reads is suggestive the TF binding site is at the USP9X edit site.

ZNF175: https://www.encodeproject.org/experiments/ENCSR011PEI/
NO

CTCF: https://www.encodeproject.org/experiments/ENCSR000EGM/
NO

E2F6: https://www.encodeproject.org/experiments/ENCSR000BLI/
YES, BUT the peak centre appears further upstream from the USP9X edit site, suggesting the TF binding site is more upstream.

ELF1: https://www.encodeproject.org/experiments/ENCSR975SSR/
NO

GABPA: https://www.encodeproject.org/experiments/ENCSR290MUH/
YES, but very broad and weak peak, suggest of perhaps not being a good candidate.

RBPJ: https://www.encodeproject.org/experiments/ENCSR740XMB/
NO

ZBTB7A: https://www.encodeproject.org/experiments/ENCSR000BME/
YES, and the peak seems to hint at plausibility the binding site is at the edit location! But is a poor motif score.

# %% [code]
print(motifs_[0]) ### Looks like a legit binding site for SP1...

# %% [code]
motifs_[0].reverse_complement()

# %% [code]
?logomaker.Logo

# %% [code]
motif.counts

# %% [code]
import logomaker

motif = motifs_[0].reverse_complement()

counts_df = pd.DataFrame(motif.pwm)

# Normalize counts to probabilities for a clearer logo
#logo_df = counts_df.div(counts_df.sum(axis=1), axis=0)

#logomaker.Logo(logo_df)
logomaker.Logo(counts_df)
plt.show()

# %% [code]
import logomaker

motif = motifs_[0]

counts_df = pd.DataFrame(motif.pwm)

# Normalize counts to probabilities for a clearer logo
#logo_df = counts_df.div(counts_df.sum(axis=1), axis=0)

#logomaker.Logo(logo_df)
logomaker.Logo(counts_df)
plt.show()

# %% [markdown]
Looking for SP1 KD data, I found this paper where the knocked-down SP1 in HEK293T cells and performed RNA-seq: 

    https://www.cell.com/molecular-cell/pdf/S1097-2765(22)00649-9.pdf

Although I cannot see the SP1 binding site in HEK293T ChIP-seq in ENCODE: 

    https://www.encodeproject.org/experiments/ENCSR906PEI/
    

In Table S3 they list the DE genes, which INCLUDES USP9X!!!!! This is the significance:

# %% [code]
ID	NAME	siMock.cRPKM	siSP1.cRPKM	siSP1.log2FC	siSP1.PValue	siSP1.FDR
ENSG00000124486	USP9X	61.53	26.735	-1.04907006220249	2.63e-08	2.86e-07

# %% [markdown]
This paper also knocked-down USP9X:
    
    https://onlinelibrary.wiley.com/doi/10.1002/mc.23591

In supp table 3 with the DE results in breast cancer. Could not see NRF1 DE, or many of the targets, but could see these targets as DE specific to USP9X down-regulation: 

    BRD2, all others I could not find.

Putting this all together in IGV to make a visualisation, can see the session here I used:

    data/edit_capture/superb_count_all10/USP9X_edit_dissection.xml
```
