## Additional Files for "Joint single-cell capture of Cas9 edits and transcriptomes reveals on- and off-target effects on gene expression": compile_baf-nurd-targets.ipynb

```
# %% [markdown]
# Compiling the BAF/NURD targets based on SMARCA4 K562 ChIP-seq and HDAC1 ChIP-seq from ENCODE:

# INPUT: 

    * NONE, will be downloaded.

# OUTPUT: 

    * targets/

# Environment Setup

# %% [code]
work_dir = './'
import os
os.chdir( work_dir )

import numpy as np
import pandas as pd

import matplotlib.pyplot as plt

bedtools_dir = 'third_party_tools/bedtools2/bin/'

data_dir = 'targets/'
data_dir2 = '231009_index/'
data_dir3 = 'edit_capture/data/DE_out_parse_corrected2/superbseq/'
data_dir4 = 'edit_capture/data/DE_out_parse_corrected2/sample/'
out_dir = data_dir

# %% [code]
!mkdir {out_plots}

# %% [code]
!mkdir {data_dir}

# %% [markdown]
# Downloading the data

# %% [code]
# SMARCA4 K562 ChIP-seq peaks.
!wget https://www.encodeproject.org/files/ENCFF267OGF/@@download/ENCFF267OGF.bed.gz -O {data_dir}smarca4_k562_peaks.bed.gz
!echo y | gzip -d {data_dir}smarca4_k562_peaks.bed.gz

# %% [code]
!head -3 {data_dir}smarca4_k562_peaks.bed

# %% [code]
!cut -f1,2,3 {data_dir}smarca4_k562_peaks.bed | sort -k1,1 -k2,2n | uniq > {data_dir}smarca4_k562_peaks.regions.bed

# %% [code]
!head -3 {data_dir}smarca4_k562_peaks.regions.bed

# %% [code]
!wget https://www.encodeproject.org/files/ENCFF432KJA/@@download/ENCFF432KJA.bed.gz -O {data_dir}hdac1_k562_peaks.bed.gz
!echo y | gzip -d {data_dir}hdac1_k562_peaks.bed.gz

# %% [code]
!cut -f1,2,3 {data_dir}hdac1_k562_peaks.bed | sort -k1,1 -k2,2n | uniq > {data_dir}hdac1_k562_peaks.regions.bed

# %% [code]
!head -3 {data_dir}hdac1_k562_peaks.regions.bed

# %% [code]
!wget https://www.encodeproject.org/files/ENCFF652ZZF/@@download/ENCFF652ZZF.bed.gz -O {data_dir}hdac2_k562_peaks.bed.gz
!echo y | gzip -d {data_dir}hdac2_k562_peaks.bed.gz

# %% [code]
!cut -f1,2,3 {data_dir}hdac2_k562_peaks.bed | sort -k1,1 -k2,2n | uniq > {data_dir}hdac2_k562_peaks.regions.bed

# %% [code]
!head -3 {data_dir}hdac2_k562_peaks.regions.bed

# %% [markdown]
***Also ATAC-seq to narrow to accessible genes!***

# %% [code]
!wget https://www.encodeproject.org/files/ENCFF333TAT/@@download/ENCFF333TAT.bed.gz -O {data_dir}atac_k562_peaks.bed.gz
!echo y | gzip -d {data_dir}atac_k562_peaks.bed.gz

# %% [code]
!cut -f1,2,3 {data_dir}atac_k562_peaks.bed | sort -k1,1 -k2,2n | uniq > {data_dir}atac_k562_peaks.regions.bed

# %% [code]
!head -3 {data_dir}atac_k562_peaks.regions.bed

# %% [code]
!wget https://www.encodeproject.org/files/ENCFF985QBS/@@download/ENCFF985QBS.bed.gz -O {data_dir}chd4_k562_peaks.bed.gz
!echo y | gzip -d {data_dir}chd4_k562_peaks.bed.gz

# %% [code]
!cut -f1,2,3 {data_dir}chd4_k562_peaks.bed | sort -k1,1 -k2,2n | uniq > {data_dir}chd4_k562_peaks.regions.bed

# %% [code]
!head -3 {data_dir}chd4_k562_peaks.regions.bed

# %% [markdown]
***NRF1 targets!!!***

# %% [code]
!wget https://www.encodeproject.org/files/ENCFF259YUE/@@download/ENCFF259YUE.bed.gz -O {data_dir}nrf1_k562_peaks.bed.gz
!echo y | gzip -d {data_dir}nrf1_k562_peaks.bed.gz

# %% [code]
!cut -f1,2,3 {data_dir}nrf1_k562_peaks.bed | sort -k1,1 -k2,2n | uniq > {data_dir}nrf1_k562_peaks.regions.bed

# %% [code]
!head -3 {data_dir}nrf1_k562_peaks.regions.bed

# %% [markdown]
# Determining gene TSS

# %% [code]
!zcat {data_dir2}Homo_sapiens.GRCh38.110.gtf.gz | grep "	gene	" | cut -f1,4,5,7,9 > {data_dir}hg38_genes.txt
!head -3 {data_dir}hg38_genes.txt

# %% [code]
#### Parsing to get transcription start sites!
gene_info = open(f'{data_dir}hg38_genes.txt', 'r')
tss_bed = open(f'{data_dir}hg38_tss.bed', 'w')

window=200
for line in gene_info:

    #### Positional information.
    info_split = line.split('\t')

    chr_ = f'chr{info_split[0]}'
    if info_split[3]=="+":
        tss_pos = int(info_split[1])
    else:
        tss_pos = int(info_split[2])

    region_start, region_end = tss_pos - window, tss_pos + window

    #### Gene information
    gene_split = info_split[-1].split(';')
    gene_name = [split_ for split_ in gene_split if 'gene_name' in split_]
    if len(gene_name) > 0:
        gene_name = gene_name[0].split('"')[1]
    else:
        gene_name = [split_ for split_ in gene_split if 'gene_id' in split_][0].split('"')[1]

    #### Writing out the information.
    tss_bed.write(f'{chr_}\t{region_start}\t{region_end}\t{gene_name}\n')

gene_info.close()
tss_bed.close()

# %% [code]
!cat {data_dir}hg38_tss.bed | sort -k1,1 -k2,2n | uniq > {data_dir}hg38_tss.sorted.bed

# %% [code]
!head {data_dir}hg38_tss.sorted.bed

# %% [code]
!wc -l {data_dir}hg38_tss.sorted.bed

# %% [markdown]
***Looking good!***

# %% [markdown]
# Intersecting to get the genes!

# %% [markdown]
***First let's get genes with an open promoter***

# %% [markdown]
BAF complex: https://www.ncbi.nlm.nih.gov/pmc/articles/PMC4968166/

# %% [code]
!{bedtools_dir}bedtools intersect -wa -a {data_dir}hg38_tss.sorted.bed -b {data_dir}atac_k562_peaks.regions.bed | sort -k1,1 -k2,2n | uniq > {data_dir}atac_k562_genes.bed
!head {data_dir}atac_k562_genes.bed

# %% [code]
!wc -l {data_dir}atac_k562_genes.bed

# %% [markdown]
***Now getting smarca4 marked genes with open promoters***

# %% [code]
!{bedtools_dir}bedtools intersect -wa -a {data_dir}atac_k562_genes.bed -b {data_dir}smarca4_k562_peaks.regions.bed | sort -k1,1 -k2,2n | uniq > {data_dir}smarca4_k562_genes.bed
!head {data_dir}smarca4_k562_genes.bed

# %% [code]
!cat {data_dir}smarca4_k562_genes.bed | cut -f4 | sort | uniq > {data_dir}smarca4_k562_genes.txt
!wc -l {data_dir}smarca4_k562_genes.txt

# %% [markdown]
Seems like a reasonable number!

***Checking these in the ENCODE genome browser verified they do have SMARCA4 K562 CHIP-seq promoter!***

# %% [markdown]
***Much more reasonable number!!!***

# %% [markdown]
***Try with chd4 only***

# %% [code]
!{bedtools_dir}bedtools intersect -wa -a {data_dir}hg38_tss.sorted.bed -b {data_dir}chd4_k562_peaks.regions.bed | sort -k1,1 -k2,2n | uniq > {data_dir}chd4_k562_genes.bed
!head {data_dir}chd4_k562_genes.bed

# %% [code]
!cat {data_dir}chd4_k562_genes.bed | cut -f4 | sort | uniq > {data_dir}chd4_k562_genes.txt
!wc -l {data_dir}chd4_k562_genes.txt

# %% [code]
!{bedtools_dir}bedtools intersect -wa -a {data_dir}hg38_tss.sorted.bed -b {data_dir}atac_k562_peaks.regions.bed | sort -k1,1 -k2,2n | uniq > {data_dir}atac_k562_genes.bed
!head {data_dir}atac_k562_genes.bed

# %% [markdown]
# Candidate targets based on enhancer - gene associations!!

Speaking with Alex, he said the SMARCA4 ChIP-seq peaks that intersect with promoters would likely enrich for p-BAF, which is one of 3 BAF complexes that is more promoter associated. SMARCA4 is found in all BAF complexes. However, ARID1A is found in the more enhancer-associated BAF complexes, so would be worth finding gene targets based on enhancers as well. SO will also try and get a set of targets based on enhancers also!!!

Reading about this, this resource appears very useful, and has cell-line / cell type specific targets:

    https://academic.oup.com/nar/article/48/D1/D58/5628925

Specifically, can get the K562 enhancer - gene associations from this file (NOTE checked the coordinates in the genome browser and look to correspond to hg19!):

    http://www.enhanceratlas.org/data/AllEPs/hs/K562_EP.txt
    

# %% [code]
!wget http://www.enhanceratlas.org/data/AllEPs/hs/K562_EP.txt -O {data_dir}K562_EP.hg19.txt

# %% [code]
!head -3 {data_dir}K562_EP.hg19.txt

# %% [code]
#### Let's determine a cutoff for a good association first!
enhancer_gene = pd.read_csv(f'{data_dir}K562_EP.hg19.txt', sep='\t', header=None)
enhancer_gene.head(3)

# %% [code]
scores = enhancer_gene[1].values
scores

# %% [code]
plt.hist(scores, bins=100)
plt.show()

# %% [code]
score_cutoff = 2
sum(scores >= score_cutoff)

# %% [code]
#### Let's load it and reformat it...
enh_to_gene_file = open(f'{data_dir}K562_EP.hg19.txt', 'r')
enh_to_gene_bed_file = open(f'{data_dir}K562_EP.hg19.bed', 'w')
for line in enh_to_gene_file:
    score = float( line.split('\t')[1] ) 
    if score < score_cutoff: # Only keep confident links.
        continue
        
    info = line.split('\t')[0].split('$')
    enh, gene = info[0], info[1]
    
    enh_chr, enh_start, enh_end = enh.split(':')[0], enh.split(':')[1].split('-')[0], enh.split(':')[1].split('-')[1].split('_')[0]

    enh_to_gene_bed_file.write(f'{enh_chr}\t{enh_start}\t{enh_end}\t{gene}\n')

enh_to_gene_bed_file.close()
enh_to_gene_file.close()

# %% [code]
!head {data_dir}K562_EP.hg19.bed

# %% [code]
!ls -s -h {data_dir}K562_EP.hg19.bed

# %% [code]
#### Used liftOver to convert to hg38 coords: https://genome.ucsc.edu/cgi-bin/hgLiftOver
!head {data_dir}K562_EP.hg19.lifted-to-hg38.bed

# %% [code]
#### Removing the extra column added by liftOver
!cut -f1-4 {data_dir}K562_EP.hg19.lifted-to-hg38.bed > {data_dir}K562_EP.hg19.lifted-to-hg38.sub.bed

# %% [code]
!head -3 {data_dir}K562_EP.hg19.lifted-to-hg38.sub.bed

# %% [code]
!{bedtools_dir}bedtools intersect -wa -a {data_dir}K562_EP.hg19.lifted-to-hg38.sub.bed -b {data_dir}atac_k562_peaks.regions.bed | sort -k1,1 -k2,2n | uniq > {data_dir}K562_EP.hg19.lifted-to-hg38.sub.atac.bed
!head -3 {data_dir}K562_EP.hg19.lifted-to-hg38.sub.atac.bed


# %% [code]
!wc -l {data_dir}K562_EP.hg19.lifted-to-hg38.sub.atac.bed

# %% [code]
!{bedtools_dir}bedtools intersect -wa -a {data_dir}K562_EP.hg19.lifted-to-hg38.sub.atac.bed -b {data_dir}smarca4_k562_peaks.regions.bed | sort -k1,1 -k2,2n | uniq > {data_dir}smarca4_k562_genes.enhancer.bed
!head {data_dir}smarca4_k562_genes.enhancer.bed

# %% [code]
!wc -l {data_dir}smarca4_k562_genes.enhancer.bed

# %% [code]
!cat {data_dir}smarca4_k562_genes.enhancer.bed | cut -f4 | sort | uniq > {data_dir}smarca4_k562_genes.enhancer.genes.txt 
!wc -l {data_dir}smarca4_k562_genes.enhancer.genes.txt

# %% [code]
!head {data_dir}smarca4_k562_genes.enhancer.genes.txt

# %% [code]
### Checking overlap with the promoter genes...
smarca4_prom_genes = set([line.strip('\n') for line in open(f'{data_dir}smarca4_k562_genes.txt', 'r')])
smarca4_enh_genes = set([line.strip('\n') for line in open(f'{data_dir}smarca4_k562_genes.enhancer.genes.txt', 'r')])

len(smarca4_prom_genes), len(smarca4_enh_genes)

# %% [code]
from matplotlib_venn import venn2

plt.figure(figsize=(8, 8))
venn = venn2([smarca4_prom_genes, smarca4_enh_genes], ('Smarca4 promoter genes', 'Smarca4 enhancer genes'))

# Display the Venn diagram
plt.title("Overlap between SMARCA4 targets by promoter versus by enhancer")
plt.show()

# %% [markdown]
***Some level of overlap, but they do look like largely unique sets of genes! Good to know***

# %% [markdown]
# NRF1 targets

# %% [code]
!{bedtools_dir}bedtools intersect -wa -a {data_dir}hg38_tss.sorted.bed -b {data_dir}nrf1_k562_peaks.regions.bed | sort -k1,1 -k2,2n | uniq > {data_dir}nrf1_k562_peaks.genes.bed
!head {data_dir}nrf1_k562_peaks.genes.bed


# %% [code]
!cut -f4 {data_dir}nrf1_k562_peaks.genes.bed | sort | uniq > {data_dir}nrf1_k562_peaks.genes.txt

# %% [code]
!head {data_dir}nrf1_k562_peaks.genes.txt

# %% [markdown]
***Trying to subset to open chromatin***

# %% [code]
!{bedtools_dir}bedtools intersect -wa -a {data_dir}nrf1_k562_peaks.genes.bed -b {data_dir}atac_k562_genes.bed | sort -k1,1 -k2,2n | uniq > {data_dir}nrf1_k562_peaks.genes.open.bed
!head {data_dir}nrf1_k562_peaks.genes.open.bed


# %% [code]
!cut -f4 {data_dir}nrf1_k562_peaks.genes.open.bed | sort | uniq > {data_dir}nrf1_k562_peaks.genes.open.txt

# %% [code]
!head {data_dir}nrf1_k562_peaks.genes.open.txt
```
