## Additional Files for "Joint single-cell capture of Cas9 edits and transcriptomes reveals on- and off-target effects on gene expression": superb_analysis_guide_inference.py

```
"""
Running the edit-to-guide inference
"""

########################################################################################################################
                                            # Environment Setup #
########################################################################################################################
work_dir = './'
import os
os.chdir( work_dir )

import numpy as np
import pandas as pd
import matplotlib.pyplot as plt

import seaborn as sb

import scanpy as sc

from collections import defaultdict

import superb_guide_inference as sgi
import importlib as imp
imp.reload( sgi )

data_dir = 'superb_count_all/'
data_dir2 = 'data/edit_capture/'
out_dir = data_dir

########################################################################################################################
                                            # Loading the data #
########################################################################################################################
data = sc.read_h5ad(f"{data_dir}superb_data.h5ad")

print(data)

print("Here")

# NOTE that data.X is the MixScape package corrected values, setting back..
print( data.layers.keys() )

data.layers['log1p_counts'] = data.X.copy()

########################################################################################################################
                                    # Performing the edit-calling correction #
########################################################################################################################
ref_fasta = f'{data_dir2}Homo_sapiens.GRCh38.dna.primary_assembly.fa'

# Specifies the name of the guide, and the sequence of the guide.
guide_seq_dict = {
    'ARID1A g05': 'AAGAACTCGAACGGGAACGCGGG',
    'ARID1A g06': 'GCTTCGGGCAACCCTACGGCCGG',
    'SMARCA4 g19': 'TGGCCGAGGAGTTCCGCCCAGGG',
    'SMARCA4 g22': 'ACCGAGGGTGGAGGTAGCGTTGG',
    'CHD3 g10': 'AATATGGAACCGGACCGGGTCGG',
    'CHD4 g12': 'CCTCACTGCCCGCCGAGCAGGGG',
    'CHD4 g14': 'GCCGGCATAAGAGCATACGCTGG'
}

# decrease query distance to 50, runs without error (reduced artifacts with distant pams)
sgi.impute_edit_guides_homology(data, ref_fasta, guide_seq_dict, n_targets=3, query_range=25, seed_size=5)

paired_guides = [('ARID1A g05', 'SMARCA4 g19'), ('CHD3 g10', 'CHD4 g14')]

# Inferring the primary on-target edit sites for each guide, as the top matching by homology edit sites.
# Alternatively, could input own custom dictionary that has the actual edit sites.
n_guides = 7
ham_df = data.uns['edit_site_info_guide_homology'].copy()
guide_to_site_dict = pd.Series(ham_df['name'].head(7).values,
                               index=ham_df['homol_guide_name'].head(7).values).to_dict()

sgi.impute_edit_guides_coinc(data, guide_seq_dict, paired_guides, guide_to_site_dict)

####### Now performing cell-to-guide imputation
# list of guide sets
guide_sets = [
    {'ARID1A g05'},
    {'ARID1A g06'},
    {'SMARCA4 g19'},
    {'SMARCA4 g22'},
    {'ARID1A g05', 'SMARCA4 g19'},
    {'CHD4 g12'},
    {'CHD4 g14'},
    {'CHD3 g10', 'CHD4 g14'},
]

##### Now at this point, with a ton of information about the edit sites, let's try and do a secondary set of
##### filtering to figure out which are real versus not. I will label cases that look very geniune based on the
##### t7 barcoded bam, and for ambiguous cases I will label as ambiguous.
manual_ambiguous_edit_names = ['hg38_9:97998566', 'hg38_17:64502467', 'hg38_7:155680729', 'hg38_10:73387744',
                               'hg38_19:3406603', 'hg38_1:246596227', 'hg38_21:34086839', 'hg38_15:42565538',
                               'hg38_21:8992513', 'hg38_22:19372513', 'hg38_2:19901462', 'hg38_19:38836517',
                               'hg38_13:18958292', 'hg38_6:33275863', 'hg38_19:975594', 'hg38_8:47862492',
                               'hg38_15:34382827', 'hg38_6:140374836', 'hg38_6:16262954', 'hg38_6:67885109',
                                ]

##### Can now plot out the key metrics gathered to see if other aspects aside from the alignment (like the guide homology)
##### can clearly separate the two!
data.uns['edit_site_info_guide_homology_and_coinc_final']['manual_edit_call'] = True
ambiguous_bool = [edit_ in manual_ambiguous_edit_names for edit_ in data.uns['edit_site_info_guide_homology_and_coinc_final']['name']]
data.uns['edit_site_info_guide_homology_and_coinc_final'].loc[ambiguous_bool, 'manual_edit_call'] = False

metrics = ['match_aln_count', 'n_cells_edited', 'stranded_edit_dist', 'abs_offset']
metric = metrics[0]

edit_info = data.uns['edit_site_info_guide_homology_and_coinc_final']

##### Saving this, so Mickey can make some figures on the alignments for the called true edit events.
edit_info.to_csv(f'{out_dir}edit_site_info.with-guide-inference.txt', sep='\t', index=False)

for metric in metrics:
    sb.stripplot(edit_info, x='manual_edit_call', y=metric, hue='manual_edit_call')
    plt.show()

plt.scatter(edit_info['match_aln_count'].values, np.log2(edit_info['n_cells_edited'].values), color='blue',
            label='manually false edit')
plt.scatter(edit_info['match_aln_count'].values[edit_info['manual_edit_call'].values],
            np.log2(edit_info['n_cells_edited'].values[edit_info['manual_edit_call'].values]), color='orange',
            label='manually called edit')
plt.xlabel('Guide and target sequence alignment match count')
plt.ylabel('Log2 number of cells edited')
plt.legend()
plt.show()

# Now can compare the manual edit site calls and the edit sites where the guide could be called confidently:
conf_guide_edits = set( list(edit_info['name'].values[ edit_info['confident_guide_call'].values ]) )
conf_true_edits = set( list(edit_info['name'].values[ edit_info['manual_edit_call'].values ]) )

from matplotlib_venn import venn2

plt.figure(figsize=(8, 8))
venn = venn2([conf_guide_edits, conf_true_edits], ('Confident edits by homology', 'Manually true edits'))

# Display the Venn diagram
plt.title("Overlap between homology edit filtering and manual true edits")
plt.show()

data.uns['edit_site_info_guide_homology_and_coinc_final'].index = \
                                                data.uns['edit_site_info_guide_homology_and_coinc_final']['name'].values

data.write_h5ad(f'{out_dir}superb_data.with-guide-imputation.h5ad', compression='gzip')
```
