## Additional Files for "Joint single-cell capture of Cas9 edits and transcriptomes reveals on- and off-target effects on gene expression": superb_differential_expression_analysis.py

```
"""
INPUT:

    * superb_count_all/superb_data.with-guide-imputation.h5ad

OUTPUT:

    * plots/edit_capture/allelic_de_v7/
    * edit_capture/allelic_de_v7/

conda setup:

    mamba create -n superb_py10 python=3.10
    conda activate superb_py10

    mamba install scanpy
"""

########################################################################################################################
                                            # Environment Setup #
########################################################################################################################
work_dir = './'
import os
os.chdir( work_dir )

import numpy as np
import pandas as pd
import matplotlib.pyplot as plt

import seaborn as sb

from collections import defaultdict

import scanpy as sc

import scipy.stats as stats
import statsmodels.stats.multitest as smm

import superb_helpers as superb_help
import importlib as imp
imp.reload( superb_help )

import utils

data_dir = 'superb_count_all/'
data_dir2 = 'targets/' # contains the gene sets compiled which have BAF/NuRD/NRF1 targets, also has G2M gene set from mSigDB
out_dir = 'edit_capture/allelic_de_v7/'
out_plots = 'plots/edit_capture/allelic_de_v7/'

########################################################################################################################
                                            # Loading the data #
########################################################################################################################
data = sc.read_h5ad(f"{data_dir}superb_data.with-guide-imputation.h5ad")

data.layers.keys()
data.X = data.layers['log1p_counts'].copy()

########################################################################################################################
                                        # Filtering to edited cells #
########################################################################################################################
on_targets = ['ARID1A', 'SMARCA4', 'CHD3', 'CHD4']

##### Determining the true off-target edits, this is based on manually checking the edit sites
edit_site_info = data.uns['edit_site_info_guide_homology_and_coinc_final']
edit_site_info_true = edit_site_info.loc[edit_site_info['manual_edit_call'].values,:]

edit_site_genes_true = edit_site_info_true['intersecting_genes']
genes_or_edit_set = []
genes_edited = []
[genes_or_edit_set.extend( genes.split(',') ) for edit, genes in edit_site_genes_true.to_dict().items() if str(genes)!='nan']
genes_edited.extend(genes_or_edit_set)
[genes_or_edit_set.append( edit ) for edit, genes in edit_site_genes_true.to_dict().items() if str(genes)=='nan']
genes_or_edit_set = list( set(genes_or_edit_set) )
genes_edited = list( set(genes_edited) )

off_targets = [gene_or_edit for gene_or_edit in genes_or_edit_set if gene_or_edit not in on_targets]
off_target_genes = [gene for gene in off_targets if gene in genes_edited]

##### Will label each cell according to the collapsed gene dosage for which cell is edited.
expr_genes = data.var_names.values[ data.var['means'].values>.2 ]

dosages_targets = data.obsm['dosages_genes'].loc[:, genes_or_edit_set]

#### Curious as to the edited genes frequencies across cells, could JUST validate this analysis by testing each effected
#### gene, and then doing fine-mapping to validate that the actual edit gene is the cause, and not the off-target.
freqs = (dosages_targets.loc[:, genes_edited].values>0).sum(axis=0)

edited_genes = dosages_targets.loc[:, genes_edited].columns.values[freqs>30]
edited_genes = [gene for gene in edited_genes if gene in expr_genes]

#### Need to subset to edited cells
n_ontargets = (dosages_targets.values > 0).sum(axis=1)

# Also need to keep the control cells
edit_cells_ = np.logical_or(n_ontargets > 0, data.obs['sample'].values=='no_ki')

edit_info = data.uns['edit_site_info_guide_homology_and_coinc_final']

########################################################################################################################
                                # R Code used for the sctransform normalisation #
########################################################################################################################
# Setting up the R environment:
# mamba create -n sctransform_py10 python=3.10
# conda activate sctransform_py10
# mamba install conda-forge::r-base
# mamba install scanpy
#
# R
# install.packages('Seurat')
# install.packages("sctransform")
# install.packages('BiocManager')
#
# To get the glmGamPoi to install, had to run the following steps:
# Go here and download the macOS arm64 version: https://www.bioconductor.org/packages/release/bioc/html/rhdf5filters.html
# Then:
# install.packages("~/Downloads/rhdf5filters_1.16.0.tgz", repos = NULL, type = "source")
# BiocManager::install("rhdf5")
# BiocManager::install("HDF5Array")
# BiocManager::install('glmGamPoi')
#
# Now loading the data in R..
# Sys.setenv(RETICULATE_PYTHON = "miniforge3/envs/sctransform_py10/bin/python") #set conda environment
# library(reticulate)
# library(Seurat)
#
# sc <- import("scanpy")
# ad <- sc$read_h5ad("superb_count_all/superb_data.with-guide-imputation.h5ad")
# counts <- ad$layers[['gene_counts']]
# rownames(counts) <- ad$obs_names$values
# colnames(counts) <- ad$var_names$values
#
# gene_n_cells <- ad$var$n_cells_by_counts
# keep_genes <- gene_n_cells > 3
# # Check keeping relevant genes:
# "CDC27" %in% ad$var_names$values[keep_genes]
# "USP9X" %in% ad$var_names$values[keep_genes]
# "SMARCA4" %in% ad$var_names$values[keep_genes]
# "ARID1A" %in% ad$var_names$values[keep_genes]
# "CHD3" %in% ad$var_names$values[keep_genes]
# "CHD4" %in% ad$var_names$values[keep_genes]
### ALL True
# sum(keep_genes)
# 23898
#
# counts <- counts[,keep_genes]
#
# counts_sparse <- as(t(counts), "dgCMatrix")
#
# dat = CreateAssayObject(counts_sparse)
# seurat_obj <- CreateSeuratObject(dat)
#
# # Need to increase this since dataset is oversize...
#
# options(future.globals.maxSize = 2 * 1024^3)  # 2 GB
# seurat_obj <- SCTransform(seurat_obj, vst.flavor = "v2", verbose = T, return.only.var.genes=F)
#
# pearson_residuals <- as.data.frame( seurat_obj@assays$SCT@scale.data )
# log_cpm <- as.data.frame( seurat_obj@assays$SCT@data )
#
# ad_scaled <- sc$AnnData( pearson_residuals )
# ad_cpm <- sc$AnnData( log_cpm )
#
# ad_scaled$write_h5ad("superb_count_all/pearsons.h5ad")
# ad_cpm$write_h5ad("superb_count_all/log-cpm.h5ad")
#
# datas <- dict("pearson_residuals"=pearson_residuals, "log_cpm"=log_cpm)
# datas_py <- r_to_py(datas)
#
# py_save_object(datas_py, 'superb_count_all/sctransform_data.pkl')
#
# py_save_object(pearson_residuals, 'superb_count_all/pearsons.pkl')
#
#  write.table(pearson_residuals, "superb_count_all/pearsons.txt",
#  quote=F, sep='\t')
########################################################################################################################

pearsons = sc.read_h5ad("superb_count_all/pearsons.h5ad").to_df().transpose()
log_cpm = sc.read_h5ad("superb_count_all/log-cpm.h5ad").to_df().transpose()

print(np.all(data.obs_names.values == log_cpm.index.values))
print(np.all(pearsons.index.values == log_cpm.index.values))
print(np.all(pearsons.columns.values == log_cpm.columns.values))

data = data[pearsons.index.values, pearsons.columns.values].copy()

data.layers['sctransform_pearsons'] = pearsons
data.layers['sctransform_log-cpm'] = log_cpm

# Subsetting the cells
data_edited = data[edit_cells_,:].copy()

dosages_targets.loc[:, genes_edited].columns.values[freqs>40]
# ALL on-targets, and a few off targets. Just 8, that we could plausibly test some effects

# min_disp=830 in the original version
sc.pp.highly_variable_genes(data, min_mean=.2, min_disp=0.6, max_mean=5)
print(sum(data.var['highly_variable'].values))
sc.pl.highly_variable_genes(data)

########################################################################################################################
     # for each one of the edits occuring in genes, test their effect on the gene's expression, to determine
     #  if the on-targets have a detectable effect on gene expression.  #
########################################################################################################################
# Filtering to edited genes that have some level of expression!
hvgs = data.var_names.values[ data.var['highly_variable'] ]

#### Also adding in to the tested genes additional genes that were determined as SMARCA4 or CHD4 regulated:
nurd_genes = set([gene.strip('\n') for gene in open(f'{data_dir2}chd4_k562_genes.txt', 'r')])
min_mean = .03
min_disp = .0
nurd_extra, nurd_hvg = superb_help.selected_extra_genes(nurd_genes, hvgs, data_edited, min_mean, min_disp)

### Should also do the same for the BAF targets, to have a consistent criteria.
baf_genes = set([gene.strip('\n') for gene in open(f'{data_dir2}smarca4_k562_genes.txt', 'r')])
min_mean = .2
min_disp = .3
baf_extra, baf_hvg = superb_help.selected_extra_genes(baf_genes, hvgs, data_edited, min_mean, min_disp)

baf_enh_genes = set([gene.strip('\n') for gene in open(f'{data_dir2}smarca4_k562_genes.enhancer.genes.txt', 'r')])
min_mean = .2
min_disp = .3
baf_enh_extra, baf_enh_hvg = superb_help.selected_extra_genes(baf_enh_genes, hvgs, data_edited, min_mean, min_disp)

genes_to_test = np.unique( list(hvgs)+list(nurd_extra)+list(baf_extra)+list(baf_enh_extra) )
# 1937 genes

### Also for the baf enhancers..
print([target in edited_genes for target in on_targets])

edit_gene_colors = {gene: 'grey' for gene in edited_genes} # non-sig so don't care.
edit_gene_colors.update({'CHD3': 'gold', 'CHD4': 'sandybrown',
                        'ARID1A': 'springgreen', 'SMARCA4': 'limegreen',
                         'USP9X': 'red', 'CDC27': 'violet', 'FBXO38-DT': 'lightyellow',
                         'CNIH3': 'deepskyblue'
                         })

# Now we are working with Pearson residuals!!
data_edited.X = pearsons.loc[data_edited.obs_names.values,:].copy()

# Specifies if the genes must belong to a certain condition...
gene_to_condition = {}
for rowi, row in edit_site_info_true.iterrows():

    row_genes = row['intersecting_genes'].split(',')
    row_guide = row['coinc_guide_name']

    for gene in row_genes:
        if gene != 'nan':
            if 'SMARCA4' in row_guide or 'ARID1A' in row_guide:
                gene_to_condition[gene] = 'baf_ki'
            else:
                gene_to_condition[gene] = 'nurd_ki'

if True: # By-passing if already calculated.
    edited_gene_stats = np.zeros((len(edited_genes), 2))
    fit_results = {}
    version_entries = [] # Pearson correlations relating to effect of allelic pairing.
    allelic_group_entries = [] # Pearson correlations relating to expression effects of allelic groups.
    for i, gene in enumerate( edited_genes ):

        # Getting the edit with the most cells for the given gene.
        gene_indices = np.where([gene in genes for genes in edit_info['intersecting_genes']])[0]
        n_cells = edit_info['n_cells_edited'].values[ gene_indices ]
        edit_name = edit_info.index.values[gene_indices][ np.argmax( n_cells ) ]

        # Checked individual levels for CHD4, still a problem... when looking at the log for the 500 cell
        # seems that it over-estimates alot for these, so I suspect ti m
        data_entries_df = superb_help.construct_allelic_pairings(data_edited, gene,
                                                                 umap_key='X_umap',
                                                                 condition_key=gene_to_condition[gene],
                                                                 dosage_key='dosages_genes_manual-edits-only',
                                                                 min_cells_with_allele=25,
                                                                 max_total_umi_dist=1000, #Says total UMI but actually means gene count
                                                                 n_reps=10,
                                                                 total_count_key='gene_count',
                                                                 method='pair_each_allele',
                                                                 )
        if len(np.unique(data_entries_df['subject_allelic_edits'].values)) == 1:
            fixed_effect_variables = ['allelic_edits']
        else: # Apply the 'gamma' correction for higher detection rate in cells with 2 alleles detected as edited.
            fixed_effect_variables = ['allelic_edits', 'subject_allelic_edits']

        alleles_beta, alleles_pval, result, model = superb_help.test_edit_and_gene(data_edited, data_entries_df, gene,
                                                                        random_effect_variable="subject_cell_index",
                                                                        fixed_effect_variables=fixed_effect_variables,
                                                                                   )

        if type(alleles_beta)!=float:
            alleles_beta = alleles_beta[0]
            alleles_pval = alleles_pval[0]

        r_total, r_genes = superb_help.plot_results(result, alleles_beta, data_entries_df, gene, data,
                                 color_violin=edit_gene_colors[gene], out_plots=out_plots, save_fig=True,
                                 show_bootstrap_trend=True)
        version_entries.append( [r_total, r_genes, gene, 'LM_final'] )

        edited_gene_stats[i, :] = [alleles_beta, alleles_pval]
        fit_results[gene] = [result, model]

        #################
        ## Plotting the raw data WITHOUT the allelic edit pairing / linear mixture modelling
        r_total_unmodelled, r_genes_unmodelled = superb_help.plot_unmodelled_alleles(data_edited, gene, gene,
                                            'dosages_genes_manual-edits-only', gene_to_condition[gene],
                                 color_violin=edit_gene_colors[gene],
                                 out_plots=out_plots+'/without_linear_mixture_modelling/', save_fig=True,
                                 show_bootstrap_trend=True)
        version_entries.append([r_total_unmodelled, r_genes_unmodelled, gene, 'noLM'])

        #################
        ## Also plot if we did NOT model the allelic edit group as a fixed effect variable, to make it clear the effect
        ## of more alleles.
        if 'subject_allelic_edits' in fixed_effect_variables:
            #### Recording the effect on the gene expression estimate for the allelic groups with / without the
            #### the adjustment...

            # Getting WITH the allelic group effect adjustment, to show we are correcting for this bias that was present.
            # Now getting it as though we did not model with allelic group effect.
            control_cells = data_entries_df['allelic_edits'].values == 0
            control_cells_edit1 = np.logical_and(control_cells, data_entries_df['subject_allelic_edits'].values==1)
            control_cells_edit2 = np.logical_and(control_cells, data_entries_df['subject_allelic_edits'].values==2)

            r_expr_genes_allelic_group_edit1, pval = stats.spearmanr(
                                                        data_entries_df[f'{gene}_corrected'].values[control_cells],
                                                        data_entries_df['subject_allelic_edits'].values[control_cells]
                                                        )
            r_expr_genes_allelic_group_edit1 = float(r_expr_genes_allelic_group_edit1)

            #### Difference between the means
            min_, max_ = (np.min(data_entries_df[f'{gene}_corrected'].values[control_cells]),
                          np.max(data_entries_df[f'{gene}_corrected'].values[control_cells]))
            range_ = max_ - min_
            mean_edit1 = (np.mean( data_entries_df[f'{gene}_corrected'].values[control_cells_edit1] ) - min_) / range_
            mean_edit2 = (np.mean( data_entries_df[f'{gene}_corrected'].values[control_cells_edit2] ) - min_) / range_
            diff_allelic_group = float(mean_edit2 - mean_edit1)

            # Run the model / correction WITHOUT the correction, this will change {gene}_corrected in data_entries_df
            alleles_beta, alleles_pval, result, model = superb_help.test_edit_and_gene(data_edited, data_entries_df,
                                                                                   gene,
                                                                                   random_effect_variable="subject_cell_index",
                                                                                   fixed_effect_variables=['allelic_edits'],
                                                                                   )

            r_total_no_alleleic_group, r_genes_no_alleleic_group = superb_help.plot_results(result, alleles_beta, data_entries_df, gene, data,
                                                        color_violin=edit_gene_colors[gene],
                                                        out_plots=out_plots + 'without_alleleic_group_effect',
                                                        save_fig=True,
                                                        show_bootstrap_trend=True)

            # Now getting it as though we did not model with allelic group effect.
            r_expr_genes_no_allelic_group_edit1, pval = stats.spearmanr(data_entries_df[f'{gene}_corrected'].values[control_cells],
                                                data_entries_df['subject_allelic_edits'].values[control_cells]
                                                                       )
            r_expr_genes_no_allelic_group_edit1 = float(r_expr_genes_no_allelic_group_edit1)

            min_, max_ = (np.min(data_entries_df[f'{gene}_corrected'].values[control_cells]),
                          np.max(data_entries_df[f'{gene}_corrected'].values[control_cells]))
            range_ = max_ - min_
            mean_edit1 = (np.mean( data_entries_df[f'{gene}_corrected'].values[control_cells_edit1] ) - min_) / range_
            mean_edit2 = (np.mean( data_entries_df[f'{gene}_corrected'].values[control_cells_edit2] ) - min_) / range_
            diff_no_allelic_group = float(mean_edit2 - mean_edit1)

            ### Now without any modelling, just the allelic edit pairing strategy:
            r_orig_expr_genes_edit1, pval = stats.spearmanr(data_entries_df[f'{gene}'].values[control_cells],
                                                data_entries_df['subject_allelic_edits'].values[control_cells]
                                                           )
            r_orig_expr_genes_edit1 = float(r_orig_expr_genes_edit1)

            min_, max_ = (np.min(data_entries_df[f'{gene}'].values[control_cells]),
                          np.max(data_entries_df[f'{gene}'].values[control_cells]))
            range_ = max_ - min_
            mean_edit1 = (np.mean( data_entries_df[f'{gene}'].values[control_cells_edit1] ) - min_) / range_
            mean_edit2 = (np.mean( data_entries_df[f'{gene}'].values[control_cells_edit2] ) - min_) / range_
            diff_orig = float(mean_edit2 - mean_edit1)

            allelic_group_entries.append([diff_allelic_group, 1, gene, 'LM_allelic_group'])

            allelic_group_entries.append([diff_no_allelic_group, 1, gene, 'LM_no_allelic_group'])

            allelic_group_entries.append([diff_orig, 1, gene, 'noLM'])

        # Checking that the selection process reliably removes the relationship with the overall detection rate...
        x, y = data.obs['gene_count'].values[data_entries_df['corr_cell_index'].values], data_entries_df['allelic_edits'].values
        print(f"{gene} correlation between gene and edit detection rate:", stats.pearsonr(x, y))
        # PearsonRResult(statistic=0.0033989763555062917, pvalue=0.8210518967747259)

        # HOWEVER, if we subset to more than 1 allelic edit if available...
        max_allelic_edits = np.max(y)
        if max_allelic_edits > 1:
            print(f"{gene} correlation between gene and >=1 allelic edit detection rate:", stats.pearsonr(x[y>0], y[y>0]))

    edited_gene_stats = pd.DataFrame(edited_gene_stats, index=edited_genes, columns=['allele_beta', 'pval'])

    # Apply FDR correction
    # `method='fdr_bh'` applies the Benjamini-Hochberg procedure
    reject, pvals_corrected, _, _ = smm.multipletests(edited_gene_stats['pval'].values, alpha=0.05, method='fdr_bh')
    edited_gene_stats['padj'] = pvals_corrected
    edited_gene_stats['significant'] = reject

    print("DONE calling edits that significantly effect gene expression")

    # Looking at the bias corrections..
    allelic_group_data = pd.DataFrame(allelic_group_entries, columns=['diff', 'subject_allelic_edits', 'gene', 'model'])
    sb.barplot(allelic_group_data, x='gene', y='diff', hue='model', legend=None)
    plt.ylabel(f"logFC (controls cells: 2 allelic edit group - 1 allelic edit group)")
    utils.dealWithPlot(True, True, True, out_plots+'model_effect_stats/',
                     'correlations_barchar.png', 300)

    version_data = pd.DataFrame(version_entries, columns=['r_genes', 'r_total', 'gene', 'model'])
    keep_bool = [gene_ in on_targets for gene_ in version_data['gene']]
    sb.barplot(version_data.loc[keep_bool,:], x='gene', y='r_genes', hue='model', legend=None)
    plt.ylabel(f"Pearson's R (allelic edits, genes detected)")
    utils.dealWithPlot(True, True, True, out_plots+'model_effect_stats/',
                     'correlations_allelic_edits_barchart.png', 300)

if True: # DEGs for other genes, the highly variable genes selected above.
    ##### Significant edits..
    sig_edits = edited_gene_stats.index.values[reject]
    # Adding back in CHD4, since is an on-target edit that we SHOULD test, since although could not detect DE have
    # qRT-PCR showing it is in-fact DE, i.e. is a false-negative.
    sig_edits = np.unique( list(sig_edits)+['CHD4'] )

    test_genes = genes_to_test

    #### Now for each of these significant edits, checking if they are significantly associated with hvgs!
    sig_edits_stats = {}
    sig_edit_fits = {}
    for i, gene in enumerate( sig_edits ):

        data_entries_df = superb_help.construct_allelic_pairings(data_edited, gene,
                                                                 umap_key='X_umap',
                                                                 condition_key=gene_to_condition[gene],
                                                                 dosage_key='dosages_genes_manual-edits-only',
                                                                 min_cells_with_allele=25,
                                                                 max_total_umi_dist=1000, #Says total UMI but actually means gene count
                                                                 n_reps=10,
                                                                 total_count_key='gene_count',
                                                                 method='pair_each_allele',
                                                                 )

        if len(np.unique(data_entries_df['subject_allelic_edits'].values)) == 1:
            fixed_effect_variables = ['allelic_edits']
        else:
            fixed_effect_variables = ['allelic_edits', 'subject_allelic_edits']

        sig_edit_stats = np.zeros((len(test_genes), 2))
        sig_edit_fit = {}
        for j, hvg in enumerate(test_genes):
            alleles_beta, alleles_pval, result, model = superb_help.test_edit_and_gene(data_edited, data_entries_df,
                                                                                       hvg,
                                                                                       random_effect_variable="subject_cell_index",
                                                                                       fixed_effect_variables=fixed_effect_variables
                                                                                       )
            if type(alleles_beta) != float:
                alleles_beta = alleles_beta[0]
                alleles_pval = alleles_pval[0]

            sig_edit_stats[j, :] = [alleles_beta, alleles_pval]
            sig_edit_fit[hvg] = [result, model]

        sig_edit_fits[gene] = sig_edit_fit

        sig_edit_stats = pd.DataFrame(sig_edit_stats, index=test_genes, columns=['allele_beta', 'pval'])
        sig_edit_stats['-log10_pval'] = -np.log10(sig_edit_stats['pval'].values)

        # Apply FDR correction
        # `method='fdr_bh'` applies the Benjamini-Hochberg procedure
        reject, pvals_corrected, _, _ = smm.multipletests(sig_edit_stats['pval'].values, alpha=0.05, method='fdr_bh')
        sig_edit_stats['padj'] = pvals_corrected
        sig_edit_stats['significant'] = reject

        sig_edits_stats[gene] = sig_edit_stats.sort_values('padj')

    print("DONE calling DE HVGs with respect to allelic edits.")

    ##### Now need to estimate the correlation between each of the edits, so can then perform fine-mapping with FINEMAP !
    utils.saveAsPickle(f'{out_dir}de_results.pkl',
                     {"sig_edits_stats": sig_edits_stats,
                      "edited_gene_stats": edited_gene_stats,
                      })
    utils.saveAsPickle(f'{out_dir}de_models.pkl',
                     {"sig_edit_fits": sig_edit_fits,
                      "fit_results": fit_results})

    print(f"Saved DE results to: {out_dir}de_results.pkl")
    print(f"Saved DE models to: {out_dir}de_models.pkl")

else:
    print(f"Loading {out_dir}de_results.pkl")
    de_data = spl.loadPickle(f'{out_dir}de_results.pkl')
    print("Finished loading")
    edited_gene_stats = de_data["edited_gene_stats"]
    sig_edits_stats = de_data["sig_edits_stats"]

    sig_edits = list(sig_edits_stats.keys())

# But isn't true for all the genes, mostly the on-targets. So it might actually be true, i.e. lots of small effects.
sig_bools = [np.logical_and(sig_edits_stats[gene_]['padj'].values<0.01,
                            np.abs(sig_edits_stats[gene_]['allele_beta'].values)>=0.2)
             for gene_ in sig_edits]
sig_edit_genes = [list(set(list(sig_edits_stats[gene_].index[ sig_bools[i] ])))
                    for i, gene_ in enumerate(sig_edits)]
print([f"{gene}: {len(edit_genes)}" for gene, edit_genes in zip(sig_edits, sig_edit_genes)])
#['ARID1A: 653', 'CHD3: 303', 'CHD4: 251', 'SMARCA4: 650', 'USP9X: 156']

min_overlap = 10
utils.upset_plot(sig_edit_genes, list(sig_edits), show=False, min_subset_size=min_overlap)
utils.dealWithPlot(True, True, True, out_plots, 'de_gene_edit_overlap.png', 300)

#### Again as svg so can make it pretty!!!
utils.upset_plot(sig_edit_genes, list(sig_edits), show=False, min_subset_size=min_overlap)
utils.dealWithPlot(True, True, True, out_plots, 'de_gene_edit_overlap.svg', 300)

utils.upset_plot(sig_edit_genes, list(sig_edits), show=False, min_subset_size=1)
utils.dealWithPlot(True, True, True, out_plots, 'de_gene_edit_overlap_full.png', 300)

#### Estimate associations between edits, the overlap of cells...
data_entries_dict = {}
for i, gene in enumerate( sig_edits ):
    data_entries_dict[gene] = superb_help.construct_allelic_pairings(data_edited, gene,
                                                                 umap_key='X_umap',
                                                                 condition_key=gene_to_condition[gene]
                                                                 )

# Counting cell barcode overlaps..
sig_edit_cells = [list(set(list(data_entries_dict[gene_]['corr_cell_barcode'].values[
                        data_entries_dict[gene_]['allelic_edits'].values>0])))
                  for gene_ in sig_edits]

utils.upset_plot(sig_edit_cells, list(sig_edits), show=False, min_subset_size=1)
utils.dealWithPlot(True, True, True, out_plots, 'edited_cells_overlap.png', 300)

# Again as svg so can modify in illustrator better!
utils.upset_plot(sig_edit_cells, list(sig_edits), show=False, min_subset_size=1)
utils.dealWithPlot(True, True, True, out_plots, 'edited_cells_overlap.svg', 300)

print("DONE looking at overlaps between DE genes AND edited cells")

import superb_guide_inference as sgi
import importlib as imp
imp.reload( sgi )

edit_cell_props = sgi.get_overlap_props(data,
                                        'dosages_genes', cols=sig_edits
                                        )

off_targets_ = [gene_ for gene_ in sig_edits if gene_ not in on_targets]

sb.heatmap(edit_cell_props, vmin=0, vmax=0.8, annot=True, fmt=".2f"
           )
utils.dealWithPlot(True, True, True, out_plots, 'edited_cells_overlap_heatmap.png', 300)

##### Let's try to split this, so can more clearly see..
ax = sb.heatmap(edit_cell_props.loc[on_targets,off_targets_], vmin=0.05, vmax=0.08, annot=True, fmt=".2f")
ax.set_yticklabels(ax.get_yticklabels(), rotation=0)
ax.set_xticklabels(ax.get_xticklabels(), rotation=0)
utils.dealWithPlot(True, True, True, out_plots,
                 'edited_cells_ontargets_overlap_heatmap.png', 300)

ax = sb.heatmap(edit_cell_props.loc[off_targets_,on_targets], vmin=0, vmax=1,
           annot=True, fmt=".2f")
ax.set_yticklabels(ax.get_yticklabels(), rotation=0)
ax.set_xticklabels(ax.get_xticklabels(), rotation=0)
utils.dealWithPlot(True, True, True, out_plots,
                 'edited_cells_offtargets_overlap_heatmap.png', 300)

# co-ocurring on-targets and off-targets
edit_cell_co_occur = edit_cell_props > .06
co_occur_indices = np.where(edit_cell_co_occur.values)

edit_to_off_targets = defaultdict(list)
[edit_to_off_targets[sig_edits[index1]].append(sig_edits[index2])
 for index1, index2 in zip(co_occur_indices[0], co_occur_indices[1])]

#########################################################################################################################
                                # The over-representation analysis #
#########################################################################################################################

####  now plot enrichment before and after of determined BAF / NuRD targets determined from ENCODE.
baf_genes = set([gene.strip('\n') for gene in open(f'{data_dir2}smarca4_k562_genes.txt', 'r')])
baf_enh_genes = set([gene.strip('\n') for gene in open(f'{data_dir2}smarca4_k562_genes.enhancer.genes.txt', 'r')])
nurd_genes = set([gene.strip('\n') for gene in open(f'{data_dir2}chd4_k562_genes.txt', 'r')])

# Subsetting to just the tested genes..
baf_genes = baf_genes.intersection(list(set(genes_to_test)))
baf_enh_genes = baf_enh_genes.intersection(list(set(genes_to_test)))
nurd_genes = nurd_genes.intersection(list(set(genes_to_test)))
len(baf_genes), len(baf_enh_genes), len(nurd_genes)

#### Visualising relationship between the mean expression of the genes, allelic beta, and significance (should not be correlated)
p_cut = .01
min_effect_cut = 0.2
for gene_, stats in sig_edits_stats.items():

    means = data[:,stats.index.values].var['means'].values
    stats['mean'] = means

    effect_sizes = stats['allele_beta'].values
    log10_padjs = stats['-log10_pval'].values
    padjs = stats['padj'].values
    sig = np.logical_and(padjs < p_cut, np.abs(effect_sizes)>=min_effect_cut)
    up_ = effect_sizes > 0

    ### MA plots
    fig, ax = plt.subplots(figsize=(6,5))
    fig.suptitle(f"{gene_} knock-down DEGs (n={sum(sig)})")
    ax.scatter(means[~sig], effect_sizes[~sig], color='grey', label='non-significant')
    ax.scatter(means[np.logical_and(sig, up_)], effect_sizes[np.logical_and(sig, up_)], color='red', label='DE up')
    ax.scatter(means[np.logical_and(sig, ~up_)], effect_sizes[np.logical_and(sig, ~up_)], color='blue', label='DE down')
    plt.xlabel('gene mean expression')
    plt.ylabel('estimated allelic effect size')
    plt.legend()
    utils.dealWithPlot(True, True, True, out_plots, f"{gene_}_MAplot_wLegend.png",300)

    fig, ax = plt.subplots(figsize=(6,5))
    fig.suptitle(f"{gene_} knock-down DEGs (n={sum(sig)})")
    ax.scatter(means[~sig], effect_sizes[~sig], color='grey', label='non-significant')
    ax.scatter(means[np.logical_and(sig, up_)], effect_sizes[np.logical_and(sig, up_)], color='red', label='DE up')
    ax.scatter(means[np.logical_and(sig, ~up_)], effect_sizes[np.logical_and(sig, ~up_)], color='blue', label='DE down')
    plt.xlabel('gene mean expression')
    plt.ylabel('estimated allelic effect size')
    utils.dealWithPlot(True, True, True, out_plots, f"{gene_}_MAplot.png",300)

    ### Volcano plot
    fig, ax = plt.subplots(figsize=(6,5))
    fig.suptitle(f"{gene_} knock-down DEGs (n={sum(sig)})")
    ax.scatter(effect_sizes[~sig], log10_padjs[~sig], color='grey', label='non-significant')
    ax.scatter(effect_sizes[np.logical_and(sig, up_)], log10_padjs[np.logical_and(sig, up_)], color='red', label='DE up')
    ax.scatter(effect_sizes[np.logical_and(sig, ~up_)], log10_padjs[np.logical_and(sig, ~up_)], color='blue', label='DE down')
    plt.xlabel('estimated allelic effect size')
    plt.ylabel('-log10(p_adj)')
    utils.dealWithPlot(True, True, True, out_plots, f"{gene_}_volcano_wLegend.png",300)

########### Calling the DE genes
p_cut = .01
min_effect_cut = 0.2

# Trying up vs down-regulated genes as well!
gene_to_marginal_de = superb_help.get_de_dict(sig_edits_stats, True,
                                              cutoff=p_cut, effect_size_cutoff=min_effect_cut,
                                              update_sig_col=True)

gene_to_marginal_de_up = superb_help.get_de_dict(sig_edits_stats, True, cutoff=p_cut, down=False,
                                                 effect_size_cutoff=min_effect_cut,
                                              update_sig_col=False)
gene_to_marginal_de_down = superb_help.get_de_dict(sig_edits_stats, True, cutoff=p_cut, up=False,
                                                   effect_size_cutoff=min_effect_cut,
                                              update_sig_col=False)

de_counts = {target: len(gene_to_marginal_de[target]) for target in gene_to_marginal_de}
print(de_counts)

####### Let's now save these as an excel sheet, this will become a supplementary file..
utils.writeDFsToExcelSheets(f"{out_dir}allelic_edit_DEGs.xlsx",
                                    [edited_gene_stats]+list(sig_edits_stats.values()),
                                       sheet_names=['edited_gene_stats']+\
                                                   [f"{gene}_DEG_stats" for gene in sig_edits_stats.keys()])

#### Saving as text files also, so can easily load into R...
for gene_, stats_ in sig_edits_stats.items():
    stats_.to_csv(f"{out_dir}{gene_}_allelic_edit_DEGs.txt", sep='\t')

###### Getting the odds-ratios...
nurd_edited = ['CHD3', 'CHD4', 'USP9X']
baf_edited = ['ARID1A', 'SMARCA4',
              ]

nurd_gene_sets = {'NURD_targets': nurd_genes}
baf_gene_sets = {'BAF_prom_targets': baf_genes,
                 'BAF_enh_targets': baf_enh_genes}

# Loading relevant gene sets to do with the off-target genes!!!
gene_set_files = [f'{data_dir2}HALLMARK_G2M_CHECKPOINT.v2024.1.Hs.gmt']
gene_set_alloc = [baf_gene_sets, nurd_gene_sets,
                  ]
for i, file_name in enumerate(gene_set_files):
    set_name = file_name.split('/')[-1].split('.')[0]
    gene_set = set(list(open(file_name, 'r'))[0].split('\t')[2:]).intersection(list(set(genes_to_test)))
    gene_set_alloc[i][set_name] = gene_set

nrf1_targets = [line.strip('\n') for line in open(f"{data_dir2}nrf1_k562_peaks.genes.open.txt", 'r')]
nrf1_targets = set(nrf1_targets).intersection(list(set(genes_to_test)))

gene_set_alloc[1]['NRF1_targets'] = nrf1_targets

# NOTE this is where I subset to the USP9X-specific genes, by taking the set difference with the DEGs associated
# with the on-target edit at CHD3 for the guide that also causes the USP9X edit.
for gene_to_marginal in [gene_to_marginal_de, gene_to_marginal_de_up, gene_to_marginal_de_down]:
    gene_to_marginal['USP9X'] = gene_to_marginal['USP9X'].difference(gene_to_marginal['CHD3'])

#### WOW quite the overlap! Very cool.
print(gene_to_marginal_de['USP9X'].intersection(nrf1_targets))
print(len(gene_to_marginal_de['USP9X'].intersection(nrf1_targets)))

tested_genes = sig_edits_stats[on_targets[0]].index.values

# Was from the overall
marginal_results, marginal_tables = superb_help.compare_de_enrichments(
                                                                    sig_edits, nurd_edited, nurd_gene_sets, baf_gene_sets,
                                                                    gene_to_marginal_de, tested_genes)

#######
marginal_results_up, marginal_tables_up, = superb_help.compare_de_enrichments(
                                                                    sig_edits, nurd_edited, nurd_gene_sets, baf_gene_sets,
                                                              gene_to_marginal_de_up,  tested_genes)

marginal_results_down, marginal_tables_down = superb_help.compare_de_enrichments(
                                                                    sig_edits, nurd_edited, nurd_gene_sets, baf_gene_sets,
                                                          gene_to_marginal_de_down, tested_genes)

print("Marginal all:", marginal_results)

print("Marginal up", marginal_results_up)

print("Marginal down", marginal_results_down)

print("Marginal all:", marginal_results)
print("Marginal down", marginal_results_down)
print("Marginal up", marginal_results_up)

### Writing these results to a supplementary excel sheet for the paper!!!
ora_stats = []
sheet_names = []
for name_, results_ in zip(['all', 'up', 'down'], [marginal_results, marginal_results_up, marginal_results_down]):

    for comparison_, stats_ in results_.items():
        ora_stats.append(stats_)
        sheet_names.append( f"{comparison_}_{name_}" )

utils.writeDFsToExcelSheets(f"{out_dir}allelic_edit_DEG_ORA_stats.xlsx", ora_stats, sheet_names=sheet_names)

#### Let's also write out the contingency tables.
cont_tables = []
cont_sheet_names = []
for name_, results_ in zip(['all', 'up', 'down'], [marginal_tables, marginal_tables_up, marginal_tables_down]):

    for comparison_, table_ in results_.items():
        gene_name = comparison_.split('_')[0]
        gene_set_ = "_".join(comparison_.split('_')[1:])
        cont_tables.append( pd.DataFrame(table_, index=[f'{gene_name}_{name_}DEG', f'{gene_name}_not{name_}DEG'],
                                         columns=[gene_set_, f"not{gene_set_}"]) )

utils.writeDFsToExcelSheets(f"{out_dir}allelic_edit_DEG_ORA_contigency_tables.xlsx",
                             cont_tables, sheet_names=list(np.array(list(range(len(cont_tables)))).astype(str))
                             )

#### Also creating a table of what these gene sets are...
gene_set_dfs = []
gene_set_df_names = ['BAF_gene_sets', 'NuRD_gene_sets']

for gene_sets_name, gene_sets_dict in zip(['BAF_gene_sets', 'NuRD_gene_sets'], gene_set_alloc):

    gene_sets_list = []
    gene_set_list_names = []
    for gene_set_name, gene_set in gene_sets_dict.items():

        gene_sets_list.append( list(gene_set) )
        gene_set_list_names.append( gene_set_name )

    gene_sets_df = form.listsToFrame(gene_sets_list, columns=gene_set_list_names)
    gene_set_dfs.append( gene_sets_df )

utils.writeDFsToExcelSheets(f"{out_dir}functional_gene_sets.xlsx",
                             gene_set_dfs, sheet_names=gene_set_df_names
                             )

# Now let's visualise these results...
def plot_fet_results(fet_results, color_dict, odds_cut,
                     xlabel='odds ratio',
                     max_error=None, rotate=False, order=None, figsize=(25, 5),
                     s=500, lw=3, log=False, label_order=None):
    if type(order)==type(None):
        if type(label_order) == type(None):
            order = np.argsort(fet_results['odds_ratio'].values)
            label_order = fet_results.index.values
        else:
            order = np.array([np.where(fet_results.index.values==label)[0][0] for label in label_order])

    else:
        label_order = fet_results.index.values[order]

    c = [color_dict[celltype] for celltype in label_order]
    x = list(range(fet_results.shape[0]))

    upper = fet_results['odds_ratio_0.95_upper'].values[order]
    lower = fet_results['odds_ratio_0.95_lower'].values[order]
    if type(max_error)!=type(None):
        upper[upper>max_error] = max_error

    odds = fet_results['odds_ratio'].values[order]
    if log:
        odds = np.log2(odds)
        upper = np.log2(upper)
        lower = np.log2(lower)
        odds_cut = np.log2(odds_cut)

        xlabel = f'log2 {xlabel}'

    if not rotate:
        fig, ax = plt.subplots(figsize=(25, 5))
        plt.vlines(odds_cut, min(x) - 0.5, max(x) + 0.5, linestyles='dashed', color='lightgrey')
        plt.scatter(odds, x,
                    s=500, c=c)
        plt.hlines(x, lower,
                   upper, colors='grey')
        plt.yticks(x, list(fet_results.index.values[order]))
        ax.spines['right'].set_visible(False)
        ax.spines['top'].set_visible(False)
        plt.xlabel(xlabel)
    else:
        fig, ax = plt.subplots(figsize=figsize)
        plt.hlines(odds_cut, min(x) - 0.5, max(x) + 0.5, linestyles='dashed', color='lightgrey', linewidth=lw)
        plt.scatter(x, odds,
                    s=s, c=c)
        plt.vlines(x, lower,
                   upper, colors='darkgrey', linewidth=lw)
        plt.xticks(x, list(fet_results.index.values[order]), rotation=90)
        ax.spines['right'].set_visible(False)
        ax.spines['top'].set_visible(False)
        plt.ylabel(xlabel)
        ax.spines['left'].set_linewidth(lw)
        ax.spines['bottom'].set_linewidth(lw)

    return fig, ax, order

color_dict = utils.getColors(list(marginal_results['nurd'].index.values)+list(marginal_results['baf'].index.values))

result_names = ['overall', 'up', 'down']
order = None
for i, (name, results) in enumerate(zip(result_names, [marginal_results, marginal_results_up, marginal_results_down])):
    fig, ax, order = plot_fet_results(results['nurd'], color_dict, odds_cut=1, max_error=20, rotate=True,
                                          order=order)
    plt.title(f'NURD {name} DE genes')
    utils.dealWithPlot(True, True, True, out_plots, f'NURD_{name}_odds-ratios.png', 300)

result_names = ['down', 'overall', 'up', ]
order = None
for i, (name, results) in enumerate(zip(result_names, [marginal_results_down, marginal_results, marginal_results_up, ])):
    fig, ax, order = plot_fet_results(results['baf'], color_dict, odds_cut=1, max_error=20, rotate=True,
                                          order=order)
    plt.title(f'BAF {name} DE genes')
    utils.dealWithPlot(True, True, True, out_plots, f'BAF_{name}_odds-ratios.png', 300)

##### Making a more square figure for the actual illustrator figure
figsize, lw, s = (10,10), 5, 1000
result_names = ['overall', 'up', 'down']
order = None
for i, (name, results) in enumerate(zip(result_names, [marginal_results, marginal_results_up, marginal_results_down])):
    fig, ax, order = plot_fet_results(results['nurd'], color_dict, odds_cut=1, max_error=20, rotate=True,
                                          order=order, figsize=figsize, lw=lw, s=s)
    plt.title(f'NURD {name} DE genes')
    utils.dealWithPlot(True, True, True, out_plots, f'NURD_{name}_odds-ratios.png', 300)

result_names = ['down', 'overall', 'up', ]
order = None
for i, (name, results) in enumerate(zip(result_names, [marginal_results_down, marginal_results, marginal_results_up, ])):
    fig, ax, order = plot_fet_results(results['baf'], color_dict, odds_cut=1, max_error=20, rotate=True,
                                          order=order, figsize=figsize, lw=lw, s=s)
    plt.title(f'BAF {name} DE genes')
    utils.dealWithPlot(True, True, True, out_plots, f'BAF_{name}_odds-ratios.png', 300)

##### This time will do the log2 odds ratio, and set the label order based on figure aesthetics.

#### make the colors more consistent...
set_names = list(marginal_results['nurd'].index.values)+list(marginal_results['baf'].index.values)
color_dict = {set_name: edit_gene_colors[set_name.split('_')[0]] for set_name in set_names}

figsize, lw, s = (10,10), 5, 1000
log_ = True
result_names = ['overall', 'up', 'down']
order = None
nurd_order = ['USP9X_NRF1_targets', 'CHD4_NRF1_targets', 'CHD3_NRF1_targets',
              'USP9X_NURD_targets', 'CHD4_NURD_targets', 'CHD3_NURD_targets'][::-1]
for i, (name, results) in enumerate(zip(result_names, [marginal_results, marginal_results_up, marginal_results_down])):
    fig, ax, order = plot_fet_results(results['nurd'], color_dict, odds_cut=1, max_error=20, rotate=True,
                                          order=order, figsize=figsize, lw=lw, s=s, log=log_, label_order=nurd_order)
    plt.title(f'NURD {name} DE genes')
    utils.dealWithPlot(True, True, True, out_plots, f'NURD_{name}_odds-ratios.png', 300)

baf_order = ['ARID1A_HALLMARK_G2M_CHECKPOINT', 'SMARCA4_HALLMARK_G2M_CHECKPOINT',
             'ARID1A_BAF_enh_targets', 'SMARCA4_BAF_enh_targets',
             'ARID1A_BAF_prom_targets', 'SMARCA4_BAF_prom_targets'][::-1]
result_names = ['down', 'overall', 'up', ]
order = None
for i, (name, results) in enumerate(zip(result_names, [marginal_results_down, marginal_results, marginal_results_up, ])):
    fig, ax, order = plot_fet_results(results['baf'], color_dict, odds_cut=1, max_error=20, rotate=True,
                                          order=order, figsize=figsize, lw=lw, s=s, log=log_, label_order=baf_order)
    plt.title(f'BAF {name} DE genes')
    utils.dealWithPlot(True, True, True, out_plots, f'BAF_{name}_odds-ratios.png', 300)
```
