## Additional Files for "Joint single-cell capture of Cas9 edits and transcriptomes reveals on- and off-target effects on gene expression": superb_guide_inference.py

```
""" Functions related to imputing
    edits-to-guides based on homology/coincidence of edits in the same cells that have
    an on-target edit

    NOTE most of these functions were written by Mickey Lorenzini, but have adapted them and optimised, particularly
    replacing the homology with an alignment-based approach..
"""

import time

import numpy as np
import pandas as pd

import pysam

import re

from collections import defaultdict

from Bio import pairwise2 # pip install biopython==1.83 # New version have changed this

# Run hamming code (homology-based guide inference) to for auto coinc_im_guide dictionary
# and for coincidence-based guide inference optimization...

# Functions to get edit site sequences with minimum hamming abs_offset to guide RNA sequences
def find_pams(query_seq, pam_regexp, is_reverse):
    """
    Find all occurrences of a PAM motif in the query sequence.
    Return two lists:
    - The 0-based position of of the 5' end of each PAM ("N" in "NGG", or "N" in "CNN" if is_reverse is True).
    - The PAM sequences (each query sequence match).
    """
    # cut position defined as the 3rd base upstream (5' direction) of the pam start position (5' end of pam)
    cut_positions = []
    pams = []

    for match in pam_regexp.finditer(query_seq):
        if is_reverse:
            # reverse strand, cut position is last matched character (pattern end -1) plus 3
            cut_positions.append(match.end(1) + 2)
            pams.append(match.group(0))
        else:
            # cut position is first matched character minus 3
            cut_positions.append(match.start(1) - 3)
            pams.append(match.group(0))

    # return cut positions as zero-based query sequence position
    return cut_positions, pams


def fetch_each_target(fasta, chrom, cut_coord, is_reverse):
    """
    Fetch a 23 base reference sequence at each PAM.
    For forward strand PAMs, fetch the 20 bases upstream + the 3 base PAM motif.
    For reverse strand PAMs, fetch the 3 base PAM motif + the 20 bases downstream, and return the reverse complement.
    """

    # helper function for reverse complements
    def rev_comp(seq):
        complement = str.maketrans('ACGTacgt', 'TGCAtgca')
        return seq.translate(complement)[::-1]

    # reverse strand behavior
    if is_reverse:
        # TEST: return just PAM
        # start = cut_coord - 5
        # end = cut_coord - 2

        # return 23 bp target sequence
        start = cut_coord - 5
        end = cut_coord + 18
        sequence = fasta.fetch(chrom, start, end)
        # Reverse complement the sequence
        sequence = rev_comp(sequence)
    # fwd strand behavior
    else:
        # TEST: return just PAM
        # start = cut_coord + 3
        # end = cut_coord + 6

        # return 23 bp target sequence
        start = cut_coord - 17
        end = cut_coord + 6
        sequence = fasta.fetch(chrom, start, end)

    return start, end, sequence

# Converts lists of values into comma-separated strings, add to storage lists, with helper function
def csv_append(input_list, storage_csv):
    values_to_str = ",".join(map(str, input_list))
    storage_csv.append(values_to_str)

def fetch_close_targets(edit_site_info_df, ref_fasta, n_targets, query_range):
    print('Getting target sequences...')
    start_time = time.time()

    # Read edit_site_info.txt into a DataFrame
    # df = pd.read_csv(edit_site_txt, sep='\t')
    # df = df.rename(columns={
    #     'name': 'edit_site', 'chr': 'chrom', 'pos': 'position', 'copy-number': 'copy_number',
    # })
    # NOTE don't like the renaming here, cause will likely get a little confusing.

    # Will subset to the relevant columns, in-case are running it under different set of settings
    new_cols = ['query_seq', 'n_close', 'close_offsets', 'abs_offsets', 'close_starts', 'close_ends', 'close_targets',
                'close_strands']
    present = [col for col in new_cols if col in edit_site_info_df.columns]
    if len(present) > 0: # Want it to only drop them if present
        df = edit_site_info_df.drop(columns=present)
    else:
        df = edit_site_info_df

    # print(f'New edit_site_info column names: {df.columns}')

    # Open the reference genome fasta file
    fasta = pysam.FastaFile(ref_fasta)

    # Initialize lists to store comma-separated values
    query_csv = []
    fwd_cuts_csv = []
    fwd_pams_csv = []
    fwd_offsets_csv = []
    rev_cuts_csv = []
    rev_pams_csv = []
    rev_offsets_csv = []

    both_offsets_csv = []
    both_strands_csv = []
    close_offsets_csv = []
    close_abs_offsets_csv = []
    close_strands_csv = []
    n_close = []

    starts_csv = []
    ends_csv = []
    seqs_csv = []

    # Compile regex patterns for forward and reverse PAMs
    # Use LOOKAHEAD ASSERTION to find overlapping matches
    fwd_pam_regexp_1 = re.compile(r'(?=([ACGT]GG))')  # Pattern to find 'NGG'
    rev_pam_regexp_1 = re.compile(r'(?=(CC[ACGT]))')  # Pattern to find 'CCN'

    fwd_pam_regexp_2 = re.compile(r'(?=([ACGT]AG))')  # Pattern to find 'NAG'
    rev_pam_regexp_2 = re.compile(r'(?=(CT[ACGT]))')  # Pattern to find 'CTN'

    # Effectively removes the pam limitation...
    fwd_pam_regexp_3 = re.compile(r'(?=([ACGT]))')  # Pattern to find 'N'
    rev_pam_regexp_3 = re.compile(r'(?=([ACGT]))')  # Pattern to find 'N'

    fwd_pam_regexps = [fwd_pam_regexp_1, fwd_pam_regexp_2,
                        #fwd_pam_regexp_3
                       ]
    rev_pam_regexps = [rev_pam_regexp_1, rev_pam_regexp_2,
                        #rev_pam_regexp_3
                       ]

    # Iterate through each row in the DataFrame
    for index, row in df.iterrows():
        name = row['name'] #row['edit_site']
        chrom = row['chr'] #row['chrom']
        position = row['pos'] #row['position']

        # Calculate the start and end positions for the query sequence (0-based, half-open)
        query_start = position - query_range
        query_end = position + query_range

        # Fetch the query sequence from the reference genome
        query_seq = fasta.fetch(chrom, query_start, query_end).upper()
        query_csv.append(query_seq)

        fwd_cuts, fwd_pams, fwd_offsets = [], [], []
        rev_cuts, rev_pams, rev_offsets = [], [], []
        for fwd_pam_regexp, rev_pam_regexp in zip(fwd_pam_regexps, rev_pam_regexps):

            # Get lists of all cut positions within query sequence (and corresponding PAMS for qc)
            # fwd strand
            fwd_cuts_, fwd_pams_ = find_pams(query_seq, fwd_pam_regexp, is_reverse=False)
            # Get offset distance between cut position and query center (edit site prediction)
            fwd_offsets_ = [cut_position - query_range for cut_position in fwd_cuts_]
            fwd_cuts.extend( fwd_cuts_ )
            fwd_pams.extend( fwd_pams_ )
            fwd_offsets.extend( fwd_offsets_ )

            rev_cuts_, rev_pams_ = find_pams(query_seq, rev_pam_regexp, is_reverse=True)
            rev_offsets_ = [cut_position - query_range for cut_position in rev_cuts_]
            rev_cuts.extend( rev_cuts_ )
            rev_pams.extend( rev_pams_ )
            rev_offsets.extend( rev_offsets_ )

        csv_append(fwd_cuts, fwd_cuts_csv)
        csv_append(fwd_pams, fwd_pams_csv)
        csv_append(fwd_offsets, fwd_offsets_csv)
        csv_append(rev_cuts, rev_cuts_csv)
        csv_append(rev_pams, rev_pams_csv)
        csv_append(rev_offsets, rev_offsets_csv)

        # PAMs and cut positions (within query) look good! continue...

        # For CLOSEST n PAMs, fetch lists of target information (start, end, sequences)
        # Determine closest n PAMs...
        # Combine fwd and rev offset lists...
        both_offsets = fwd_offsets + rev_offsets
        # Track strand to apply proper coordinate ranges later
        both_strands = ['+' for _ in fwd_offsets] + ['-' for _ in rev_offsets]
        # Generate a tuple of PAM info, ordered by closest to edit site prediction (absolute offset)
        tuple_by_closest = sorted(zip(both_offsets, both_strands), key=lambda x: abs(x[0]))
        # Handle ties for the nth closest sequence
        # Initialize lists
        close_offsets = []
        close_abs_offsets = []
        close_strands = []
        current_abs_offset = set()
        # get closest targets
        for offset, strand in tuple_by_closest:
            if len(close_offsets) < n_targets or abs(offset) in current_abs_offset:
                close_offsets.append(offset)
                close_abs_offsets.append(abs(offset))
                close_strands.append(strand)
                current_abs_offset.add(abs(offset))
            else:
                break
        csv_append(both_offsets, both_offsets_csv)
        csv_append(both_strands, both_strands_csv)
        csv_append(close_offsets, close_offsets_csv)
        csv_append(close_abs_offsets, close_abs_offsets_csv)
        csv_append(close_strands, close_strands_csv)
        n_close.append(len(close_offsets))

        # Fetch start, end, sequence for each of closest n PAMs
        # initialize storage lists
        starts = []
        ends = []
        seqs = []  # length 23, includes pam
        # Convert cut position (wrt query) to genome coordinate
        cut_coords = [position + offset for offset in close_offsets]
        # Convert strand symbol to boolean for is_reverse
        is_reverse = [strand == '-' for strand in close_strands]
        # Get target info per closest PAM from tuple of coords and strands
        for cut_coord, is_rev in zip(cut_coords, is_reverse):
            start, end, sequence = fetch_each_target(fasta, chrom, cut_coord, is_rev)
            starts.append(start)
            ends.append(end)
            seqs.append(sequence)

        csv_append(starts, starts_csv)
        csv_append(ends, ends_csv)
        csv_append(seqs, seqs_csv)

    # Close the reference genome file
    fasta.close()

    # Dataframe outputs
    df['query_seq'] = query_csv
    # df['fwd_cuts'] = fwd_cuts_csv
    # df['fwd_pams'] = fwd_pams_csv
    # df['rev_cuts'] = rev_cuts_csv
    # df['rev_pams'] = rev_pams_csv
    # df['fwd_offsets'] = fwd_offsets_csv
    # df['rev_offsets'] = rev_offsets_csv
    # df['both_offsets'] = both_offsets_csv
    # df['both_strands'] = both_strands_csv
    # df['close_offsets'] = close_offsets_csv
    df['n_close'] = n_close
    df['close_offsets'] = close_offsets_csv
    df['abs_offsets'] = close_abs_offsets_csv
    df['close_starts'] = starts_csv
    df['close_ends'] = ends_csv
    df['close_targets'] = seqs_csv
    df['close_strands'] = close_strands_csv

    # Calculate and print the total run time
    end_time = time.time()
    total_sec = (end_time - start_time)
    total_min = total_sec / 60
    print(f"Done. Run time: {total_min:.2f} min ({total_sec:.2f} s)")

    return df


# now define function that makes long form of target df for input into variant seq generator...
def expand_targets(df):
    print('Expanding target sequences...')
    start_time = time.time()

    # Split targets, pams, and strands into character lists
    targets_split = df['close_targets'].str.split(',')
    pams_split = df['close_targets'].apply(lambda x: [item[-3:] for item in x.split(',')])
    strands_split = df['close_strands'].str.split(',')

    # split offsets, starts, ends into integer lists
    offsets_split = df['close_offsets'].str.split(',').apply(lambda x: list(map(int, x)))
    abs_offsets_split = df['abs_offsets'].str.split(',').apply(lambda x: list(map(int, x)))
    starts_split = df['close_starts'].str.split(',').apply(lambda x: list(map(int, x)))
    ends_split = df['close_ends'].str.split(',').apply(lambda x: list(map(int, x)))

    # Flatten the list of lists into a single list
    seqs_flat = [item.strip() for sublist in targets_split for item in sublist]
    pams_flat = [item.strip() for sublist in pams_split for item in sublist]
    strands_flat = [item.strip() for sublist in strands_split for item in sublist]
    offsets_flat = [item for sublist in offsets_split for item in sublist]
    abs_offsets_flat = [item for sublist in abs_offsets_split for item in sublist]
    starts_flat = [item for sublist in starts_split for item in sublist]
    ends_flat = [item for sublist in ends_split for item in sublist]

    # Repeat each row to match the length of split targets
    long_df = df.loc[df.index.repeat(targets_split.apply(len))].reset_index(drop=True)

    # Generate 'target_name' by combining 'edit_site' and a 1-based integer index
    target_names = [f"{name}_{i + 1}" for name, count in zip(df.index.values, targets_split.apply(len)) for i in
                    range(count)]

    # Make output dataframe
    long_df['target_name'] = target_names
    long_df['target_start'] = starts_flat
    long_df['target_end'] = ends_flat
    long_df['target_seq'] = seqs_flat
    long_df['target_pam'] = pams_flat
    long_df['strand'] = strands_flat
    long_df['offset'] = offsets_flat
    long_df['abs_offset'] = abs_offsets_flat
    long_df = long_df.sort_values('target_name').reset_index(drop=True)

    # Calculate and print the total run time
    end_time = time.time()
    total_sec = (end_time - start_time)
    total_min = total_sec / 60
    print(f"Done. Run time: {total_min:.2f} min ({total_sec:.2f} s)")

    return long_df


def make_1bp_indels(df, seed_size=5):
    print('Generating 1 bp indel variant sequences...')
    start_time = time.time()

    # Lists to store new variant information
    variant_info = []

    # Iterate over each row in the DataFrame
    for index, row in df.iterrows():
        key = row['target_name']
        ref_seq = row['target_seq'][:-3].upper()

        # Add the reference sequence itself
        variant_info.append((key, f"{key}_ref_0", ref_seq))

        # Generate positional variants up to defined seed region size, by scanning a 1-bp indel from 5' end toward seed
        # generate variants that MAINTAIN SEED ALIGNMENT
        # start with second position in guide, first position is non-bulge sequence
        for i in range(1, len(ref_seq) - seed_size):
            # 1-bp deletions, add terminal 'N' for equal length for Hamming distances (1-based positions)
            del_seq = '-' + ref_seq[:i] + ref_seq[i + 1:]
            # 1-bp insertions by inserting 'N', trim terminal base for equal length
            ins_seq = ref_seq[1:i + 1] + '-' + ref_seq[i + 1:]
            # add data to storage list
            variant_info.append((key, f"{key}_del_{i + 1}", del_seq))
            variant_info.append((key, f"{key}_ins_{i + 1}", ins_seq))

            # Create a DataFrame from the variant information
    variant_df = pd.DataFrame(variant_info, columns=['target_name', 'variant_name', 'variant_seq'])

    # Merge with the original DataFrame on 'target_name'
    # Removing any existing columns in-common between these...
    intersect_cols = [col for col in variant_df.columns if col in df.columns if col!='target_name']
    if len(intersect_cols) > 0:
        df = df.drop(columns=intersect_cols)
    merged_df = df.merge(variant_df, on='target_name', how='inner')

    # Calculate and print the total run time
    end_time = time.time()
    total_sec = (end_time - start_time)
    total_min = total_sec / 60
    print(f"Done. Run time: {total_min:.2f} min ({total_sec:.2f} s)")

    return merged_df


def hamming_vectorized(seq1, seq2):
    """Calculate the Hamming distance between two sequences."""
    return np.sum(np.array(list(seq1)) != np.array(list(seq2)))


def calc_hamming(df, guide_to_seq_dict):
    print('Calculating Hamming distances...')
    start_time = time.time()

    # Lists to store Hamming distance results
    hamming_list = []

    # # Iterate over each combination of variant and guide sequences
    # for (var_index, var_row), (guide_index, guide_row) in itertools.product(df.iterrows(), guides_df.iterrows()):
    #     variant_name = var_row['variant_name']
    #     variant_seq = var_row['variant_seq']
    #     guide_name = guide_row['homol_guide_name']
    #     guide_seq = guide_row['homol_guide_seq']
    #     # Calculate Hamming distance using the vectorized function
    #     try:
    #         hamming = hamming_vectorized(variant_seq, guide_seq)
    #         hamming_list.append((variant_name, guide_name, guide_seq, hamming))
    #     except ValueError as e:
    #         print(f"Skipping comparison due to error: {e}")

    # Input guide-to-sequence dictionary instead of df
    # Iterate over each variant sequence and guide sequence
    for var_index, var_row in df.iterrows():
        variant_name = var_row['variant_name']
        variant_seq = var_row['variant_seq']
        for guide_name, sequence in guide_to_seq_dict.items():
            # Calculate Hamming distance using the vectorized function
            guide_seq = sequence[:20]
            pam_seq = sequence[-3:]
            try:
                hamming = hamming_vectorized(variant_seq, guide_seq)
                hamming_list.append((variant_name, guide_name, guide_seq, pam_seq, hamming))
            except ValueError as e:
                print(f"Skipping comparison due to error: {e}")

    # Create a DataFrame from the Hamming distance results
    ham_df = pd.DataFrame(hamming_list, columns=[
        'variant_name', 'homol_guide_name', 'homol_guide_seq', 'homol_guide_pam', 'hamming_distance'
    ])

    # Merge the ham_df with the original indel_df on 'variant_name'
    intersect_cols = [col for col in ham_df.columns if col in df.columns if col!='variant_name']
    if len(intersect_cols) > 0:
        df = df.drop(columns=intersect_cols)
    all_df = df.merge(ham_df, on='variant_name', how='left').sort_values(['chr', 'pos']).reset_index(drop=True)

    # select lowest ham distance candidate per edit site
    # group by 'edit_site', index by first occurence of min ham distance
    # closest ham distance per variant
    top_df = all_df.loc[all_df.groupby('name')['hamming_distance'].idxmin()]
    top_df = top_df.sort_values('hamming_distance').reset_index(drop=True)

    # format top_df
    top_df = top_df.drop(columns=['query_seq', 'n_close', 'close_offsets', 'abs_offsets', 'close_starts', 'close_ends',
                                  'close_targets', 'close_strands'])
    # Think just re-ordering the columns.
    # top_df = top_df[[
    #     'edit_site', 'chrom', 'position', 'start_window', 'end_window',
    #     'intersecting_genes', 'copy_number', 'n_cells_edited', 'stranded_edit_dist',
    #     'homol_guide_name', 'homol_guide_seq', 'homol_guide_pam', 'hamming_distance',
    #     'target_start', 'target_end', 'target_seq', 'target_pam', 'strand', 'abs_offset', 'variant_name', 'variant_seq'
    # ]]

    # format all_df
    all_df = all_df.drop(columns=['close_offsets', 'abs_offsets', 'close_starts', 'close_ends', 'close_targets',
                                  'close_strands'])

    # Calculate and print the total run time
    end_time = time.time()
    total_sec = (end_time - start_time)
    total_min = total_sec / 60
    print(f"Done. Run time: {total_min:.2f} min ({total_sec:.2f} s)")

    return top_df, all_df

def remove_consec_gaps(seq):
    aln_target_consecs = re.findall(r"((.)\2{1,})", seq)
    aln_target_consecs = np.array([consec[0] for consec in aln_target_consecs if '-' in consec[0]])

    # Need to order these to do a proper replacement!
    consec_sizes = np.array([len(consec) for consec in aln_target_consecs])
    consec_order = np.argsort( -consec_sizes )
    aln_target_consecs_ordered = aln_target_consecs[consec_order]

    seqA_no_consecs = seq
    for consec in aln_target_consecs_ordered:
        seqA_no_consecs = seqA_no_consecs.replace(consec, '-')

    return seqA_no_consecs

def get_indel_len_and_count(seq):
    aln_single_indel = remove_consec_gaps(seq)
    indel_count = aln_single_indel.count('-')
    indel_len = seq.count('-')
    return indel_count, indel_len

def matches_vectorized(seq1, seq2):
    """Calculate the Hamming distance between two sequences."""
    return np.sum(np.array(list(seq1)) == np.array(list(seq2)))

def alignment_based_hamming(df, guide_to_seq_dict):
    print('Calculating Global Alignment Hamming distances...')
    start_time = time.time()

    # Lists to store Hamming distance results
    hamming_list = []

    # # Iterate over each combination of variant and guide sequences
    # for (var_index, var_row), (guide_index, guide_row) in itertools.product(df.iterrows(), guides_df.iterrows()):
    #     variant_name = var_row['variant_name']
    #     variant_seq = var_row['variant_seq']
    #     guide_name = guide_row['homol_guide_name']
    #     guide_seq = guide_row['homol_guide_seq']
    #     # Calculate Hamming distance using the vectorized function
    #     try:
    #         hamming = hamming_vectorized(variant_seq, guide_seq)
    #         hamming_list.append((variant_name, guide_name, guide_seq, hamming))
    #     except ValueError as e:
    #         print(f"Skipping comparison due to error: {e}")

    # Input guide-to-sequence dictionary instead of df
    # Iterate over each variant sequence and guide sequence
    for target_index, target_row in df.iterrows():
        target_name = target_row['target_name']
        target_seq = target_row['target_seq'][:-3] # Target_seq includes the PAM sequence, so chop this off
        for guide_name, sequence in guide_to_seq_dict.items():
            # Calculate Hamming distance using the vectorized function
            guide_seq = sequence[:20]
            pam_seq = sequence[-3:]

            # Aligning the guide sequence with the candidate cut site, to see if it makes sense.
            aln = pairwise2.align.localms(target_seq, guide_seq,
                                          1,  # score for match
                                          -1,  # mismatch penalty
                                          -.8, #-.5,  # gap-open penalty
                                          -.5,  # gap-extension penalty
                                          one_alignment_only=True)[0]

            try:
                hamming_aln = hamming_vectorized(aln.seqA, aln.seqB)

                # Now calculating hamming where indel mismatch has penalty of 1
                target_indel_count, target_indel_len = get_indel_len_and_count( aln.seqA )
                guide_indel_count, guide_indel_len = get_indel_len_and_count( aln.seqB )
                # Correction to the above hamming distance, whereby the mismatches due to gaps at indels are only
                # counting as 1 mismatch, as opposed to multiple mismatches
                indel_correction = (target_indel_len-target_indel_count) + (guide_indel_len - guide_indel_count)

                hamming_aln_indel_corrected = hamming_aln - indel_correction

                match_count = matches_vectorized(aln.seqA, aln.seqB)

                hamming_list.append((target_name, guide_name, guide_seq, pam_seq, aln.seqA, aln.seqB, hamming_aln,
                                     hamming_aln_indel_corrected, match_count))

            except ValueError as e:
                print(f"Skipping comparison due to error: {e}")

    # Create a DataFrame from the Hamming distance results
    ham_df = pd.DataFrame(hamming_list, columns=[
        'target_name', 'homol_guide_name', 'homol_guide_seq', 'homol_guide_pam', 'target_aln', 'guide_aln',
        'hamming_aln_distance', 'hamming_aln_indel_corrected', 'match_aln_count',
    ])

    # Merge the ham_df with the original indel_df on 'variant_name'
    intersect_cols = [col for col in ham_df.columns if col in df.columns if col!='target_name']
    if len(intersect_cols) > 0:
        df = df.drop(columns=intersect_cols)
    all_df = df.merge(ham_df, on='target_name', how='left').sort_values(['chr', 'pos']).reset_index(drop=True)

    # select lowest ham distance candidate per edit site
    # group by 'edit_site', index by first occurence of min ham distance
    # closest ham distance per variant
    top_df = all_df.loc[all_df.groupby('name')['match_aln_count'].idxmax()]
    top_df = top_df.sort_values('match_aln_count', ascending=False).reset_index(drop=True)

    # format top_df
    top_df = top_df.drop(columns=['query_seq', 'n_close', 'close_offsets', 'abs_offsets', 'close_starts', 'close_ends',
                                  'close_targets', 'close_strands',])
    # Think just re-ordering the columns.
    # top_df = top_df[[
    #     'edit_site', 'chrom', 'position', 'start_window', 'end_window',
    #     'intersecting_genes', 'copy_number', 'n_cells_edited', 'stranded_edit_dist',
    #     'homol_guide_name', 'homol_guide_seq', 'homol_guide_pam', 'hamming_distance',
    #     'target_start', 'target_end', 'target_seq', 'target_pam', 'strand', 'abs_offset', 'variant_name', 'variant_seq'
    # ]]

    # format all_df
    all_df = all_df.drop(columns=['close_offsets', 'abs_offsets', 'close_starts', 'close_ends', 'close_targets',
                                  'close_strands'])

    # Calculate and print the total run time
    end_time = time.time()
    total_sec = (end_time - start_time)
    total_min = total_sec / 60
    print(f"Done. Run time: {total_min:.2f} min ({total_sec:.2f} s)")

    return top_df, all_df


# helper function that automatically generates an edit_site_info_closest.txt post_hoc output
# edit_site_info.txt expanded with closest aligned sequence...
def impute_edit_to_homol_guide(edit_site_info_df, ref_fasta, guide_to_seq_dict,
                               n_targets=5, query_range=100, seed_size=5):

    targets_df = fetch_close_targets(edit_site_info_df, ref_fasta, n_targets, query_range)
    targets_long_df = expand_targets(targets_df)
    var_df = make_1bp_indels(targets_long_df, seed_size)
    #ham_df, all_ham_df = calc_hamming(var_df, guide_to_seq_dict)
    # For this added version, I will also try an alignment-based method for trying to match the guide-sequence and the
    # edit site
    # We will just use the targets, no 1bp variants, since global align will do this.
    aln_ham_df, all_aln_ham_df = alignment_based_hamming(targets_long_df, guide_to_seq_dict)

    #### Let's add to the aln_ham_df and all_aln_ham_df the hamming information allowing for a 1bp mismatch..
    # Just for sanity checking....
    # intersect_cols = [col for col in ham_df.columns if col in aln_ham_df.columns if col!='target_name']
    # if len(intersect_cols) > 0:
    #     aln_ham_df_ = aln_ham_df.drop(columns=intersect_cols)
    # aln_ham_df_ = aln_ham_df_.merge(ham_df, on='target_name', how='left')#.sort_values(['chr', 'pos']).reset_index(drop=True)

    #return ham_df, all_ham_df
    return aln_ham_df, all_aln_ham_df

def impute_edit_guides_homology(data, ref_fasta, guide_seq_dict, n_targets=3, query_range=50, seed_size=5):

    edit_site_info_df = data.uns['edit_site_info'].copy()

    ham_df, all_ham_df = impute_edit_to_homol_guide(edit_site_info_df, ref_fasta, guide_seq_dict,
                                                    n_targets=n_targets, query_range=query_range, seed_size=seed_size)

    data.uns['edit_site_info_guide_homology'] = ham_df
    data.uns['edit_site_info_all_guide_homology'] = all_ham_df
    print(f"Added data.uns['edit_site_info_guide_homology'], with extra columns with homology-based guide inference of each edit site.")
    print(f"Added data.uns['edit_site_info_all_guide_info'], with ALL information about homology-based guide "
          f"inference of each edit site."
          )

# Causal guide imputation by maximum on-target coincidence

# Function to map edits to causal guides
def impute_edit_guides_coinc(data, guide_to_seq_dict, paired_guides, guide_to_site_dict,
                             max_ontarget_edits=1,
                             #hamming_conf_cutoff = 4, # cutoff beyond which may not consider a very confident guide call.
                             match_aln_cutoff = 15,
                             coinc_frac_cutoff = 0.7, # cutoff below which may consider the coincidence call not very confident.
):
    """
    Imput guide for each edit site based maximum coincident on-target count.
    For edit sites without any coincident on-target counts, guide is imputed based on homology.

    Parameters:
    combined_outputs_dir (str): Path to sheriff combined outputs (chromosome-merged).
    guide_to_site_dict (dict): Dictionary of edit site names (keys) and guide RNA names (values).
    paired_guides (list): List of tuples of guide names that were multiplexed, to remove ambiguous multiplex-edited cells from calculation.
    output_dir (str): Path to output directory for.

    Returns:
    imp_all (dict): Dictionary of dfs with keys 'umi', 'allele', and 'cell' to mapping and coincidence counts (scores) of all edit sites to all on-target guides.
    imp (df): Dictionary of subset of imp_all dfs of edit-to-guide imputation by maximum coincidence count, without scaling.
    imp2 (df): Dictionary of subset of imp_all dfs of edit-to-guide imputation by maximum coincidence of scaled counts, normalized to each on-target site's count coverage.
    edit_to_guide_all.pq (file): Parquet file of flattened imp_all dict, with key as first column
    edit_to_guide_unscaled.pq (file): Parquet file of flattened imp dict, with key as first column
    edit_to_guide_scaled.pq (file): Parquet file of flattened imp2 dict, with key as first column
    edit_site_imputed.txt (file): Expanded edit_site_info.txt with imputed guide information (coincidence and homology)
    """

    if 't7_umis' not in data.obsm or 'dosages' not in data.obsm:
        raise Exception("Did not find superb-seq specific data attached to AnnData, "
                        "so does not appear to be from a superb-seq experiment.")

    if 'edit_site_info_guide_homology' not in data.uns:
        raise Exception("Missing homology-based edit-to-guide information, please run impute_edit_guides_homology() first.")

    cell_umi = data.obsm['t7_umis'].copy()
    cell_allele = data.obsm['dosages'].copy()

    # counting the number of on-target edits the cells receive, ideally we just want cells that have only a limited
    # number of multiple guides to make this calculation, otherwise the confouding of multiple guides per cell will be
    # a problem...
    # on_target_edits = list(guide_to_site_dict.values())
    # cell_ontarget_counts = cell_umi.loc[:, on_target_edits].sum(axis=1)
    # keep_cells = cell_ontarget_counts.values <= max_ontarget_edits
    #
    # cell_umi = cell_umi.loc[keep_cells, :] #remove_dki_cells(cell_umi, edit_pairs)
    # cell_allele = cell_allele.loc[keep_cells, :]  #remove_dki_cells(cell_allele, edit_pairs)
    #
    # print(f"NOTE: using {sum(keep_cells)} to calculated edit coincidence, since have at most "
    #       f"{max_ontarget_edits} guide edits.")

    # remove "double knock-in" cells with multiplexed on-target counts
    # edit_pairs = []
    # for pair in paired_guides:
    #     pair_new = (guide_to_site_dict[pair[0]], guide_to_site_dict[pair[1]])
    #     edit_pairs.append(pair_new)
    #
    # def remove_dki_cells(count_df, edit_pairs):
    #     for col1, col2 in edit_pairs:
    #         condition = (count_df[col1] == 0) | (count_df[col2] == 0)
    #         count_df = count_df[condition]
    #     return count_df
    #
    # cell_umi = remove_dki_cells(cell_umi, edit_pairs)
    # cell_allele = remove_dki_cells(cell_allele, edit_pairs)

    cell_bin = (cell_allele > 0).astype(int)

    level_matrices = {'umi': cell_umi, 'allele': cell_allele, 'cell': cell_bin}

    # # make coincidence matrices
    # think the above, based on un-normalised dot product, will be heavily susceptible to the sequencing depth of a cell,
    # indeed, the allelic calls and the umis will be heavily correlated with the total cell UMIs, so to remove this
    # factor of scaling, will normalise the dot-products so is the angle, which should be scale-invariant.
    from scipy.spatial.distance import pdist, squareform

    coinc_imp = {}
    for level, matrix in level_matrices.items():
        cosine_distances = pdist(level_matrices[level].T, metric='cosine')
        cosine_similarity_matrix = 1 - squareform(cosine_distances)
        coinc_imp[level] = pd.DataFrame(cosine_similarity_matrix, index=level_matrices[level].columns.values,
                                        columns=level_matrices[level].columns.values)
        # Some issue in this version of the cosine similarity, cases where diagonal was not 1 for some reason.
        # dot_product = level_matrices[level].T.dot( level_matrices[level] )
        # norms = np.linalg.norm(level_matrices[level], axis=0)
        # norms[norms == 0] = 1e-10
        # cosine_similarity = dot_product / np.outer(norms, norms)
        # coinc_imp[level] = cosine_similarity

    # guide imputation based on most coincident of on-target events
    imp_all = {}
    imp = {}
    imp2 = {}
    edit_guide = {}
    edit_guide2 = {}
    max_coinc_set = {}
    max_coinc_set2 = {}

    # levels = ['umi', 'allele', 'cell', 'binary']
    levels = ['umi', 'allele', 'cell']
    on_df = pd.DataFrame(list(guide_to_site_dict.items()), columns=['guide_name', 'on_target'])

    # # read in balance.pq to get edit sites above balance threshold (0.8), for set filtering...
    # post_hoc = '/iblm/netapp/data4/mlorenzini/superb_seq/data/sheriff/t7_processing_v10/post_hoc/'
    # bal_pq = post_hoc+'balance.pq'
    # bal_df = pd.read_parquet(bal_pq)
    # sites_above_thresh = bal_df[bal_df['above_balance_thresh'] == True]['edit_site'].tolist()

    for l in levels:
        # calculate on-target coincidence for each edit site
        imp_all[l] = coinc_imp[l].copy().reset_index().rename(columns={'index': 'site1'})
        imp_all[l] = imp_all[l].melt(id_vars='site1', var_name='site2', value_name='coinc_count').query(
            'coinc_count != 0')
        imp_all[l] = imp_all[l][imp_all[l]['site2'].isin(on_df['on_target'])]
        imp_all[l] = imp_all[l].sort_values('site1').reset_index(drop=True)
        imp_all[l] = imp_all[l].rename(columns={'site1': 'edit_site', 'site2': 'on_target'})
        imp_all[l] = imp_all[l].merge(on_df, how='left', on='on_target')
        imp_all[l]['on_sum'] = imp_all[l].groupby('on_target')['coinc_count'].transform('sum')
        imp_all[l]['on_scale'] = imp_all[l]['on_sum'] / imp_all[l]['on_sum'].min()
        imp_all[l]['count_scaled'] = imp_all[l]['coinc_count'] / imp_all[l]['on_scale']
        imp_all[l]['sum'] = imp_all[l].groupby('edit_site')['coinc_count'].transform('sum')
        imp_all[l]['sum_scaled'] = imp_all[l].groupby('edit_site')['count_scaled'].transform('sum')
        imp_all[l]['coinc_fraction'] = imp_all[l]['coinc_count'] / imp_all[l]['sum']
        imp_all[l]['coinc_frac_scaled'] = imp_all[l]['count_scaled'] / imp_all[l]['sum_scaled']
        imp_all[l] = imp_all[l].rename(columns={'guide_name': 'coinc_guide_name'})

        # impute guides by max on-target coincidence count
        imp[l] = imp_all[l].copy().sort_values(['edit_site', 'coinc_count'], ascending=[True, False]).groupby(
            'edit_site').first().reset_index()
        imp[l] = imp[l].sort_values(['coinc_guide_name', 'coinc_count'], ascending=[True, False]).reset_index(drop=True)
        imp[l] = imp[l].sort_values('coinc_count', ascending=False).reset_index(drop=True)
        imp2[l] = imp_all[l].copy().sort_values(['edit_site', 'count_scaled'], ascending=[True, False]).groupby(
            'edit_site').first().reset_index()
        imp2[l] = imp2[l].sort_values(['coinc_guide_name', 'count_scaled'], ascending=[True, False]).reset_index(
            drop=True)
        imp2[l] = imp2[l].sort_values('count_scaled', ascending=False).reset_index(drop=True)

    # flatten dictionaries to dataframes
    for l in levels:
        imp_all[l].insert(0, 'level', l)
        imp[l].insert(0, 'level', l)
        imp2[l].insert(0, 'level', l)
    imp_all_df = pd.concat(imp_all.values(), ignore_index=True)
    imp_df = pd.concat(imp.values(), ignore_index=True)
    imp2_df = pd.concat(imp2.values(), ignore_index=True)

    # Add UNSCALED (max coinc_coint imputation), UMI-based guide imputation to edit_site_info_homology.txt
    # add columns on_target, guide_name, GUIDE SEQUENCE, coinc_count, coinc_fraction,
    # Make edit-to-guide dictionary from this, for cell-to-guide imputation and count correction...
    imp_merge = imp['umi'].copy()
    imp_merge['coinc_guide_seq'] = imp_merge['coinc_guide_name'].map(guide_to_seq_dict)
    imp_merge = imp_merge[['edit_site', 'coinc_guide_name', 'coinc_guide_seq', 'coinc_count', 'coinc_fraction']]

    imp_merge = imp_merge.rename({'edit_site': 'name'}, axis='columns')

    ham_df = data.uns['edit_site_info_guide_homology'].copy()
    edit_imp = ham_df.merge(imp_merge, how='left', on='name').reset_index(drop=True)

    ########## Now making a final call on which edit_to_guide, and seeing which are confident!!!!
    homol_df_all = data.uns['edit_site_info_all_guide_homology']
    imp_all_df_umi = imp_all_df.loc[imp_all_df['level'].values=='umi',:]
    imp_all_df_umi.index = [f'{edit_site}---{on_target}' for edit_site, on_target
                            in zip(imp_all_df_umi['edit_site'], imp_all_df_umi['on_target'])]
    site_to_guide_dict = {edit_site: guide for guide, edit_site in guide_to_site_dict.items()}

    edit_imp_confident = [] # whether or not we think is confiden imputation..
    edit_scores = []

    edit_imp_final = edit_imp.copy()
    coinc_col_indices = [i for i, col in enumerate(edit_imp_final.columns) if col.startswith('coinc')]
    for rowi, row in edit_imp.iterrows():
        edit_site = row['name']
        homol_guide, coinc_guide = row['homol_guide_name'], row['coinc_guide_name']
        homol_matches, coinc_frac = row['match_aln_count'], row['coinc_fraction']
        # OLD version where was using hamming distance based on a 1bp variant of the target genome sequence.
        #homol_hamming, coinc_frac = row['hamming_distance'], row['coinc_fraction']
        #homol_guide_len, coinc_guide_len = len(row['homol_guide_seq']), len(row['coinc_guide_seq'])

        # Handling case where homology and guide edit-site coincidence in cells agrees
        if homol_guide == coinc_guide:
            #homol_score = (1 - (homol_hamming / hamming_conf_cutoff)) # old method
            homol_score = (homol_matches / match_aln_cutoff) ** 10 - 1
            coinc_score = coinc_frac / coinc_frac_cutoff

            guide_score = homol_score + coinc_score
            edit_scores.append( guide_score )

        elif str(coinc_guide) == 'nan' or str(coinc_frac)=='nan': # Is no guide call based on coincidence, never co-occurs within on-target edit.
            # Best call is that the causal guide based on homology.
            edit_imp_final.iloc[rowi, coinc_col_indices] = [homol_guide, row['homol_guide_seq'], 0, 0]

            #homol_score = (1 - (homol_hamming / hamming_conf_cutoff)) # OLD method
            # Calibrating the score, to try and really favour homology if very high, and be less favourable as it gets lower
            #import matplotlib.pyplot as plt
            #scores = [(x / match_aln_cutoff) ** 10 - 1 for x in range(21)]
            #plt.plot(scores)
            #plt.show()
            homol_score = (homol_matches / match_aln_cutoff) ** 10 - 1
            edit_scores.append( homol_score ) # since the coinc_score will be 0, can just add the homol_score as the guide_score

        else:
            # Handling case where they disagree:
            # Where they disagree, will handle by
            # 1) getting the coincidence scores for the best guide by homology.
            # 2) getting the homology scores for the best guide by coincidence.
            # 3) a joint score for each candidate guide, to decide which is the causal guide for the edit site
            # 1) getting the coincidence scores for the best guide by homology.
            homol_guide_edit_site = guide_to_site_dict[homol_guide]
            entry_name = f'{edit_site}---{homol_guide_edit_site}'
            if entry_name in imp_all_df_umi.index: # If is present, had a non-zero coinc_count
                homol_coinc = imp_all_df_umi.loc[entry_name,:]
                homol_coinc_frac = homol_coinc['coinc_fraction']

            else: # had a 0 coinc_count, with the inferred edit_site, so all scores are 0
                homol_coinc_frac = 0

            # 2) getting the homology scores for the best guide by coincidence.
            top_df = homol_df_all.loc[homol_df_all['name'].values==edit_site, :]
            top_df = top_df.loc[top_df['homol_guide_name'].values==coinc_guide, :]
            # coinc_homol = top_df.loc[top_df['hamming_distance'].idxmin()] # old method
            # coinc_hamming = coinc_homol['hamming_distance']
            coinc_homol = top_df.loc[top_df['match_aln_count'].idxmax()]
            coinc_homol_matches = coinc_homol['match_aln_count']

            # 3) a joint score for each candidate guide, to decide which is the causal guide for the edit site
            #homol_hamming, homol_coinc_frac, coinc_hamming, coinc_frac

            ### First, let's use our confidence cutoffs, to see if that breaks the disagreement.
            # homol_guide_homol_score = (1 - (homol_hamming / homol_guide_len)) / ((1 - (hamming_conf_cutoff / homol_guide_len)))
            # coinc_guide_homol_score = (1 - (coinc_hamming / coinc_guide_len)) / ((1 - (hamming_conf_cutoff / homol_guide_len)))
            #
            # homol_guide_homol_score = (1 - (homol_hamming / hamming_conf_cutoff)) # old method
            # coinc_guide_homol_score = (1 - (coinc_hamming / hamming_conf_cutoff))
            homol_matches, homol_coinc_frac, coinc_homol_matches, coinc_frac

            homol_guide_homol_score = (homol_matches / match_aln_cutoff) ** 10 - 1
            coinc_guide_homol_score = (coinc_homol_matches / match_aln_cutoff) ** 10 - 1

            homol_guide_coinc_score = homol_coinc_frac / coinc_frac_cutoff
            coinc_guide_coinc_score = coinc_frac / coinc_frac_cutoff

            # homology guide has high hamming and no coincidence, go for highest coincidence
            # if homol_guide_homol_score < 0 and homol_guide_coinc_score == 0 and coinc_frac > 0:
            #     homol_guide_score = 0
            #     coinc_guide_score = 1
            # else: # Need to balance the score to select the causal guide.
            homol_guide_score = homol_guide_homol_score + homol_guide_coinc_score
            coinc_guide_score = coinc_guide_homol_score + coinc_guide_coinc_score

            # if rowi == 33:
            #     print("here")

            if homol_guide_score > coinc_guide_score: # Homology guide scored better, so make this our causal guide call.
                edit_scores.append( homol_guide_score )

                #### Update the final information to reflect only the homology-based guide.
                if homol_coinc_frac == 0: # Is 0, so don't have values for this, so just replace with 0..
                    edit_imp_final.iloc[rowi, coinc_col_indices] = [homol_guide, row['homol_guide_seq'], 0, 0]

                else:
                    intersect_cols = list(set(list(homol_coinc.index)).intersection(set(list(edit_imp_final.columns))))
                    col_indices = [i for i, col in enumerate(edit_imp_final.columns) if col in intersect_cols]
                    intersect_cols = [edit_imp_final.columns.values[i] for i in col_indices]
                    edit_imp_final.iloc[rowi, col_indices] = homol_coinc.loc[intersect_cols].values

            else: # coinc guide scored better!
                edit_scores.append( coinc_guide_score )

                intersect_cols = list(set(list(coinc_homol.index)).intersection(set(list(edit_imp_final.columns))))
                col_indices = [i for i, col in enumerate(edit_imp_final.columns) if col in intersect_cols]
                intersect_cols = [edit_imp_final.columns.values[i] for i in col_indices]
                edit_imp_final.iloc[rowi, col_indices] = coinc_homol.loc[intersect_cols].values

        row_new = edit_imp_final.iloc[rowi, :]
        #homol_hamming_new, coinc_frac_new = row_new['hamming_distance'], row_new['coinc_fraction'] # old method
        homol_matches_new, coinc_frac_new = row_new['match_aln_count'], row_new['coinc_fraction']

        # Reflects how confident we are in the causal guide call for a given edit site.
        if homol_matches_new >= match_aln_cutoff:
            edit_imp_confident.append( True )
        else:
            edit_imp_confident.append( False )

    edit_imp_final['guide_score'] = edit_scores
    edit_imp_final['confident_guide_call'] = edit_imp_confident

    data.uns['edit_site_info_guide_homology_and_coinc'] = edit_imp
    data.uns['edit_site_info_guide_homology_and_coinc_final'] = edit_imp_final
    print(f"Added data.uns['edit_site_info_guide_homology_and_coinc'], with extra columns indicating best guide by edit-site coincidence with guide on-target.")
    print(f"Added data.uns['edit_site_info_guide_homology_and_coinc_final'], final calls on causal guide, with confident calls indicated.")
```
