## Additional Files for "Joint single-cell capture of Cas9 edits and transcriptomes reveals on- and off-target effects on gene expression": superb_helpers.py

```
"""Helper functions for performing the superb-seq analysis.
"""

import numpy as np
import pandas as pd
from collections import namedtuple

import scipy.stats as stats
import statsmodels.formula.api as smf

import statsmodels.stats.multitest as smm

import scipy.stats
from scipy.stats.contingency import odds_ratio

from collections import defaultdict
import beautifulcells.preprocessing.load_data.simple_pickle as spl

from sklearn.neighbors import NearestNeighbors

import gseapy as gp

import matplotlib.pyplot as plt


def construct_allelic_pairings(data_edited, edited_gene, n_reps=1,
                               umap_key='X_umap_log1p', total_count_key='log1p_total_counts',
                               condition_key=None, # Specifies if the positive cells must belong to a certain condition
                               dosage_key='dosages_genes',
                               edit_name=None, #If want to make a call for a particular edit.
                               min_cells_with_allele=50,
                               max_total_umi_dist=None, # Maximum number of total UMIs that can be different between the cells
                               method='mode_allele_anchor',
                               ):
    """ Constructs a dataframe of pairings representing groups of 0, 1, 2, etc allelic edited cells, which aside from
        having different allelic edit counts have the same library size and cell state.

        Assumes AnnData already subsetted to just edited cells and control cells.
    """
    supported_methods = ['mode_allele_anchor', 'pair_each_allele']
    if method == 'mode_allele_anchor':
        return construct_allelic_pairings_mode_allele_group_anchor(data_edited, edited_gene, n_reps=n_reps,
                               umap_key=umap_key, total_count_key=total_count_key,
                               condition_key=condition_key, # Specifies if the positive cells must belong to a certain condition
                               dosage_key=dosage_key,
                               edit_name=edit_name, #If want to make a call for a particular edit.
                               min_cells_with_allele=min_cells_with_allele,
                               max_total_umi_dist=max_total_umi_dist, # Maximum number of total UMIs that can be different between the cells
                            )
    elif method == 'pair_each_allele':
        return construct_allelic_pairings_for_each_allele(data_edited, edited_gene, n_reps=n_reps,
                                                                   umap_key=umap_key, total_count_key=total_count_key,
                                                                   condition_key=condition_key,
                                                                   # Specifies if the positive cells must belong to a certain condition
                                                                   dosage_key=dosage_key,
                                                                   edit_name=edit_name,
                                                                   # If want to make a call for a particular edit.
                                                                   min_cells_with_allele=min_cells_with_allele,
                                                                   max_total_umi_dist=max_total_umi_dist,
                                                                   # Maximum number of total UMIs that can be different between the cells
                                                                   )
    else:
        raise Exception(f"Inputted method not supported. Selected one of: {supported_methods}")

def construct_allelic_pairings_for_each_allele(data_edited, edited_gene,
                                    n_reps=1, umap_key='X_umap_log1p', total_count_key='log1p_total_counts',
                                    condition_key=None, # Specifies if the positive cells must belong to a certain condition
                                    dosage_key='dosages_genes', edit_name=None, # If want to make a call for a particular edit.
                                    min_cells_with_allele=50,
                                    max_total_umi_dist=None, # Maximum number of total UMIs that can be different between the cells
                                            ):
            """ Allelic groups are constructed by selecting a no-edit cell for each edited cell with a similar
            total UMI and cell state to the edited cell, so that each group forms a pair of cells with unique
            cell state and UMI, BUT we still modelling allelic counts!
            """

            # subject_cell_index corr_cell_index allelic_edits subject_cell_index_counts subject_cell_index_UMAP corr_cell_index_counts corr_cell_index_UMAP gene_expr1 .. gene_exprn
            Entry = namedtuple('Entry',
                               ['group', 'subject_cell_index', 'corr_cell_index',
                                'subject_cell_bc', 'corr_cell_barcode',
                                'subject_allelic_edits', 'allelic_edits',
                                'subject_counts', 'subject_UMAP1', 'subject_UMAP2',
                                'corr_counts', 'corr_UMAP1', 'corr_UMAP2'])

            if type(edit_name) == type(None):
                edit_name = edited_gene

            # We only want cells with evidence of being edited.
            condition = data_edited.obsm[dosage_key].loc[:, edit_name].values > 0
            if type(condition_key) != type(None):
                condition = np.logical_and(condition, data_edited.obs['sample'].values == condition_key)
            # And true negatives.
            unedited_bool = data_edited.obs['sample'].values == 'no_ki'
            condition_bool = np.logical_or(condition, unedited_bool)

            # AnnData of the cells with the particular edit.
            adata = data_edited[condition_bool, :]  #
            cell_barcodes = adata.obs_names.values

            # Classify cells into edited and unedited
            on_target_alleles = adata.obsm[dosage_key].loc[:, edit_name].values

            no_edit_bool = on_target_alleles == 0
            no_edit_indices = np.where(no_edit_bool)[0]
            edit_indices = np.where(~no_edit_bool)[0]

            # Removing edit_indices corresponding to very rare alleles.
            allele_set = np.unique(on_target_alleles)
            allele_indices = [np.where(on_target_alleles==allele_)[0] for allele_ in allele_set]
            allele_counts = np.array([len(allele_indices_) for allele_indices_ in allele_indices])
            remove_allele_indices = np.where(allele_counts < min_cells_with_allele)[0]
            for remove_allele_index in remove_allele_indices: # If no alleles to remove this won't iterate.
                edit_indices = np.array([index for index in edit_indices # Filters out edit cell indices with this allele
                                         if index not in allele_indices[remove_allele_index]])

            # Extracting features we want to select the cells to have equivalent of:
            umap_ = adata.obsm[umap_key]
            sizes_ = adata.obs[total_count_key].values
            cell_features = np.hstack((umap_, sizes_.reshape(-1,1)))

            # Z-scaling the features:
            means = cell_features.mean(axis=0)
            stds = cell_features.std(axis=0)
            cell_features = np.subtract(cell_features, means)
            cell_features = np.divide(cell_features, stds)

            # Selecting example no-edit cell for each edited cell!
            control_cell_features = cell_features[no_edit_indices,:]
            edit_cell_features = cell_features[edit_indices,:]

            # Find nearest neighbors for matching based on sequencing depth and cell state
            nn = NearestNeighbors(n_neighbors=n_reps, metric='manhattan')
            nn.fit(control_cell_features)

            distances, indices = nn.kneighbors(edit_cell_features) # Getting closest no-edit cells!

            # Storing the pairing information!
            data_entries = []
            # Randomising the order!
            for i, row in enumerate(indices):

                edit_index = edit_indices[i]

                selected = False
                for ci, control_index_ in enumerate(row):
                    control_index = no_edit_indices[control_index_]  # Was relative to subset
                    group_str = f"{edit_index}-{ci}"

                    # Will only add as an entry IF the library sizes are comparable...
                    if type(max_total_umi_dist) != type(None):
                        subject_counts = sizes_[edit_index]
                        corr_counts = sizes_[control_index]
                        diff_counts = subject_counts - corr_counts
                        if abs(diff_counts) > max_total_umi_dist:  # TOO different to make a comparison.
                            continue

                    # Another entry
                    data_entries.append(
                        Entry(group=group_str, subject_cell_index=edit_index, corr_cell_index=control_index,
                              subject_cell_bc=cell_barcodes[edit_index],
                              corr_cell_barcode=cell_barcodes[control_index],
                              subject_allelic_edits=on_target_alleles[edit_index],
                              allelic_edits=on_target_alleles[control_index], subject_counts=sizes_[edit_index],
                              subject_UMAP1=umap_[edit_index, 0], subject_UMAP2=umap_[edit_index, 1],
                              corr_counts=sizes_[control_index],
                              corr_UMAP1=umap_[control_index, 0], corr_UMAP2=umap_[control_index, 1]
                              )
                        )

                    selected = True
                    break # Don't look through any more control cells as a potential candidate!

                # This means the edited cell data entry!
                if selected: # We could find a control cell that has similar technical measures as this cell, so add!
                    base_entry = Entry(group=group_str, subject_cell_index=edit_index, corr_cell_index=edit_index,
                                       subject_cell_bc=cell_barcodes[edit_index],
                                       corr_cell_barcode=cell_barcodes[edit_index],
                                       subject_allelic_edits=on_target_alleles[edit_index],
                                       allelic_edits=on_target_alleles[edit_index], subject_counts=sizes_[edit_index],
                                       subject_UMAP1=umap_[edit_index, 0], subject_UMAP2=umap_[edit_index, 1],
                                       corr_counts=sizes_[edit_index],
                                       corr_UMAP1=umap_[edit_index, 0], corr_UMAP2=umap_[edit_index, 1]
                                       )
                    data_entries.append(base_entry)

            data_entries_df = pd.DataFrame(data_entries).sort_values(by='group')
            print("Done pairing cells")

            return data_entries_df

def construct_allelic_pairings_mode_allele_group_anchor(data_edited, edited_gene, n_reps=1,
                               umap_key='X_umap_log1p', total_count_key='log1p_total_counts',
                               condition_key=None, # Specifies if the positive cells must belong to a certain condition
                               dosage_key='dosages_genes',
                               edit_name=None, #If want to make a call for a particular edit.
                               min_cells_with_allele=50,
                               max_total_umi_dist=None, # Maximum number of total UMIs that can be different between the cells
                               ):
    """ Allelic groups are constructed by selecting the most frequent allelic edit dosage as an anchor, then
        choosing no-edits cells and other allelic edits that have the same total UMIs and cell state!
    """

    # subject_cell_index corr_cell_index allelic_edits subject_cell_index_counts subject_cell_index_UMAP corr_cell_index_counts corr_cell_index_UMAP gene_expr1 .. gene_exprn
    Entry = namedtuple('Entry',
                       ['group', 'subject_cell_index', 'corr_cell_index',
                        'subject_cell_bc', 'corr_cell_barcode', 'allelic_edits',
                        'subject_counts', 'subject_UMAP1', 'subject_UMAP2',
                        'corr_counts', 'corr_UMAP1', 'corr_UMAP2'])

    if type(edit_name) == type(None):
        edit_name = edited_gene

    # We only want cells with evidence of being edited.
    condition = data_edited.obsm[ dosage_key ].loc[:, edit_name].values > 0
    if type(condition_key)!=type(None):
        condition = np.logical_and(condition, data_edited.obs['sample'].values == condition_key)
    # And true negatives.
    unedited_bool = data_edited.obs['sample'].values == 'no_ki'
    condition_bool = np.logical_or(condition, unedited_bool)

    # AnnData of the cells with the particular edit.
    adata = data_edited[condition_bool, :]  #
    cell_barcodes = adata.obs_names.values

    # Classify cells into edited and unedited
    on_target_alleles = adata.obsm[ dosage_key ].loc[:, edit_name].values

    mode_allele = stats.mode(on_target_alleles[on_target_alleles > 0])[0]
    all_alleles = np.unique(on_target_alleles)
    other_alleles = np.array([allele for allele in all_alleles if allele != mode_allele])

    mode_allele_indices = np.where(on_target_alleles == mode_allele)[0]

    umap_ = adata.obsm[umap_key]
    sizes_ = adata.obs[total_count_key].values

    mode_allele_umap = umap_[mode_allele_indices, :]
    mode_allele_sizes = sizes_[mode_allele_indices, np.newaxis]
    mode_allele_features = np.hstack((mode_allele_umap, mode_allele_sizes))
    # Standardising, so that scaling does not lend more weight to the umap coords or the total counts
    means = mode_allele_features.mean(axis=0)
    stds = mode_allele_features.std(axis=0)
    mode_allele_features = np.subtract(mode_allele_features, means)
    mode_allele_features = np.divide(mode_allele_features, stds)

    # Storing the pairing information!
    data_entries = []
    n_subjects = 0
    for allele in other_alleles:
        allele_indices = np.where(on_target_alleles == allele)[0]

        if len(allele_indices) < min_cells_with_allele:  # Too few cells edited to make a call
            continue

        allele_umap = umap_[allele_indices, :]
        allele_sizes = sizes_[allele_indices]
        allele_features = np.hstack((allele_umap, allele_sizes[:, np.newaxis]))
        allele_features = np.subtract(allele_features, means)
        allele_features = np.divide(allele_features, stds)

        # Find nearest neighbors for matching based on sequencing depth and cell state
        nn = NearestNeighbors(n_neighbors=n_reps, metric='manhattan')
        nn.fit(allele_features)

        distances, indices = nn.kneighbors(mode_allele_features)

        # Randomising the order!
        for i, row in enumerate(indices):
            np.random.shuffle(row)

            mode_index = mode_allele_indices[i]

            for ai, allele_index_ in enumerate(row):
                allele_index = allele_indices[allele_index_]  # Was relative to subset
                group_str = f"{mode_index}-{ai}"

                base_entry = Entry(group=group_str, subject_cell_index=mode_index, corr_cell_index=mode_index,
                                   subject_cell_bc=cell_barcodes[mode_index], corr_cell_barcode=cell_barcodes[mode_index],
                                   allelic_edits=mode_allele, subject_counts=sizes_[mode_index],
                                   subject_UMAP1=umap_[mode_index, 0], subject_UMAP2=umap_[mode_index, 1],
                                   corr_counts=sizes_[mode_index],
                                   corr_UMAP1=umap_[mode_index, 0], corr_UMAP2=umap_[mode_index, 1]
                                   )
                if base_entry not in data_entries:
                    data_entries.append(base_entry)

                # Will only add as an entry IF the library sizes are comparable...
                if type(max_total_umi_dist) != type(None):
                    subject_counts = sizes_[mode_index]
                    corr_counts = sizes_[allele_index]
                    diff_counts = subject_counts - corr_counts
                    if abs(diff_counts) > max_total_umi_dist: # TOO different to make a comparison.
                        continue

                # Another entry
                data_entries.append(Entry(group=group_str, subject_cell_index=mode_index, corr_cell_index=allele_index,
                                          subject_cell_bc=cell_barcodes[mode_index],
                                          corr_cell_barcode=cell_barcodes[allele_index],
                                          allelic_edits=allele, subject_counts=sizes_[mode_index],
                                          subject_UMAP1=umap_[mode_index, 0], subject_UMAP2=umap_[mode_index, 1],
                                          corr_counts=sizes_[allele_index],
                                          corr_UMAP1=umap_[allele_index, 0], corr_UMAP2=umap_[allele_index, 1]
                                          )
                                    )

    data_entries_df = pd.DataFrame(data_entries).sort_values(by='group')
    print("Done pairing cells")

    return data_entries_df

def test_edit_and_gene(data_edited, data_entries_df, gene,
                       fixed_effect_variables=['allelic_edits'], random_effect_variable='subject_cell_index'):
    """Tests association between an edit event and the gene expression
    """

    # Adding the gene to the data frame
    gene_expr = data_edited[data_entries_df['corr_cell_barcode'].values, gene].X.toarray()[:, 0]

    # Need to account for problem with lm fit
    gene = gene.replace('-', '_')

    data_entries_df[gene] = gene_expr

    # Fitting the linear mixed-effects model
    n_groups = len(np.unique(data_entries_df[random_effect_variable].values))
    if n_groups > 1:
        model = smf.mixedlm(f"{gene} ~ {'+'.join(fixed_effect_variables)}", data_entries_df,
                            groups=data_entries_df[random_effect_variable],
                            )

        result = model.fit()

        if len(fixed_effect_variables) == 1:  # Just fitting one fixed effect, easy to extract
            alleles_beta = float(result.summary().tables[1].values[1, 0])
            alleles_pval = result.pvalues.loc[fixed_effect_variables[0]]
            print("Overall reduction for 2 alleles:", np.e ** (alleles_beta * 2))

            output = alleles_beta, alleles_pval, result, model

        else:
            allele_betas = result.summary().tables[1].loc[fixed_effect_variables, 'Coef.'].values.astype(float)
            allele_pvals = result.pvalues.loc[fixed_effect_variables].values
            output = allele_betas, allele_pvals, result, model

    else: # No groups..
        model = smf.ols(f"{gene} ~ {'+'.join(fixed_effect_variables)}", data_entries_df,
                            )

        result = model.fit()

        allele_betas = result.params['allelic_edits']
        allele_pvals = result.pvalues['allelic_edits']
        output = allele_betas, allele_pvals, result, model

    print(result.summary())

    print("DONE")
    return output


def plot_results(result, alleles_beta, data_entries_df, gene, data,
                 color_violin='blue', save_fig = False, out_plots='./',
                 show_bootstrap_trend=True):
    """ Plots the results.
    """
    # Need to account for problem with lm fit
    gene = gene.replace('-', '_')

    # Residual analysis
    residuals = result.resid
    fitted = result.fittedvalues
    fitted_without_alleles = fitted - (alleles_beta * data_entries_df['allelic_edits'].values)
    actual_with_alleles_without_covars = data_entries_df[gene].values - fitted_without_alleles

    # Residual vs. Fitted Plot
    plt.scatter(fitted, residuals)
    plt.axhline(0, linestyle='--', color='red')
    plt.xlabel('Fitted values')
    plt.ylabel('Residuals')
    plt.title('Residual vs. Fitted Plot')
    plt.show()

    plt.scatter(data_entries_df[gene].values, fitted)
    plt.xlabel("Actual CHD4 expression")
    plt.ylabel("Predicted CHD4 expression")
    plt.show()

    data_entries_df[f'{gene}_corrected'] = actual_with_alleles_without_covars
    import seaborn as sb

    # Going to deduplicate BEFORE plotting, so more accurately reflects total cells in each category.
    unique_indices = np.unique(data_entries_df['corr_cell_index'].values)
    unique_indexes = [np.where(data_entries_df['corr_cell_index'].values == index)[0][0] for index in unique_indices]
    data_entries_df_derep = data_entries_df.iloc[unique_indexes, :]

    sb.violinplot(data_entries_df_derep, x='allelic_edits', y=f'{gene}_corrected',
                  color=color_violin)
    sb.stripplot(data_entries_df_derep, x='allelic_edits', y=f'{gene}_corrected',
                 alpha=.2, edgecolor='k', linewidth=1, color=color_violin)
    plt.plot(data_entries_df_derep['allelic_edits'].values, data_entries_df_derep['allelic_edits'].values * alleles_beta,
             c='red')
    if save_fig:
        plt.savefig(f'{out_plots}/{gene}-expr_allelic-edits_violin.png', dpi=300)
    plt.show()

    # Creating a boot-strapped mean representation:
    gene_counts = data.obs.loc[data_entries_df['corr_cell_barcode'].values, 'gene_count'].values
    log1p_total_counts = data.obs.loc[data_entries_df['corr_cell_barcode'].values, 'log1p_total_counts'].values

    data_entries_df[ 'log1p_total_counts' ] = log1p_total_counts
    data_entries_df['gene_counts'] = gene_counts

    ### Total counts
    r_total, pval = stats.pearsonr(data_entries_df['allelic_edits'].values, data_entries_df['log1p_total_counts'].values)
    sb.violinplot(data_entries_df, x='allelic_edits', y='log1p_total_counts',
                  color=color_violin)
    plt.title(f"Pearson R: {round(r_total, 5)}")
    if save_fig:
        plt.savefig(f'{out_plots}/log1p_total_counts_{gene}-allelic-edits_violin.png', dpi=300)
    plt.show()

    ### Genes detected
    r_genes, pval = stats.pearsonr(data_entries_df['allelic_edits'].values, data_entries_df['gene_counts'].values)
    sb.violinplot(data_entries_df, x='allelic_edits', y='gene_counts',
                  color=color_violin)
    plt.title(f"Pearson R: {round(r_genes, 5)}")
    if save_fig:
        plt.savefig(f'{out_plots}/gene_counts_{gene}-allelic-edits_violin.png', dpi=300)
    plt.show()

    ### Bootstrapped versions
    expr_ = data_entries_df[f'{gene}_corrected'].values
    allele_values = data_entries_df['allelic_edits'].values

    allele_set = np.unique(allele_values)
    allele_indices = [np.where(allele_values==allele_)[0] for allele_ in allele_set]

    bootstrap_entries = []
    n_straps = 10_000
    for strap_i in range(n_straps):
        for indices_, allele_ in zip(allele_indices, allele_set):

            n_to_select = len(indices_)
            select_indices = np.random.choice(indices_, n_to_select, replace=True)
            median_expr = np.median(expr_[select_indices])
            mean_expr = np.mean(expr_[select_indices])

            mean_total_counts = np.mean( log1p_total_counts[select_indices] )
            mean_gene_counts = np.mean( gene_counts[select_indices] )

            bootstrap_entries.append( [float(mean_expr), int(allele_), float(median_expr),
                                       float(mean_total_counts), float(mean_gene_counts)] )

    line_color='k'
    bootstrap_data = pd.DataFrame(bootstrap_entries, columns=[f'{gene}_corrected', 'allelic_edits',
                                                              f'{gene}_corrected_median',
                                                              'log1p_total_counts', 'gene_counts',])

    sb.violinplot(bootstrap_data, x='allelic_edits', y=f'{gene}_corrected',
                  color=color_violin, inner=None, linecolor=line_color)
    sb.boxplot(bootstrap_data, x='allelic_edits', y=f'{gene}_corrected',
                  color=line_color, whis=(2.5, 97.5), fliersize=0, width=.1, fill=False)
    if show_bootstrap_trend:
        plt.plot(bootstrap_data['allelic_edits'].values,
                 (bootstrap_data['allelic_edits'].values * alleles_beta),
                 c='red')
    if save_fig:
        plt.savefig(f'{out_plots}/{gene}-expr_allelic-edits_bootstrapped_violin.png', dpi=300)
    plt.show()

    #### Looking at total library size.
    sb.violinplot(bootstrap_data, x='allelic_edits', y=f'log1p_total_counts',
                  color=color_violin, inner=None, linecolor=line_color)
    sb.boxplot(bootstrap_data, x='allelic_edits', y=f'log1p_total_counts',
                  color=line_color, whis=(2.5, 97.5), fliersize=0, width=.1, fill=False)
    if save_fig:
        plt.savefig(f'{out_plots}/log1p_total_counts_{gene}-allelic-edits_bootstrapped_violin.png', dpi=300)
    plt.show()

    #### Looking at total gene counts.
    sb.violinplot(bootstrap_data, x='allelic_edits', y=f'gene_counts',
                  color=color_violin, inner=None, linecolor=line_color)
    sb.boxplot(bootstrap_data, x='allelic_edits', y=f'gene_counts',
               color=line_color, whis=(2.5, 97.5), fliersize=0, width=.1, fill=False)
    if save_fig:
        plt.savefig(f'{out_plots}/gene_counts_{gene}-allelic-edits_bootstrapped_violin.png', dpi=300)
    plt.show()

    return float(r_total), float(r_genes)

def plot_unmodelled_alleles(data_edited, gene, edit_name, dosage_key, condition_key,
                 color_violin='blue', save_fig = False, out_plots='./',
                 show_bootstrap_trend=True, min_cells_with_allele=25):
    """ Plots the results.
    """

    # We only want cells with evidence of being edited.
    condition = data_edited.obsm[dosage_key].loc[:, edit_name].values > 0
    if type(condition_key) != type(None):
        condition = np.logical_and(condition, data_edited.obs['sample'].values == condition_key)
    # And true negatives.
    unedited_bool = data_edited.obs['sample'].values == 'no_ki'
    condition_bool = np.logical_or(condition, unedited_bool)

    # AnnData of the cells with the particular edit.
    adata = data_edited[condition_bool, :]  #
    cell_barcodes = adata.obs_names.values

    # Classify cells into edited and unedited
    on_target_alleles = adata.obsm[dosage_key].loc[:, edit_name].values

    no_edit_bool = on_target_alleles == 0
    no_edit_indices = np.where(no_edit_bool)[0]
    edit_indices = np.where(~no_edit_bool)[0]

    # Removing edit_indices corresponding to very rare alleles.
    allele_set = np.unique(on_target_alleles)
    allele_indices = [np.where(on_target_alleles == allele_)[0] for allele_ in allele_set]
    allele_counts = np.array([len(allele_indices_) for allele_indices_ in allele_indices])
    remove_allele_indices = np.where(allele_counts < min_cells_with_allele)[0]
    for remove_allele_index in remove_allele_indices:  # If no alleles to remove this won't iterate.
        edit_indices = np.array([index for index in edit_indices  # Filters out edit cell indices with this allele
                                 if index not in allele_indices[remove_allele_index]])

    # Now getting the indices for all relevant cells:
    relevant_cell_indices = np.array(list(no_edit_indices)+list(edit_indices))

    gene_expr = adata[relevant_cell_indices, gene].X[:,0]
    allele_values = on_target_alleles[relevant_cell_indices]
    data_entries_df = pd.DataFrame([allele_values, gene_expr],
                                   index=['allelic_edits', f"{gene}_expression"],
                                   columns=cell_barcodes[relevant_cell_indices]).transpose()

    import seaborn as sb

    sb.violinplot(data_entries_df, x='allelic_edits', y=f'{gene}_expression',
                  color=color_violin)
    sb.stripplot(data_entries_df, x='allelic_edits', y=f'{gene}_expression',
                 alpha=.2, edgecolor='k', linewidth=1, color=color_violin)
    if save_fig:
        plt.savefig(f'{out_plots}/{gene}-expr_allelic-edits_violin.png', dpi=300)
    plt.show()

    # Creating a boot-strapped mean representation:
    gene_counts = adata.obs['gene_count'].values[relevant_cell_indices]
    log1p_total_counts = adata.obs['log1p_total_counts'].values[relevant_cell_indices]

    data_entries_df['gene_counts'] = gene_counts
    data_entries_df['log1p_total_counts'] = log1p_total_counts

    ### Calculating pearson
    r_total, pval = stats.pearsonr(data_entries_df['allelic_edits'].values, data_entries_df['log1p_total_counts'].values)
    sb.violinplot(data_entries_df, x='allelic_edits', y='log1p_total_counts',
                  color=color_violin)
    plt.title(f"Pearson R: {round(r_total,5)}")
    if save_fig:
        plt.savefig(f'{out_plots}/log1p_total_counts_{gene}-allelic-edits_violin.png', dpi=300)
    plt.show()

    ### Genes detected
    r_genes, pval = stats.pearsonr(data_entries_df['allelic_edits'].values, data_entries_df['gene_counts'].values)
    sb.violinplot(data_entries_df, x='allelic_edits', y='gene_counts',
                  color=color_violin)
    plt.title(f"Pearson R: {round(r_genes,5)}")
    if save_fig:
        plt.savefig(f'{out_plots}/gene_counts_{gene}-allelic-edits_violin.png', dpi=300)
    plt.show()

    ### Now boot-strapped version
    allele_set = np.unique(allele_values)
    allele_indices = [np.where(allele_values==allele_)[0] for allele_ in allele_set]

    bootstrap_entries = []
    n_straps = 10_000
    for strap_i in range(n_straps):
        for indices_, allele_ in zip(allele_indices, allele_set):

            n_to_select = len(indices_)
            select_indices = np.random.choice(indices_, n_to_select, replace=True)
            median_expr = np.median(gene_expr[select_indices])
            mean_expr = np.mean(gene_expr[select_indices])

            mean_total_counts = np.mean( log1p_total_counts[select_indices] )
            mean_gene_counts = np.mean( gene_counts[select_indices] )

            bootstrap_entries.append( [float(mean_expr), int(allele_), float(median_expr),
                                       float(mean_total_counts), float(mean_gene_counts)] )

    line_color='k'
    bootstrap_data = pd.DataFrame(bootstrap_entries,
                                  columns=[f'{gene}_expression', 'allelic_edits', f'{gene}_expression_median',
                                           'log1p_total_counts', 'gene_counts', ])

    sb.violinplot(bootstrap_data, x='allelic_edits', y=f'{gene}_expression',
                  color=color_violin, inner=None, linecolor=line_color)
    sb.boxplot(bootstrap_data, x='allelic_edits', y=f'{gene}_expression',
                  color=line_color, whis=(2.5, 97.5), fliersize=0, width=.1, fill=False)
    if save_fig:
        plt.savefig(f'{out_plots}/{gene}-expr_allelic-edits_bootstrapped_violin.png', dpi=300)
    plt.show()

    #### Looking at total library size.
    sb.violinplot(bootstrap_data, x='allelic_edits', y=f'log1p_total_counts',
                  color=color_violin, inner=None, linecolor=line_color)
    sb.boxplot(bootstrap_data, x='allelic_edits', y=f'log1p_total_counts',
                  color=line_color, whis=(2.5, 97.5), fliersize=0, width=.1, fill=False)
    if save_fig:
        plt.savefig(f'{out_plots}/log1p_total_counts_{gene}-allelic-edits_bootstrapped_violin.png', dpi=300)
    plt.show()

    #### Looking at total gene counts.
    sb.violinplot(bootstrap_data, x='allelic_edits', y=f'gene_counts',
                  color=color_violin, inner=None, linecolor=line_color)
    sb.boxplot(bootstrap_data, x='allelic_edits', y=f'gene_counts',
               color=line_color, whis=(2.5, 97.5), fliersize=0, width=.1, fill=False)
    if save_fig:
        plt.savefig(f'{out_plots}/gene_counts_{gene}-allelic-edits_bootstrapped_violin.png', dpi=300)
    plt.show()

    ### Now trying with the median instead.
    sb.violinplot(bootstrap_data, x='allelic_edits', y=f'{gene}_expression_median',
                  color=color_violin, inner=None, linecolor=line_color)
    sb.boxplot(bootstrap_data, x='allelic_edits', y=f'{gene}_expression_median',
                  color=line_color, whis=(2.5, 97.5), fliersize=0, width=.1, fill=False)
    if save_fig:
        plt.savefig(f'{out_plots}/{gene}-expr_allelic-edits_bootstrapped-medians_violin.png', dpi=300)
    plt.show()

    return float(r_total), float(r_genes)

########################################################################################################################
                            # Functions related to downstream enrichment analysis #
########################################################################################################################
def get_cont_table(de_genes, marked_genes, hvgs):
    cont_table = np.zeros((2,2), dtype=int) #rows: de, no de; cols: marked_gene, nonmarked_gene

    cont_table[0, 0] = len(de_genes.intersection(marked_genes))
    cont_table[1, 0] = len(marked_genes) - cont_table[0, 0]
    cont_table[0, 1] = len(de_genes) - cont_table[0, 0]
    cont_table[1, 1] = len(hvgs) - cont_table.sum()

    return cont_table

def get_fet_stats(cont_table):
    # FET test
    fet_test = scipy.stats.fisher_exact(cont_table)
    pval, odds_ratio_ = fet_test.pvalue, fet_test.statistic

    res = odds_ratio(cont_table)
    conf_int = res.confidence_interval(confidence_level=0.95)
    lower_odds = conf_int.low
    upper_ods = conf_int.high

    return pval, odds_ratio_, lower_odds, upper_ods

def compare_de_enrichments(on_targets, nurd_edited, nurd_gene_sets, baf_gene_sets, # dicts
                              gene_to_marginal_de, tested_genes):
    baf_edited = [gene for gene in on_targets if gene not in nurd_edited]
    baf_tests = len(baf_edited)*len(baf_gene_sets)
    nurd_tests = len(nurd_edited)*len(nurd_gene_sets)

    marginal_results = {'baf': np.zeros((baf_tests, 5)),
                        'nurd': np.zeros((nurd_tests, 5))} #cols: pval, odds_ratio, 0.95 CI lower bound, 0.95 CI upper bound, padj
    marginal_tables = {}
    test_n_baf = 0
    test_n_nurd = 0
    test_names_baf = []
    test_name_nurd = []
    for i, ontarget in enumerate(on_targets):

        de_genes_marginal = gene_to_marginal_de[ontarget]

        if ontarget in nurd_edited:
            #marked_genes = nurd_genes
            gene_sets = nurd_gene_sets
        else:
            #marked_genes = baf_genes
            gene_sets = baf_gene_sets

        for gene_set_name, marked_genes in gene_sets.items():
            test_name = f'{ontarget}_{gene_set_name}'

            marginal_cont_table = get_cont_table(de_genes_marginal, marked_genes, tested_genes)
            marginal_stats = get_fet_stats(marginal_cont_table)

            marginal_tables[test_name] = marginal_cont_table

            if ontarget in nurd_edited:
                marginal_results_ = marginal_results['nurd']

                marginal_results_[test_n_nurd, 0:4] = marginal_stats

                test_n_nurd += 1
                test_name_nurd.append(test_name)

            else:
                marginal_results_ = marginal_results['baf']

                marginal_results_[test_n_baf, 0:4] = marginal_stats

                test_n_baf += 1
                test_names_baf.append(test_name)

    test_names = {'baf': test_names_baf, 'nurd': test_name_nurd}
    for key_, test_names_ in test_names.items():

        marginal_results_ = marginal_results[key_]

        reject, pvals_corrected, _, _ = smm.multipletests(marginal_results_[:,0], alpha=0.05, method='fdr_bh')
        marginal_results_[:,-1] = pvals_corrected

        colnames = ['pval', 'odds_ratio', 'odds_ratio_0.95_lower', 'odds_ratio_0.95_upper', 'padj']
        marginal_results_ = pd.DataFrame(marginal_results_, index=test_names_,
                                        columns=colnames)

        marginal_results[key_] = marginal_results_

    return marginal_results, marginal_tables

def get_variable_genes_of_set(data_edited, gene_set, min_mean, min_disp):
    # Let's get the nurd_genes with best signal-to-noise ratio.
    means = data_edited[:, gene_set].var['means'].values
    disps = data_edited[:, gene_set].var['dispersions_norm'].values

    keep_bool = np.logical_and(means > min_mean, disps > min_disp)

    print("Genes selected:", sum(keep_bool))
    plt.scatter(means[~keep_bool], disps[~keep_bool], c='blue', s=1)
    plt.scatter(means[keep_bool], disps[keep_bool], c='red', s=1)
    plt.xlabel('mean')
    plt.ylabel('normalised dispersion')
    plt.show()

    return np.array(list(gene_set))[keep_bool]

def get_de_dict(sig_edits_stats, marginal, cutoff=None, up=True, down=True, get_all_genes=False,
                effect_size_cutoff=None, update_sig_col=False):

    gene_to_marginal_de = {}
    for ontarget in sig_edits_stats:

        de_df = sig_edits_stats[ontarget]

        if not get_all_genes:
            all_genes = [ontarget]
        else:
            padj_cols = [col for col in sig_edits_stats[ontarget].columns if col.endswith('padj')]
            all_genes = [col.split('_')[0] for col in padj_cols]

        for gene in all_genes:
            prefix = ''
            if not marginal:
                prefix = f'{gene}_'

            keep_bool = np.full((de_df.shape[0]), fill_value=False)
            if up:
                keep_bool = de_df[f'{prefix}allele_beta'].values > 0
            if down:
                keep_bool = np.logical_or(keep_bool, de_df[f'{prefix}allele_beta'].values < 0)

            if type(cutoff)==type(None):
                keep_bool = np.logical_and(keep_bool, de_df[f'{prefix}significant'].values )
            else:
                keep_bool = np.logical_and(keep_bool, de_df[f'{prefix}padj'].values < cutoff)

            if type(effect_size_cutoff)!=type(None):
                keep_bool = np.logical_and(keep_bool, np.abs(de_df[f'{prefix}allele_beta'].values) >= effect_size_cutoff)

            if update_sig_col:
                de_df[f'{prefix}significant'] = keep_bool
                print(f"Updated {prefix}significant")

            de_genes = de_df.index.values[keep_bool]
            gene_to_marginal_de[gene] = set(list(de_genes))

    return gene_to_marginal_de

def selected_extra_genes(nurd_genes, hvgs, data_edited, min_mean, min_disp):
    nurd_tested = nurd_genes.intersection(hvgs)
    nurd_genes_not_tested = nurd_genes.difference(hvgs)
    nurd_gene_candidates = list(set(list(data_edited.var_names.values)).intersection(nurd_genes_not_tested))

    extra_nurd_genes =get_variable_genes_of_set(data_edited, nurd_gene_candidates, min_mean, min_disp)

    return extra_nurd_genes, nurd_tested
```
