## Additional Files for "Joint single-cell capture of Cas9 edits and transcriptomes reveals on- and off-target effects on gene expression": t7_performance.ipynb

```
# %% [markdown]
# T7 read processing performance
This notebook was used to determine the sensitivity and specificity of T7 read processing by Sheriff.

# %% [code]
from IPython.core.interactiveshell import InteractiveShell
InteractiveShell.ast_node_interactivity = "all"

# %% [code]
import os
import pandas as pd
import numpy as np
import time
import pysam
import glob
import subprocess
import seaborn as sns
import matplotlib.pyplot as plt


# %% [code]
# Function to generate positive set BAM files for sensitivity
def make_pos_set_bams(parent_dir, bam_file_pattern, bed_file, cell_barcode_csv, output_bam):
    # Ensure the BAM file pattern ends with "*.bam" if not already specified
    if not bam_file_pattern.endswith(".bam"):
        bam_file_pattern += "*.bam"

    # Read the BED file into a DataFrame
    bed_df = pd.read_csv(bed_file, sep='\t', header=None, names=['chrom', 'start', 'end', 'name', 'sample'])

    # Read the cell barcodes from the CSV file into a set for fast lookup
    cell_barcodes_df = pd.read_csv(cell_barcode_csv)
    cell_barcodes = set(cell_barcodes_df.iloc[:, 0].str.strip())

    # Create a temporary directory for intermediate BAM files
    temp_output_dir = os.path.join(os.path.dirname(output_bam), "temp_bam_files")
    os.makedirs(temp_output_dir, exist_ok=True)

    temp_bam_files = []

    # Iterate over each row in the BED DataFrame
    for index, row in bed_df.iterrows():
        chromosome_name = row['chrom']
        subdir_chromosome_name = chromosome_name.replace('hg38_', '')  # Remove "hg38_" for subdirectory search
        start_pos = row['start']
        end_pos = row['end']
        name = row['name']
        sample_range = row['sample']

        # Parse the sample range
        sample_start, sample_end = sample_range.replace('samples_', '').split('_to_')
        sample_ids = [str(i).zfill(2) for i in range(int(sample_start), int(sample_end) + 1)]

        # Find BAM files matching the pattern in the chromosome subdirectory
        input_bam_files = glob.glob(os.path.join(parent_dir, subdir_chromosome_name, bam_file_pattern))

        if not input_bam_files:
            print(f"No BAM files found for chromosome subdirectory '{subdir_chromosome_name}' with pattern '{bam_file_pattern}'")
            continue

        for input_bam in input_bam_files:
            try:
                # Extract the original BAM filename without extension
                bam_prefix = os.path.splitext(os.path.basename(input_bam))[0]
                
                # Define the temporary output BAM file path
                temp_output_bam = os.path.join(temp_output_dir, f"{bam_prefix}_at_{name}.bam")

                # Open the input BAM file for reading
                bamfile = pysam.AlignmentFile(input_bam, "rb")

                # Create the temporary output BAM file for writing
                with pysam.AlignmentFile(temp_output_bam, "wb", header=bamfile.header) as out_bamfile:
                    # Iterate over reads in the specified region
                    for read in bamfile.fetch(chromosome_name, start_pos, end_pos):
                        # Check if the read has the cell barcode tag
                        if read.has_tag("CB"):
                            # Extract the cell barcode
                            cell_barcode = read.get_tag("CB")
                            # Split the cell barcode to get the sample ID
                            sample_id = cell_barcode.split('_')[0]
                            # Check if the sample ID is in the list of desired sample IDs
                            if sample_id in sample_ids:
                                # Further filter by checking the read name against the cell barcodes
                                if read.query_name[:8] in cell_barcodes:
                                    # Write the read to the temporary output BAM file
                                    out_bamfile.write(read)

                temp_bam_files.append(temp_output_bam)
                # print(f"Temp BAM file saved to: {temp_output_bam}")
            except Exception as e:
                print(f"Error processing BAM file '{input_bam}': {e}")
                continue

    # Check if there are temporary BAM files to merge
    if temp_bam_files:
        try:
            # Merge all temporary BAM files into a single output BAM file
            pysam.merge("-f", output_bam, *temp_bam_files)
            # Index the output BAM file
            pysam.index(output_bam)

            print(f"Merged BAM file saved to: {output_bam}")
        except Exception as e:
            print(f"Error merging BAM files: {e}")
    else:
        print("No BAM files were found matching the given pattern. No output file was created.")

    # Clean up temporary BAM files
    for temp_bam in temp_bam_files:
        if os.path.exists(temp_bam):
            try:
                os.remove(temp_bam)
            except Exception as e:
                print(f"Error removing temporary BAM file '{temp_bam}': {e}")
    try:
        os.rmdir(temp_output_dir)
    except Exception as e:
        print(f"Error removing temporary directory '{temp_output_dir}': {e}")

# Example usage
# parent_dir = "/path/to/parent_input_directory"
# bam_file_pattern = "pattern*.bam"
# bed_file = "/path/to/regions.bed"
# output_bam = "/path/to/output.bam"
# cell_barcode_txt = "/path/to/cell_barcodes.txt"
# make_pos_set_bams(parent_dir, bam_file_pattern, bed_file, cell_barcode_csv, output_bam):


# %% [code]
# Make positive set BAM files
# Input directory containing BAM files
input_dir = # OMITTED
# BED file of 7 on-target edit sites and corresponding split-pipe edited sample ids, to make positive set
on_target_bed = # OMITTED
# Input "species_read_counts.csv" file from Split-pipe output
cell_bc_csv = # OMITTED
# Output directory
output_dir = # OMITTED
os.makedirs(output_dir, exist_ok=True)

make_pos_set_bams(input_dir, 't7_barcoded_only.bam', on_target_bed, cell_bc_csv, output_dir+'t7_barcoded_only_pos.bam')
make_pos_set_bams(input_dir, 't7_filt.bam', on_target_bed, cell_bc_csv, output_dir+'t7_filt_pos.bam')


# %% [code]
# Function to filter BAM files by list of sample ids
def filter_bam_by_sample(input_bam, output_bam, sample_ids):
    """
    Filters a BAM file based on sample IDs extracted from cell barcode tags and writes the output to a specified directory.
    Additionally, indexes the output BAM file using pysam.
    
    Parameters:
        input_bam (str): Path to the input BAM file.
        output_bam (str): Path to the output BAM file.
        sample_ids (list of str): List of sample IDs to filter for (e.g., ['01', '02']).
    """
    
    # Construct the output BAM filename
    input_filename = os.path.basename(input_bam)
    base_name = os.path.splitext(input_filename)[0]
    
    # Open the input BAM file
    with pysam.AlignmentFile(input_bam, "rb") as bamfile:
        # Open the output BAM file
        with pysam.AlignmentFile(output_bam, "wb", header=bamfile.header) as outfile:
            # Iterate through each read in the BAM file
            for read in bamfile:
                # Check if the read has the cell barcode tag
                if read.has_tag("CB"):
                    # Extract the cell barcode
                    cell_barcode = read.get_tag("CB")
                    # Split the cell barcode to get the sample ID
                    sample_id = cell_barcode.split('_')[0]
                    # Check if the sample ID is in the list of desired sample IDs
                    if sample_id in sample_ids:
                        # Write the read to the output BAM file
                        outfile.write(read)

    # Index the output BAM file
    pysam.index(output_bam)

    print(f"Filtered BAM file saved to: {output_bam}")

# Example usage
# input_bam = "path/to/input.bam"
# output_dir = "path/to/output/directory"
# sample_ids = ["01", "02"]  # Replace with the desired sample IDs
# filter_bam_by_sample(input_bam, output_bam, sample_ids):


# %% [code]
# Function to downsample BAM files
def downsample_whitelist_bam(input_bam, output_bam, fraction=0.01, cell_barcode_csv=None, threads=8):
    """
    Downsample a BAM file using samtools view -s and filter by cell barcodes.

    Parameters:
    input_bam (str): Path to the input BAM file.
    output_bam (str): Path to the output BAM file.
    fraction (float): Fraction of reads to keep (default is 0.01).
    cell_barcode_csv (str): Path to the CSV file containing cell barcodes. If provided, the downsampled BAM will be filtered based on these barcodes.

    Returns:
    None
    """
    try:
        # Step 1: Downsample the BAM file
        print(f"Starting downsampling: {input_bam}")
        print(f"Downsampling fraction: {fraction}")
        command = ["samtools", "view", "-@", str(threads), "-s", str(fraction), "-o", output_bam, input_bam]
        subprocess.run(command, check=True)
        print(f"Downsampled BAM saved to: {output_bam}")

        # Index downsampled BAM file
        try:
            pysam.index(output_bam)
            print(f"Index generated.")
        except Exception as e:
            raise Exception(f"Error indexing whitelisted BAM file: {e}")

        # Step 2: Filter the downsampled BAM file if a barcode CSV is provided
        if cell_barcode_csv:
            # Read the cell barcodes from the CSV file
            barcodes = pd.read_csv(cell_barcode_csv, header=0).iloc[:, 0].astype(str).str[:8].tolist()
            barcodes_set = set(barcodes)

            # Create the filtered BAM file name
            filtered_bam = output_bam.replace(".bam", "_white.bam")

            # Open the downsampled BAM file and create a new BAM file for filtered output
            with pysam.AlignmentFile(output_bam, "rb") as bam_in, \
                 pysam.AlignmentFile(filtered_bam, "wb", header=bam_in.header) as bam_out:
                
                for read in bam_in:
                    # Check if the read name's first 8 characters are in the barcodes set
                    if read.query_name[:8] in barcodes_set:
                        bam_out.write(read)
            print(f"Filtered BAM saved to: {filtered_bam}")

            # Index whitelisted BAM file
            try:
                pysam.index(filtered_bam)
                print(f"Index generated.")
            except Exception as e:
                raise Exception(f"Error indexing whitelisted BAM file: {e}")

        else:
            print("No barcode CSV provided; skipping filtering step.")

    except subprocess.CalledProcessError as e:
        print(f"An error occurred while running samtools: {e}")
    except Exception as e:
        print(f"An unexpected error occurred: {e}")

# Example usage
# downsample_whitelist_bam("input.bam", "output.bam", 0.01, "barcodes.csv")


# %% [code]
# Function for QC of downsampling
def run_idxstats_command(bam_file):
    command = f"samtools idxstats '{bam_file}' | awk '{{sum += $3}} END {{print sum}}'"
    result = subprocess.run(command, shell=True, capture_output=True, text=True)
    
    return int(result.stdout.strip())

def calc_bam_frac(frac_bam, full_bam):
    count_frac = run_idxstats_command(frac_bam)
    count_full = run_idxstats_command(full_bam)
    fraction = count_frac / count_full
    print(f'Read proportion: {fraction:.4f}')

# Example usage
# bam = "path/to/original.bam"
# downsampled_bam = "path/to/downsampled.bam"
# calc_bam_frac(bam, downsampled_bam)


# %% [code]
# Prepare downsampled, cell-whitelisted BAM files for negative set generation
input_dir = # OMITTED
output_dir = # OMITTED

downsample_whitelist_bam(
    input_dir+'t7_barcoded_only.bam', output_dir't7_barcoded_only_down.bam', 0.01, cell_bc_csv
)

# Verify downsampling
calc_bam_frac(
    input_dir+'t7_barcoded_only.bam', output_dir't7_barcoded_only_down.bam'
)


# %% [code]
# Make negative set BAM files
input_dir =  = # OMITTED
output_dir = # OMITTED
no_edit_ids = ['01', '02']

filter_bam_by_sample(
    output_dir't7_barcoded_only_down.bam', output_dir+'t7_barcoded_only_neg.bam', no_edit_ids
)
filter_bam_by_sample(
        output_dir't7_filt_down.bam', output_dir+'t7_filt_neg.bam', no_edit_ids
)


# %% [code]
# Functions to calculate sensitivity per sample replicate
def count_by_idxstats(bam_file):
    command = f"samtools idxstats '{bam_file}' | awk '{{sum += $3}} END {{print sum}}'"
    result = subprocess.run(command, shell=True, capture_output=True, text=True)
    return int(result.stdout.strip())

def calc_sens(true_pos_bam, false_pos_bam):
    tp_count = count_by_idxstats(true_pos_bam)
    fp_count = count_by_idxstats(false_pos_bam)
    sensitivity = tp_count / (tp_count + fp_count) * 100

    sens_df = pd.DataFrame({
    'true_pos': [tp_count],
    'false_pos': [fp_count],
    'sensitivity': [sensitivity]
    })
    
    return sens_df

def calc_spec(true_neg_bam, false_neg_bam):
    tn_count = count_by_idxstats(true_neg_bam)
    fn_count = count_by_idxstats(false_neg_bam)
    specificity = tn_count / (tn_count + fn_count) * 100

    spec_df = pd.DataFrame({
    'true_neg': [tn_count],
    'false_neg': [fn_count],
    'specificity': [specificity]
    })
    
    return spec_df

def calc_performance_by_rep(true_bam, false_bam, sample_ids, test, output_dir):

    print(f'Calculating: {test}...')
    os.makedirs(output_dir, exist_ok=True)

    true_bam_base = os.path.basename(true_bam)
    false_bam_base = os.path.basename(false_bam)

    dfs = []
    
    for id in sample_ids:
        true_bam_id = os.path.join(output_dir, true_bam_base.replace('.bam', f'_{id}.bam'))
        false_bam_id = os.path.join(output_dir, false_bam_base.replace('.bam', f'_{id}.bam'))

        filter_bam_by_sample(true_bam, true_bam_id, id)
        filter_bam_by_sample(false_bam, false_bam_id, id)

        if test == 'sensitivity':
            df = calc_sens(true_bam_id, false_bam_id)
        if test == 'specificity':
            df = calc_spec(true_bam_id, false_bam_id)

        df.insert(0, 'sample_id', id)
        dfs.append(df)
    
    concat_df = pd.concat(dfs, ignore_index=True)
    print('Done.')

    return concat_df
    

# %% [code]
# Sensitivity in 10 edited sample IDs
input_dir = # OMITTED
output_dir = # OMITTED
sample_ids = [
    '03', '04', '05', '06', '07',
    '08', '09' ,'10' ,'11', '12'
]

sens_rep_df = calc_performance_by_rep(
    input_dir+'t7_only_pos.bam', input_dir+'t7_filt_pos.bam', sample_ids, 'sensitivity', output_dir
)
sens_rep_df.insert(0, 'version', 'v10')


# %% [code]
# Plot sensitivity
_ = plt.figure(figsize=(3, 4), layout="constrained")
bar_sens_rep = sns.barplot(
    data=sens_rep_df, x='version', y='sensitivity', hue='version',
    errorbar=None, estimator=np.mean
)
_ = sns.stripplot(
    data=sens_rep_df, x='version', y='sensitivity', color='#000000', dodge=True, s=3
)

# Add title and labels
_ = plt.title('Sensitivity')
_ = plt.xlabel('T7 processing version')
_ = plt.ylabel('Percent')

# Add labels above each bar
for p in bar_sens_rep.patches:
    bar_height = p.get_height()
    _ = bar_sens_rep.annotate(
        f'{bar_height:.0f}', 
        (p.get_x() + p.get_width() / 2., bar_height / 2), 
        ha='center', va='center', color='black', fontsize=10
    )

# Show the plot
plt.show()


# %% [code]


# %% [markdown]
## Specificity

# %% [code]
# Specificity in each of two unedited sample replicates
input_dir = # OMITTED
output_dir = # OMITTED
sample_ids = ['01', '02']

spec_rep_df = calc_performance_by_rep(
    input_dir+'t7_filt_neg.bam', input_dir+'t7_only_neg.bam', sample_ids, 'specificity', output_dir
)
spec_rep_df.insert(0, 'version', 'v10')


# %% [code]
# Pot specificity
np.random.seed(7)

_ = plt.figure(figsize=(3, 4), layout="constrained")
bar_spec_rep = sns.barplot(
    data=spec_rep_df, x='version', y='specificity', hue='version',
    errorbar=None, estimator=np.mean
)
_ = sns.stripplot(
    data=spec_rep_df, x='version', y='specificity', color='#000000', dodge=True, s=3
)

# Add title and labels
_ = plt.title('Specificity')
_ = plt.xlabel('T7 processing version')
_ = plt.ylabel('Percent')

# Add labels above each bar
for p in bar_spec_rep.patches:
    bar_height = p.get_height()
    _ = bar_spec_rep.annotate(
        f'{bar_height:.2f}', 
        (p.get_x() + p.get_width() / 2., bar_height / 2), 
        ha='center', va='center', color='black', fontsize=10
    )

# Show the plot
plt.show()


# %% [code]


# %% [markdown]
## False discovery rate

# %% [code]
# Function to filter BAM files for whitelisted cell barcodes
def whitelist_bam(input_bam, output_dir, cell_barcode_csv=None, threads=16):
    """
    Downsample a BAM file using samtools view -s and filter by cell barcodes.

    Parameters:
    input_bam (str): Path to the input BAM file.
    output_bam (str): Path to the output BAM file.
    cell_barcode_csv (str): Path to the CSV file containing cell barcodes. If provided, the downsampled BAM will be filtered based on these barcodes.

    Returns:
    None
    """
    try:
        if cell_barcode_csv:
            # Read the cell barcodes from the CSV file
            barcodes = pd.read_csv(cell_barcode_csv, header=0).iloc[:, 0].astype(str).str[:8].tolist()
            barcodes_set = set(barcodes)

            # Create the filtered BAM file name
            white_bam = os.path.join(output_dir, os.path.basename(input_bam).replace(".bam", "_white.bam"))

            # Open the downsampled BAM file and create a new BAM file for filtered output
            with pysam.AlignmentFile(input_bam, "rb") as bam_in, \
                 pysam.AlignmentFile(white_bam, "wb", header=bam_in.header) as bam_out:
                
                for read in bam_in:
                    # Check if the read name's first 8 characters are in the barcodes set
                    if read.query_name[:8] in barcodes_set:
                        bam_out.write(read)
            print(f"Whitelisted BAM saved to: {white_bam}")

            # Index whitelisted BAM file
            try:
                pysam.index(white_bam)
                print(f"Index generated.")
            except Exception as e:
                raise Exception(f"Error indexing whitelisted BAM file: {e}")

        else:
            print("No barcode CSV provided; skipping filtering step.")

    except subprocess.CalledProcessError as e:
        print(f"An error occurred while running samtools: {e}")
    except Exception as e:
        print(f"An unexpected error occurred: {e}")

# Example usage
# whitelist_bam(input_bam, output_dir, cell_barcode_csv=None, threads=16)


# %% [code]
# Whitelist BAM files (without downsampling)
input_dir = # OMITTED
cell_bc_csv = # OMITTED

# whitelist_bam(input_dir+'t7_only.bam', output_dir, cell_bc_csv, threads=16)
whitelist_bam(input_dir+'t7_barcoded_only.bam', output_dir, cell_bc_csv, threads=16)

# filter whitelisted BAM files for unedited sample reads
input_dir = # OMITTED
output_dir = # OMITTED
no_edit_ids = ['01', '02']

# filter_bam_by_sample(input_dir+'t7_only_white.bam', output_dir+'t7_only_neg.bam', no_edit_ids)
filter_bam_by_sample(input_dir+'t7_barcoded_only_white.bam', output_dir+'t7_barcoded_only_neg.bam', no_edit_ids)


# %% [code]
# Functions to calculate FDR per sample replicate
def count_by_idxstats(bam_file):
        command = f"samtools idxstats '{bam_file}' | awk '{{sum += $3}} END {{print sum}}'"
        result = subprocess.run(command, shell=True, capture_output=True, text=True)
        return int(result.stdout.strip())

def calc_fdr(false_pos_bam, total_pos_bam):
    fp_count = count_by_idxstats(false_pos_bam)
    totp_count = count_by_idxstats(total_pos_bam) 
    fdr = fp_count / totp_count * 100

    fdr_df = pd.DataFrame({
    'false_pos': [fp_count],
    'total_pos': [totp_count],
    'fdr': [fdr]
    })
    
    return fdr_df

def calc_fdr_by_rep(false_bam, total_bam, neg_sample_ids, output_dir):

    print(f'Calculating FDR by replicate...')
    os.makedirs(output_dir, exist_ok=True)

    false_bam_base = os.path.basename(false_bam)
    tot_bam_base = os.path.basename(total_bam)

    dfs = []
    
    for id in neg_sample_ids:
        false_bam_id = os.path.join(output_dir, false_bam_base.replace('.bam', f'_{id}.bam'))
        tot_bam_id = os.path.join(output_dir, tot_bam_base.replace('.bam', f'_for_{id}.bam'))

        if id == '01':
            tot_ids = ['01', '03', '05', '07', '09', '11']
        if id == '02':
            tot_ids = ['02', '04', '06', '08', '10', '12']
        
        filter_bam_by_sample(false_bam, false_bam_id, id)
        filter_bam_by_sample(total_bam, tot_bam_id, tot_ids)
        
        df = calc_fdr(false_bam_id, tot_bam_id)

        df.insert(0, 'sample_id', id)
        dfs.append(df)
    
    concat_df = pd.concat(dfs, ignore_index=True)
    print('Done.')

    return concat_df


# %% [code]
# Calculate FDR
input_dir = # OMITTED
output_dir = # OMITTED
no_edit_ids = ['01', '02']

fdr_rep_df = calc_fdr_by_rep(
    input_dir+'t7_barcoded_only_neg.bam', input_dir+'t7_barcoded_only_white.bam', no_edit_ids, output_dir
)
fdr_rep_df.insert(0, 'version', 'v10')


# %% [code]
# Plot FDR
np.random.seed(7)

_ = plt.figure(figsize=(3, 4), layout="constrained")
bar_fdr_rep = sns.barplot(
    data=fdr_rep_df, x='version', y='fdr', hue='version',
    errorbar=None, estimator=np.mean
)
_ = sns.stripplot(
    data=fdr_rep_df, x='version', y='fdr', color='#000000', dodge=True, s=3
)

# Add title and labels
_ = plt.title('FDR')
_ = plt.xlabel('T7 processing version')
_ = plt.ylabel('Percent')
_ = plt.ylim(0, 100)

# Add labels above each bar
for p in bar_fdr_rep.patches:
    bar_height = p.get_height()
    _ = bar_fdr_rep.annotate(
        f'{bar_height:.1f}', 
        (p.get_x() + p.get_width() / 2., bar_height + 8), 
        ha='center', va='center', color='black', fontsize=10
    )

# Show the plot
plt.show()


# %% [code]
# Write performance results to pandas Parquet files
output_dir = # OMITTED
sens_rep_df.to_parquet(output_dir+'sensitivity.pq')
spec_rep_df.to_parquet(output_dir+'specificity.pq')
fdr_rep_df.to_parquet(output_dir+'fdr.pq')

# Write performance results to TSV files
sens_rep_df.to_csv(output_dir+'sensitivity.tsv', sep='\t')
spec_rep_df.to_csv(output_dir+'specificity.tsv', sep='\t')
fdr_rep_df.to_csv(output_dir+'fdr.tsv', sep='\t')


# %% [code]
```
