## Additional Files for "Joint single-cell capture of Cas9 edits and transcriptomes reveals on- and off-target effects on gene expression": utils.py

```
import os
import pickle

import numpy
import numpy as np
import pandas
import pandas as pd

import upsetplot
import matplotlib
import matplotlib.pyplot as plt

def dealWithPlot(savePlot, showPlot, closePlot, folder, plotName, dpi,
				 tightLayout=True):
	""" Deals with the current matplotlib.pyplot.
	"""

	if tightLayout:
		plt.tight_layout()

	if savePlot:
		plt.savefig(folder+plotName, dpi=dpi,
					format=plotName.split('.')[-1])

	if showPlot:
		plt.show()

	if closePlot:
		plt.close()

def saveAsPickle(pickleName, singleCellAnalysisObjects,
                 max_bytes = 2 ** 31 - 1):

	## write
	bytes_out = pickle.dumps(singleCellAnalysisObjects,
							 protocol=pickle.HIGHEST_PROTOCOL)
	with open(pickleName, 'wb') as f_out:
		for idx in range(0, len(bytes_out), max_bytes):
			f_out.write(bytes_out[idx:idx + max_bytes])

def loadPickle(pickleName, loadType='fast'):

	if loadType=='slow':
		"""
		This is a defensive way to write pickle.load, allowing for very large 
		files on all platforms.
		"""
		max_bytes = 2 ** 31 - 1
		try:
			input_size = os.path.getsize(pickleName)
			bytes_in = bytearray(0)
			with open(pickleName, 'rb') as f_in:
				for _ in range(0, input_size, max_bytes):
					bytes_in += f_in.read(max_bytes)
			obj = pickle.loads(bytes_in)
		except:
			return None
		return obj

	else:
		with open(pickleName, 'rb') as input:
			return pickle.load(input)

def get_upset_df(obj_lists, group_names):
    """ Creates the necessary input to draw an upset plot for visualising overlaps
        between multiple groups!
    Args:
        obj_lists (list<list<object>>): List of items in different groups want \
                                          to generate upset-plot for to compare.
        group_names (list<str>): List of strings indicating the names for the \
                                                               different groups.
    Returns:
        pd.DataFrame: This is a dataframe formatted in the required formatted \
                    for input to upsetplot so can visualise multi-overlaps.
    """
    all_hs_genes = []
    [all_hs_genes.extend(de_hs_) for de_hs_ in obj_lists]
    all_hs_genes = np.unique(all_hs_genes)

    de_hs_genes = obj_lists
    samples = group_names

    de_hs_vals = np.zeros((len(samples), len(all_hs_genes)))
    for i, samp in enumerate(samples):
        for j, gene in enumerate(all_hs_genes):
            if gene in de_hs_genes[i]:
                de_hs_vals[i, j] = 1
    de_hs_df = pd.DataFrame(de_hs_vals.transpose(),
                            index=all_hs_genes, columns=samples)

    upset_df = pd.DataFrame()
    col_names = samples
    for idx, col in enumerate(de_hs_df[samples]):
        temp = []
        for i in de_hs_df[col]:
            if i != 0:
                temp.append(True)
            else:
                temp.append(False)
        upset_df[col_names[idx]] = temp

    upset_df['c'] = 1
    example = upset_df.groupby(col_names).count().sort_values('c')

    return example

def upset_plot(obj_lists, group_names=None, fig_title='', min_subset_size=1,
               sort_by="cardinality", sort_groups_by=None, show=True):
    """ Creates an upset plot, a visualisation which is useful for comparing \
        overlaps between multiple groups when have more than one group.

    Args:
        obj_lists (list<list<object>>): List of items in different groups want \
                                          to generate upset-plot for to compare.
        group_names (list<str>): List of strings indicating the names for the \
                                                               different groups.
    """
    obj_lists = obj_lists[::-1]
    if type(group_names)==type(None):
        group_names = [f'group_{i}' for i in range(len(obj_lists))]
    else:
        group_names = group_names[::-1]

    upset_df = get_upset_df(obj_lists, group_names)

    upsetplot.plot(upset_df['c'], sort_by=sort_by,
                   sort_categories_by=sort_groups_by,
                   min_subset_size=min_subset_size)
    plt.title(fig_title, loc='left')
    if show:
        plt.show()

def writeDFsToExcelSheets(excel_name, data_frames, sheet_names,
						  read_files=False, index=True):
	""" Takes in a list of dataframe fileNames, \
	and writes these to an excel sheet. Automatically adds an extra column to \
	DESingle output called Type_renamed, making the following conversion from \
	the Type column: typeToType2 = {'DEs': 'DEa', 'DEa': 'DEm', 'DEg': 'Limma_DE',
														   numpy.nan: numpy.nan}

	Args:
		excel_name (str): Specifies the name of the excel sheet.

		data_frames (list-like<str> or list-like(pandas.DataFrame)):
									List of strings specifying locations of \
									dataframes to read if read_files=True; \
									otherwise list of dataframes to write \
									into excel sheets.

		sheet_names (list-like<str>): List of strings specifying the names of \
															 excel sheet to add.

		read_files (bool): Whether to read in the file names as pandas data \
						frames, or these are already pandas data frames.
	"""

	# Create a Pandas Excel writer using XlsxWriter as the engine.
	writer = pandas.ExcelWriter(excel_name,
								engine='xlsxwriter')
	workbook = writer.book

	for i, data_frame in enumerate(data_frames):
		if read_files:
			df = pandas.read_csv(data_frame, sep='\t', header=0)

		else:
			df = data_frame

		sheetName = sheet_names[i]

		# Convert the dataframe to an XlsxWriter Excel object.
		df.to_excel(writer, sheet_name=sheetName, startrow=0,
					index=index)

		# Extracting the sheet for extra formatting
		sheet = writer.sheets[sheetName]

	# Close the Pandas Excel writer and output the Excel file.
	writer._save()

def getOrderedLabelSet(labels):
	""" Gets the set of labels associated with labels ordered from most to \
		least frequent.
	"""
	labelSet = list(set(labels))
	freqs = numpy.array([len(numpy.where(labels == label)[0])
						 for label in labelSet])
	order = numpy.array(list((-freqs).argsort()))
	labelSetOrdered = list(numpy.array(labelSet)[order])

	return labelSetOrdered

def getColors(labels, labelSet=None, colorMap='tab20', rgb=False):
	""" Gets an OrderedDict of colors; the order indicates the frequency of \
	labels from largest to smallest.

	Args:
		labels (numpy.array<str>): Indicates a set of labels for observations.

		labelSet (list-like<str>): Indicates the set of labels in labels. \
									If None, calculated based on labels.

		colorMap (str): A matplotlib colormap.

		rgb (bool): If True, colors indicated by rgb value, if false hexcode.

	Returns:
		dict<str, tuple or str>: An ordered dict indicating the labels which \
					occur most to least frequently and the associated colors.
	"""
	# Determining the set of labels #
	labelSet = input.returnDefaultIfNone(labelSet,
										 getOrderedLabelSet(labels))

	# Initialising the ordered dict #
	cellTypeColors = {}

	# Ordering the cells according to their frequency and obtaining colors #
	nLabels = len(labelSet)
	cmap = plt.cm.get_cmap(colorMap, nLabels)
	rgbs = [cmap(i)[:3] for i in range(nLabels)]
	#rgbs = list(numpy.array(rgbs)[order]) # Make sure color order is the same.

	# Populating the color dictionary with rgb values or hexcodes #
	for i in range(len(labelSet)):
		cellType = labelSet[i]
		rgbi = rgbs[i]
		if not rgb:
			cellTypeColors[cellType] = matplotlib.colors.rgb2hex(rgbi)
		else:
			cellTypeColors[cellType] = rgbi

	return cellTypeColors
```
