## Additional Files for "Joint single-cell capture of Cas9 edits and transcriptomes reveals on- and off-target effects on gene expression": blacklist.ipynb

```
# %% [markdown]
# Blacklist
This notebook was used to generate a blacklist of genome regions that contain likely mapping artifacts.

# %% [code]
from IPython.core.interactiveshell import InteractiveShell
InteractiveShell.ast_node_interactivity = "all"

# %% [code]
# Function to filter input bam by split-pipe sample id(s)
import os
import glob
import pysam

def filter_bam_by_spipe_sample_id(input_dir, bam_file_pattern, sample_ids, output_dir):
    """
    Filters BAM files in a directory based on sample IDs extracted from cell barcode tags and writes the output to a specified directory.
    
    Parameters:
        input_dir (str): Path to the directory containing input BAM files.
        bam_file_pattern (str): Pattern to match BAM filenames (e.g., "*.bam").
        sample_ids (list of str): List of sample IDs to filter for (e.g., ['01', '02']).
        output_dir (str): Path to the output directory.
    """
    # Ensure the BAM file pattern ends with "*.bam" if not already specified
    if not bam_file_pattern.endswith(".bam"):
        bam_file_pattern += "*.bam"

    # Ensure the output directory exists
    os.makedirs(output_dir, exist_ok=True)
    
    # Find BAM files matching the pattern in the input directory
    input_bam_files = glob.glob(os.path.join(input_dir, bam_file_pattern))

    if not input_bam_files:
        print(f"No BAM files found in directory '{input_dir}' with pattern '{bam_file_pattern}'")
        return

    for input_bam in input_bam_files:
        try:
            # Construct the output BAM filename
            input_filename = os.path.basename(input_bam)
            base_name = os.path.splitext(input_filename)[0]
            output_filename = f"{base_name}_samples_{sample_ids[0]}_to_{sample_ids[-1]}.bam"
            output_bam = os.path.join(output_dir, output_filename)
            
            # Open the input BAM file
            with pysam.AlignmentFile(input_bam, "rb") as bamfile:
                # Open the output BAM file
                with pysam.AlignmentFile(output_bam, "wb", header=bamfile.header) as outfile:
                    # Iterate through each read in the BAM file
                    for read in bamfile:
                        # Check if the read has the cell barcode tag
                        if read.has_tag("CB"):
                            # Extract the cell barcode
                            cell_barcode = read.get_tag("CB")
                            # Split the cell barcode to get the sample ID
                            sample_id = cell_barcode.split('_')[0]
                            # Check if the sample ID is in the list of desired sample IDs
                            if sample_id in sample_ids:
                                # Write the read to the output BAM file
                                outfile.write(read)
            
            print(f"Filtered BAM file saved to: {output_bam}")
        
        except Exception as e:
            print(f"Error processing BAM file '{input_bam}': {e}")

# Example usage
# input_dir = "/path/to/input_directory"
# bam_file_pattern = "*.bam"
# sample_ids = ["01", "02", "03"]  # Replace with the desired sample IDs
# output_dir = "/path/to/output_directory"
# filter_bam_by_spipe_sample_id(input_dir, bam_file_pattern, sample_ids, output_dir)


# %% [code]
input_dir = # OMITTED
# Split-pipe aligned reads, 10k cell library, downsampled 100x
spipe_100x_down_bam = # OMITTED
sample_ids = ['01', '02']
output_dir = # OMITTED

filter_bam_by_spipe_sample_id(input_dir, spipe_100x_down_bam, sample_ids, output_dir)


# %% [code]
# Function to convert genome file to parse chrom names
import os
import pandas as pd

def genome_to_parse_chroms(input_genome_file, output_dir):
    # Read the input genome file
    genome_df = pd.read_csv(input_genome_file, sep='\t', header=None)
    
    # Perform the string processing on the first column
    genome_df[0] = genome_df[0].str.replace('chr', 'hg38_')
    genome_df[0] = genome_df[0].str.replace(r'_(.*?)_G', '_G', regex=True)
    genome_df[0] = genome_df[0].str.replace(r'_(.*?)_K', '_K', regex=True)
    genome_df[0] = genome_df[0].str.replace('Un_', '')
    genome_df[0] = genome_df[0].str.replace('v', '.')
    genome_df[0] = genome_df[0].str.replace('_alt', '')
    genome_df[0] = genome_df[0].str.replace('_random', '')

    # Extract the input filename and construct the output filename
    input_filename = os.path.basename(input_genome_file)
    output_filename = input_filename.replace('.genome', '_parse_chroms.genome')
    output_file_path = os.path.join(output_dir, output_filename)

    # Write the processed data to the output file
    genome_df.to_csv(output_file_path, sep='\t', header=False, index=False)

    print(f"Processed genome file saved to: {output_file_path}")

# Example usage
# genome_to_parse_chroms('input.genome', 'output_directory')


# %% [code]
input_genome = # OMITTED
output_dir = # OMITTED

genome_to_parse_chroms(input_genome, output_dir)


# %% [code]
# Function that calls (blacklist) peaks from input bam
import os
import subprocess
import pandas as pd

def bam_to_blacklist_peaks(bam_file, output_dir, output_prefix=""):
    
    # Create output directory if it doesn't exist
    if not os.path.exists(output_dir):
        os.makedirs(output_dir)

    # Define MACS2 command   
    # --nomodel,       Skip dynamic lambda shifting model
    # --extsize 98,    Manual alignment length for pileup when --nomodel set, 3' extension from 5' shift position
    # --max-gap 3000,  Peaks within this distance will be merged
    # --keep-dup all,  Keep all reads
    # -q 1e-50, Q-val  sig cutoff for peaks
    # -g hs,           Human genome size, for calculating background signal
    # -n,              Prefix for output files
    # -t,              Treatment bam
    # --outdir,        Output directory
    # -B --SPMR,       Output bedGraphs (fragment pileup per M reads)
    # --trackline,     Include UCSC genome browser header line in outputs
    macs2_command = f"macs2 callpeak --nomodel --extsize 98 --max-gap 3000 --keep-dup all -q 1e-50 -g hs -n {output_prefix} -t {bam_file} --outdir {output_dir} -B --SPMR --trackline"
    
    # Run MACS2 command
    subprocess.run(macs2_command, shell=True, check=True)

    # File paths for narrowPeak calling
    bed_file = os.path.join(output_dir, f'{output_prefix}_summits.bed')
    narrowpeak_file = os.path.join(output_dir, f'{output_prefix}_peaks.narrowPeak')
    treatment_bedgraph_file = os.path.join(output_dir, f'{output_prefix}_treat_pileup.bdg')
    control_bedgraph_file = os.path.join(output_dir, f'{output_prefix}_control_lambda.bdg')

    # Function to process the file: replace strings, filter lines, and modify track name
    def process_file(input_file, output_file, output_prefix):
        input_filename = os.path.basename(input_file)
        track_name = input_filename.split('_')[-1].split('.')[0]
        new_track_name = f"{output_prefix}_{track_name}"

        with open(input_file, 'r') as file:
            lines = file.readlines()

        processed_lines = []
        for i, line in enumerate(lines):
            if i == 0 and line.startswith('track'):
                # Replace the entire name field in the trackline
                line = line.replace('name="', f'name="{new_track_name}"')
                processed_lines.append(line)
                continue
            # Split the line into columns
            columns = line.strip().split('\t')
            if not (columns[0].startswith('hg38_G') or columns[0].startswith('hg38_K')):
                processed_line = line.replace('hg38_', 'chr').replace('chrMT', 'chrM')
                processed_lines.append(processed_line)

        with open(output_file, 'w') as file:
            file.writelines(processed_lines)

    # Process and write new files with "ucsc" appended
    process_file(bed_file, os.path.join(output_dir, f'{output_prefix}_summits_ucsc.bed'), output_prefix)
    process_file(narrowpeak_file, os.path.join(output_dir, f'{output_prefix}_peaks_ucsc.narrowPeak'), output_prefix)
    process_file(treatment_bedgraph_file, os.path.join(output_dir, f'{output_prefix}_treat_pileup_ucsc.bdg'), output_prefix)
    process_file(control_bedgraph_file, os.path.join(output_dir, f'{output_prefix}_control_lambda_ucsc.bdg'), output_prefix)

    # Sort .bed file by the fifth column in descending order
    sorted_bed_file = os.path.join(output_dir, f'{output_prefix}_summits_by_qval.bed')
    bed_df = pd.read_csv(bed_file, sep='\t', header=None, skiprows=1)
    bed_df[4] = pd.to_numeric(bed_df[4], errors='coerce')
    bed_df_sorted = bed_df.sort_values(by=4, ascending=False)
    bed_df_sorted.to_csv(sorted_bed_file, sep='\t', header=False, index=False)

    # Sort .narrowPeak file by descending q-value 9th col (instead of fold enrich 7th col)
    sorted_narrowpeak_file = os.path.join(output_dir, f'{output_prefix}_peaks_by_qval.narrowPeak')
    narrowpeak_df = pd.read_csv(narrowpeak_file, sep='\t', header=None, skiprows=1)
    narrowpeak_df[8] = pd.to_numeric(narrowpeak_df[8], errors='coerce')
    narrowpeak_df_sorted = narrowpeak_df.sort_values(by=8, ascending=False)
    narrowpeak_df_sorted.to_csv(sorted_narrowpeak_file, sep='\t', header=False, index=False)

    # Print total number of peaks
    total_peaks = len(narrowpeak_df_sorted)
    print(f"Total number of peaks: {total_peaks}")

    # Read the sorted bed file for q values
    sorted_bed_df = pd.read_csv(sorted_bed_file, sep='\t', header=None)
    total_q_peaks = len(sorted_bed_df)

    # Print the top three peaks with the strongest q values
    print("Top 3 peaks with strongest q values:")
    for i in range(min(3, total_q_peaks)):
        row = sorted_bed_df.iloc[i]
        location = f"{row[0]}:{row[1]}-{row[2]}"
        q_value = row[4]
        print(f"Peak {i+1}: Location = {location}, Q Value = {q_value}")

# Example usage
# bam_to_blacklist_peaks('input.bam', 'output_directory')
# bam_to_blacklist_peaks('input.bam', 'output_directory', 'output_prefix')


# %% [code]
# Split-pipe aligned reads, 10k cell library, downsampled 100x, unedited sample
# Generated with filter_bam_by_spipe_sample_id()
spipe_100x_down_no_edit_bam = # OMITTED
output_dir = # OMITTED

bam_to_blacklist_peaks(spipe_100x_down_no_edit_bam, output_dir, output_prefix='black_100x')

# %% [code]
####### Code snippets then used to add the polyN sites >50bp in length, noting that additional sites were also added afterward.
### Converting the narrowPeak file from the macs2 peak calling above to a standard bed file
# cut -f1-6 black_100x_peaks_by_qval.narrowPeak > black_100x_peaks_by_qval.bed

######## Downloading repeat masker simple repeat annotations from IGV host
# wget https://s3.amazonaws.com/igv.org.genomes/hg38/rmsk/hg38_rmsk_Simple_repeat.bed.gz
# gzip -d hg38_rmsk_Simple_repeat.bed.gz

# ALSO subset the repeats to just those that have a >60 repeats, which likely effects mapping of the 100bp reads,
# especially when they have the soft-clippped sequence and potentially the TSO!!
# awk '($2 - $1) > 70' hg38_rmsk_Simple_repeat.bed > hg38_rmsk_Simple_repeat.70N.bed
# awk '{gsub(/chr/, "hg38_"); print}' hg38_rmsk_Simple_repeat.70N.bed > hg38_rmsk_Simple_repeat.70N.renamed.bed
# cat black_100x_peaks_by_qval.bed hg38_rmsk_Simple_repeat.70N.renamed.bed > black_100x_peaks_by_qval.simple_repeats_70N.bed
#blacklist_file = f"{data_dir}black_100x_peaks_by_qval.simple_repeats_70N.bed"

# Turns out this blacklist bed is WAYY to broad. Main issues seems to be the VERY simple repeats, like for example (T)n
# Will just limit it to those cases!!!
# awk '{gsub(/chr/, "hg38_"); print}' hg38_rmsk_Simple_repeat.bed > hg38_rmsk_Simple_repeat.renamed.bed
# grep -E '\(T\)n|\(A\)n|\(C\)n|\(G\)n' hg38_rmsk_Simple_repeat.renamed.bed > hg38_rmsk_Simple_repeat.renamed.polyN.bed
# awk -F'\t' '($3 - $2) > 50' hg38_rmsk_Simple_repeat.renamed.polyN.bed > hg38_rmsk_Simple_repeat.renamed.polyN.50N.bed

#### Check by printing diffs:
# awk  -F'\t' '{print $3 - $2}' hg38_rmsk_Simple_repeat.renamed.polyN.50N.bed

### Make it so they are just 1 bp peaks, so need most of the simple repeat to overlap the edit region!
# awk -F'\t' '{print $1 "\t" int($2 + (($3 - $2) / 2)) "\t" int(($2 + (($3 - $2) / 2))+1)}' hg38_rmsk_Simple_repeat.renamed.polyN.50N.bed > hg38_rmsk_Simple_repeat.renamed.polyN.50N.1bp.bed
# cat black_100x_peaks_by_qval.min.bed hg38_rmsk_Simple_repeat.renamed.polyN.50N.1bp.bed > black_100x_peaks_by_qval.simple_repeats_50N.bed

# Renamed this to the chr convention, so can look at these in IGV:
# awk '{gsub("hg38_", "chr"); print}' black_100x_peaks_by_qval.simple_repeats_50N.bed > black_100x_peaks_by_qval.simple_repeats_50N.renamed.bed

#### Now making adding the additional region which corresponded to a reference genome bias that looked like barcodes!:
# This region added at top:
# hg38_12 56112901    56112902
# hg38_12 56112801    56113001
# black_100x_peaks_by_qval.simple_repeats_50N.EXTRA.bed
#
# Renaming for visualisation purposes:
# awk '{gsub("hg38_", "chr"); print}' black_100x_peaks_by_qval.simple_repeats_50N.EXTRA.bed > black_100x_peaks_by_qval.simple_repeats_50N.EXTRA.renamed.bed
```
