## Additional Files for "Joint single-cell capture of Cas9 edits and transcriptomes reveals on- and off-target effects on gene expression": combine_results.py

```
# Combined results from the super_count processing across multiple chromosomes.
# Assumes being run from the directory where the outputs from superb_count_multi.sh are directed!
# Example for running:
# cd superb_count_all
# conda activate t7_variant_calling
# nohup python combine_results.py 1>combine.out 2>combine.err &

import os, sys
import subprocess

import pandas as pd

import scanpy as sc

import time

data_dir3 = 'DGE_filtered/'
out_dir = 'combined/'

########## Making an output directory to retrieve the results.
os.system('mkdir combined')

########## First need to get a list of chromosomes, have a little bit of python code to achieve this.

# Command to list the chromosomes, excluding specified patterns
# Decided to remove mitochondrial genome, since cas9 allegedly cannot get into here anyhow, and takes forever to process
command = "ls | grep -vE '(KI2|GL0|MT|superb_count|superb_data|combined|.err|.out|combine)'"

# Use subprocess.run to execute the command and capture the output
result = subprocess.run(command, shell=True, stdout=subprocess.PIPE, stderr=subprocess.PIPE, text=True)

# Capture the output
chroms = result.stdout.split()

##################### Now compiling all the bam files!!!
print(f"Compiling bams across chromosomes, including renaming to standard chromosome names so can visualise with normal genome browsers.", file=sys.stdout, flush=True)
bam_file_names = ['t7_barcoded_only.bam']#, 't7_non-barcoded_only.bam', 't7_only.bam', 't7_filt.bam']
print(" ".join(bam_file_names), file=sys.stdout, flush=True)
def join_bams(bam_file_names):
    for bam_file_name in bam_file_names:
        chrom_files = [f'{chrom}/{bam_file_name}' for chrom in chroms]
        os.system(f"samtools cat -o {out_dir}{bam_file_name} {' '.join(chrom_files)}")
        os.system(f"samtools index {out_dir}{bam_file_name}")

        # Renaming to normalise chromosome naming conventions for visualisation of edits.
        if bam_file_name == 't7_barcoded_only.bam': # Just doing this bam, since this operation takes alot of time.
            awk_str = "awk "+"'{ gsub("+'"hg38_"'+', "chr", $0); print }'+"'"
            os.system(f"samtools view -h {out_dir}{bam_file_name} | {awk_str} > {out_dir}{bam_file_name.replace('.bam', '.renamed.sam')}")
            os.system(f"samtools view -h -b {out_dir}{bam_file_name.replace('.bam', '.renamed.sam')} > {out_dir}{bam_file_name.replace('.bam', '.renamed.bam')}")
            os.system(f"rm {out_dir}{bam_file_name.replace('.bam', '.renamed.sam')}")
            os.system(f"samtools index {out_dir}{bam_file_name.replace('.bam', '.renamed.bam')}")

join_bams(bam_file_names)

########### Now let's combine the different dataframes!!!
dosages_genes_list = [] # concatted
gene_counts_list = [] # concatted
edit_site_infos = [] # concatted
t7_umis_list = [] # concatted
t7_no_bc_list = [] # concatted
dosages_list = [] # concatted 
t7_edits_list = [] # concatted

engine = 'pyarrow' #'fastparquet'
start = time.time()
for chrom in chroms:
    n_edits = list(open(f'{chrom}/edit_sites.bed', 'r'))
    if n_edits == 0:
        print(f"NO edit for chrom, so bypassing compiling the edit information: {chrom}")

    else:
        print(f"Loading data for chrom: {chrom}")
        dosages_genes_chrom = pd.read_parquet(f'{chrom}/cell_allelic_dosage.canonical-edit-sites.gene-collapsed.parquet.gz', engine=engine)
        dosages_chrom = pd.read_parquet(f'{chrom}/cell_allelic_dosage.canonical-edit-sites.parquet.gz', engine=engine)
        edit_site_info_chrom = pd.read_csv(f'{chrom}/edit_site_info.txt', sep='\t')
        t7_umis_chrom = pd.read_parquet(f'{chrom}/t7_barcoded_counts.parquet.gz', engine=engine)
        t7_no_bc_chrom = pd.read_parquet(f'{chrom}/t7_nonbarcoded_counts.parquet.gz', engine=engine)
        t7_edits_chrom = pd.read_csv(f'{chrom}/t7_barcode_edits.tsv', sep='\t')

        dosages_genes_list.append( dosages_genes_chrom )
        edit_site_infos.append( edit_site_info_chrom )
        t7_umis_list.append( t7_umis_chrom )
        dosages_list.append( dosages_chrom )
        t7_no_bc_list.append( t7_no_bc_chrom )
        t7_edits_list.append( t7_edits_chrom )

    gene_counts_chrom = pd.read_parquet(f'{chrom}/cell_gene_mrna_counts.parquet.gz', engine=engine)
    gene_counts_list.append(gene_counts_chrom)

    end = time.time()
    print(f"Finished loading data for chrom {chrom} in {round((end-start)/50, 3)} minutes", file=sys.stdout, flush=True)

###### Compiling the results...
# edit_site_info, load to make sure all the outputs have a consistent ordering of the edits!
edit_site_info = pd.concat(edit_site_infos)
edit_site_info.index = edit_site_info['name'].values
edit_site_info.to_csv(f'{out_dir}edit_site_info.txt', sep='\t')

# gene_counts, do this first and make sure everything else ordered the same way!!
print(f"Compiling and saving compiled matrices across chromosomes.", file=sys.stdout, flush=True)
gene_counts = gene_counts_list[0]
for gene_counts_ in gene_counts_list[1:]:
    gene_counts += gene_counts_
gene_counts.to_parquet(f'{out_dir}cell_gene_mrna_counts.parquet.gz', compression="gzip")

# Checked and the below is order ordered correctly according to the cell ordering and edit ordering!
cell_whitelist = gene_counts.index.values
edit_names = edit_site_info.index.values

# Particular edit information
t7_edits = pd.concat(t7_edits_list, axis=0)
t7_edits.to_csv(f'{out_dir}t7_barcode_edits.tsv', sep='\t')

# t7 UMIs
t7_umis = pd.concat(t7_umis_list, axis=1)
t7_no_bc_umis = pd.concat(t7_no_bc_list, axis=1)
t7_all_counts = t7_umis + t7_no_bc_umis

t7_umis.to_parquet(f'{out_dir}t7_barcoded_counts.parquet.gz', compression="gzip")
t7_no_bc_umis.to_parquet(f'{out_dir}t7_nonbarcoded_counts.parquet.gz', compression="gzip")
t7_all_counts.to_parquet(f'{out_dir}t7_all_counts.parquet.gz', compression="gzip")

# allelic dosages
dosages_genes = pd.concat(dosages_genes_list, axis=1)
dosages = pd.concat(dosages_list, axis=1)

dosages.to_parquet(f'{out_dir}cell_allelic_dosage.canonical-edit-sites.parquet.gz', compression="gzip")
dosages_genes.to_parquet(f'{out_dir}cell_allelic_dosage.canonical-edit-sites.gene-collapsed.parquet.gz', compression="gzip")

######## Now creating an AnnData object with most of the analysis performed!
print(f"Creating AnnData of compiled matrices.", file=sys.stdout, flush=True)
cell_metadata = pd.read_csv(data_dir3+'cell_metadata.csv')
cell_metadata.index = cell_metadata['bc_wells']
cell_metadata = cell_metadata.loc[cell_whitelist, :]

data = sc.AnnData(gene_counts, obs=cell_metadata)
data.layers['gene_counts'] = data.X.copy()
data.obsm['dosages_genes'] = dosages_genes
data.obsm['dosages'] = dosages
data.obsm['t7_umis'] = t7_umis
data.obsm['t7_no_bc_umis'] = t7_no_bc_umis
data.obsm['t7_all'] = t7_all_counts

for i, col in enumerate(edit_site_info.columns):
    if col == 'chr':
        edit_site_info[col] = edit_site_info[col].values.astype(str)
        edit_site_info[col] = edit_site_info[col].astype( str )
        continue

    edit_site_info[col] = edit_site_info[col].astype( type(edit_site_info.values[0,i]) ) 

data.uns['edit_site_info'] = edit_site_info

print(f"Basic single cell processing on the AnnData.", file=sys.stdout, flush=True)
sc.pp.calculate_qc_metrics(data, inplace=True)

sc.pp.normalize_total(data)
sc.pp.log1p(data)

sc.pp.highly_variable_genes(data)

sc.pp.pca(data)

sc.pp.neighbors(data, metric="cosine")

sc.tl.umap(data)

print(f"Saving AnnData with structure:", file=sys.stdout, flush=True)
print(data, file=sys.stdout, flush=True)
print(f"Saving AnnData to : {out_dir}superb_data.h5ad", file=sys.stdout, flush=True)
data.write_h5ad(f"{out_dir}superb_data.h5ad", compression='gzip')

##################### Compiling the less important bams!
print(f"Compiling bams across chromosomes, including renaming to standard chromosome names so can visualise with normal genome browsers.",
    file=sys.stdout, flush=True)
bam_file_names = ['t7_non-barcoded_only.bam', 't7_only.bam', 't7_filt.bam']
print(" ".join(bam_file_names), file=sys.stdout, flush=True)
join_bams(bam_file_names)

##################### Now just concatenating outputs that have no header.
print(f"Concatentating across chromosomes output files with no header:", file=sys.stdout, flush=True)
file_names = ['t7_barcoded_reads.txt', 't7_reads.txt', 't7_non-barcoded_reads.txt', 'edit_sites.bed']
print(" ".join(file_names), file=sys.stdout, flush=True)
for file_name in file_names:
    chrom_files = [f'{chrom}/{file_name}' for chrom in chroms]
    os.system(f'cat {" ".join(chrom_files)} > {out_dir}{file_name}')

print(f"DONE compiling data across chromosomes!!", file=sys.stdout, flush=True)
 
```
