## Additional Files for "Joint single-cell capture of Cas9 edits and transcriptomes reveals on- and off-target effects on gene expression": create_anndata.py

```
""" Built for when have run the t7 read processing on JUST the full bam, i.e. not split by chromosome.
Therefore is no need to actually rejoin everything, can just creating the AnnData base on the output files.

Setup:

	nohup python create_anndata.py 1>create_anndata.500cell1.out 2>create_anndata.500cell1.err &
"""

import os, sys

import pandas as pd

import scanpy as sc

import time

out_dir = './' # Just to satisfy previous versions!

##################### Now compiling all the bam files!!!
print(f"Compiling bams across chromosomes, including renaming to standard chromosome names so can visualise with normal genome browsers.", file=sys.stdout, flush=True)
bam_file_names = ['t7_barcoded_only.bam']#, 't7_non-barcoded_only.bam', 't7_only.bam', 't7_filt.bam']
print(" ".join(bam_file_names), file=sys.stdout, flush=True)
def rename_bams(bam_file_names):
    for bam_file_name in bam_file_names:
        # Renaming to normalise chromosome naming conventions for visualisation of edits.
        awk_str = "awk "+"'{ gsub("+'"hg38_"'+', "chr", $0); print }'+"'"
        os.system(f"samtools view -h {out_dir}{bam_file_name} | {awk_str} > {out_dir}{bam_file_name.replace('.bam', '.renamed.sam')}")
        os.system(f"samtools view -h -b {out_dir}{bam_file_name.replace('.bam', '.renamed.sam')} > {out_dir}{bam_file_name.replace('.bam', '.renamed.bam')}")
        os.system(f"rm {out_dir}{bam_file_name.replace('.bam', '.renamed.sam')}")
        os.system(f"samtools index {out_dir}{bam_file_name.replace('.bam', '.renamed.bam')}")

rename_bams(bam_file_names)

######### Now compiling the AnnData
engine = 'pyarrow' #'fastparquet'
start = time.time()

print("Reading in the necessary data to create the anndata...", file=sys.stdout, flush=True)

dosages_genes = pd.read_parquet(f'cell_allelic_dosage.canonical-edit-sites.gene-collapsed.parquet.gz', engine=engine)
dosages = pd.read_parquet(f'cell_allelic_dosage.canonical-edit-sites.parquet.gz', engine=engine)
edit_site_info = pd.read_csv(f'edit_site_info.txt', sep='\t')
t7_umis = pd.read_parquet(f't7_barcoded_counts.parquet.gz', engine=engine)
t7_no_bc_umis = pd.read_parquet(f't7_nonbarcoded_counts.parquet.gz', engine=engine)
t7_all_counts = t7_umis + t7_no_bc_umis
t7_edits = pd.read_csv(f't7_barcode_edits.tsv', sep='\t')
gene_counts = pd.read_parquet(f'cell_gene_mrna_counts.parquet.gz', engine=engine)

# Checked and the below is order ordered correctly according to the cell ordering and edit ordering!
cell_whitelist = gene_counts.index.values
edit_names = edit_site_info.index.values

######## Now creating an AnnData object with most of the analysis performed!
print(f"Creating AnnData of compiled matrices.", file=sys.stdout, flush=True)

data = sc.AnnData(gene_counts)
data.layers['gene_counts'] = data.X.copy()
data.obsm['dosages_genes'] = dosages_genes
data.obsm['dosages'] = dosages
data.obsm['t7_umis'] = t7_umis
data.obsm['t7_no_bc_umis'] = t7_no_bc_umis
data.obsm['t7_all'] = t7_all_counts

for i, col in enumerate(edit_site_info.columns):
    if col == 'chr':
        edit_site_info[col] = edit_site_info[col].values.astype(str)
        edit_site_info[col] = edit_site_info[col].astype( str )
        continue

    edit_site_info[col] = edit_site_info[col].astype( type(edit_site_info.values[0,i]) )

data.uns['edit_site_info'] = edit_site_info

print(f"Basic single cell processing on the AnnData.", file=sys.stdout, flush=True)
sc.pp.calculate_qc_metrics(data, inplace=True)

sc.pp.normalize_total(data)
sc.pp.log1p(data)

sc.pp.highly_variable_genes(data)

sc.pp.pca(data)

sc.pp.neighbors(data, metric="cosine")

sc.tl.umap(data)

print(f"Saving AnnData with structure:", file=sys.stdout, flush=True)
print(data, file=sys.stdout, flush=True)
print(f"Saving AnnData to : {out_dir}superb_data.h5ad", file=sys.stdout, flush=True)
data.write_h5ad(f"{out_dir}superb_data.h5ad", compression='gzip')

print(f"DONE! Finished creating AnnData", file=sys.stdout, flush=True)
```
