## Additional Files for "Joint single-cell capture of Cas9 edits and transcriptomes reveals on- and off-target effects on gene expression": superb_count_multi.sh

```
#!/bin/bash
#$ -q iblm.q
#$ -t 1-194
#$ -tc 10
#$ -V
#$ -N superb_count_all
#$ -j y
#$ -o superb_count_all/

source conda.sh
conda activate superb_count

# Input Data
data_dir="data"

# File that maps chrom name on bam to ref, and provides paths chrom split bam and ref
chrom_map="combined_chrom_split_map_noheader.renamed.fixed.tsv"

# READ Specific line from file
input_array=($(awk "NR==$SGE_TASK_ID {print;exit}" ${chrom_map}))

chrom_bam=${input_array[0]} # chrom name in bam
chrom_ref=${input_array[1]} # chrom name in reference genome
split_bam=${input_array[2]} # chrom subset bam
split_ref=${input_array[3]} # chrom subset reference

# Other required/standard inputs
barcode_file="${data_dir}/barcode_whitelist.txt"
gtf_file="${data_dir}/Homo_sapiens.GRCh38.110.gtf"
cnv_file="${data_dir}/K562_CNV_hg19.tsv"
blacklist_file="/black_100x_peaks_by_qval.simple_repeats_50N.EXTRA.bed"
blacklist_seqs="blacklist_seqs.txt"
t7_barcode="GGGAGAGTAT"

# Options and Params
k=6 # kmer match value
edit_dist=140 # +/- distance from canonical edit site to collapse edits to that edit site.
stranded_edit_dist=15 # distance between the closest forward and reverse reads from the canonical edit site to classify as a true edit event (real edits tend to have a higly localised cut sequence!)
ploidy=2 # expected number of genome copies
mrna_count_mode='polyT'

# output directory
outdir="superb_count_all/${chrom_ref}"

# Run with options and params denoted
python superb_count/src ${split_bam} ${split_ref} ${barcode_file} ${gtf_file} --edit_dist ${edit_dist} --stranded_edit_dist ${stranded_edit_dist} --t7 ${t7_barcode} --kmer ${k} --ploidy ${ploidy} --cnv_file ${cnv_file} --blacklist_file ${blacklist_file} --blacklist_seqs ${blacklist_seqs} --mrna_count_mode ${mrna_count_mode} -o ${outdir}

## End-of-job summary, if running as a job
[[ -n "$JOB_ID" ]] && qstat -j "$JOB_ID"
```
