## Additional Files for "Joint single-cell capture of Cas9 edits and transcriptomes reveals on- and off-target effects on gene expression": superb_count_multi_500-cells.sh

```
# Running the superb-seq processing on the 500-cell library

# Run with:
# nohup ./superb_count_multi_500-cells.sh 1>500-cells_run1.out 2>500-cells_run1.err &

# Input Data
data_dir="500_cell/"
data_dir2="t7_indel_calling/"

# Other required/standard inputs
barcode_file="${data_dir}barcode_whitelist.500-cell.txt"
bam_file="${data_dir}barcode_headAligned_anno.sorted.bam"
## For testing
#bam_file="${data_dir}barcode_headAligned_anno.sorted.hg38_12.bam"

ref_file="${data_dir2}Homo_sapiens.GRCh38.dna.primary_assembly.fa"
gtf_file="${data_dir2}Homo_sapiens.GRCh38.110.gtf"
cnv_file="${data_dir2}K562_CNV_hg19.tsv"
blacklist_file="${data_dir2}black_100x_peaks_by_qval.simple_repeats_50N.EXTRA.bed"
blacklist_seqs="${data_dir2}blacklist_seqs.txt"
t7_barcode="GGGAGAGTAT"

# Options and Params
k=6 # kmer match value
edit_dist=140 # +/- distance from canonical edit site to collapse edits to that edit site.
stranded_edit_dist=15 # distance between the closest forward and reverse reads from the canonical edit site to classify as a true edit event (real edits tend to have a higly localised cut sequence!)
ploidy=2 # expected number of genome copies
mrna_count_mode='all'

##### Settings that differ from the 10k library!
edit_site_min_cells=1
whitelist_file="${data_dir}edit_site_white_list.txt"

# output directory
outdir="superb_count_500_cells1/"

# Run with options and params denoted
python superb_count/src ${bam_file} ${ref_file} ${barcode_file} ${gtf_file} --edit_dist ${edit_dist} --stranded_edit_dist ${stranded_edit_dist} --t7 ${t7_barcode} --kmer ${k} --ploidy ${ploidy} --cnv_file ${cnv_file} --blacklist_file ${blacklist_file} --blacklist_seqs ${blacklist_seqs} --mrna_count_mode ${mrna_count_mode} --edit_site_min_cells ${edit_site_min_cells} --whitelist_file ${whitelist_file} -o ${outdir}
```
