## Additional Files for "Joint single-cell capture of Cas9 edits and transcriptomes reveals on- and off-target effects on gene expression": split_pipe_500_cell.sh

```
#!/bin/bash
# Task ID range (e.g. 1-6)
#$ -t 1
# Max parallel tasks
#$ -tc 1
# Concatenate stdout, stderr
#$ -j y
# Job name
#$ -N spipe-500
# Job stdout directory
#$ -o # [OMITTED]

# SPLIT-PIPE PROCESSING OF SUPERB-SEQ READS
# 500 CELL LIBRARY
# Split-pipe v1.1.1
# Run time ~90 minutes with 24 threads

# Run name
runName="split_pipe_500_cell"
# Parent directory
parentDir= # [OMITTED]
# Input file paths (input cBC.bam)
inDir= # [OMITTED]
# Output directory
outDir= # [OMITTED]
# BWA index directory
hg38Dir= # [OMITTED]

# Source environment
source # [OMITTED]
mamba activate # [ENV NAME]
# Make variables
inR1=$inDir/*R1.fastq.gz
inR2=$inDir/*R2.fastq.gz

# SPLIT-PIPE COMMAND
# --mode all,     Run all pipeline steps
# --nthreads 24,  Threads
# --chemistry v2, Kit version
# --sample,       Sample names and bacroding round 1 well ranges
date
echo "Running split-pipe..."
echo "Processing $inR1 and $inR2..."
split-pipe \
    --mode all --nthreads 24 --chemistry v2 \
    --genome_dir $hg38Dir --output_dir $outDir \
    --fq1 $inR1 --fq2 $inR2 \
    --sample no_ki A1-A2 \
    --sample baf_ki A3-A7 \
    --sample nurd_ki A8-A12
echo "Done"
date
```
