## Additional Files for "Joint single-cell capture of Cas9 edits and transcriptomes reveals on- and off-target effects on gene expression": splitpipe_bench10k_combine.sh

```
#!/bin/bash

# activate spipe_v160 env

# set variables
threads=24
parentDir='/PATH/TO/PARENT'
outDir="/PATH/TO/OUTPUT"
lib3Dir="$parentDir/library3"
lib4Dir="$parentDir/library4"

# Run split-pipe comb, to combine sublibrary outputs
date
echo 'ML starting split-pipe v1.6.0 ...'

split-pipe \
    --mode comb \
    --nthreads $threads \
    --output_dir $outDir \
    --sublibraries $lib3Dir $lib4Dir

echo 'ML done.'
date
```
