## Additional Files for "Joint single-cell capture of Cas9 edits and transcriptomes reveals on- and off-target effects on gene expression": splitpipe_bench10k_mkref.sh

```
#!/bin/bash

# activate spipe_v160 env

# Set threads
threads=32

# Input and output directories
genomeDir='/PATH/TO/GENOME'
           /Homo_sapiens.GRCh38.114.chr.gtf.gz
date
echo 'Running split-pipe mkref v1.6.0 ...'

# Make indexed reference genome
split-pipe \
    --mode mkref \
    --genome_name hg38 \
    --nthreads $threads \
    --fasta "$genomeDir/Homo_sapiens.GRCh38.dna.primary_assembly.fa.gz" \
    --genes "$genomeDir/Homo_sapiens.GRCh38.114.gtf.gz" \
    --output_dir $genomeDir
 
date
echo 'Done.'
```
