## Additional Files for "Joint single-cell capture of Cas9 edits and transcriptomes reveals on- and off-target effects on gene expression": splitpipe_bench10k_rep2.sh

```
#!/bin/bash

# activate spipe_v160 env

# set variables
threads=24
runName='lib3_all_250821'
parentDir='/PATH/TO/PARENT'
genomeDir="/PATH/TO/GENOME"
R1_nova1="$parentDir/fastq_nova1_250715/Library3_S1_L008_R1_001.fastq.gz"
R2_nova1="$parentDir/fastq_nova1_250715/Library3_S1_L008_R2_001.fastq.gz"
R1_nova2="$parentDir/fastq_nova2_250821/Lib3Nova2_S1_L005_R1_001.fastq.gz"
R2_nova2="$parentDir/fastq_nova2_250821/Lib3Nova2_S1_L005_R2_001.fastq.gz"
R1_aviti="$parentDir/fastq_aviti_250815/Samples/McVicker_Mickey/Library3_R1.fastq.gz"
R2_aviti="$parentDir/fastq_aviti_250815/Samples/McVicker_Mickey/Library3_R2.fastq.gz"
outDir="$parentDir/splitpipe_all_250821/library3"
sampleList="$outDir/../benchmarking_sample_list.txt"

# Run full pipeline, sample definitions from list file
date
echo 'ML starting split-pipe v1.6.0 ...'

split-pipe \
    --mode all \
    --chemistry v3 \
    --kit WT_mini \
    --nthreads $threads \
    --run_name $runName\
    --genome_dir $genomeDir \
    --fq1 $R1_nova1 $R1_nova2 $R1_aviti \
    --fq2 $R2_nova1 $R2_nova2 $R2_aviti \
    --output_dir $outDir \
    --samp_list $sampleList

echo 'ML done.'
date
```
