## Additional Files for "Joint single-cell capture of Cas9 edits and transcriptomes reveals on- and off-target effects on gene expression": SuppMaterial1.pdf

manual edit call:True loc: chr1;26696719 genes=ARID1A

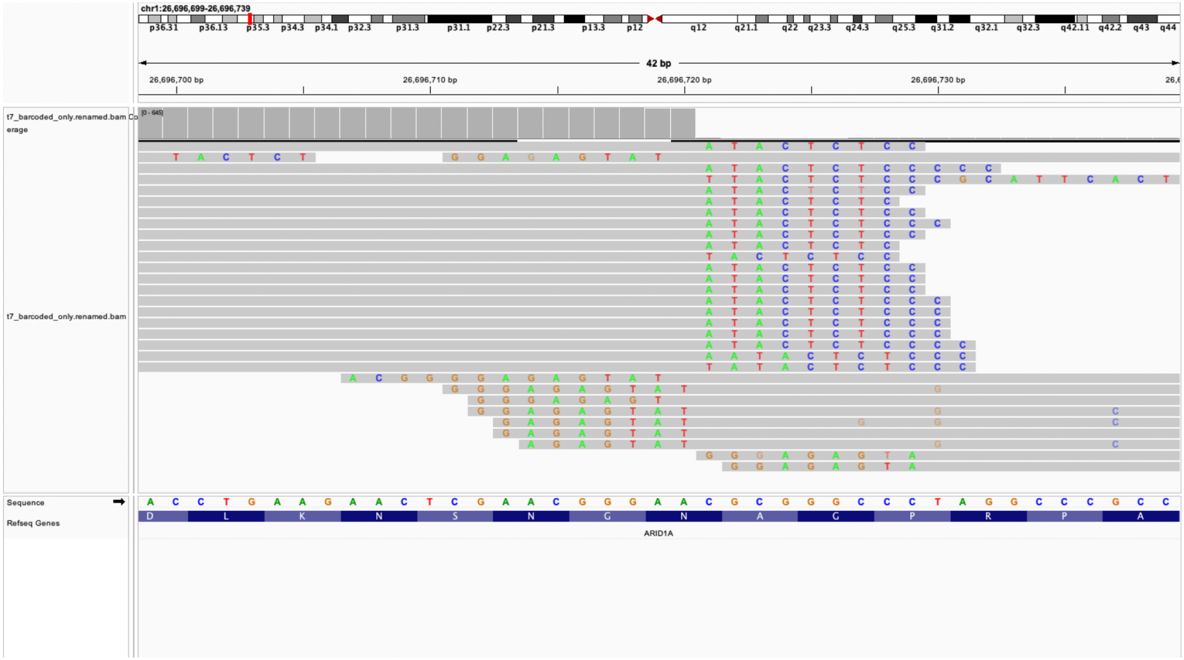

manual edit call:True loc: chr1;26696862 genes=ARID1A

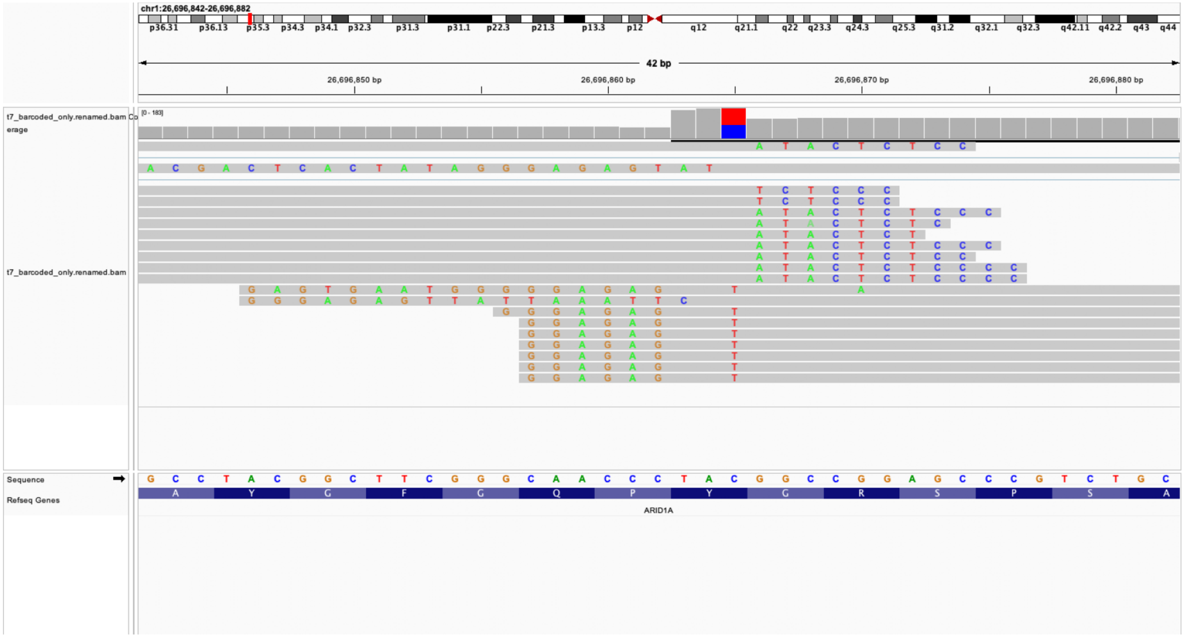

manual edit call:True loc: chr12;6602172 genes=ENSG00000285238;CH

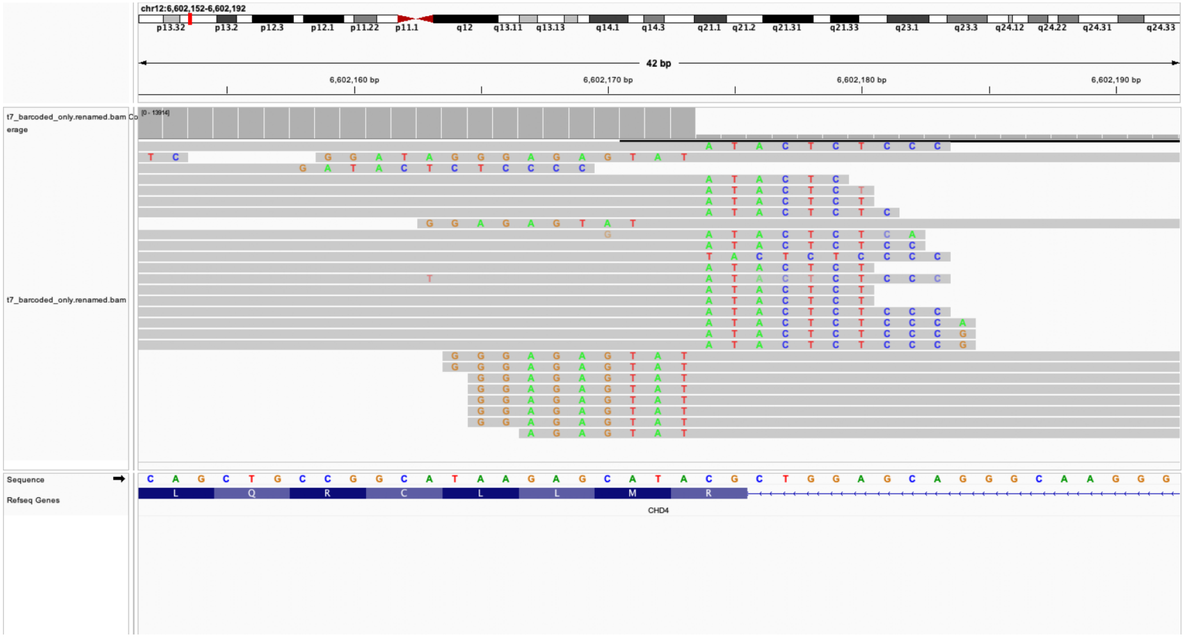

manual edit call:True loc: chr12;6606340 genes=ENSG00000285238;CH

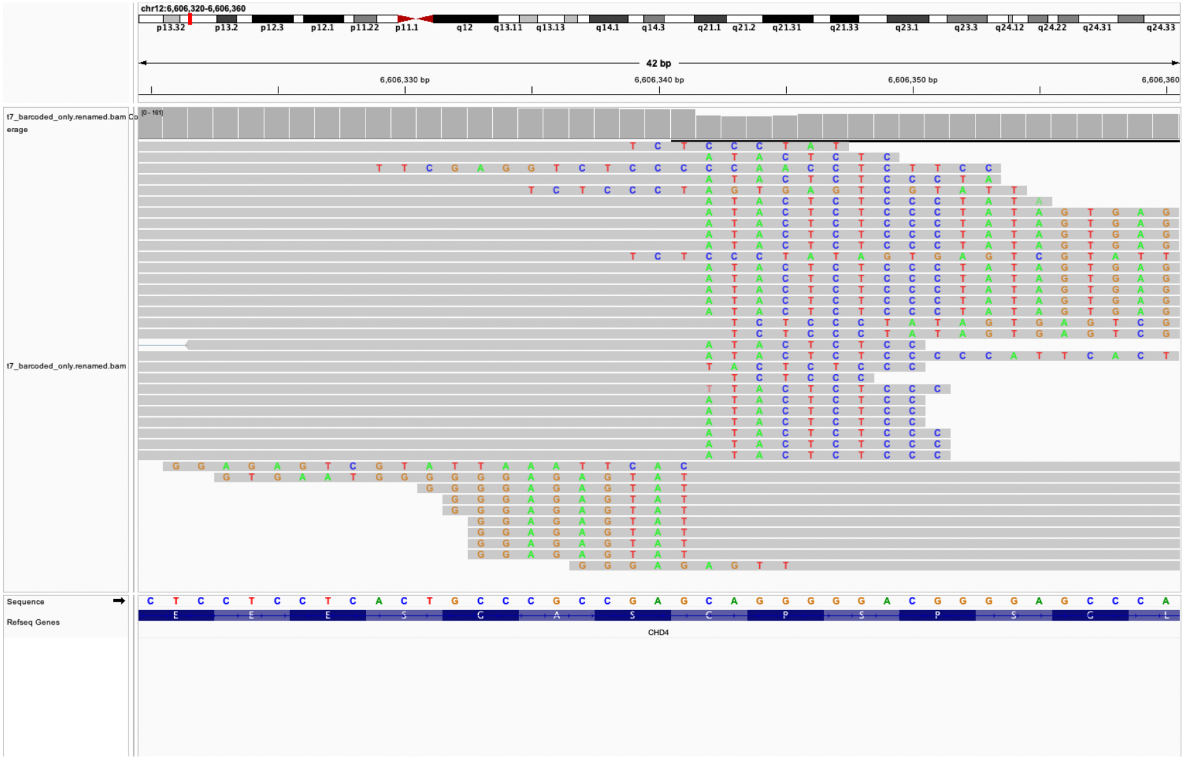

### manual edit call:True loc: chr19;10986490 genes=SMARCA4

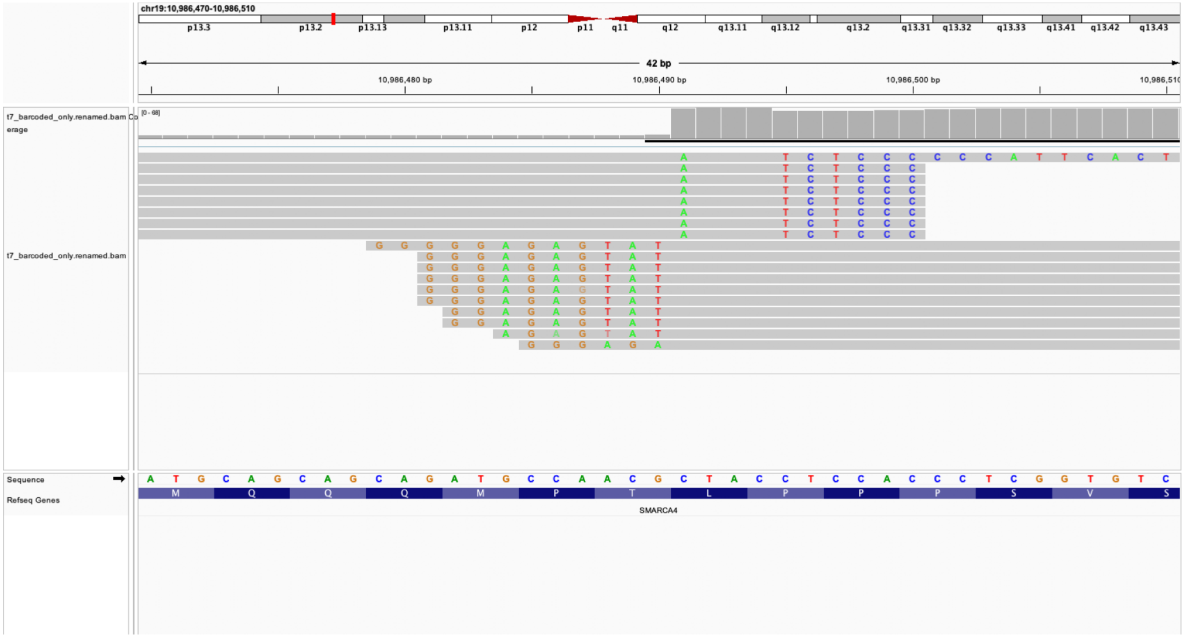

manual edit call:True loc: chr19;10984175 genes=SMARCA4

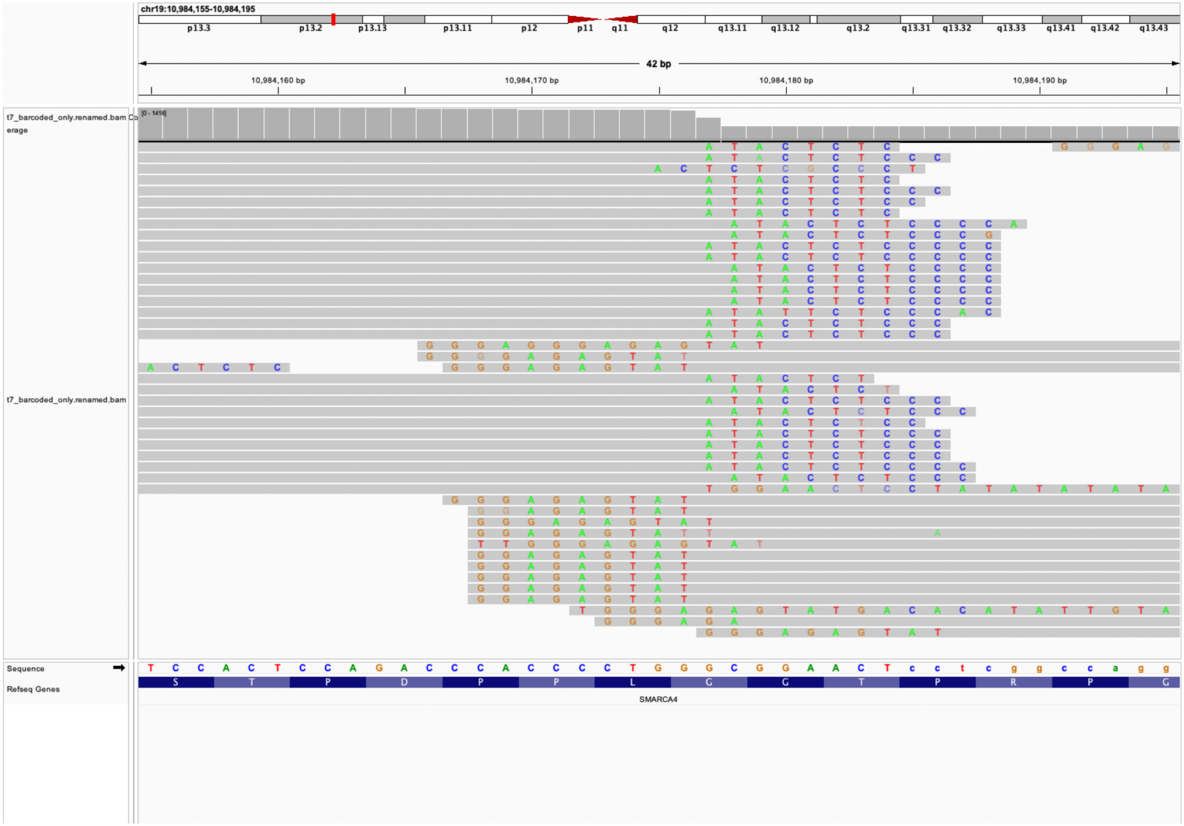

### manual edit call:True loc: chr17;7890653 genes=CHD3

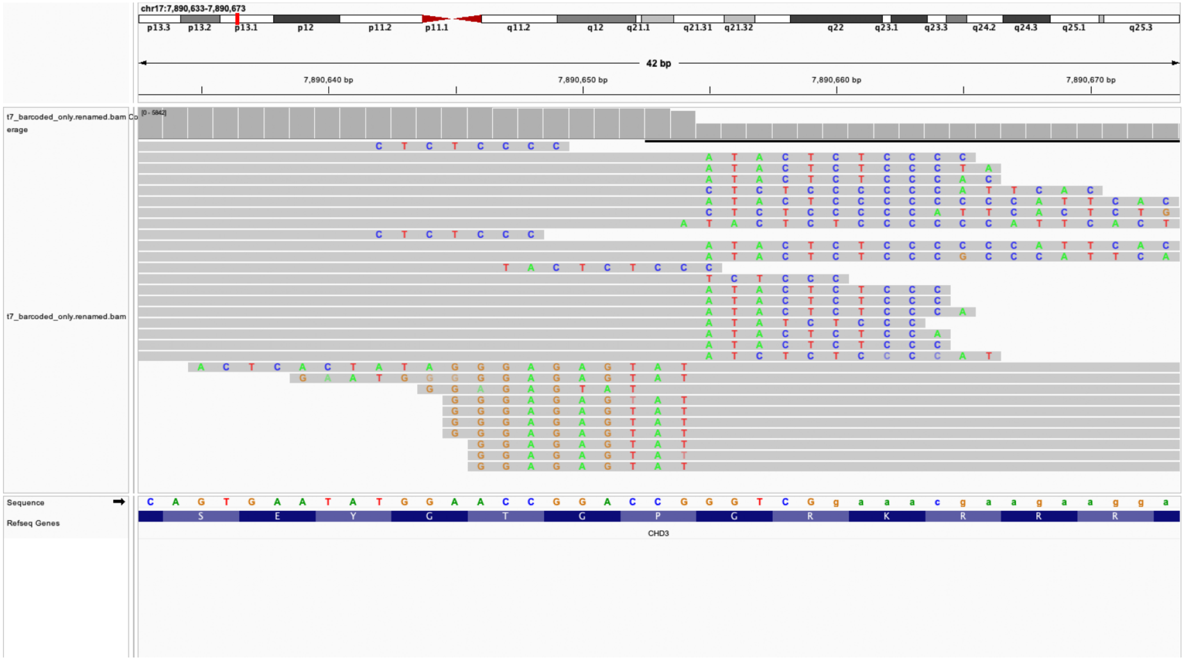

anual edit call:True loc: chr21;37649273 genes=ENSG00000286717;KC

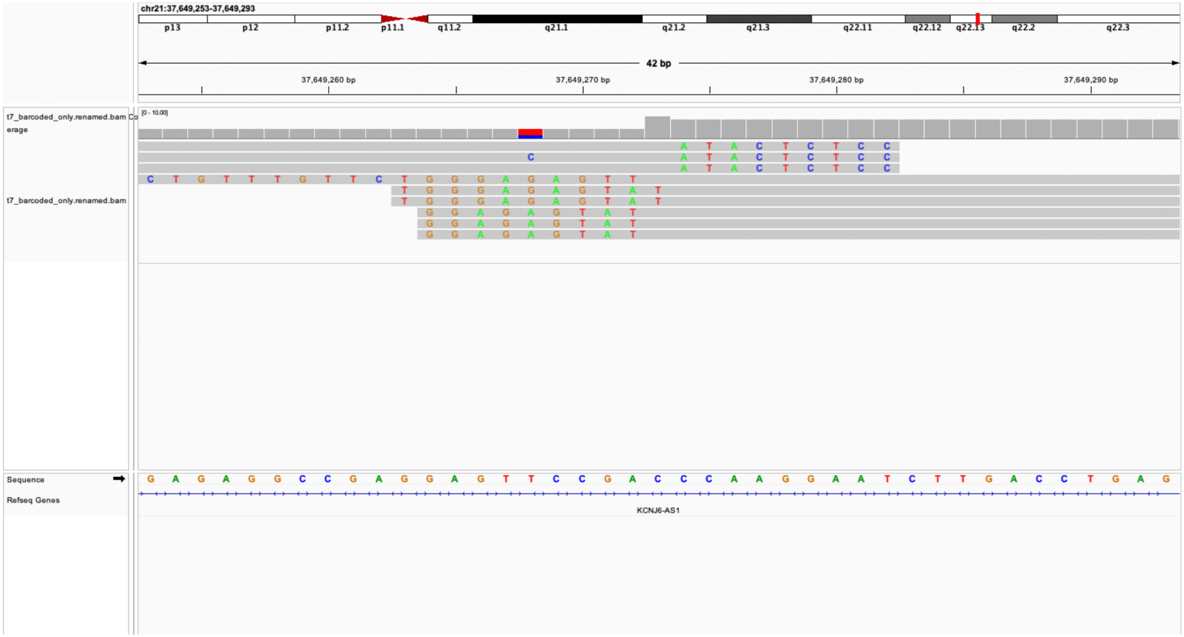

### manual edit call:True loc: chr19;6212712 genes=MLLT1

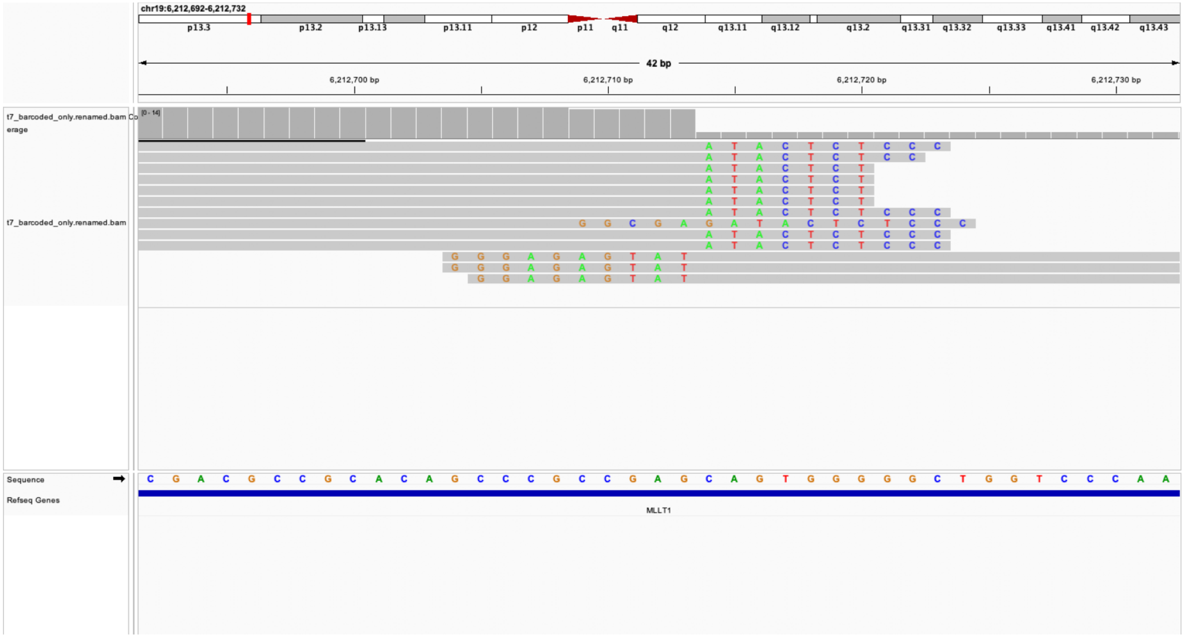

### manual edit call:True loc: chr6;1397192 genes=nan

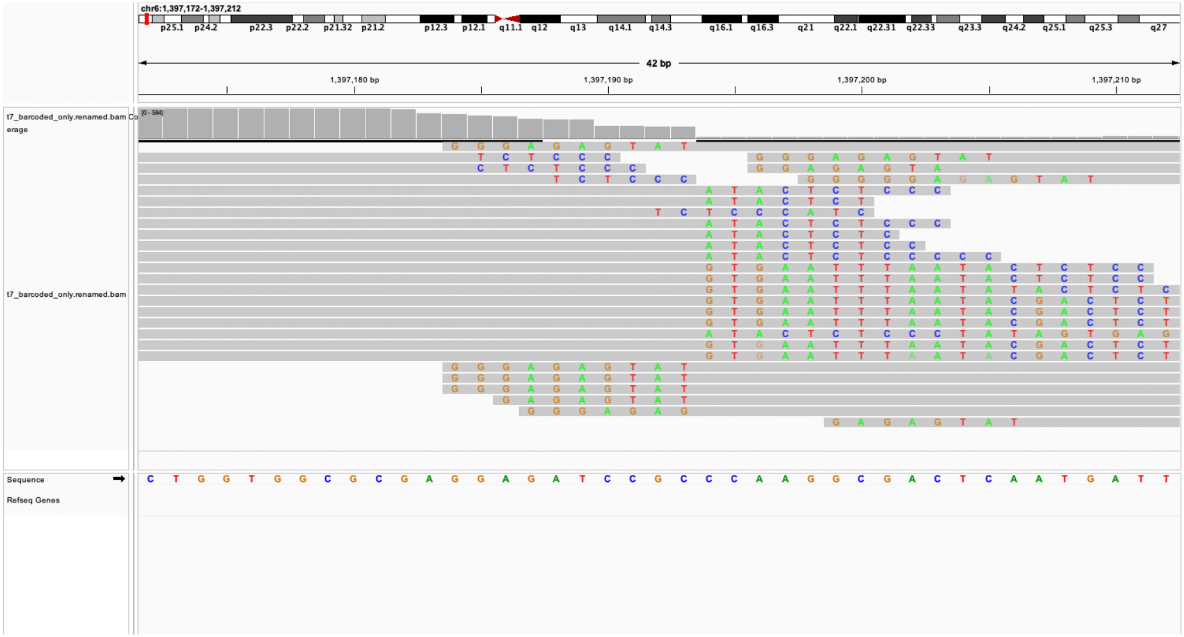

### manual edit call:True loc: chrX;41086519 genes=USP9X

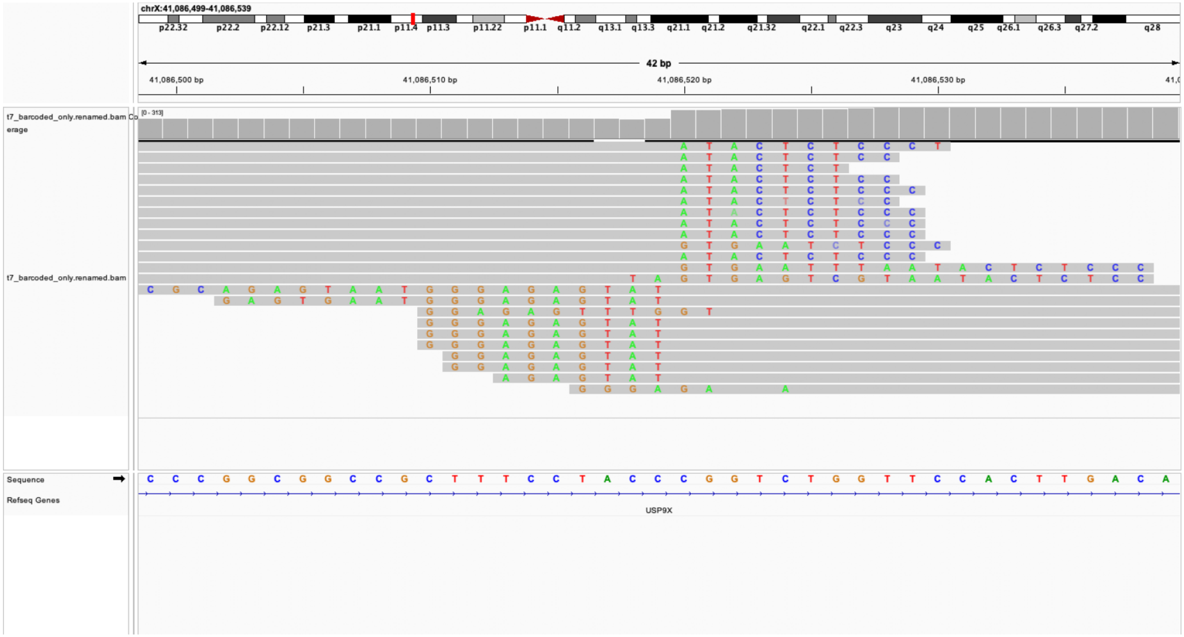

### manual edit call:True loc: chr20;14937204 genes=MACROD2

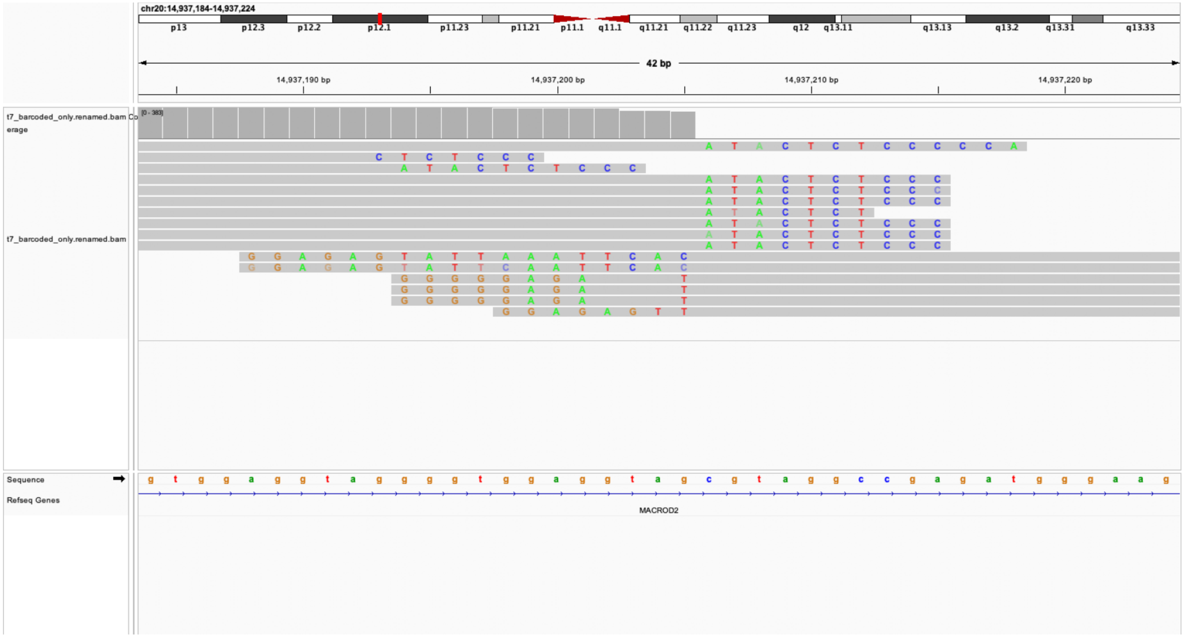

### manual edit call:True loc: chr18;3115881 genes=MYOM1

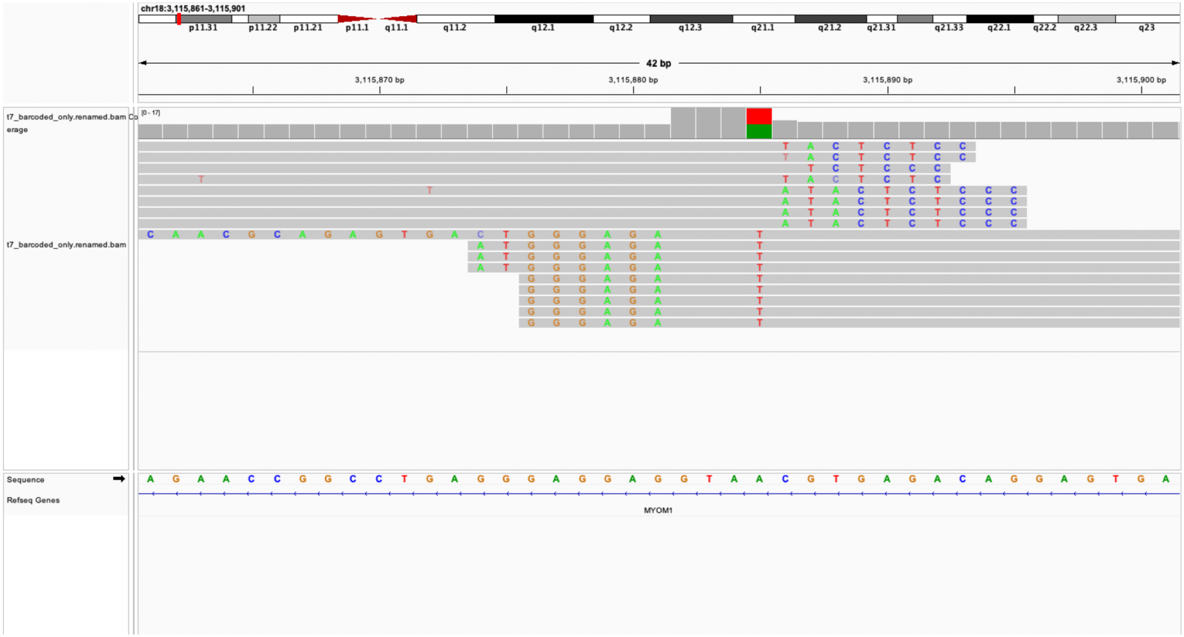

### manual edit call:True loc: chr4;14359416 genes=ENSG00000287360

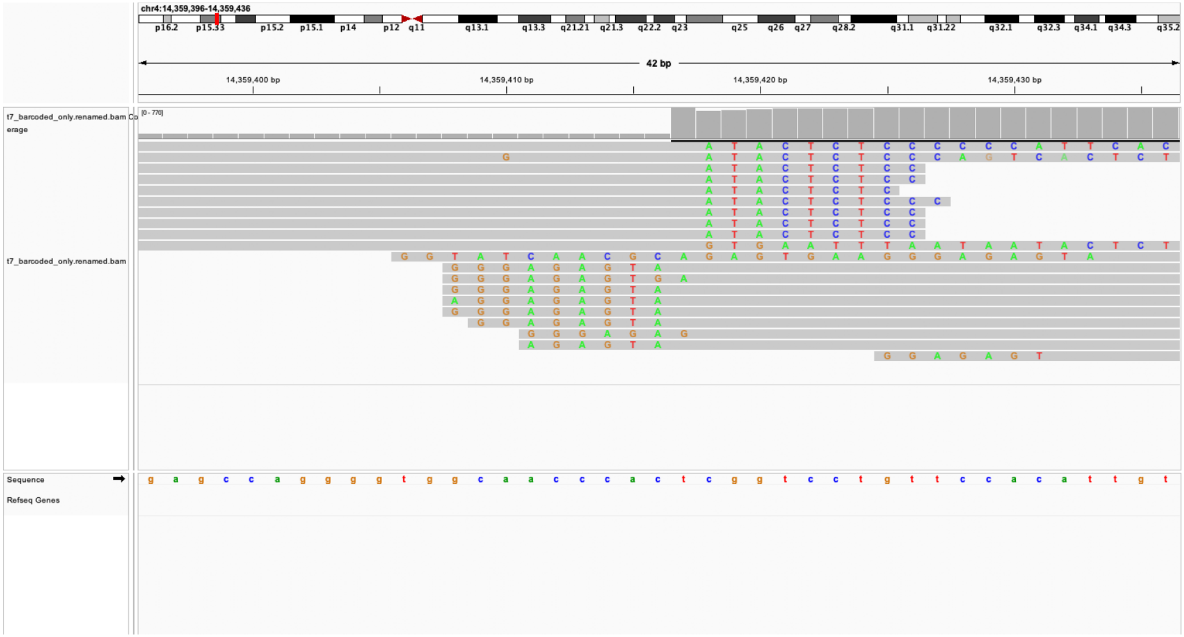

manual edit call:True loc: chr13;34520796 genes=LINC02343;LINC0045

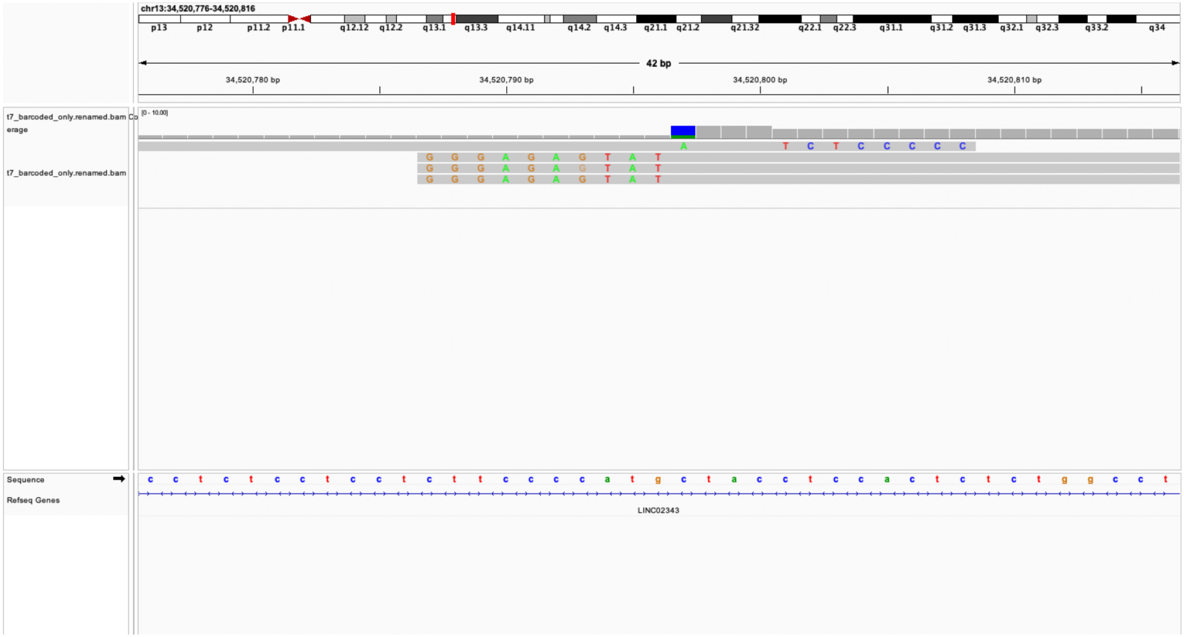

manual edit call:True loc: chr12;68620881 genes=RAP1B

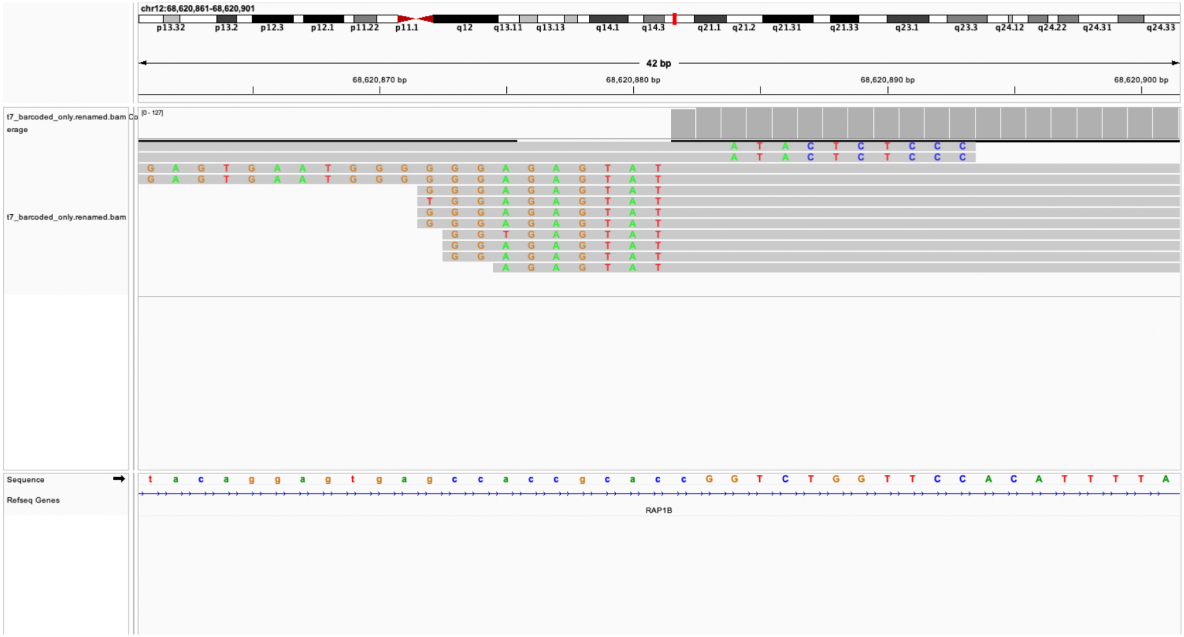

### manual edit call:True loc: chr1;11841677 genes=CLCN6

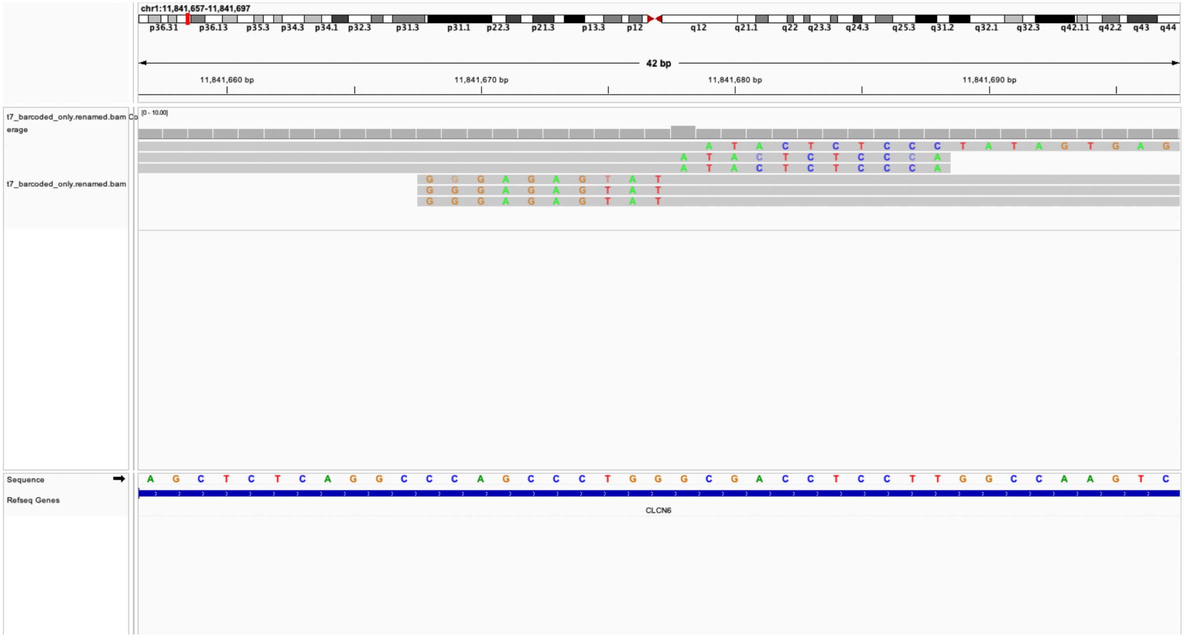

manual edit call:True loc: chr9;37594631 genes=ENSG00000255872

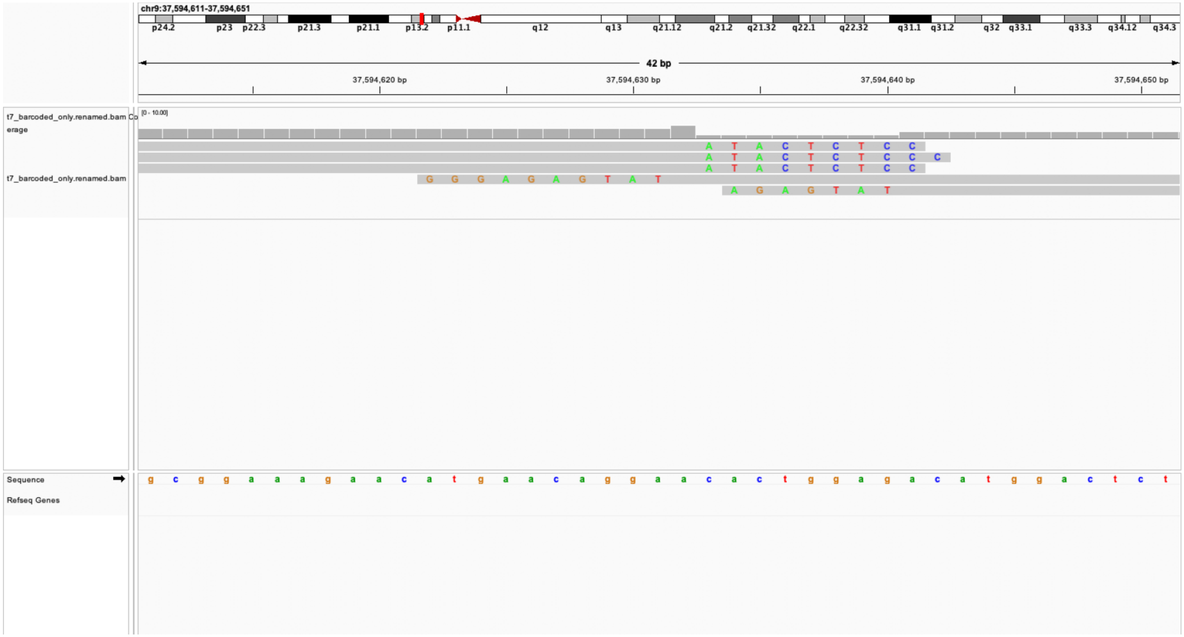

manual edit call:True loc: chr1;245947392 genes=SMYD3

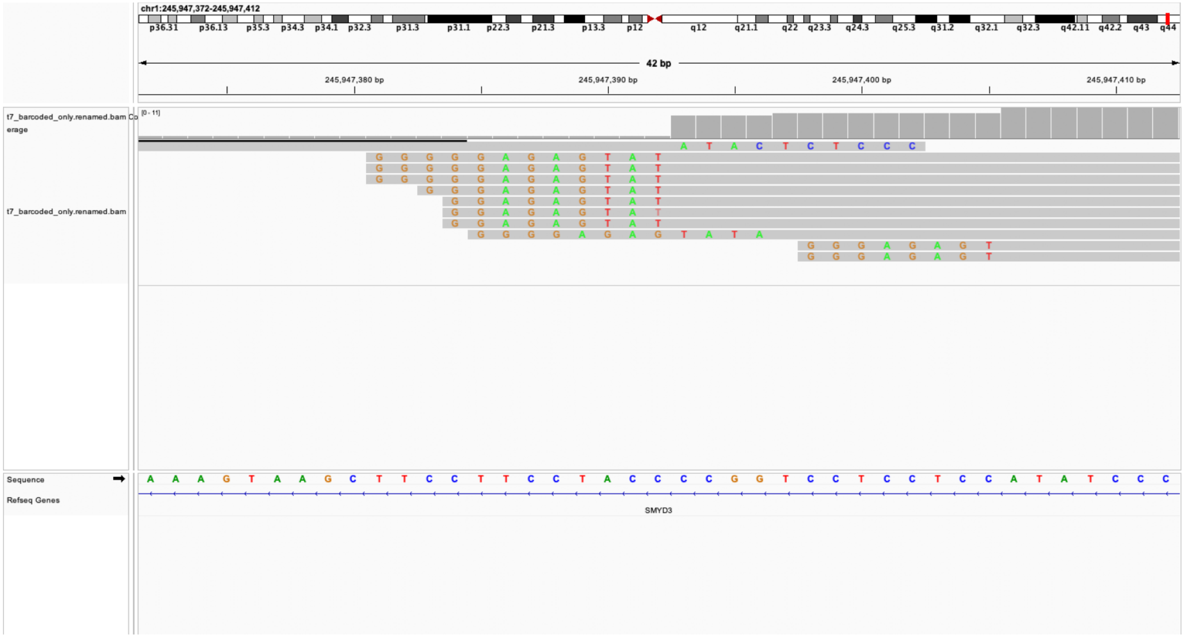

### manual edit call:True loc: chr10;93632366 genes=PDE6C

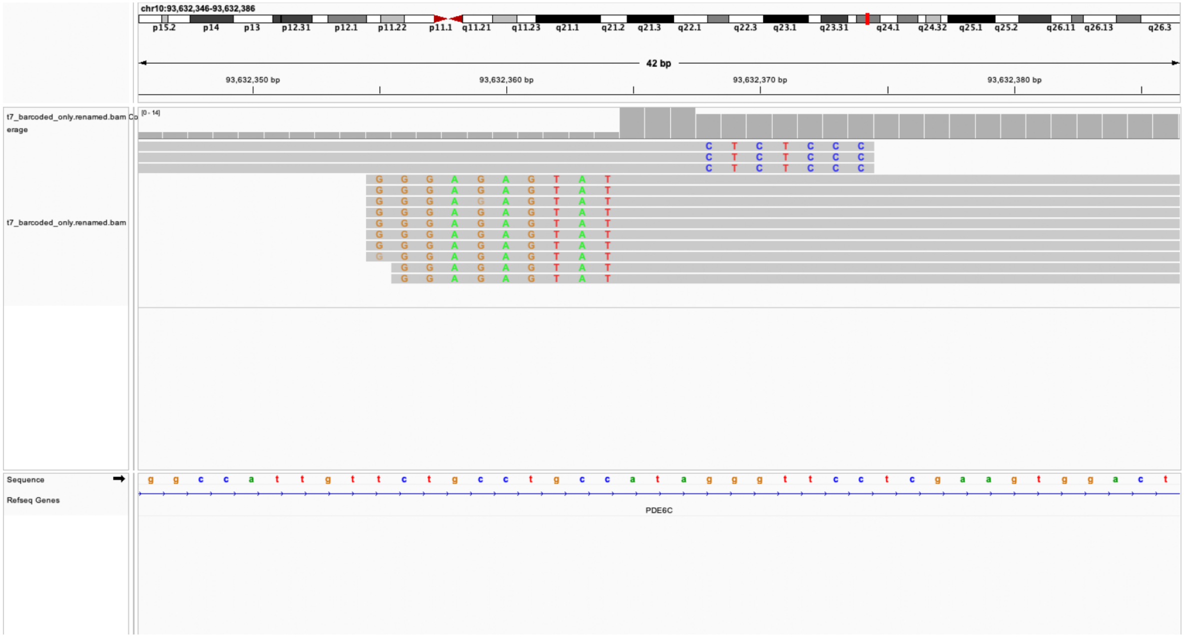

### manual edit call:True loc: chr3;25497711 genes=RARB

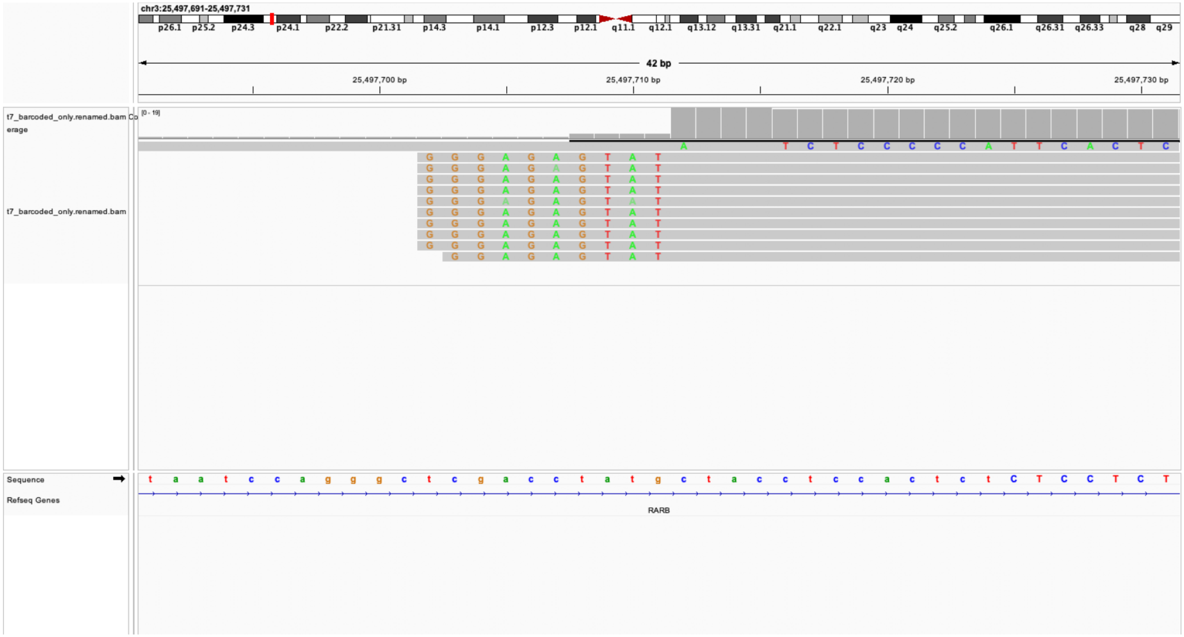

ual edit call:True loc: chr3;194003698 genes=LINC02026;ENSG0000023

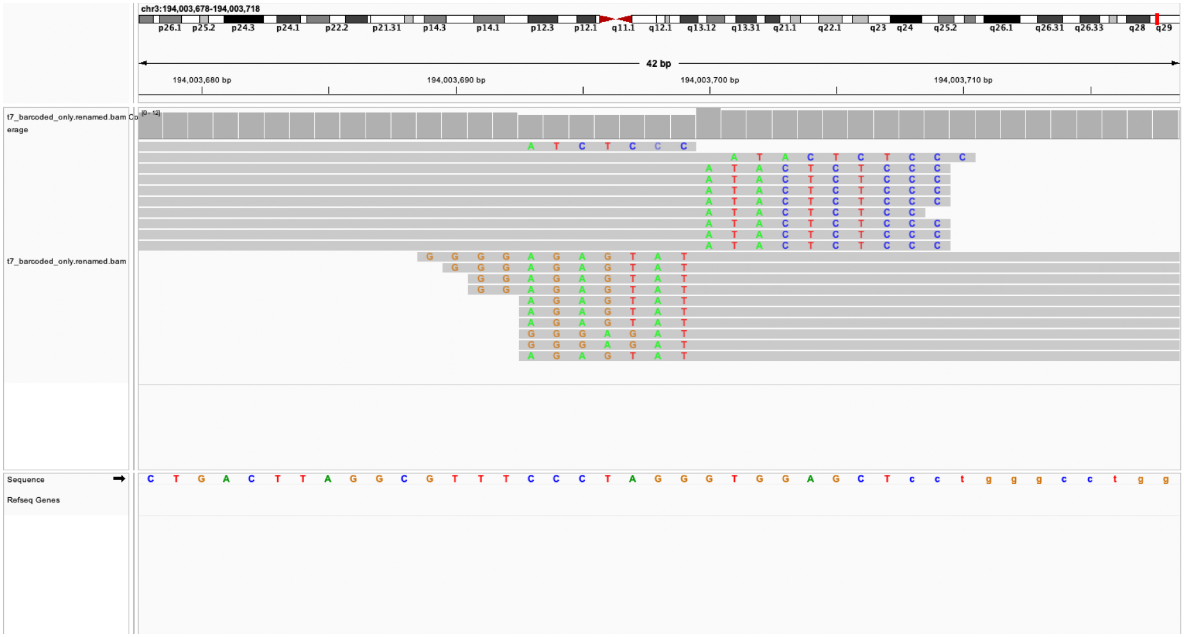

### manual edit call:True loc: chr3;31532824 genes=STT3B

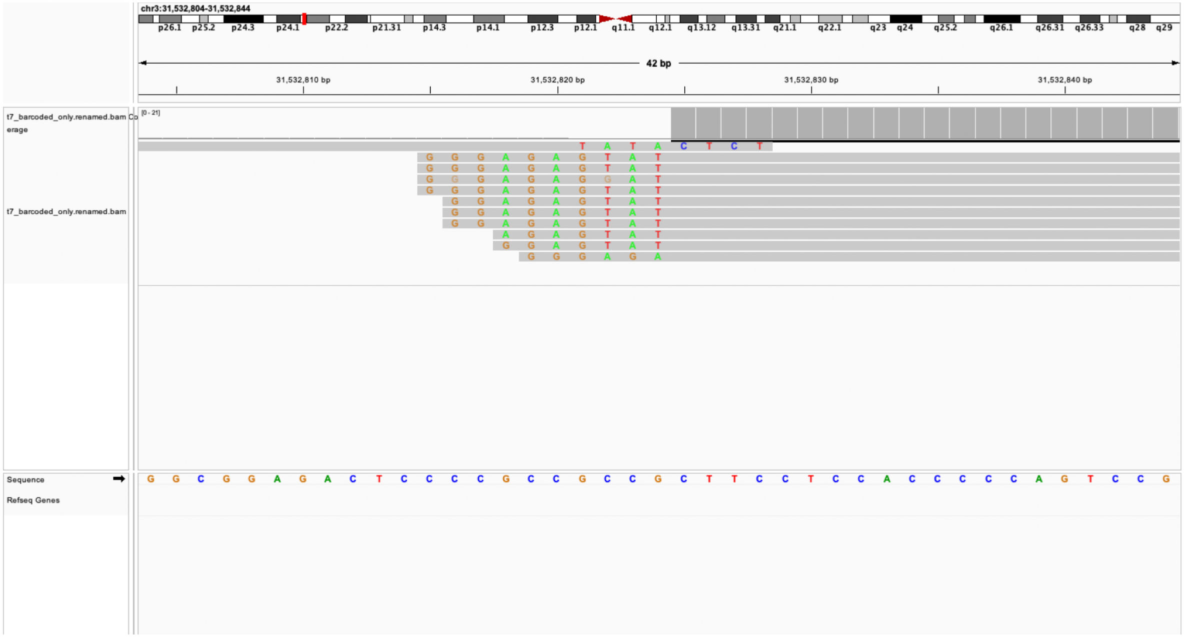

### manual edit call:True loc: chr4;79849103 genes=PCAT4

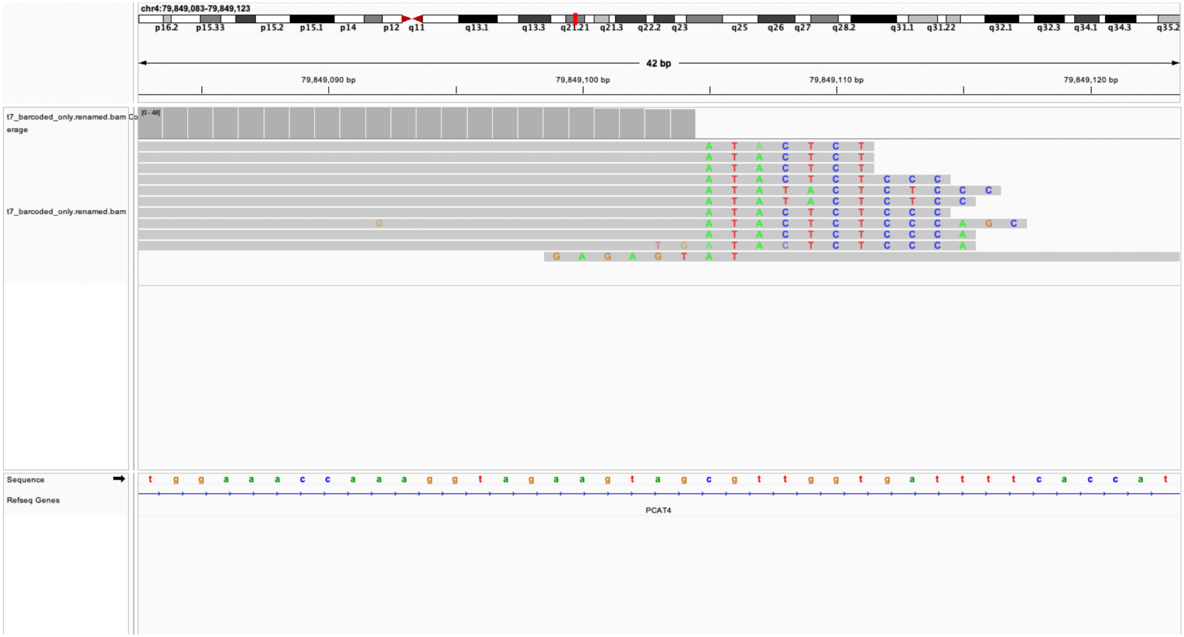

### manual edit call:True loc: chr4;88368039 genes=nan

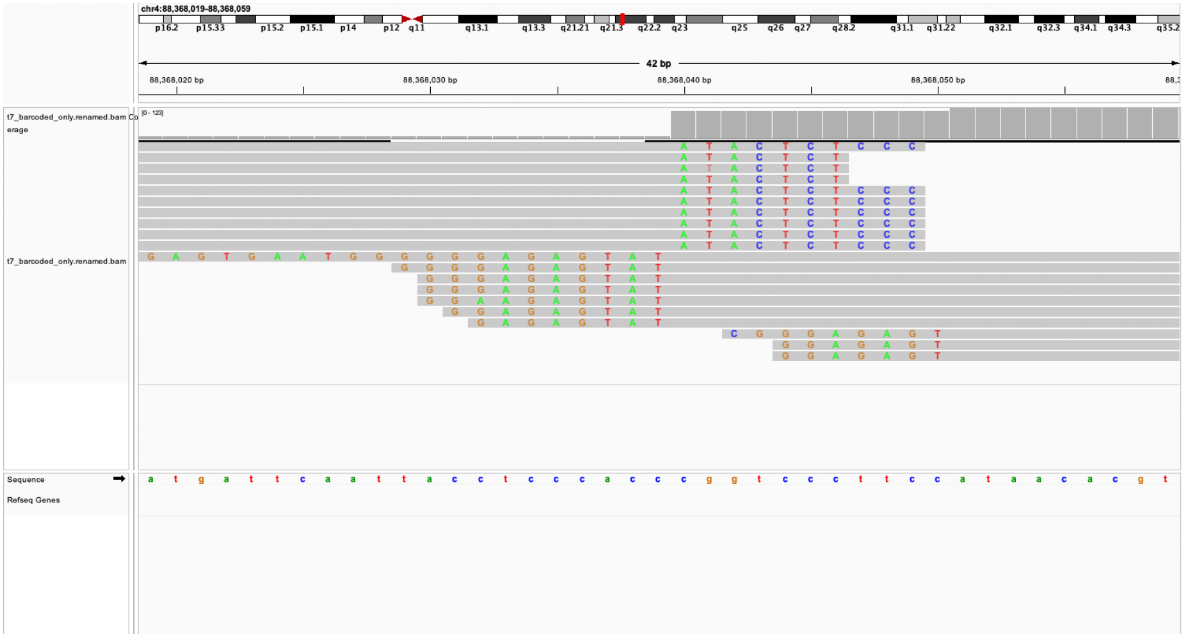

### manual edit call:True loc: chr6;57807628 genes=nan

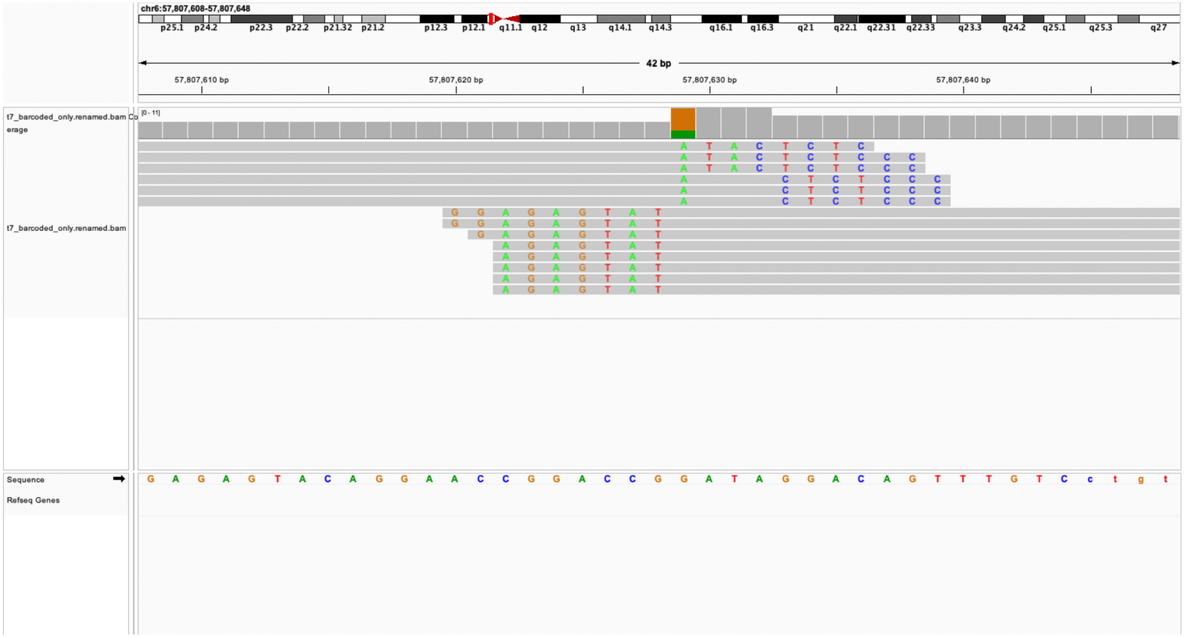

### manual edit call:True loc: chr1;108747053 genes=STXBP3

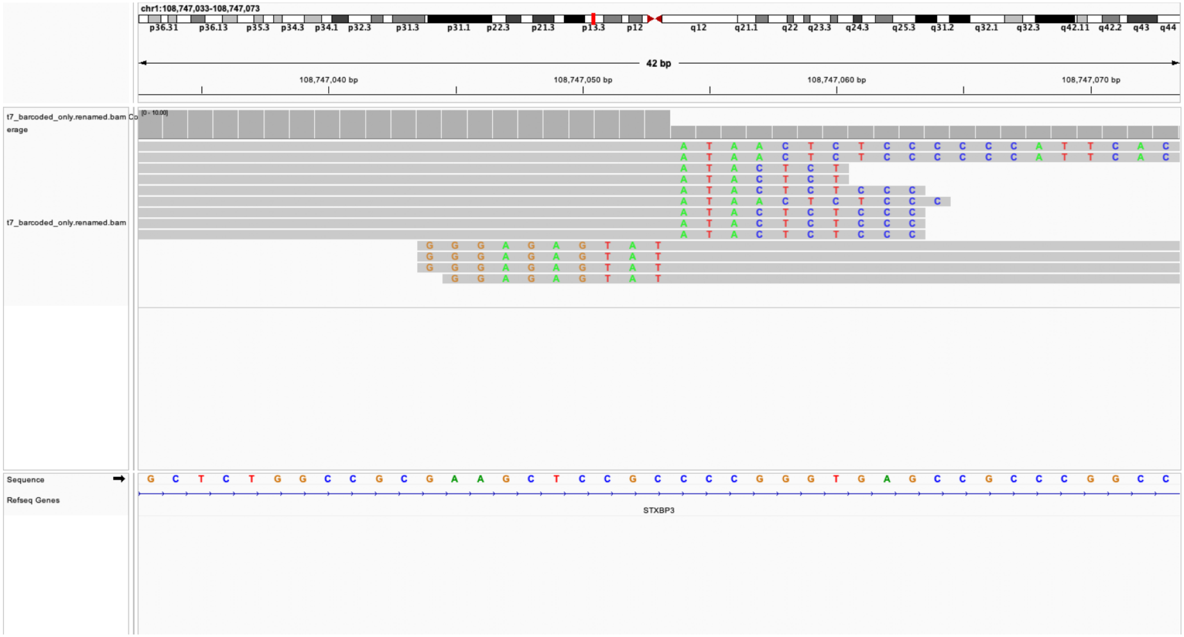

manual edit call:True loc: chr1;224544957 genes=CNIH3

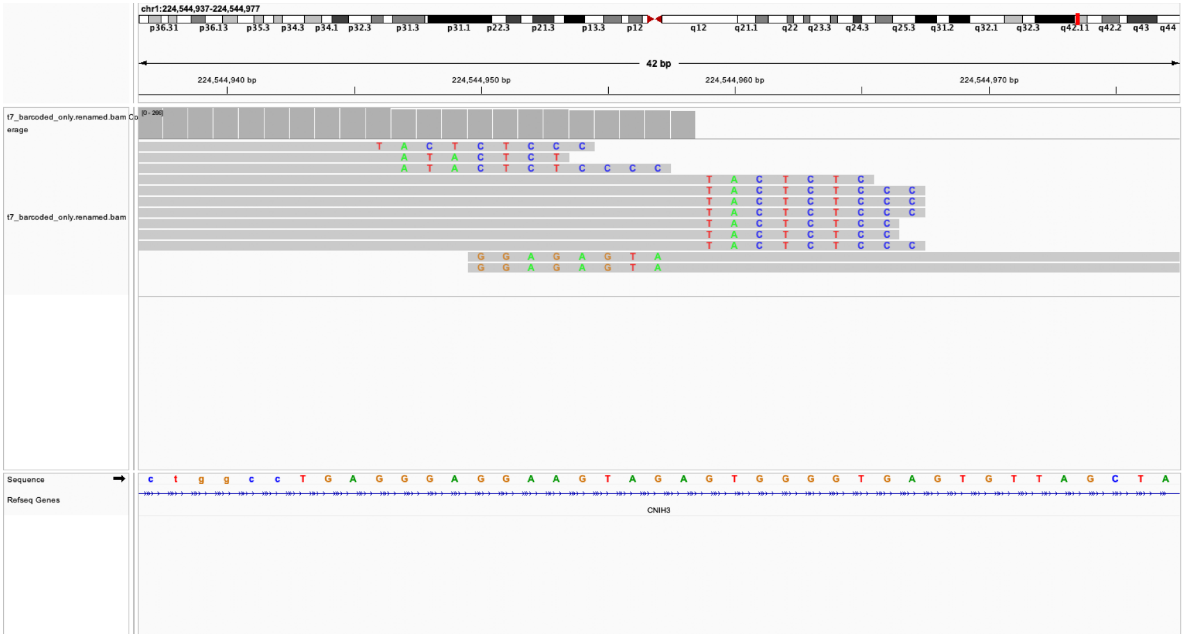

### manual edit call:True loc: chr11;70522075 genes=SHANK2

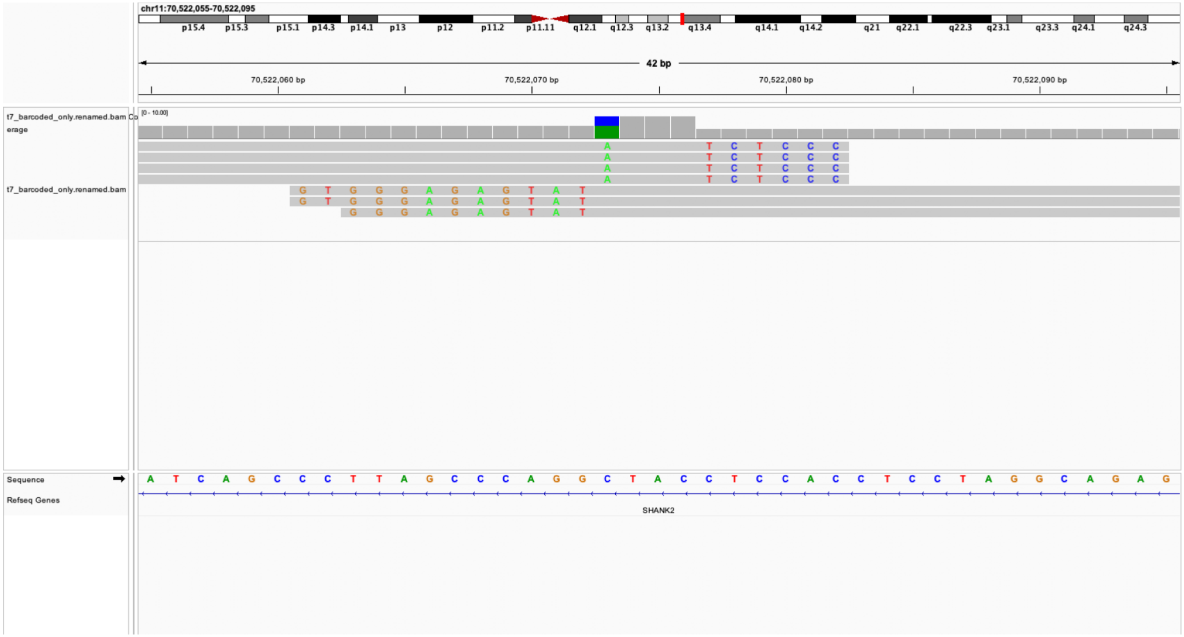

manual edit call:True loc: chr16;81393308 genes=nan

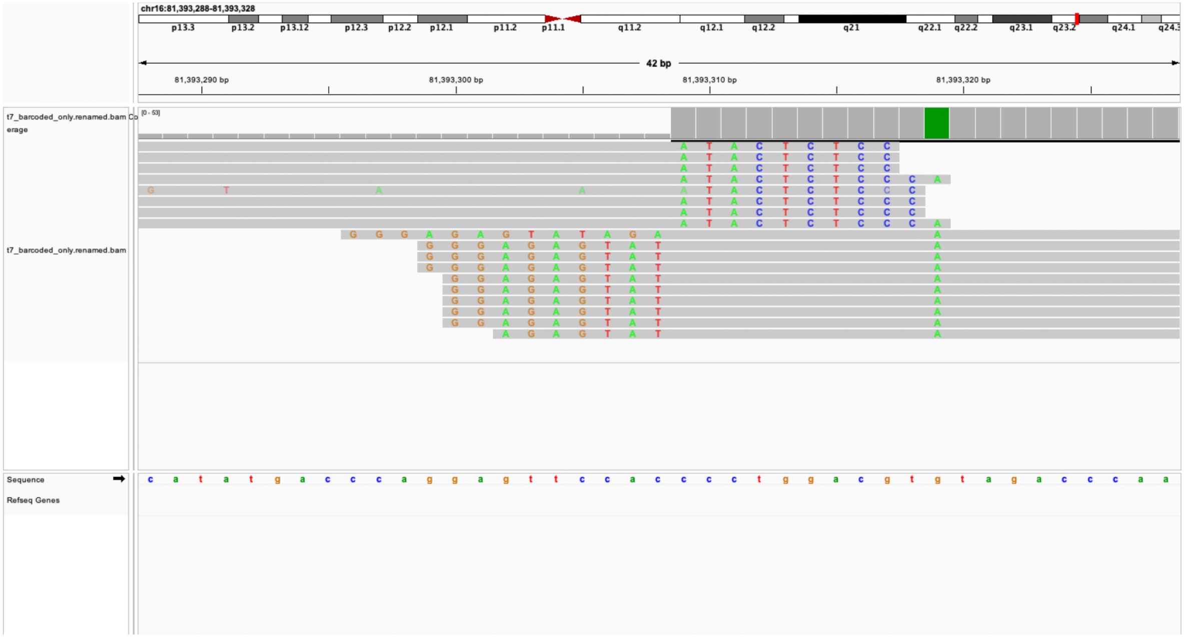

### manual edit call:True loc: chr19;10293417 genes=ICAM5

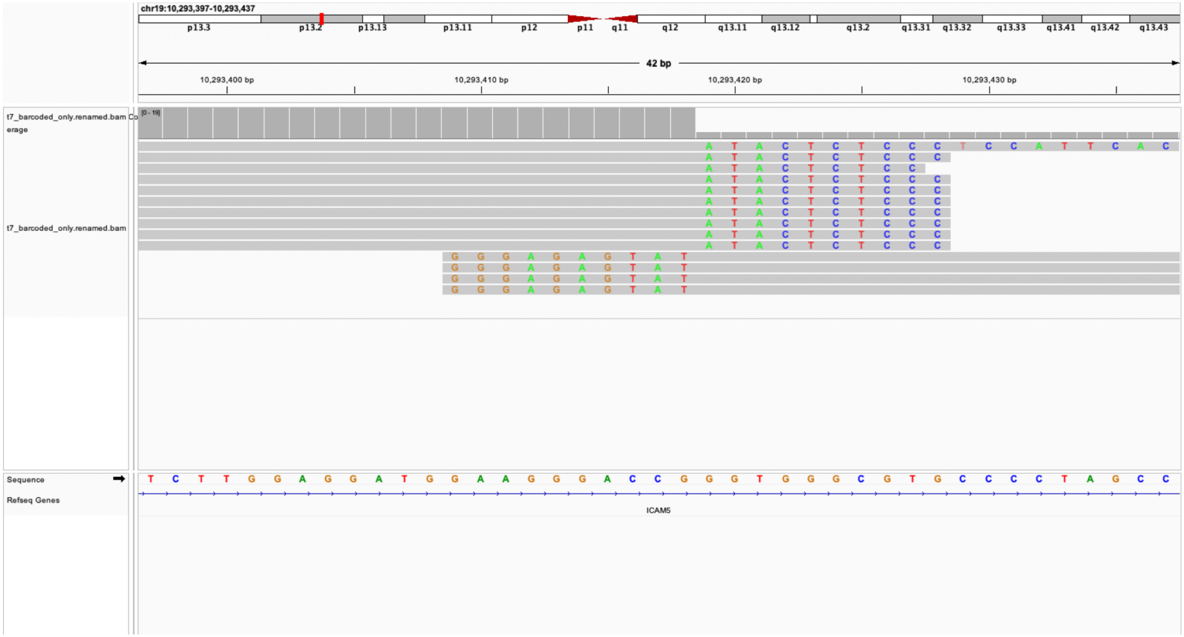

### manual edit call:True loc: chr14;104729499 genes=ADSS1

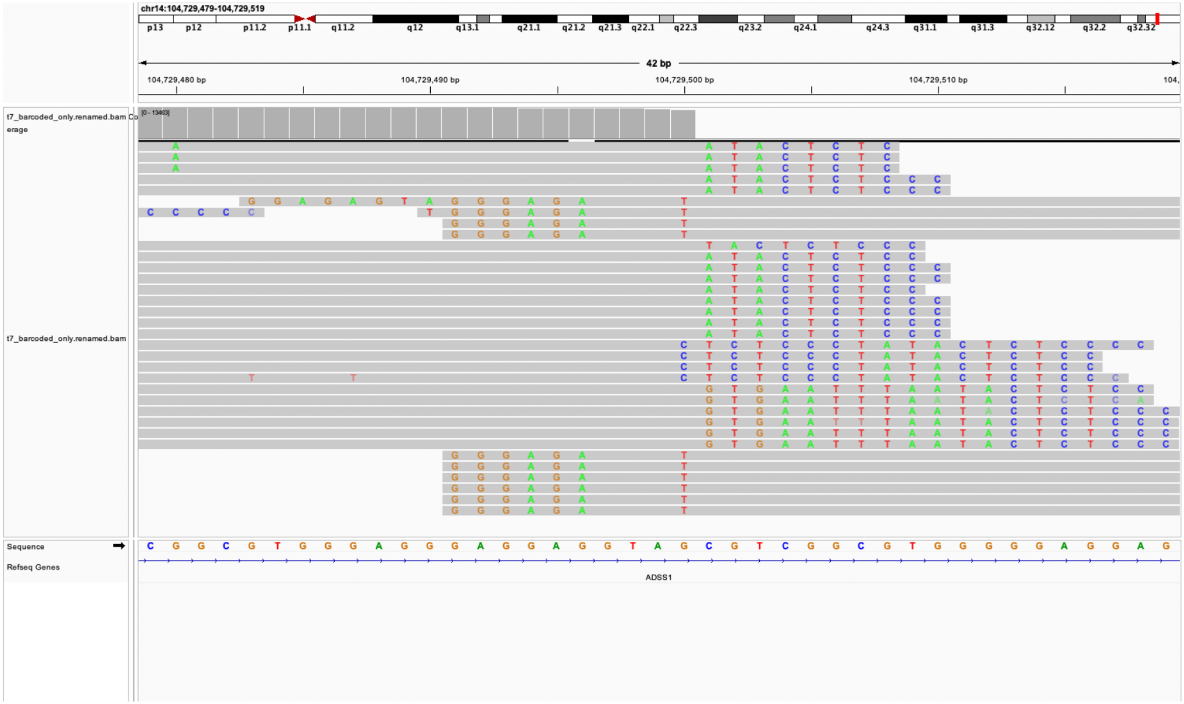

### manual edit call:True loc: chr17;47188686 genes=CDC27

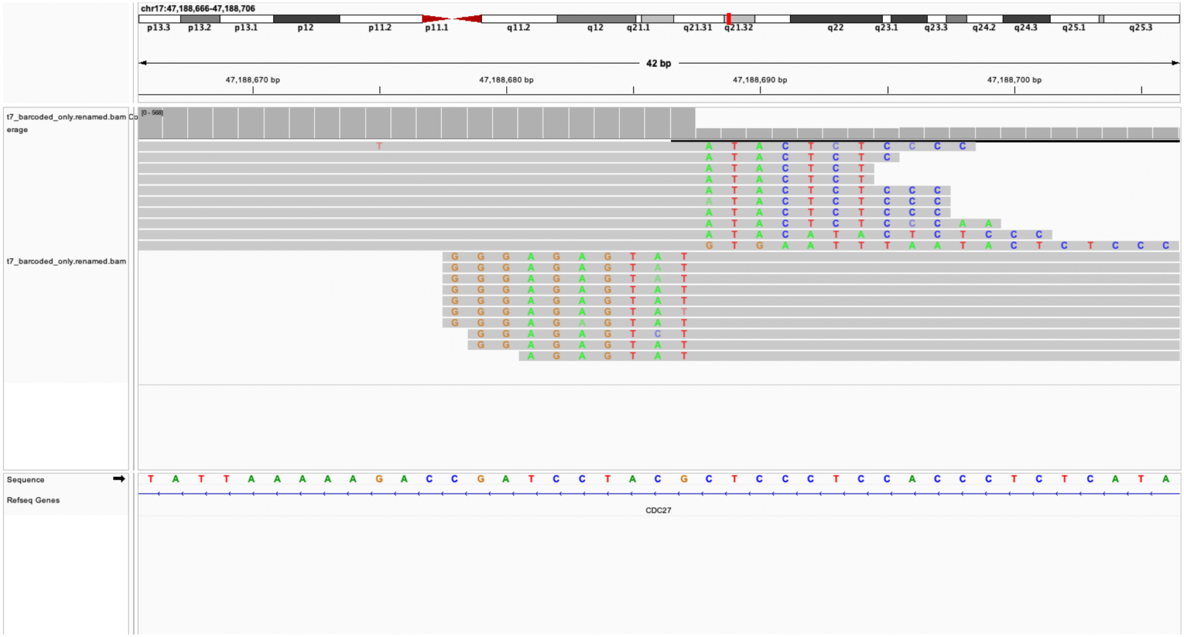

### manual edit call:True loc: chr4;59298822 genes=nan

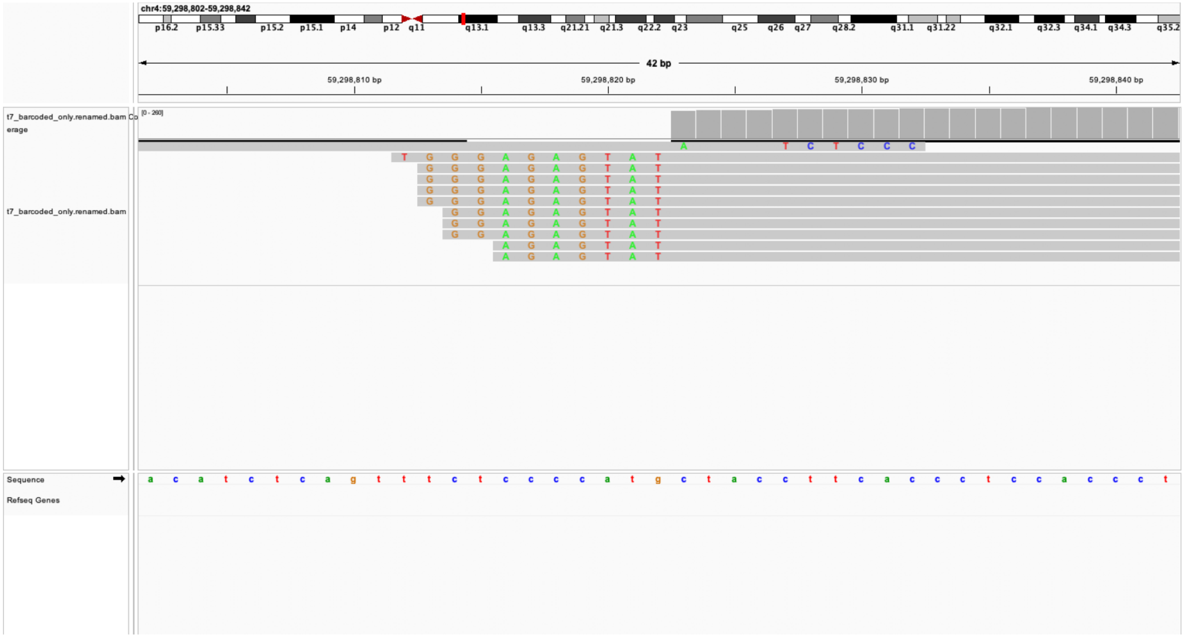

manual edit call:True loc: chr12;110404170 genes=ENSG0000025821[illegible]

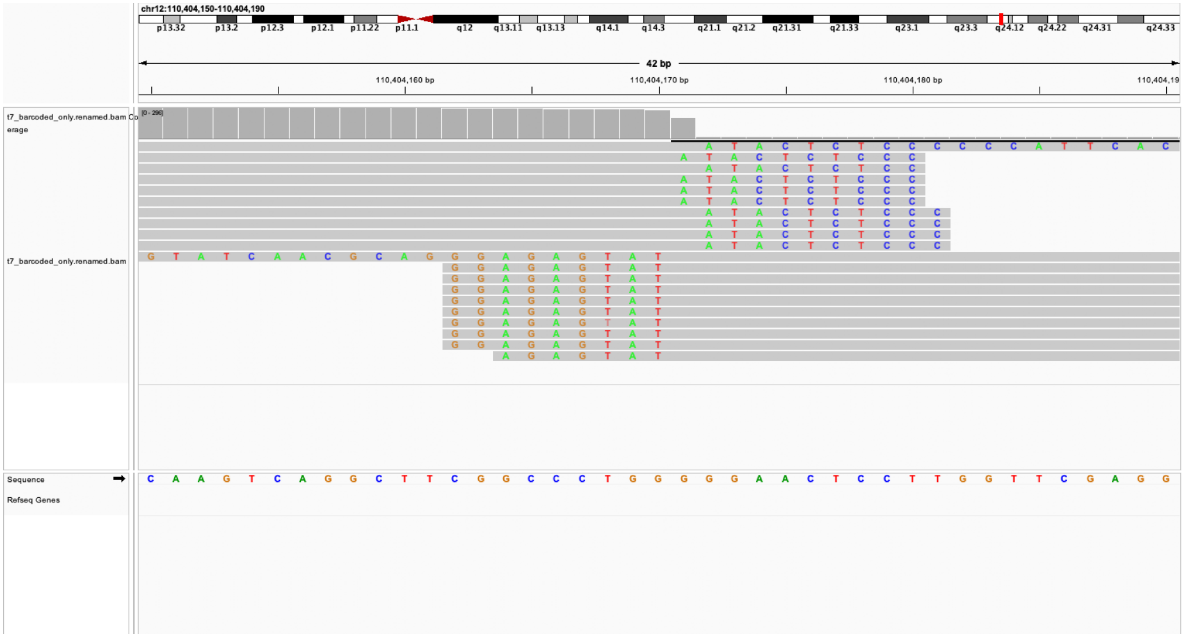

manual edit call:True loc: chr5;148237295 genes=FBXO38-DT;MARCOL

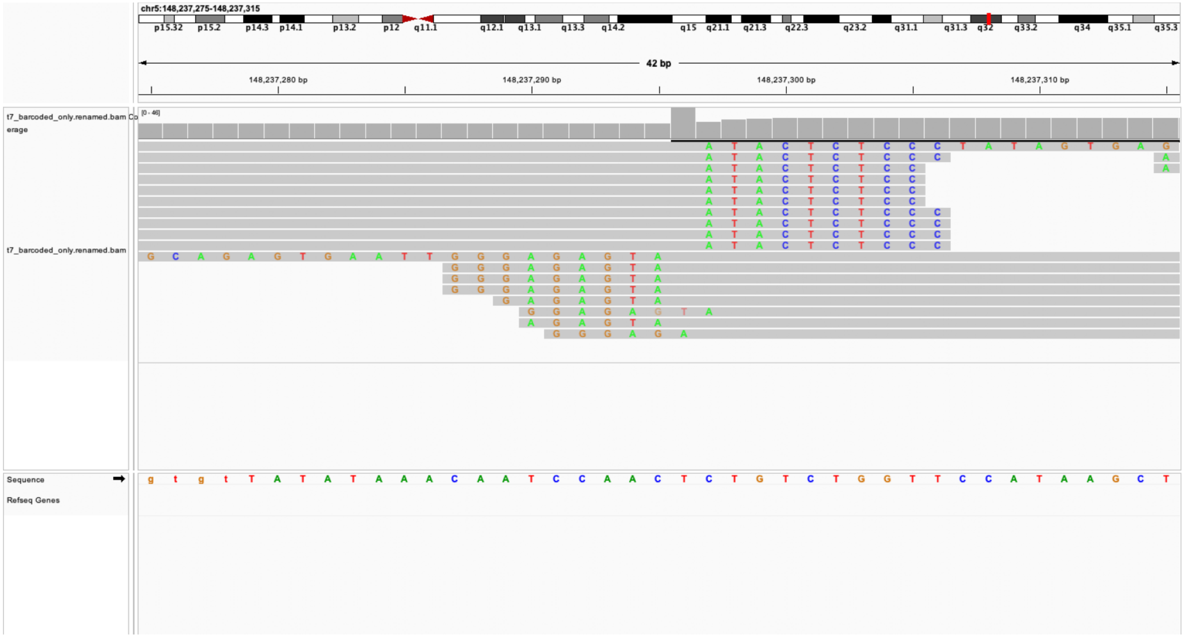

### manual edit call:True loc: chr13;110873574 genes=nan

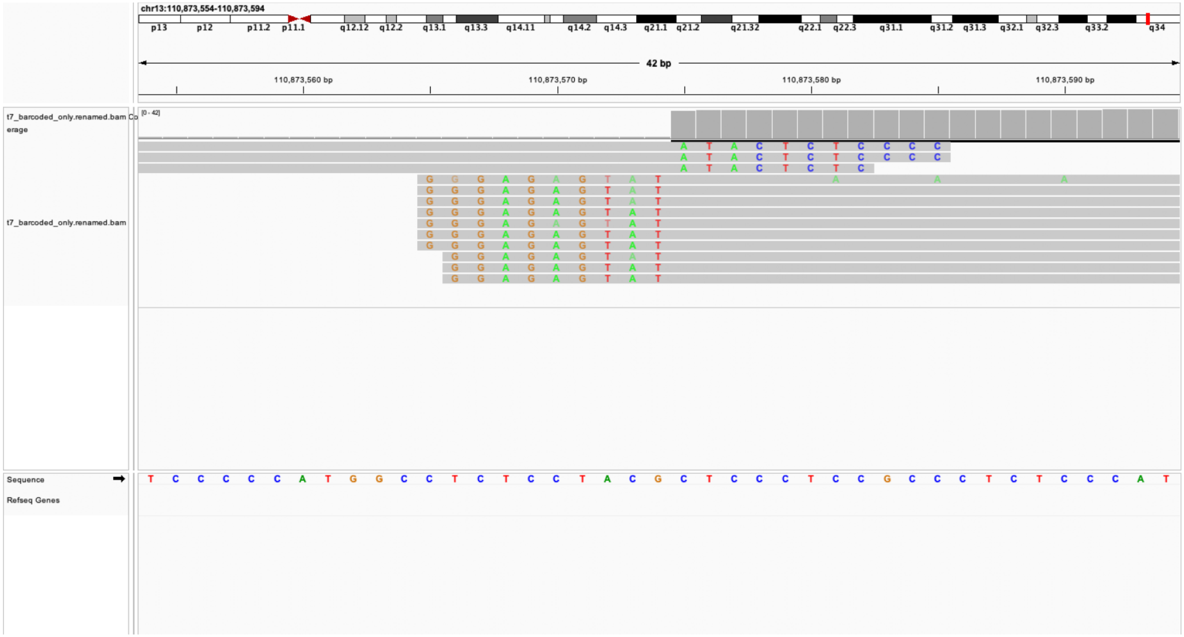

### manual edit call:True loc: chr17;3029640 genes=RAP1GAP2

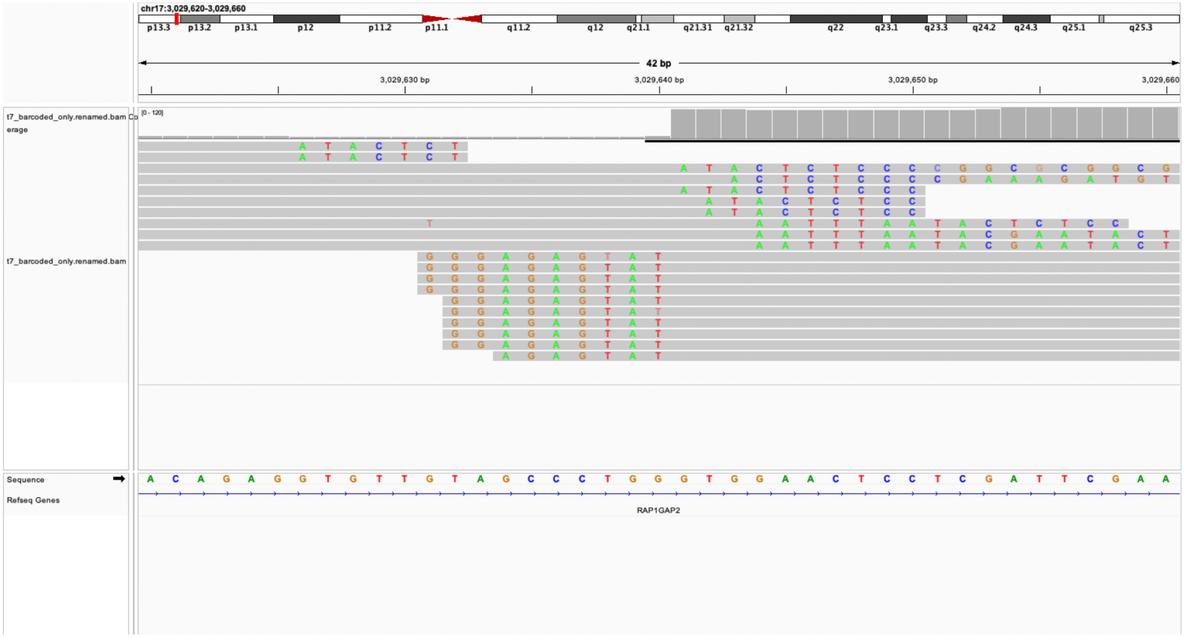

### manual edit call:True loc: chr9;113111494 genes=FAM225B

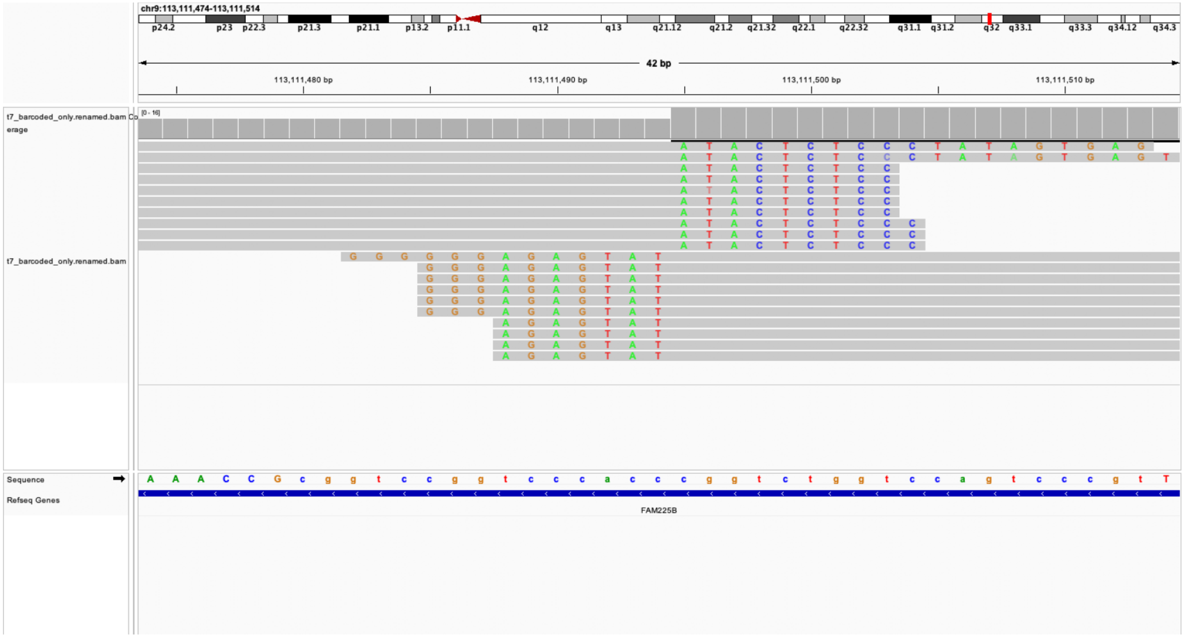

manual edit call:True loc: chr11;68841673 genes=CPT1A

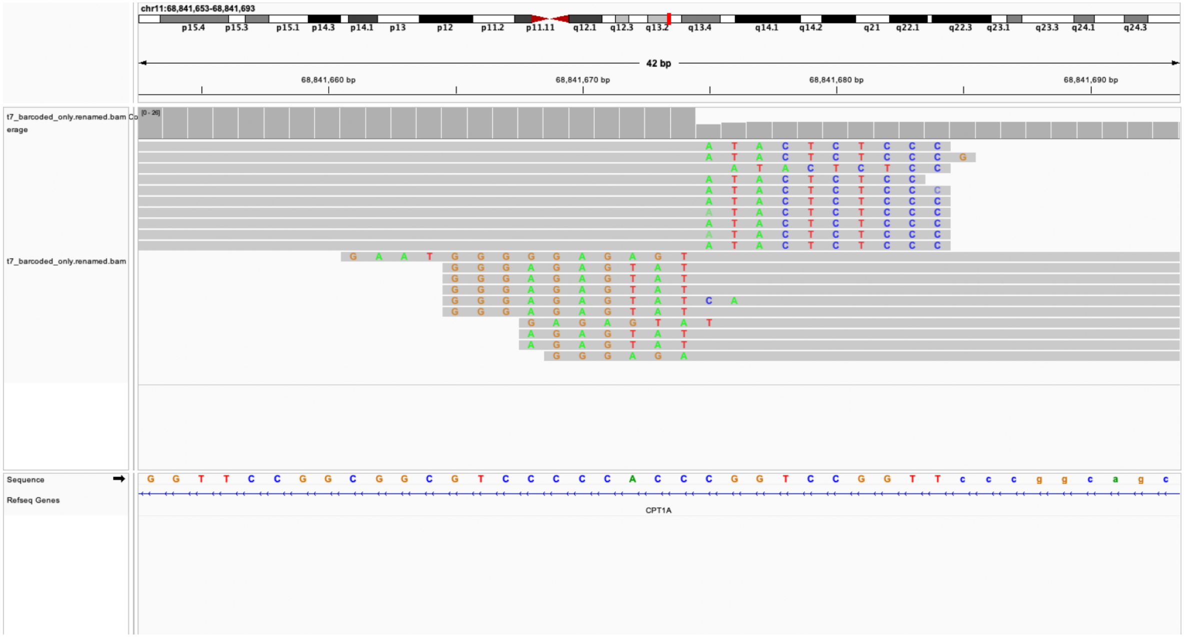

edit call:False loc: chr6;67885109 genes=ENSG00000227706;ENSG0000

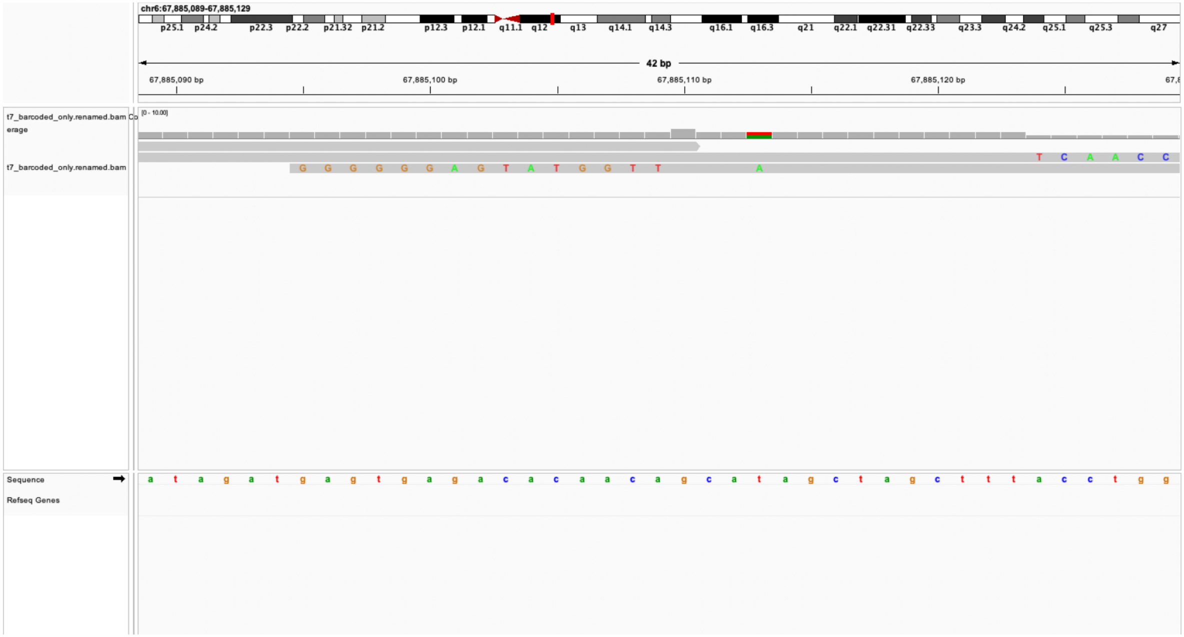

### manual edit call:True loc: chr13;113608067 genes=TFDP1

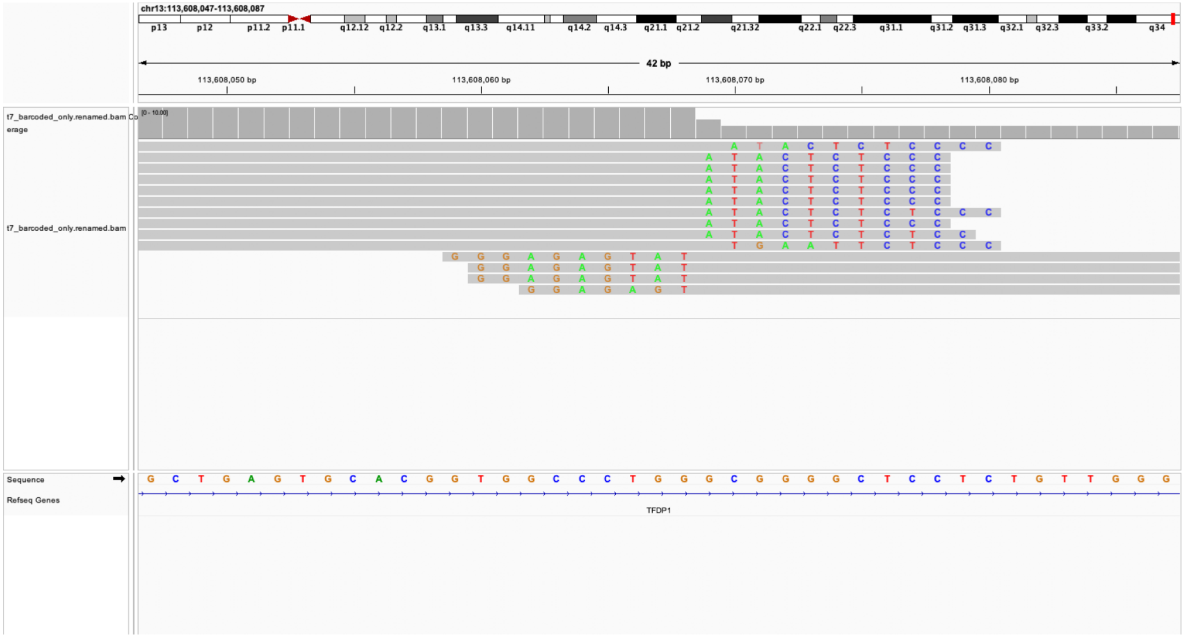

nanual edit call:False loc: chr6;16262954 genes=GMPR;ENSG0000282C

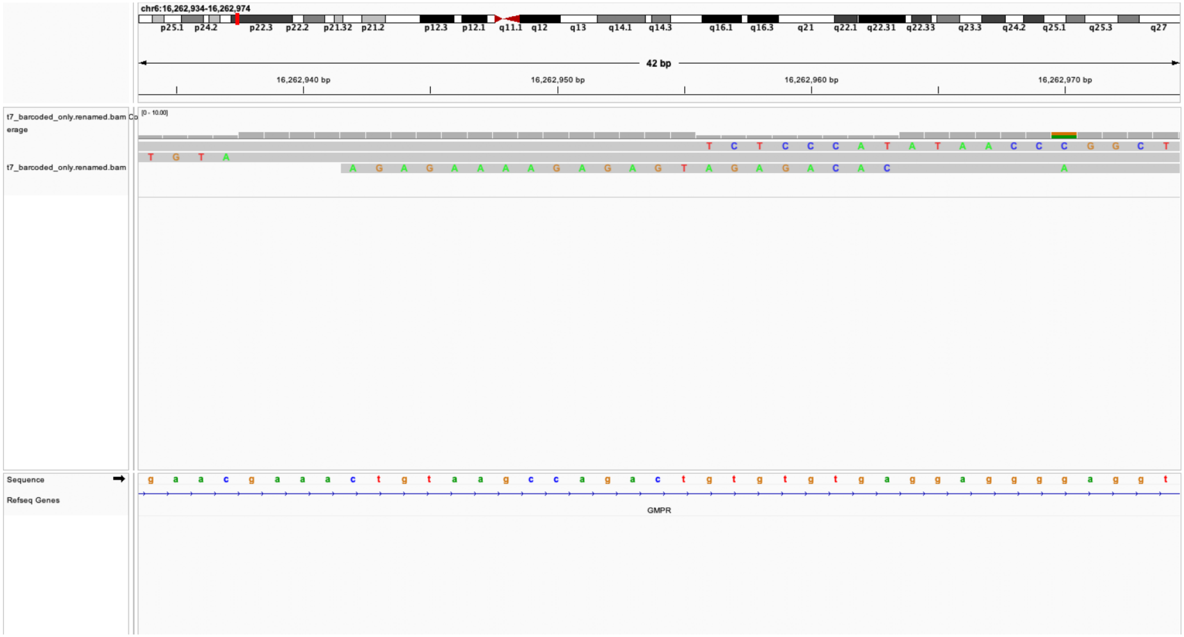

manual edit call:False loc: chr6;140374836 genes=ENSG00000288714

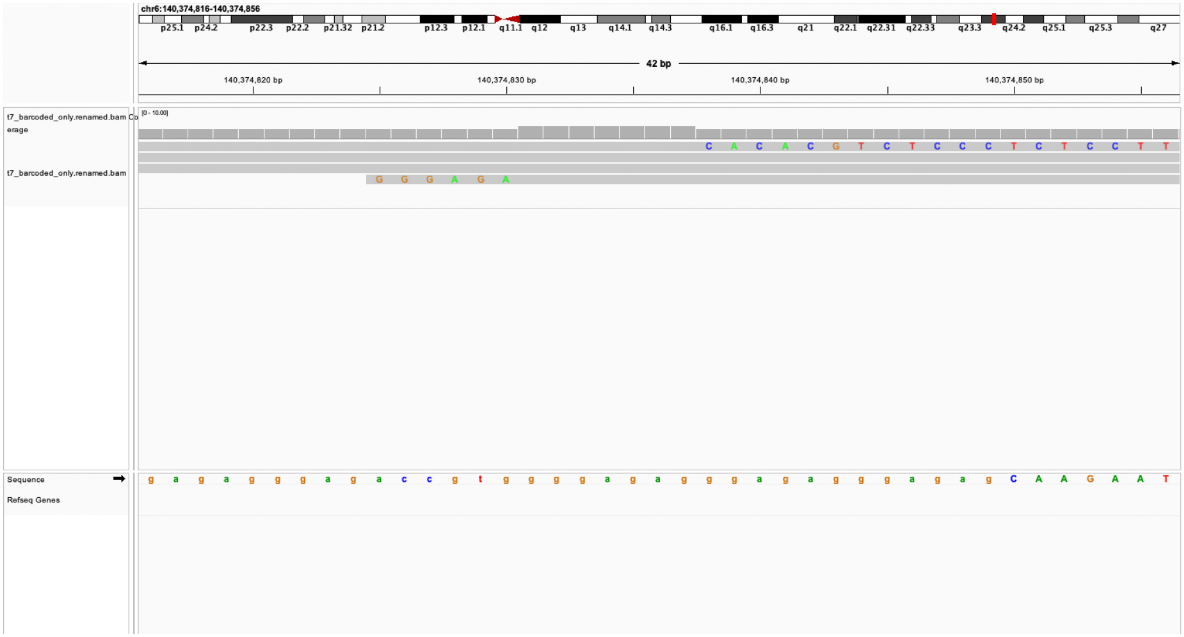

### manual edit call:False loc: chr15;34382827 genes=GOLGA8A

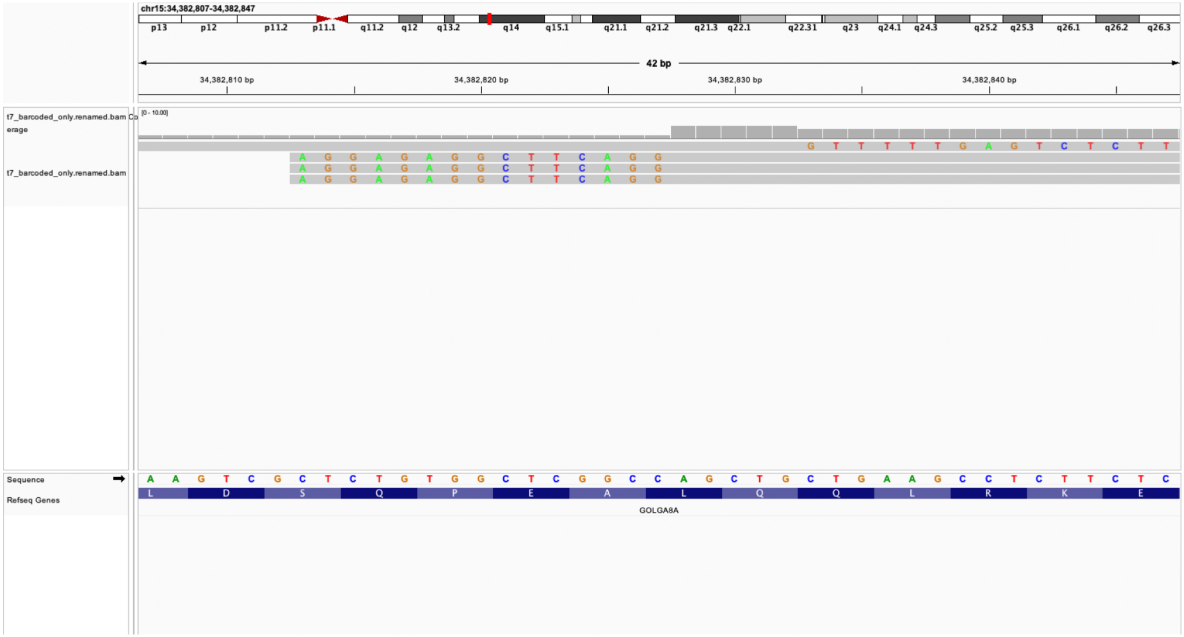

manual edit call:True loc: chr14;71885108 genes=ENSG00000266869

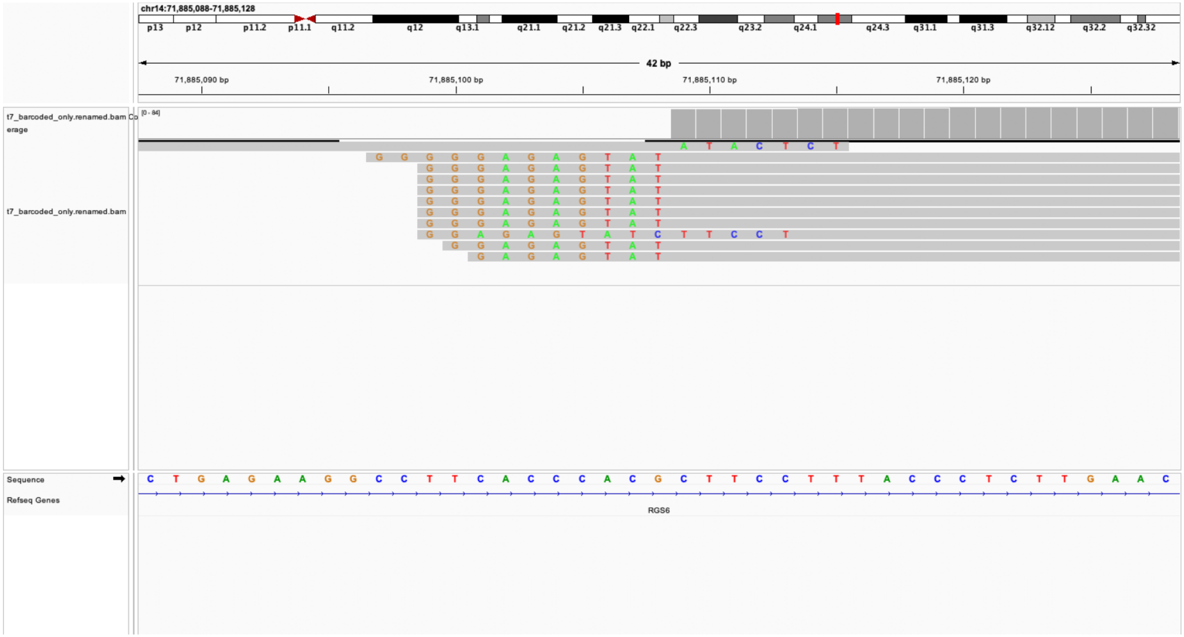

### manual edit call:False loc: chr8;47862492 genes=PRKDC

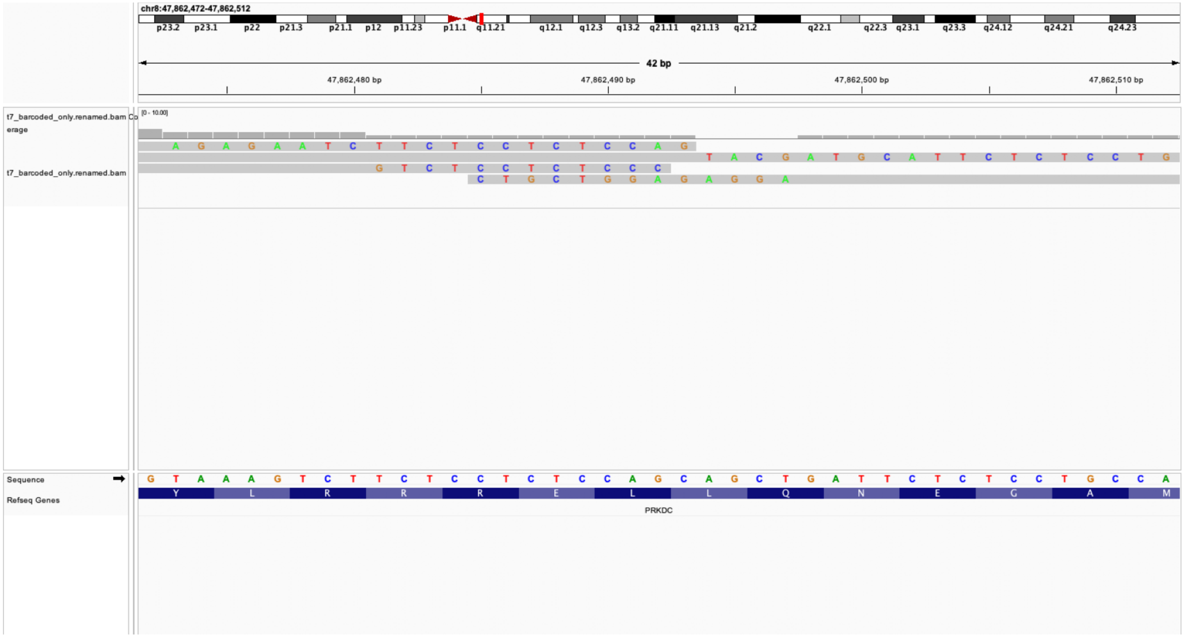

### manual edit call:True loc: chr4;7542884 genes=SORCS2

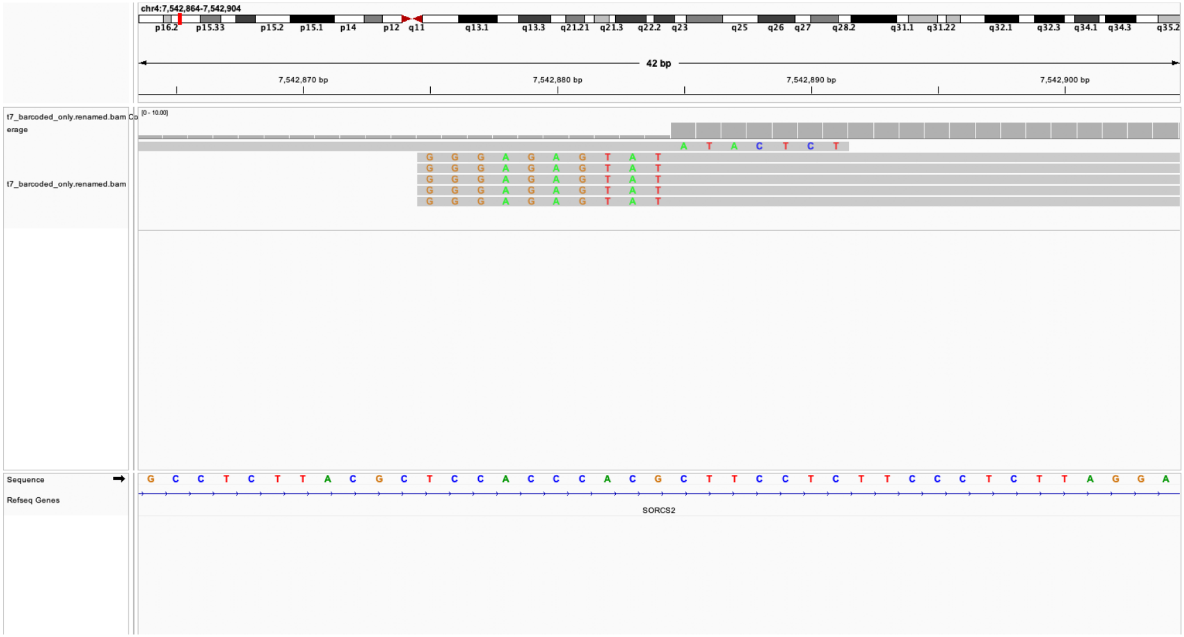

manual edit call:False loc: chr19;975594 genes=ARID3A

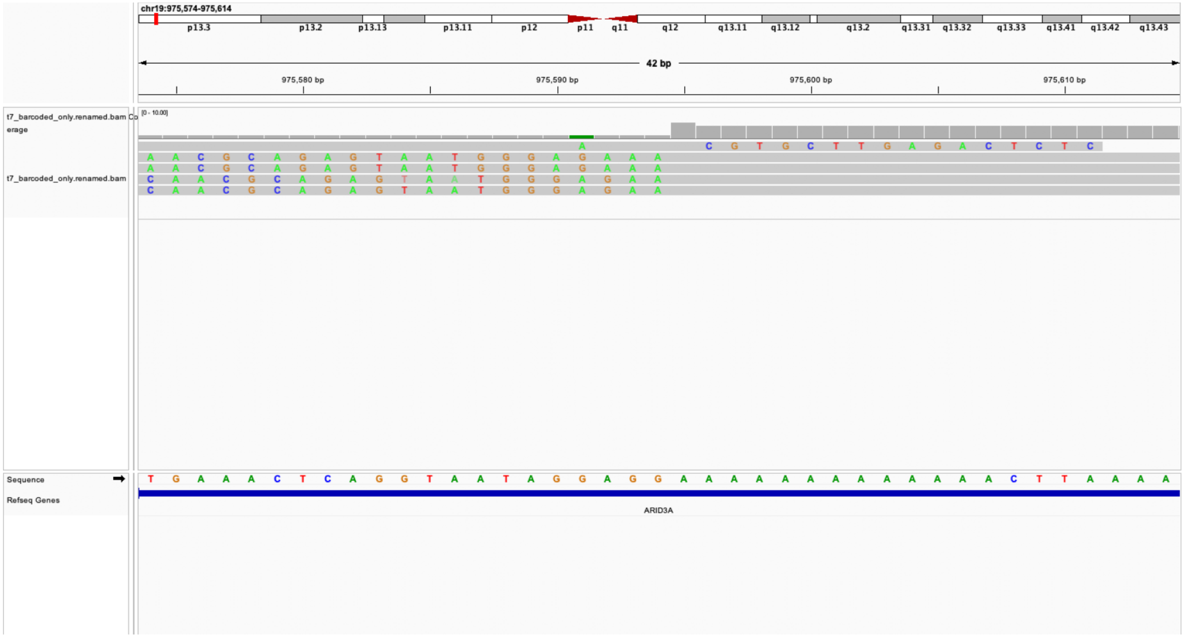

### manual edit call:False loc: chr6;33275863 genes=RPS18

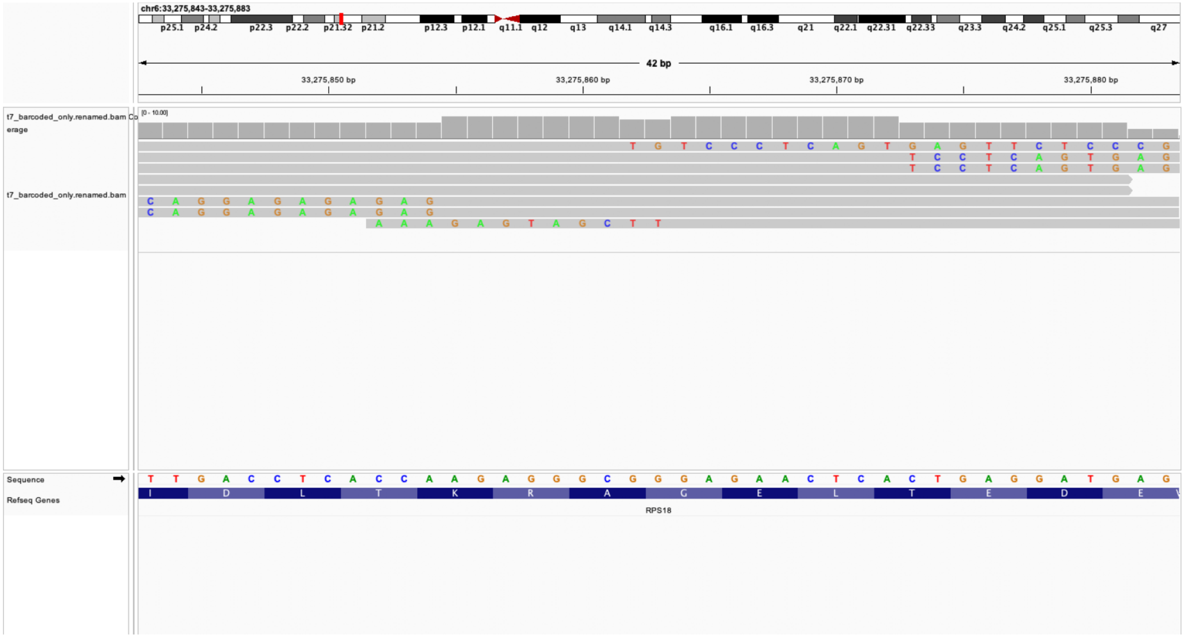

### manual edit call:False loc: chr13;18958292 genes=PHF2P2

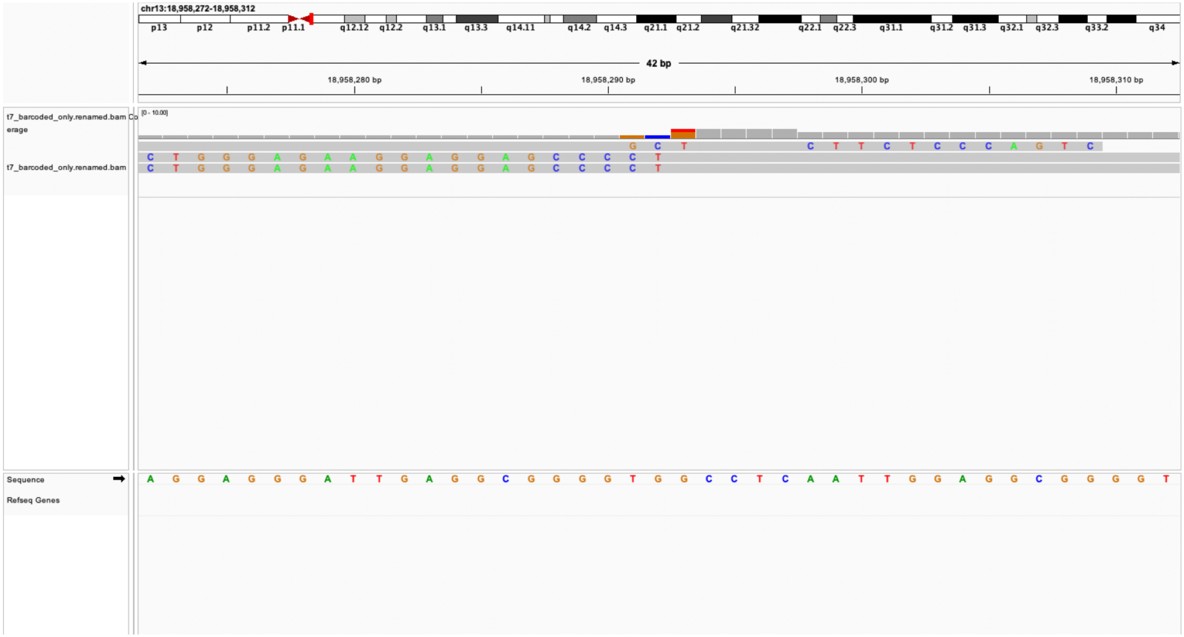

### manual edit call:False loc: chr2;19901462 genes=TTC32

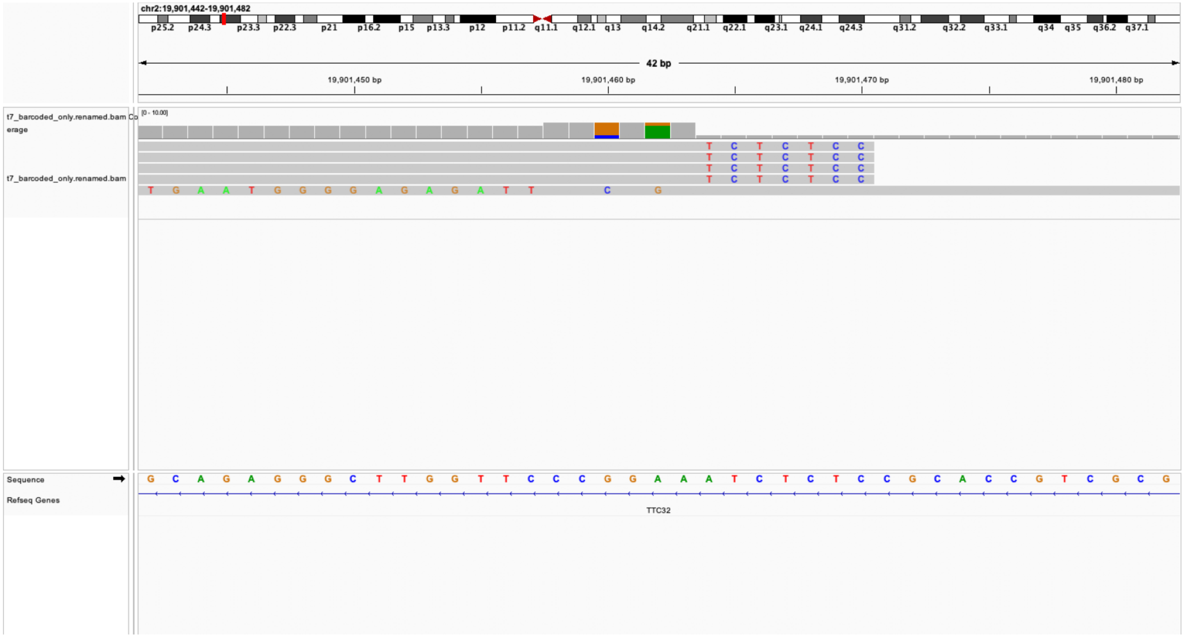

manual edit call:False loc: chr22;19372513 genes=HIRA;C22orf39

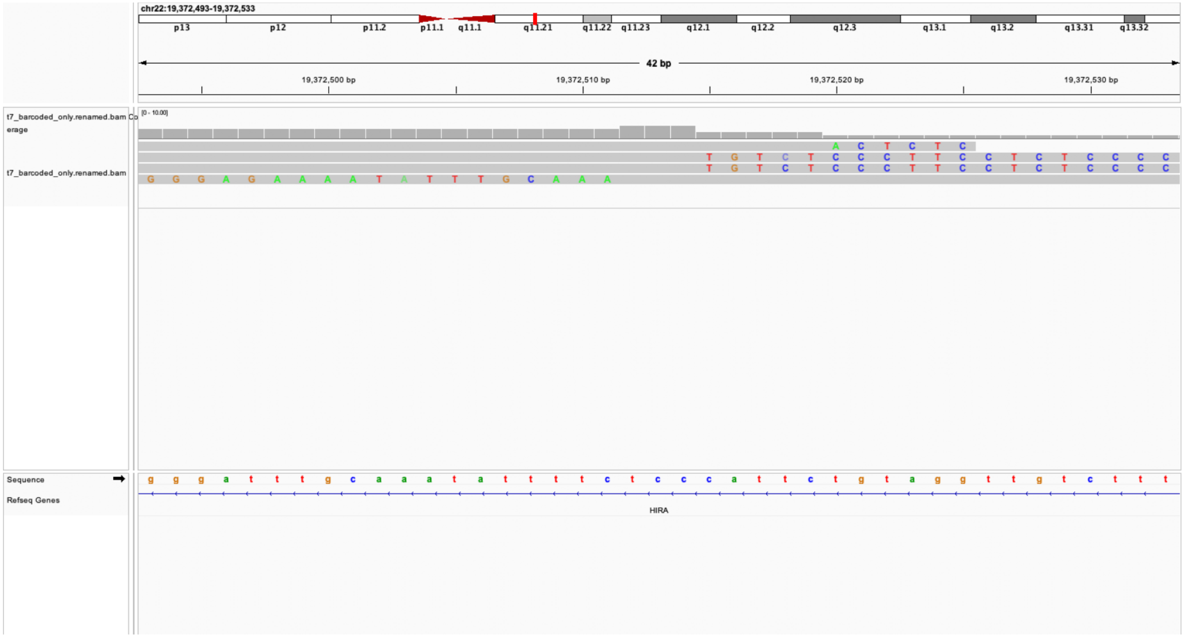

### manual edit call:False loc: chr21;8992513 genes=nan

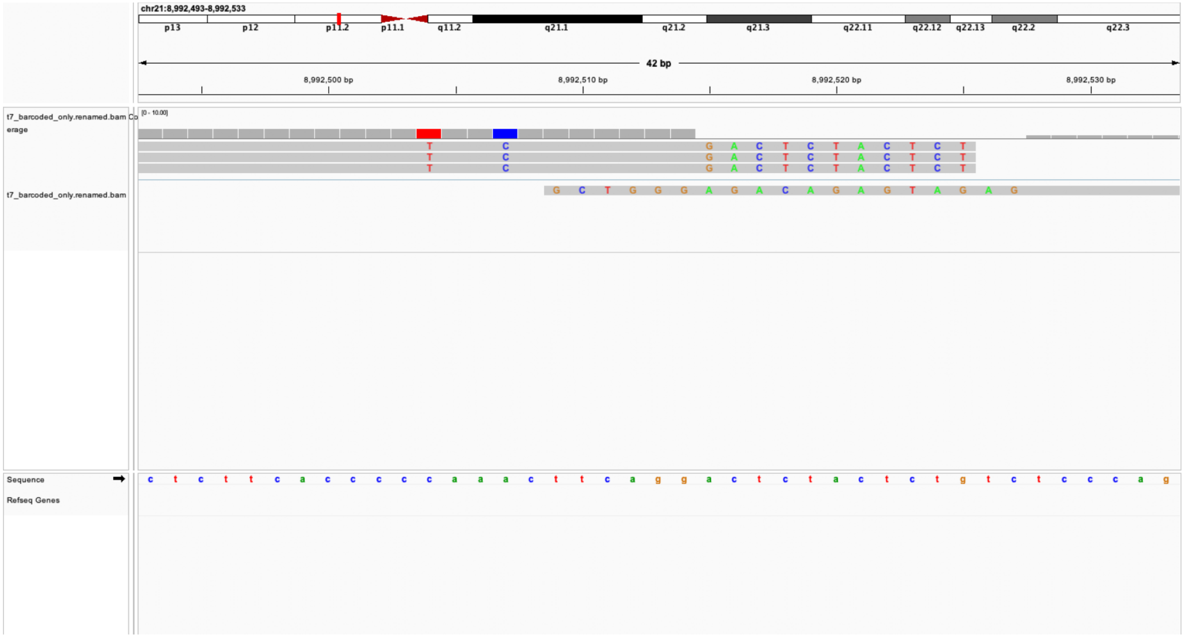

al edit call:False loc: chr21;34086839 genes=MRPS6;SLC5A3;ENSG0000[illegible]

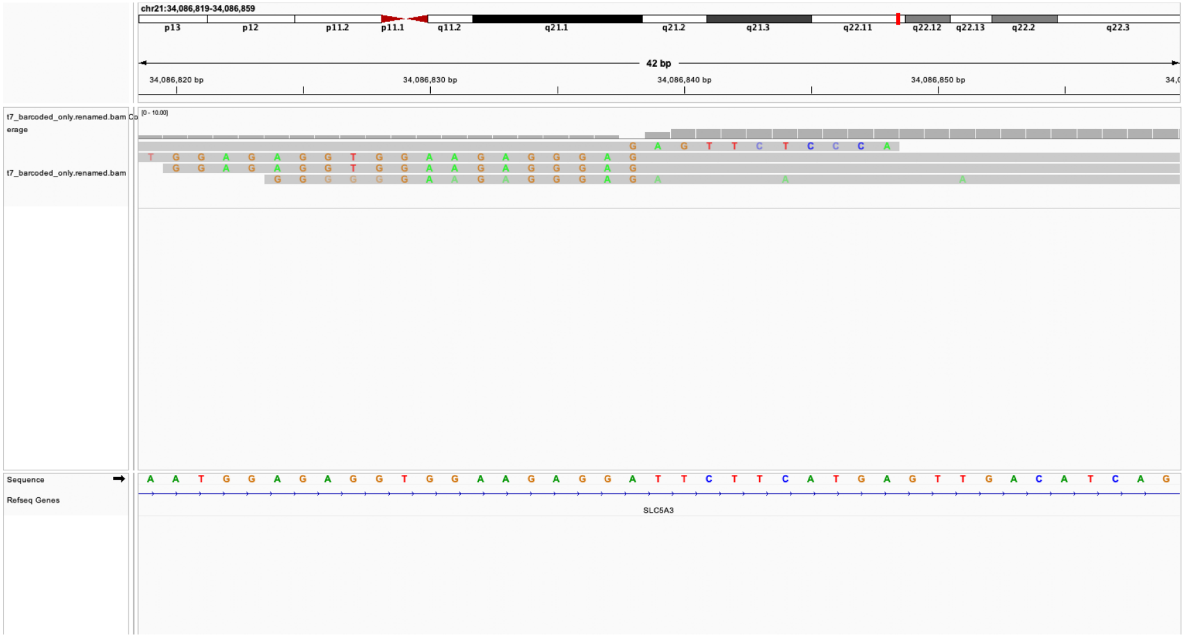

### manual edit call:False loc: chr1;246596227 genes=CNST

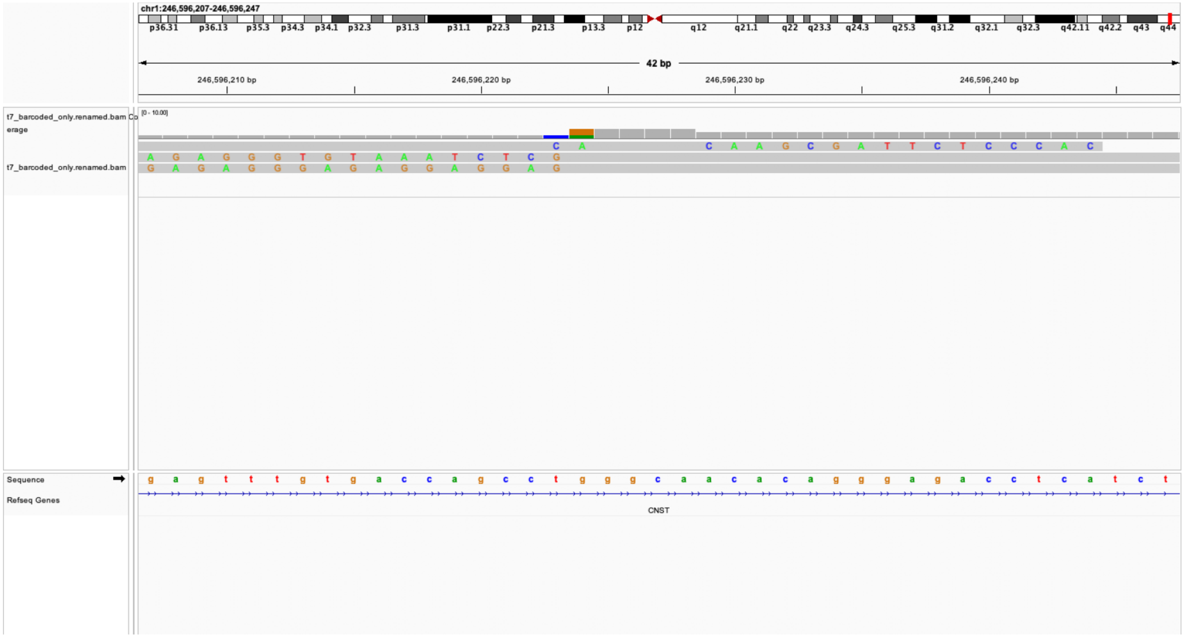

manual edit call:False loc: chr19;3406603 genes=NFIC

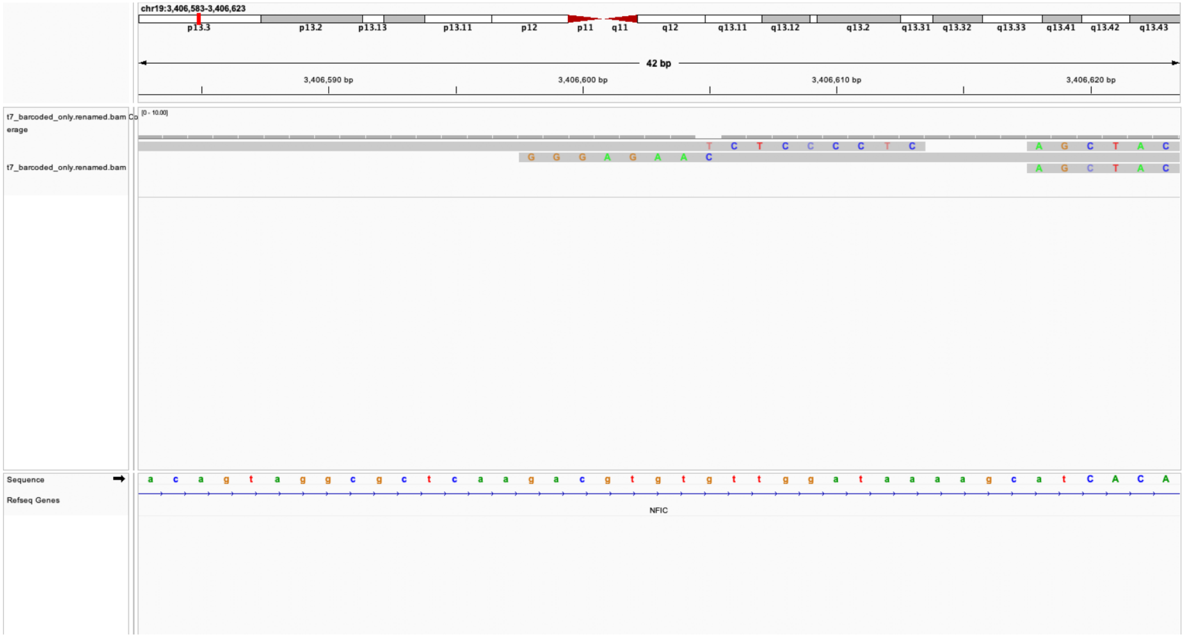

manual edit call:False loc: chr10;73387744 genes=ANXA7

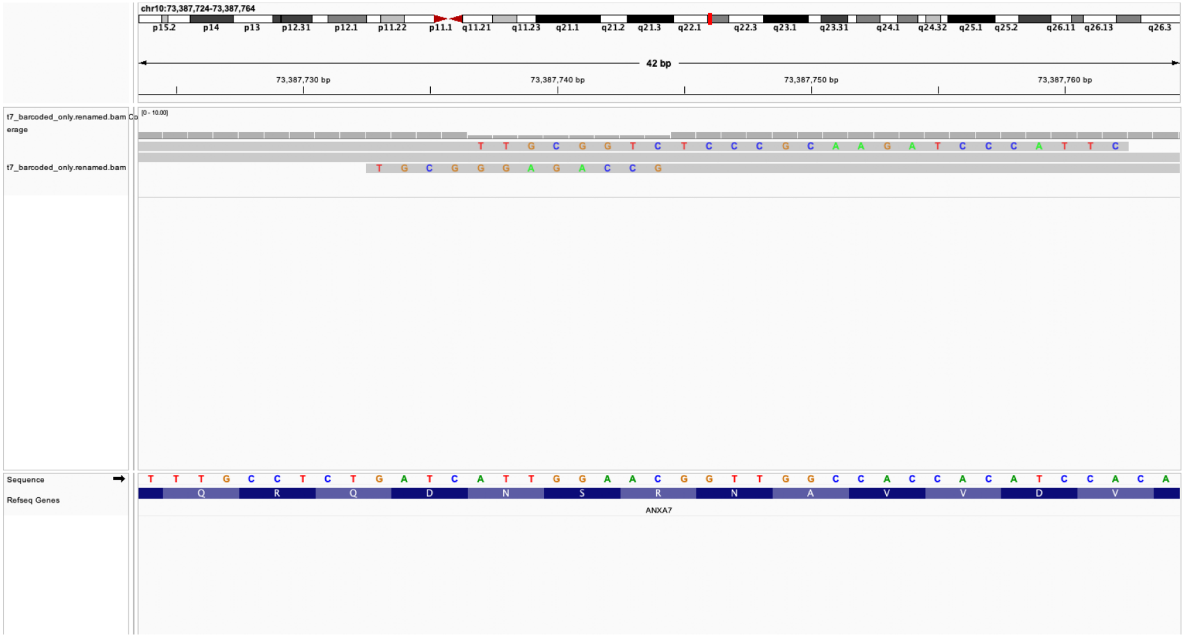

nual edit call:False loc: chr19;38836517 genes=ENSG00000268083;HNI

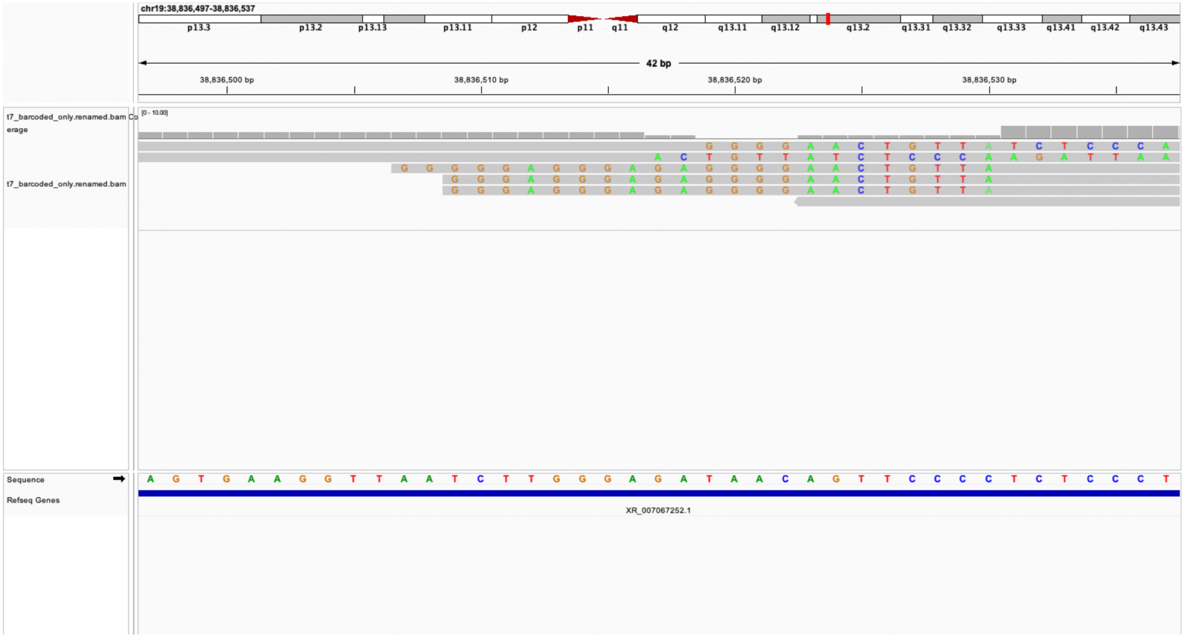

manual edit call:False loc: chr15;42565538 genes=HAUS2

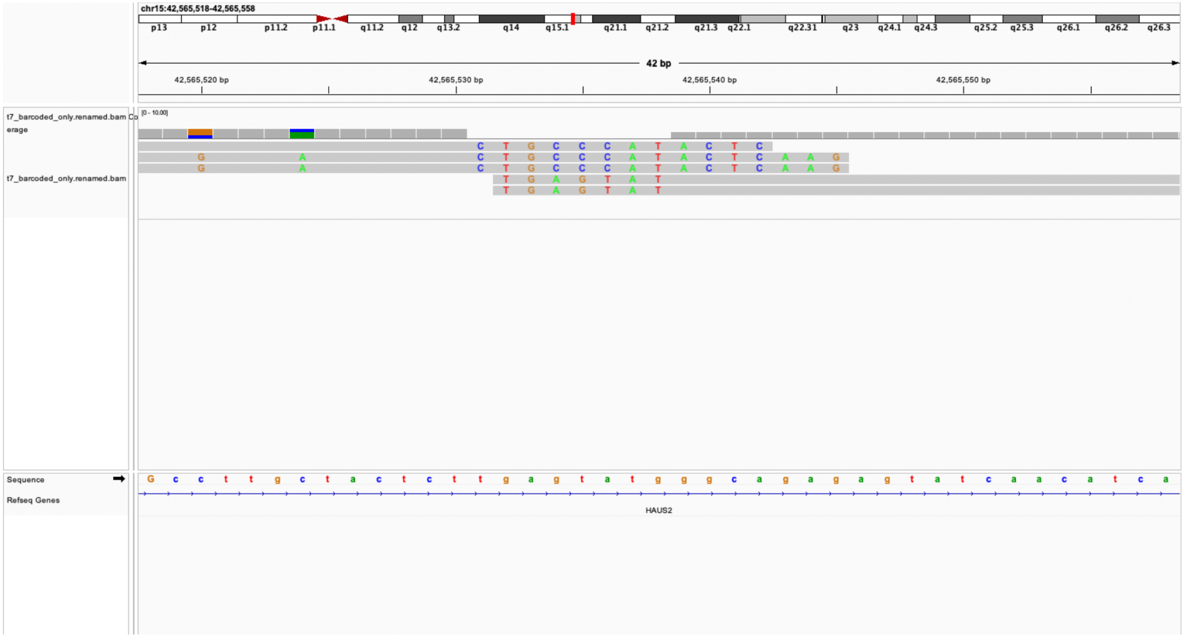

manual edit call:False loc: chr17;64502467 genes=DDX5

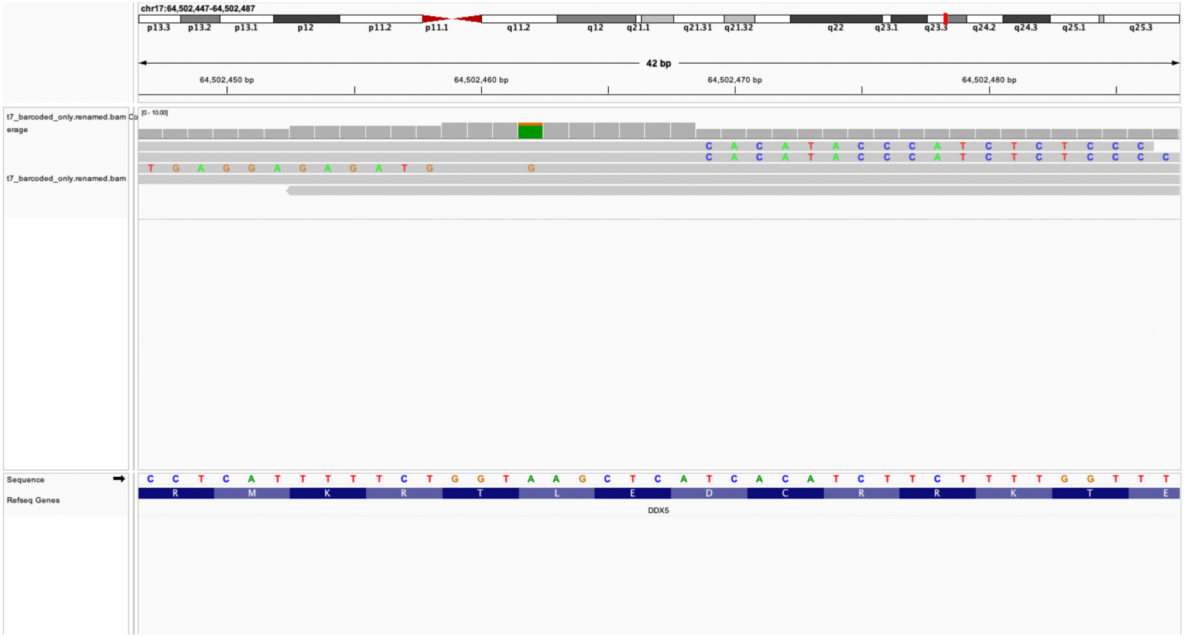

manual edit call:False loc: chr9;97998566 genes=ANP32B

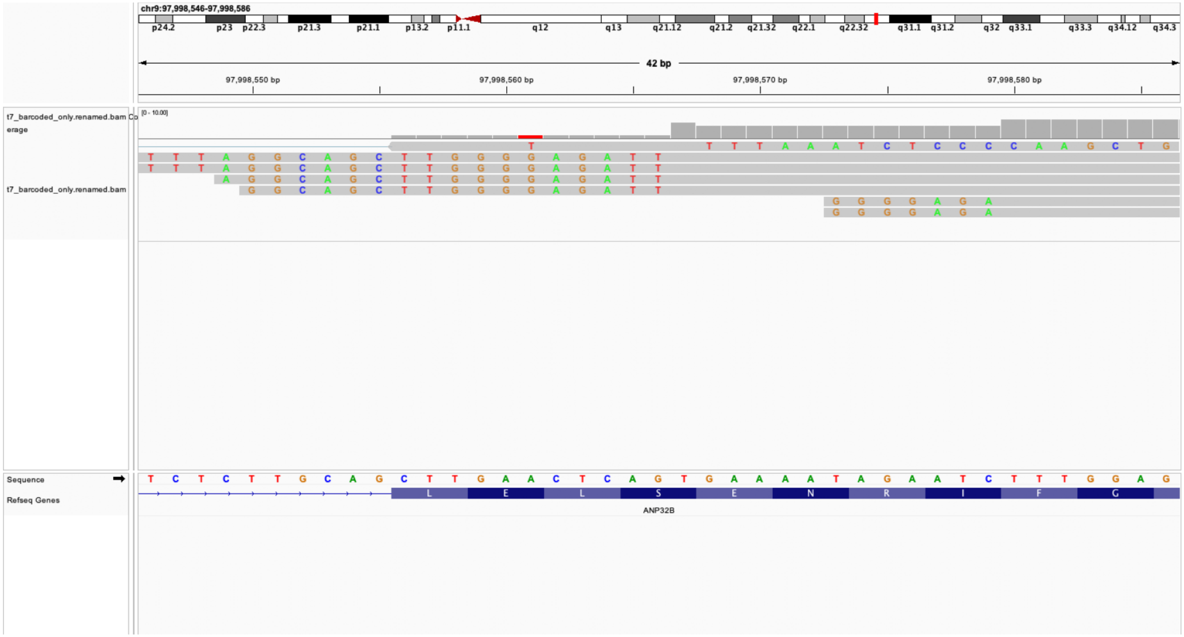

### manual edit call:False loc: chr7;155680729 genes=RBM33

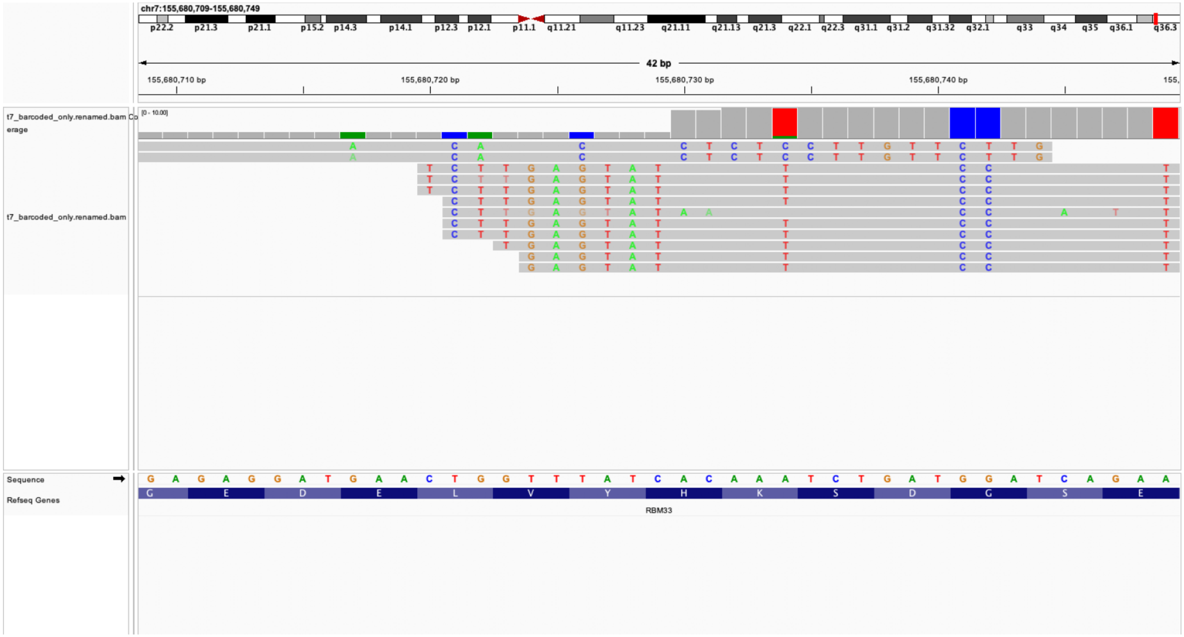
