## Supplementary figures and images for "Joint single-cell capture of Cas9 edits and transcriptomes reveals on- and off-target effects on gene expression"

### sheriff2.png

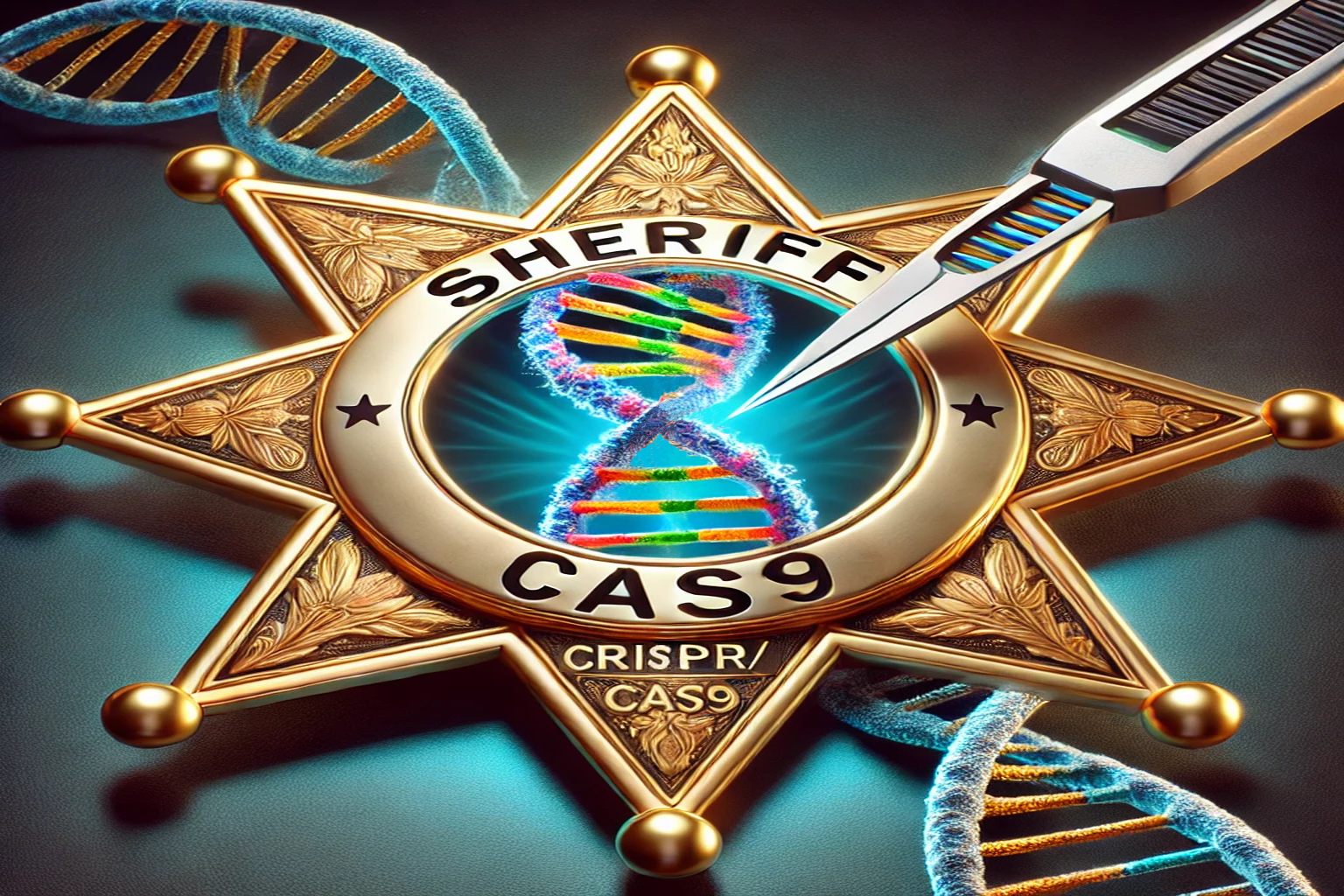

### sheriff.png

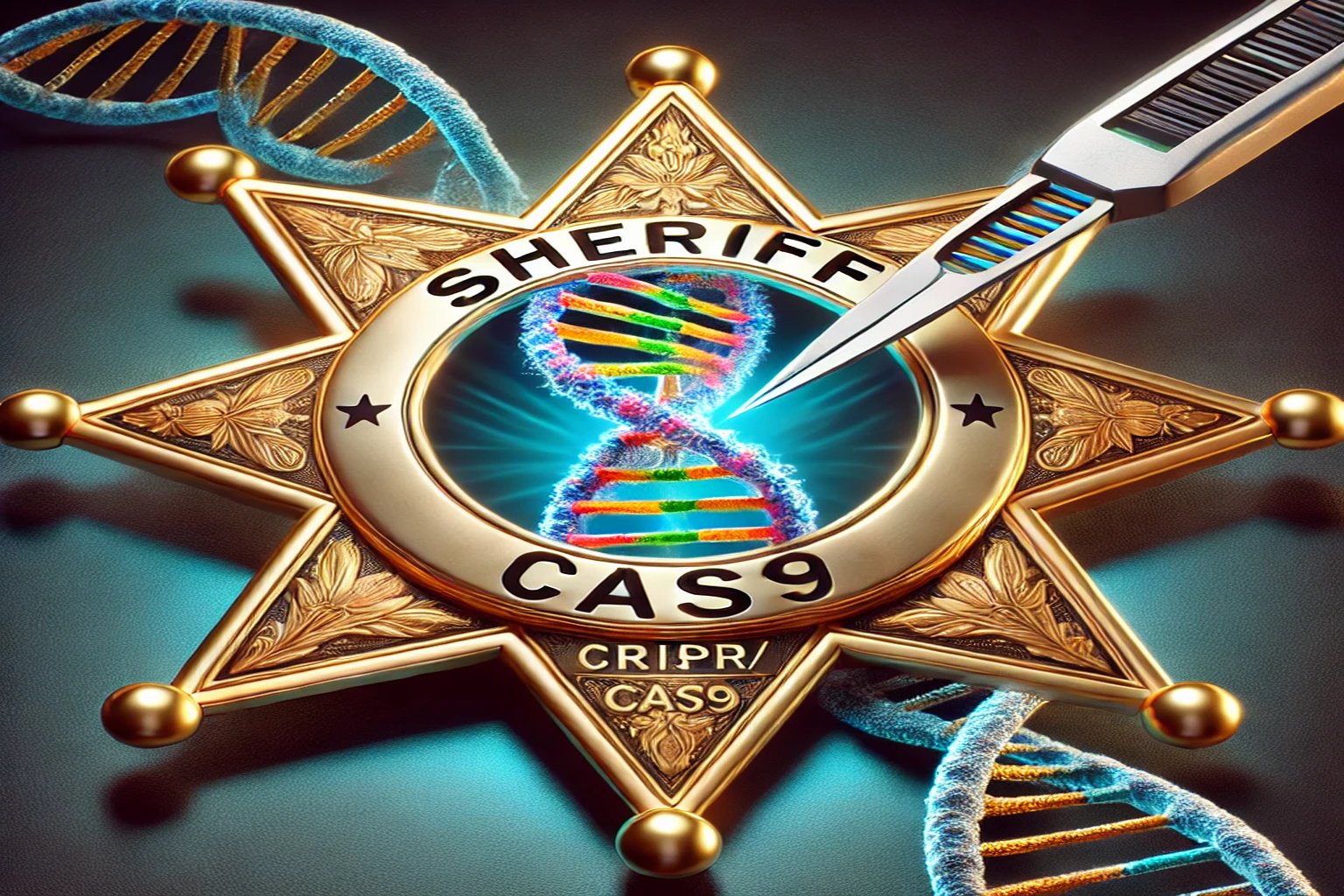

### sheriff_orig.png

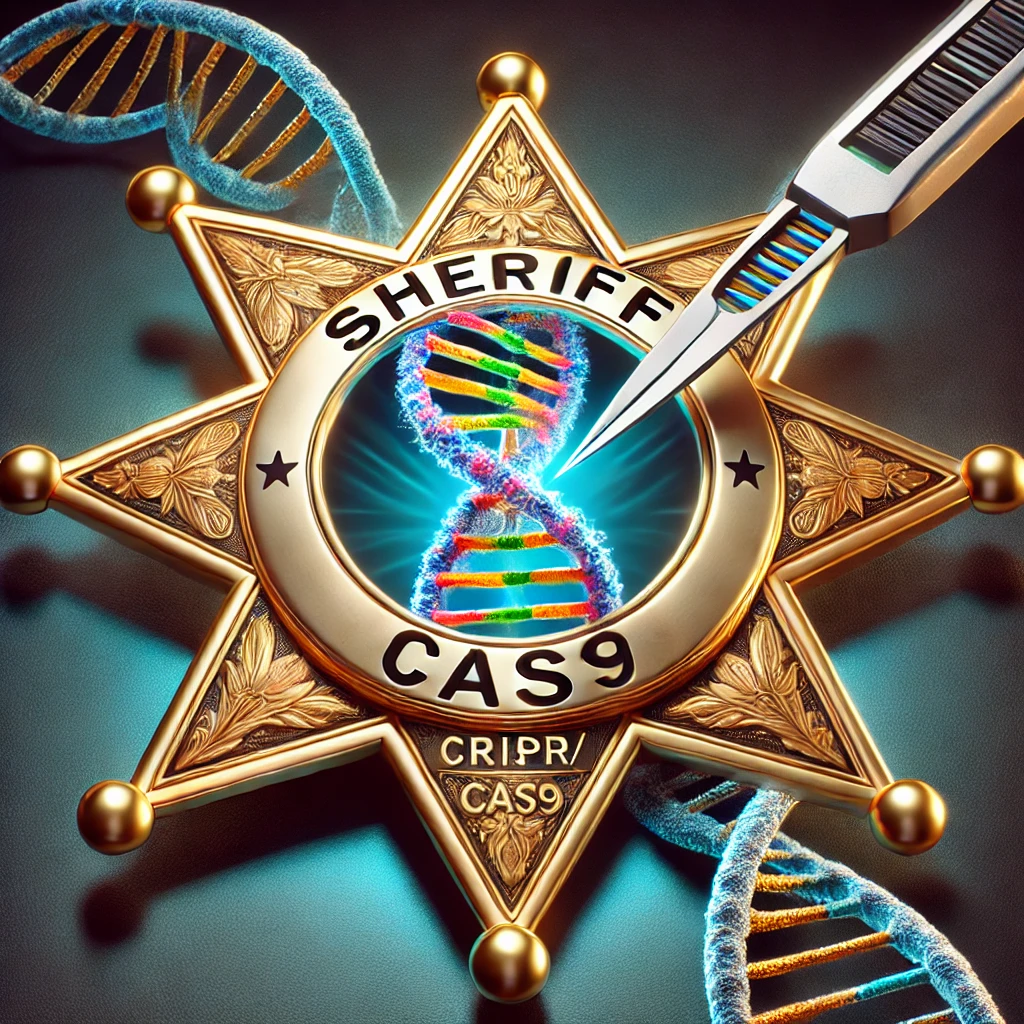
